# Proteomic and mitochondrial adaptations to early-life stress are distinct in juveniles and adults

**DOI:** 10.1101/2020.07.30.229690

**Authors:** Kathie L. Eagleson, Miranda Villaneuva, Rebecca M. Southern, Pat Levitt

## Abstract

Exposure to early-life stress (ELS) increases risk for poor mental and physical health outcomes that emerge at different stages across the lifespan. Yet, how age interacts with ELS to impact the expression of specific phenotypes remains largely unknown. An established limited-bedding paradigm was used to induce ELS in mouse pups over the early postnatal period. Initial analyses focused on the hippocampus, based on documented sensitivity to ELS in humans and various animal models, and the large body of data reporting anatomical and physiological outcomes in this structure using this ELS paradigm. An unbiased discovery proteomics approach revealed distinct adaptations in the non-nuclear hippocampal proteome in male versus female offspring at two distinct developmental stages: juvenile and adult. Gene ontology and KEGG pathway analyses revealed significant enrichment in proteins associated with mitochondria and the oxidative phosphorylation (OXPHOS) pathway in response to ELS in female hippocampus only. To determine whether the protein adaptations to ELS reflected altered function, mitochondrial respiration (driven through complexes II-IV) and complex I activity were measured in isolated hippocampal mitochondria using a Seahorse X96 Flux analyzer and immunocapture ELISA, respectively. ELS had no effect on basal respiration in either sex at either age. In contrast, ELS increased OXPHOS capacity in juvenile males and females, and reduced OXPHOS capacity in adult females but not adult males. A similar pattern of ELS-induced changes was observed for complex I activity. These data suggest that initial adaptations in juvenile hippocampus due to ELS were not sustained in adults. Mitochondrial adaptations to ELS were also exhibited peripherally by liver. Overall, the temporal distinctions in mitochondrial responses to ELS show that ELS-generated adaptations and outcomes are complex over the lifespan. This may contribute to differences in the timing of appearance of mental and physical disturbances, as well as potential sex differences that influence only select outcomes.

## 1. INTRODUCTION

Adverse childhood experiences (ACEs) contribute to negative outcomes in cognitive, social and emotional functioning, and long-term physical health (Anda et al., 1999; Anda et al., 2006; Danese et al., 2009; Dube et al., 2009; Dube et al., 2003; Edwards et al., 2003; Felitti et al., 1998). Outcomes emerge over the lifespan, often after ACEs exposure (Caspi et al., 2020). Mounting epidemiologic evidence indicates that negative health outcomes are influenced by a combination of direct effects of stressors on the developing brain and peripheral organs and altered life styles, including increased substance and alcohol abuse (National Academies of Sciences, 2019). There are limited data, however, on the molecular and cellular mechanisms that link ACEs to later-emerging outcomes. Indeed, the development of pathophysiologies prior to the expression of overt functional disruption is difficult to address in clinical studies.

There are several well-characterized preclinical models of ELS. Two rodent ELS models, classic maternal separation (Nishi et al., 2014) and the introduction of limited nesting and bedding materials (Walker et al., 2017), generate their effects by disrupting patterns of maternal care during the early postnatal period and are thus highly relevant to human ACEs. While both models induce stress in the pups, variability between studies is reduced because there is no daily human handling of the dam or pups in the limited bedding paradigm. In this paradigm, the dam makes repeated entries and exits off the nest, although total time on the nest and duration of nursing are not affected (for example, (Gilles et al., 1996; Heun-Johnson and Levitt, 2016; Rice et al., 2008)). This altered maternal behavior results in increased vocalization rates in the pups and increased basal plasma glucocorticoid levels at the end of the stress period (Heun-Johnson and Levitt, 2016; Naninck et al., 2015; Rice et al., 2008; Wang et al., 2011a). Mice stressed between postnatal days (P) 2-9 exhibit 1) reduced body weight gain over the stress period that normalizes post-pubertal; 2) increased plasma glucocorticoids and hypertrophied adrenal glands at the end of the ELS period; 3) alterations in hippocampal neuronal architecture and long-term potentiation; and 4) disruption in cognitive and pleasure/reward function, but no changes in anxiety and locomotor activity (Arp et al., 2016; Bolton et al., 2018a; Bolton et al., 2018b; Brunson et al., 2005; Heun-Johnson and Levitt, 2017; Ivy et al., 2010; Lesuis et al., 2019; Mitchell et al., 2018; Molet et al., 2016; Naninck et al., 2015; Naninck et al., 2017; Rice et al., 2008; Wang et al., 2011a; Yam et al., 2017a; Yam et al., 2019). When the same phenotypes were considered, there were similar results using a P4-11 ELS period, including reduced weight gain over the ELS period, disruption in cognitive and reward function, and reductions in glucocorticoid and mineralcorticoid receptor expression (Bath et al., 2016; Bath et al., 2017; Goodwill et al., 2019; Goodwill et al., 2018; Manzano-Nieves et al., 2018; Manzano Nieves et al., 2019). Peripheral measures also are impacted, including changes in adipose tissue, plasma leptin and ghrelin levels, visceral hyperalgesia, fecal microbial diversity, and responses to a high fat/high sugar diet (Guo et al., 2015; Maniam et al., 2015a; Maniam et al., 2015b; Moussaoui et al., 2017; Yam et al., 2017a; Yam et al., 2017b). Notably, different outcomes emerge at different times across the life span. While no animal model can replicate all variables that define human ACEs, these behavioral and physiological outcomes highlight the translational opportunities provided by the limited-bedding model.

The current study uses the limited-bedding paradigm to investigate molecular and functional adaptations to ELS in juveniles and adults. We focus on the hippocampus, based on documented sensitivity to ELS in humans and various animal models (Delpech et al., 2016; Derks et al., 2016; Hanson et al., 2015; Pillai et al., 2018; Suri and Vaidya, 2015; Teicher et al., 2018; Wei et al., 2014; Wei et al., 2015), and the wealth of anatomical and physiological data generated using this ELS paradigm (Brunson et al., 2005; Fenoglio et al., 2006; Heun-Johnson and Levitt, 2017; Ivy et al., 2010; Lesuis et al., 2019; Maniam et al., 2016; Molet et al., 2016; Naninck et al., 2015; Naninck et al., 2017; Rice et al., 2008; Walker et al., 2017; Wang et al., 2011b; Yam et al., 2019; Youssef et al., 2019). We first used an unbiased discovery approach to identify age-selective changes in the non-nuclear hippocampal proteome in juveniles and young adulthood. Based on these findings, we measured potential functional adaptations in the mitochondrial respiratory chain in isolated hippocampal mitochondria. The disease burden following ACEs often includes comorbid brain and metabolic dysfunction, suggesting that shared cellular mechanisms may underlie the emergence of peripheral and brain-related disorders. To address potential peripheral mitochondrial adaptations to ELS, we extend our functional analyses to mitochondria isolated from the liver, a key peripheral metabolic organ vulnerable to stress (Dong et al., 2003; Vasquez et al., 2014). We also included sex as a variable because epidemiological data in humans and basic studies using this ELS paradigm indicate both sexes are susceptible to ELS in overlapping and sometimes distinct ways (Arp et al., 2016; Bath et al., 2016; Bath et al., 2017; Goodwill et al., 2019; Goodwill et al., 2018; Guo et al., 2015; Knop et al., 2019; Maniam et al., 2015a; Maniam et al., 2015b; Manzano-Nieves et al., 2018; Manzano Nieves et al., 2019; Moussaoui et al., 2017; Naninck et al., 2015; Prusator and Greenwood-Van Meerveld, 2015, 2016; Rasmussen et al., 2019; Sumner et al., 2019; Walker et al., 2017; Yam et al., 2017a; Yam et al., 2017b; Youssef et al., 2019).

## 2. METHODS AND MATERIALS

Male and female C57Bl/6J mice were shipped from The Jackson Laboratory (Bar Harbor, ME) at 8-10 weeks of age. The Baram laboratory developed the limited-bedding ELS model in the C57Bl/6J strain (Rice et al., 2008). This strain has a large-scale deletion in the nicotinamide nucleotide transhydrogenase gene (Freeman et al., 2006), which encodes a protein located at the inner mitochondrial membrane that is important to redox balance in mitochondria. In the present study, there were no differences in basal mitochondrial respiration between maternal care conditions (see Results), although mice carrying this mutation have been reported to exhibit increased baseline reactive oxygen species, with negative physiological outcomes only apparent with additional genetic or environmental challenges (Francisco et al., 2018; Kim et al., 2010; Leskov et al., 2017; Meimaridou et al., 2018; Ripoll et al., 2012; Ronchi et al., 2013; Vozenilek et al., 2018). Mice were housed in a temperature (20-22°C) and humidity (40-60%) controlled vivarium under at 12-hour light/dark cycle, in standard ventilated mouse cages (Allentown, NJ), with Alpha-Dri bedding (Shepherd Specialty Papers, MI) and one standard cotton fiber nestlet square (Ancare Corp., NY). Food and water were provided ad libitum. All mouse procedures were approved by the Institutional Animal Care and Use Committee at the University of Southern California and conformed to NIH guidelines. All chemicals and reagents were from Sigma (St Louis, MO) unless otherwise noted.

### 2.1. Early-Life Stress Paradigm

A limited-bedding paradigm was used to induce ELS in mice. Detailed methods were as described previously (Heun-Johnson and Levitt, 2016), with minor modifications. Briefly, each cohort comprised 12 dams and 3 sires. The second litter of each dam was used, with the day of birth designated P0. Dams were assigned alternately to control or ELS conditions based on litter date of birth. On P2, litters were culled to 6 pups (sex ratio, 3:3 or 4:2) and exposed to ELS- or control-rearing until P9. Mice were weaned at P21 and housed with same-sex littermates. Analyses were performed at two ages to identify changes in response to ELS prior to (P21) or following (P90-120) onset of pubertal gonadal hormone exposure. Tissue used in this study was derived from 96 litters (sex ratio 3:3 - 89 litters, or 4:2 – 3 control, 4 ELS litters) generated over 18 independent cohorts between July 2016 and December 2019. Other litters generated were used for analyses not reported in this manuscript.

### 2.2. Tissue

Mice were anesthetized using 4% isoflurane, decapitated and the hippocampus and liver dissected. Hippocampal tissue (90-100mg) from 3 male or 3 female pups within a single litter was pooled to generate a single biological sample. A liver sample (80-120mg) from one male and female pup was also assayed; liver samples from the remaining pups in the litter were used in other studies. Tissue was either stored at −80°C until use (proteomics) or assayed immediately (mitochondrial respiration and complex I activity). We recognize that, in addition to hippocampus and liver, multiple brain regions, including prefrontal cortex and select nuclei in the hypothalamus and amygdala, and peripheral organs, including endocrine pancreas and white adipose tissue, are sensitive to early life stress. For the current study, in addition to the biological rationale noted above, we focused on the hippocampus and liver for technical advantages, including 1) the functional mitochondrial assays required a minimum of 70mg of fresh tissue for each sample, and 2) rapid, consistent dissection was optimal for these two tissues.

### 2.3 Proteomics screen

Hippocampi were homogenized at 4°C in homogenization buffer (320mM sucrose, 0.1mM CaCl_2_, 1.0mM MgCl_2_) containing protease inhibitor cocktail. The homogenate was centrifuged at 1000g for 10 minutes at 4°C and the supernatant collected. The pellet was resuspended in homogenization buffer, centrifuged at 1000g for 5 minutes at 4°C and the supernatant collected; this step was repeated once and the three supernatants pooled to generate a non-nuclear fraction that was precipitated with ice-cold acetone overnight at −20°C. The precipitated samples were centrifuged at 14,000g for 10 minutes at 4°C, dried and solubilized in 1% PPS Silent surfactant (Expedeon, San Diego, CA)/50mM Triethyl Ammonium Bicarbonate. Samples were assayed for protein concentration by the Pierce BCA protein assay kit (Thermo Fisher, Waltham, MA), and stored at −80°C prior to shipment to Vanderbilt University.

Proteomics procedures were performed at the Mass Spectrometry Research Center Proteomics Core at the Vanderbilt University School of Medicine. Three 4-plex isobaric tag for relative and absolute quantitation (iTRAQ) experiments, using independent cohorts, were performed at each age. Each experiment included: male control, female control, male ELS, female ELS. Mass spectrometry methods and data analyses were performed similarly to those described previously (Voss et al., 2015), with some minor modifications. One unit of labeling reagent (reporter tags: 114, 115, 116, 117) was used for 25μg protein. MudPIT analysis was performed using either a Q Exactive, Q Exactive Plus, or Q Exactive HF mass spectrometer. Samples analyzed on a Q Exactive were analyzed similar to Voss *et al.*, with a few differences. The Q Exactive instrument was operated in data-dependent mode acquiring HCD MS/MS scans after each MS1 scan on the 20 most abundant ions using an MS2 target of 1 × 10^5^ ions. The HCD-normalized collision energy was set to 30, and dynamic exclusion was set to 30 s. For experiments conducted on a QE Plus instrument, the instrument was operated in data-dependent mode acquiring HCD MS/MS scans on the 15 most abundant ions using an MS1 ion target of 3 × 10^6^ ions and an MS2 target of 1 × 10^5^ ions. For experiments conducted on a QE HF mass spectrometer, the instrument was operated in data-dependent mode acquiring HCD MS/MS scans at *R* = 15,000 on the 15 most abundant ions. For experiments on both the QE Plus and QE HF instruments, a Dionex Ultimate 3000 nano LC and autosampler were used, and peptides were gradient-eluted from the reverse analytical column at a flow rate of 350nL/min. For the peptides from the first 11 strong cation exchange (SCX) fractions, the reverse phase gradient consisted of 2–50% solvent B (0.1% formic acid in acetonitrile) in 83 min, followed by a 10 min equilibration at 2% solvent B. For the last 2 SCX-eluted peptide fractions, the peptides were eluted from the reverse phase analytical column using a gradient of 2-98% solvent B in 83 min, followed by a 10 min equilibration at 2% solvent B. Peptide/protein identifications and quantitative analysis were performed using Spectrum Mill (Agilent Technologies, Santa Clara, CA) as described previously. MS/MS spectra were searched against a subset of the UniProt KB protein database (www.uniprot.org) containing *Mus musculus* proteins. Autovalidation procedures in Spectrum Mill were used to filter the data rigorously to <1% false discovery rates at the protein and peptide level. For each group in each experiment, Log2 protein ratios were fit to a normal distribution using non-linear (least squares) regression. The calculated mean derived from the Gaussian fit was used to normalize individual log2 ratios for each quantified protein.

The mass spectrometry proteomics data were deposited to the ProteomeXchange Consortium via the PRIDE partner repository with the dataset identifier PXD013460 (Perez-Riverol et al., 2019). File names correspond to the following experiments: P21, 3906 (experiment 1), 4298 (experiment 2) and 4374 (experiment 3); adult, 4660 (experiment 1), 4787 (experiment 2) and 4932 (experiment 3). In each experiment, the following quantitative ratios for each protein were calculated: 1) male ELS:control and 2) female ELS:control – to determine sex-selective adaptations to ELS; 3) control male:female and 4) ELS male:female – to determine sex differences present normally in the hippocampus and if ELS alters this. Gene ontology (GO), using cellular component as the criteria, and Kyoto Encyclopedia of Genes and Genomes (KEGG) pathway enrichment analyses were performed using the Database for Annotation, Visualization, and Integrated Discovery (DAVID, version 6.8) web server (Huang et al., 2009) with default settings.

### 2.4. Mitochondrial isolation

Mitochondria were isolated from fresh hippocampus and liver using the MITOISO1 kit following the manufacturer’s protocol. Mitochondrial protein content was determined using the BCA protein assay kit (BioRad, Hercules CA). In pilot studies, mitochondrial integrity was confirmed by measuring citrate synthase activity in the presence and absence of CellLytic M, which lyses the mitochondria, following the manufacturer’s protocol (Cayman Chemical, Ann Arbor, MI; supplementary Figure S1). There was insufficient material generated to allow citrate synthase assays to be performed in preparations used for complex I and respiration assays.

### 2.5. Complex I activity

Mitochondrial complex I activity was measured using an immunocapture ELISA complex I enzyme activity assay kit, according to the manufacturer’s protocol (Abcam, Cambridge, MA). For each age, a single experiment included liver and hippocampal samples, representing one biological sample of each sex/stress group. To ensure that all assays were in the linear range, three dilutions of mitochondrial protein (20-80μg) from each group were plated per well. Measurements were done in duplicates that were averaged. Activity was expressed as change in absorbance/minute/ug protein.

### 2.6. Mitochondrial respiration

Respiration assays using isolated mitochondria were performed in the USC Leonard Davis School of Gerontology Seahorse Core Facility using a Seahorse XF96 flux analyzer (Agilent Technologies, Santa Clara, CA) as described (Agrawal et al., 2020; Rogers et al., 2011). For each age, a single microplate included liver and hippocampal samples, representing one biological sample of each sex/stress group. We note that, in one P21 and one adult respiration assay, only hippocampal mitochondria were assayed. To ensure that all assays were in the linear range, three dilutions of mitochondrial protein (1-6μg) from each group were plated per well. Bioenergic profiles were generated by measuring oxygen consumption rate (OCR) using 10mM succinate as a substrate in the presence of 2μM rotenone (driving respiration through complex II-IV activity). The assay was initiated in 4mM adenosine diphosphate (ADP), which provides a measure of oxidative phosphorylation (OXPHOS) capacity, followed by sequential addition of 1) 2.5μg/ml oligomycin (oligo, which inhibits complex V and evaluates proton leakage), 2) 4μM carbonyl cyanide-4-(trifluoromethoxy)phenylhydrazone (FCCP), which measures uncoupled respiration), and 3) 4μM antimycin A (AA, which inhibits respiration, providing a measure of non-respiratory oxygen consumption). In all but two experiments, basal respiration was measured in replicate wells on the same microplate. Measurements were done in duplicates that were averaged and then normalized to mitochondrial protein to give a final value. OCR was expressed as pmol/min/ug protein.

### 2.7. Statistical Analysis

Data are reported as mean and 95% confidence interval (CI), which was calculated using a *Z* statistic for n>30 and a *t* statistic for n<30. Individual measures for each assay are reported in the graphs and supplemental tables. For each assay and sex, an individual litter represents a single sample. For all tests, *p* values are reported to the third decimal place; *a priori* α = 0.050.

#### 2.7.1. Body weight

For analyses of offspring body weight, weights of male or female pups were averaged across a litter to generate a single value for each sex in each litter at each age. Comparisons between groups were performed at each age using a two-way analysis of variance (ANOVA, with sex and rearing as between subjects factors; offspring body weight) or an unpaired two-tailed *t-*test (dam body weight).

#### 2.7.2. Proteomics

Two by two χ^2^ analyses were performed to determine significant differences in the number of differentially-expressed proteins detected following ELS in 1) males versus females at P21; 2) males versus females in adults; 3) males at P21 versus adults; and 4) females at P21 versus adults. In juveniles and adults, significant differences in the distribution of protein ratios between the three experiments for male ELS versus control, and for female ELS versus control were determined using χ^2^ test.

#### 2.7.3. Mitochondrial assays

The studies were designed to detect large effects sizes in mitochondrial adaptations in the hippocampus and liver at each age at α=0.05 and 1-β=0.8. Effect sizes were determined by calculating omega squared (ω^2^), with small, medium and large effects classified as 0.010, 0.060, and > 0.140, respectively (Cohen, 1988; Lakens, 2013). We did not know *a priori* expected means and standard deviations for the complex I and Seahorse assays. Therefore, to determine sample sizes, power analyses were conducted using mean values and standard deviations observed following 6 experiments using G*Power, version 3.1.9.6 (Heinrich-Heine-Universitat Dusseldorf, Dusseldorf, Germany). To provide a complete overview of the data and to allow for power analyses in future studies, we report small, medium and large effect sizes. In addition, comparisons between groups were performed using a two-way ANOVA (sex and rearing as between-subjects factors), with significant interactions followed by Bonferroni/Dunn *post-hocs* (StatView, version 5.0.1; SAS Institute Inc, Cary, NC, USA). A Hedge’s *g*, appropriate for small sample sizes, was calculated as an estimate of effect size for each post hoc comparison. Final sample sizes, F-statistics and p-values are reported in Tables 3, 4 and 5.

## 3. RESULTS

We previously validated dam behavior in the limited-bedding paradigm in our laboratory, also replicating reduced pup weight gain over the stress period and cortisol increases at the end of the ELS period (Heun-Johnson and Levitt, 2016). We used pup weight measures to validate the impact of the ELS manipulation in the current study for each litter (Supplemental Figure S2; Supplemental Table S2).

### 3.1. Age- and sex-selective adaptations in the hippocampal proteome in response to ELS

To date, there has not been an in depth characterization of molecular changes due to ELS. Therefore, we applied a discovery-driven proteomics approach, measuring thousands of proteins in each sample, to determine differences in the P21 and adult non-nuclear hippocampal proteome following ELS (see Supplemental Tables S3 (P21) and S4 (adult) for complete list of proteins). Significantly more proteins were detected at P21 (3299.0 [95% CI: 2046.65, 4551.35]) compared to adults (2007.33 [95% CI: 956.93, 3057.73]; *t*_4_=3.4, *p*=0.027). We note that the absence of a protein cannot be distinguished from a low level of expression; thus, these data reflect differences in detection of different proteins across the two ages. In subsequent analyses, only proteins detected in at least two of the three experiments at each age (P21, 3216 proteins; adult, 1860 proteins; Supplemental Table S5) were considered. Based on this criterion, approximately 52% of proteins were detected at both ages (1766 proteins), 45% unique at P21 (1450) and 3% unique in the adult (94). These data indicate that in the hippocampus, a large proportion of proteins were downregulated, with only a relatively small number upregulated, over the course of juvenile to adult development.

We next determined adaptive changes in the non-nuclear hippocampal proteome in response to ELS in juveniles and adults, considering each sex independently. For these analyses, a protein ratio of 1.0 indicated no difference in expression between control- and ELS-reared mice. At each age, the distribution of protein species following ELS was overlapping but not identical across experiments (Figures 1A, B). Nevertheless, in each experiment, the distribution of protein ratios in males was significantly different than females (Table 1). To account for the heterogeneity in response, for each sex at each age, a protein was reported as differentially expressed if it met two criteria: 1) average fold-change across the three experiments was <0.8 or >1.2, similar to a previous report (Palmfeldt et al., 2016), and 2) this fold-change difference was obtained in at least two experiments. There were four key findings. First, there were more differentially-expressed proteins in adults than juveniles following ELS (males, ∼2.5-fold; females, ∼3.7-fold; Table 1), with most proteins exhibiting differences due to rearing distinct at the two ages (Supplemental Tables S6, S7). Second, almost all proteins (∼95%) were upregulated in response to ELS at P21, but surprisingly, the majority (∼60%) was downregulated in the adult, consistent with a striking age-dependent adaptation in the hippocampal proteome following ELS. Third, significantly more proteins were expressed differentially in response to ELS in females than males at both ages (Figure 1C, D; Table 1). Fourth, the many of the proteins exhibiting altered expression following ELS differed between males and females (Figure 1E, F; Supplemental Tables S6, S7). Thus, while both males and females exhibit changes in protein expression due to ELS, each sex has a unique response to the early environmental challenge.

**Figure 1.**
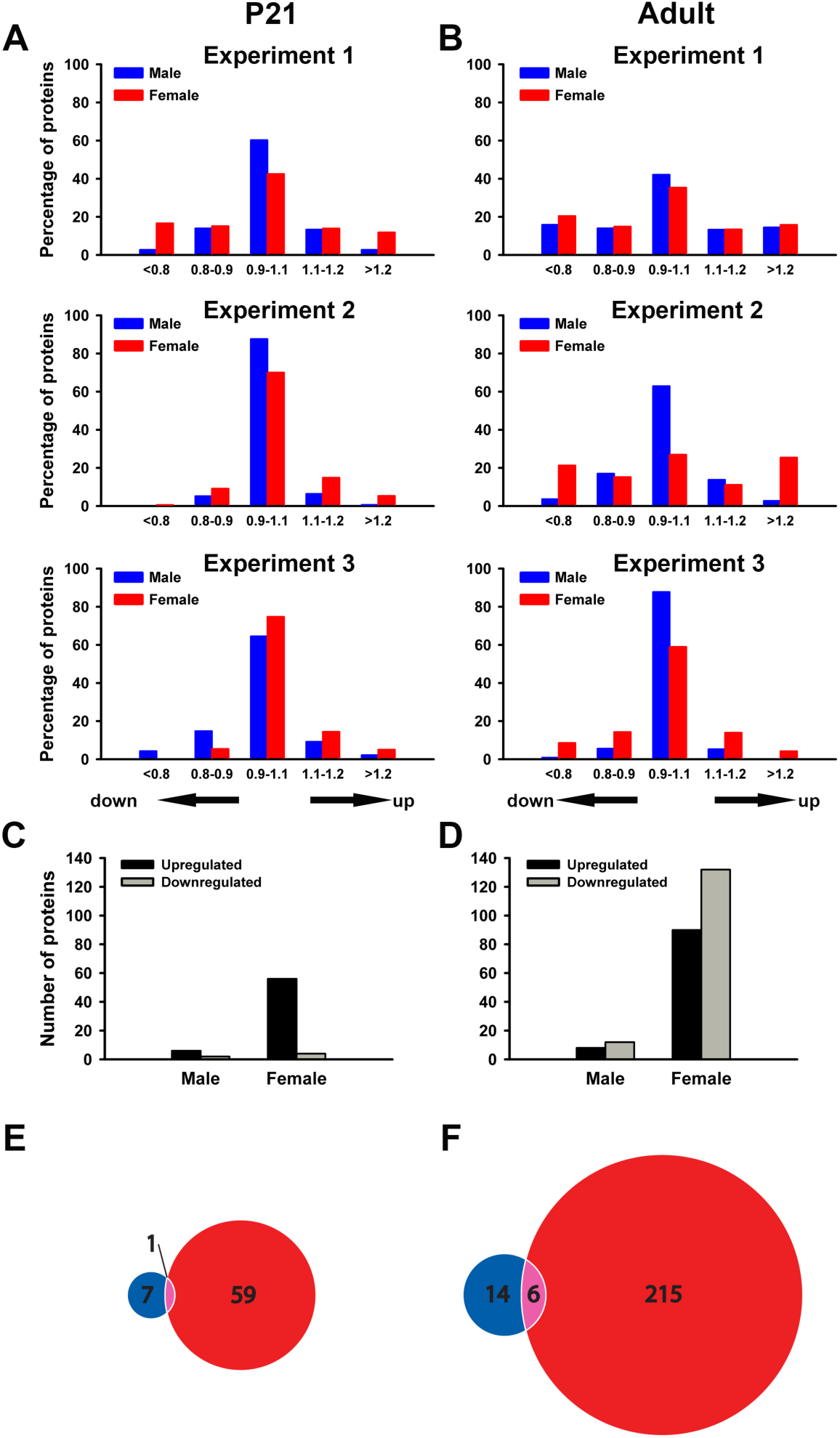
Females exposed to ELS exhibit more changes in the non-nuclear hippocampal proteome than males. (A, B) Chi-squared analyses of the distribution of protein ratios reveal significant differences in the response of males and females to ELS in each experiment at P21 (A) and in adults (B). (C, D) Graphs illustrate the number of hippocampal proteins that reach criterion for differential expression (see Results) following ELS across the three experiments at each age. Chi-squared analyses reveal that significantly more proteins are altered in response to ELS in the female compared to male hippocampus at P21 (C) and in adults (D). Statistical details are provided in Table 1. (E, F) Venn diagrams displaying the number of common and unique proteins altered in response to ELS in the male versus female hippocampus at P21 (E) and in adults (F).

**Table 1.**
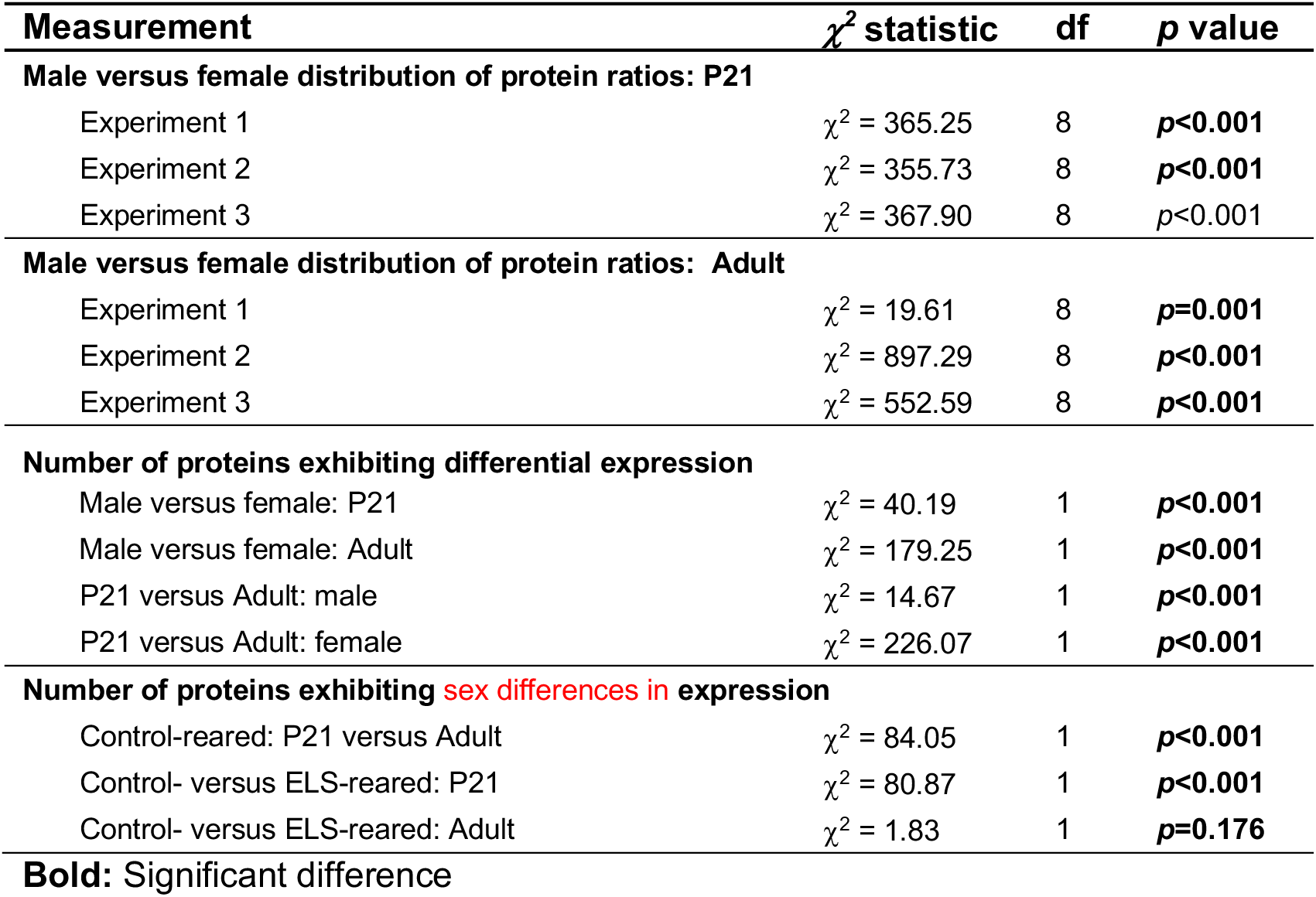
Statistical test details of proteomics measurements

The 4-plex iTRAQ design enabled us to examine sex differences in the hippocampal non-nuclear proteome before and after puberty. In control-reared mice, there was a 14- fold increase in the number of proteins exhibiting sex differences in expression in adult compared to P21 hippocampus (Figure 2A; Table 1). Such differences in expression were exaggerated following ELS at P21; further, only one of four proteins exhibiting sex differences in expression in control-reared mice at P21 continued to do so following ELS (Figure 2B). In adults, the number of proteins exhibiting sex differences in expression was similar on control-versus ELS-reared mice (Figure 2A; Table 1), but the identity of these proteins was largely non-overlapping (Figure 2C). These findings again highlight the different ELS-induced adaptations occurring in males versus females over time in the hippocampus.

**Figure 2.**
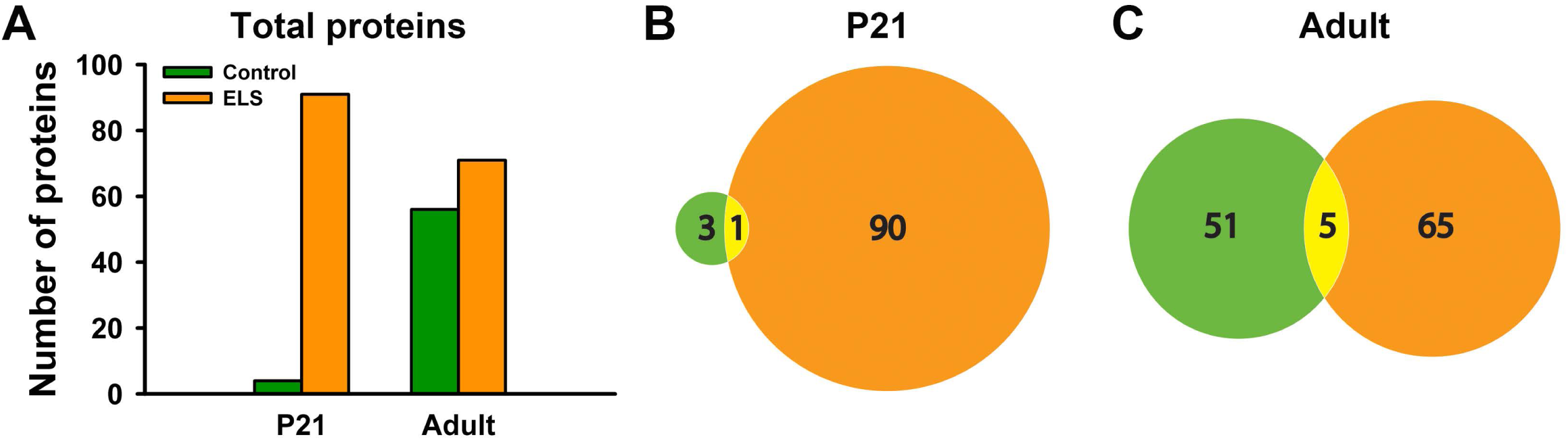
Sex differences in the non-nuclear hippocampal proteome. (A) Graph illustrates the number of proteins expressed differentially between males and females at each age following control- and ELS-rearing. Chi-squared analyses reveal that there is a significant increase in the number of proteins differentially expressed in control-reared mice in adults compared to P21, and that ELS increases the number of proteins exhibiting differential expression at P21 but not in the adult. Statistical details are provided in Table 1. (B, C) Venn diagrams displaying the number of common and unique proteins exhibiting sex differences in expression in the hippocampus of control- and ELS-reared mice at P21 (B) and in adults (C).

Changes in expression for individual proteins reported here typically were only in the 20-30% range. We therefore used GO enrichment analyses to identify if changes in protein expression in the hippocampus following ELS are associated with specific cellular components. Initial analyses, conducted for each sex at each age, considered all changed proteins as a single dataset. Five terms, including mitochondrion, were enriched significantly in P21 females (Figure 3A); forty-one terms were enriched in adult females, including mitochondrial inner membrane, mitochondrion, respiratory chain and mitochondrial respiration chain complex I (Figure 3E; Supplementary Table S8). No terms were enriched in P21 males; two terms, myelin sheath and extracellular exosome, were enriched in adult males (Figure 3C). A list of proteins associated with the mitochondrion, defined by GO analyses, altered in response to ELS is provided in Supplemental Tables S9 (P21) and S10 (adult). We next conducted GO analyses on subsets of differentially expressed proteins following ELS. Upregulated proteins demonstrated enrichment only in females, with 4 terms - including mitochondrion – returned at P21 (Figure 3B) and 25 terms - including mitochondrion and mitochondrial inner membrane - in adults (Figure 3F; Supplementary Table S8). In contrast, analysis of proteins downregulated in response to ELS returned a set of enriched GO terms in adults only; specifically, enrichment in one term, extracellular exosome, in adult males (Figure 3D) and 6 terms in adult females (Figure 3G).

**Figure 3.**
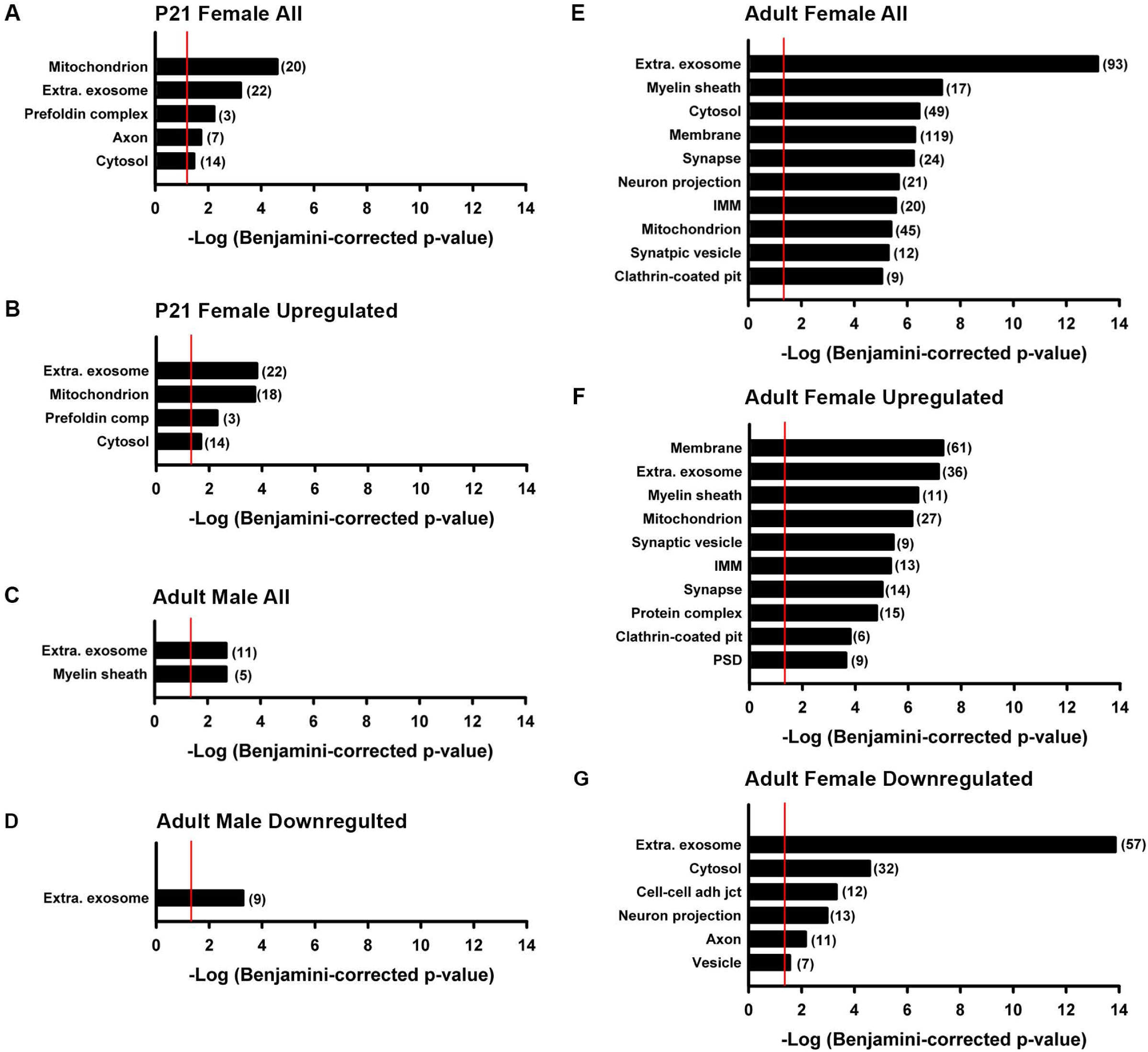
Gene ontology – cellular component - analyses of proteins expressed differentially in male and female control-versus ELS-reared hippocampus. When considering all differentially expressed proteins, there is enrichment in mitochondrial proteins in the P21 female (A) and adult female (E), but not the adult male, hippocampus (C). No terms are enriched in P21 males (not shown). Similar analyses considering upregulated proteins independently return enriched GO terms in P21 (B) and adult (F) female hippocampus only, while downregulated proteins return enriched GO terms only in adult male (D) and female (G) hippocampus. Note that for the adult female analyses including all and upregulated proteins, only the top 10 enriched terms are illustrated. The complete set is listed in Supplementary Table S8. Numbers in parentheses indicate the number of proteins for each term. Red line indicates p=0.05.

KEGG pathway analyses of the same datasets returned terms only in female hippocampus (Figure 4; Supplementary Table S11). When considering all differentially expressed proteins following ELS as a single dataset, there is enrichment in 2 KEGG pathways in females at P21 (Figure 4A), including one related to pyruvate metabolism, and 40 KEGG pathways in adults, including OXPHOS and the TCA cycle (Figure 4C; Supplementary Table S11). Upregulated proteins returned 3 terms in females at P21 (Figure 4B) and 35 pathways in adults (Figure 4E, Supplementary Table S11). In contrast, analysis of downregulated proteins returned an enrichment in 3 KEGG pathways in adult females only (Figure 4D).

**Figure 4.**
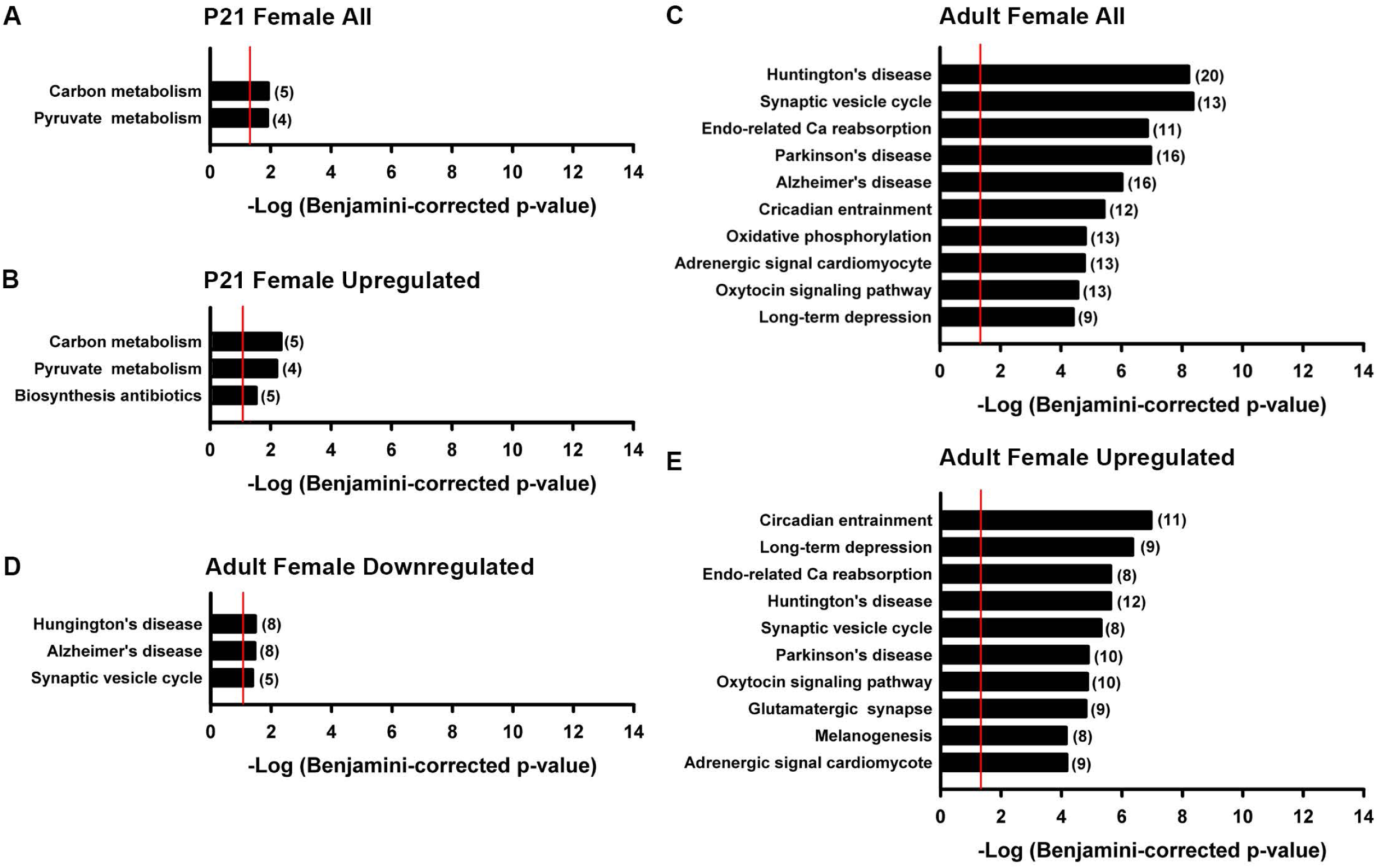
KEGG pathway analyses of proteins expressed differentially in male and female control-versus ELS-reared hippocampus. There is enrichment in KEGG pathways in females only. Analyses including all differentially expressed proteins returns 2 pathways at P21 (A) and 40 pathways in adults (C). Similar analyses considering upregulated proteins independently return 3 enriched pathways in P21 (B) and 35 pathways in adult (C) hippocampus, while downregulated proteins return enriched pathways only in adult hippocampus (D). Note that for the adult analyses including all and upregulated proteins, only the top 10 enriched terms are illustrated. The complete set is listed in Supplementary Table S11. Numbers in parentheses indicate the number of proteins for each pathway. Red line indicates p=0.05.

### 3.2. Age-selective adaptations in hippocampal mitochondrial function in response to ELS

The proteomics analyses revealed alterations in the expression levels of five subunits of complex I in adult female hippocampus following ELS (Table 2); alterations in complex I expression/activity have been associated with the pathophysiology of several brain-based disorders (Gu et al., 2013; Holper et al., 2018) and observed in animal models following adult metabolic or chronic stress (Emmerzaal et al., 2020; Toniazzo et al., 2019). Based on the iTRAQ discoveries, we next explored whether the protein adaptations to ELS reflected altered function, using an enzyme activity assay in mitochondria isolated from the hippocampus in juveniles and adults. At P21, there was a significant large effect of rearing (ω^2^ = 0.288) on complex I activity (Figure 5A), with complex I activity significantly higher following ELS (control: 5.34 [95% CI: 4.76, 5.92], ELS: 7.66 [95% CI: 6.76, 8.56]; Table 3). Sex had a small effect on complex I activity at this age (ω^2^ = 0.028) and there was no sex*ELS interaction (ω^2^ = 0.007). There was also a significant large effect of rearing on complex I activity (ω^2^ = 0.421) in adults; strikingly, in contrast to juveniles, complex I activity was reduced following ELS (control: 7.14 [95% CI: 6.47, 7.82], ELS: 4.85 [95% CI: 4.36, 5.34]; Figure 5B; Table 3). There was no main effect of sex (ω^2^ = −0.011), but the rearing effect was moderated by a significant interaction between sex and rearing (ω^2^ = 0.082; Table 3). Post-hoc analyses revealed a significant reduction in complex I activity in females (control: 7.71 [95% CI: 6.67, 8.75], ELS: 4.35 [95% CI: 3.91, 4.79]) but not males (control: 6.57 [95% CI: 5.68, 7.46], ELS: 5.34 [95% CI: 4.48, 6.20]) when corrected for multiple comparisons (Table 3).

**Figure 5.**
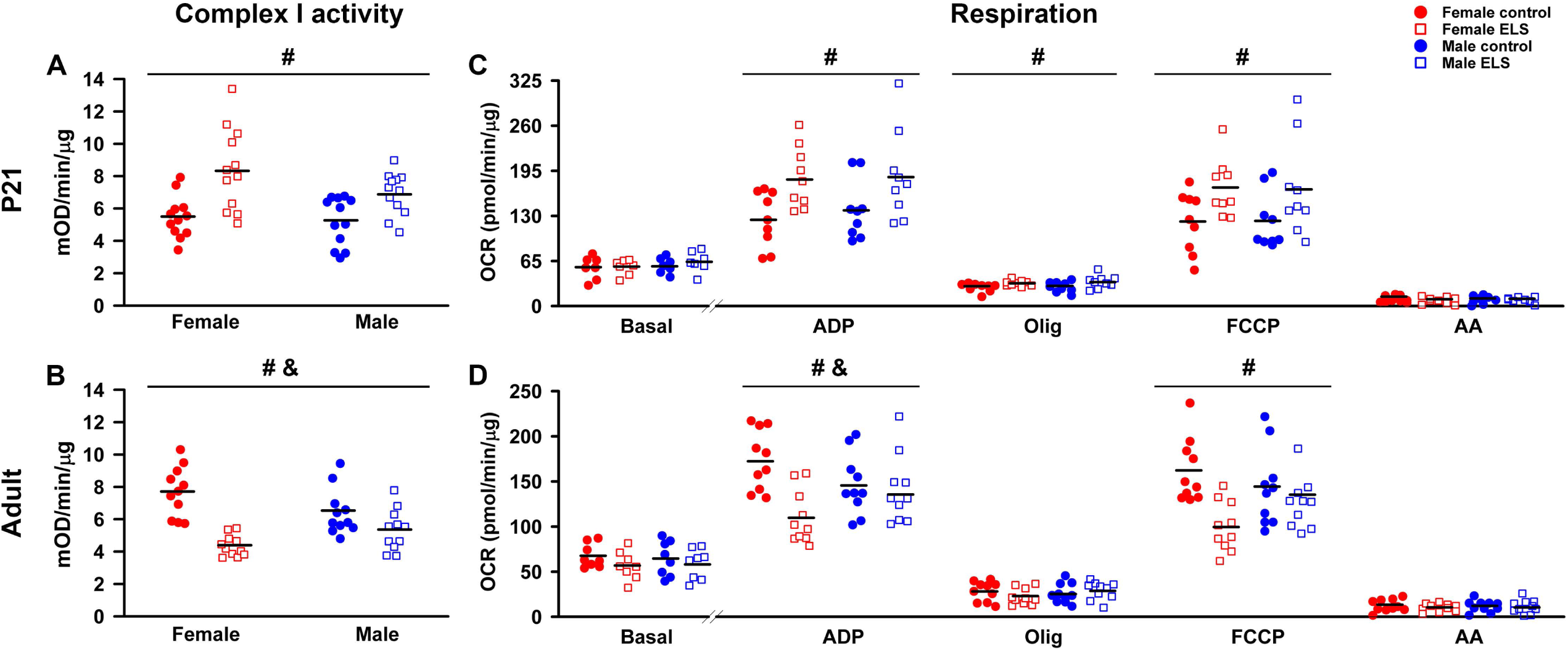
Age-dependent changes in mitochondrial function in hippocampus following ELS. Complex I activity (A, B) and oxygen consumption rate (OCR) driven through complex II-IV (C, D) were measured in mitochondria isolated from P21 (top panels) or adult hippocampus (bottom panels). ELS increases complex I activity in males and females at P21 (A), and decreases activity in females in the adult (B). (C, D) OCR was measured under basal conditions or in sister wells following the sequential addition of adenosine diphosphate (ADP), oligomycin (olig), FCCP and antimycin A (AA). At P21, OCR is increased in the presence of ADP, olig and FCP (C). In contrast, OCR is decreased in the presence of ADP and FCCP in adults (D). ^#^main effect of ELS; ^&^sex*ELS interaction by two-way ANOVA. Details of statistical analyses and sample sizes are provided in Tables 3 and 4. The raw data contributing to these figures are found in Supplementary Tables S12-S15. Each symbol represents an independent sample, with the bar representing the group mean.

**Table 2.** Expression of OXPHOS proteins altered by ELS in adult female hippocampus

| **Protein** | **Direction of change** |
| --- | --- |
| ATP synthase subunit b, mitochondrial | Up |
| ATP synthase subunit O, mitochondrial | Up |
| Cytochrome c oxidase subunit 2 | Up |
| Cytochrome c oxidase subunit 6C | Up |
| NADH dehydrogenase [ubiquinone] 1 alpha subcomplex subunit 12<sup>a</sup> | Up |
| NADH dehydrogenase [ubiquinone] 1 alpha subcomplex subunit 9<sup>a</sup> | Up |
| V-type proton ATPase subunit d 1 | Up |
| Cytochrome b-c1 complex subunit 6, mitochondrial | Down |
| Cytochrome c oxidase subunit 6B1 | Down |
| NADH dehydrogenase [ubiquinone] 1 alpha subcomplex subunit 2<sup>a</sup> | Down |
| NADH dehydrogenase [ubiquinone] iron-sulfur protein 4, mitochondrial<sup>a</sup> | Down |
| NADH dehydrogenase [ubiquinone] flavoprotein 2, mitochondrial<sup>a</sup> | Down |
| V-type proton ATPase subunit F | Down |

<sup>a</sup>Subunit of complex I

**Table 3.** Statistical test details of Complex I measurements

|  |  | Test statistic | p value | Effect size |
| --- | --- | --- | --- | --- |
| **P21 hippocampus (n=12/group)** |  |  |  |  |
| Two-way ANOVA | Sex | $F(1,44) = 2.98$ | 0.091 | $\omega^2 = 0.028^b$ |
|  | **Rearing** | **$F(1,44) = 21.38$** | **<0.001** | **$\omega^2 = 0.288$** |
| | Sex*rearing | $F(1,44) = 1.52$ | 0.224 | $\omega^2 = 0.007$ |
| **Adult hippocampus (n=11/group)** |  |  |  |  |
| Two-way ANOVA | Sex | $F(1,40) = 0.04$ | 0.837 | $\omega^2 = -0.011$ |
|  | **Rearing** | **$F(1,40) = 37.44$** | **<0.001** | **$\omega^2 = 0.421$** |
|  | **Sex\*rearing** | **$F(1,40) = 8.11$** | **0.007** | **$\omega^2 = 0.082$** |
| Post-hoc<br>(sex*rearing) | **Female control** | **Female ELS** | **<0.001<sup>a</sup>** | **$g = 2.730$** |
| | Male control | Male ELS | 0.026 | $g = 0.907$ |
| | Female control | Male control | 0.037 | $g = 0.768$ |
| | Female ELS | Male ELS | 0.069 | $g = 0.937$ |
| **P21 liver (n=12/group)** |  |  |  |  |
| Two-way ANOVA | Sex | $F(1,44) = 2.54$ | 0.118 | $\omega^2 = 0.023^b$ |
|  | **Rearing** | **$F(1,44) = 14.16$** | **<0.001** | **$\omega^2 = 0.200$** |
| | Sex*rearing | $F(1,44) = 4.01$ | 0.051 | $\omega^2 = 0.046^b$ |
| **Adult liver (n=11/group)** |  |  |  |  |
| Two-way ANOVA | Sex | $F(1,40) = 0.02$ | 0.893 | $\omega^2 = -0.024$ |
| | Rearing | $F(1,40) = 0.03$ | 0.874 | $\omega^2 = -0.024$ |
| | Sex*rearing | $F(1,40) < 0.01$ | 0.964 | $\omega^2 = -0.024$ |

Bold, significantly different by two-way ANOVA at $1-\beta=0.8$ ; <sup>a</sup>significantly different means by Bonferroni/Dunn post-hocs, following correction for multiple comparisons where appropriate

**Table 4.**
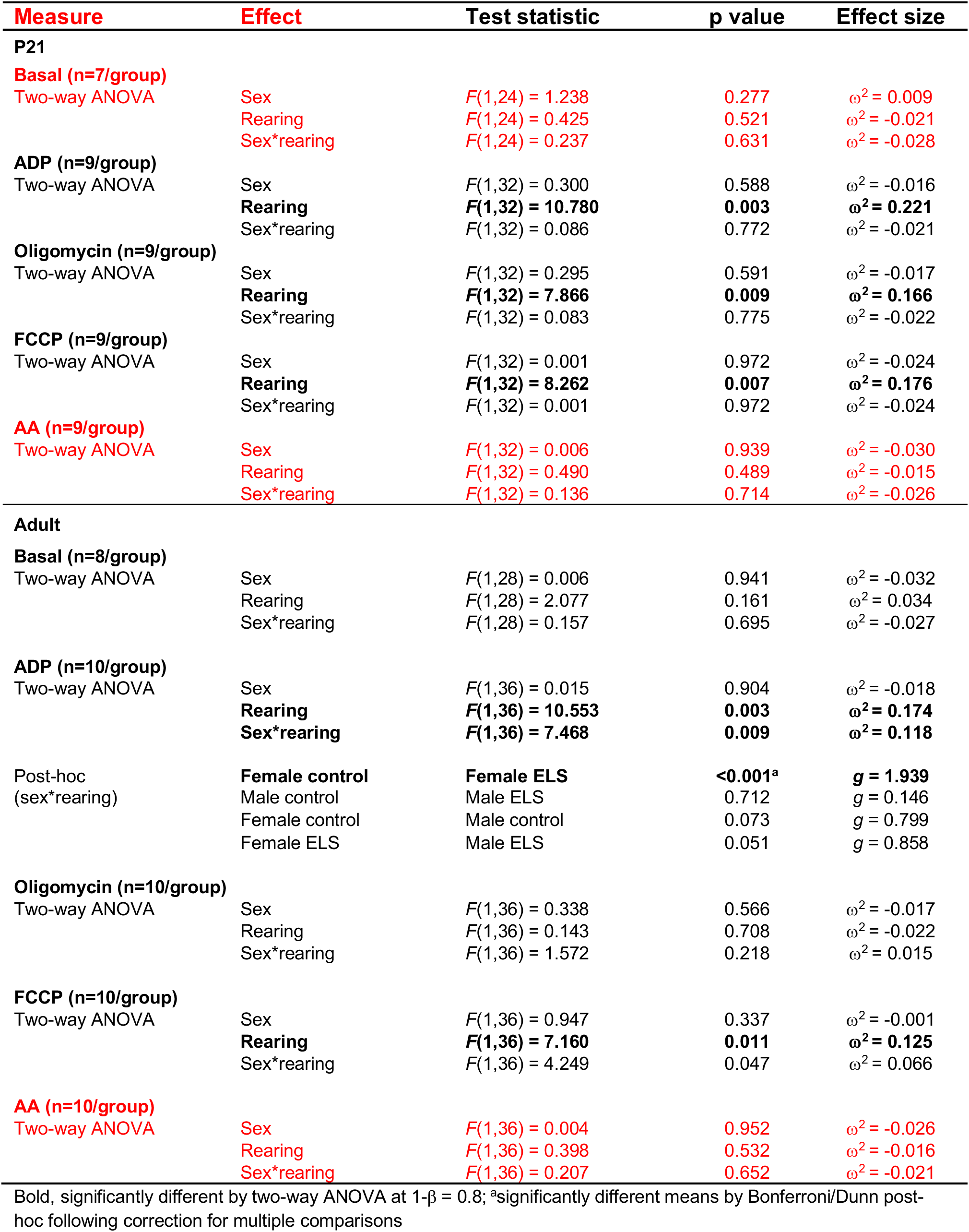
Statistical test details of hippocampal respiration measurements^a^

| **Measure** | **Effect** | **Test statistic** | **p value** | **Effect size** |
| --- | --- | --- | --- | --- |
| **P21** |  |  |  |  |
| **Basal (n=7/group)** |  |  |  |  |
| Two-way ANOVA | Sex | $F(1,24) = 1.238$ | 0.277 | $\omega^2 = 0.009$ |
| | Rearing | $F(1,24) = 0.425$ | 0.521 | $\omega^2 = -0.021$ |
| | Sex*rearing | $F(1,24) = 0.237$ | 0.631 | $\omega^2 = -0.028$ |
| **ADP (n=9/group)** |  |  |  |  |
| Two-way ANOVA | Sex | $F(1,32) = 0.300$ | 0.588 | $\omega^2 = -0.016$ |
|  | **Rearing** | **$F(1,32) = 10.780$** | **0.003** | **$\omega^2 = 0.221$** |
| | Sex*rearing | $F(1,32) = 0.086$ | 0.772 | $\omega^2 = -0.021$ |
| **Oligomycin (n=9/group)** |  |  |  |  |
| Two-way ANOVA | Sex | $F(1,32) = 0.295$ | 0.591 | $\omega^2 = -0.017$ |
|  | **Rearing** | **$F(1,32) = 7.866$** | **0.009** | **$\omega^2 = 0.166$** |
| | Sex*rearing | $F(1,32) = 0.083$ | 0.775 | $\omega^2 = -0.022$ |
| **FCCP (n=9/group)** |  |  |  |  |
| Two-way ANOVA | Sex | $F(1,32) = 0.001$ | 0.972 | $\omega^2 = -0.024$ |
|  | **Rearing** | **$F(1,32) = 8.262$** | **0.007** | **$\omega^2 = 0.176$** |
| | Sex*rearing | $F(1,32) = 0.001$ | 0.972 | $\omega^2 = -0.024$ |
| **AA (n=9/group)** |  |  |  |  |
| Two-way ANOVA | Sex | $F(1,32) = 0.006$ | 0.939 | $\omega^2 = -0.030$ |
| | Rearing | $F(1,32) = 0.490$ | 0.489 | $\omega^2 = -0.015$ |
| | Sex*rearing | $F(1,32) = 0.136$ | 0.714 | $\omega^2 = -0.026$ |
| **Adult** |  |  |  |  |
| **Basal (n=8/group)** |  |  |  |  |
| Two-way ANOVA | Sex | $F(1,28) = 0.006$ | 0.941 | $\omega^2 = -0.032$ |
| | Rearing | $F(1,28) = 2.077$ | 0.161 | $\omega^2 = 0.034$ |
| | Sex*rearing | $F(1,28) = 0.157$ | 0.695 | $\omega^2 = -0.027$ |
| **ADP (n=10/group)** |  |  |  |  |
| Two-way ANOVA | Sex | $F(1,36) = 0.015$ | 0.904 | $\omega^2 = -0.018$ |
|  | **Rearing** | **$F(1,36) = 10.553$** | **0.003** | **$\omega^2 = 0.174$** |
|  | **Sex\*rearing** | **$F(1,36) = 7.468$** | **0.009** | **$\omega^2 = 0.118$** |
| Post-hoc<br>(sex*rearing) | **Female control** | **Female ELS** | **<0.001<sup>a</sup>** | **$g = 1.939$** |
| | Male control | Male ELS | 0.712 | $g = 0.146$ |
| | Female control | Male control | 0.073 | $g = 0.799$ |
| | Female ELS | Male ELS | 0.051 | $g = 0.858$ |
| **Oligomycin (n=10/group)** |  |  |  |  |
| Two-way ANOVA | Sex | $F(1,36) = 0.338$ | 0.566 | $\omega^2 = -0.017$ |
| | Rearing | $F(1,36) = 0.143$ | 0.708 | $\omega^2 = -0.022$ |
| | Sex*rearing | $F(1,36) = 1.572$ | 0.218 | $\omega^2 = 0.015$ |
| **FCCP (n=10/group)** |  |  |  |  |
| Two-way ANOVA | Sex | $F(1,36) = 0.947$ | 0.337 | $\omega^2 = -0.001$ |
|  | **Rearing** | **$F(1,36) = 7.160$** | **0.011** | **$\omega^2 = 0.125$** |
| | Sex*rearing | $F(1,36) = 4.249$ | 0.047 | $\omega^2 = 0.066$ |
| **AA (n=10/group)** |  |  |  |  |
| Two-way ANOVA | Sex | $F(1,36) = 0.004$ | 0.952 | $\omega^2 = -0.026$ |
| | Rearing | $F(1,36) = 0.398$ | 0.532 | $\omega^2 = -0.016$ |
| | Sex*rearing | $F(1,36) = 0.207$ | 0.652 | $\omega^2 = -0.021$ |

Bold, significantly different by two-way ANOVA at $1-\beta = 0.8$ ; <sup>a</sup>significantly different means by Bonferroni/Dunn post-hoc following correction for multiple comparisons

In the adult female hippocampus, KEGG pathway analyses revealed that 11/45 (∼25%) mitochondrial proteins exhibiting increased/decreased expression following ELS were involved in OXPHOS (Table 2). To investigate functionally the differential expression of OXPHOS proteins, we conducted Seahorse respiration experiments measuring OCR. OCR was unstable over time with complex I substrates. Therefore, mitochondria were incubated with succinate, which drives respiration through complex II, and rotenone, which inhibits complex I. At P21, there was a significant large effect of rearing on OCR in the presence of ADP (ω^2^ = 0.221), oligomycin (ω^2^ = 0.166) and FCCP (ω^2^ = 0.176), but no effect of sex or sex*rearing interaction in the presence of any drug (Figure 5C; Table 4). Specifically, OCR was significantly higher following ELS in the presence of ADP (control: 131.61 [95% CI: 111.45, 151.77], ELS: 185.31 [95% CI: 158.33, 212.29]), oligomycin (control: 27.32 [95% CI: 24.02, 30.62], ELS: 33.84 [95% CI: 30.38, 37.30]) and FCCP (control: 120.85 [95% CI: 100.97, 140.73], ELS: 169.44 [95% CI: 142.46, 196.42]). All main effects and interactions were non-significant for OCR under basal conditions and in the presence of antimycin A (Table 4). These observations indicate an increase in maximal respiration in male and female hippocampal mitochondria following ELS that potentially could enhance the organelle’s ability to respond during an increase in energy demand. Further, the increase in OCR in the presence of oligomycin suggests a mild OXPHOS uncoupling following ELS. In the adult, there was a significant large effect of rearing (ω^2^ = 0.174), but no effect of sex (ω^2^ = 0.018), on OCR in the presence of ADP (Figure 5D; Table 4). The rearing effect was moderated by a significant sex*rearing interaction, with medium effect (ω^2^ = 0.118; Table 4). Similar to observations with Complex I, post-hoc analyses indicated that there was a significant reduction in OCR in ADP in females (control: 173.91 [95% CI: 150.12, 197.70], ELS: 110.22 [95% CI: 89.07, 131.37]), but not males (control: 146.09 [95% CI: 122.15, 170.03], ELS: 140.64 [95% CI: 113.57, 167.71]), following ELS (Table 4). There was also a significant medium effect of rearing (ω^2^ = 0.125), but no effect of sex (ω^2^ = −0.001), in the presence of FCCP (Figure 4D; Table 4), with OCR significantly lower following ELS (control: 152.01 [95% CI: 132.47, 171.55], ELS: 117.62 [95% CI: 100.34, 134.91]). This suggested that, in contrast to juveniles, there was a reduction in maximal respiration in adult hippocampal mitochondria following ELS, consistent with a reduced ability to respond to an increase in energy demand. We note that for adult respiration, the data also revealed a sex*rearing interaction with medium effect (ω^2^ = 0.125) in the presence of FCCP (Table 4), small effect of ELS on basal respiration (ω^2^ = 0.034) and a sex*rearing interaction with small effect (ω^2^ = 0.015) in the presence of oligomycin (Table 4), but the significance of these effects will require further investigation in future studies. All other main effects and interactions were non-significant for OCR (Figure 5D; Table 4).

Together, the complex I and respiration analyses indicate an age-selective response in hippocampal mitochondrial function in response to ELS. Although there is no significant main effect of sex on any measure, in the adult, the impact of ELS on complex I activity and OCR in the presence of ADP is moderated by sex.

### 3.3. Liver mitochondria are also impacted by ELS

It has been suggested that shared cellular mechanisms may underlie the emergence of peripheral and brain-related disorders. Thus, we wondered if similar changes in respiratory chain function could be observed peripherally. Analyses were performed simultaneously in mitochondria isolated from livers of the same animals used in the hippocampal assays. Similar to hippocampus, there was a significant large effect of rearing (ω^2^ = 0.200) on complex I activity at P21 (Figure 6A), with complex I activity significantly higher following ELS (control: 5.46 [95% CI: 5.11, 5.81], ELS: 6.83 [95% CI: 6.12, 7.54]; Table 3). Sex had a small effect on complex I activity at this age (ω^2^ = 0.023) and there was a small sex*ELS interaction (ω^2^ = 0.046; Table 3). In contrast to the adult hippocampus, however, there was no effect of ELS (ω^2^ = −0.024) or sex (ω^2^ = - 0.024), or sex*ELS interaction (ω^2^ = −0.024), on adult liver mitochondrial complex I activity (Figure 6B; Table 3). We next measured respiration driven through complex II-IV. At P21, all main effects and interactions were non-significant for OCR 1 (Figure 6C; Table 5), although we note a small effect of sex (ω^2^ = 0.062) and ELS (ω^2^ = 0.083) on basal respiration (Table 5). In the adult liver, there were significant medium effects of ELS on OCR in the presence of ADP (ω^2^ = 0.124) and FCCP (ω^2^ = 0.122; Figure 6D, Table 5), with OCR lower following ELS (ADP, control: 80.15 [95% CI: 62.80, 97.50], ELS: 64.11 [95% CI: 52.93, 75.29]; FCCP, control: 84.30 [95% CI: 66.39, 102.21], ELS: 67.01 [95% CI: 54.40, 79.61]). There was also a small effect of sex (ω^2^ = 0.019) on basal OCR (Table 5). All other main effects and interactions were non-significant (Table 5). The findings in liver are consistent with mitochondria being a common cellular target of ELS centrally and peripherally.

**Figure 6.**
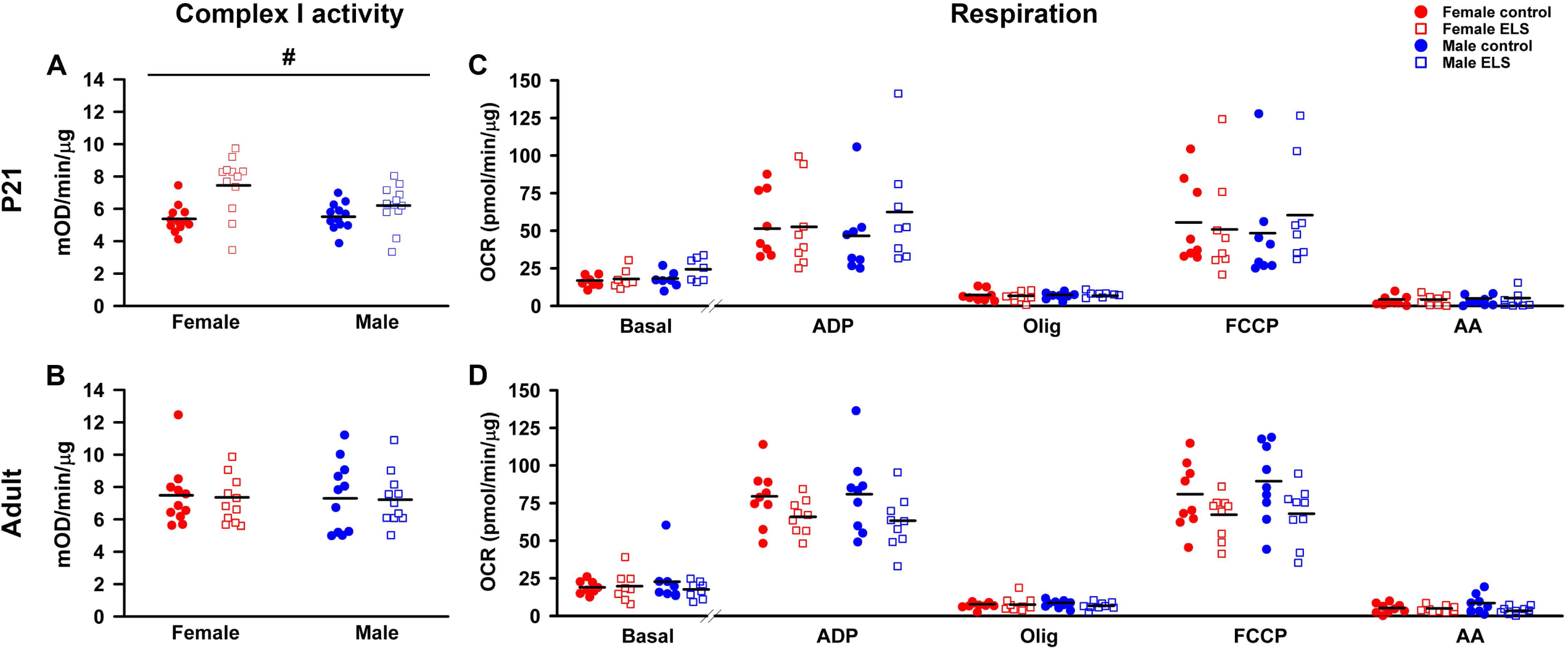
Alterations in mitochondrial function in liver following ELS. Complex I activity (A, B) and oxygen consumption rate (OCR) driven through complex II-IV (C, D) were measured in mitochondria isolated from P21 (top panels) or adult liver (bottom panels). ELS increases complex I activity in males and females at P21 (A), but there is no change in activity in the adult (B). (C, D) OCR was measured under basal conditions or in sister wells following the sequential addition of adenosine diphosphate (ADP), oligomycin (olig), FCCP and antimycin A (AA). There was no effect of sex or ELS, or sex*ELS interaction on any measure in juveniles and adults. ^#^main effect of ELS by two-way ANOVA. Details of statistical analyses and sample sizes are provided in Tables 3 and 5. The raw data contributing to these figures are found in Supplementary Tables S12-S15. Each symbol represents an independent sample, with the bar representing the group mean.

**Table 4.**
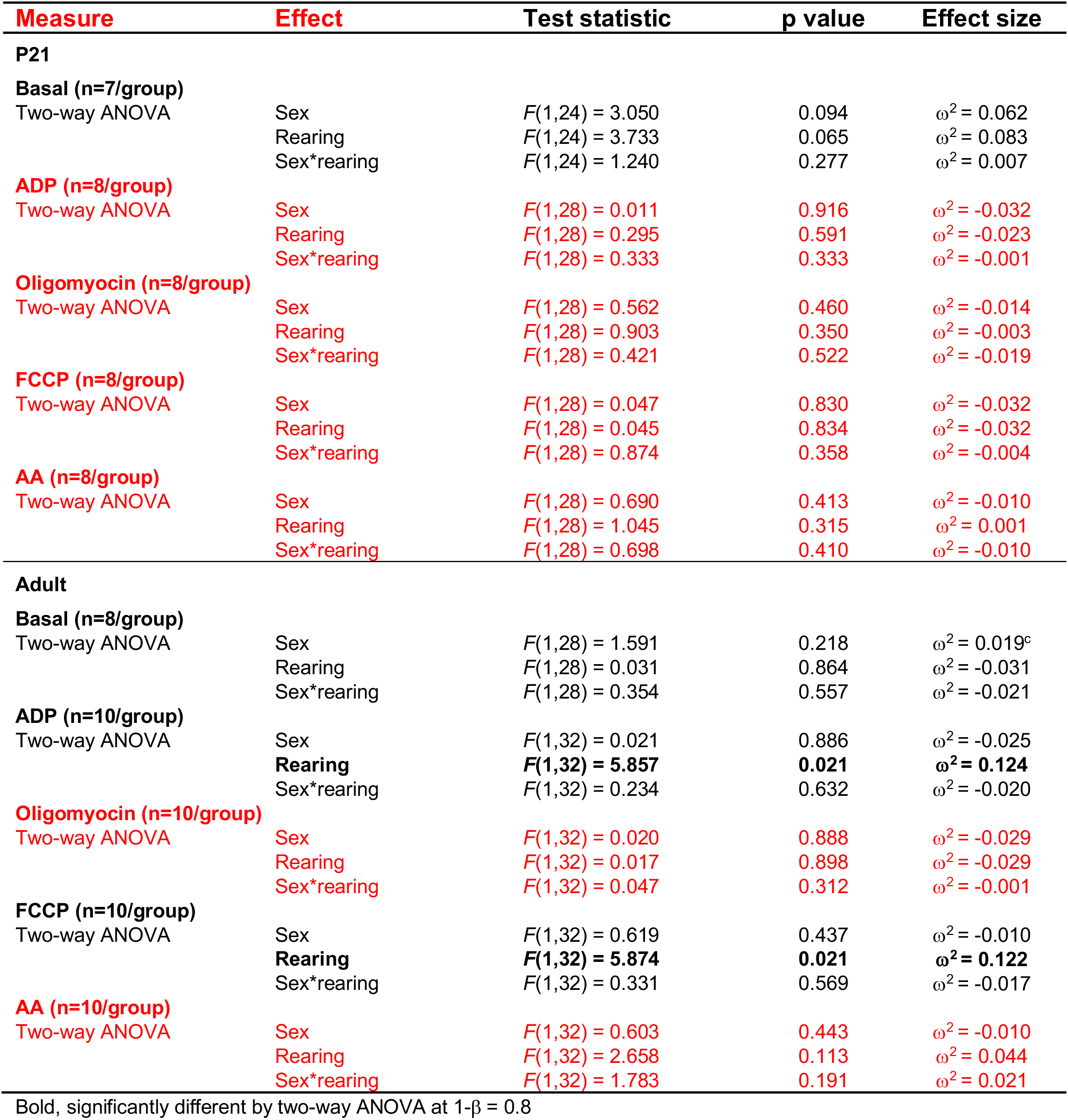
Statistical test details of liver respiration measurements^a^

## 4. DISCUSSION

A central hypothesis of the impact of stress on health implicates disruption of energetics through altered mitochondrial function (Picard et al., 2018). This has been investigated mostly in adult models of stress. The present study, incorporating a model of ELS with an unbiased discovery strategy and functional assays, revealed a substantial impact on the hippocampal proteome as a whole, as well as large-medium effects on mitochondrial respiration and protein composition. We hypothesize that these adaptations contribute to the well-known cellular and circuit dysfunctions caused by ELS. Importantly, the developmental and adult analyses demonstrated distinct changes following ELS in juveniles and adults. The adaptations displayed three temporal patterns. First, certain functional measures were increased in juveniles but then flipped, decreasing in adults. Hippocampal neurogenesis and cognition exhibit similar reciprocal responses following maternal separation, with increased hippocampal neurogenesis and enhanced spatial learning at P21 and decreased neurogenesis and deficits in spatial memory in middle-aged adult rats (Suri et al., 2013). The patterns are thought to represent initial advantageous adaptations to ELS, which are not sustained long term. Second, adaptations of the proteome observed in juveniles became more pronounced in adults, reflected in the increased number of differentially-expressed proteins in the adult hippocampus following ELS. Third, certain ELS-induced changes measured in juveniles were no longer present in adults. For example, many proteins exhibiting altered expression at P21 normalized in adults. Similar transient alterations have been reported at the transcript level, including *Bdnf* in rodent hippocampus following maternal separation (Gross et al., 2012; Kuma et al., 2004). The present data, together with a growing body of literature (for example, (Derks et al., 2016; Goodwill et al., 2019; Loi et al., 2014; Ruiz et al., 2018; Yam et al., 2017b)), highlight the complexity of the biological response to ELS, emphasizing the importance of measuring responses to environmental perturbations across the lifespan.

The design of the iTRAQ proteomics studies facilitated identification of potential sex differences in protein expression in the juvenile and adult hippocampus under different rearing conditions. The studies revealed a marked increase in the number of proteins exhibiting sex differences in control-reared animals between P21 and adults. This likely reflects the influence of gonadal pubertal hormones. This could be addressed in future studies, including determination of the estrus cycle at time of tissue dissection. An increase in differentially-expressed transcripts in the male and female hippocampus after puberty has been reported (Bundy et al., 2017). More dramatically, following exposure to ELS, we observed a large ∼14-fold increase in the number of proteins exhibiting sex differences in expression in the juvenile hippocampus. Such an increase was not observed in the adult; however, the identity of proteins exhibiting differential expression in the adult hippocampus following ELS is different than that in control-reared mice. This discovery emphasizes that while both male and female mice are responsive to disrupted maternal care, there are sex-specific adaptations in the hippocampal proteome.

Increases and decreases in expression for individual proteins following ELS reported here typically were in the 20-30% range, with a striking enrichment in proteins associated with certain cellular compartments and pathways. In adults, proteins associated with the myelin sheath and extracellular exosome were enriched in both sexes. Alterations in myelination have been reported previously in ELS models, including the paradigm used here (Bath et al., 2016; Carlyle et al., 2012; Miki et al., 2014; Yang et al., 2017). In juveniles and adults, there also was an enrichment in mitochondria-associated proteins in the female hippocampus. Further, in adults, almost half of these proteins were associated with the inner mitochondrial membrane, including subunits of the respiratory chain. This was linked functionally, by the mitochondrial respiratory measures, to the enrichment in altered protein expression associated with the OXPHOS pathway. Previous proteomics studies are consistent with our findings, reporting effects of pre- and early post-natal stress on protein expression associated with hippocampal mitochondria and energy metabolism (Focking et al., 2014; Mairesse et al., 2012; Marais et al., 2009; Wei et al., 2015); however, the studies reported data at one age and in one sex, precluding assessment of temporal- and sex-selective adaptive changes across the lifespan. Further, to our knowledge, no direct measures of mitochondrial function have been performed following these early life stressors. Such analyses, as presented here, are critical given the accumulating evidence that mitochondrial dysfunction plays a direct role in central and peripheral disease pathology (Akhter et al., 2017; Ben-Shachar, 2016; Erpapazoglou et al., 2017; Giannoccaro et al., 2017; Patti et al., 2003; Siddiqui et al., 2016).

One of the key observations in the current study was the upregulation in respiratory chain function in juvenile hippocampus, contrasted by a pronounced reduction in function in adults. The upregulation in maximal respiration and complex I activity observed in the juvenile hippocampus may represent an initial positive adaptation to ELS. A recent study using an adolescent stressor drew this same conclusion, and further suggested that the initial adaptive increase would result in an excess of free radical production that then leads to reduced mitochondrial functioning over time (Nold et al., 2019). The mild OXPHOS uncoupling at P21 and the large-medium adult reduction in respiratory chain functioning reported here is consistent with this hypothesis. Given the female-selective alterations in the hippocampal mitochondrial proteome, we were surprised to observe no main effect of sex on hippocampal respiration and complex I measures. We note, however, that the effects of ELS on complex I activity and OCR in the presence of ADP in the adult were moderated by a sex, with significant reductions in females, but not males. We speculate that the relative sparing of mitochondrial function in males may be due to the male-selective up-regulation of the antioxidant superoxide dismutase [Cu-Zn].

Two additional observations in the current study are worth noting. First, the inter-individual differences observed in mitochondrial respiration likely arise through a combination of experimental variability in the measures, including the documented variation in OCR measures by Seahorse across plates and experimental day (Sakamuri 2019; Yepez, 2018; Hubbard, 2018), and by well-known heterogeneous responses of different litters to ELS, as reported using different measures, including behavioral outcomes (for example(Bolton et al., 2018b; Goodwill et al., 2019; Heun-Johnson and Levitt, 2017; McIlwrick et al., 2016). We emphasize, however, that the statistically determined large effect sizes (ω^2^), even in light of such variation, indicate the robustness of the ELS adaptations reported here. Second, central and peripheral mitochondrial function was impacted by ELS in the current model, supporting the framework advanced by Picard and McEwen that links stress to mitochondrial allostatic load across organ systems (Picard and McEwen, 2018). Many measures are affected similarly in hippocampus and liver, but with some notable differences: 1) in the adult, ELS has no effect on complex I activity in the liver, but reduces activity in the hippocampus, and 2) at P21, maximal OXPHOS capacity when respiration is driven through complex II is increased in the hippocampus, but unaltered in liver. Future studies can address the biological significance of these differences, as well as identifying the response of mitochondria in other stress-sensitive brain areas and peripheral organ systems.

There are some limitations to the current study. First, the iTRAQ proteomics method readily detects more abundantly expressed proteins. It is possible that low abundance proteins also are altered in response to ELS. Second, outcomes of the limited-bedding paradigm have been attributed primarily to unpredictable fragmented maternal care during the stress period combined with subsequent alterations in the pup HPA axis (Davis et al., 2017; Heun-Johnson and Levitt, 2016; Naninck et al., 2015; Rice et al., 2008; Wang et al., 2011a). It is possible that a combination of factors during the stress period influence pup outcome, including mild cold exposure and reported altered micronutritional intake (Naninck et al., 2017; Yam et al., 2015). We note, however, that in the ELS paradigm used here, core body temperature and utilization of brown adipose tissue does not vary across rearing conditions suggesting that the pups do not experience hypothermia (Bolton et al., 2019).

Among the most important discoveries of the current study are the adaptive mitochondrial functional changes that support the conclusion of Suri et al (Suri et al., 2013). They state that in response to ELS “potentially adaptive outcomes shift to deleterious consequences” in the hippocampus later in life. Based on the mounting epidemiologic evidence of the powerful impact that ELS has on lifespan health outcomes, greater insight will derive from an emphasis on understanding the changing developmental context in which adaptations to ELS occur. Such data will contribute to a more complete understanding of biological adaptations that may be responsive to therapeutic interventions applied differently across the lifespan.

## Supporting information

Supplemental results

## ACKNOWLEDGEMENTS

We acknowledge Dr. Kristie Lindsey Rose, Salisha Hill and Amanda Hachey (Mass Spectrometry Research Center (MSRC) Proteomics Core at the Vanderbilt University School of Medicine) for performing the iTRAQ protocols and generating the iTRAQ datasets, Hemal H. Mehta (USC Leonard Davis School of Gerontology Seahorse Core Facility) for assistance with the Seahorse assay and Drs. Allison Knoll and Lisa Schlueter for critical reading of the manuscript.

Funding: This work was supported by The JPB Foundation through a grant to The JPB Research Network on Toxic Stress: A Project of the Center on the Developing Child at Harvard University (grant agreement #1025), the Simms/Mann Chair in Developmental Neurogenetics, the WM Keck Chair in Neurogenetics (P.L.), and the The Saban Research Institute Program in Developmental Neuroscience and Neurogenetics.

All authors report no biomedical financial interests or potential conflicts of interest.

