## Supplemental results for "Proteomic and mitochondrial adaptations to early-life stress are distinct in juveniles and adults"

There was no difference in dam and pup body weight at P2, the start of the ELS period (Supplemental Figures S2A, D; Supplemental Tables S1, S2). Rearing had no effect on body weight gain in dams over the stress period (Supplemental Figure S2B; Supplemental Table S1). In contrast, there was a large effect of rearing ($F(1,188) = 49.91$, $p < .001$, $\omega^2 = .191$), but no effect of sex or sex*rearing interaction, on weight gain in pups over the same period (Supplemental Figure S2E; Supplemental Table S2). ELS pups weighed ~7% less than control pups at P9 (Supplemental Figure S2F; Supplemental Table S2). We also measured offspring weight at P21 and adults, the ages at which our analyses were performed. There was a small effect of rearing ($F(1,188) = 8.67$, $p = .004$, $\omega^2 = .038$), but no effect of sex or sex*rearing interaction, on weight gain between P9 and P21 (Supplemental Figure S2G; Supplemental Table S2); thus ELS pups still weighed ~7% less than control pups at P21 (Supplemental Figure S2H; Supplemental Table S2). By adulthood, there is a large effect of sex ($F(1,92) = 305.426$, $p < .001$, $\omega^2 = .763$), but no effect of rearing or sex*rearing interaction, on offspring weight (Supplemental Figure S2I; Supplemental Table S2), consistent with previous reports (1-4).

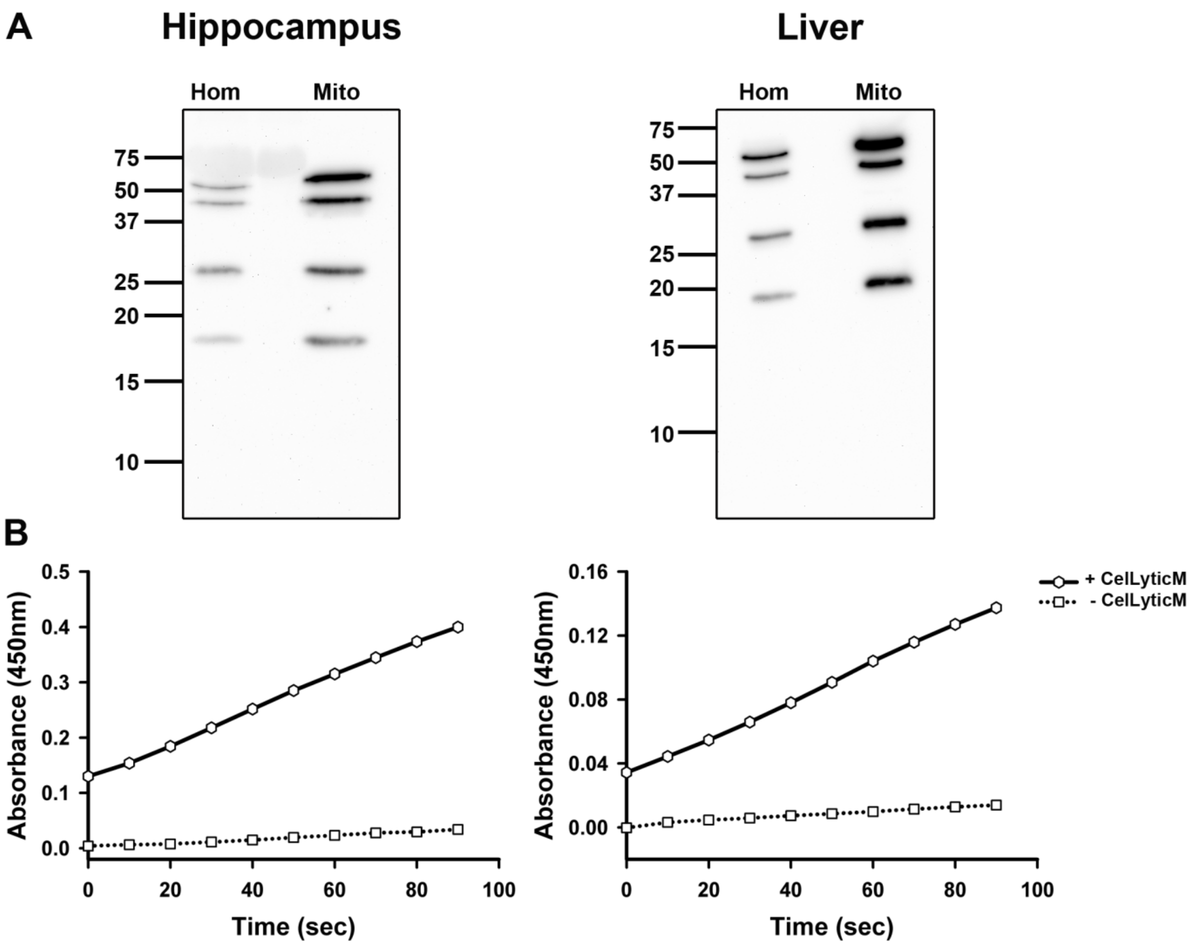

**Figure S1.** Characterization of the isolated mitochondrial preparation in pilot studies. **(A)** Western blot analysis confirms enrichment of OXPHOS complexes in isolated mitochondria (Mito) compared to non-nuclear homogenates (Hom) of hippocampus and liver. **(B)** Mitochondrial integrity was confirmed by measuring citrate synthase activity in the presence and absence of CellLytic M, which lyses the mitochondria. Only lysed mitochondria display CS activity.

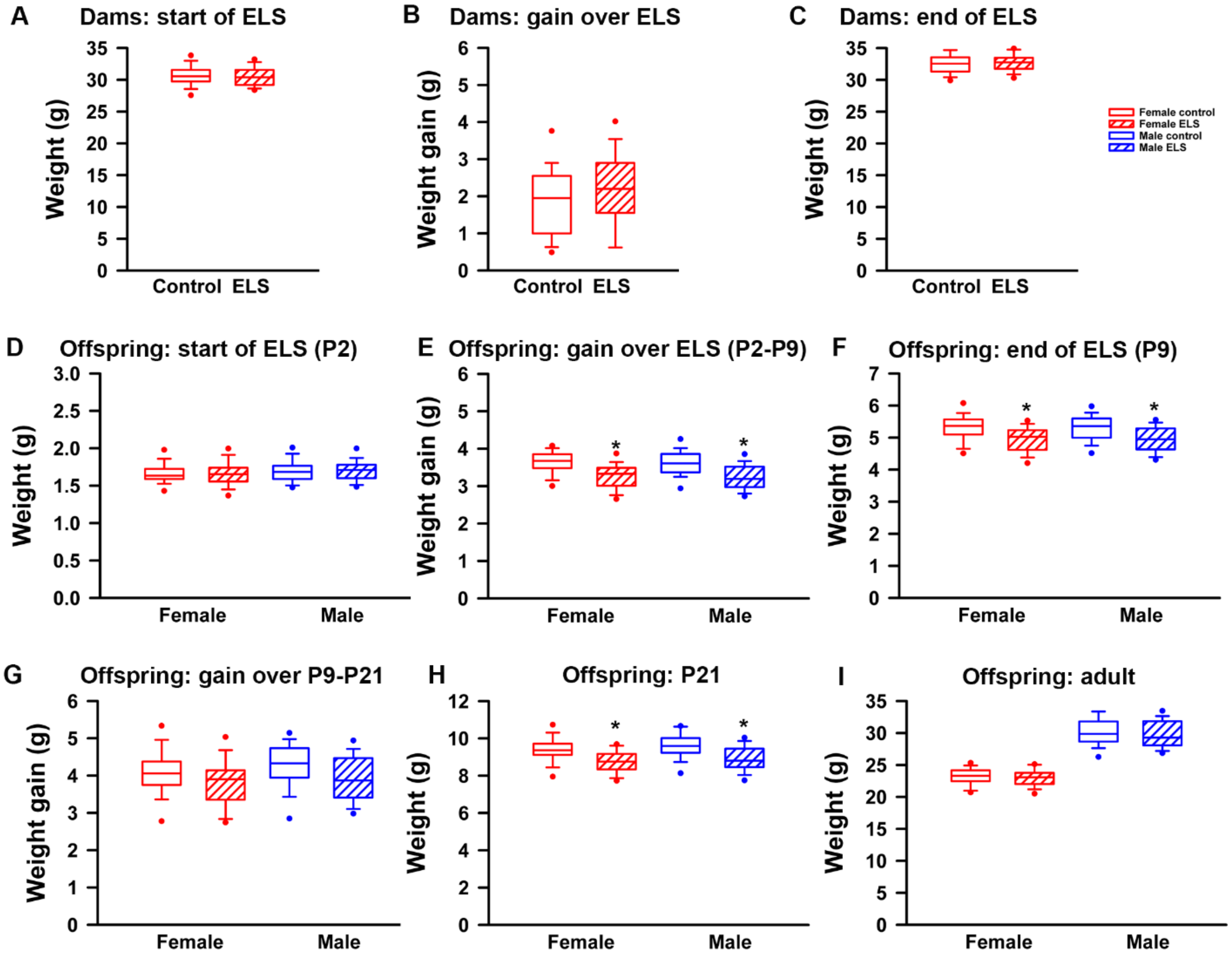

**Figure S2.** Effects of ELS on body weight gain and total body weight during the stress period and at the age of analysis. For dams, there is no difference in **(A)** body weight at the start of the ELS period, **(B)** body weight gain over the ELS period, or **(C)** body weight at the end of the ELS period. For offspring, there is no difference in body weight at the start of the ELS period **(D)**. Body weight gain over the ELS period is reduced in ELS males and females **(E)**, but returns to control-reared levels after the stress period ends **(G)**. Male and female ELS pups weigh less than control-reared pups at the end of the ELS period **(D)** and at weaning **(E)**.  Weights have normalized by adulthood **(F)**.

Data are presented as box and whisker plots. The box represents the 25$^{th}$ and 75 percentiles, the whiskers denote the 10$^{th}$ and 90$^{th}$ percentiles, and the dots represent the 5$^{th}$ and 95$^{th}$ percentiles. The line bisecting each box is the median. *Significantly different from control-reared mice of the same sex. Raw data and statistical details are provided in Supplementary Tables S1, S2.

### Table S1. Body weight (g) of control and ELS dams

| Control (n=48) | | | ELS (n=48) | | |
|---|---|---|---|---|---|
| P2 | P9 | P9-P2 | P2 | P9 | P9-P2 |
| 30.4 | 31.4 | 1.0 | 31.0 | 32.8 | 1.8 |
| 30.8 | 32.9 | 2.1 | 33.1 | 34.6 | 1.5 |
| 29.9 | 32.8 | 2.9 | 29.7 | 32.4 | 2.7 |
| 31.6 | 33.4 | 1.8 | 28.7 | 32.0 | 3.3 |
| 27.1 | 30.3 | 3.2 | 32.3 | 34.8 | 2.5 |
| 30.9 | 33.5 | 2.6 | 28.4 | 32.1 | 3.7 |
| 28.9 | 31.3 | 2.4 | 29.2 | 30.4 | 1.2 |
| 28.8 | 30.7 | 1.9 | 30.4 | 33.5 | 3.1 |
| 32.4 | 33.0 | 0.6 | 30.0 | 33.4 | 3.4 |
| 30.6 | 31.6 | 1.0 | 31.8 | 33.1 | 1.3 |
| 32.0 | 32.5 | 0.5 | 30.7 | 33.3 | 2.6 |
| 27.6 | 28.1 | 0.5 | 32.9 | 34.7 | 1.8 |
| 31.5 | 31.8 | 0.3 | 30.4 | 34.9 | 4.5 |
| 25.1 | 26.2 | 1.1 | 30.2 | 31.8 | 1.6 |
| 32.5 | 34.7 | 2.2 | 29.2 | 29.2 | 0.0 |
| 28.9 | 32.6 | 3.7 | 33.2 | 35.4 | 2.2 |
| 31.1 | 33.6 | 2.5 | 33.6 | 33.1 | -0.5 |
| 33.2 | 35.4 | 2.2 | 30.8 | 32.7 | 1.9 |
| 29.9 | 30.3 | 0.4 | 28.8 | 30.3 | 1.5 |
| 28.0 | 30.1 | 2.1 | 31.8 | 35.2 | 3.4 |
| 33.6 | 34.6 | 1.0 | 31.3 | 33.2 | 1.9 |
| 31.5 | 33.4 | 1.9 | 31.8 | 31.5 | -0.3 |
| 30.8 | 33.7 | 2.9 | 33.3 | 34.9 | 1.6 |
| 30.1 | 32.9 | 2.8 | 29.2 | 32.1 | 2.9 |
| 32.3 | 33.9 | 1.6 | 31.8 | 34.2 | 2.4 |
| 30.3 | 31.3 | 1.0 | 29.4 | 32.1 | 2.7 |
| 30.3 | 31.5 | 1.2 | 30.3 | 32.9 | 2.6 |
| 33.7 | 34.6 | 0.9 | 29.2 | 32.1 | 2.9 |
| 30.8 | 32.2 | 1.4 | 28.8 | 32.8 | 4.0 |
| 31.1 | 33.2 | 2.1 | 31.3 | 33.3 | 2.0 |
| 32.4 | 34.3 | 1.9 | 31.2 | 31.3 | 0.1 |
| 29.3 | 31.9 | 2.6 | 30.3 | 33.5 | 3.2 |
| 29.8 | 32.1 | 2.3 | 31.2 | 32.2 | 1.0 |
| 29.9 | 31.9 | 2.0 | 30.7 | 31.7 | 1.0 |

| | | | | | | |
|---|---|---|---|---|---|---|
| | 31.3 | 32.9 | 1.6 | 29.9 | 31.9 | 2.0 |
| | 29.7 | 32.5 | 2.8 | 31.9 | 32.8 | 0.9 |
| | 30.3 | 31.0 | 0.7 | 30.9 | 33.1 | 2.2 |
| | 34.9 | 35.7 | 0.8 | 32.5 | 33.0 | 0.5 |
| | 32.2 | 34.8 | 2.6 | 28.6 | 30.8 | 2.2 |
| | 30.6 | 31.7 | 1.1 | 28.5 | 31.0 | 2.5 |
| | 30.6 | 31.3 | 0.7 | 30.0 | 31.7 | 1.7 |
| | 35.4 | 39.8 | 4.4 | 28.1 | 30.2 | 2.1 |
| | 28.5 | 30.7 | 2.2 | 29.5 | 31.6 | 2.1 |
| | 30.5 | 33.3 | 2.8 | 29.0 | 31.3 | 2.3 |
| | 28.7 | 31.1 | 2.4 | 28.3 | 32.5 | 4.2 |
| | 30.2 | 32.6 | 2.4 | 30.6 | 34.2 | 3.6 |
| | 30.1 | 31.2 | 1.1 | 29.4 | 32.7 | 3.3 |
| | 29.3 | 33.6 | 4.3 | 30.8 | 33.6 | 2.8 |
| **Mean** | **30.61** | **32.50** | **1.89** | **30.50** | **32.66** | **2.16** |
| **Standard deviation** | **1.91** | **2.07** | **0.99** | **1.47** | **1.40** | **1.14** |
| **95% CI** | **30.07, 31.15** | **32.01, 33.19** | **1.61, 2.17** | **30.08, 30.92** | **32.26, 33.06** | **1.84, 2.48** |

### Two-tailed unpaired *t* -test

**P2**

| | Mean Differer | DF | t-Value | P-Value |
|---|---|---|---|---|
| C, ELS | 0.113 | 94 | 0.324 | 0.747 |

**P9**

| | Mean Differer | DF | t-Value | P-Value |
|---|---|---|---|---|
| C, ELS | 0.167 | 94 | -0.462 | 0.645 |

**P9-P2**

| | Mean Differer | DF | t-Value | P-Value |
|---|---|---|---|---|
| C, ELS | 0.279 | 94 | -1.283 | 0.203 |

**Table S2. Body weight (g) of control and ELS offspring**

ND, not determined because litter used for P21 studies

| | Control | | | | | | | | | | | |
|---|---|---|---|---|---|---|---|---|---|---|---|---|
| | Male | | | | | | Female | | | | | |
| | **P2** | **P9** | **P9-P2** | **P21** | **P21-P9** | **Adult** | **P2** | **P9** | **P9-P2** | **P21** | **P21-P9** | **Adult** |
| | 1.73 | 5.44 | 3.71 | 10.20 | 4.76 | ND | 1.72 | 5.40 | 3.68 | 9.00 | 3.60 | ND |
| | 1.59 | 4.83 | 3.24 | 9.23 | 4.40 | ND | 1.65 | 5.07 | 3.42 | 8.60 | 3.53 | ND |
| | 2.00 | 5.71 | 3.71 | 9.59 | 3.88 | ND | 2.07 | 5.92 | 3.85 | 9.78 | 3.86 | ND |
| | 1.54 | 4.73 | 3.19 | 9.38 | 4.65 | 28.73 | 1.56 | 4.70 | 3.14 | 9.25 | 4.55 | 23.59 |
| | 1.57 | 3.87 | 2.30 | 7.83 | 3.96 | 31.63 | 1.60 | 3.80 | 2.20 | 7.97 | 4.17 | 24.23 |
| | 1.56 | 5.44 | 3.88 | 10.00 | 4.56 | 28.59 | 1.43 | 5.19 | 3.76 | 9.52 | 4.33 | 20.98 |
| | 1.70 | 5.43 | 3.73 | 9.65 | 4.22 | ND | 1.62 | 5.13 | 3.51 | 9.36 | 4.23 | ND |
| | 1.69 | 5.35 | 3.66 | 9.87 | 4.52 | ND | 1.71 | 5.37 | 3.66 | 8.68 | 3.31 | ND |
| | 1.73 | 5.28 | 3.55 | 10.10 | 4.82 | 28.82 | 1.69 | 5.70 | 4.01 | 9.65 | 3.95 | 20.83 |
| | 1.71 | 5.26 | 3.55 | 9.73 | 4.47 | 31.31 | 1.69 | 5.26 | 3.57 | 9.35 | 4.09 | 23.51 |
| | 2.01 | 6.60 | 4.59 | 9.53 | 2.93 | ND | 1.82 | 7.09 | 5.27 | 8.05 | 0.96 | ND |
| | 1.69 | 5.08 | 3.39 | 11.63 | 6.55 | ND | 1.63 | 5.15 | 3.52 | 10.80 | 5.65 | ND |
| | 2.09 | 5.37 | 3.28 | 9.97 | 4.60 | 36.38 | 1.97 | 5.53 | 3.56 | 10.10 | 4.57 | 23.19 |
| | 1.44 | 4.35 | 2.91 | 10.06 | 5.71 | 32.00 | 1.59 | 4.60 | 3.01 | 9.90 | 5.30 | 24.44 |
| | 1.65 | 5.58 | 3.93 | 8.92 | 3.34 | 28.49 | 1.58 | 5.53 | 3.95 | 9.11 | 3.58 | 25.31 |
| | 1.59 | 4.53 | 2.94 | 9.60 | 5.07 | ND | 1.59 | 4.54 | 2.95 | 9.51 | 4.97 | ND |
| | 1.51 | 4.88 | 3.37 | 9.19 | 4.31 | ND | 1.50 | 5.05 | 3.55 | 9.29 | 4.24 | ND |
| | 1.73 | 5.60 | 3.87 | 10.03 | 4.43 | 27.76 | 1.67 | 5.50 | 3.83 | 9.50 | 4.00 | 22.47 |
| | 1.70 | 5.68 | 3.98 | 10.00 | 4.32 | 32.09 | 1.73 | 5.66 | 3.93 | 9.47 | 3.81 | 24.46 |
| | 1.66 | 5.14 | 3.48 | 9.33 | 4.19 | ND | 1.63 | 5.24 | 3.61 | 9.17 | 3.93 | ND |
| | 1.93 | 6.47 | 4.54 | 9.33 | 2.86 | ND | 1.79 | 5.65 | 3.86 | 9.35 | 3.70 | ND |
| | 1.92 | 5.78 | 3.86 | 10.63 | 4.85 | 29.01 | 1.81 | 5.79 | 3.98 | 10.73 | 4.94 | 21.12 |
| | 1.71 | 5.54 | 3.83 | 9.47 | 3.93 | 29.76 | 1.57 | 5.38 | 3.81 | 9.95 | 4.57 | 22.45 |
| | 1.68 | 5.58 | 3.90 | 9.37 | 3.79 | 29.95 | 1.59 | 5.61 | 4.02 | 9.55 | 3.94 | 23.82 |
| | 1.66 | 5.60 | 3.94 | 10.63 | 5.03 | 26.43 | 1.68 | 5.51 | 3.83 | 9.03 | 3.52 | 22.95 |
| | 1.59 | 5.17 | 3.58 | 9.37 | 4.20 | 25.88 | 1.59 | 5.30 | 3.71 | 9.13 | 3.83 | 20.41 |
| | 1.29 | 4.57 | 3.28 | 9.43 | 4.86 | ND | 1.20 | 4.21 | 3.01 | 9.46 | 5.25 | ND |
| | 1.61 | 5.68 | 4.07 | 9.67 | 3.99 | ND | 1.62 | 5.66 | 4.04 | 10.00 | 4.34 | ND |
| | 1.50 | 5.09 | 3.59 | 9.43 | 4.34 | ND | 1.52 | 5.15 | 3.63 | 10.97 | 5.82 | ND |
| | 1.67 | 5.41 | 3.74 | 8.17 | 2.76 | ND | 1.68 | 5.41 | 3.73 | 7.63 | 2.22 | ND |
| | 1.55 | 5.04 | 3.49 | 9.23 | 4.19 | 28.47 | 1.59 | 5.33 | 3.74 | 9.13 | 3.80 | 23.00 |
| | 1.88 | 5.66 | 3.78 | 10.37 | 4.71 | 31.03 | 1.86 | 5.72 | 3.86 | 9.77 | 4.05 | 21.93 |
| | 1.72 | 5.08 | 3.36 | 8.73 | 3.65 | 28.90 | 1.63 | 5.13 | 3.50 | 9.27 | 4.14 | 23.43 |
| | 1.78 | 5.49 | 3.71 | 10.00 | 4.51 | 33.20 | 1.70 | 5.36 | 3.66 | 9.37 | 4.01 | 23.00 |
| | 1.66 | 5.92 | 4.26 | 11.00 | 5.08 | 34.73 | 1.61 | 5.64 | 4.03 | 9.87 | 4.23 | 25.37 |
| | 1.48 | 4.79 | 3.31 | 9.27 | 4.48 | 33.03 | 1.43 | 4.63 | 3.20 | 9.40 | 4.77 | 23.17 |
| | 1.58 | 5.42 | 3.84 | 9.70 | 4.28 | 30.75 | 1.58 | 5.38 | 3.80 | 9.58 | 4.20 | 24.13 |
| | 1.81 | 5.87 | 4.06 | 10.63 | 4.76 | 29.37 | 1.88 | 6.33 | 4.45 | 10.40 | 4.07 | 24.87 |
| | 2.01 | 5.77 | 3.76 | 10.63 | 4.86 | 30.03 | 2.08 | 6.05 | 3.97 | 10.47 | 4.42 | 23.83 |
| | 1.82 | 5.44 | 3.62 | 10.27 | 4.83 | ND | 1.86 | 5.53 | 3.67 | 9.53 | 4.00 | ND |
| | 1.81 | 5.24 | 3.43 | 9.80 | 4.56 | ND | 1.74 | 5.19 | 3.45 | 9.67 | 4.48 | ND |
| | 1.50 | 4.89 | 3.39 | 9.07 | 4.18 | ND | 1.58 | 5.05 | 3.47 | 9.13 | 4.08 | ND |
| | 1.63 | 4.94 | 3.31 | 7.57 | 2.63 | ND | 1.67 | 4.93 | 3.26 | 7.77 | 2.84 | ND |
| | 1.78 | 5.67 | 3.89 | 9.68 | 4.01 | ND | 1.73 | 5.45 | 3.72 | 9.30 | 3.85 | ND |
| | 1.75 | 5.35 | 3.60 | 9.20 | 3.85 | ND | 1.64 | 4.98 | 3.34 | 8.40 | 3.42 | ND |
| | 1.61 | 4.93 | 3.32 | 8.73 | 3.80 | ND | 1.63 | 5.00 | 3.37 | 8.57 | 3.57 | ND |
| | 1.62 | 5.11 | 3.49 | 9.20 | 4.09 | ND | 1.67 | 5.18 | 3.51 | 9.30 | 4.12 | ND |
| | 1.70 | 4.96 | 3.26 | 8.63 | 3.67 | ND | 1.54 | 5.39 | 3.85 | 8.73 | 3.34 | ND |
| **Mean** | **1.69** | **5.30** | **3.61** | **9.61** | **4.30** | **30.26** | **1.67** | **5.32** | **3.66** | **9.37** | **4.04** | **23.19** |
| **Standard deviation** | **0.16** | **0.49** | **0.40** | **0.76** | **0.71** | **2.49** | **0.15** | **0.52** | **0.44** | **0.72** | **0.79** | **1.38** |
| **95% CI** | **1.64, 1.74** | **5.16, 5.44** | **3.50, 3.72** | **9.40, 9.82** | **4.10, 4.50** | **29.56, 30.96** | **1.63, 1.71** | **5.17, 5.47** | **3.54, 3.78** | **9.17, 9.57** | **3.82, 4.26** | **22.80, 23.58** |

| | ELS | | | | | | | | | | | |
|---|---|---|---|---|---|---|---|---|---|---|---|---|
| | Male | | | | | | Female | | | | | |
| | **P2** | **P9** | **P9-P2** | **P21** | **P21-P9** | **Adult** | **P2** | **P9** | **P9-P2** | **P21** | **P21-P9** | **Adult** |
| | 1.88 | 4.88 | 3.00 | 6.57 | 1.69 | ND | 1.98 | 5.21 | 3.23 | 7.80 | 2.59 | ND |
| | 2.07 | 5.74 | 3.67 | 8.73 | 2.99 | ND | 1.89 | 5.52 | 3.63 | 8.30 | 2.78 | ND |
| | 1.66 | 5.33 | 3.67 | 8.75 | 3.42 | ND | 1.52 | 4.85 | 3.33 | 8.8 | 3.95 | ND |
| | 1.49 | 4.30 | 2.81 | 8.80 | 4.50 | 31.29 | 1.45 | 4.05 | 2.60 | 8.70 | 4.65 | 23.56 |
| | 1.61 | 4.67 | 3.06 | 8.87 | 4.20 | 28.28 | 1.51 | 4.45 | 2.94 | 9.15 | 4.70 | 19.80 |
| | 1.38 | 4.06 | 2.68 | 9.85 | 5.79 | 28.18 | 1.37 | 4.06 | 2.69 | 9.48 | 5.42 | 23.38 |
| | 2.34 | 5.40 | 3.06 | 8.81 | 3.41 | ND | 2.31 | 5.10 | 2.79 | 8.89 | 3.79 | ND |
| | 1.57 | 4.54 | 2.97 | 9.76 | 5.22 | ND | 1.53 | 4.60 | 3.07 | 9.69 | 5.09 | ND |
| | 1.74 | 4.78 | 3.04 | 8.93 | 4.15 | 27.39 | 1.62 | 4.89 | 3.27 | 9.10 | 4.21 | 23.53 |
| | 1.77 | 5.44 | 3.67 | 9.93 | 4.49 | 29.01 | 1.68 | 5.29 | 3.61 | 9.35 | 4.06 | 21.23 |
| | 1.51 | 4.44 | 2.93 | 8.07 | 3.63 | ND | 1.57 | 4.90 | 3.33 | 8.40 | 3.50 | ND |
| | 1.75 | 5.15 | 3.40 | 8.53 | 3.38 | ND | 1.79 | 5.18 | 3.39 | 8.40 | 3.22 | ND |
| | 1.84 | 4.37 | 2.53 | 7.90 | 3.53 | 28.00 | 2.06 | 5.70 | 3.64 | 8.55 | 2.85 | 22.97 |
| | 1.64 | 4.39 | 2.75 | 8.92 | 4.53 | 32.56 | 1.57 | 4.98 | 3.41 | 8.71 | 3.73 | 24.89 |
| | 1.51 | 4.31 | 2.80 | 8.72 | 4.41 | 28.12 | 1.62 | 5.09 | 3.47 | 9.19 | 4.10 | 25.17 |
| | 1.54 | 4.71 | 3.17 | 8.64 | 3.93 | ND | 1.32 | 4.40 | 3.08 | 8.54 | 4.14 | ND |
| | 1.61 | 5.54 | 3.93 | 9.22 | 3.68 | ND | 1.62 | 5.59 | 3.97 | 8.42 | 2.83 | ND |
| | 1.79 | 5.40 | 3.61 | 10.03 | 4.63 | 27.69 | 1.73 | 5.38 | 3.65 | 9.63 | 4.25 | 23.55 |
| | 1.76 | 5.07 | 3.31 | 9.73 | 4.66 | 27.22 | 1.72 | 5.20 | 3.48 | 10.23 | 5.03 | 23.98 |
| | 1.85 | 5.70 | 3.85 | 8.97 | 3.27 | ND | 1.99 | 5.27 | 3.28 | 9.40 | 4.13 | ND |
| | 1.76 | 4.74 | 2.98 | 8.40 | 3.66 | ND | 1.69 | 4.56 | 2.87 | 8.50 | 3.94 | ND |
| | 1.72 | 4.94 | 3.22 | 9.03 | 4.09 | 30.31 | 1.77 | 5.12 | 3.35 | 8.90 | 3.78 | 23.45 |
| | 1.72 | 5.19 | 3.47 | 9.03 | 3.84 | 26.93 | 1.70 | 5.05 | 3.35 | 9.20 | 4.15 | 20.81 |
| | 1.71 | 4.62 | 2.91 | 8.07 | 3.45 | 32.80 | 1.58 | 4.52 | 2.94 | 7.77 | 3.25 | 22.68 |
| | 1.91 | 4.64 | 2.73 | 9.07 | 4.43 | 26.67 | 1.86 | 4.47 | 2.61 | 8.27 | 3.80 | 21.64 |
| | 1.70 | 4.78 | 3.08 | 8.03 | 3.25 | 32.39 | 1.62 | 4.37 | 2.75 | 8.03 | 3.66 | 22.33 |
| | 1.71 | 4.73 | 3.02 | 9.40 | 4.67 | ND | 1.75 | 4.78 | 3.03 | 9.10 | 4.32 | ND |
| | 1.52 | 5.00 | 3.48 | 8.54 | 3.54 | ND | 1.44 | 4.69 | 3.25 | 8.60 | 3.91 | ND |
| | 1.66 | 4.46 | 2.80 | 9.20 | 4.74 | ND | 1.50 | 4.41 | 2.91 | 9.15 | 4.74 | ND |
| | 1.61 | 5.13 | 3.52 | 8.17 | 3.04 | ND | 1.64 | 5.02 | 3.38 | 7.33 | 2.31 | ND |
| | 1.80 | 4.86 | 3.06 | 8.67 | 3.81 | 29.53 | 1.76 | 4.87 | 3.11 | 8.37 | 3.50 | 21.7 |
| | 1.83 | 5.48 | 3.65 | 10.33 | 4.85 | 28.43 | 1.77 | 5.30 | 3.53 | 9.70 | 4.40 | 22.65 |
| | 1.72 | 5.15 | 3.43 | 9.53 | 4.38 | 30.17 | 1.68 | 5.05 | 3.37 | 9.17 | 4.12 | 21.63 |
| | 1.75 | 5.20 | 3.45 | 9.50 | 4.30 | 31.83 | 1.66 | 5.29 | 3.63 | 9.67 | 4.38 | 25.03 |
| | 1.72 | 4.78 | 3.06 | 8.60 | 3.82 | 31.77 | 1.65 | 4.64 | 2.99 | 8.67 | 4.03 | 24.1 |
| | 1.62 | 5.33 | 3.71 | 9.63 | 4.30 | 32.63 | 1.69 | 5.19 | 3.50 | 9.03 | 3.84 | 23.13 |
| | 1.62 | 5.53 | 3.91 | 10.10 | 4.57 | 34.97 | 1.48 | 5.46 | 3.98 | 9.57 | 4.11 | 25.1 |
| | 1.99 | 5.25 | 3.26 | 9.70 | 4.45 | 31.87 | 1.92 | 5.25 | 3.33 | 9.17 | 3.92 | 23 |
| | 1.59 | 4.39 | 2.80 | 7.80 | 3.41 | 29.03 | 1.59 | 4.25 | 2.66 | 7.77 | 3.52 | 22.4 |
| | 1.64 | 5.16 | 3.52 | 8.33 | 3.17 | ND | 1.55 | 5.41 | 3.86 | 8.17 | 2.76 | ND |
| | 1.43 | 4.56 | 3.13 | 8.17 | 3.61 | ND | 1.34 | 4.88 | 3.54 | 8.03 | 3.15 | ND |
| | 1.50 | 4.41 | 2.91 | 7.30 | 2.89 | ND | 1.45 | 4.22 | 2.77 | 7.17 | 2.95 | ND |
| | 1.82 | 5.43 | 3.61 | 8.67 | 3.24 | ND | 1.72 | 5.44 | 3.72 | 8.90 | 3.46 | ND |
| | 1.71 | 4.96 | 3.25 | 9.87 | 4.91 | ND | 1.72 | 5.05 | 3.33 | 9.07 | 4.02 | ND |
| | 1.76 | 5.22 | 3.46 | 8.30 | 3.08 | ND | 1.68 | 4.90 | 3.22 | 8.07 | 3.17 | ND |
| | 1.56 | 4.70 | 3.14 | 8.60 | 3.90 | ND | 1.61 | 5.03 | 3.42 | 8.93 | 3.90 | ND |
| | 1.84 | 5.35 | 3.51 | 8.67 | 3.32 | ND | 1.70 | 5.05 | 3.35 | 8.10 | 3.05 | ND |
| | 1.56 | 5.15 | 3.59 | 9.13 | 3.98 | ND | 1.56 | 4.98 | 3.42 | 8.53 | 3.55 | ND |
| **Mean** | **1.70** | **4.95** | **3.24** | **8.87** | **3.92** | **29.88** | **1.67** | **4.94** | **3.27** | **8.74** | **3.81** | **22.99** |
| **Standard deviation** | **0.17** | **0.42** | **0.36** | **0.75** | **0.73** | **2.30** | **0.19** | **0.41** | **0.35** | **0.65** | **0.67** | **1.40** |
| **95% CI** | **1.65, 1.75** | **4.83, 5.07** | **3.14, 3.34** | **8.66, 9.08** | **3.71, 4.13** | **29.23, 30.53** | **1.62, 1.72** | **4.82, 5.06** | **3.17, 3.37** | **8.56, 8.92** | **3.62, 4.00** | **22.59, 23.39** |

**Two-way ANOVA (sex and rearing between subjects factors)**

## P2

| | DF | SS | MS | F-value | P-Value | $h^2$ | $w^2$ |
|---|---|---|---|---|---|---|---|
| **Sex** | 1 | 0.049 | 0.049 | 1.753 | 0.187 | 0.009 | 0.004 |
| **Rearing** | 1 | 0.002 | 0.002 | 0.080 | 0.777 | 0.000 | -0.005 |
| **Sex*Rearing** | 1 | 0.002 | 0.002 | 0.076 | 0.784 | 0.000 | -0.005 |
| **Residual** | 188 | 5.3 | 0.028 | | | | |

## P9

| | DF | SS | MS | F-value | [a]P-Value | $h^2$ | [b]$w^2$ |
|---|---|---|---|---|---|---|---|
| **Sex** | 1 | 0.001 | 0.001 | 0.003 | 0.955 | 0.000 | -0.005 |
| **Rearing** | 1 | 6.623 | 6.623 | 31.041 | **<.001** | 0.142 | **0.137** |
| **Sex*Rearing** | 1 | 0.007 | 0.007 | 0.033 | 0.856 | 0.000 | -0.004 |
| **Residual** | 188 | 40.112 | 0.213 | | | | |

[a]values in red = significant
[b]values in blue = medium effect size

## P2-P9

| | DF | SS | MS | F-value | [a]P-Value | $h^2$ | [b]$w^2$ |
|---|---|---|---|---|---|---|---|
| **Sex** | 1 | 0.062 | 0.062 | 0.412 | 0.522 | 0.002 | -0.002 |
| **Rearing** | 1 | 6.871 | 6.871 | 45.914 | **<.001** | 0.196 | **0.191** |
| **Sex*Rearing** | 1 | 0.001 | 0.001 | 0.009 | 0.923 | 0.000 | -0.004 |
| **Residual** | 188 | 28.132 | 0.150 | | | | |

[a]values in red = significant
[b]values in red = large effect size

## P21

| | DF | SS | MS | F-value | P-Value | $h^2$ | [b]$w^2$ |
|---|---|---|---|---|---|---|---|
| **Sex** | 1 | 1.582 | 1.582 | 3.048 | 0.083 | 0.013 | 0.009 |
| **Rearing** | 1 | 22.229 | 22.229 | 42.825 | **<.001** | 0.183 | **0.178** |
| **Sex*Rearing** | 1 | 0.165 | 0.165 | 0.318 | 0.574 | 0.001 | -0.003 |
| **Residual** | 188 | 97.585 | 0.519 | | | | |

[a]values in red = significant
[b]values in red = large effect size

## P21-P9

| | DF | SS | MS | F-value | [a]P-Value | $h^2$ | [b]$w^2$ |
|---|---|---|---|---|---|---|---|
| **Sex** | 1 | 1.648 | 1.648 | 3.117 | 0.079 | 0.016 | 0.011 |
| **Rearing** | 1 | 4.585 | 4.585 | 8.669 | **0.004** | 0.043 | **0.038** |
| **Sex*Rearing** | 1 | 0.24 | 0.240 | 0.454 | 0.501 | 0.002 | -0.003 |
| **Residual** | 188 | 99.426 | 0.529 | | | | |

[a]values in red = significant
[b]values in green = small effect size

### Adult

| | DF | SS | MS | F-value | [a]P-Value | $h^2$ | [b]$w^2$ |
|---|---|---|---|---|---|---|---|
| **Sex** | 1 | 1170.477 | 1170.477 | 305.426 | **<.001** | 0.767 | **0.763** |
| **Rearing** | 1 | 2.056 | 2.056 | 0.537 | 0.466 | 0.001 | -0.001 |
| **Sex*Rearing** | 1 | 0.210 | 0.210 | 0.055 | 0.815 | 0.000 | -0.002 |
| **Residual** | 92 | 352.570 | 3.832 | | | | |

[a]values in red = significant
[b]values in red = large effect size

**Table S3. Proteins detected by iTRAQ at P21**

| Detected in all experiments - 2236 proteins | |
|---|---|
| **Protein Name** | **Uniprot ID** |
| [Protein ADP-ribosylarginine] hydrolase | P54923 |
| [Pyruvate dehydrogenase [lipoamide]] kinase isoz | Q9JK42 |
| 1,5-anhydro-D-fructose reductase | Q9DCT1 |
| 10 kDa heat shock protein, mitochondrial | Q64433 |
| 14 kDa phosphohistidine phosphatase | Q9DAK9 |
| 14-3-3 protein beta/alpha | Q9CQV8 |
| 14-3-3 protein epsilon | P62259 |
| 14-3-3 protein eta | P68510 |
| 14-3-3 protein gamma | P61982 |
| 14-3-3 protein theta | P68254 |
| 14-3-3 protein zeta/delta | P63101 |
| 182 kDa tankyrase-1-binding protein | P58871 |
| 1-acyl-sn-glycerol-3-phosphate acyltransferase g | Q9D517 |
| 1-phosphatidylinositol 3-phosphate 5-kinase | Q9Z1T6 |
| 1-phosphatidylinositol 4,5-bisphosphate phospho | Q9Z1B3 |
| 1-phosphatidylinositol 4,5-bisphosphate phospho | Q8R3B1 |
| 1-phosphatidylinositol 4,5-bisphosphate phospho | A2AP18 |
| 1-phosphatidylinositol 4,5-bisphosphate phospho | Q62077 |
| 2',3'-cyclic-nucleotide 3'-phosphodiesterase | P16330 |
| 26S protease regulatory subunit 10B | P62334 |
| 26S protease regulatory subunit 4 | P62192 |
| 26S protease regulatory subunit 6A | O88685 |
| 26S protease regulatory subunit 6B | P54775 |
| 26S protease regulatory subunit 7 | P46471 |
| 26S protease regulatory subunit 8 | P62196 |
| 26S proteasome non-ATPase regulatory subunit | Q3TXS7 |
| 26S proteasome non-ATPase regulatory subunit | Q8BG32 |
| 26S proteasome non-ATPase regulatory subunit | Q9D8W5 |
| 26S proteasome non-ATPase regulatory subunit | Q9WVJ2 |
| 26S proteasome non-ATPase regulatory subunit | Q8VDM4 |
| 26S proteasome non-ATPase regulatory subunit | P14685 |
| 26S proteasome non-ATPase regulatory subunit | O35226 |
| 26S proteasome non-ATPase regulatory subunit | Q8BJY1 |
| 26S proteasome non-ATPase regulatory subunit | Q99JI4 |
| 26S proteasome non-ATPase regulatory subunit | P26516 |
| 26S proteasome non-ATPase regulatory subunit | Q9CX56 |
| 26S proteasome non-ATPase regulatory subunit | Q9CR00 |
| 28 kDa heat- and acid-stable phosphoprotein | Q3UHX2 |
| 28S ribosomal protein S27, mitochondrial | Q8BK72 |
| 28S ribosomal protein S31, mitochondrial | Q61733 |
| 2-aminoethanethiol dioxygenase | Q6PDY2 |
| 2-oxoglutarate dehydrogenase, mitochondrial | Q60597 |
| 3'(2'),5'-bisphosphate nucleotidase 1 | Q9Z0S1 |
| 39S ribosomal protein L1, mitochondrial | Q99N96 |
| 39S ribosomal protein L12, mitochondrial | Q9DB15 |
| 39S ribosomal protein L39, mitochondrial | Q9JKF7 |
| 39S ribosomal protein L46, mitochondrial | Q9EQI8 |
| 3-hydroxyacyl-CoA dehydrogenase type-2 | O08756 |
| 3-hydroxyisobutyrate dehydrogenase, mitochond | Q99L13 |
| 3-hydroxyisobutyryl-CoA hydrolase, mitochondria | Q8QZS1 |
| 3-ketoacyl-CoA thiolase, mitochondrial | Q8BWT1 |
| 3-mercaptopyruvate sulfurtransferase | Q99J99 |
| 3-phosphoinositide-dependent protein kinase 1 | Q9Z2A0 |
| 40S ribosomal protein S10 | P63325 |
| 40S ribosomal protein S11 | P62281 |
| 40S ribosomal protein S12 | P63323 |
| 40S ribosomal protein S13 | P62301 |
| 40S ribosomal protein S15a | P62245 |
| 40S ribosomal protein S16 | P14131 |
| 40S ribosomal protein S17 | P63276 |
| 40S ribosomal protein S18 | P62270 |
| 40S ribosomal protein S19 | Q9CZX8 |
| 40S ribosomal protein S2 | P25444 |
| 40S ribosomal protein S20 | P60867 |
| 40S ribosomal protein S21 | Q9CQR2 |
| 40S ribosomal protein S24 | P62849 |
| 40S ribosomal protein S25 | P62852 |
| 40S ribosomal protein S26 | P62855 |
| 40S ribosomal protein S27 | Q6ZWU9 |
| 40S ribosomal protein S28 | P62858 |
| 40S ribosomal protein S3 | P62908 |
| 40S ribosomal protein S3a | P97351 |
| 40S ribosomal protein S4, X isoform | P62702 |
| 40S ribosomal protein S5 | P97461 |
| 40S ribosomal protein S6 | P62754 |
| 40S ribosomal protein S7 | P62082 |
| 40S ribosomal protein S8 | P62242 |
| 40S ribosomal protein S9 | Q6ZWN5 |
| 40S ribosomal protein SA | P14206 |
| 4-aminobutyrate aminotransferase, mitochondrial | P61922 |
| 4F2 cell-surface antigen heavy chain | P10852 |
| 4-trimethylaminobutyraldehyde dehydrogenase | Q9JLJ2 |
| 5'(3')-deoxyribonucleotidase, cytosolic type | Q9JM14 |
| 5'-AMP-activated protein kinase catalytic subunit | Q5EG47 |
| 5'-AMP-activated protein kinase catalytic subunit | Q8BRK8 |
| 5'-AMP-activated protein kinase subunit gamma- | O54950 |
| 5'-AMP-activated protein kinase subunit gamma- | Q91WG5 |

| Detected in 2 experiments - 980 proteins | |
|---|---|
| **Protein Name** | **Uniprot ID** |
| **Detected in experiments 1&2 - 168 proteins** | |
| 1-phosphatidylinositol 4,5-bisphosphate phospho | Q8CIH5 |
| 28S ribosomal protein S26, mitochondrial | Q80ZS3 |
| 39S ribosomal protein L9, mitochondrial | Q99N94 |
| 40S ribosomal protein S14 | P62264 |
| 40S ribosomal protein S27-like | Q6ZWY3 |
| 5-azacytidine-induced protein 1 | Q62036 |
| 60S ribosomal protein L35a | O55142 |
| Acidic leucine-rich nuclear phosphoprotein 32 fan | Q9EST5 |
| Actin-related protein 2/3 complex subunit 5-like p | Q9D898 |
| Acyl-CoA-binding domain-containing protein 6 | Q9D061 |
| Acyl-coenzyme A thioesterase 2, mitochondrial | Q9QYR9 |
| Adrenodoxin-like protein, mitochondrial | Q9CPW2 |
| Aftiphilin | Q80WT5 |
| Alpha-1-antitrypsin 1-4 | Q00897 |
| Apolipoprotein A-IV | P06728 |
| Apoptosis regulator BAX | Q07813 |
| Aquaporin-4 | P55088 |
| Aromatic-L-amino-acid decarboxylase | O88533 |
| Aspartate--tRNA ligase, mitochondrial | Q8BIP0 |
| Biorientation of chromosomes in cell division prot | Q5SQY2 |
| Brain-specific angiogenesis inhibitor 3 | Q80ZF8 |
| BTB/POZ domain-containing protein KCTD8 | Q50H33 |
| Calcium/calmodulin-dependent protein kinase typ | Q91YS8 |
| cAMP-dependent protein kinase type I-beta regul | P12849 |
| cAMP-regulated phosphoprotein 21 | Q9DCB4 |
| cAMP-specific 3',5'-cyclic phosphodiesterase 4D | Q01063 |
| Caytaxin | Q8BHE3 |
| Centrosomal protein of 290 kDa | Q6A078 |
| Charged multivesicular body protein 1a | Q921W0 |
| Charged multivesicular body protein 2a | Q9DB34 |
| Coiled-coil and C2 domain-containing protein 1B | Q8BRN9 |
| Coiled-coil domain-containing protein 124 | Q9D8X2 |
| Collagen type IV alpha-3-binding protein | Q9EQG9 |
| Conserved oligomeric Golgi complex subunit 5 | Q8C0L8 |
| Conserved oligomeric Golgi complex subunit 7 | Q3UM29 |
| Cytohesin-1 | Q9QX11 |
| DBF4-type zinc finger-containing protein 2 homol | Q5SS00 |
| Dehydrogenase/reductase SDR family member 7 | Q99J47 |
| Dipeptidyl peptidase 8 | Q80YA7 |
| Disheveled-associated activator of morphogenesi | Q80U19 |
| DnaJ homolog subfamily B member 2 | Q9QYI5 |
| Dynactin subunit 5 | Q9QZB9 |
| E3 ubiquitin-protein ligase RNF181 | Q9CY62 |
| E3 ubiquitin-protein ligase RNF31 | Q924T7 |
| Elongator complex protein 4 | Q9ER73 |
| Enhancer of yellow 2 transcription factor homolog | Q9JIX0 |
| Epoxide hydrolase 2 | P34914 |
| Espin-like protein | Q3UYR4 |
| Exocyst complex component 6 | Q8R313 |
| FGFR1 oncogene partner 2 homolog | Q9CRA9 |
| FK506-binding protein 15 | Q6P9Q6 |
| GA-binding protein alpha chain | Q00422 |
| Gamma-tubulin complex component 2 | Q921G8 |
| Gamma-tubulin complex component 3 | P58854 |
| General transcription factor II-I | Q9ESZ8 |
| Glia maturation factor beta | Q9CQI3 |
| Glucosamine--fructose-6-phosphate aminotransfe | P47856 |
| Glucose-fructose oxidoreductase domain-contain | Q3UHD2 |
| Glypican-5 | Q8CAL5 |
| GSK3-beta interaction protein | Q8BGR8 |
| GTPase NRas | P08556 |
| GTP-binding protein SAR1a | P36536 |
| Guanine nucleotide-binding protein G(s) subunit a | Q6R0H7 |
| Histidine triad nucleotide-binding protein 2, mitoc | Q9D0S9 |
| Histone H3.1 | P68433 |
| Inactive phospholipase C-like protein 1 | Q3USB7 |
| Isoamyl acetate-hydrolyzing esterase 1 homolog | Q9DB29 |
| L-2-hydroxyglutarate dehydrogenase, mitochondr | Q91YP0 |
| Legumain | O89017 |
| Lipopolysaccharide-responsive and beige-like an | Q9ESE1 |
| Luc7-like protein 3 | Q5SUF2 |
| Lysosome membrane protein 2 | O35114 |
| Metabotropic glutamate receptor 2 | Q14BI2 |
| Metaxin-1 | P47802 |
| Methylcrotonoyl-CoA carboxylase subunit alpha, | Q99MR8 |
| Methylthioribose-1-phosphate isomerase | Q9CQT1 |
| Microtubule-associated proteins 1A/1B light chain | Q91VR7 |
| MIP18 family protein FAM96A | Q9DCL2 |
| Mitochondrial carnitine/acylcarnitine carrier protei | Q9Z2Z6 |
| Molybdopterin synthase sulfur carrier subunit | Q9Z224 |
| Myotrophin | P62774 |
| N-alpha-acetyltransferase 38, NatC auxiliary subu | Q6ZWM4 |
| Nuclear fragile X mental retardation-interacting pr | Q5F2E7 |
| Nuclear receptor-binding protein | Q99J45 |
| OTU domain-containing protein 6B | Q8K2H2 |
| Oxysterol-binding protein 2 | Q5QNQ6 |

| Detected in 1 experiment - 1230 proteins | |
|---|---|
| **Protein Name** | **Uniprot ID** |
| **Detected in experiment 1 only - 187 proteins** | |
| 1,4-alpha-glucan-branching enzyme | Q9D6Y9 |
| 1-phosphatidylinositol 4,5-bisphosphate phosphoc | Q4KWH5 |
| 39S ribosomal protein L23, mitochondrial | O35972 |
| Acyl-coenzyme A thioesterase 4 | Q8BWN8 |
| Adenylate cyclase type 1 | O88444 |
| Adenylosuccinate synthetase isozyme 1 | P28650 |
| A-kinase anchor protein 8-like | Q9R0L7 |
| Ankyrin repeat domain-containing protein 13A | Q80UP5 |
| Ankyrin repeat domain-containing protein 27 | Q3UMR0 |
| Annexin A2 | P07356 |
| Apolipoprotein D | P51910 |
| ATP-binding cassette sub-family B member 6, mit | Q9DC29 |
| Axonemal dynein light chain domain-containing p | Q3UZ57 |
| Baculoviral IAP repeat-containing protein 1b | Q9QUK4 |
| Baculoviral IAP repeat-containing protein 1f | Q9JIB6 |
| Beta-taxilin | Q8VBT1 |
| Bromodomain and WD repeat-containing protein | Q921C3 |
| Cadherin-23 | Q99PF4 |
| Calpastatin | P51125 |
| Carboxylesterase 1C | P23953 |
| Caspase recruitment domain-containing protein 1 | Q99KF0 |
| Centriolar coiled-coil protein of 110 kDa | Q7TSH4 |
| Centrosomal protein of 112 kDa | Q5PR68 |
| Chromatin modification-related protein MEAF6 | Q2VPQ9 |
| Class E basic helix-loop-helix protein 40 | O35185 |
| Coiled-coil domain-containing protein 51 | Q3URS9 |
| Coiled-coil domain-containing protein 57 | Q6PHN1 |
| Coiled-coil domain-containing protein 88B | Q4QRL3 |
| COMM domain-containing protein 9 | Q8K2Q0 |
| Conserved oligomeric Golgi complex subunit 1 | Q9Z160 |
| Conserved oligomeric Golgi complex subunit 6 | Q8R3I3 |
| Creatine kinase S-type, mitochondrial | Q6P8J7 |
| Cullin-associated NEDD8-dissociated protein 2 | Q6ZQ73 |
| DENN domain-containing protein 5B | A2RSQ0 |
| Dimethylaniline monooxygenase [N-oxide-forming | Q8VHG0 |
| Diphosphoinositol polyphosphate phosphohydrola | Q8R2U6 |
| DNA polymerase epsilon catalytic subunit A | Q9WVF7 |
| DNA repair protein XRCC2 | Q9CX47 |
| DNA topoisomerase 2-alpha | Q01320 |
| DNA2-like helicase | Q6ZQJ5 |
| DnaJ homolog subfamily B member 12 | Q9QYI4 |
| Dynein heavy chain 2, axonemal | P0C6F1 |
| Dynein light chain roadblock-type 1 | P62627 |
| E3 ubiquitin-protein ligase BRE1A | Q5DTM8 |
| E3 ubiquitin-protein ligase RNF123 | Q5XPI3 |
| E3 ubiquitin-protein ligase UBR1 | O70481 |
| Ethanolamine kinase 1 (Fragment) | Q9D4V0 |
| Eukaryotic translation initiation factor 2-alpha kina | Q9Z2B5 |
| Eukaryotic translation initiation factor 4E-binding | Q80VV3 |
| Eukaryotic translation initiation factor 6 | O55135 |
| F-box only protein 28 | Q8BIG4 |
| Fibrinogen gamma chain | Q8VCM7 |
| Forkhead box protein K1 | P42128 |
| Formin-binding protein 1-like | Q8K012 |
| FUN14 domain-containing protein 2 | Q9D6K8 |
| Gamma-aminobutyric acid receptor-associated pr | Q8R3R8 |
| Ganglioside GM2 activator | Q60648 |
| General transcription factor IIF subunit 2 | Q8R0A0 |
| Glutamate receptor delta-1 subunit | Q61627 |
| Glutamate receptor, ionotropic kainate 5 | Q61626 |
| GMP reductase 2 | Q99L27 |
| G-protein coupled receptor family C group 5 mem | Q923Z0 |
| Group XV phospholipase A2 | Q8VEB4 |
| Growth hormone-inducible transmembrane protei | Q91VC9 |
| Growth/differentiation factor 2 | Q9WV56 |
| HCLS1-binding protein 3 | Q3TC93 |
| Histone acetyltransferase KAT6B | Q8BRB7 |
| Histone lysine demethylase PHF8 | Q80TJ7 |
| Histone-binding protein RBBP7 | Q60973 |
| Inositol hexakisphosphate kinase 1 | Q6PD10 |
| Inositol-tetrakisphosphate 1-kinase | Q8BYN3 |
| Interferon regulatory factor 2-binding protein-like | Q8K3X4 |
| Interferon-induced guanylate-binding protein 2 | Q9Z0E6 |
| Intraflagellar transport protein 88 homolog | Q61371 |
| Kinesin-like protein KIF2B | Q8C0N1 |
| Laminin subunit alpha-3 | Q61789 |
| Laminin subunit alpha-4 | P97927 |
| Leucine-rich repeat and guanylate kinase domain | Q9D5S7 |
| Leucine-rich repeat-containing protein 20 | Q8CI70 |
| Leucine-rich repeat-containing protein 8D | Q8BGR2 |
| Low-density lipoprotein receptor-related protein 1 | Q8CB67 |
| Low-density lipoprotein receptor-related protein 2 | A2ARV4 |
| MAP6 domain-containing protein 1 | Q14BB9 |
| Membrane transport protein XK | Q9QXY7 |
| Mesencephalic astrocyte-derived neurotrophic fac | Q9CXI5 |
| MICAL-like protein 1 | Q8BGT6 |

| | | | | | |
|---|---|---|---|---|---|
| 5'-nucleotidase domain-containing protein 3 | Q3UHB1 | Pantothenate kinase 4 | Q80YV4 | Microtubule-associated proteins 1A/1B light chain | Q9CQV6 |
| 5-oxoprolinase | Q8K010 | Peptidyl-prolyl cis-trans isomerase-like 1 | Q9D0W5 | Mitochondrial fission process protein 1 | Q9CRB8 |
| 60 kDa heat shock protein, mitochondrial | P63038 | Peroxisomal targeting signal 1 receptor | O09012 | Mitogen-activated protein kinase kinase kinase kin | P97820 |
| 60 kDa SS-A/Ro ribonucleoprotein | O08848 | PH domain leucine-rich repeat-containing protein | Q8CHE4 | MKL/myocardin-like protein 1 | Q8K4J6 |
| 60S acidic ribosomal protein P0 | P14869 | Phosphatidylinositol 4,5-bisphosphate 3-kinase ca | Q8BTI9 | Multimerin-2 | A6H6E2 |
| 60S acidic ribosomal protein P2 | P99027 | Phospholipase DDHD1 | Q80YA3 | Myosin-binding protein C, cardiac-type | O70468 |
| 60S ribosomal protein L10 | Q6ZWV3 | Phospholipid hydroperoxide glutathione peroxidas | O70325 | NACHT, LRR and PYD domains-containing protei | Q8R4B8 |
| 60S ribosomal protein L10a | P53026 | Plexin-C1 | Q9QZC2 | Nicotinate phosphoribosyltransferase | Q8CC86 |
| 60S ribosomal protein L11 | Q9CXW4 | Poly(rC)-binding protein 2 | Q61990 | Nidogen-1 | P10493 |
| 60S ribosomal protein L12 | P35979 | Potassium voltage-gated channel subfamily A me | P16390 | NK-tumor recognition protein | P30415 |
| 60S ribosomal protein L13 | P47963 | Potassium voltage-gated channel subfamily A me | Q61423 | Nucleolar protein 14 | Q8R3N1 |
| 60S ribosomal protein L13a | P19253 | Pre-mRNA branch site protein p14 | P59708 | Peroxisomal N(1)-acetyl-spermine/spermidine oxi | Q8C0L6 |
| 60S ribosomal protein L14 | Q9CR57 | Profilin-1 | P62962 | Peroxisome proliferator-activated receptor gamma | Q8VHJ7 |
| 60S ribosomal protein L15 | Q9CZM2 | Proteasome activator complex subunit 4 | Q5SSW2 | Phosphatidylinositol 4-kinase type 2-alpha | Q2TBE6 |
| 60S ribosomal protein L17 | Q9CPR4 | Proteasome subunit beta type-2 | Q9R1P3 | Phosphoprotein associated with glycosphingolipid | Q3U1F9 |
| 60S ribosomal protein L18 | P35980 | Protein 4.1 | P48193 | Pleckstrin homology domain-containing family A m | Q8BUL6 |
| 60S ribosomal protein L18a | P62717 | Protein canopy homolog 3 | Q9DAU1 | Poliovirus receptor-related protein 2 | P32507 |
| 60S ribosomal protein L19 | P84099 | Protein cereblon | Q8C7D2 | Poliovirus receptor-related protein 3 | Q9JLB9 |
| 60S ribosomal protein L21 | O09167 | Protein lin-7 homolog B | O88951 | Poly(A)-specific ribonuclease PARN | Q8VDG3 |
| 60S ribosomal protein L22 | P67984 | Protein LSM12 homolog | Q9D0R8 | Potassium voltage-gated channel subfamily D me | Q9Z0V2 |
| 60S ribosomal protein L23a | P62751 | Protein LYRIC | Q80WJ7 | Pre-mRNA-processing factor 39 | Q8K2Z2 |
| 60S ribosomal protein L26 | P61255 | Protein OS-9 | Q8K2C7 | Probable ATP-dependent RNA helicase DDX28 | Q9CWT6 |
| 60S ribosomal protein L27 | P61358 | Protein phosphatase Slingshot homolog 2 | Q5SW75 | Probable cysteine--tRNA ligase, mitochondrial | Q8BYM8 |
| 60S ribosomal protein L27a | P14115 | Protein quaking | Q9QYS9 | Prolyl 3-hydroxylase 2 | Q8CG71 |
| 60S ribosomal protein L28 | P41105 | Protocadherin-19 | Q80TF3 | Protein asunder homolog | Q8QZV7 |
| 60S ribosomal protein L3 | P27659 | Pterin-4-alpha-carbinolamine dehydratase | P61458 | Protein C10 | O35127 |
| 60S ribosomal protein L30 | P62889 | Putative peptidyl-tRNA hydrolase PTRHD1 | D3Z4S3 | Protein CIP2A | Q8BWY9 |
| 60S ribosomal protein L31 | P62900 | Quinone oxidoreductase | P47199 | Protein dpy-30 homolog | Q99LT0 |
| 60S ribosomal protein L35 | Q6ZWV7 | Rab-3A-interacting protein | Q68EF0 | Protein ERGIC-53 | Q9D0F3 |
| 60S ribosomal protein L38 | Q9JJI8 | Ras-related GTP-binding protein B | Q6NTA4 | Protein FAM203A | Q8C3I8 |
| 60S ribosomal protein L4 | Q9D8E6 | Ras-related protein Rab-8B | P61028 | Protein FAM214A | Q69ZK7 |
| 60S ribosomal protein L5 | P47962 | Ras-related protein Ral-B | Q9JIW9 | Protein FAM55B | Q3U095 |
| 60S ribosomal protein L6 | P47911 | Receptor expression-enhancing protein 1 | Q8BGH4 | Protein FAM63B | Q6PDI6 |
| 60S ribosomal protein L7 | P14148 | Regulation of nuclear pre-mRNA domain-containi | Q8VDS4 | Protein IWS1 homolog | Q8C1D8 |
| 60S ribosomal protein L7a | P12970 | Reticulocalbin-1 | Q05186 | Protein KIBRA | Q5SXA9 |
| 60S ribosomal protein L8 | P62918 | Retinal-specific ATP-binding cassette transporter | O35600 | Protein kinase C delta type | P28867 |
| 60S ribosomal protein L9 | P51410 | Rho GTPase-activating protein 26 | Q6ZQ82 | Protein kinase C zeta type | Q02956 |
| 6-phosphofructokinase type C | Q9WUA3 | Rho GTPase-activating protein 33 | Q80YF9 | Protein midA homolog, mitochondrial | Q9CWG8 |
| 6-phosphofructokinase, liver type | P12382 | Ribosomal protein S6 kinase alpha-2 | Q9WUT3 | Protein NipSnap homolog 2 | O55126 |
| 6-phosphofructokinase, muscle type | P47857 | RNA-binding protein 12 | Q8R4X3 | Protein phosphatase Slingshot homolog 1 | Q76I79 |
| 6-phosphogluconate dehydrogenase, decarboxyl | Q9DCD0 | Sarcosine dehydrogenase, mitochondrial | Q99LB7 | Protein tyrosine phosphatase type IVA 1 | Q63739 |
| 6-phosphogluconolactonase | Q9CQ60 | Selenium-binding protein 2 | Q63836 | Putative RNA polymerase II subunit B1 CTD phos | Q8VC34 |
| 78 kDa glucose-regulated protein | P20029 | Serine/arginine-rich splicing factor 3 | P84104 | Rab9 effector protein with kelch motifs | Q8VCH5 |
| Abelson tyrosine-protein kinase 2 | Q4JIM5 | Serine/threonine-protein kinase B-raf | P28028 | Rap guanine nucleotide exchange factor-like 1 | Q68EF8 |
| Abhydrolase domain-containing protein 16A | Q9Z1Q2 | Serine/threonine-protein kinase Nek7 | Q9ES74 | Ras association domain-containing protein 5 | Q5EBH1 |
| Abl interactor 1 | Q8CBW3 | Serine/threonine-protein kinase TAO2 | Q6ZQ29 | Ras association domain-containing protein 8 | Q8CJ96 |
| Abl interactor 2 | P62484 | Serine/threonine-protein phosphatase 4 regulator | Q8C0Y0 | Rho guanine nucleotide exchange factor 18 | Q6P9R4 |
| Acetoacetyl-CoA synthetase | Q9D2R0 | SID1 transmembrane family member 2 | Q8CIF6 | Rho guanine nucleotide exchange factor 33 | Q8BW86 |
| Acetyl-CoA acetyltransferase, cytosolic | Q8CAY6 | Signal peptidase complex catalytic subunit SEC1 | Q9R0P6 | Rho guanine nucleotide exchange factor 6 | Q8K4I3 |
| Acetyl-CoA acetyltransferase, mitochondrial | Q8QZT1 | Sister chromatid cohesion protein PDS5 homolog | Q4VA53 | Ribonuclease ZC3H12A | Q5D1E7 |
| Acetyl-CoA carboxylase 1 | Q5SWU9 | Small VCP/p97-interacting protein | Q3UZP4 | RING finger protein 157 | Q3TEL6 |
| Acetyl-coenzyme A synthetase 2-like, mitochondr | Q99NB1 | S-methyl-5'-thioadenosine phosphorylase | Q9CQ65 | Ryanodine receptor 1 | E9PZQ0 |
| Acetyl-coenzyme A synthetase, cytoplasmic | Q9QXG4 | SNARE-associated protein Snapin | Q9Z266 | Secreted frizzled-related sequence protein 4 | Q9Z1N6 |
| Acidic leucine-rich nuclear phosphoprotein 32 fan | O35381 | Solute carrier family 2, facilitated glucose transpo | P17809 | Separin | P60330 |
| Aconitate hydratase, mitochondrial | Q99KI0 | Spastin | Q9QYY8 | Serine protease inhibitor A3F | Q80X76 |
| Actin, aortic smooth muscle | P62737 | Spermatogenesis-associated protein 7 homolog | Q80VP2 | Serine/arginine-rich splicing factor 9 | Q9D0B0 |
| Actin, cytoplasmic 2 | P63260 | Sterile alpha and TIR motif-containing protein 1 | Q6PDS3 | Serine/threonine-protein kinase 25 | Q9Z2W1 |
| Actin-binding LIM protein 1 | Q8K4G5 | Succinate dehydrogenase assembly factor 1, mito | Q3U276 | Serine/threonine-protein kinase 31 | Q99MW1 |
| Actin-binding LIM protein 2 | Q8BL65 | Synapse-associated protein 1 | Q9D5V6 | Serine/threonine-protein kinase MST4 | Q99JT2 |
| Actin-related protein 10 | Q9QZB7 | TBC1 domain family member 15 | Q9CXF4 | Serine/threonine-protein kinase tousled-like 1 | Q8C0V0 |
| Actin-related protein 2 | P61161 | Terminal uridylyltransferase 7 | Q5BLK4 | Serine/threonine-protein phosphatase 4 regulator | Q8K2V1 |
| Actin-related protein 2/3 complex subunit 1A | Q9R0Q6 | TGF-beta-activated kinase 1 and MAP3K7-bindin | Q8CF89 | Serine/threonine-protein phosphatase 6 regulator | Q505D1 |
| Actin-related protein 2/3 complex subunit 2 | Q9CVB6 | Thioredoxin reductase 2, mitochondrial | Q9JLT4 | Signal peptidase complex subunit 3 | Q9D365 |
| Actin-related protein 2/3 complex subunit 3 | Q9JM76 | Transcription factor E3 | Q64092 | Signal transducer and activator of transcription 2 | Q9WVL2 |
| Actin-related protein 2/3 complex subunit 4 | P59999 | Transcription factor HIVEP2 | Q3UHF7 | Signal transducer and activator of transcription 3 | P42227 |
| Actin-related protein 3 | Q99JY9 | Transmembrane protein 2 | Q5FWI3 | Sphingosine kinase 2 | Q9JIA7 |
| Actin-related protein 3B | Q641P0 | Transmembrane protein 65 | Q4VAE3 | StAR-related lipid transfer protein 7, mitochondria | Q8R1R3 |
| Activated RNA polymerase II transcriptional coact | P11031 | Transportin-2 | Q99LG2 | Synaptogyrin-1 | O55100 |
| Activator of 90 kDa heat shock protein ATPase h | Q8BK64 | tRNA (cytosine(34)-C(5))-methyltransferase | Q1HFZ0 | TBC1 domain family member 10A | P58802 |
| Active breakpoint cluster region-related protein | Q5SSL4 | TSC22 domain family protein 3 | Q9Z2S7 | T-complex protein 11-like protein 2 | Q8K1H7 |
| Acyl carrier protein, mitochondrial | Q9CR21 | Tubulin polymerization-promoting protein family m | Q9CRB6 | Telomerase protein component 1 | P97499 |
| Acylamino-acid-releasing enzyme | Q8R146 | Tubulin--tyrosine ligase | P38585 | Tetraspanin-7 | Q62283 |
| Acyl-CoA dehydrogenase family member 9, mitoc | Q8JZN5 | Ubiquitin carboxyl-terminal hydrolase 11 | Q99K46 | Tetratricopeptide repeat protein 39B | Q8BYY4 |
| Acyl-CoA synthetase family member 2, mitochond | Q8VCW8 | Ubiquitin-conjugating enzyme E2 variant 1 | Q9CZY3 | Tetratricopeptide repeat protein 9B | Q9D6E4 |
| Acyl-CoA-binding protein | P31786 | UDP-N-acetylhexosamine pyrophosphorylase-like | Q3TW96 | TOX high mobility group box family member 4 | Q8BU11 |
| Acyl-coenzyme A thioesterase 1 | O55137 | Uncharacterized membrane protein C1orf95 hom | Q0VBF8 | Transcription initiation factor IIA subunit 1 | Q99PM3 |
| Acyl-coenzyme A thioesterase 13 | Q9CQR4 | Uncharacterized protein C18orf25 homolog | Q8BH50 | Transmembrane and coiled-coil domains protein 3 | Q8R310 |
| Acyl-coenzyme A thioesterase 9, mitochondrial | Q9R0X4 | Uncharacterized protein C6orf174 homolog | Q6NZL0 | Transmembrane protein 163 | Q8C996 |
| Acylglycerol kinase, mitochondrial | Q9ESW4 | UPF0505 protein C16orf62 homolog | Q8BWQ6 | TREMBL:A2A5Y0 Tax_Id=10090 Gene_Symbol= | Fragment\|Cl |
| Acylphosphatase-1 | P56376 | Vacuolar protein sorting-associated protein 28 ho | Q9D1C8 | tRNA dimethylallyltransferase, mitochondrial | Q80UN9 |
| Acylphosphatase-2 | P56375 | Vacuolar protein sorting-associated protein VTA1 | Q9CR26 | Tubulin alpha-1B chain | P05213 |
| Acyl-protein thioesterase 2 | Q9WTL7 | Vacuolar-sorting protein SNF8 | Q9CZ28 | Ubiquitin carboxyl-terminal hydrolase 12 | Q9D9M2 |
| Adapter molecule crk | Q64010 | **Detected in experiments 1&3 - 132 proteins** | | Ubiquitin domain-containing protein 2 | Q6PGH0 |
| Adaptin ear-binding coat-associated protein 1 | Q9CR95 | 26S proteasome non-ATPase regulatory subunit | O35593 | Ubiquitin thioesterase OTUB2 | Q9CQX0 |
| Adenomatous polyposis coli protein | Q61315 | 28S ribosomal protein S5, mitochondrial | Q99N87 | Ubiquitin-associated domain-containing protein 2 | Q8R1K1 |
| Adenosine kinase | P55264 | 39S ribosomal protein L37, mitochondrial | Q921S7 | Ubiquitin-fold modifier 1 | P61961 |
| Adenosylhomocysteinase | P50247 | Abnormal spindle-like microcephaly-associated pr | Q8CJ27 | Uncharacterized protein C3orf26 homolog | Q9CZT6 |
| Adenylate cyclase type 9 | P51830 | ADP-ribosylation factor 6 | P62331 | Uncharacterized protein C4orf29 homolog | Q8C1A9 |
| Adenylate kinase 2, mitochondrial | Q9WTP6 | Alanine aminotransferase 1 | Q8QZR5 | Unconventional myosin-Vb | P21271 |
| Adenylate kinase isoenzyme 1 | Q9R0Y5 | Amyloid beta A4 precursor protein-binding family | Q9QXJ1 | UPF0235 protein C15orf40 homolog | Q9CRC3 |

| | |
|---|---|
| Adenylate kinase isoenzyme 4, mitochondrial | Q9WUR9 |
| Adenylate kinase isoenzyme 5 | Q920P5 |
| Adenylosuccinate lyase | P54822 |
| Adenylosuccinate synthetase isozyme 2 | P46664 |
| Adenylyl cyclase-associated protein 1 | P40124 |
| Adenylyl cyclase-associated protein 2 | Q9CYT6 |
| Adipocyte plasma membrane-associated protein | Q9D7N9 |
| ADP/ATP translocase 1 | P48962 |
| ADP/ATP translocase 2 | P51881 |
| ADP/ATP translocase 4 | Q3V132 |
| ADP-ribose pyrophosphatase, mitochondrial | Q8BVU5 |
| ADP-ribosylation factor 2 | Q8BSL7 |
| ADP-ribosylation factor 3 | P61205 |
| ADP-ribosylation factor 4 | P61750 |
| ADP-ribosylation factor 5 | P84084 |
| ADP-ribosylation factor GTPase-activating protei | Q9EPJ9 |
| ADP-ribosylation factor-binding protein GGA1 | Q8R0H9 |
| ADP-ribosylation factor-binding protein GGA3 | Q8BMI3 |
| ADP-ribosylation factor-like protein 3 | Q9WUL7 |
| ADP-ribosylation factor-like protein 8A | Q8VEH3 |
| ADP-ribosylation factor-like protein 8B | Q9CQW2 |
| Afadin | Q9QZQ1 |
| AFG3-like protein 2 | Q8JZQ2 |
| Aflatoxin B1 aldehyde reductase member 2 | Q8CG76 |
| AH receptor-interacting protein | O08915 |
| A-kinase anchor protein 12 | Q9WTQ5 |
| A-kinase anchor protein 5 | D3YVF0 |
| A-kinase anchor protein 9 | Q70FJ1 |
| A-kinase anchor protein SPHKAP | Q6NSW3 |
| Alanine--tRNA ligase, cytoplasmic | Q8BGQ7 |
| Alanyl-tRNA editing protein Aarsd1 | Q3THG9 |
| Alcohol dehydrogenase [NADP(+)] | Q9JII6 |
| Alcohol dehydrogenase class-3 | P28474 |
| Aldehyde dehydrogenase X, mitochondrial | Q9CZS1 |
| Aldehyde dehydrogenase, mitochondrial | P47738 |
| Aldose reductase | P45376 |
| Alpha-1,3/1,6-mannosyltransferase ALG2 | Q9DBE8 |
| Alpha-1-syntrophin | Q61234 |
| Alpha-2-HS-glycoprotein | P29699 |
| Alpha-2-macroglobulin | Q61838 |
| Alpha-2-macroglobulin receptor-associated protei | P55302 |
| Alpha-actinin-1 | Q7TPR4 |
| Alpha-actinin-2 | Q9JI91 |
| Alpha-actinin-4 | P57780 |
| Alpha-adducin | Q9QYC0 |
| Alpha-aminoadipic semialdehyde dehydrogenase | Q9DBF1 |
| Alpha-centractin | P61164 |
| Alpha-endosulfine | P60840 |
| Alpha-enolase | P17182 |
| Alpha-internexin | P46660 |
| Alpha-ketoglutarate-dependent dioxygenase FTC | Q8BGW1 |
| Alpha-mannosidase 2C1 | Q91W89 |
| Alpha-soluble NSF attachment protein | Q9DB05 |
| Alpha-synuclein | O55042 |
| Alpha-taxilin | Q6PAM1 |
| Alpha-tubulin N-acetyltransferase | Q8K341 |
| Alsin | Q920R0 |
| Amine oxidase [flavin-containing] A | Q64133 |
| Amine oxidase [flavin-containing] B | Q8BW75 |
| Aminoacyl tRNA synthase complex-interacting mu | P31230 |
| Aminoacyl tRNA synthase complex-interacting mu | Q8R010 |
| Aminoacylase-1 | Q99JW2 |
| Aminopeptidase B | Q8VCT3 |
| AMP deaminase 2 | Q9DBT5 |
| AMP deaminase 3 | O08739 |
| Amphiphysin | Q7TQF7 |
| Amyloid beta A4 precursor protein-binding family | B2RUJ5 |
| Amyloid beta A4 protein | P12023 |
| Amyloid-like protein 1 | Q03157 |
| Anaphase-promoting complex subunit 2 | Q8BZQ7 |
| Ankyrin repeat and FYVE domain-containing prot | Q810B6 |
| Ankyrin repeat and MYND domain-containing pro | Q3TPE9 |
| Ankyrin repeat and sterile alpha motif domain-cor | Q8BIZ1 |
| Ankyrin repeat domain-containing protein 17 | Q99NH0 |
| Ankyrin-1 | Q02357 |
| Ankyrin-2 | Q8C8R3 |
| Annexin A11 | P97384 |
| Annexin A3 | O35639 |
| Annexin A5 | P48036 |
| Annexin A6 | P14824 |
| Annexin A7 | Q07076 |
| AP-1 complex subunit beta-1 | O35643 |
| AP-1 complex subunit gamma-1 | P22892 |
| AP-1 complex subunit mu-1 | P35585 |
| AP-1 complex subunit sigma-1A | P61967 |
| AP-2 complex subunit alpha-1 | P17426 |
| AP-2 complex subunit alpha-2 | P17427 |
| AP-2 complex subunit beta | Q9DBG3 |
| AP-2 complex subunit mu | P84091 |
| AP-2 complex subunit sigma | P62743 |
| Ankyrin repeat domain-containing protein 32 | Q8R3P9 |
| Anoctamin-1 | Q8BHY3 |
| Arf-GAP with GTPase, ANK repeat and PH doma | Q8BXK8 |
| ATP synthase subunit f, mitochondrial | P56135 |
| ATP-binding cassette sub-family F member 3 | Q8K268 |
| ATP-dependent RNA helicase A | O70133 |
| Basic leucine zipper and W2 domain-containing p | Q2L4X1 |
| Bcl-2-like protein 1 | Q64373 |
| Beta-crystallin B1 | Q9WVJ5 |
| Bisphosphoglycerate mutase | P15327 |
| BRI3-binding protein | Q8BXV2 |
| Calmodulin-regulated spectrin-associated protein | A2AHC3 |
| Calponin-3 | Q9DAW9 |
| cAMP-dependent protein kinase inhibitor alpha | P63248 |
| Centrosomal protein of 128 kDa | Q8BI22 |
| Charged multivesicular body protein 1b-1 | Q99LU0 |
| Chromodomain-helicase-DNA-binding protein 8 | Q09XV5 |
| Coiled-coil domain-containing protein 25 | Q78PG9 |
| Complement C1q subcomponent subunit B | P14106 |
| Complement C1q tumor necrosis factor-related pr | Q8R066 |
| Complement regulatory protein Crry | Q64735 |
| Contactin-2 | Q61330 |
| Cullin-4A | Q3TCH7 |
| Cyclin-dependent kinase 17 | Q8K0D0 |
| Cytohesin-3 | O08967 |
| Cytosolic 5'-nucleotidase III-like protein | Q3UFY7 |
| Cytosolic carboxypeptidase 1 | Q641K1 |
| Delta-1-pyrroline-5-carboxylate synthase | Q9Z110 |
| Dipeptidyl peptidase 9 | Q8BVG4 |
| Double zinc ribbon and ankyrin repeat-containing | Q8C008 |
| Dual specificity mitogen-activated protein kinase | Q63932 |
| E3 ubiquitin-protein ligase HECTD3 | Q3U487 |
| E3 ubiquitin-protein ligase RNF14 | Q9JI90 |
| ELAV-like protein 4 | Q61701 |
| Enhancer of rudimentary homolog | P84089 |
| Envoplakin | Q9D952 |
| Epsin-3 | Q91W69 |
| Eukaryotic peptide chain release factor GTP-bind | Q149F3 |
| FAD synthase | Q8R123 |
| Fascin-2 | Q32M02 |
| Fatty acyl-CoA reductase 1 | Q922J9 |
| F-box/SPRY domain-containing protein 1 | Q8K3B1 |
| Fibrous sheath-interacting protein 2 | A2ARZ3 |
| Flavin reductase (NADPH) | Q923D2 |
| G protein-coupled receptor kinase 6 | O70293 |
| Galectin-related protein A | Q8VED9 |
| Gamma-aminobutyric acid receptor subunit alpha | P62812 |
| Gamma-tubulin complex component 4 | Q9D4F8 |
| Glutamate--cysteine ligase regulatory subunit | O09172 |
| Golgi SNAP receptor complex member 2 | O35166 |
| GTP-binding protein 1 | O08582 |
| GTP-binding protein SAR1b | Q9CQC9 |
| Guanine nucleotide-binding protein G(I)/G(S)/G(C | P63213 |
| HBS1-like protein | Q69ZS7 |
| Heterogeneous nuclear ribonucleoprotein H2 | P70333 |
| Histone H2B type 1-P | Q8CGP2 |
| Huntingtin-interacting protein K | Q9CR41 |
| Inosine triphosphate pyrophosphatase | Q9D892 |
| Inosine-5'-monophosphate dehydrogenase 1 | P50096 |
| Intraflagellar transport protein 172 homolog | Q6VH22 |
| Isopentenyl-diphosphate Delta-isomerase 1 | P58044 |
| Katanin p80 WD40-containing subunit B1 | Q8BG40 |
| KH domain-containing, RNA-binding, signal trans | Q60749 |
| KH domain-containing, RNA-binding, signal trans | Q9R226 |
| Kinesin-like protein KIFC2 | O08672 |
| Large proline-rich protein BAG6 | Q9Z1R2 |
| Leucine-rich repeat and fibronectin type III domai | Q2WF71 |
| LIM domain-containing protein 2 | Q8BGB5 |
| Metastasis-associated protein MTA3 | Q924K8 |
| Mitochondrial import inner membrane translocase | Q9D880 |
| Mitotic checkpoint protein BUB3 | Q9WVA3 |
| Mitotic spindle-associated MMXD complex subuni | Q9D187 |
| Myotubularin-related protein 9 | Q9Z2D0 |
| N-acetylglucosamine-6-sulfatase | Q8BFR4 |
| NADH dehydrogenase [ubiquinone] flavoprotein 3 | Q8BK30 |
| NHS-like protein 1 | Q8CAF4 |
| Osteoclast-stimulating factor 1 | Q62422 |
| Phosphatidylinositol 3,4,5-trisphosphate 3-phospl | O08586 |
| Phosphatidylinositol 3-kinase regulatory subunit b | O08908 |
| Polyubiquitin-C | P0CG50 |
| Potassium/sodium hyperpolarization-activated cyc | O88703 |
| Potassium-transporting ATPase alpha chain 2 | Q9Z1W8 |
| Pre-mRNA-processing-splicing factor 8 | Q99PV0 |
| Probable E3 ubiquitin-protein ligase HERC4 | Q6PAV2 |
| Probable histidine--tRNA ligase, mitochondrial | Q99KK9 |
| Protein EFR3 homolog A | Q8BG67 |
| Protein FAM59A | Q3UFT3 |
| Protein phosphatase 1 regulatory subunit 37 | Q8BKR5 |
| Protein-L-isoaspartate O-methyltransferase doma | P59913 |
| RAF proto-oncogene serine/threonine-protein kin | Q99N57 |
| UPF0258 protein KIAA1024 | Q8K3V7 |
| UPF0368 protein Cxorf26 homolog | Q9D0B6 |
| UPF0515 protein C19orf66 homolog | Q8CAK3 |
| UPF0693 protein C10orf32 homolog | Q9CRC6 |
| Uridine 5'-monophosphate synthase | P13439 |
| Uridine diphosphate glucose pyrophosphatase | Q9D142 |
| Vesicle transport through interaction with t-SNARI | O89116 |
| Villin-1 | Q62468 |
| WD repeat-containing protein 41 | Q3UDP0 |
| X-linked retinitis pigmentosa GTPase regulator | Q9R0X5 |
| Zinc finger protein ZIC 5 | Q7TQ40 |
| **Detected in experiment 2 only - 436 proteins** | |
| [Pyruvate dehydrogenase [lipoamide]] kinase isoz | Q8BFP9 |
| 15 kDa selenoprotein | Q9ERR7 |
| 1-acyl-sn-glycerol-3-phosphate acyltransferase de | Q8K4X7 |
| 28S ribosomal protein S33, mitochondrial | Q9D2R8 |
| 28S ribosomal protein S34, mitochondrial | Q9JIK9 |
| 2-oxoglutarate and iron-dependent oxygenase do | Q3U0K8 |
| 2-oxoisovalerate dehydrogenase subunit alpha, m | P50136 |
| 2-oxoisovalerate dehydrogenase subunit beta, mi | Q6P3A8 |
| 39S ribosomal protein L43, mitochondrial | Q99N89 |
| 39S ribosomal protein L48, mitochondrial | Q8JZS9 |
| 39S ribosomal protein L50, mitochondrial | Q8VDT9 |
| 5'-3' exoribonuclease 1 | P97789 |
| 60S ribosomal protein L23 | P62830 |
| Abhydrolase domain-containing protein 10, mitoch | Q6PE15 |
| Abhydrolase domain-containing protein 11 | Q8K4F5 |
| Acidic leucine-rich nuclear phosphoprotein 32 fam | P97822 |
| Activating signal cointegrator 1 complex subunit 1 | Q9D8Z1 |
| Activator of 90 kDa heat shock protein ATPase ho | Q8N9S3 |
| Acyl-coenzyme A thioesterase 11 | Q8VHQ9 |
| Adenosine deaminase domain-containing protein | Q5SUE7 |
| Adenosine deaminase-like protein | Q80SY6 |
| ADP-ribosylation factor-like protein 1 | P61211 |
| ADP-ribosylation factor-like protein 15 | Q8BGR6 |
| ADP-ribosylation factor-like protein 6 | O88848 |
| ADP-sugar pyrophosphatase | Q9JKX6 |
| AFG3-like protein 1 | Q920A7 |
| A-kinase anchor protein 2 | O54931 |
| A-kinase anchor protein 7 | O55074 |
| Alanine aminotransferase 2 | Q8BGT5 |
| Alanine--tRNA ligase, mitochondrial | Q14CH7 |
| Aldose reductase-related protein 1 | P21300 |
| Allograft inflammatory factor 1 | O70200 |
| Alpha-(1,6)-fucosyltransferase | Q9WTS2 |
| Alpha-1-antitrypsin 1-1 | P07758 |
| Alpha-mannosidase 2x | Q8BRK9 |
| Amyloid beta A4 precursor protein-binding family | Q8R5A3 |
| Amyloid protein-binding protein 2 | Q9DAX9 |
| AN1-type zinc finger protein 5 | O88878 |
| Anion exchange protein 3 | P16283 |
| Ankyrin repeat and BTB/POZ domain-containing p | Q6GQW0 |
| Apolipoprotein O-like | Q78IK4 |
| Apoptosis-stimulating of p53 protein 2 | Q8CG79 |
| Arginase-2, mitochondrial | O08691 |
| Armadillo repeat-containing protein 8 | Q9DBR3 |
| Arsenite methyltransferase | Q91WU5 |
| Aspartoacylase | Q8R3P0 |
| Ataxin-1-like | P0C7T6 |
| ATP-binding cassette sub-family A member 13 | Q5SSE9 |
| ATP-binding cassette sub-family B member 7, mit | Q61102 |
| ATP-binding cassette sub-family G member 2 | Q7TMS5 |
| ATP-dependent RNA helicase DDX25 | Q9QY15 |
| AT-rich interactive domain-containing protein 4B | A2CG63 |
| Baculoviral IAP repeat-containing protein 2 | Q62210 |
| B-cell CLL/lymphoma 9-like protein | Q67FY2 |
| Beclin-1 | O88597 |
| Beta-1,3-glucosyltransferase | Q8BHT6 |
| Beta-2-syntrophin | Q61235 |
| Branched-chain-amino-acid aminotransferase, mi | O35855 |
| Bromodomain and WD repeat-containing protein : | A2AHJ4 |
| BTB/POZ domain-containing protein KCTD3 | Q8BFX3 |
| BTB/POZ domain-containing protein KCTD4 | Q9D7X1 |
| Cadherin EGF LAG seven-pass G-type receptor 2 | Q9R0M0 |
| Cadherin-6 | P97326 |
| Calcitonin gene-related peptide type 1 receptor | Q9R1W5 |
| Calmodulin-like protein 3 | Q9D6P8 |
| cAMP and cAMP-inhibited cGMP 3',5'-cyclic phos | Q8CA95 |
| Carboxypeptidase M | Q80V42 |
| Carcinoembryonic antigen-related cell adhesion m | Q3UKK2 |
| Caseinolytic peptidase B protein homolog | Q60649 |
| Catechol O-methyltransferase | O88587 |
| Cathepsin F | Q9R013 |
| CCR4-NOT transcription complex subunit 7 | Q60809 |
| CD109 antigen | Q8R422 |
| CD2 antigen cytoplasmic tail-binding protein 2 | Q9CWK3 |
| CDKN2AIP N-terminal-like protein | Q9D211 |
| Cell differentiation protein RCD1 homolog | Q9JKY0 |
| Centrosomal protein of 95 kDa | Q8BVV7 |
| Ceruloplasmin | Q61147 |

| | |
|---|---|
| AP2-associated protein kinase 1 | Q3UHJ0 |
| AP-3 complex subunit beta-1 | Q9Z1T1 |
| AP-3 complex subunit beta-2 | Q9JME5 |
| AP-3 complex subunit delta-1 | O54774 |
| AP-3 complex subunit sigma-1 | Q9DCR2 |
| Apolipoprotein A-I | Q00623 |
| Apolipoprotein E | P08226 |
| Apolipoprotein O | Q9DCZ4 |
| Apoptosis inhibitor 5 | O35841 |
| Apoptosis-inducing factor 1, mitochondrial | Q9Z0X1 |
| Apoptosis-stimulating of p53 protein 1 | Q62415 |
| ARF GTPase-activating protein GIT1 | Q68FF6 |
| Arfaptin-2 | Q8K221 |
| Arf-GAP domain and FG repeat-containing protei | Q8K2K6 |
| Arf-GAP with coiled-coil, ANK repeat and PH don | Q6ZQK5 |
| Arf-GAP with GTPase, ANK repeat and PH doma | Q3UHD9 |
| Arf-GAP with GTPase, ANK repeat and PH doma | Q8VHH5 |
| Arf-GAP with SH3 domain, ANK repeat and PH d | Q9QWY8 |
| Arf-GAP with SH3 domain, ANK repeat and PH d | Q7SIG6 |
| Arginine--tRNA ligase, cytoplasmic | Q9D0I9 |
| Argininosuccinate lyase | Q91YI0 |
| Argininosuccinate synthase | P16460 |
| Arginyl-tRNA--protein transferase 1 | Q9Z2A5 |
| Armadillo repeat-containing protein 6 | Q8BNU0 |
| Arylsulfatase B | P50429 |
| Asparagine synthetase [glutamine-hydrolyzing] | Q61024 |
| Asparagine--tRNA ligase, cytoplasmic | Q8BP47 |
| Aspartate aminotransferase, cytoplasmic | P05201 |
| Aspartate aminotransferase, mitochondrial | P05202 |
| Aspartate--tRNA ligase, cytoplasmic | Q922B2 |
| Aspartyl aminopeptidase | Q9Z2W0 |
| Astrocytic phosphoprotein PEA-15 | Q62048 |
| Astrotactin-1 | Q61137 |
| Ataxin-10 | P28658 |
| Ataxin-2 | O70305 |
| Ataxin-2-like protein | Q7TQH0 |
| Atlastin-1 | Q8BH66 |
| Atlastin-2 | Q6PA06 |
| ATP synthase subunit alpha, mitochondrial | Q03265 |
| ATP synthase subunit b, mitochondrial | Q9CQQ7 |
| ATP synthase subunit beta, mitochondrial | P56480 |
| ATP synthase subunit d, mitochondrial | Q9DCX2 |
| ATP synthase subunit delta, mitochondrial | Q9D3D9 |
| ATP synthase subunit e, mitochondrial | Q06185 |
| ATP synthase subunit epsilon, mitochondrial | P56382 |
| ATP synthase subunit g, mitochondrial | Q9CPQ8 |
| ATP synthase subunit gamma, mitochondrial | Q91VR2 |
| ATP synthase subunit O, mitochondrial | Q9DB20 |
| ATP synthase-coupling factor 6, mitochondrial | P97450 |
| ATPase Asna1 | O54984 |
| ATPase family AAA domain-containing protein 1 | Q9D5T0 |
| ATPase family AAA domain-containing protein 3 | Q925I1 |
| ATPase inhibitor, mitochondrial | O35143 |
| ATP-binding cassette sub-family D member 3 | P55096 |
| ATP-binding cassette sub-family E member 1 | P61222 |
| ATP-binding cassette sub-family F member 1 | Q6P542 |
| ATP-binding cassette sub-family F member 2 | Q99LE6 |
| ATP-citrate synthase | Q91V92 |
| ATP-dependent (S)-NAD(P)H-hydrate dehydratas | Q9CZ42 |
| ATP-dependent RNA helicase DDX1 | Q91VR5 |
| ATP-dependent RNA helicase DDX19A | Q61655 |
| ATP-dependent RNA helicase DDX3X | Q62167 |
| ATP-dependent RNA helicase Dhx29 | Q6PGC1 |
| Autophagy-related protein 16-1 | Q8C0J2 |
| Autophagy-related protein 2 homolog B | Q80XK6 |
| Baculoviral IAP repeat-containing protein 6 | O88738 |
| BAG family molecular chaperone regulator 2 | Q91YN9 |
| BAG family molecular chaperone regulator 4 | Q8CI61 |
| BAG family molecular chaperone regulator 5 | Q8CI32 |
| Band 4.1-like protein 1 | Q9Z2H5 |
| Band 4.1-like protein 2 | O70318 |
| Band 4.1-like protein 3 | Q9WV92 |
| Basic leucine zipper and W2 domain-containing p | Q9CQC6 |
| Basigin | P18572 |
| B-cell receptor-associated protein 31 | Q61335 |
| Bcl-2-like protein 13 | P59017 |
| BDNF/NT-3 growth factors receptor | P15209 |
| Beta-adducin | Q9QYB8 |
| Beta-adrenergic receptor kinase 1 | Q99MK8 |
| Beta-arrestin-1 | Q8BWG8 |
| Beta-centractin | Q8R5C5 |
| Beta-galactosidase | P23780 |
| Beta-glucuronidase | P12265 |
| Beta-hexosaminidase subunit beta | P20060 |
| Beta-soluble NSF attachment protein | P28663 |
| Beta-synuclein | Q91ZZ3 |
| Bifunctional 3'-phosphoadenosine 5'-phosphosulf | Q60967 |
| Bifunctional ATP-dependent dihydroxyacetone ki | Q8VC30 |
| Bifunctional glutamate/proline--tRNA ligase | Q8CGC7 |
| Bifunctional protein NCOAT | Q9EQQ9 |

| | |
|---|---|
| RalBP1-associated Eps domain-containing protei | Q80XA6 |
| Ran-binding protein 6 | Q8BIV3 |
| Ras-related protein Rab-33B | O35963 |
| Receptor-type tyrosine-protein phosphatase gam | Q05909 |
| Ribose-phosphate pyrophosphokinase 1 | Q9D7G0 |
| Ribosyldihydronicotinamide dehydrogenase [quin | Q9JI75 |
| RUN and FYVE domain-containing protein 1 | Q8BIJ7 |
| Secernin-3 | Q3TMH2 |
| Serine/arginine-rich splicing factor 5 | O35326 |
| Serine/threonine-protein kinase 38-like | Q7TSE6 |
| Serine/threonine-protein kinase LMTK2 | Q3TYD6 |
| Serine/threonine-protein phosphatase 6 regulator | Q922D4 |
| Shootin-1 | Q8K2Q9 |
| Short/branched chain specific acyl-CoA dehydrog | Q9DBL1 |
| Signal transducer and activator of transcription 5E | P42232 |
| SNF-related serine/threonine-protein kinase | Q8VDU5 |
| Solute carrier family 12 member 6 | Q924N4 |
| Spermatogenesis-defective protein 39 homolog | Q8BGQ1 |
| SRA stem-loop-interacting RNA-binding protein, r | Q9D8T7 |
| Syntaxin-binding protein 3 | Q60770 |
| Target of rapamycin complex subunit LST8 | Q9DCJ1 |
| TBC1 domain family member 7 | Q9D0K0 |
| Tensin-3 | Q5SSZ5 |
| Tetratricopeptide repeat protein 9C | Q810A3 |
| Thioredoxin-related transmembrane protein 4 | Q8C0L0 |
| Thymocyte nuclear protein 1 | Q91YJ3 |
| Transgelin | P37804 |
| Trypsin - Sus scrofa (Pig). | P00761 |
| Tubulin beta-6 chain | Q922F4 |
| UPF0552 protein C15orf38 homolog | Q9D0A3 |
| UPF0670 protein C8orf55 homolog | Q80ZW2 |
| Vacuolar protein sorting-associated protein 37C | Q8R105 |
| Vacuolar protein sorting-associated protein 4B | P46467 |
| VPS10 domain-containing receptor SorCS3 | Q8VI51 |
| YTH domain family protein 3 | Q8BYK6 |
| **Detected in experiments 2&3 - 680 proteins** | |
| [Pyruvate dehydrogenase [acetyl-transferring]]-ph | Q3UV70 |
| [Pyruvate dehydrogenase [lipoamide]] kinase isoz | Q922H2 |
| 1-phosphatidylinositol 4,5-bisphosphate phospho | P51432 |
| 2,4-dienoyl-CoA reductase, mitochondrial | Q9CQ62 |
| 28S ribosomal protein S16, mitochondrial | Q9CPX7 |
| 28S ribosomal protein S2, mitochondrial | Q924T2 |
| 28S ribosomal protein S22, mitochondrial | Q9CXW2 |
| 28S ribosomal protein S23, mitochondrial | Q8VE22 |
| 28S ribosomal protein S29, mitochondrial | Q9ER88 |
| 28S ribosomal protein S30, mitochondrial | Q9D0G0 |
| 2-C-methyl-D-erythritol 4-phosphate cytidylyltrans | Q5RJG7 |
| 2-hydroxyacylsphingosine 1-beta-galactosyltransf | Q64676 |
| 2-methoxy-6-polyprenyl-1,4-benzoquinol methylas | Q9CXI0 |
| 39S ribosomal protein L11, mitochondrial | Q9CQF0 |
| 39S ribosomal protein L15, mitochondrial | Q9CPR5 |
| 39S ribosomal protein L19, mitochondrial | Q9D338 |
| 39S ribosomal protein L2, mitochondrial | Q9D773 |
| 39S ribosomal protein L22, mitochondrial | Q8BU88 |
| 39S ribosomal protein L38, mitochondrial | Q8K2M0 |
| 39S ribosomal protein L40, mitochondrial | Q9Z2Q5 |
| 39S ribosomal protein L41, mitochondrial | Q9CQN7 |
| 39S ribosomal protein L49, mitochondrial | Q9CQ40 |
| 3-hydroxyacyl-CoA dehydratase 3 | Q8K2C9 |
| 40S ribosomal protein S29 | P62274 |
| 5'(3')-deoxyribonucleotidase, mitochondrial | Q8VCE6 |
| 60S ribosomal protein L22-like 1 | Q9D7S7 |
| 60S ribosomal protein L24 | Q8BP67 |
| 60S ribosomal protein L32 | P62911 |
| 60S ribosomal protein L36 | P47964 |
| 6-phosphofructo-2-kinase/fructose-2,6-bisphosph | P70265 |
| Abhydrolase domain-containing protein 14B | Q8VCR7 |
| Abhydrolase domain-containing protein 4 | Q8VD66 |
| Acid ceramidase | Q9WV54 |
| Actin-related protein 2/3 complex subunit 1B | Q9WV32 |
| Actin-related protein 2/3 complex subunit 5 | Q9CPW4 |
| Acyl-CoA synthetase family member 3, mitochonc | Q3URE1 |
| Adenine phosphoribosyltransferase | P08030 |
| Adenosine deaminase | P03958 |
| Adenylyltransferase and sulfurtransferase MOCS | A2BDX3 |
| ADP-ribosylation factor GTPase-activating protei | Q99K28 |
| Aggrecan core protein | Q61282 |
| Agrin | A2ASQ1 |
| A-kinase anchor protein 10, mitochondrial | O88845 |
| Alkyldihydroxyacetonephosphate synthase, perox | Q8C0I1 |
| AMSH-like protease | Q76N33 |
| Amyloid beta A4 precursor protein-binding family | P98084 |
| Amyloid-like protein 2 | Q06335 |
| AN1-type zinc finger protein 2B | Q91X58 |
| Anaphase-promoting complex subunit 4 | Q91W96 |
| Ankyrin repeat domain-containing protein 13D | Q6PD24 |
| AP-3 complex subunit mu-1 | Q9JKC8 |
| AP-3 complex subunit mu-2 | Q8R2R9 |
| AP-3 complex subunit sigma-2 | Q8BSZ2 |
| APC membrane recruitment protein 2 | Q8CCJ4 |

| | |
|---|---|
| cGMP-dependent protein kinase 1 | P0C605 |
| Chaperone activity of bc1 complex-like, mitochonc | Q60936 |
| Charged multivesicular body protein 1b-2 | Q9CQD4 |
| Chromobox protein homolog 1 | P83917 |
| Chromosome-associated kinesin KIF4 | P33174 |
| Cleft lip and palate transmembrane protein 1 hom | Q8VBZ3 |
| Coiled-coil domain-containing protein 39 | Q9D5Y1 |
| Coiled-coil domain-containing protein 75 | Q3UFS4 |
| Coiled-coil domain-containing protein 85A | Q5SP85 |
| Cold-inducible RNA-binding protein | P60824 |
| Complement C3 | P01027 |
| Copine-3 | Q8BT60 |
| Copper chaperone for superoxide dismutase | Q9WU84 |
| COX assembly mitochondrial protein homolog | Q9CPZ8 |
| C-type lectin domain family 2 member L | P0C7M9 |
| CUE domain-containing protein 1 | Q8R3V6 |
| Cyclin-dependent kinase inhibitor 1B | P46414 |
| Cytochrome b-c1 complex subunit 8 | Q9CQ69 |
| Cytochrome c-type heme lyase | P53702 |
| Cytochrome P450 2S1 | Q9DBX6 |
| Cytohesin-2 | P63034 |
| Cytosolic carboxypeptidase 4 | Q09M05 |
| Cytospin-A | Q2KN98 |
| DAZ-associated protein 1 | Q9JII5 |
| DDB1- and CUL4-associated factor 6 | Q9DC22 |
| Dedicator of cytokinesis protein 11 | A2AF47 |
| Dihydroorotate dehydrogenase (quinone), mitocho | O35435 |
| DNA ligase 3 | P97386 |
| DNA replication licensing factor MCM8 | Q9CWV1 |
| DNA topoisomerase 3-beta-1 | Q9Z321 |
| DNA-directed RNA polymerase II subunit RPB9 | P60898 |
| DNA-directed RNA polymerases I and III subunit F | P52432 |
| DnaJ homolog subfamily C member 2 | P54103 |
| Double-stranded RNA-binding protein Staufen ho | Q9Z108 |
| Dr1-associated corepressor | Q9D6N5 |
| Dynactin subunit 6 | Q9WUB4 |
| Dynein heavy chain 8, axonemal | Q91XQ0 |
| Dynein light chain 1, axonemal | Q05A62 |
| E3 ubiquitin-protein ligase BRE1B | Q3U319 |
| E3 ubiquitin-protein ligase MGRN1 | Q9D074 |
| E3 ubiquitin-protein ligase MIB1 | Q80SY4 |
| E3 ubiquitin-protein ligase Praja-1 | O55176 |
| E3 ubiquitin-protein ligase RBX1 | P62878 |
| E3 ubiquitin-protein ligase RNF187 | Q8BFX1 |
| E3 ubiquitin-protein ligase SH3RF1 | Q69ZI1 |
| Ectopic P granules protein 5 homolog | Q80TA9 |
| EGF-like repeat and discoidin I-like domain-contai | O35474 |
| ELMO domain-containing protein 1 | Q3V1U8 |
| Emerin | O08579 |
| EMI domain-containing protein 1 | Q91VF5 |
| Endoplasmic reticulum lectin 1 | Q8VEH8 |
| Endoplasmic reticulum-Golgi intermediate compar | Q9CQE7 |
| Enhancer of filamentation 1 | O35177 |
| Enhancer of mRNA-decapping protein 3 | Q8K2D3 |
| Ephrin type-A receptor 7 | Q61772 |
| Eukaryotic translation initiation factor 1A | Q60872 |
| Eukaryotic translation initiation factor 2 subunit 3, | Q9Z0N2 |
| Exonuclease 3'-5' domain-containing protein 2 | Q8VEG4 |
| Exosome complex component MTR3 | Q8BTW3 |
| Exosome complex exonuclease RRP44 | Q9CSH3 |
| Extended synaptotagmin-1 | Q3U7R1 |
| FACT complex subunit SSRP1 | Q08943 |
| FAD-linked sulfhydryl oxidase ALR | P56213 |
| Fatty acid desaturase 1 | Q920L1 |
| Fatty acid desaturase 2 | Q9Z0R9 |
| F-box only protein 42 | Q6PDJ6 |
| F-box only protein 7 | Q3U7U3 |
| F-box/LRR-repeat protein 15 | Q91W61 |
| F-box-like/WD repeat-containing protein TBL1XR: | Q8BHJ5 |
| FERM domain-containing protein 8 | Q3UFK8 |
| Fibrocystin-L | Q80ZA4 |
| Flap endonuclease 1 | P39749 |
| Forkhead-associated domain-containing protein 1 | A6PWD2 |
| FXYD domain-containing ion transport regulator 7 | P59648 |
| GAS2-like protein 3 | Q3UWW6 |
| GDNF family receptor alpha-2 | O08842 |
| Gem-associated protein 5 | Q8BX17 |
| Glutamyl-tRNA(Gln) amidotransferase subunit C, | Q8CBY0 |
| Glutathione peroxidase 1 | P11352 |
| Glycogen [starch] synthase, muscle | Q9Z1E4 |
| Glypican-6 | Q9R087 |
| Golgin subfamily A member 5 | Q9QYE6 |
| Golgin subfamily A member 7 | Q91W53 |
| GPI transamidase component PIG-T | Q8BXQ2 |
| Guanine nucleotide exchange factor MSS4 | Q91X96 |
| Guanine nucleotide-binding protein G(I)/G(S)/G(O | P50153 |
| Guanine nucleotide-binding protein G(I)/G(S)/G(O | Q80SZ7 |
| Guanine nucleotide-binding protein subunit alpha- | P27600 |
| H(+)/Cl(-) exchange transporter 3 | P51791 |
| Heat shock 70 kDa protein 1-like | P16627 |

| | |
|---|---|
| Bifunctional purine biosynthesis protein PURH | Q9CWJ9 |
| Biliverdin reductase A | Q9CY64 |
| Bleomycin hydrolase | Q8R016 |
| BolA-like protein 2 | Q8BGS2 |
| Brain acid soluble protein 1 | Q91XV3 |
| Brain-specific angiogenesis inhibitor 1 | Q3UHD1 |
| Brain-specific angiogenesis inhibitor 1-associatec | Q8BKX1 |
| Brain-specific angiogenesis inhibitor 2 | Q8CGM1 |
| Branched-chain-amino-acid aminotransferase, cy | P24288 |
| Breast carcinoma-amplified sequence 1 homolog | Q80YN3 |
| Brefeldin A-inhibited guanine nucleotide-exchang | Q3UGY8 |
| Brevican core protein | Q61361 |
| BRO1 domain-containing protein BROX | Q8K2Q7 |
| BTB/POZ domain-containing protein 17 | Q9DB72 |
| BTB/POZ domain-containing protein KCTD12 | Q6WVG3 |
| C-1-tetrahydrofolate synthase, cytoplasmic | Q922D8 |
| C2 domain-containing protein 2-like | Q80X80 |
| Cadherin-13 | Q9WTR5 |
| Cadherin-2 | P15116 |
| Calbindin | P12658 |
| Calcineurin subunit B type 1 | Q63810 |
| Calcineurin-like phosphoesterase domain-contair | Q8BFS6 |
| Calcipressin-1 | Q9JHG6 |
| Calcium uniporter protein, mitochondrial | Q3UMR5 |
| Calcium/calmodulin-dependent 3',5'-cyclic nucleo | Q61481 |
| Calcium/calmodulin-dependent 3',5'-cyclic nucleo | Q01065 |
| Calcium/calmodulin-dependent protein kinase kin | Q8VBY2 |
| Calcium/calmodulin-dependent protein kinase kin | Q8C078 |
| Calcium/calmodulin-dependent protein kinase typ | Q8BW96 |
| Calcium/calmodulin-dependent protein kinase typ | P11798 |
| Calcium/calmodulin-dependent protein kinase typ | P28652 |
| Calcium/calmodulin-dependent protein kinase typ | Q6PHZ2 |
| Calcium/calmodulin-dependent protein kinase typ | Q923T9 |
| Calcium/calmodulin-dependent protein kinase typ | P08414 |
| Calcium-binding and coiled-coil domain-containin | Q8CGU1 |
| Calcium-binding mitochondrial carrier protein Aral | Q8BH59 |
| Calcium-binding mitochondrial carrier protein SCa | Q6GQS1 |
| Calcium-binding protein 39 | Q06138 |
| Calcium-binding protein p22 | P61022 |
| Calcium-dependent secretion activator 1 | Q80TJ1 |
| Calcium-dependent secretion activator 2 | Q8BYR5 |
| Calcium-transporting ATPase type 2C member 1 | Q80XR2 |
| Calcyclin-binding protein | Q9CXW3 |
| Calmodulin | P62204 |
| Calmodulin-regulated spectrin-associated protein | Q8C1B1 |
| Calmodulin-regulated spectrin-associated protein | Q80VC9 |
| Calnexin | P35564 |
| Calpain small subunit 1 | O88456 |
| Calpain-2 catalytic subunit | O08529 |
| Calpain-5 | O08688 |
| Calreticulin | P14211 |
| Calretinin | Q08331 |
| Calsyntenin-1 | Q9EPL2 |
| Calumenin | O35887 |
| CaM kinase-like vesicle-associated protein | Q3UHL1 |
| cAMP-dependent protein kinase catalytic subunit | P68181 |
| cAMP-dependent protein kinase type I-alpha regu | Q9DBC7 |
| cAMP-dependent protein kinase type II-alpha reg | P12367 |
| cAMP-dependent protein kinase type II-beta regu | P31324 |
| CAP-Gly domain-containing linker protein 1 | Q922J3 |
| CAP-Gly domain-containing linker protein 2 | Q9Z0H8 |
| Caprin-1 | Q60865 |
| Carbonic anhydrase 2 | P00920 |
| Carbonyl reductase [NADPH] 1 | P48758 |
| Carbonyl reductase [NADPH] 3 | Q8K354 |
| Carboxypeptidase E | Q00493 |
| Carnitine O-palmitoyltransferase 1, brain isoform | Q8BGD5 |
| Cartilage acidic protein 1 | Q8R555 |
| Casein kinase I isoform alpha | Q8BK63 |
| Casein kinase I isoform epsilon | Q9JMK2 |
| Casein kinase I isoform gamma-1 | Q8BTH8 |
| Casein kinase II subunit alpha | Q60737 |
| Casein kinase II subunit alpha' | O54833 |
| Casein kinase II subunit beta | P67871 |
| Caskin-1 | Q6P9K8 |
| Caspase-3 | P70677 |
| Catalase | P24270 |
| Catenin alpha-2 | Q61301 |
| Catenin beta-1 | Q02248 |
| Catenin delta-1 | P30999 |
| Catenin delta-2 | O35927 |
| Cathepsin D | P18242 |
| Cation-dependent mannose-6-phosphate recepto | P24668 |
| Cation-independent mannose-6-phosphate recep | Q07113 |
| CB1 cannabinoid receptor-interacting protein 1 | Q5M8N0 |
| CCA tRNA nucleotidyltransferase 1, mitochondria | Q8K1J6 |
| CCR4-NOT transcription complex subunit 1 | Q6ZQ08 |
| CD166 antigen | Q61490 |
| CDP-diacylglycerol--glycerol-3-phosphate 3-phos | Q8BHF7 |
| Cell adhesion molecule 1 | Q8R5M8 |
| Arginine and glutamate-rich protein 1 | Q3UL36 |
| Armadillo repeat protein deleted in velo-cardio-fac | P98203 |
| Armadillo repeat-containing protein 1 | Q9D7A8 |
| Armadillo repeat-containing X-linked protein 3 | Q8BHS6 |
| ATPase WRNIP1 | Q91XU0 |
| ATP-binding cassette sub-family A member 2 | P41234 |
| ATP-binding cassette sub-family A member 7 | Q91V24 |
| ATP-binding cassette sub-family D member 2 | Q61285 |
| ATP-dependent Clp protease ATP-binding subun | Q9JHS4 |
| Autophagy protein 5 | Q99J83 |
| BAG family molecular chaperone regulator 1 | Q60739 |
| Barrier-to-autointegration factor | O54962 |
| B-cell receptor-associated protein 29 | Q61334 |
| Bcl-2-associated transcription factor 1 | Q8K019 |
| Beta-adrenergic receptor kinase 2 | Q3UYH7 |
| Beta-hexosaminidase subunit alpha | P29416 |
| Beta-lactamase-like protein 2 | Q99KR3 |
| Bifunctional arginine demethylase and lysyl-hydrc | Q9ERI5 |
| Bifunctional coenzyme A synthase | Q9DBL7 |
| Bis(5'-nucleosyl)-tetraphosphatase [asymmetrical | P56380 |
| BolA-like protein 1 | Q9D8S9 |
| Brain protein 44-like protein | P63030 |
| Brain-enriched guanylate kinase-associated prote | Q68EF6 |
| Breakpoint cluster region protein | Q6PAJ1 |
| Breast cancer anti-estrogen resistance protein 1 | Q61140 |
| Breast carcinoma-amplified sequence 3 homolog | Q8CCN5 |
| BTB/POZ domain-containing protein KCTD16 | Q5DTY9 |
| C2 domain-containing protein 3 | Q52KB6 |
| CAAX prenyl protease 1 homolog | Q80W54 |
| Cadherin-10 | P70408 |
| Cadherin-11 | P55288 |
| Cadherin-8 | P97291 |
| Cadherin-9 | P70407 |
| Calcipressin-2 | Q9JHG2 |
| Calcium uptake protein 1, mitochondrial | Q8VCX5 |
| Calcium/calmodulin-dependent protein kinase II ir | Q6QWF9 |
| Calcium/calmodulin-dependent protein kinase typ | Q9QYK9 |
| Calcium-activated potassium channel subunit alpl | Q08460 |
| Calcium-independent phospholipase A2-gamma | Q8K1N1 |
| Calsyntenin-3 | Q99JH7 |
| CAP-Gly domain-containing linker protein 3 | B9EHT4 |
| Cap-specific mRNA (nucleoside-2'-O-)-methyltran | Q9DBC3 |
| Carbonic anhydrase 4 | Q64444 |
| Carboxymethylenebutenolidase homolog | Q8R1G2 |
| Carboxypeptidase Q | Q9WVJ3 |
| Carnitine O-acetyltransferase | P47934 |
| Carnitine O-palmitoyltransferase 2, mitochondrial | P52825 |
| Casein kinase I isoform delta | Q9DC28 |
| Catechol O-methyltransferase domain-containing | Q8BIG7 |
| Catenin alpha-1 | P26231 |
| Cathepsin B | P10605 |
| Cathepsin L1 | P06797 |
| CCR4-NOT transcription complex subunit 3 | Q8K0V4 |
| Cdc42 effector protein 4 | Q9JM96 |
| CDGSH iron-sulfur domain-containing protein 1 | Q91WS0 |
| CDGSH iron-sulfur domain-containing protein 2 | Q9CQB5 |
| Cell division cycle protein 23 homolog | Q8BGZ4 |
| Cell division cycle protein 27 homolog | A2A6Q5 |
| Cellular nucleic acid-binding protein | P53996 |
| Centromere protein V | Q9CXS4 |
| Centromere-associated protein E | Q6RT24 |
| Cerebellin-1 | Q9R171 |
| Charged multivesicular body protein 6 | P0C0A3 |
| Chloride intracellular channel protein 1 | Q9Z1Q5 |
| Chloride transport protein 6 | O35454 |
| Cholecystokinin | P09240 |
| Choline O-acetyltransferase | Q03059 |
| Choline transporter-like protein 2 | Q8BY89 |
| Choline/ethanolamine kinase | O55229 |
| Choline-phosphate cytidylyltransferase B | Q811Q9 |
| Chromobox protein homolog 3 | P23198 |
| Chromobox protein homolog 5 | Q61686 |
| Chromogranin-A | P26339 |
| Citron Rho-interacting kinase | P49025 |
| Clusterin-associated protein 1 | Q8R3P7 |
| Coatomer subunit zeta-1 | P61924 |
| Cob(I)yrinic acid a,c-diamide adenosyltransferase | Q9D273 |
| Coiled-coil domain-containing protein 127 | Q3TC33 |
| Coiled-coil domain-containing protein 136 | Q3TVA9 |
| Coiled-coil domain-containing protein 165 | Q3UHU5 |
| Coiled-coil domain-containing protein 47 | Q9D024 |
| Coiled-coil domain-containing protein 50 | Q810U5 |
| Coiled-coil domain-containing protein 58 | Q8R3Q6 |
| Coiled-coil domain-containing protein 90B, mitoch | Q8C3X2 |
| Coiled-coil domain-containing protein 91 | Q9D8L5 |
| Coiled-coil-helix-coiled-coil-helix domain-containir | Q9D1L0 |
| COMM domain-containing protein 2 | Q8BXC6 |
| Complement component 1 Q subcomponent-bind | O35658 |
| Contactin-6 | Q9JMB8 |
| Copper transport protein ATOX1 | O08997 |
| Hematological and neurological expressed 1 prote | P97825 |
| Heparan-sulfate 6-O-sulfotransferase 3 | Q9QYK4 |
| Hepatocyte growth factor receptor | P16056 |
| Highly divergent homeobox | Q14B70 |
| Histone deacetylase 2 | P70288 |
| Histone H1.1 | P43275 |
| Histone H1.3 | P43277 |
| Histone H2B type 1-M | P10854 |
| Histone-lysine N-methyltransferase EHMT2 | Q9Z148 |
| Hsp90 co-chaperone Cdc37-like 1 | Q9CZP7 |
| IgE-binding protein | P03975 |
| Importin subunit alpha-4 | O35343 |
| Integral membrane protein 2B | O89051 |
| Integrin-linked protein kinase | O55222 |
| Intraflagellar transport protein 122 homolog | Q6NWV3 |
| Iron-sulfur cluster co-chaperone protein HscB, mit | Q8K3A0 |
| Junctional adhesion molecule B | Q9JI59 |
| Kelch-like ECH-associated protein 1 | Q9Z2X8 |
| Kelch-like protein 3 | E0CZ16 |
| Keratin, type I cytoskeletal 18 | P05784 |
| KH domain-containing, RNA-binding, signal transo | Q9WU01 |
| Kinesin light chain 4 | Q9DBS5 |
| Kinesin-like protein KIF27 | Q7M6Z4 |
| Kininogen-1 | O08677 |
| Lamin-B2 | P21619 |
| Lanosterol 14-alpha demethylase | Q8K0C4 |
| La-related protein 4 | Q8BWW4 |
| Latexin | P70202 |
| Leptin receptor | P48356 |
| Leucine zipper putative tumor suppressor 1 | P60853 |
| Leucine-rich repeat and fibronectin type-III domai | Q80XU8 |
| Leucine-rich repeat neuronal protein 1 | Q61809 |
| Leucine-rich repeat-containing protein 8B | Q5DU41 |
| Leucine-rich repeat-containing protein 8C | Q8R502 |
| Lipid phosphate phosphatase-related protein type | Q7TPB0 |
| LMBR1 domain-containing protein 2 | Q8C561 |
| Low-density lipoprotein receptor-related protein 4 | Q8VI56 |
| Lymphotoxin-alpha | P09225 |
| Lysosomal protective protein | P16675 |
| Macrophage colony-stimulating factor 1 receptor | P09581 |
| Magnesium transporter protein 1 | Q9CQY5 |
| Metastasis-associated protein MTA2 | Q9R190 |
| Methyl-CpG-binding protein 2 | Q9Z2D6 |
| Methylmalonyl-CoA epimerase, mitochondrial | Q9D1I5 |
| MHC class II transactivator | P79621 |
| Microtubule-associated serine/threonine-protein k | Q811L6 |
| Mitochondrial import receptor subunit TOM6 homo | Q9CQN3 |
| Mitochondrial inner membrane protein OXA1L | Q8BGA9 |
| Mitogen-activated protein kinase kinase kinase 5 | O35099 |
| Mitogen-activated protein kinase kinase kinase 7 | Q62073 |
| MOB kinase activator 2 | Q8VI63 |
| Molybdopterin synthase catalytic subunit | Q9Z223 |
| Multiple epidermal growth factor-like domains prot | P60882 |
| Muscarinic acetylcholine receptor M1 | P12657 |
| Myb/SANT-like DNA-binding domain-containing pr | Q91YU3 |
| Myosin light chain 6B | Q8CI43 |
| NACHT, LRR and PYD domains-containing protei | Q91WS2 |
| NEDD4-like E3 ubiquitin-protein ligase WWP2 | Q9DBH0 |
| Netrin-G1 | Q8R4G0 |
| Netrin-G2 | Q8R4F1 |
| Neutrophil cytosol factor 2 | O70145 |
| NHP2-like protein 1 | Q9D0T1 |
| Niban-like protein 1 | Q8R1F1 |
| NIPA-like protein 3 | Q8BGN5 |
| Non-specific lipid-transfer protein | P32020 |
| Non-syndromic hearing impairment protein 5 hom | Q9Z2D3 |
| Nuclear receptor-interacting protein 3 | Q9JJR9 |
| Nuclease-sensitive element-binding protein 1 | P62960 |
| Nucleolar transcription factor 1 | P25976 |
| O-acetyl-ADP-ribose deacetylase MACROD2 | Q3UYG8 |
| Oligophrenin-1 | Q99J31 |
| Opioid growth factor receptor-like protein 1 | Q8VE52 |
| Optic atrophy 3 protein homolog | Q505D7 |
| PAB-dependent poly(A)-specific ribonuclease sub | Q640Q5 |
| Paralemmin-3 | A2TJV2 |
| PDZ and LIM domain protein 4 | P70271 |
| Peptidyl-prolyl cis-trans isomerase-like 3 | Q9D6L8 |
| Peroxisome assembly factor 2 | Q99LC9 |
| Peroxisome biogenesis factor 1 | Q5BL07 |
| PERQ amino acid-rich with GYF domain-containir | Q6Y7W8 |
| PH and SEC7 domain-containing protein 2 | Q6P1I6 |
| Phenylalanine--tRNA ligase, mitochondrial | Q99M01 |
| Phosphoethanolamine/phosphocholine phosphata | Q8R2H9 |
| Phospholipase D1 | Q9Z280 |
| Phospholipase DDHD2 | Q80Y98 |
| Phosphorylase b kinase gamma catalytic chain, sl | P07934 |
| Phosphoserine phosphatase | Q99LS3 |
| Pleckstrin homology domain-containing family A n | Q3UIL6 |
| Pleckstrin homology domain-containing family G r | Q66T02 |
| Polypyrimidine tract-binding protein 1 | P17225 |

| | |
|---|---|
| Cell adhesion molecule 2 | Q8BLQ9 |
| Cell adhesion molecule 3 | Q99N28 |
| Cell adhesion molecule 4 | Q8R464 |
| Cell cycle exit and neuronal differentiation protein | Q9JKC6 |
| Cell division control protein 42 homolog | P60766 |
| Cell surface glycoprotein MUC18 | Q8R2Y2 |
| Centrin-2 | Q9R1K9 |
| Centrosomal protein of 170 kDa | Q6A065 |
| cGMP-dependent 3',5'-cyclic phosphodiesterase | Q922S4 |
| Charged multivesicular body protein 3 | Q9CQ10 |
| Charged multivesicular body protein 4b | Q9D8B3 |
| Charged multivesicular body protein 5 | Q9D7S9 |
| Charged multivesicular body protein 7 | Q8R1T1 |
| Chitinase domain-containing protein 1 | Q922Q9 |
| Chloride intracellular channel protein 4 | Q9QYB1 |
| Cholesterol 24-hydroxylase | Q9WVK8 |
| Choline-phosphate cytidylyltransferase A | P49586 |
| Chondroitin sulfate proteoglycan 4 | Q8VHY0 |
| Chondroitin sulfate proteoglycan 5 | Q71M36 |
| Citrate lyase subunit beta-like protein, mitochondi | Q8R4N0 |
| Citrate synthase, mitochondrial | Q9CZU6 |
| C-Jun-amino-terminal kinase-interacting protein 3 | Q9ESN9 |
| C-Jun-amino-terminal kinase-interacting protein 4 | Q58A65 |
| Clathrin coat assembly protein AP180 | Q61548 |
| Clathrin heavy chain 1 | Q68FD5 |
| Clathrin interactor 1 | Q99KN9 |
| Clathrin light chain A | O08585 |
| Clathrin light chain B | Q6IRU5 |
| Clavesin-1 | Q9D4C9 |
| Clavesin-2 | Q8BG92 |
| CLIP-associating protein 1 | Q80TV8 |
| CLIP-associating protein 2 | Q8BRT1 |
| Clusterin | Q06890 |
| Coactosin-like protein | Q9CQI6 |
| Coatomer subunit alpha | Q8CIE6 |
| Coatomer subunit beta | Q9JIF7 |
| Coatomer subunit beta' | O55029 |
| Coatomer subunit delta | Q5XJY5 |
| Coatomer subunit epsilon | O89079 |
| Coatomer subunit gamma-1 | Q9QZE5 |
| Coatomer subunit gamma-2 | Q9QXK3 |
| Cofilin-1 | P18760 |
| Cofilin-2 | P45591 |
| Coiled-coil and C2 domain-containing protein 1A | Q8K1A6 |
| Coiled-coil domain-containing protein 104 | Q8C6E0 |
| Coiled-coil domain-containing protein 132 | Q8CI71 |
| Coiled-coil domain-containing protein 22 | Q9JIG7 |
| Coiled-coil domain-containing protein 43 | Q9CR29 |
| Coiled-coil domain-containing protein 6 | D3YZP9 |
| Coiled-coil domain-containing protein 73 | Q8CDM4 |
| Coiled-coil domain-containing protein 92 | Q8VDN4 |
| Coiled-coil domain-containing protein 93 | Q7TQK5 |
| Coiled-coil-helix-coiled-coil-helix domain-containir | Q9CRB9 |
| Coiled-coil-helix-coiled-coil-helix domain-containir | Q91VN4 |
| Cold shock domain-containing protein E1 | Q91W50 |
| COMM domain-containing protein 3 | Q63829 |
| Complexin-1 | P63040 |
| Complexin-2 | P84086 |
| Connector enhancer of kinase suppressor of ras | Q80YA9 |
| Constitutive coactivator of PPAR-gamma-like pro | Q6A0A9 |
| Constitutive coactivator of PPAR-gamma-like pro | Q8C3F2 |
| Contactin-1 | P12960 |
| Contactin-associated protein 1 | O54991 |
| Contactin-associated protein-like 2 | Q9CPW0 |
| COP9 signalosome complex subunit 1 | Q99LD4 |
| COP9 signalosome complex subunit 2 | P61202 |
| COP9 signalosome complex subunit 3 | O88543 |
| COP9 signalosome complex subunit 4 | O88544 |
| COP9 signalosome complex subunit 5 | O35864 |
| COP9 signalosome complex subunit 6 | O88545 |
| COP9 signalosome complex subunit 7a | Q9CZ04 |
| Copine-1 | Q8C166 |
| Copine-2 | P59108 |
| Copine-4 | Q8BLR2 |
| Copine-6 | Q9Z140 |
| Copine-7 | Q0VE82 |
| Copine-8 | Q9DC53 |
| Coronin-1A | O89053 |
| Coronin-1B | Q9WUM3 |
| Coronin-1C | Q9WUM4 |
| Coronin-2B | Q8BH44 |
| Coronin-7 | Q9D2V7 |
| Coxsackievirus and adenovirus receptor homolog | P97792 |
| Creatine kinase B-type | Q04447 |
| Creatine kinase M-type | P07310 |
| Creatine kinase U-type, mitochondrial | P30275 |
| CREB-regulated transcription coactivator 1 | Q68ED7 |
| C-terminal-binding protein 1 | O88712 |
| CTP synthase 1 | P70698 |
| CUGBP Elav-like family member 2 | Q9Z0H4 |
| Coproporphyrinogen-III oxidase, mitochondrial | P36552 |
| Coronin-2A | Q8C0P5 |
| Corticotropin-releasing factor-binding protein | Q60571 |
| Craniofacial development protein 1 | O88271 |
| Crk-like protein | P47941 |
| C-terminal-binding protein 2 | P56546 |
| CTP synthase 2 | P70303 |
| CUB and sushi domain-containing protein 1 | Q923L3 |
| CUGBP Elav-like family member 4 | Q7TSY6 |
| Cyclin-dependent kinase 9 | Q99J95 |
| Cyclin-dependent kinase-like 5 | Q3UTQ8 |
| Cyclin-Y | Q8BGU5 |
| Cysteine and histidine-rich domain-containing pro | Q9D1P4 |
| Cysteine-rich with EGF-like domain protein 1 | Q91XD7 |
| Cytochrome b-c1 complex subunit 9 | Q8R1I1 |
| Cytoplasmic dynein 2 heavy chain 1 | Q45VK7 |
| Cytoplasmic phosphatidylinositol transfer protein | Q8K4R4 |
| Cytoplasmic polyadenylation element-binding pro | Q7TN99 |
| Cytoplasmic protein NCK2 | O55033 |
| Cytosolic purine 5'-nucleotidase | Q3V1L4 |
| DCN1-like protein 1 | Q9QZ73 |
| Dedicator of cytokinesis protein 10 | Q8BZN6 |
| Dedicator of cytokinesis protein 7 | Q8R1A4 |
| Dehydrogenase/reductase SDR family member 4 | Q99LB2 |
| Dendrin | Q80TS7 |
| Density-regulated protein | Q9CQJ6 |
| Deoxyhypusine synthase | Q3TXU5 |
| Diablo homolog, mitochondrial | Q9JIQ3 |
| Diphosphomevalonate decarboxylase | Q99JF5 |
| Disco-interacting protein 2 homolog A | Q8BWT5 |
| Disintegrin and metalloproteinase domain-contair | Q9R1V4 |
| Disintegrin and metalloproteinase domain-contair | Q9R1V7 |
| DNA topoisomerase 1 | Q04750 |
| DNA topoisomerase 2-beta | Q64511 |
| DNA-binding protein A | Q9JKB3 |
| DNA-directed RNA polymerase I subunit RPA1 | O35134 |
| DNA-directed RNA polymerase II subunit RPB2 | Q8CFI7 |
| DnaJ homolog subfamily B member 14 | Q149L6 |
| DnaJ homolog subfamily B member 5 | O89114 |
| DnaJ homolog subfamily C member 10 | Q9DC23 |
| DnaJ homolog subfamily C member 3 | Q91YW3 |
| Dolichyl-diphosphooligosaccharide--protein glyco: | Q9DBG6 |
| Dolichyl-diphosphooligosaccharide--protein glyco: | P46978 |
| Dolichyl-diphosphooligosaccharide--protein glyco: | Q3TDQ1 |
| Down syndrome critical region protein 3 homolog | O35075 |
| Dual specificity mitogen-activated protein kinase l | Q8CE90 |
| Dystrophia myotonica WD repeat-containing prote | Q08274 |
| E3 ubiquitin-protein ligase HECW1 | Q8K4P8 |
| E3 ubiquitin-protein ligase MIB2 | Q8R516 |
| E3 ubiquitin-protein ligase TRIM23 | Q8BGX0 |
| E3 ubiquitin-protein ligase UBR5 | Q80TP3 |
| EF-hand domain-containing family member A2 | Q9CTY5 |
| Electroneutral sodium bicarbonate exchanger 1 | Q8JZR6 |
| Elongator complex protein 2 | Q91WG4 |
| Embigin | P21995 |
| Ena/VASP-like protein | P70429 |
| Endoplasmic reticulum-Golgi intermediate compa | Q9DC16 |
| Endothelial differentiation-related factor 1 | Q9JMG1 |
| Enoyl-CoA delta isomerase 2, mitochondrial | Q9WUR2 |
| Ensconsin | O88735 |
| Ephexin-1 | Q8CHT1 |
| Ephrin type-B receptor 2 | P54763 |
| Ephrin-B2 | P52800 |
| Erlin-1 | Q91X78 |
| Estradiol 17-beta-dehydrogenase 11 | Q9EQ06 |
| Estradiol 17-beta-dehydrogenase 8 | P50171 |
| Etoposide-induced protein 2.4 | Q61070 |
| Eukaryotic translation initiation factor 3 subunit G | Q9Z1D1 |
| Eukaryotic translation initiation factor 4E | P63073 |
| Eukaryotic translation initiation factor 5B | Q05D44 |
| Exocyst complex component 5 | Q3TPX4 |
| Extended synaptotagmin-2 | Q3TZZ7 |
| Fas apoptotic inhibitory molecule 1 | Q9WUD8 |
| Fasciculation and elongation protein zeta-1 | Q8K0X8 |
| F-box only protein 41 | Q6NS60 |
| FERM, RhoGEF and pleckstrin domain-containing | Q91VS8 |
| Ferrochelatase, mitochondrial | P22315 |
| Fibroblast growth factor 1 | P61148 |
| Formin-2 | Q9JL04 |
| Four and a half LIM domains protein 2 | O70433 |
| Fractalkine | O35188 |
| FTS and Hook-interacting protein | Q3U2I3 |
| Fucose mutarotase | Q8R2K1 |
| Fumarylacetoacetate hydrolase domain-containin | Q3TC72 |
| FYVE, RhoGEF and PH domain-containing protei | Q91ZT5 |
| G patch domain-containing protein 8 | A2A6A1 |
| G protein-regulated inducer of neurite outgrowth : | Q8BWS5 |
| Galactokinase | Q9R0N0 |
| Gamma-aminobutyric acid receptor subunit alpha | Q8BHJ7 |
| Gamma-aminobutyric acid receptor subunit beta-: | P63137 |
| Potassium voltage-gated channel subfamily H me | Q60603 |
| Potassium voltage-gated channel subfamily KQT | Q9Z351 |
| Potassium voltage-gated channel subfamily KQT | Q9JK97 |
| Pre-B-cell leukemia transcription factor-interacting | Q3TVI8 |
| Probable global transcription activator SNF2L1 | Q6PGB8 |
| Probable hydrolase PNKD | Q69ZP3 |
| Proline-rich transmembrane protein 3 | Q6PE13 |
| Prominin-1 | O54990 |
| Prostaglandin F2 receptor negative regulator | Q9WV91 |
| Protein canopy homolog 4 | Q8BQ47 |
| Protein cappuccino | Q8VED2 |
| Protein CASC4 | Q6P2L7 |
| Protein dopey-1 | Q8BL99 |
| Protein Dos | Q66L44 |
| Protein FAM115A | Q8BNE1 |
| Protein FAM118A | Q91YN1 |
| Protein FAM131B | Q3TY60 |
| Protein FAM133B | Q9CVI2 |
| Protein FAM177A1 | Q8BR63 |
| Protein FAM19A2 | Q7TPG7 |
| Protein FAM50B | Q9WTJ8 |
| Protein FAM91A1 | Q3UVG3 |
| Protein FAM98A | Q3TJZ6 |
| Protein jagunal homolog 1 | Q5XKN4 |
| Protein LAP2 | Q80TH2 |
| Protein Njmu-R1 | Q9CYI0 |
| Protein odr-4 homolog | Q4PJX1 |
| Protein phosphatase 1 regulatory subunit 12B | Q8BG95 |
| Protein phosphatase 1 regulatory subunit 14C | Q8R4S0 |
| Protein phosphatase 1L | Q8BHN0 |
| Protein syndesmos | Q8VHN8 |
| Protein transport protein Sec61 subunit alpha isof | P61620 |
| Protein-glutamine gamma-glutamyltransferase 2 | P21981 |
| Protocadherin Fat 3 | Q8BNA6 |
| Putative E3 ubiquitin-protein ligase UBR7 | Q8BU04 |
| Putative hydrolase RBBP9 | O88851 |
| Putative transferase CAF17 homolog, mitochondr | Q8CAK1 |
| R3H and coiled-coil domain-containing protein 1 | Q8BSI6 |
| Rab11 family-interacting protein 1 | Q9D620 |
| Ragulator complex protein LAMTOR1 | Q9CQ22 |
| Ras-related GTP-binding protein C | Q99K70 |
| Ras-related GTP-binding protein D | Q7TT45 |
| Ras-related protein Rab-2B | P59279 |
| Ras-related protein Rab-31 | Q921E2 |
| Ras-specific guanine nucleotide-releasing factor F | A2AR50 |
| Receptor tyrosine-protein kinase erbB-2 | P70424 |
| Receptor-type tyrosine-protein phosphatase beta | B2RU80 |
| Regulator of G-protein signaling 17 | Q9QZB0 |
| Regulator of G-protein signaling 7-binding protein | Q8BQP9 |
| Reticulon-4 receptor | Q99PI8 |
| Retinal rod rhodopsin-sensitive cGMP 3',5'-cyclic ; | O55057 |
| Retinoid-inducible serine carboxypeptidase | Q920A5 |
| Rho guanine nucleotide exchange factor 40 | Q3UPH7 |
| Rho-related GTP-binding protein RhoG | P84096 |
| Ribose-phosphate pyrophosphokinase 2 | Q9CS42 |
| Ribosomal biogenesis protein LAS1L | A2BE28 |
| Ribosomal protein S6 kinase alpha-5 | Q8C050 |
| Ribulose-phosphate 3-epimerase | Q8VEE0 |
| RING finger and CCCH-type zinc finger domain-c | P0C090 |
| RING finger protein 11 | Q9QYK7 |
| RING finger protein 112 | Q96DY5 |
| RING finger protein 141 | Q99MB7 |
| RNA polymerase II-associated protein 1 | Q80TE0 |
| RNA-binding motif protein, X chromosome | Q9WV02 |
| RNA-binding protein 8A | Q9CWZ3 |
| RNA-binding protein Nova-1 | Q9JKN6 |
| RWD domain-containing protein 4 | Q9CPR1 |
| Secretory carrier-associated membrane protein 2 | Q9ERN0 |
| Selenoprotein O | Q9DBC0 |
| Sequestosome-1 | Q64337 |
| Serine protease HTRA2, mitochondrial | Q9JIY5 |
| Serine/threonine-protein kinase 4 | Q9JI11 |
| Serine/threonine-protein kinase Chk1 | O35280 |
| Serine/threonine-protein kinase D1 | Q62101 |
| Serine/threonine-protein kinase Nek1 | P51954 |
| Serine/threonine-protein kinase Nek2 | O35942 |
| Serine/threonine-protein kinase PRP4 homolog | Q61136 |
| Serine/threonine-protein kinase TBK1 | Q9WUN2 |
| Serine/threonine-protein phosphatase 2A 55 kDa | Q925E7 |
| SH3 and PX domain-containing protein 2A | O89032 |
| SH3 domain-binding protein 4 | Q921I6 |
| SH3 domain-binding protein 5 | Q9Z131 |
| SH3 domain-binding protein 5-like | Q99LH9 |
| SID1 transmembrane family member 1 | Q6AXF6 |
| Sigma non-opioid intracellular receptor 1 | O55242 |
| Signal transducer and activator of transcription 1 | P42225 |
| Sjoegren syndrome/scleroderma autoantigen 1 hc | P56873 |
| SLAIN motif-containing protein 2 | Q8CI08 |
| Small nuclear ribonucleoprotein Sm D2 | P62317 |
| Small nuclear ribonucleoprotein Sm D3 | P62320 |

| | |
|---|---|
| Cullin-1 | Q9WTX6 |
| Cullin-2 | Q9D4H8 |
| Cullin-3 | Q9JLV5 |
| Cullin-4B | A2A432 |
| Cullin-5 | Q9D5V5 |
| Cullin-9 | Q80TT8 |
| Cullin-associated NEDD8-dissociated protein 1 | Q6ZQ38 |
| Cyclin-dependent kinase 14 | O35495 |
| Cyclin-dependent kinase 16 | Q04735 |
| Cyclin-dependent kinase 5 | P49615 |
| Cyclin-G-associated kinase | Q99KY4 |
| Cystathionine beta-synthase | Q91WT9 |
| Cystathionine gamma-lyase | Q8VCN5 |
| Cystatin-B | Q62426 |
| Cystatin-C | P21460 |
| Cysteine and glycine-rich protein 1 | P97315 |
| Cysteine desulfurase, mitochondrial | Q9Z1J3 |
| Cysteine sulfinic acid decarboxylase | Q9DBE0 |
| Cysteine-rich protein 2 | Q9DCT8 |
| Cysteine--tRNA ligase, cytoplasmic | Q9ER72 |
| Cytochrome b5 | P56395 |
| Cytochrome b5 type B | Q9CQX2 |
| Cytochrome b-c1 complex subunit 1, mitochondri | Q9CZ13 |
| Cytochrome b-c1 complex subunit 2, mitochondri | Q9DB77 |
| Cytochrome b-c1 complex subunit 6, mitochondri | P99028 |
| Cytochrome b-c1 complex subunit 7 | Q9D855 |
| Cytochrome b-c1 complex subunit Rieske, mitoch | Q9CR68 |
| Cytochrome c oxidase subunit 2 | P00405 |
| Cytochrome c oxidase subunit 4 isoform 1, mitocl | P19783 |
| Cytochrome c oxidase subunit 5A, mitochondrial | P12787 |
| Cytochrome c oxidase subunit 5B, mitochondrial | P19536 |
| Cytochrome c oxidase subunit 6B1 | P56391 |
| Cytochrome c oxidase subunit 6C | Q9CPQ1 |
| Cytochrome c oxidase subunit 7C, mitochondrial | P17665 |
| Cytochrome c, somatic | P62897 |
| Cytochrome c1, heme protein, mitochondrial | Q9D0M3 |
| Cytoplasmic aconitate hydratase | P28271 |
| Cytoplasmic dynein 1 heavy chain 1 | Q9JHU4 |
| Cytoplasmic dynein 1 intermediate chain 1 | O88485 |
| Cytoplasmic dynein 1 intermediate chain 2 | O88487 |
| Cytoplasmic dynein 1 light intermediate chain 1 | Q8R1Q8 |
| Cytoplasmic dynein 1 light intermediate chain 2 | Q6PDL0 |
| Cytoplasmic FMR1-interacting protein 1 | Q7TMB8 |
| Cytoplasmic FMR1-interacting protein 2 | Q5SQX6 |
| Cytoskeleton-associated protein 4 | Q8BMK4 |
| Cytoskeleton-associated protein 5 | A2AGT5 |
| Cytosol aminopeptidase | Q9CPY7 |
| Cytosolic 10-formyltetrahydrofolate dehydrogena | Q8R0Y6 |
| Cytosolic acyl coenzyme A thioester hydrolase | Q91V12 |
| Cytosolic non-specific dipeptidase | Q9D1A2 |
| Cytospin-B | Q5SXY1 |
| D-3-phosphoglycerate dehydrogenase | Q61753 |
| D-beta-hydroxybutyrate dehydrogenase, mitocho | Q80XN0 |
| DBIRD complex subunit KIAA1967 homolog | Q8VDP4 |
| DCC-interacting protein 13-alpha | Q8K3H0 |
| DCC-interacting protein 13-beta | Q8K3G9 |
| DDB1- and CUL4-associated factor 7 | P61963 |
| D-dopachrome decarboxylase | O35215 |
| Death-associated protein kinase 1 | Q80YE7 |
| Dedicator of cytokinesis protein 3 | Q8CIQ7 |
| Dedicator of cytokinesis protein 4 | P59764 |
| Dedicator of cytokinesis protein 9 | Q8BIK4 |
| Dehydrogenase/reductase SDR family member 1 | Q99L04 |
| Delta(3,5)-Delta(2,4)-dienoyl-CoA isomerase, mit | O35459 |
| Delta-1-pyrroline-5-carboxylate dehydrogenase, r | Q8CHT0 |
| Delta-aminolevulinic acid dehydratase | P10518 |
| Dematin | Q9WV69 |
| Desmoplakin | E9Q557 |
| Destrin | Q9R0P5 |
| Deubiquitinating protein VCIP135 | Q8CDG3 |
| Developmentally-regulated GTP-binding protein 1 | P32233 |
| Developmentally-regulated GTP-binding protein 2 | Q9QXB9 |
| Diacylglycerol kinase beta | Q6NS52 |
| Diacylglycerol kinase epsilon | Q9R1C6 |
| Diacylglycerol kinase gamma | Q91WG7 |
| Diacylglycerol kinase theta | Q6P5E8 |
| Diacylglycerol kinase zeta | Q80UP3 |
| Dihydrolipoyl dehydrogenase, mitochondrial | O08749 |
| Dihydrolipoyllysine-residue acetyltransferase com | Q8BMF4 |
| Dihydrolipoyllysine-residue succinyltransferase co | Q9D2G2 |
| Dihydropteridine reductase | Q8BVI4 |
| Dihydropyrimidinase-related protein 1 | P97427 |
| Dihydropyrimidinase-related protein 2 | O08553 |
| Dihydropyrimidinase-related protein 3 | Q62188 |
| Dihydropyrimidinase-related protein 4 | O35098 |
| Dihydropyrimidinase-related protein 5 | Q9EQF6 |
| Dipeptidyl aminopeptidase-like protein 6 | Q9Z218 |
| Dipeptidyl peptidase 3 | Q99KK7 |
| Diphosphoinositol polyphosphate phosphohydrola | Q9JI46 |
| Diphosphoinositol polyphosphate phosphohydrola | P0C027 |

| | |
|---|---|
| Gap junction alpha-1 protein | P23242 |
| General receptor for phosphoinositides 1-associa | Q9JJA9 |
| General transcription factor IIF subunit 1 | Q3THK3 |
| Glucosamine-6-phosphate isomerase 2 | Q9CRC9 |
| Glucosylceramidase | P17439 |
| Glutamate carboxypeptidase 2 | O35409 |
| Glutamate receptor 3 | Q9Z2W9 |
| Glutamate receptor, ionotropic kainate 2 | P39087 |
| Glutamate receptor-interacting protein 1 | Q925T6 |
| Glutaryl-CoA dehydrogenase, mitochondrial | Q60759 |
| Glutathione reductase, mitochondrial | P47791 |
| Glycine amidinotransferase, mitochondrial | Q9D964 |
| Golgi membrane protein 1 | Q91XA2 |
| Golgi reassembly-stacking protein 2 | Q99JX3 |
| Golgin subfamily A member 1 | Q9CW79 |
| GRAM domain-containing protein 1B | Q80TI0 |
| GRIP and coiled-coil domain-containing protein 1 | Q9D4H2 |
| Growth arrest and DNA damage-inducible protein | Q9CR59 |
| GTPase-activating Rap/Ran-GAP domain-like pro | Q3V0G7 |
| Guanine nucleotide-binding protein subunit beta-4 | P29387 |
| Guanylate kinase | Q64520 |
| Haloacid dehalogenase-like hydrolase domain-co | Q3UGR5 |
| Haloacid dehalogenase-like hydrolase domain-co | Q9CYW4 |
| Hamartin | Q9EP53 |
| HCLS1-associated protein X-1 | O35387 |
| Hemopexin | Q91X72 |
| Heparan sulfate 2-O-sulfotransferase 1 | Q8R3H7 |
| Heterogeneous nuclear ribonucleoprotein A0 | Q9CX86 |
| Heterogeneous nuclear ribonucleoprotein H | O35737 |
| Heterogeneous nuclear ribonucleoproteins C1/C2 | Q9Z204 |
| Hexokinase-2 | O08528 |
| Hexokinase-3 | Q3TRM8 |
| High mobility group protein B1 | P63158 |
| Histone deacetylase 11 | Q91WA3 |
| Histone deacetylase 4 | Q6NZM9 |
| Histone deacetylase 5 | Q9Z2V6 |
| Histone-binding protein RBBP4 | Q60972 |
| Histone-lysine N-methyltransferase MLL2 | Q6PDK2 |
| Homer protein homolog 2 | Q9QWW1 |
| Huntingtin-associated protein 1 | O35668 |
| Hydroxyacylglutathione hydrolase-like protein | Q9DB32 |
| Hydroxymethylglutaryl-CoA lyase, mitochondrial | P38060 |
| Hydroxymethylglutaryl-CoA synthase, mitochondr | P54869 |
| Hydroxysteroid dehydrogenase-like protein 2 | Q2TPA8 |
| IgLON family member 5 | Q8HW98 |
| Immunoglobulin superfamily containing leucine-ri | Q5RKR3 |
| Inactive hydroxysteroid dehydrogenase-like prote | Q8BTX9 |
| Inositol hexakisphosphate and diphosphoinositol- | A2ARP1 |
| Inositol hexakisphosphate and diphosphoinositol- | Q6ZQB6 |
| Inositol monophosphatase 3 | Q80V26 |
| Insulin receptor substrate 2 | P81122 |
| Integrin beta-1 | P09055 |
| Integrin-alpha FG-GAP repeat-containing protein | Q91WI7 |
| Interferon-inducible double stranded RNA-depend | Q9WTX2 |
| Intersectin-2 | Q9Z0R6 |
| Junction-mediating and -regulatory protein | Q9QXM1 |
| Junctophilin-3 | Q9ET77 |
| Junctophilin-4 | Q80WT0 |
| Katanin p60 ATPase-containing subunit A-like 1 | Q8K0T4 |
| Ketosamine-3-kinase | Q8K274 |
| Kinesin-like protein KIF13A | Q9EQW7 |
| Kinesin-like protein KIF21B | Q9QXL1 |
| Kv channel-interacting protein 2 | Q9JJ69 |
| Lanosterol synthase | Q8BLN5 |
| La-related protein 4B | Q6A0A2 |
| La-related protein 6 | Q8BN59 |
| La-related protein 7 | Q05CL8 |
| Latrophilin-2 (Fragment) | Q8JZZ7 |
| LDLR chaperone MESD | Q9ERE7 |
| Leucine zipper protein 1 | Q8R4U7 |
| Leucine zipper protein 2 | Q8BGY3 |
| Leucine-rich repeat and immunoglobulin-like dom | Q3URE9 |
| Leucine-rich repeat and WD repeat-containing pro | Q6P5U7 |
| Leucine-rich repeat flightless-interacting protein 2 | Q91WK0 |
| Leucine-rich repeat-containing protein 16B | Q3UFQ8 |
| Leucine-rich repeat-containing protein 8A | Q80WG5 |
| LIM domain and actin-binding protein 1 | Q9ERG0 |
| Long-chain fatty acid transport protein 4 | Q91VE0 |
| Long-chain-fatty-acid--CoA ligase 4 | Q9QUJ7 |
| Lymphocyte antigen 6H | Q9WUC3 |
| LysM and putative peptidoglycan-binding domain- | Q9D0E3 |
| LysM and putative peptidoglycan-binding domain- | Q9D7V2 |
| Lysophosphatidic acid phosphatase type 6 | Q8BP40 |
| Lysosomal acid phosphatase | P24638 |
| Lysosomal-trafficking regulator | P97412 |
| Lysosome-associated membrane glycoprotein 1 | P11438 |
| Macrophage erythroblast attacher | Q4VC33 |
| MAGUK p55 subfamily member 3 | O88910 |
| MAGUK p55 subfamily member 5 | Q9JLB2 |
| Mannosyl-oligosaccharide glucosidase | Q80UM7 |

| | |
|---|---|
| Small nuclear ribonucleoprotein-associated protei | P27048 |
| Small subunit processome component 20 homolo | Q5XG71 |
| Sodium channel protein type 8 subunit alpha | Q9WTU3 |
| Sodium/hydrogen exchanger 1 | Q61165 |
| Sorting nexin-29 | Q9D3S3 |
| Sphingolipid delta(4)-desaturase DES1 | O09005 |
| Sphingomyelin phosphodiesterase 2 | O70572 |
| Sphingomyelin phosphodiesterase 3 | Q9JJY3 |
| Stromal cell-derived factor 2 | Q9DCT5 |
| Structural maintenance of chromosomes protein 1 | Q9CU62 |
| Sulfhydryl oxidase 1 | Q8BND5 |
| Synaptosomal-associated protein 23 | O09044 |
| Syndecan-4 | O35988 |
| Syntaxin-17 | Q9D0I4 |
| Syntaxin-5 | Q8K1E0 |
| TATA element modulatory factor | B9EKI3 |
| Tax1-binding protein 3 | Q9DBG9 |
| TBC domain-containing protein kinase-like proteir | Q8BM85 |
| TBC1 domain family member 1 | Q60949 |
| Testican-1 | Q62288 |
| Tetraspanin-18 | Q80WR1 |
| Tetratricopeptide repeat protein 39C | Q8VE09 |
| Thioredoxin domain-containing protein 16 | Q7TN22 |
| Thyrotropin receptor | P47750 |
| Tigger transposable element derived 5 | Q499M4 |
| TNFAIP3-interacting protein 1 | Q9WUU8 |
| Tomoregulin-2 | Q9QYM9 |
| Trafficking protein particle complex subunit 1 | Q5NCF2 |
| Trafficking protein particle complex subunit 11 | B2RXC1 |
| Transcription elongation factor SPT6 | Q62383 |
| Transcription factor BTF3 | Q64152 |
| Transcription initiation factor IIB | P62915 |
| Trans-Golgi network integral membrane protein 1 | Q62313 |
| Transient receptor potential cation channel subfar | Q2TV84 |
| Translocation protein SEC62 | Q8BU14 |
| Translocon-associated protein subunit alpha | Q9CY50 |
| Transmembrane and coiled-coil domains protein 2 | Q80W04 |
| Transmembrane emp24 domain-containing protei | Q8R1V4 |
| Transmembrane protein 126B | Q9D1R1 |
| Transmembrane protein 160 | Q9D938 |
| Transmembrane protein 9 | Q9CR23 |
| Transmembrane protein 9B | Q9JJR8 |
| Tripartite motif-containing protein 66 | Q924W6 |
| tRNA (guanine(26)-N(2))-dimethyltransferase | Q3TX08 |
| tRNA methyltransferase 112 homolog | Q9DCG9 |
| Tubulin alpha-1A chain | P68369 |
| Tubulin gamma-2 chain | Q8VCK3 |
| Tudor domain-containing protein 7 | Q8K1H1 |
| Tumor suppressor candidate 3 | Q8BTV1 |
| Type II inositol 1,4,5-trisphosphate 5-phosphatase | Q8K337 |
| Tyrosine-protein kinase Srms | Q62270 |
| U2 small nuclear ribonucleoprotein A' | P57784 |
| U4/U6.U5 tri-snRNP-associated protein 1 | Q9Z315 |
| Ubiquitin carboxyl-terminal hydrolase 37 | Q8C0R0 |
| Ubiquitin carboxyl-terminal hydrolase 46 | P62069 |
| Ubiquitin carboxyl-terminal hydrolase 8 | Q80U87 |
| Ubiquitin domain-containing protein UBFD1 | Q78JW9 |
| Ubiquitin-60S ribosomal protein L40 | P62984 |
| Ubiquitin-conjugating enzyme E2 E2 | Q91W82 |
| Ubiquitin-like domain-containing CTD phosphatas | Q8BGR9 |
| Ufm1-specific protease 2 | Q99K23 |
| Uncharacterized protein C16orf45 homolog | Q8R1Y2 |
| Uncharacterized protein C19orf43 homolog | Q9D735 |
| Uncharacterized protein C2orf47 homolog, mitoch | Q8BHE8 |
| Uncharacterized protein KIAA0528 | Q7TPS5 |
| UPF0317 protein C14orf159 homolog, mitochond | Q8BH86 |
| UPF0485 protein C1orf144 homolog | Q6NXN1 |
| UPF0565 protein C2orf69 homolog | Q9D9H8 |
| UPF0585 protein C16orf13 homolog | Q9DCS2 |
| UPF0668 protein C10orf76 homolog | Q6PD19 |
| Uridine-cytidine kinase 1 | P52623 |
| Uveal autoantigen with coiled-coil domains and ar | Q8CGB3 |
| Vam6/Vps39-like protein | Q8R5L3 |
| Vesicle-associated membrane protein 3 | P63024 |
| V-set and transmembrane domain-containing prot | Q8R0A6 |
| WAS protein family homolog 1 | Q8VDD8 |
| WD repeat-containing protein 61 | Q9ERF3 |
| WD repeat-containing protein 73 | Q9CWR1 |
| WD repeat-containing protein 91 | Q7TMQ7 |
| WD repeat-containing protein C2orf44 homolog | Q6NV72 |
| Xin actin-binding repeat-containing protein 2 | Q4U4S6 |
| YTH domain family protein 1 | P59326 |
| Zinc finger CCCH domain-containing protein 15 | Q3TIV5 |
| Zinc finger SWIM domain-containing protein 4 | Q8C7B8 |
| Zinc finger SWIM domain-containing protein 5 | Q80TC6 |
| Zinc finger SWIM domain-containing protein KIAA | Q3UHH1 |
| Zinc transporter 9 | Q5IRJ6 |
| Zinc transporter ZIP10 | Q6P5F6 |
| **Detected in experiment 3 only - 607 proteins** | |
| 116 kDa U5 small nuclear ribonucleoprotein comp | O08810 |

| | |
|---|---|
| DIS3-like exonuclease 2 | Q8CI75 |
| Disabled homolog 2-interacting protein | Q3UHC7 |
| Disco-interacting protein 2 homolog B | Q3UH60 |
| Disheveled-associated activator of morphogenes | Q8BPM0 |
| Disintegrin and metalloproteinase domain-contair | Q9R1V6 |
| Disks large homolog 1 | Q811D0 |
| Disks large homolog 2 | Q91XM9 |
| Disks large homolog 3 | P70175 |
| Disks large homolog 4 | Q62108 |
| Disks large-associated protein 1 | Q9D415 |
| Disks large-associated protein 2 | Q8BJ42 |
| Disks large-associated protein 3 | Q6PFD5 |
| Disks large-associated protein 4 | B1AZP2 |
| DmX-like protein 2 | Q8BPN8 |
| DNA damage-binding protein 1 | Q3U1J4 |
| DNA-(apurinic or apyrimidinic site) lyase | P28352 |
| DnaJ homolog subfamily A member 1 | P63037 |
| DnaJ homolog subfamily A member 2 | Q9QYJ0 |
| DnaJ homolog subfamily A member 3, mitochond | Q99M87 |
| DnaJ homolog subfamily A member 4 | Q9JMC3 |
| DnaJ homolog subfamily B member 1 | Q9QYJ3 |
| DnaJ homolog subfamily B member 11 | Q99KV1 |
| DnaJ homolog subfamily B member 4 | Q9D832 |
| DnaJ homolog subfamily B member 6 | O54946 |
| DnaJ homolog subfamily C member 11 | Q5U458 |
| DnaJ homolog subfamily C member 7 | Q9QYI3 |
| DnaJ homolog subfamily C member 8 | Q6NZB0 |
| Dolichol-phosphate mannosyltransferase | O70152 |
| Dolichyl-diphosphooligosaccharide--protein glyco | O54734 |
| Dolichyl-diphosphooligosaccharide--protein glyco | Q91YQ5 |
| Drebrin | Q9QXS6 |
| Drebrin-like protein | Q62418 |
| D-tyrosyl-tRNA(Tyr) deacylase 1 | Q9DD18 |
| Dual specificity mitogen-activated protein kinase | P31938 |
| Dual specificity mitogen-activated protein kinase | P47809 |
| Dual specificity mitogen-activated protein kinase | P70236 |
| Dual specificity protein phosphatase 3 | Q9D7X3 |
| Dynactin subunit 1 | O08788 |
| Dynactin subunit 2 | Q99KJ8 |
| Dynactin subunit 3 | Q9Z0Y1 |
| Dynactin subunit 4 | Q8CBY8 |
| Dynamin-1 | P39053 |
| Dynamin-1-like protein | Q8K1M6 |
| Dynamin-2 | P39054 |
| Dynamin-3 | Q8BZ98 |
| Dynamin-like 120 kDa protein, mitochondrial | P58281 |
| Dynein light chain 1, cytoplasmic | P63168 |
| Dynein light chain 2, cytoplasmic | Q9D0M5 |
| Dyslexia-associated protein KIAA0319-like protei | Q8K135 |
| Dystonin | Q91ZU6 |
| Dystrobrevin alpha | Q9D2N4 |
| Dystroglycan | Q62165 |
| Dystrophin | P11531 |
| E3 ubiquitin-protein ligase HECTD1 | Q69ZR2 |
| E3 ubiquitin-protein ligase HECW2 | Q6I6G8 |
| E3 ubiquitin-protein ligase HUWE1 | Q7TMY8 |
| E3 ubiquitin-protein ligase LRSAM1 | Q80ZI6 |
| E3 ubiquitin-protein ligase NEDD4 | P46935 |
| E3 ubiquitin-protein ligase NEDD4-like | Q8CFI0 |
| E3 ubiquitin-protein ligase TRIM32 | Q8CH72 |
| E3 ubiquitin-protein ligase TRIM9 | Q8C7M3 |
| E3 ubiquitin-protein ligase UBR3 | Q5U430 |
| E3 ubiquitin-protein ligase UBR4 | A2AN08 |
| Early endosome antigen 1 | Q8BL66 |
| Echinoderm microtubule-associated protein-like 1 | Q05BC3 |
| Echinoderm microtubule-associated protein-like 2 | Q7TNG5 |
| Echinoderm microtubule-associated protein-like 4 | Q3UMY5 |
| Ectonucleoside triphosphate diphosphohydrolase | O55026 |
| Ectonucleotide pyrophosphatase/phosphodiester | Q8BGN3 |
| EF-hand domain-containing protein D2 | Q9D8Y0 |
| EH domain-binding protein 1 | Q69ZW3 |
| EH domain-containing protein 1 | Q9WVK4 |
| EH domain-containing protein 3 | Q9QXY6 |
| ELAV-like protein 1 | P70372 |
| ELAV-like protein 2 | Q60899 |
| ELAV-like protein 3 | Q60900 |
| Electrogenic sodium bicarbonate cotransporter 1 | O88343 |
| Electron transfer flavoprotein subunit alpha, mitoc | Q99LC5 |
| Electron transfer flavoprotein subunit beta | Q9DCW4 |
| Electron transfer flavoprotein-ubiquinone oxidore | Q921G7 |
| ELKS/Rab6-interacting/CAST family member 1 | Q99MI1 |
| Elongation factor 1-alpha 1 | P10126 |
| Elongation factor 1-alpha 2 | P62631 |
| Elongation factor 1-beta | O70251 |
| Elongation factor 1-delta | P57776 |
| Elongation factor 1-gamma | Q9D8N0 |
| Elongation factor 2 | P58252 |
| Elongation factor G, mitochondrial | Q8K0D5 |
| Elongation factor Tu GTP-binding domain-contair | Q8C0D5 |
| Elongation factor Tu, mitochondrial | Q8BFR5 |
| MAP/microtubule affinity-regulating kinase 3 | Q03141 |
| MAP/microtubule affinity-regulating kinase 4 | Q8CIP4 |
| Melanoma-associated antigen E1 | Q6PCZ4 |
| Membrane-associated guanylate kinase, WW anc | Q6RHR9 |
| Membrane-associated guanylate kinase, WW anc | Q9EQJ9 |
| Metabotropic glutamate receptor 7 | Q68ED2 |
| Metal transporter CNNM3 | Q32NY4 |
| Metallo-beta-lactamase domain-containing protei | Q8BL86 |
| Metallothionein-3 | P28184 |
| Metaxin-2 | O88441 |
| Methionine aminopeptidase 1 | Q8BP48 |
| Methionine aminopeptidase 2 | O08663 |
| Methylosome subunit pICln | Q61189 |
| Microtubule-associated protein 9 | Q3TRR0 |
| Microtubule-associated serine/threonine-protein k | Q3U214 |
| Mitochondrial 10-formyltetrahydrofolate dehydrog | Q8K009 |
| Mitochondrial import inner membrane translocase | P62075 |
| Mitochondrial import inner membrane translocase | Q9CQV1 |
| Mitochondrial import inner membrane translocase | Q9WVA2 |
| Mitochondrial import inner membrane translocase | Q9WV98 |
| Mitochondrial import inner membrane translocase | Q9WV96 |
| Mitochondrial import receptor subunit TOM22 hon | Q9CPQ3 |
| Mitochondrial intermediate peptidase | A6H611 |
| Mitochondrial peptide methionine sulfoxide reduct | Q9D6Y7 |
| Mitogen-activated protein kinase 10 | Q61831 |
| Mitogen-activated protein kinase 8 | Q91Y86 |
| Mitogen-activated protein kinase kinase kinase ki | Q99JP0 |
| Mitogen-activated protein kinase kinase kinase ki | Q8BPM2 |
| MOB kinase activator 1B | Q8BPB0 |
| Monoacylglycerol lipase ABHD6 | Q8R2Y0 |
| MOSC domain-containing protein 2, mitochondria | Q922Q1 |
| MTSS1-like protein | Q6P9S0 |
| Multiple epidermal growth factor-like domains pro | Q8BH27 |
| Myosin phosphatase Rho-interacting protein | P97434 |
| Myosin regulatory light polypeptide 9 | Q9CQ19 |
| Myotubularin-related protein 2 | Q9Z2D1 |
| Myotubularin-related protein 6 | Q8VE11 |
| N-acetylneuraminate lyase | Q9DCJ9 |
| N-acylneuraminate cytidylyltransferase | Q99KK2 |
| NADH dehydrogenase [ubiquinone] 1 alpha subc | Q9DCJ5 |
| NADH dehydrogenase [ubiquinone] 1 beta subco | Q9CQZ6 |
| NADH dehydrogenase [ubiquinone] 1 beta subco | Q3UIU2 |
| NADH dehydrogenase [ubiquinone] 1 beta subco | Q9CQJ8 |
| NADH dehydrogenase [ubiquinone] iron-sulfur pr | Q99LY9 |
| NADH-cytochrome b5 reductase 1 | Q9DB73 |
| N-chimaerin | Q91V57 |
| Neighbor of COX4 | O70378 |
| Neuritin | Q8CFV4 |
| Neuroendocrine convertase 2 | P21661 |
| Neuroendocrine protein 7B2 | P12961 |
| Neurogranin | P60761 |
| Neuropilin-1 | P97333 |
| Neuropilin-2 | O35375 |
| Neuroserpin | O35684 |
| NLR family member X1 | Q3TL44 |
| NT-3 growth factor receptor | Q6VNS1 |
| Nuclear ubiquitous casein and cyclin-dependent k | Q80XU3 |
| Nucleolar protein 58 | Q6DFW4 |
| Nucleolysin TIAR | P70318 |
| OCIA domain-containing protein 1 | Q9CRD0 |
| OCIA domain-containing protein 2 | Q9D8W7 |
| O-phosphoseryl-tRNA(Sec) selenium transferase | Q6P6M7 |
| OX-2 membrane glycoprotein | O54901 |
| Oxysterol-binding protein-related protein 11 | Q8CI95 |
| Oxysterol-binding protein-related protein 6 | Q8BXR9 |
| Oxysterol-binding protein-related protein 9 | A2A8Z1 |
| Palmitoyl-protein thioesterase 1 | O88531 |
| Palmitoyltransferase ZDHHC5 | Q8VDZ4 |
| Paraplegin | Q3ULF4 |
| PDZ domain-containing protein GIPC1 | Q9Z0G0 |
| Peflin | Q8BFY6 |
| Pentatricopeptide repeat-containing protein 3, mit | Q14C51 |
| Peptidyl-glycine alpha-amidating monooxygenase | P97467 |
| Peptidyl-prolyl cis-trans isomerase FKBP1A | P26883 |
| Peptidyl-tRNA hydrolase 2, mitochondrial | Q8R2Y8 |
| Perilipin-3 | Q9DBG5 |
| Peroxiredoxin-4 | O08807 |
| Peroxisomal 2,4-dienoyl-CoA reductase | Q9WV68 |
| Peroxisomal carnitine O-octanoyltransferase | Q9DC50 |
| Peroxisomal membrane protein PEX14 | Q9R0A0 |
| PEST proteolytic signal-containing nuclear protein | Q6P8I4 |
| Phosphatase and actin regulator 3 | Q8BYK5 |
| Phosphatidate phosphatase LPIN2 | Q99PI5 |
| Phosphatidylinositol 3,4,5-trisphosphate-depende | Q3LAC4 |
| Phosphatidylinositol transfer protein beta isoform | P53811 |
| Pinin | O35691 |
| Plasminogen receptor (KT) | Q9D3P8 |
| Platelet-activating factor acetylhydrolase IB subu | Q61206 |
| Platelet-activating factor acetylhydrolase IB subu | Q61205 |
| Platelet-derived growth factor subunit A | P20033 |
| 14-3-3 protein sigma | O70456 |
| 2',5'-phosphodiesterase 12 | Q3TIU4 |
| 26S proteasome non-ATPase regulatory subunit 1 | Q9Z2X2 |
| 28S ribosomal protein S15, mitochondrial | Q9DC71 |
| 28S ribosomal protein S17, mitochondrial | Q9CQE3 |
| 28S ribosomal protein S36, mitochondrial | Q9CQX8 |
| 28S ribosomal protein S9, mitochondrial | Q9D7N3 |
| 2-amino-3-ketobutyrate coenzyme A ligase, mitoc | O88986 |
| 39S ribosomal protein L14, mitochondrial | Q9D1I6 |
| 39S ribosomal protein L18, mitochondrial | Q9CQL5 |
| 39S ribosomal protein L4, mitochondrial | Q9DCU6 |
| 39S ribosomal protein L44, mitochondrial | Q9CY73 |
| 39S ribosomal protein L47, mitochondrial | Q8K2Y7 |
| 3-ketoacyl-CoA thiolase A, peroxisomal | Q921H8 |
| 3-oxoacyl-[acyl-carrier-protein] synthase, mitochor | Q9D404 |
| 40S ribosomal protein S23 | P62267 |
| 45 kDa calcium-binding protein | Q61112 |
| 5'-3' exoribonuclease 2 | Q9DBR1 |
| 55 kDa erythrocyte membrane protein | P70290 |
| 5'-AMP-activated protein kinase subunit beta-2 | Q6PAM0 |
| 5-hydroxytryptamine receptor 6 | Q9R1C8 |
| 60S ribosomal protein L29 | P47915 |
| 80 kDa MCM3-associated protein | Q9WUU9 |
| Acetylcholinesterase | P21836 |
| Actin filament-associated protein 1 | Q80YS6 |
| Actin-binding LIM protein 3 | Q69ZX8 |
| Actin-binding Rho-activating protein | Q8BUZ1 |
| Activating signal cointegrator 1 complex subunit 3 | E9PZJ8 |
| Acyl-CoA dehydrogenase family member 10 | Q8K370 |
| Acyl-CoA dehydrogenase family member 11 | Q80XL6 |
| Adenylate cyclase type 3 | Q8VHH7 |
| Adenylate cyclase type 6 | Q01341 |
| Adenylate cyclase type 8 | P97490 |
| ADP-ribosylation factor GTPase-activating protein | Q9D8S3 |
| A-kinase anchor protein 8 | Q9DBR0 |
| Alkaline phosphatase, tissue-nonspecific isozyme | P09242 |
| Alpha-1-antitrypsin 1-2 | P22599 |
| Alpha-1-antitrypsin 1-3 | Q00896 |
| Alpha-methylacyl-CoA racemase | O09174 |
| Alpha-tocopherol transfer protein-like | Q9D3D0 |
| Anamorsin | Q8WTY4 |
| Anaphase-promoting complex subunit 1 | P53995 |
| Anaphase-promoting complex subunit 5 | Q8BTZ4 |
| Angiotensin-converting enzyme | P09470 |
| Ankycorbin | Q9EP71 |
| Anoctamin-3 | A2AHL1 |
| AP-5 complex subunit zeta-1 | Q3U829 |
| Apoptotic chromatin condensation inducer in the r | Q9JIX8 |
| ARF GTPase-activating protein GIT2 | Q9JLQ2 |
| Arf-GAP with dual PH domain-containing protein 2 | Q8R2V5 |
| Armadillo repeat-containing protein 10 | Q9D0L7 |
| Armadillo repeat-containing X-linked protein 2 | Q6A058 |
| Aryl hydrocarbon receptor nuclear translocator 2 | Q61324 |
| Aspartate beta-hydroxylase domain-containing pr | Q80VP9 |
| Aspartyl/asparaginyl beta-hydroxylase | Q8BSY0 |
| Astrotactin-2 | Q80Z10 |
| ATP synthase mitochondrial F1 complex assembl | Q811I0 |
| ATP-binding cassette sub-family A member 6 | Q8K441 |
| ATP-binding cassette sub-family B member 10, m | Q9JI39 |
| ATP-dependent RNA helicase DDX3Y | Q62095 |
| ATP-dependent RNA helicase SUPV3L1, mitocho | Q80YD1 |
| ATP-dependent zinc metalloprotease YME1L1 | O88967 |
| Attractin | Q9WU60 |
| Autophagy-related protein 101 | Q9D8Z6 |
| Autophagy-related protein 13 | Q91YI1 |
| Autophagy-related protein 2 homolog A | Q6P4T0 |
| Axin interactor, dorsalization-associated protein | Q8C4Q6 |
| Bardet-Biedl syndrome 7 protein homolog | Q8K2G4 |
| B-cell CLL/lymphoma 7 protein family member A | Q9CXE2 |
| Beta-arrestin-2 | Q91YI4 |
| Beta-catenin-like protein 1 | Q9CWL8 |
| Bifunctional polynucleotide phosphatase/kinase | Q9JLV6 |
| BPI fold-containing family B member 4 | A2BGH0 |
| Brain mitochondrial carrier protein 1 | Q9Z2B2 |
| BRCA1-associated protein | Q99MP8 |
| BRISC complex subunit Abro1 | Q3TCJ1 |
| BTB/POZ domain-containing protein KCTD5 | Q8VC57 |
| C2 calcium-dependent domain-containing protein | Q5HZI2 |
| Calcium homeostasis endoplasmic reticulum prote | Q8CGZ0 |
| Calcium-binding mitochondrial carrier protein SCa | A2ASZ8 |
| Calpain-1 catalytic subunit | O35350 |
| Calpain-7 | Q9R1S8 |
| Calpain-9 | Q9D805 |
| cAMP-dependent protein kinase catalytic subunit | P05132 |
| cAMP-regulated phosphoprotein 19 | P56212 |
| cAMP-specific 3',5'-cyclic phosphodiesterase 4A | O89084 |
| Cannabinoid receptor 1 | P47746 |
| Carbonyl reductase family member 4 | Q91VT4 |
| Carboxypeptidase D | O89001 |
| Carnitine O-palmitoyltransferase 1, liver isoform | P97742 |

| | | | | | |
|---|---|---|---|---|---|
| Elongator complex protein 1 | Q7TT37 | Pleckstrin homology domain-containing family A r | Q7TQG1 | CCR4-NOT transcription complex subunit 2 | Q8C5L3 |
| Elongator complex protein 3 | Q9CZX0 | Pleiotrophin | P63089 | CCR4-NOT transcription complex subunit 6-like | Q8VEG6 |
| Endonuclease domain-containing 1 protein | Q8C522 | Pleiotropic regulator 1 | Q922V4 | CDK5 regulatory subunit-associated protein 2 | Q8K389 |
| Endophilin-A1 | Q62420 | Plexin-B1 | Q8CJH3 | CDK5 regulatory subunit-associated protein 3 | Q99LM2 |
| Endophilin-A2 | Q62419 | Poly [ADP-ribose] polymerase 1 | P11103 | CDP-diacylglycerol--inositol 3-phosphatidyltransfe | Q8VDP6 |
| Endophilin-A3 | Q62421 | Poly(U)-binding-splicing factor PUF60 | Q3UEB3 | Centrosome-associated protein CEP250 | Q60952 |
| Endophilin-B1 | Q9JK48 | Polymerase delta-interacting protein 2 | Q91VA6 | Cerebellin-4 | Q8BME9 |
| Endophilin-B2 | Q8R3V5 | Polypyrimidine tract-binding protein 2 | Q91Z31 | Chloride channel CLIC-like protein 1 | Q99LI2 |
| Endoplasmic reticulum metallopeptidase 1 | Q3UVK0 | Polyribonucleotide nucleotidyltransferase 1, mitoc | Q8K1R3 | Choline transporter-like protein 1 | Q6X893 |
| Endoplasmic reticulum resident protein 29 | P57759 | Potassium voltage-gated channel subfamily KQT | Q8K3F6 | Chromodomain-helicase-DNA-binding protein 4 | Q6PDQ2 |
| Endoplasmic reticulum resident protein 44 | Q9D1Q6 | Potassium/sodium hyperpolarization-activated cy | O88704 | Chromodomain-helicase-DNA-binding protein 9 | Q8BYH8 |
| Endoplasmin | P08113 | PRA1 family protein 2 | Q9JIG8 | Cleavage and polyadenylation specificity factor su | Q9CQF3 |
| Engulfment and cell motility protein 2 | Q8BHL5 | PRA1 family protein 3 | Q8R5J9 | C-Myc-binding protein | Q9EQS3 |
| Enhancer of mRNA-decapping protein 4 | Q3UJB9 | Pre-mRNA-processing factor 6 | Q91YR7 | Coiled-coil and C2 domain-containing protein 2A | Q8CFW7 |
| Enolase-phosphatase E1 | Q8BGB7 | Prenylcysteine oxidase-like | Q8C7K6 | Coiled-coil domain-containing protein 12 | Q8R344 |
| Enoyl-CoA delta isomerase 1, mitochondrial | P42125 | Probable arginine--tRNA ligase, mitochondrial | Q3U186 | Coiled-coil domain-containing protein 158 | Q8CDI6 |
| Enoyl-CoA hydratase, mitochondrial | Q8BH95 | Probable ATP-dependent RNA helicase DDX46 | Q569Z5 | Coiled-coil domain-containing protein 41 | Q9D5R3 |
| Ephrin type-A receptor 4 | Q03137 | Probable ATP-dependent RNA helicase DHX36 | Q8VHK9 | Coiled-coil domain-containing protein 85B | Q6PDY0 |
| Ephrin type-A receptor 6 | Q62413 | Probable cation-transporting ATPase 13A1 | Q9EPE9 | Coiled-coil domain-containing protein 85C | E9Q6B2 |
| Ephrin type-B receptor 1 | Q8CBF3 | Probable D-tyrosyl-tRNA(Tyr) deacylase 2 | Q8BHA3 | Coiled-coil domain-containing protein 9 | Q8VC31 |
| Epidermal growth factor receptor | Q01279 | Probable E3 ubiquitin-protein ligase MYCBP2 | Q7TPH6 | COMM domain-containing protein 1 | Q8K4M5 |
| Epidermal growth factor receptor substrate 15 | P42567 | Probable fructose-2,6-bisphosphatase TIGAR | Q8BZA9 | COMM domain-containing protein 8 | Q9CZG3 |
| Epidermal growth factor receptor substrate 15-like | Q60902 | Probable glutamate--tRNA ligase, mitochondrial | Q9CXJ1 | Conserved oligomeric Golgi complex subunit 4 | Q8R1U1 |
| EPM2A-interacting protein 1 | Q8VEH5 | Probable tRNA threonylcarbamoyladenosine bios | Q8BWU5 | Contactin-associated protein-like 4 | Q99P47 |
| Epsin-1 | Q80VP1 | Pro-interleukin-16 | O54824 | COP9 signalosome complex subunit 8 | Q8VBV7 |
| Epsin-2 | Q8CHU3 | Prolactin regulatory element-binding protein | Q9WUQ2 | Copine-5 | Q8JZW4 |
| ERC protein 2 | Q6PH08 | Proline-rich AKT1 substrate 1 | Q9D1F4 | Cyclin-dependent kinase 18 | Q04899 |
| Erlin-2 | Q8BFZ9 | Prostaglandin E synthase 3 | Q9R0Q7 | Cytoplasmic protein NCK1 | Q99M51 |
| Ermin | Q5EBJ4 | Prostaglandin-H2 D-isomerase | O09114 | DDB1- and CUL4-associated factor 5 | Q80T85 |
| ES1 protein homolog, mitochondrial | Q9D172 | Protein arginine N-methyltransferase 8 | Q6PAK3 | DDB1- and CUL4-associated factor 8 | Q8N7N5 |
| Ester hydrolase C11orf54 homolog | Q91V76 | Protein argonaute-1 | Q8CJG1 | DDRGK domain-containing protein 1 | Q80WW9 |
| Estradiol 17-beta-dehydrogenase 12 | O70503 | Protein argonaute-3 | Q8CJF9 | Death domain-associated protein 6 | O35613 |
| Ethanolamine-phosphate cytidylyltransferase | Q922E4 | Protein bicaudal D homolog 2 | Q921C5 | Dedicator of cytokinesis protein 1 | Q8BUR4 |
| Ethylmalonyl-CoA decarboxylase | Q9D9V3 | Protein DDI1 homolog 2 | A2ADY9 | Deleted in bladder cancer protein 1 homolog | Q920P3 |
| Eukaryotic initiation factor 4A-I | P60843 | Protein diaphanous homolog 1 | O08808 | DENN domain-containing protein 5A | Q6PAL8 |
| Eukaryotic initiation factor 4A-II | P10630 | Protein EFR3 homolog B | Q6ZQ18 | Deoxyhypusine hydroxylase | Q99LN9 |
| Eukaryotic initiation factor 4A-III | Q91VC3 | Protein FAM114A2 | Q8VE88 | Dephospho-CoA kinase domain-containing protei | Q8BHC4 |
| Eukaryotic peptide chain release factor GTP-bind | Q8R050 | Protein FAM126B | Q8C729 | Differentially expressed in FDCP 6 | Q8C2K1 |
| Eukaryotic peptide chain release factor subunit 1 | Q8BWY3 | Protein FAM13C | Q9DBR2 | Dihydropyrimidine dehydrogenase [NADP(+)] | Q8CHR6 |
| Eukaryotic translation initiation factor 1A, X-chrom | Q8BMJ3 | Protein FAM54B | Q9CWE0 | Dihydroxyacetone phosphate acyltransferase | P98192 |
| Eukaryotic translation initiation factor 2 subunit 1 | Q6ZWX6 | Protein FAM5B | Q6DFY8 | Dipeptidyl peptidase 1 | P97821 |
| Eukaryotic translation initiation factor 2 subunit 2 | Q99L45 | Protein FAM65A | Q68FE6 | Dipeptidyl peptidase 2 | Q9ET22 |
| Eukaryotic translation initiation factor 2 subunit 3, | Q9Z0N1 | Protein FAM65B | Q80U16 | Discoidin domain-containing receptor 2 | Q62371 |
| Eukaryotic translation initiation factor 2A | Q8BJW6 | Protein FAM81A | Q3UXZ6 | DNA excision repair protein ERCC-6-like | Q8BHK9 |
| Eukaryotic translation initiation factor 3 subunit A | P23116 | Protein farnesyltransferase subunit beta | Q8K2I1 | DNA-directed RNA polymerase II subunit RPB1 | P08775 |
| Eukaryotic translation initiation factor 3 subunit B | Q8JZQ9 | Protein HEXIM1 | Q8R409 | DNA-directed RNA polymerase III subunit RPC6 | Q921X6 |
| Eukaryotic translation initiation factor 3 subunit C | Q8R1B4 | Protein Hook homolog 1 | Q8BIL5 | DNA-directed RNA polymerase, mitochondrial | Q8BKF1 |
| Eukaryotic translation initiation factor 3 subunit D | O70194 | Protein kinase C alpha type | P20444 | DnaJ homolog subfamily C member 5 | P60904 |
| Eukaryotic translation initiation factor 3 subunit E | P60229 | Protein misato homolog 1 | Q2YDW2 | Double-stranded RNA-binding protein Staufen ho | Q8CJ67 |
| Eukaryotic translation initiation factor 3 subunit F | Q9DCH4 | Protein MON2 homolog | Q80TL7 | Dynein heavy chain 12, axonemal | Q3V0Q1 |
| Eukaryotic translation initiation factor 3 subunit H | Q91WK2 | Protein O-glucosyltransferase 1 | Q8BYB9 | Dynein heavy chain 17, axonemal | Q69Z23 |
| Eukaryotic translation initiation factor 3 subunit I | Q9QZD9 | Protein phosphatase 1 regulatory subunit 12C | Q3UMT1 | Dysferlin | Q9ESD7 |
| Eukaryotic translation initiation factor 3 subunit J | Q66JS6 | Protein phosphatase 1 regulatory subunit 28 | Q8C8T7 | E3 SUMO-protein ligase RanBP2 | Q9ERU9 |
| Eukaryotic translation initiation factor 3 subunit K | Q9DBZ5 | Protein PRRC2A | Q7TSC1 | E3 ubiquitin-protein ligase CBL-B | Q3TTA7 |
| Eukaryotic translation initiation factor 3 subunit L | Q8QZY1 | Protein QIL1 | Q8R404 | E3 ubiquitin-protein ligase HERC2 | Q4U2R1 |
| Eukaryotic translation initiation factor 4 gamma 1 | Q6NZJ6 | Protein RCC2 | Q8BK67 | E3 ubiquitin-protein ligase KCMF1 | Q80UY2 |
| Eukaryotic translation initiation factor 4 gamma 2 | Q62448 | Protein S100-A1 | P56565 | E3 ubiquitin-protein ligase pellino homolog 2 | Q8BST6 |
| Eukaryotic translation initiation factor 4 gamma 3 | Q80XI3 | Protein S100-A6 | P14069 | E3 ubiquitin-protein ligase RNF149 | Q3U2C5 |
| Eukaryotic translation initiation factor 4B | Q8BGD9 | Protein SCO1 homolog, mitochondrial | Q5SUC9 | E3 ubiquitin-protein ligase RNF213 | E9Q555 |
| Eukaryotic translation initiation factor 4H | Q9WUK2 | Protein SEC13 homolog | Q9D1M0 | E3 ubiquitin-protein ligase RNF34 | Q99KR6 |
| Eukaryotic translation initiation factor 5 | P59325 | Protein SZT2 | A2A9C3 | E3 ubiquitin-protein ligase synoviolin | Q9DBY1 |
| Eukaryotic translation initiation factor 5A-1 | P63242 | Protein transport protein Sec23B | Q9D662 | E3 ubiquitin-protein ligase TRIM33 | Q99PP7 |
| Excitatory amino acid transporter 1 | P56564 | Protein transport protein Sec61 subunit beta | Q9CQS8 | E3 ubiquitin-protein ligase TTC3 | O88196 |
| Excitatory amino acid transporter 2 | P43006 | Protein tweety homolog 1 | Q9D3A9 | E3 UFM1-protein ligase 1 | Q8CCJ3 |
| Exocyst complex component 1 | Q8R3S6 | Protein tweety homolog 3 | Q6P5F7 | Ectoderm-neural cortex protein 1 | O35709 |
| Exocyst complex component 2 | Q9D4H1 | Protein unc-79 homolog | Q0KK59 | Ectonucleotide pyrophosphatase/phosphodiestera | Q9EQG7 |
| Exocyst complex component 3 | Q6KAR6 | Protein very KIND | Q0KK55 | EF-hand domain-containing family member A1 | Q8CD10 |
| Exocyst complex component 4 | O35382 | Protein WWC2 | Q6NXJ0 | Egl nine homolog 1 | Q91YE3 |
| Exocyst complex component 6B | A6H5Z3 | Protein XRP2 | Q9EPK2 | EMILIN-3 | Q61062 |
| Exocyst complex component 7 | O35250 | Protein zer-1 homolog | Q80ZJ6 | Endonuclease/exonuclease/phosphatase family d | Q3TGW2 |
| Exocyst complex component 8 | Q6PGF7 | Proton myo-inositol cotransporter | Q3UHK1 | Endoplasmic reticulum aminopeptidase 1 | Q9EQH2 |
| Exportin-1 | Q6P5F9 | Pumilio homolog 2 | Q80U58 | Endoribonuclease Dicer | Q8R418 |
| Exportin-2 | Q9ERK4 | Purine-rich element-binding protein gamma | Q8R4E6 | Engulfment and cell motility protein 1 | Q8BPU7 |
| Exportin-5 | Q924C1 | Putative ATP-dependent RNA helicase DHX30 | Q99PU8 | Ephrin type-B receptor 3 | P54754 |
| Exportin-7 | Q9EPK7 | Putative polypeptide N-acetylgalactosaminyltrans | Q7TT15 | Ephrin-B3 | O35393 |
| Exportin-T | Q9CRT8 | Putative RNA-binding protein 3 | O89086 | Epidermal growth factor receptor kinase substrate | Q08509 |
| Ezrin | P26040 | Putative RNA-binding protein Luc7-like 2 | Q7TNC4 | Epididymal secretory protein E1 | Q9Z0J0 |
| F-actin-capping protein subunit alpha-1 | P47753 | Pyrroline-5-carboxylate reductase 2 | Q922Q4 | Epimerase family protein SDR39U1 | Q5M8N4 |
| F-actin-capping protein subunit alpha-2 | P47754 | Rab GTPase-binding effector protein 2 | Q91WG2 | Eukaryotic translation elongation factor 1 epsilon- | Q9D1M4 |
| F-actin-capping protein subunit beta | P47757 | Rab proteins geranylgeranyltransferase compone | Q9QXG2 | Excitatory amino acid transporter 4 | O35544 |
| Far upstream element-binding protein 1 | Q91WJ8 | Rab11 family-interacting protein 5 | Q8R361 | Exosome component 10 | P56960 |
| Far upstream element-binding protein 2 | Q3U0V1 | Ral GTPase-activating protein subunit beta | Q8BQZ4 | Exostosin-like 2 | Q9ES89 |
| Farnesyl pyrophosphate synthase | Q920E5 | Rapamycin-insensitive companion of mTOR | Q6QI06 | Fanconi-associated nuclease 1 | Q69ZT1 |
| FAS-associated factor 1 | P54731 | Ras GTPase-activating protein 3 | Q60790 | FAS-associated factor 2 | Q3TDN2 |
| Fascin | Q61553 | RAS guanyl-releasing protein 1 | Q9Z1S3 | Fatty acid-binding protein 12 | Q9DAK4 |
| Fatty acid synthase | P19096 | Ras-related protein M-Ras | O08989 | F-box only protein 4 | Q8CHQ0 |
| Fatty acid-binding protein, brain | P51880 | Ras-related protein Rab-22A | P35285 | FCH domain only protein 2 | Q3UQN2 |
| Fatty acid-binding protein, epidermal | Q05816 | Ras-related protein Rab-24 | P35290 | FERM and PDZ domain-containing protein 4 | A2AFR3 |
| Fatty acid-binding protein, heart | P11404 | Ras-related protein Rab-27B | Q99P58 | Fermitin family homolog 3 | Q8K1B8 |
| Fatty aldehyde dehydrogenase | P47740 | Ras-related protein Rab-39B | Q8BHC1 | Filamin-A | Q8BTM8 |

| | |
|---|---|
| Fatty-acid amide hydrolase 1 | O08914 |
| F-box only protein 2 | Q80UW2 |
| F-box only protein 22 | Q78JE5 |
| F-box/LRR-repeat protein 16 | A2RT62 |
| FERM domain-containing protein 4A | Q8BIE6 |
| Fermitin family homolog 2 | Q8CIB5 |
| Ferritin heavy chain | P09528 |
| Fibrinogen C domain-containing protein 1 | A2AV25 |
| Fibronectin type III and SPRY domain-containing | Q7TPM6 |
| Flotillin-1 | O08917 |
| Flotillin-2 | Q60634 |
| Focal adhesion kinase 1 | P34152 |
| Formin-binding protein 1 | Q80TY0 |
| Formin-like protein 1 | Q9JL26 |
| Formin-like protein 2 | A2APV2 |
| Fragile X mental retardation protein 1 homolog | P35922 |
| Fragile X mental retardation syndrome-related pro | Q61584 |
| Fragile X mental retardation syndrome-related pro | Q9WVR4 |
| Fructose-bisphosphate aldolase A | P05064 |
| Fructose-bisphosphate aldolase C | P05063 |
| FSD1-like protein | Q8BYN5 |
| Fumarate hydratase, mitochondrial | P97807 |
| Fumarylacetoacetase | P35505 |
| G protein-regulated inducer of neurite outgrowth | Q3UNH4 |
| Galectin-1 | P16045 |
| Gamma-adducin | Q9QYB5 |
| Gamma-aminobutyric acid receptor subunit beta- | P63080 |
| Gamma-aminobutyric acid receptor-associated pr | P60521 |
| Gamma-aminobutyric acid type B receptor subun | Q9WV18 |
| Gamma-aminobutyric acid type B receptor subun | Q80T41 |
| Gamma-enolase | P17183 |
| Gamma-glutamylcyclotransferase | Q9D7X8 |
| Gamma-glutamyltransferase 7 | Q99JP7 |
| Gamma-soluble NSF attachment protein | Q9CWZ7 |
| Gamma-synuclein | Q9Z0F7 |
| Ganglioside-induced differentiation-associated pr | O88741 |
| Ganglioside-induced differentiation-associated pr | Q8VE33 |
| GDP-D-glucose phosphorylase C15orf58 homolo | Q3TLS3 |
| GDP-L-fucose synthase | P23591 |
| GDP-mannose 4,6 dehydratase | Q8K0C9 |
| Gelsolin | P13020 |
| General vesicular transport factor p115 | Q9Z1Z0 |
| Gephyrin | Q8BUV3 |
| Geranylgeranyl pyrophosphate synthase | Q9WTN0 |
| Geranylgeranyl transferase type-2 subunit alpha | Q9JHK4 |
| Geranylgeranyl transferase type-2 subunit beta | P53612 |
| Girdin | Q5SNZ0 |
| Glia-derived nexin | Q07235 |
| Glial fibrillary acidic protein | P03995 |
| Glucosamine-6-phosphate isomerase 1 | O88958 |
| Glucose 1,6-bisphosphate synthase | Q8CAA7 |
| Glucose-6-phosphate 1-dehydrogenase X | Q00612 |
| Glucose-6-phosphate isomerase | P06745 |
| Glucosidase 2 subunit beta | O08795 |
| Glutamate [NMDA] receptor subunit epsilon-1 | P35436 |
| Glutamate [NMDA] receptor subunit epsilon-2 | Q01097 |
| Glutamate [NMDA] receptor subunit zeta-1 | P35438 |
| Glutamate decarboxylase 1 | P48318 |
| Glutamate decarboxylase 2 | P48320 |
| Glutamate dehydrogenase 1, mitochondrial | P26443 |
| Glutamate receptor 1 | P23818 |
| Glutamate receptor 2 | P23819 |
| Glutamate--cysteine ligase catalytic subunit | P97494 |
| Glutamine synthetase | P15105 |
| Glutaredoxin-3 | Q9CQM9 |
| Glutaredoxin-related protein 5, mitochondrial | Q80Y14 |
| Glutathione S-transferase A4 | P24472 |
| Glutathione S-transferase Mu 1 | P10649 |
| Glutathione S-transferase Mu 5 | P48774 |
| Glutathione S-transferase Mu 7 | Q80W21 |
| Glutathione S-transferase omega-1 | O09131 |
| Glutathione S-transferase P 1 | P19157 |
| Glutathione synthetase | P51855 |
| Glyceraldehyde-3-phosphate dehydrogenase | P16858 |
| Glycerol kinase | Q64516 |
| Glycerol-3-phosphate dehydrogenase [NAD(+)], c | P13707 |
| Glycerol-3-phosphate dehydrogenase 1-like prote | Q3ULJ0 |
| Glycerol-3-phosphate dehydrogenase, mitochond | Q64521 |
| Glycerophosphocholine phosphodiesterase GPC | Q8C0L9 |
| Glycerophosphodiester phosphodiesterase 1 | Q9JL56 |
| Glycerophosphodiester phosphodiesterase doma | Q9CRY7 |
| Glycine dehydrogenase [decarboxylating], mitoch | Q91W43 |
| Glycine--tRNA ligase | Q9CZD3 |
| Glycogen phosphorylase, brain form | Q8CI94 |
| Glycogen phosphorylase, muscle form | Q9WUB3 |
| Glycogen synthase kinase-3 alpha | Q2NL51 |
| Glycogen synthase kinase-3 beta | Q9WV60 |
| Glycogenin-1 | Q9R062 |
| Glycolipid transfer protein | Q9JL62 |
| Glycylpeptide N-tetradecanoyltransferase 1 | O70310 |
| Ras-related protein Rab-4A | P56371 |
| Ras-related protein Rab-9B | Q8BHH2 |
| Receptor expression-enhancing protein 5 | Q60870 |
| Receptor-type tyrosine-protein phosphatase epsil | P49446 |
| Regulating synaptic membrane exocytosis protein | Q9EQZ7 |
| Regulator of G-protein signaling 10 | Q9CQE5 |
| Regulator of G-protein signaling 12 | Q8CGE9 |
| Regulator of G-protein signaling 14 | P97492 |
| Regulator of microtubule dynamics protein 1 | Q9DCV4 |
| Rho GTPase-activating protein 44 | Q5SSM3 |
| Rho GTPase-activating protein 5 | P97393 |
| Rho guanine nucleotide exchange factor 9 | Q3UTH8 |
| Ribosomal protein S6 kinase delta-1 | Q8BLK9 |
| RING finger protein 126 | Q91YL2 |
| RNA polymerase-associated protein RTF1 homol | A2AQ19 |
| RNA-binding protein 26 | Q6NZN0 |
| RNA-binding protein Musashi homolog 2 | Q920Q6 |
| Rootletin | Q8CJ40 |
| Roundabout homolog 1 | O89026 |
| Roundabout homolog 2 | Q7TPD3 |
| Sacsin | Q9JLC8 |
| SAM domain and HD domain-containing protein 1 | Q60710 |
| Sarcolemmal membrane-associated protein | Q3URD3 |
| SCY1-like protein 2 | Q8CFE4 |
| Sec1 family domain-containing protein 1 | Q8BRF7 |
| Sel1 repeat-containing protein 1 | Q921H9 |
| Selenocysteine-specific elongation factor | Q9JHW4 |
| Semaphorin-4A | Q62178 |
| Semaphorin-7A | Q9QUR8 |
| Serine beta-lactamase-like protein LACTB, mitoch | Q9EP89 |
| Serine protease HTRA1 | Q9R118 |
| Serine/arginine-rich splicing factor 1 | Q6PDM2 |
| Serine/arginine-rich splicing factor 4 | Q8VE97 |
| Serine/arginine-rich splicing factor 7 | Q8BL97 |
| Serine/threonine-protein kinase 24 | Q99KH8 |
| Serine/threonine-protein kinase A-Raf | P04627 |
| Serine/threonine-protein kinase LMTK3 | Q5XJV6 |
| Serine/threonine-protein kinase MARK2 | Q05512 |
| Serine/threonine-protein kinase TAO3 | Q8BYC6 |
| Serine/threonine-protein kinase ULK3 | Q3U3Q1 |
| Serine/threonine-protein kinase WNK3 | Q80XP9 |
| Serine/threonine-protein phosphatase 4 regulator | Q6P2K6 |
| Serine/threonine-protein phosphatase 6 regulator | Q8R3Q2 |
| Serine--tRNA ligase, mitochondrial | Q9JJL8 |
| SHC-transforming protein 3 | Q61120 |
| Signal peptidase complex catalytic subunit SEC1 | Q9D8V7 |
| Signal peptidase complex subunit 2 | Q9CYN2 |
| Signal transducing adapter molecule 2 | O88811 |
| Signal-induced proliferation-associated 1-like prot | Q80TE4 |
| SLIT and NTRK-like protein 1 | Q810C1 |
| SLIT and NTRK-like protein 3 | Q810B9 |
| SLIT-ROBO Rho GTPase-activating protein 1 | Q91Z69 |
| Small acidic protein | Q9R0P4 |
| Small glutamine-rich tetratricopeptide repeat-cont | Q8VD33 |
| Small ubiquitin-related modifier 3 | Q9Z172 |
| Smith-Magenis syndrome chromosomal region ca | Q3UMB5 |
| Sn1-specific diacylglycerol lipase alpha | Q6WQJ1 |
| Sodium- and chloride-dependent GABA transport | P31648 |
| Sodium channel protein type 9 subunit alpha | Q62205 |
| Sodium/potassium/calcium exchanger 4 | Q8CGQ8 |
| Sorting and assembly machinery component 50 h | Q8BGH2 |
| Sorting nexin-16 | Q8C080 |
| Sorting nexin-17 | Q8BVL3 |
| SPARC | P07214 |
| Spartin | Q8R1X6 |
| Spermatogenesis-associated serine-rich protein 2 | Q8K1N4 |
| Sphingomyelin phosphodiesterase | Q04519 |
| Splicing factor 1 | Q64213 |
| Splicing factor 3A subunit 1 | Q8K4Z5 |
| Splicing factor 3A subunit 2 | Q62203 |
| Splicing factor 3A subunit 3 | Q9D554 |
| SPRY domain-containing protein 4 | Q91WK1 |
| Steroid receptor RNA activator 1 | Q80VJ2 |
| Sterol regulatory element-binding protein 2 | Q3U1N2 |
| Striated muscle-specific serine/threonine-protein k | Q62407 |
| Stromal interaction molecule 1 | P70302 |
| Stromal interaction molecule 2 | P83093 |
| Structural maintenance of chromosomes protein 3 | Q9CW03 |
| Sulfite oxidase, mitochondrial | Q8R086 |
| Synapsin-3 | Q8JZP2 |
| Synaptosomal-associated protein 29 | Q9ERB0 |
| Synaptotagmin-13 | Q9EQT6 |
| Syntabulin | Q8BHS8 |
| Syntaxin-binding protein 6 | Q8R3T5 |
| Syntenin-1 | O08992 |
| Teneurin-2 | Q9WTS5 |
| Tescalcin | Q9JKL5 |
| Testican-2 | Q9ER58 |
| Testis-expressed sequence 2 protein | Q6ZPJ0 |
| Testis-specific Y-encoded-like protein 4 | Q8VD63 |
| Filamin-C | Q8VHX6 |
| Formin-1 | Q05860 |
| Frataxin, mitochondrial | O35943 |
| Friend virus susceptibility protein 1 | P70213 |
| Fructosamine-3-kinase | Q9ER35 |
| FYVE, RhoGEF and PH domain-containing protei | P52734 |
| Galactose-3-O-sulfotransferase 3 | P61315 |
| Galactosylgalactosylxylosylprotein 3-beta-glucuro | P58158 |
| Gamma-aminobutyric acid receptor subunit beta-1 | P50571 |
| Gamma-aminobutyric acid receptor subunit gamm | P22723 |
| GATS-like protein 2 | Q8CAB8 |
| GC-rich sequence DNA-binding factor 2 | Q8BKT3 |
| General transcription factor 3C polypeptide 4 | Q8BMQ2 |
| General transcription factor IIE subunit 2 | Q9D902 |
| Glutamyl-tRNA(Gln) amidotransferase subunit A, | Q9CZN8 |
| Glutaredoxin-1 | Q9QUH0 |
| Glutathione S-transferase kappa 1 | Q9DCM2 |
| Glycosylphosphatidylinositol anchor attachment 1 | Q9WTK3 |
| Golgi SNAP receptor complex member 1 | O88630 |
| Golgi-associated PDZ and coiled-coil motif-contai | Q8BH60 |
| GPI transamidase component PIG-S | Q6PD26 |
| GTP-binding nuclear protein Ran, testis-specific is | Q61820 |
| GTP-binding protein Rit1 | P70426 |
| GTP-binding protein Rit2 | P70425 |
| Guanine nucleotide-binding protein G(I)/G(S)/G(O | Q9JMF3 |
| Guanine nucleotide-binding protein G(s) subunit a | P63094 |
| Guanine nucleotide-binding protein subunit alpha- | P30677 |
| Guanylate cyclase soluble subunit alpha-3 | Q9ERL9 |
| HEAT repeat-containing protein 5B | Q8C547 |
| Heat shock 70 kDa protein 13 | Q8BM72 |
| Heme-binding protein 2 | Q9WU63 |
| Hexaprenyldihydroxybenzoate methyltransferase, | Q8BMS4 |
| Histone deacetylase complex subunit SAP18 | O55128 |
| Histone H1.4 | P43274 |
| Histone H1.5 | P43276 |
| Histone H2A.x | P27661 |
| Histone H2B type 3-B | Q8CGP0 |
| Histone-lysine N-methyltransferase MLL3 | Q8BRH4 |
| Hsp70-binding protein 1 | Q99P31 |
| Hyaluronan and proteoglycan link protein 4 | Q80WM4 |
| Immunoglobulin-like and fibronectin type III domai | Q3KNY0 |
| Inhibitor of nuclear factor kappa-B kinase subunit | O88351 |
| Integral membrane protein 2C | Q91VK4 |
| Integrin alpha-3 | Q62470 |
| Inter-alpha-trypsin inhibitor heavy chain H3 | Q61704 |
| Interferon regulatory factor 2-binding protein 1 | Q8R3Y8 |
| Interleukin-1 receptor accessory protein-like 1 | P59823 |
| Iron-sulfur cluster assembly 1 homolog, mitochon | Q9D924 |
| Iron-sulfur protein NUBPL | Q9CWD8 |
| Isochorismatase domain-containing protein 2A, m | P85094 |
| Junction plakoglobin | Q02257 |
| Kaptin | Q8VCX6 |
| Kazrin | Q69ZS8 |
| Kelch domain-containing protein 2 | Q4G5Y1 |
| Kin of IRRE-like protein 3 | Q8BR86 |
| Kinesin-like protein KIF1B | Q60575 |
| Kinesin-like protein KIF20B | Q80WE4 |
| Kinesin-like protein KIF22 | Q3V300 |
| Kinesin-like protein KIF9 | Q9WV04 |
| Kinesin-like protein KIFC3 | O35231 |
| Krev interaction trapped protein 1 | Q6S5J6 |
| Lamin-B1 | P14733 |
| Laminin subunit alpha-1 | P19137 |
| Leucine-rich repeat LGI family member 2 | Q8K4Z0 |
| Leucine-rich repeat protein SHOC-2 | O88520 |
| Leucine-rich repeat-containing protein 14 | Q8VC16 |
| Lipid phosphate phosphatase-related protein type | Q8VCY8 |
| Lipoyl synthase, mitochondrial | Q99M04 |
| Lys-63-specific deubiquitinase BRCC36 | P46737 |
| Lysine-specific histone demethylase 1A | Q6ZQ88 |
| Lysine-specific histone demethylase 1B | Q8CIG3 |
| Lysophospholipase-like protein 1 | Q3UFF7 |
| Lysosomal alpha-mannosidase | O09159 |
| Lysosomal Pro-X carboxypeptidase | Q7TMR0 |
| Lysosome-associated membrane glycoprotein 2 | P17047 |
| Macrophage-capping protein | P24452 |
| Maleylacetoacetate isomerase | Q9WVL0 |
| Malignant T-cell-amplified sequence 1 | Q9DB27 |
| Malonyl-CoA decarboxylase, mitochondrial | Q99J39 |
| Malonyl-CoA-acyl carrier protein transacylase, mit | Q8R3F5 |
| MAM domain-containing glycosylphosphatidylinos | Q0PMG2 |
| MAM domain-containing glycosylphosphatidylinos | P60755 |
| Melanoma-associated antigen D1 | Q9QYH6 |
| Metabotropic glutamate receptor 4 | Q68EF4 |
| Metal transporter CNNM1 | Q0GA42 |
| Metal transporter CNNM2 | Q3TWN3 |
| Metalloproteinase inhibitor 2 | P25785 |
| Methionine--tRNA ligase, mitochondrial | Q499X9 |
| Microtubule-associated tumor suppressor 1 homo | Q5HZI1 |
| Mitochondrial carrier triple repeat protein 1 | Q5HZI9 |

| | |
|---|---|
| Glycylpeptide N-tetradecanoyltransferase 2 | O70311 |
| Glyoxalase domain-containing protein 4 | Q9CPV4 |
| Glyoxylate reductase/hydroxypyruvate reductase | Q91Z53 |
| Glypican-1 | Q9QZF2 |
| Glypican-4 | P51655 |
| GMP reductase 1 | Q9DCZ1 |
| GMP synthase [glutamine-hydrolyzing] | Q3THK7 |
| Golgi apparatus protein 1 | Q61543 |
| Golgi resident protein GCP60 | Q8BMP6 |
| Golgi to ER traffic protein 4 homolog | Q9D1H7 |
| Golgin subfamily A member 2 | Q921M4 |
| Golgin subfamily A member 3 | P55937 |
| Golgin subfamily A member 4 | Q91VW5 |
| G-protein coupled receptor-associated sorting pro | Q5U4C1 |
| G-protein-signaling modulator 1 | Q6IR34 |
| GRIP and coiled-coil domain-containing protein 2 | Q8CHG3 |
| GRIP1-associated protein 1 | Q8VD04 |
| Growth arrest-specific protein 7 | Q60780 |
| Growth factor receptor-bound protein 2 | Q60631 |
| GrpE protein homolog 1, mitochondrial | Q99LP6 |
| GTP:AMP phosphotransferase, mitochondrial | Q9WTP7 |
| GTPase HRas | Q61411 |
| GTPase KRas | P32883 |
| GTPase-activating protein and VPS9 domain-con | Q6PAR5 |
| GTP-binding nuclear protein Ran | P62827 |
| GTP-binding protein Di-Ras1 | Q91Z61 |
| GTP-binding protein Di-Ras2 | Q5PR73 |
| GTP-binding protein REM 2 | Q8VEL9 |
| GTP-binding protein Rheb | Q921J2 |
| Guanine deaminase | Q9R111 |
| Guanine nucleotide-binding protein G(i) subunit a | B2RSH2 |
| Guanine nucleotide-binding protein G(i) subunit a | P08752 |
| Guanine nucleotide-binding protein G(I)/G(S)/G(C | Q9DAS9 |
| Guanine nucleotide-binding protein G(I)/G(S)/G(C | P63216 |
| Guanine nucleotide-binding protein G(I)/G(S)/G(C | Q61016 |
| Guanine nucleotide-binding protein G(I)/G(S)/G(T | P62874 |
| Guanine nucleotide-binding protein G(I)/G(S)/G(T | P62880 |
| Guanine nucleotide-binding protein G(k) subunit a | Q9DC51 |
| Guanine nucleotide-binding protein G(o) subunit a | P18872 |
| Guanine nucleotide-binding protein G(olf) subunit | Q8CGK7 |
| Guanine nucleotide-binding protein G(q) subunit a | P21279 |
| Guanine nucleotide-binding protein G(z) subunit a | O70443 |
| Guanine nucleotide-binding protein subunit alpha | P21278 |
| Guanine nucleotide-binding protein subunit alpha | P27601 |
| Guanine nucleotide-binding protein subunit beta- | P68040 |
| Guanine nucleotide-binding protein subunit beta- | P62881 |
| Guanine nucleotide-binding protein-like 1 | P36916 |
| Guanylate cyclase soluble subunit beta-1 | O54865 |
| HD domain-containing protein 2 | Q3SXD3 |
| Heat shock 70 kDa protein 12A | Q8K0U4 |
| Heat shock 70 kDa protein 14 | Q99M31 |
| Heat shock 70 kDa protein 1B | P17879 |
| Heat shock 70 kDa protein 4 | Q61316 |
| Heat shock 70 kDa protein 4L | P48722 |
| Heat shock cognate 71 kDa protein | P63017 |
| Heat shock factor-binding protein 1 | Q9CQZ1 |
| Heat shock protein 105 kDa | Q61699 |
| Heat shock protein 75 kDa, mitochondrial | Q9CQN1 |
| Heat shock protein HSP 90-alpha | P07901 |
| Heat shock protein HSP 90-beta | P11499 |
| Heat shock-related 70 kDa protein 2 | P17156 |
| Heme oxygenase 2 | O70252 |
| Heme-binding protein 1 | Q9R257 |
| Hemoglobin subunit alpha | P01942 |
| Hemoglobin subunit beta-1 | P02088 |
| Hepatocyte cell adhesion molecule | Q640R3 |
| Hepatocyte growth factor-regulated tyrosine kinas | Q99LI8 |
| Hepatoma-derived growth factor | P51859 |
| Hepatoma-derived growth factor-related protein 2 | Q3UMU9 |
| Hepatoma-derived growth factor-related protein 3 | Q9JMG7 |
| Heterochromatin protein 1-binding protein 3 | Q3TEA8 |
| Heterogeneous nuclear ribonucleoprotein A/B | Q99020 |
| Heterogeneous nuclear ribonucleoprotein A1 | P49312 |
| Heterogeneous nuclear ribonucleoprotein A3 | Q8BG05 |
| Heterogeneous nuclear ribonucleoprotein D0 | Q60668 |
| Heterogeneous nuclear ribonucleoprotein F | Q9Z2X1 |
| Heterogeneous nuclear ribonucleoprotein K | P61979 |
| Heterogeneous nuclear ribonucleoprotein L | Q8R081 |
| Heterogeneous nuclear ribonucleoprotein L-like | Q921F4 |
| Heterogeneous nuclear ribonucleoprotein M | Q9D0E1 |
| Heterogeneous nuclear ribonucleoprotein Q | Q7TMK9 |
| Heterogeneous nuclear ribonucleoprotein U | Q8VEK3 |
| Heterogeneous nuclear ribonucleoprotein U-like p | Q8VDM6 |
| Heterogeneous nuclear ribonucleoprotein U-like p | Q00PI9 |
| Heterogeneous nuclear ribonucleoproteins A2/B1 | O88569 |
| Hexokinase-1 | P17710 |
| Hippocalcin-like protein 1 | P62748 |
| Hippocalcin-like protein 4 | Q8BGZ1 |
| Histamine N-methyltransferase | Q91VF2 |
| Histidine triad nucleotide-binding protein 1 | P70349 |
| Thioesterase superfamily member 4 | Q3UUI3 |
| Thiopurine S-methyltransferase | O55060 |
| Thioredoxin-related transmembrane protein 2 | Q9D710 |
| THO complex subunit 4 | O08583 |
| Threonine synthase-like 1 | Q8BH55 |
| Thrombospondin type-1 domain-containing protei | Q69ZU6 |
| Thyroid hormone receptor-associated protein 3 | Q569Z6 |
| Thyrotropin-releasing hormone-degrading ectoen | Q8K093 |
| TNF receptor-associated factor 3 | Q60803 |
| Tomoregulin-1 | Q6PFE7 |
| TP53-regulated inhibitor of apoptosis 1 | Q9D8Z2 |
| Trafficking protein particle complex subunit 10 | Q3TLI0 |
| Transcription elongation factor A protein 1 | P10711 |
| Transcription factor RFX3 | P48381 |
| Transcriptional regulator ATRX | Q61687 |
| Transformer-2 protein homolog beta | P62996 |
| Transient receptor potential cation channel subfa | Q9WTR1 |
| Translation initiation factor eIF-2B subunit beta | Q99LD9 |
| Translation initiation factor eIF-2B subunit delta | Q61749 |
| Translin | Q62348 |
| Translin-associated protein X | Q9QZE7 |
| Translocon-associated protein subunit delta | Q62186 |
| tRNA (adenine(58)-N(1))-methyltransferase non-c | Q8CE96 |
| tRNA-dihydrouridine(47) synthase [NAD(P)(+)]-lik | Q91XI1 |
| Trophoblast glycoprotein | Q9Z0L0 |
| Tropomodulin-1 | P49813 |
| Tryptophan 5-hydroxylase 2 | Q8CGV2 |
| Tubby protein | P50586 |
| Tubulin-specific chaperone cofactor E-like protein | Q8C5W3 |
| Tumor suppressor p53-binding protein 1 | P70399 |
| Tyrosine-protein phosphatase non-receptor type 9 | O35239 |
| U1 small nuclear ribonucleoprotein 70 kDa | Q62376 |
| U2 small nuclear ribonucleoprotein B'' | Q9CQI7 |
| U6 snRNA-associated Sm-like protein LSm1 | Q8VC85 |
| U6 snRNA-associated Sm-like protein LSm2 | O35900 |
| U6 snRNA-associated Sm-like protein LSm3 | P62311 |
| U6 snRNA-associated Sm-like protein LSm4 | Q9QXA5 |
| U6 snRNA-associated Sm-like protein LSm7 | Q9CQQ8 |
| Ubiquitin-associated protein 2 | Q91VX2 |
| Ubiquitin-conjugating enzyme E2 Q1 | Q7TSS2 |
| Ubiquitin-fold modifier-conjugating enzyme 1 | Q9CR09 |
| Ubiquitin-protein ligase E3B | Q9ES34 |
| UBX domain-containing protein 2B | Q0KL01 |
| UBX domain-containing protein 7 | Q6P5G6 |
| Uncharacterized glycosyltransferase AGO61 | Q8BW41 |
| Uncharacterized protein C11orf96 homolog | Q3UPL5 |
| Uncharacterized protein C1orf43 homolog | Q8R092 |
| Uncharacterized protein KIAA1107 | Q80TK0 |
| Uncharacterized protein KIAA1671 | Q8BRV5 |
| Unconventional myosin-Ib | P46735 |
| Unconventional myosin-Id | Q5SYD0 |
| Unconventional myosin-IXa | Q8C170 |
| UPF0600 protein C5orf51 homolog | Q8BR90 |
| UPF0687 protein C20orf27 homolog | Q9D1K7 |
| Uridine-cytidine kinase-like 1 | Q91YL3 |
| Vacuolar protein sorting-associated protein 13A | Q5H8C4 |
| Vacuolar protein sorting-associated protein 16 ho | Q920Q4 |
| Vacuolar protein sorting-associated protein 33B | P59016 |
| Vacuolar protein sorting-associated protein 41 ho | Q5KU39 |
| Vacuolar protein sorting-associated protein 53 ho | Q8CCB4 |
| Vacuolar protein sorting-associated protein 8 hom | Q0P5W1 |
| Vesicle-associated membrane protein 7 | P70280 |
| Vesicular integral-membrane protein VIP36 | Q9DBH5 |
| Voltage-dependent L-type calcium channel subur | Q8R0S4 |
| V-type proton ATPase subunit S1 | Q9R1Q9 |
| WASH complex subunit 7 | Q3UMB9 |
| WD and tetratricopeptide repeats protein 1 | Q80ZK9 |
| WD repeat and FYVE domain-containing protein | Q6VNB8 |
| WD repeat domain phosphoinositide-interacting p | Q9CR39 |
| WD repeat domain phosphoinositide-interacting p | Q91VM3 |
| WD repeat-containing protein 11 | Q8K1X1 |
| WD repeat-containing protein 18 | Q4VBE8 |
| WD repeat-containing protein 26 | Q8C6G8 |
| WD repeat-containing protein 34 | Q5U4F6 |
| WD repeat-containing protein 48 | Q8BH57 |
| WD repeat-containing protein 59 | Q8C0M0 |
| WD repeat-containing protein 70 | Q3TWF6 |
| WD repeat-containing protein 82 | Q8BFQ4 |
| WD repeat-containing protein 92 | Q8BGF3 |
| WW domain-binding protein 11 | Q923D5 |
| XK-related protein 4 | Q5GH67 |
| X-linked retinitis pigmentosa GTPase regulator-in | Q9EPQ2 |
| Zinc finger C2HC domain-containing protein 1A | Q8BJH1 |
| Zinc finger FYVE domain-containing protein 1 | Q810J8 |
| Zinc finger ZZ-type and EF-hand domain-containi | Q5SSH7 |
| Zinc transporter SLC39A7 | Q31125 |
| Mitochondrial fission 1 protein | Q9CQ92 |
| Mitochondrial import inner membrane translocase | Q9CQV7 |
| Mitochondrial import receptor subunit TOM20 hom | Q9DCC8 |
| Mitochondrial Rho GTPase 2 | Q8JZN7 |
| Mitogen-activated protein kinase 14 | P47811 |
| Mitogen-activated protein kinase kinase kinase 3 | Q61084 |
| MMS19 nucleotide excision repair protein homolo | Q9D071 |
| MORN repeat-containing protein 4 | Q6PGF2 |
| mTERF domain-containing protein 3, mitochondria | Q8BKY8 |
| Multidrug resistance protein 1A | P21447 |
| Multiple inositol polyphosphate phosphatase 1 | Q9Z2L6 |
| Multiple PDZ domain protein | Q8VBX6 |
| Myosin-1 | Q5SX40 |
| Myosin-4 | Q5SX39 |
| Myosin-6 | Q02566 |
| Myosin-7 | Q91Z83 |
| Myosin-IIIa | Q8K3H5 |
| Myotubularin-related protein 12 | Q80TA6 |
| Na(+)/H(+) exchange regulatory cofactor NHE-RF | Q9JHL1 |
| N-acetylgalactosamine kinase | Q68FH4 |
| N-acetyltransferase 14 | Q8BVG8 |
| NADH dehydrogenase [ubiquinone] 1 alpha subco | Q9D8B4 |
| NADH dehydrogenase [ubiquinone] 1 beta subcor | O09111 |
| NADP-dependent malic enzyme, mitochondrial | Q8BMF3 |
| N-alpha-acetyltransferase 50 | Q6PGB6 |
| NEDD8 ultimate buster 1 | P54729 |
| Negative elongation factor B | Q8C4Y3 |
| Netrin receptor UNC5A | Q8K1S4 |
| Neuron navigator 3 | Q80TN7 |
| Neuronal tyrosine-phosphorylated phosphoinositic | Q8BM65 |
| Neuropilin and tolloid-like protein 2 | Q8BNJ6 |
| Neutral ceramidase | Q9JHE3 |
| NF-kappa-B inhibitor-interacting Ras-like protein 1 | Q8CEC5 |
| NFX1-type zinc finger-containing protein 1 | Q8R151 |
| NmrA-like family domain-containing protein 1 | Q8K2T1 |
| Non-lysosomal glucosylceramidase | Q69ZF3 |
| Non-receptor tyrosine-protein kinase TYK2 | Q9R117 |
| Nuclear receptor coactivator 5 | Q91W39 |
| Nuclear receptor coactivator 7 | Q6DFV7 |
| Nuclear receptor corepressor 2 | Q9WU42 |
| Nuclear receptor-interacting protein 2 | Q9JHR9 |
| Nucleolar MIF4G domain-containing protein 1 | Q3UFM5 |
| Nucleoporin NUP188 homolog | Q6ZQH8 |
| Nucleoredoxin | P97346 |
| NudC domain-containing protein 1 | Q6PIP5 |
| O-acetyl-ADP-ribose deacetylase MACROD1 | Q922B1 |
| Olfactomedin-like protein 2B | Q3V1G4 |
| Optineurin | Q8K3K8 |
| Oral-facial-digital syndrome 1 protein homolog | Q80Z25 |
| OTU domain-containing protein 7A | Q8R554 |
| Pantothenate kinase 1 | Q8K4K6 |
| Pecanex-like protein 1 | Q9QYC1 |
| Peptide-N(4)-(N-acetyl-beta-glucosaminyl)asparag | Q9JI78 |
| Peptidyl-prolyl cis-trans isomerase FKBP2 | P45878 |
| Peptidyl-tRNA hydrolase ICT1, mitochondrial | Q8R035 |
| Pericentriolar material 1 protein | Q9R0L6 |
| Peroxisomal membrane protein 11B | Q9Z210 |
| Pescadillo homolog | Q9EQ61 |
| Phosducin-like protein 3 | Q8BVF2 |
| Phosphatidylinositol 3-kinase catalytic subunit typ | Q6PF93 |
| Phosphatidylinositol 4,5-bisphosphate 3-kinase ca | O35904 |
| Phosphatidylinositol 4-kinase beta | Q8BKC8 |
| Phosphatidylinositol 4-phosphate 3-kinase C2 dor | O70167 |
| Phosphatidylinositol 4-phosphate 5-kinase type-1 | P70182 |
| Phosphatidylserine synthase 1 | Q99LH2 |
| Phospholipase D4 | Q8BG07 |
| Phospholysine phosphohistidine inorganic pyroph | Q9D7I5 |
| Phosphorylase b kinase gamma catalytic chain, te | Q9DB30 |
| Phosphorylase b kinase regulatory subunit alpha, | Q8BWJ3 |
| Platelet-activating factor acetylhydrolase | Q60963 |
| Pleckstrin homology domain-containing family F m | Q91WB4 |
| Pleckstrin homology-like domain family B member | Q8K1N2 |
| Plexin-A2 | P70207 |
| Plexin-A3 | P70208 |
| Plexin-D1 | Q3UH93 |
| Poly(ADP-ribose) glycohydrolase | O88622 |
| Polycomb protein SCMH1 | Q8K214 |
| Potassium voltage-gated channel subfamily A me | P63141 |
| Potassium voltage-gated channel subfamily A me | Q61923 |
| Potassium voltage-gated channel subfamily C me | P15388 |
| Potassium voltage-gated channel subfamily H me | Q920E3 |
| Potassium voltage-gated channel subfamily H me | P59111 |
| Prelamin-A/C | P48678 |
| pre-mRNA 3' end processing protein WDR33 | Q8K4P0 |
| Presqualene diphosphate phosphatase | Q9D4F2 |
| Probable alpha-ketoglutarate-dependent dioxyger | Q8K2U2 |
| Probable ATP-dependent RNA helicase DDX49 | Q4FZF3 |
| Probable ATP-dependent RNA helicase YTHDC2 | B2RR83 |
| Probable cytosolic iron-sulfur protein assembly pro | Q99KN2 |
| Probable G-protein coupled receptor 62 | Q80UC6 |

| | |
|---|---|
| Histidine--tRNA ligase, cytoplasmic | Q61035 |
| Histone deacetylase 6 | Q9Z2V5 |
| Histone H1.0 | P10922 |
| Histone H4 | P62806 |
| Histone-arginine methyltransferase CARM1 | Q9WVG6 |
| Histone-lysine N-methyltransferase setd3 | Q91WC0 |
| Histone-lysine N-methyltransferase SETD7 | Q8VHL1 |
| HIV Tat-specific factor 1 homolog | Q8BGC0 |
| Homeodomain-only protein | Q8R1H0 |
| Homer protein homolog 1 | Q9Z2Y3 |
| Homer protein homolog 3 | Q99JP6 |
| Hsc70-interacting protein | Q99L47 |
| Hsp90 co-chaperone Cdc37 | Q61081 |
| Huntingtin | P42859 |
| Huntingtin-interacting protein 1 | Q8VD75 |
| Huntingtin-interacting protein 1-related protein | Q9JKY5 |
| Hyaluronan and proteoglycan link protein 1 | Q9QUP5 |
| Hydroxyacyl-coenzyme A dehydrogenase, mitoch | Q61425 |
| Hydroxyacylglutathione hydrolase, mitochondrial | Q99KB8 |
| Hydroxymethylglutaryl-CoA synthase, cytoplasmi | Q8JZK9 |
| Hypoxanthine-guanine phosphoribosyltransferase | P00493 |
| Hypoxia up-regulated protein 1 | Q9JKR6 |
| Immunity-related GTPase family Q protein | Q8VIM9 |
| Immunoglobulin superfamily member 21 | Q7TNR6 |
| Immunoglobulin superfamily member 8 | Q8R366 |
| Immunoglobulin-binding protein 1 | Q61249 |
| Importin subunit alpha-3 | O35344 |
| Importin subunit alpha-7 | O35345 |
| Importin subunit beta-1 | P70168 |
| Importin-4 | Q8VI75 |
| Importin-5 | Q8BKC5 |
| Importin-7 | Q9EPL8 |
| Importin-9 | Q91YE6 |
| Inactive dipeptidyl peptidase 10 | Q6NXK7 |
| Inactive phospholipase C-like protein 2 | Q8K394 |
| Inorganic pyrophosphatase | Q9D819 |
| Inorganic pyrophosphatase 2, mitochondrial | Q91VM9 |
| Inosine-5'-monophosphate dehydrogenase 2 | P24547 |
| Inositol 1,4,5-trisphosphate receptor type 1 | P11881 |
| Inositol monophosphatase 1 | O55023 |
| Inositol polyphosphate 1-phosphatase | P49442 |
| Inositol-3-phosphate synthase 1 | Q9JHU9 |
| Inositol-trisphosphate 3-kinase A | Q8R071 |
| Insulin-degrading enzyme | Q9JHR7 |
| Integrin alpha-6 | Q61739 |
| Integrin alpha-M | P05555 |
| Integrin alpha-V | P43406 |
| Intercellular adhesion molecule 5 | Q60625 |
| Interleukin enhancer-binding factor 2 | Q9CXY6 |
| Interleukin enhancer-binding factor 3 | Q9Z1X4 |
| Intersectin-1 | Q9Z0R4 |
| Intracellular hyaluronan-binding protein 4 | Q9JKS5 |
| Inverted formin-2 | Q0GNC1 |
| IQ motif and SEC7 domain-containing protein 1 | Q8R0S2 |
| IQ motif and SEC7 domain-containing protein 2 | Q5DU25 |
| IQ motif and SEC7 domain-containing protein 3 | Q3TES0 |
| Iron-sulfur cluster assembly 2 homolog, mitochon | Q9DCB8 |
| Iron-sulfur cluster assembly enzyme ISCU, mitocl | Q9D7P6 |
| Islet cell autoantigen 1 | P97411 |
| Isobutyryl-CoA dehydrogenase, mitochondrial | Q9D7B6 |
| Isochorismatase domain-containing protein 1 | Q91V64 |
| Isocitrate dehydrogenase [NAD] subunit alpha, m | Q9D6R2 |
| Isocitrate dehydrogenase [NAD] subunit gamma : | P70404 |
| Isocitrate dehydrogenase [NADP] cytoplasmic | O88844 |
| Isocitrate dehydrogenase [NADP], mitochondrial | P54071 |
| Isoleucine--tRNA ligase, cytoplasmic | Q8BU30 |
| Isoleucine--tRNA ligase, mitochondrial | Q8BIJ6 |
| Isovaleryl-CoA dehydrogenase, mitochondrial | Q9JHI5 |
| IST1 homolog | Q9CX00 |
| Janus kinase and microtubule-interacting protein | Q8BVL9 |
| Janus kinase and microtubule-interacting protein | Q5DTN8 |
| Kalirin | A2CG49 |
| Kelch repeat and BTB domain-containing protein | Q8BNW9 |
| Kelch-like protein 22 | Q99JN2 |
| KIF1-binding protein | Q6ZPU9 |
| Kinase suppressor of Ras 1 | Q61097 |
| Kinectin | Q61595 |
| Kinesin heavy chain isoform 5A | P33175 |
| Kinesin heavy chain isoform 5C | P28738 |
| Kinesin light chain 1 | O88447 |
| Kinesin light chain 2 | O88448 |
| Kinesin-1 heavy chain | Q61768 |
| Kinesin-associated protein 3 | P70188 |
| Kinesin-like protein KIF1A | P33173 |
| Kinesin-like protein KIF21A | Q9QXL2 |
| Kinesin-like protein KIF2A | P28740 |
| Kinesin-like protein KIF3A | P28741 |
| Kinesin-like protein KIF3B | Q61771 |
| Kinesin-like protein KIF3C | O35066 |
| Kv channel-interacting protein 4 | Q6PHZ8 |
| Probable tRNA (uracil-O(2)-)-methyltransferase | Q9D2Q2 |
| Probable tRNA threonylcarbamoyladenosine bios | Q61136 |
| Probable Xaa-Pro aminopeptidase 3 | B7ZMP1 |
| Programmed cell death protein 4 | Q61823 |
| Proliferating cell nuclear antigen | P17918 |
| Protein bicaudal D homolog 1 | Q8BR07 |
| Protein C20orf11 homolog | Q9D7M1 |
| Protein CWC15 homolog | Q9JHS9 |
| Protein diaphanous homolog 3 | Q9Z207 |
| Protein disulfide-isomerase TMX3 | Q8BXZ1 |
| Protein Dr1 | Q91WV0 |
| Protein FAM162A | Q9D6U8 |
| Protein FAM171A2 | A2A699 |
| Protein FAM50A | Q9WV03 |
| Protein FAM63A | Q76LS9 |
| Protein FAM73B | Q8BK03 |
| Protein hairless | Q61645 |
| Protein hid-1 homolog | Q8R1F6 |
| Protein mago nashi homolog | P61327 |
| Protein MEMO1 | Q91VH6 |
| Protein NDRG1 | Q62433 |
| Protein NOV homolog | Q64299 |
| Protein OSCP1 | Q8BHW2 |
| Protein PRRC1 | Q3UPH1 |
| Protein PRRC2B | Q7TPM1 |
| Protein prune homolog 2 | Q52KR3 |
| Protein PTHB1 | Q811G0 |
| Protein Red | Q9Z1M8 |
| Protein RMD5 homolog A | Q80YQ8 |
| Protein SCO2 homolog, mitochondrial | Q8VCL2 |
| Protein scribble homolog | Q80U72 |
| Protein sidekick-2 | Q6V4S5 |
| Protein TBRG4 | Q91YM4 |
| Protein unc-80 homolog | Q8BLN6 |
| Protein VAC14 homolog | Q80WQ2 |
| Protein-L-isoaspartate O-methyltransferase doma | Q8BHD8 |
| Protocadherin beta-14 | Q6PB90 |
| Protoporphyrinogen oxidase | P51175 |
| Pseudopodium-enriched atypical kinase 1 | Q69Z38 |
| Pterin-4-alpha-carbinolamine dehydratase 2 | Q9CZL5 |
| Putative deoxyribonuclease TATDN1 | Q6P8M1 |
| Putative deoxyribose-phosphate aldolase | Q91YP3 |
| Putative polypeptide N-acetylgalactosaminyltransf | Q9JJ61 |
| Pyrroline-5-carboxylate reductase 3 | Q9DCC4 |
| Queuine tRNA-ribosyltransferase subunit QTRTD | B8ZXI1 |
| Rab11 family-interacting protein 3 | Q8CHD8 |
| Raftlin-2 | Q8CHX7 |
| Ragulator complex protein LAMTOR2 | Q9JHS3 |
| RanBP-type and C3HC4-type zinc finger-containii | Q9WUB0 |
| Ras-related GTP-binding protein A | Q80X95 |
| Ras-related protein Rab-13 | Q9DD03 |
| Ras-related protein Rab-23 | P35288 |
| Ras-related protein Rab-27A | Q9ERI2 |
| Ras-related protein Rab-28 | Q99KL7 |
| Ras-related protein Rab-33A | P97950 |
| Ras-related protein Rab-40C | Q8VHQ4 |
| Ras-related protein Rap-2c | Q8BU31 |
| Ras-related protein R-Ras | P10833 |
| Ras-specific guanine nucleotide-releasing factor 2 | P70392 |
| Receptor tyrosine-protein kinase erbB-4 | Q61527 |
| Receptor-type tyrosine-protein phosphatase C | P06800 |
| Receptor-type tyrosine-protein phosphatase F | A2A8L5 |
| Receptor-type tyrosine-protein phosphatase O | E9Q612 |
| Receptor-type tyrosine-protein phosphatase-like N | Q60673 |
| Reelin | Q60841 |
| Regulator of G-protein signaling 20 | Q9QZB1 |
| Regulator of microtubule dynamics protein 3 | Q3UJU9 |
| Renin receptor | Q9CYN9 |
| Replication protein A 70 kDa DNA-binding subuni | Q8VEE4 |
| Reticulon-4 receptor-like 1 | Q8K0S5 |
| Retinol dehydrogenase 13 | Q8CEE7 |
| Retrovirus-related Pol polyprotein LINE-1 | P11369 |
| Rho GTPase-activating protein 25 | Q8BYW1 |
| Ribose-5-phosphate isomerase | P47968 |
| Ribosomal protein 63, mitochondrial | Q9CQF8 |
| Ribosomal protein S6 kinase beta-1 | Q8BSK8 |
| Ribosomal protein S6 kinase beta-2 | Q9Z1M4 |
| RNA binding motif protein, X-linked-like-1 | Q91VM5 |
| RNA binding protein fox-1 homolog 1 | Q9JJ43 |
| RNA exonuclease 1 homolog | Q7TT28 |
| RNA-binding protein 14 | Q8C2Q3 |
| RNA-binding protein 25 | B2RY56 |
| RNA-binding protein 27 | Q5SFM8 |
| RNA-binding protein Musashi homolog 1 | Q61474 |
| RNA-binding protein Raly | Q64012 |
| Ryanodine receptor 3 | A2AGL3 |
| SAM and SH3 domain-containing protein 1 | P59808 |
| Scaffold attachment factor B1 | D3YXK2 |
| Secernin-2 | Q8VCA8 |
| Seizure protein 6 | Q7TSK2 |

| | | | |
|---|---|---|---|
| Lactoylglutathione lyase | Q9CPU0 | Selenium-binding protein 1 | P17563 |
| Lambda-crystallin homolog | Q99KP3 | Semaphorin-4D | O09126 |
| LanC-like protein 1 | O89112 | Semaphorin-4F | Q9Z123 |
| LanC-like protein 2 | Q9JJK2 | Septin-10 | Q8C650 |
| La-related protein 1 | Q6ZQ58 | Septin-4 | P28661 |
| L-asparaginase | Q8C0M9 | Serine/arginine repetitive matrix protein 1 | Q52KI8 |
| Late secretory pathway protein AVL9 homolog | Q80U56 | Serine/arginine repetitive matrix protein 2 | Q8BTI8 |
| Latrophilin-1 | Q80TR1 | Serine/threonine-protein kinase LMTK1 | Q80YE4 |
| Latrophilin-3 | Q80TS3 | Serine/threonine-protein kinase Nek6 | Q9ES70 |
| Lethal(2) giant larvae protein homolog 1 | Q80Y17 | Serine/threonine-protein kinase PAK 7 | Q8C015 |
| LETM1 and EF-hand domain-containing protein 1 | Q9Z2I0 | Serine/threonine-protein phosphatase 2A 55 kDa | Q6ZWR4 |
| Leucine zipper transcription factor-like protein 1 | Q9JHQ5 | Serine/threonine-protein phosphatase 2A 55 kDa | Q8BG02 |
| Leucine-rich glioma-inactivated protein 1 | Q9JIA1 | Serine/threonine-protein phosphatase 2A catalytic | P62715 |
| Leucine-rich PPR motif-containing protein, mitoch | Q6PB66 | Serine/threonine-protein phosphatase 6 catalytic s | Q9CQR6 |
| Leucine-rich repeat and immunoglobulin-like dom | Q9D1T0 | Serine/threonine-protein phosphatase 6 regulator | Q7TSI3 |
| Leucine-rich repeat flightless-interacting protein 1 | Q3UZ39 | Serum paraoxonase/arylesterase 2 | Q62086 |
| Leucine-rich repeat-containing protein 16A | Q6EDY6 | SH2 domain-containing protein 5 | Q8JZW5 |
| Leucine-rich repeat-containing protein 16C | Q3V3V9 | SH3 and PX domain-containing protein 2B | A2AAY5 |
| Leucine-rich repeat-containing protein 40 | Q9CRC8 | Short transient receptor potential channel 1 | Q61056 |
| Leucine-rich repeat-containing protein 47 | Q505F5 | Short-chain specific acyl-CoA dehydrogenase, mi | Q07417 |
| Leucine-rich repeat-containing protein 4B | P0C192 | Sia-alpha-2,3-Gal-beta-1,4-GlcNAc-R:alpha 2,8-si | Q64689 |
| Leucine-rich repeat-containing protein 57 | Q9D1G5 | Signal recognition particle receptor subunit alpha | Q9DBG7 |
| Leucine-rich repeat-containing protein 59 | Q922Q8 | Sister chromatid cohesion protein PDS5 homolog | Q6A026 |
| Leucine-rich repeat-containing protein 7 | Q80TE7 | Slit homolog 1 protein | Q80TR4 |
| Leucine--tRNA ligase, cytoplasmic | Q8BMJ2 | Slit homolog 2 protein | Q9R1B9 |
| Leucyl-cystinyl aminopeptidase | Q8C129 | Small conductance calcium-activated potassium c | P58390 |
| Leukocyte surface antigen CD47 | Q61735 | Small nuclear ribonucleoprotein Sm D1 | P62315 |
| Leukotriene A-4 hydrolase | P24527 | Small nuclear ribonucleoprotein-associated protei | P63163 |
| LIM and calponin homology domains-containing p | Q3UH68 | SNW domain-containing protein 1 | Q9CSN1 |
| LIM and SH3 domain protein 1 | Q61792 | Sodium channel subunit beta-2 | Q56A07 |
| Limbic system-associated membrane protein | Q8BLK3 | Sodium/potassium-transporting ATPase subunit b | P97370 |
| Lipid phosphate phosphatase-related protein type | Q7TME0 | Sodium-dependent proline transporter | Q6PGE7 |
| Lipid phosphate phosphohydrolase 3 | Q99JY8 | Solute carrier family 25 member 35 | Q5SWT3 |
| Lipoamide acyltransferase component of branche | P53395 | Solute carrier family 25 member 46 | Q9CQS4 |
| Liprin-alpha-2 | Q8BSS9 | Somatostatin | P60041 |
| Liprin-alpha-3 | P60469 | Sorting nexin-10 | Q9CWT3 |
| LisH domain and HEAT repeat-containing protein | Q148V7 | Sorting nexin-7 | Q9CY18 |
| L-lactate dehydrogenase A chain | P06151 | Spatacsin | Q3UHA3 |
| L-lactate dehydrogenase B chain | P16125 | Spectrin alpha chain, erythrocyte | P08032 |
| Lon protease homolog, mitochondrial | Q8CGK3 | Spermatid perinuclear RNA-binding protein | Q91WM1 |
| Long-chain specific acyl-CoA dehydrogenase, mi | P51174 | Stabilin-2 | Q8R4U0 |
| Long-chain-fatty-acid--CoA ligase 1 | P41216 | Stathmin-4 | P63042 |
| Long-chain-fatty-acid--CoA ligase 3 | Q9CZW4 | Sterol 26-hydroxylase, mitochondrial | Q9DBG1 |
| Long-chain-fatty-acid--CoA ligase 6 | Q91WC3 | Stonin-2 | Q8BZ60 |
| Long-chain-fatty-acid--CoA ligase ACSBG1 | Q99PU5 | Stromal membrane-associated protein 1 | Q91VZ6 |
| Low molecular weight phosphotyrosine protein ph | Q9D358 | Sulfhydryl oxidase 2 | Q3TMX7 |
| Low-density lipoprotein receptor-related protein 1 | Q9JI18 | Sulfotransferase 1A1 | P52840 |
| Lupus La protein homolog | P32067 | SUMO-conjugating enzyme UBC9 | P63280 |
| Lysine--tRNA ligase | Q99MN1 | SWI/SNF-related matrix-associated actin-depend | Q91ZW3 |
| Lysophospholipid acyltransferase LPCAT4 | Q6NVG1 | Synaptic vesicle 2-related protein | Q8BFT9 |
| Lysosomal alpha-glucosidase | P70699 | Synaptic vesicle glycoprotein 2C | Q69ZS6 |
| Macrophage migration inhibitory factor | P34884 | Synaptonemal complex protein 1 | Q62209 |
| Magnesium-dependent phosphatase 1 | Q9D967 | Synaptotagmin-11 | Q9R0N3 |
| MAGUK p55 subfamily member 2 | Q9WV34 | TAF6-like RNA polymerase II p300/CBP-associate | Q8R2K4 |
| MAGUK p55 subfamily member 6 | Q9JLB0 | Tau-tubulin kinase 1 | Q6PCN3 |
| Major prion protein | P04925 | Tax_Id=10090 Gene_Symbol=Krt33b keratin com | S.avidinii |
| Malate dehydrogenase, cytoplasmic | P14152 | TBC1 domain family member 13 | Q8R3D1 |
| Malate dehydrogenase, mitochondrial | P08249 | TBC1 domain family member 23 | Q8K0F1 |
| Malcavernin | Q8K2Y9 | T-cell activation Rho GTPase-activating protein | B2RWW0 |
| Malectin | Q6ZQI3 | Teneurin-1 | Q9WTS4 |
| Mammalian ependymin-related protein 1 | Q99M71 | Tetratricopeptide repeat protein 12 | Q8BW49 |
| Mannose-1-phosphate guanyltransferase beta | Q8BTZ7 | Tetratricopeptide repeat protein 5 | Q99LG4 |
| Mannose-6-phosphate isomerase | Q924M7 | Thioredoxin-related transmembrane protein 1 | Q8VBT0 |
| MAP kinase-activating death domain protein | Q80U28 | THO complex subunit 2 | B1AZI6 |
| MAP7 domain-containing protein 1 | A2AJI0 | THO complex subunit 6 homolog | Q5U4D9 |
| MAP7 domain-containing protein 2 | A2AG50 | THUMP domain-containing protein 3 | P97770 |
| MARCKS-related protein | P28667 | Thymidine kinase 2, mitochondrial | Q9R088 |
| Mast/stem cell growth factor receptor Kit | P05532 | Thyroglobulin | O08710 |
| Matrilin-4 | O89029 | Thyroid hormone receptor alpha | P63058 |
| Matrin-3 | Q8K310 | Tight junction protein ZO-1 | P39447 |
| Medium-chain specific acyl-CoA dehydrogenase, | P45952 | Tissue factor | P20352 |
| Megakaryocyte-associated tyrosine-protein kinas | P41242 | TLD domain-containing protein KIAA1609 | Q8K0P3 |
| Melanoma inhibitory activity protein 3 | Q8BI84 | Toll-like receptor 13 | Q6R5N8 |
| Membrane-associated guanylate kinase, WW and | Q9WVQ1 | Torsin-1A | Q9ER39 |
| Membrane-associated phosphatidylinositol transf | O35954 | Trafficking kinesin-binding protein 1 | Q6PD31 |
| Membrane-associated phosphatidylinositol transf | Q6ZPQ6 | Trafficking protein particle complex subunit 6B | Q9D289 |
| Membrane-associated progesterone receptor com | O55022 | Trans-1,2-dihydrobenzene-1,2-diol dehydrogenas | Q9DBB8 |
| Membrane-associated progesterone receptor com | Q80UU9 | Trans-2,3-enoyl-CoA reductase | Q9CY27 |
| Metabotropic glutamate receptor 1 | P97772 | Transcription factor jun-D | P15066 |
| Metabotropic glutamate receptor 3 | Q9QYS2 | Transformer-2 protein homolog alpha | Q6PFR5 |
| Metabotropic glutamate receptor 5 | Q3UVX5 | Transforming growth factor beta-1-induced transc | Q62219 |
| Methionine adenosyltransferase 2 subunit beta | Q99LB6 | Transforming growth factor-beta receptor-associa | Q3UR70 |
| Methionine--tRNA ligase, cytoplasmic | Q68FL6 | Translation initiation factor IF-2, mitochondrial | Q91YJ5 |
| Methylcrotonoyl-CoA carboxylase beta chain, mit | Q3ULD5 | Translocation protein SEC63 homolog | Q8VHE0 |
| Methylglutaconyl-CoA hydratase, mitochondrial | Q9JLZ3 | Transmembrane 9 superfamily member 2 | P58021 |
| Methylmalonate-semialdehyde dehydrogenase [a | Q9EQ20 | Transmembrane and coiled-coil domains protein 1 | Q69ZZ6 |
| Methylmalonic aciduria type A homolog, mitochon | Q8C7H1 | Transmembrane and TPR repeat-containing prote | Q8BRH0 |
| Methylmalonyl-CoA mutase, mitochondrial | P16332 | Transmembrane emp24 domain-containing protei | Q9R0Q3 |
| Methylosome protein 50 | Q99J09 | Transmembrane prolyl 4-hydroxylase | Q8BG58 |
| Microtubule-actin cross-linking factor 1 | Q9QXZ0 | Transmembrane protein 131 | O70472 |

| | |
|---|---|
| Microtubule-associated protein 1A | Q9QYR6 |
| Microtubule-associated protein 1B | P14873 |
| Microtubule-associated protein 1S | Q8C052 |
| Microtubule-associated protein 2 | P20357 |
| Microtubule-associated protein 4 | P27546 |
| Microtubule-associated protein 6 | Q7TSJ2 |
| Microtubule-associated protein RP/EB family mer | Q61166 |
| Microtubule-associated protein RP/EB family mer | Q8R001 |
| Microtubule-associated protein RP/EB family mer | Q6PER3 |
| Microtubule-associated protein tau | P10637 |
| Microtubule-associated serine/threonine-protein k | Q9R1L5 |
| Microtubule-associated tumor suppressor candida | Q3UHD3 |
| Mimitin, mitochondrial | Q59J78 |
| Misshapen-like kinase 1 | Q9JM52 |
| Mitochondrial 2-oxoglutarate/malate carrier protei | Q9CR62 |
| Mitochondrial carrier homolog 1 | Q791T5 |
| Mitochondrial carrier homolog 2 | Q791V5 |
| Mitochondrial chaperone BCS1 | Q9CZP5 |
| Mitochondrial fission factor | Q6PCP5 |
| Mitochondrial glutamate carrier 1 | Q9D6M3 |
| Mitochondrial glutamate carrier 2 | Q9DB41 |
| Mitochondrial import inner membrane translocase | P62073 |
| Mitochondrial import inner membrane translocase | O35857 |
| Mitochondrial import inner membrane translocase | P62077 |
| Mitochondrial import receptor subunit TOM34 | Q9CYG7 |
| Mitochondrial import receptor subunit TOM40 hor | Q9QYA2 |
| Mitochondrial import receptor subunit TOM70 | Q9CZW5 |
| Mitochondrial inner membrane protein | Q8CAQ8 |
| Mitochondrial Rho GTPase 1 | Q8BG51 |
| Mitochondrial-processing peptidase subunit alpha | Q9DC61 |
| Mitochondrial-processing peptidase subunit beta | Q9CXT8 |
| Mitofusin-1 | Q811U4 |
| Mitofusin-2 | Q80U63 |
| Mitogen-activated protein kinase 1 | P63085 |
| Mitogen-activated protein kinase 3 | Q63844 |
| Mitogen-activated protein kinase 9 | Q9WTU6 |
| MKL/myocardin-like protein 2 | P59759 |
| MOB-like protein phocein | Q6PEB6 |
| Moesin | P26041 |
| Monoacylglycerol lipase ABHD12 | Q99LR1 |
| Monofunctional C1-tetrahydrofolate synthase, mit | Q3V3R1 |
| Monoglyceride lipase | O35678 |
| mRNA cap guanine-N7 methyltransferase | Q9D0L8 |
| mRNA export factor | Q8C570 |
| Multifunctional protein ADE2 | Q9DCL9 |
| Myb-binding protein 1A | Q7TPV4 |
| Myc box-dependent-interacting protein 1 | O08539 |
| Myelin basic protein | P04370 |
| Myelin expression factor 2 | Q8C854 |
| Myelin proteolipid protein | P60202 |
| Myelin-associated glycoprotein | P20917 |
| Myelin-oligodendrocyte glycoprotein | Q61885 |
| Myeloid leukemia factor 2 | Q99KX1 |
| Myeloid-associated differentiation marker | O35682 |
| Myomegalin | Q80YT7 |
| Myosin light polypeptide 6 | Q60605 |
| Myosin regulatory light chain 12B | Q3THE2 |
| Myosin-10 | Q61879 |
| Myosin-14 | Q6URW6 |
| Myosin-9 | Q8VDD5 |
| Myotubularin-related protein 1 | Q9Z2C4 |
| Myotubularin-related protein 3 | Q8K296 |
| Myotubularin-related protein 5 | Q6ZPE2 |
| Myotubularin-related protein 7 | Q9Z2C9 |
| Myristoylated alanine-rich C-kinase substrate | P26645 |
| N(G),N(G)-dimethylarginine dimethylaminohydrola | Q9CWS0 |
| N(G),N(G)-dimethylarginine dimethylaminohydrola | Q99LD8 |
| Na(+)/H(+) exchange regulatory cofactor NHE-RF | P70441 |
| N-acetyl-D-glucosamine kinase | Q9QZ08 |
| NAD kinase domain-containing protein 1 | Q8C5H8 |
| NAD-dependent malic enzyme, mitochondrial | Q99KE1 |
| NAD-dependent protein deacetylase sirtuin-2 | Q8VDQ8 |
| NAD-dependent protein deacylase sirtuin-5, mito | Q8K2C6 |
| NADH dehydrogenase [ubiquinone] 1 alpha subc | Q9D1H6 |
| NADH dehydrogenase [ubiquinone] 1 alpha subc | Q99LC3 |
| NADH dehydrogenase [ubiquinone] 1 alpha subc | Q7TMF3 |
| NADH dehydrogenase [ubiquinone] 1 alpha subc | Q9ERS2 |
| NADH dehydrogenase [ubiquinone] 1 alpha subc | Q9CQ75 |
| NADH dehydrogenase [ubiquinone] 1 alpha subc | Q62425 |
| NADH dehydrogenase [ubiquinone] 1 alpha subc | Q9CPP6 |
| NADH dehydrogenase [ubiquinone] 1 alpha subc | Q9CQZ5 |
| NADH dehydrogenase [ubiquinone] 1 alpha subc | Q9Z1P6 |
| NADH dehydrogenase [ubiquinone] 1 alpha subc | Q9DC69 |
| NADH dehydrogenase [ubiquinone] 1 beta subco | Q9DCS9 |
| NADH dehydrogenase [ubiquinone] 1 beta subco | Q9CQC7 |
| NADH dehydrogenase [ubiquinone] 1 beta subco | Q9CQH3 |
| NADH dehydrogenase [ubiquinone] 1 beta subco | Q9CR61 |
| NADH dehydrogenase [ubiquinone] 1 beta subco | Q9D6J5 |
| NADH dehydrogenase [ubiquinone] 1 subunit C2 | Q9CQ54 |
| NADH dehydrogenase [ubiquinone] flavoprotein 1 | Q91YT0 |

| | |
|---|---|
| Transmembrane protein 132A | Q922P8 |
| Transmembrane protein 132E | Q6IEE6 |
| Transmembrane protein 214 | Q8BM55 |
| Transmembrane protein 245 | B1AZA5 |
| Transmembrane protein 35 | Q9D328 |
| Tropomodulin-3 | Q9JHJ0 |
| Tubulin alpha-3 chain | P05214 |
| Tubulin beta-1 chain | A2AQ07 |
| Tudor and KH domain-containing protein | Q80VL1 |
| Tudor domain-containing protein 3 | Q91W18 |
| Tumor protein p53-inducible protein 11 | Q4QQM4 |
| Type II inositol 3,4-bisphosphate 4-phosphatase | Q6P1Y8 |
| Tyrosine-protein kinase BAZ1B | Q9Z277 |
| Tyrosine-protein kinase Fer | P70451 |
| Tyrosine-protein kinase Lyn | P25911 |
| Tyrosine-protein kinase receptor TYRO3 | P55144 |
| Tyrosine-protein phosphatase non-receptor type 1 | P35821 |
| Tyrosine-protein phosphatase non-receptor type 2 | Q06180 |
| Tyrosine-protein phosphatase non-receptor type 3 | A2ALK8 |
| U4/U6 small nuclear ribonucleoprotein Prp4 | Q9DAW6 |
| Ubiquitin carboxyl-terminal hydrolase 19 | Q3UJD6 |
| Ubiquitin carboxyl-terminal hydrolase 30 | Q3UN04 |
| Ubiquitin-conjugating enzyme E2 H | P62257 |
| Ubiquitin-conjugating enzyme E2 variant 2 | Q9D2M8 |
| Ubiquitin-conjugating enzyme E2 Z | Q3UE37 |
| Ubiquitin-like modifier-activating enzyme 5 | Q8VE47 |
| Ubiquitin-like protein ATG12 | Q9CQY1 |
| UDP-glucose 4-epimerase | Q8R059 |
| Uncharacterized aarF domain-containing protein k | Q9D0L4 |
| Uncharacterized protein C12orf4 homolog | Q91YN0 |
| Uncharacterized protein C12orf43 homolog | Q3UY34 |
| Uncharacterized protein C18orf8 homolog | Q8VC42 |
| Uncharacterized protein C19orf79 homolog | P0DJE0 |
| Uncharacterized protein C1orf50 homolog | Q5EBG8 |
| Uncharacterized protein C8orf46 homolog | Q8BG31 |
| Uncharacterized protein C9orf172 homolog | A2AJA9 |
| Uncharacterized protein C9orf72 homolog | Q6DFW0 |
| Uncharacterized protein KIAA0090 | Q8C7X2 |
| Uncharacterized protein KIAA1109 | A2AAE1 |
| Unconventional myosin-If | P70248 |
| Unconventional myosin-XV | Q9QZZ4 |
| Unconventional myosin-XVI | Q5DU14 |
| Unconventionnal myosin-X | F8VQB6 |
| UPF0760 protein C2orf29 homolog | Q9CWN7 |
| Upstream-binding protein 1 | Q811S7 |
| Vacuolar fusion protein CCZ1 homolog | Q8C1Y8 |
| Vacuolar fusion protein MON1 homolog A | Q6PDG8 |
| Vasodilator-stimulated phosphoprotein | P70460 |
| Very low-density lipoprotein receptor | P98156 |
| Vesicle-associated membrane protein 4 | O70480 |
| Vezatin | Q3ZK22 |
| Voltage-dependent calcium channel gamma-2 sul | O88602 |
| Voltage-dependent L-type calcium channel subun | Q99246 |
| Voltage-dependent L-type calcium channel subun | Q8CC27 |
| Voltage-gated potassium channel subunit beta-1 | P63143 |
| von Willebrand factor A domain-containing proteir | Q3UVV9 |
| WD repeat-containing protein 27 | Q8C5V5 |
| WD repeat-containing protein 5 | P61965 |
| WD repeat-containing protein 6 | Q99ME2 |
| Wiskott-Aldrich syndrome protein family member 2 | Q8BH43 |
| Zinc finger and BTB domain-containing protein 43 | Q9DAI4 |
| Zinc finger protein 638 | Q61464 |
| Zinc finger Ran-binding domain-containing proteir | Q9R020 |
| Zinc finger SWIM domain-containing protein 6 (Fr | Q80TB7 |
| Zinc transporter 1 | Q60738 |
| Zinc transporter 3 | P97441 |

| | |
|---|---|
| NADH dehydrogenase [ubiquinone] flavoprotein 2 | Q9D6J6 |
| NADH dehydrogenase [ubiquinone] iron-sulfur pr | Q91WD5 |
| NADH dehydrogenase [ubiquinone] iron-sulfur pr | Q9DCT2 |
| NADH dehydrogenase [ubiquinone] iron-sulfur pr | Q9CXZ1 |
| NADH dehydrogenase [ubiquinone] iron-sulfur pr | P52503 |
| NADH dehydrogenase [ubiquinone] iron-sulfur pr | Q9DC70 |
| NADH dehydrogenase [ubiquinone] iron-sulfur pr | Q8K3J1 |
| NADH-cytochrome b5 reductase 3 | Q9DCN2 |
| NADH-ubiquinone oxidoreductase 75 kDa subuni | Q91VD9 |
| NADP-dependent malic enzyme | P06801 |
| NADPH:adrenodoxin oxidoreductase, mitochondr | Q61578 |
| NADPH--cytochrome P450 reductase | P37040 |
| N-alpha-acetyltransferase 15, NatA auxiliary subu | Q80UM3 |
| N-alpha-acetyltransferase 25, NatB auxiliary subu | Q8BWZ3 |
| Nardilysin | Q8BHG1 |
| Nasal embryonic luteinizing hormone-releasing ho | Q99NF2 |
| Nascent polypeptide-associated complex subunit | P70670 |
| Nck-associated protein 1 | P28660 |
| NCK-interacting protein with SH3 domain | Q9ESJ4 |
| NEDD8 | P29595 |
| NEDD8-activating enzyme E1 catalytic subunit | Q8C878 |
| NEDD8-activating enzyme E1 regulatory subunit | Q8VBW6 |
| NEDD8-conjugating enzyme Ubc12 | P61082 |
| Neogenin | P97798 |
| Nesprin-1 | Q6ZWR6 |
| Netrin receptor DCC | P70211 |
| Neudesin | Q9CQ45 |
| Neurabin-2 | Q6R891 |
| Neural cell adhesion molecule 1 | P13595 |
| Neural cell adhesion molecule 2 | O35136 |
| Neural cell adhesion molecule L1 | P11627 |
| Neural cell adhesion molecule L1-like protein | P70232 |
| Neural Wiskott-Aldrich syndrome protein | Q91YD9 |
| Neurexin-1-alpha | Q9CS84 |
| Neurexin-3-alpha | Q6P9K9 |
| Neurobeachin | Q9EPN1 |
| Neurocalcin-delta | Q91X97 |
| Neurocan core protein | P55066 |
| Neurochondrin | Q9Z0E0 |
| Neurofascin | Q810U3 |
| Neurofibromin | Q04690 |
| Neurofilament heavy polypeptide | P19246 |
| Neurofilament light polypeptide | P08551 |
| Neurofilament medium polypeptide | P08553 |
| Neuroligin-2 | Q69ZK9 |
| Neuroligin-3 | Q8BYM5 |
| Neurolysin, mitochondrial | Q91YP2 |
| Neuromodulin | P06837 |
| Neuron navigator 1 | Q8CH77 |
| Neuronal calcium sensor 1 | Q8BNY6 |
| Neuronal cell adhesion molecule | Q810U4 |
| Neuronal growth regulator 1 | Q80Z24 |
| Neuronal membrane glycoprotein M6-a | P35802 |
| Neuronal membrane glycoprotein M6-b | P35803 |
| Neuronal migration protein doublecortin | O88809 |
| Neuronal pentraxin receptor | Q99J85 |
| Neuronal pentraxin-1 | Q62443 |
| Neuronal pentraxin-2 | O70340 |
| Neuronal proto-oncogene tyrosine-protein kinase | P05480 |
| Neuronal-specific septin-3 | Q9Z1S5 |
| Neuron-specific calcium-binding protein hippocalc | P84075 |
| Neuroplastin | P97300 |
| Neurotrimin | Q99PJ0 |
| Neutral alpha-glucosidase AB | Q8BHN3 |
| Neutral amino acid transporter A | O35874 |
| Neutral cholesterol ester hydrolase 1 | Q8BLF1 |
| NF-kappa-B essential modulator | O88522 |
| NFU1 iron-sulfur cluster scaffold homolog, mitoch | Q9QZ23 |
| Nicalin | Q8VCM8 |
| Nicastrin | P57716 |
| Nicotinamide phosphoribosyltransferase | Q99KQ4 |
| NIF3-like protein 1 | Q9EQ80 |
| Nischarin | Q80TM9 |
| Nitric oxide synthase, brain | Q9Z0J4 |
| Nitrilase homolog 1 | Q8VDK1 |
| Nodal modulator 1 | Q6GQT9 |
| Noelin | O88998 |
| Noelin-2 | Q8BM13 |
| Non-POU domain-containing octamer-binding pro | Q99K48 |
| NSFL1 cofactor p47 | Q9CZ44 |
| N-terminal EF-hand calcium-binding protein 1 | Q8BG18 |
| N-terminal EF-hand calcium-binding protein 2 | Q91ZP9 |
| Nuclear autoantigenic sperm protein | Q99MD9 |
| Nuclear cap-binding protein subunit 1 | Q3UYV9 |
| Nuclear distribution protein nudE-like 1 | Q9ERR1 |
| Nuclear export mediator factor Nemf | Q8CCP0 |
| Nuclear migration protein nudC | O35685 |
| Nuclear protein localization protein 4 homolog | P60670 |
| Nuclear receptor-binding protein 2 | Q91V36 |
| Nuclear transport factor 2 | P61971 |

| | |
|---|---|
| Nuclease EXOG, mitochondrial | Q8C163 |
| Nucleobindin-1 | Q02819 |
| Nucleobindin-2 | P81117 |
| Nucleolar protein 3 | Q9D1X0 |
| Nucleolin | P09405 |
| Nucleophosmin | Q61937 |
| Nucleoside diphosphate kinase 3 | Q9WV85 |
| Nucleoside diphosphate kinase A | P15532 |
| Nucleoside diphosphate kinase B | Q01768 |
| Nucleosome assembly protein 1-like 1 | P28656 |
| Nucleosome assembly protein 1-like 4 | Q78ZA7 |
| NudC domain-containing protein 3 | Q8R1N4 |
| Numb-like protein | O08919 |
| Obg-like ATPase 1 | Q9CZ30 |
| Oligodendrocyte-myelin glycoprotein | Q63912 |
| Oligoribonuclease, mitochondrial | Q9D8S4 |
| Omega-amidase NIT2 | Q9JHW2 |
| Opioid growth factor receptor | Q99PG2 |
| Ornithine aminotransferase, mitochondrial | P29758 |
| Oxidation resistance protein 1 | Q4KMM3 |
| Oxysterol-binding protein 1 | Q3B7Z2 |
| Oxysterol-binding protein-related protein 1 | Q91XL9 |
| Oxysterol-binding protein-related protein 2 | Q8BX94 |
| Palmdelphin | Q9JHU2 |
| Paralemmin-1 | Q9Z0P4 |
| Paralemmin-2 | Q8BR92 |
| Paraspeckle component 1 | Q8R326 |
| Parathymosin | Q9D0J8 |
| Parkinson disease 7 domain-containing protein 1 | Q8BFQ8 |
| Partner of Y14 and mago | Q8CHP5 |
| Parvalbumin alpha | P32848 |
| PC4 and SFRS1-interacting protein | Q99JF8 |
| Peptidyl-prolyl cis-trans isomerase A | P17742 |
| Peptidyl-prolyl cis-trans isomerase B | P24369 |
| Peptidyl-prolyl cis-trans isomerase D | Q9CR16 |
| Peptidyl-prolyl cis-trans isomerase F, mitochondri | Q99KR7 |
| Peptidyl-prolyl cis-trans isomerase FKBP3 | Q62446 |
| Peptidyl-prolyl cis-trans isomerase FKBP4 | P30416 |
| Peptidyl-prolyl cis-trans isomerase FKBP5 | Q64378 |
| Peptidyl-prolyl cis-trans isomerase FKBP8 | O35465 |
| Peptidyl-prolyl cis-trans isomerase H | Q9D868 |
| Peptidyl-prolyl cis-trans isomerase NIMA-interacti | Q9QUR7 |
| Pericentrin | P48725 |
| Peripheral plasma membrane protein CASK | O70589 |
| Peroxiredoxin-1 | P35700 |
| Peroxiredoxin-2 | Q61171 |
| Peroxiredoxin-5, mitochondrial | P99029 |
| Peroxiredoxin-6 | O08709 |
| Peroxisomal acyl-coenzyme A oxidase 1 | Q9R0H0 |
| Peroxisomal biogenesis factor 19 | Q8VCI5 |
| Peroxisomal multifunctional enzyme type 2 | P51660 |
| PEX5-related protein | Q8C437 |
| PH and SEC7 domain-containing protein 1 | Q5DTT2 |
| PH and SEC7 domain-containing protein 3 | Q2PFD7 |
| Phenylalanine--tRNA ligase alpha subunit | Q8C0C7 |
| Phenylalanine--tRNA ligase beta subunit | Q9WUA2 |
| Phosphatase and actin regulator 1 | Q2M3X8 |
| Phosphate carrier protein, mitochondrial | Q8VEM8 |
| Phosphatidate cytidylyltransferase 2 | Q99L43 |
| Phosphatidylethanolamine-binding protein 1 | P70296 |
| Phosphatidylinositide phosphatase SAC1 | Q9EP69 |
| Phosphatidylinositide phosphatase SAC2 | Q8CDA1 |
| Phosphatidylinositol 3,4,5-trisphosphate-depende | Q69ZK0 |
| Phosphatidylinositol 3-kinase regulatory subunit a | P26450 |
| Phosphatidylinositol 4-phosphate 5-kinase type-1 | O70161 |
| Phosphatidylinositol 5-phosphate 4-kinase type-2 | O70172 |
| Phosphatidylinositol 5-phosphate 4-kinase type-2 | Q80XI4 |
| Phosphatidylinositol 5-phosphate 4-kinase type-2 | Q91XU3 |
| Phosphatidylinositol transfer protein alpha isoforr | P53810 |
| Phosphatidylinositol-binding clathrin assembly pro | Q7M6Y3 |
| Phosphatidylserine decarboxylase proenzyme | Q8BSF4 |
| Phosphoacetylglucosamine mutase | Q9CYR6 |
| Phosphoenolpyruvate carboxykinase [GTP], mito | Q8BH04 |
| Phosphofurin acidic cluster sorting protein 1 | Q8K212 |
| Phosphofurin acidic cluster sorting protein 2 | Q3V3Q7 |
| Phosphoglucomutase-1 | Q9D0F9 |
| Phosphoglucomutase-2 | Q7TSV4 |
| Phosphoglycerate kinase 1 | P09411 |
| Phosphoglycerate mutase 1 | Q9DBJ1 |
| Phosphoglycerate mutase 2 | O70250 |
| Phosphoglycolate phosphatase | Q8CHP8 |
| Phosphoinositide 3-kinase regulatory subunit 4 | Q8VD65 |
| Phospholipase A-2-activating protein | P27612 |
| Phospholipase D3 | O35405 |
| Phosphomannomutase 1 | O35621 |
| Phosphomannomutase 2 | Q9Z2M7 |
| Phosphoribosyl pyrophosphate synthase-associa | Q9D0M1 |
| Phosphoribosyl pyrophosphate synthase-associa | Q8R574 |
| Phosphoribosylformylglycinamidine synthase | Q5SUR0 |
| Phosphorylase b kinase regulatory subunit alpha, | P18826 |

| | |
|---|---|
| Phosphorylase b kinase regulatory subunit beta | Q7TSH2 |
| Phosphoserine aminotransferase | Q99K85 |
| Phytanoyl-CoA hydroxylase-interacting protein | Q8K0S0 |
| Phytanoyl-CoA hydroxylase-interacting protein-lik | Q8BGT8 |
| PI-PLC X domain-containing protein 3 | Q8BLJ3 |
| PITH domain-containing protein 1 | Q8BWR2 |
| Plakophilin-4 | Q68FH0 |
| Plasma membrane calcium-transporting ATPase | Q9R0K7 |
| Plasminogen activator inhibitor 1 RNA-binding pro | Q9CY58 |
| Plastin-2 | Q61233 |
| Plastin-3 | Q99K51 |
| Platelet-activating factor acetylhydrolase IB subu | P63005 |
| Plectin | Q9QXS1 |
| Plexin-A1 | P70206 |
| Plexin-A4 | Q80UG2 |
| Plexin-B2 | B2RXS4 |
| Poliovirus receptor-related protein 1 | Q9JKF6 |
| Poly(ADP-ribose) glycohydrolase ARH3 | Q8CG72 |
| Poly(rC)-binding protein 1 | P60335 |
| Poly(rC)-binding protein 3 | P57722 |
| Polyadenylate-binding protein 1 | P29341 |
| Polyadenylate-binding protein-interacting protein | Q8VE62 |
| Porphobilinogen deaminase | P22907 |
| Prefoldin subunit 1 | Q9CWM4 |
| Prefoldin subunit 2 | O70591 |
| Prefoldin subunit 3 | P61759 |
| Prefoldin subunit 5 | Q9WU28 |
| Prefoldin subunit 6 | Q03958 |
| Pre-mRNA-processing factor 19 | Q99KP6 |
| Prenylcysteine oxidase | Q9CQF9 |
| Pre-rRNA-processing protein TSR2 homolog | Q8C8T8 |
| Presequence protease, mitochondrial | Q8K411 |
| Prickle-like protein 2 | Q80Y24 |
| PRKCA-binding protein | Q62083 |
| Probable ATP-dependent RNA helicase DDX17 | Q501J6 |
| Probable ATP-dependent RNA helicase DDX5 | Q61656 |
| Probable ATP-dependent RNA helicase DDX6 | P54823 |
| Probable cationic amino acid transporter | Q8BXR1 |
| Probable global transcription activator SNF2L2 | Q6DIC0 |
| Probable G-protein coupled receptor 158 | Q8C419 |
| Probable phospholipid-transporting ATPase IA | P70704 |
| Probable phospholipid-transporting ATPase IIA | O70228 |
| Probable threonine--tRNA ligase 2, cytoplasmic | Q8BLY2 |
| Probable ubiquitin carboxyl-terminal hydrolase FA | P70398 |
| Profilin-2 | Q9JJV2 |
| Programmed cell death 6-interacting protein | Q9WU78 |
| Programmed cell death protein 10 | Q8VE70 |
| Programmed cell death protein 5 | P56812 |
| Programmed cell death protein 6 | P12815 |
| Prohibitin | P67778 |
| Prohibitin-2 | O35129 |
| Proliferation-associated protein 2G4 | P50580 |
| Proline dehydrogenase 1, mitochondrial | Q9WU79 |
| Proline synthase co-transcribed bacterial homolo | Q9Z2Y8 |
| Proline-rich transmembrane protein 1 | O35449 |
| Proline-rich transmembrane protein 2 | E9PUL5 |
| Prolow-density lipoprotein receptor-related protei | Q91ZX7 |
| Prolyl endopeptidase | Q9QUR6 |
| Prolyl endopeptidase-like | Q8C167 |
| Propionyl-CoA carboxylase alpha chain, mitochon | Q91ZA3 |
| Propionyl-CoA carboxylase beta chain, mitochon | Q99MN9 |
| ProSAAS | Q9QXV0 |
| ProSAP-interacting protein 1 | A2AHG0 |
| Prostaglandin E synthase 2 | Q8BWM0 |
| Prostamide/prostaglandin F synthase | Q9DB60 |
| Proteasomal ubiquitin receptor ADRM1 | Q9JKV1 |
| Proteasome activator complex subunit 1 | P97371 |
| Proteasome activator complex subunit 2 | P97372 |
| Proteasome activator complex subunit 3 | P61290 |
| Proteasome subunit alpha type-1 | Q9R1P4 |
| Proteasome subunit alpha type-2 | P49722 |
| Proteasome subunit alpha type-3 | O70435 |
| Proteasome subunit alpha type-4 | Q9R1P0 |
| Proteasome subunit alpha type-5 | Q9Z2U1 |
| Proteasome subunit alpha type-6 | Q9QUM9 |
| Proteasome subunit alpha type-7 | Q9Z2U0 |
| Proteasome subunit beta type-1 | O09061 |
| Proteasome subunit beta type-3 | Q9R1P1 |
| Proteasome subunit beta type-4 | P99026 |
| Proteasome subunit beta type-5 | O55234 |
| Proteasome subunit beta type-6 | Q60692 |
| Proteasome subunit beta type-7 | P70195 |
| Proteasome-associated protein ECM29 homolog | Q6PDI5 |
| Protein arginine N-methyltransferase 1 | Q9JIF0 |
| Protein arginine N-methyltransferase 5 | Q8CIG8 |
| Protein argonaute-2 | Q8CJG0 |
| Protein bassoon | O88737 |
| Protein BRICK1 | Q91VR8 |
| Protein C8orf37 homolog | Q3UJP5 |
| Protein canopy homolog 2 | Q9QXT0 |

| | |
|---|---|
| Protein CDV3 | Q4VAA2 |
| Protein CutA | Q9CQ89 |
| Protein deltex-3 | Q80V91 |
| Protein diaphanous homolog 2 | O70566 |
| Protein disulfide-isomerase | P09103 |
| Protein disulfide-isomerase A3 | P27773 |
| Protein disulfide-isomerase A4 | P08003 |
| Protein disulfide-isomerase A6 | Q922R8 |
| Protein DJ-1 | Q99LX0 |
| Protein dopey-2 | Q3UHQ6 |
| Protein enabled homolog | Q03173 |
| Protein ETHE1, mitochondrial | Q9DCM0 |
| Protein FAM134C | Q9CQV4 |
| Protein FAM136A | Q9CR98 |
| Protein FAM160B1 | Q8CDM8 |
| Protein FAM169A | Q5XG69 |
| Protein FAM171B | Q14CH0 |
| Protein FAM192A | Q91WE2 |
| Protein FAM40A | Q8C079 |
| Protein FAM49A | Q8BHZ0 |
| Protein FAM49B | Q921M7 |
| Protein FAM98B | Q80VD1 |
| Protein farnesyltransferase/geranylgeranyltransfe | Q61239 |
| Protein fat-free homolog | Q3UVL4 |
| Protein flightless-1 homolog | Q9JJ28 |
| Protein Hook homolog 3 | Q8BUK6 |
| Protein IMPACT | O55091 |
| Protein KIAA0284 | Q80U49 |
| Protein KIAA0664 | Q5SW19 |
| Protein KIAA1045 | Q80TL4 |
| Protein kinase C and casein kinase substrate in r | Q61644 |
| Protein kinase C and casein kinase substrate in r | Q9WVE8 |
| Protein kinase C beta type | P68404 |
| Protein kinase C epsilon type | P16054 |
| Protein kinase C gamma type | P63318 |
| Protein kinase C-binding protein NELL2 | Q61220 |
| Protein lin-7 homolog A | Q8JZS0 |
| Protein lin-7 homolog C | O88952 |
| Protein lunapark | Q7TQ95 |
| Protein LZIC | Q8K3C3 |
| Protein NDRG2 | Q9QYG0 |
| Protein NDRG3 | Q9QYF9 |
| Protein NDRG4 | Q8BTG7 |
| Protein NipSnap homolog 1 | O55125 |
| Protein NipSnap homolog 3B | Q9CQE1 |
| Protein phosphatase 1 regulatory inhibitor subuni | Q8VHQ3 |
| Protein phosphatase 1 regulatory subunit 12A | Q9DBR7 |
| Protein phosphatase 1 regulatory subunit 1A | Q9ERT9 |
| Protein phosphatase 1 regulatory subunit 21 | Q3TDD9 |
| Protein phosphatase 1 regulatory subunit 29 | Q68FM6 |
| Protein phosphatase 1 regulatory subunit 7 | Q3UM45 |
| Protein phosphatase 1A | P49443 |
| Protein phosphatase 1B | P36993 |
| Protein phosphatase 1E | Q80TL0 |
| Protein phosphatase 1F | Q8CGA0 |
| Protein phosphatase 1G | Q61074 |
| Protein phosphatase 1H | Q3UYC0 |
| Protein phosphatase methylesterase 1 | Q8BVQ5 |
| Protein piccolo | Q9QYX7 |
| Protein PRRC2C | Q3TLH4 |
| Protein prune homolog | Q8BIW1 |
| Protein rogdi homolog | Q3TDK6 |
| Protein RUFY3 | Q9D394 |
| Protein S100-A13 | P97352 |
| Protein SCAI | Q8C8N2 |
| Protein SET | Q9EQU5 |
| Protein shisa-6 homolog | Q3UH99 |
| Protein shisa-7 | Q8C3Q5 |
| Protein Shroom2 | A2ALU4 |
| Protein spire homolog 1 | Q52KF3 |
| Protein TANC1 | Q0VGY8 |
| Protein TANC2 | A2A690 |
| Protein transport protein Sec23A | Q01405 |
| Protein transport protein Sec31A | Q3UPL0 |
| Protein TSSC1 | Q8K0G5 |
| Protein unc-13 homolog A | Q4KUS2 |
| Protein-arginine deiminase type-2 | Q08642 |
| Protein-L-isoaspartate(D-aspartate) O-methyltran | P23506 |
| Protein-methionine sulfoxide oxidase MICAL3 | Q8CJ19 |
| Protein-tyrosine kinase 2-beta | Q9QVP9 |
| Prothymosin alpha | P26350 |
| Pumilio homolog 1 | Q80U78 |
| Purine nucleoside phosphorylase | P23492 |
| Purkinje cell protein 4 | P63054 |
| Puromycin-sensitive aminopeptidase | Q11011 |
| Putative adenosylhomocysteinase 2 | Q80SW1 |
| Putative adenosylhomocysteinase 3 | Q68FL4 |
| Putative ATP-dependent Clp protease proteolytic | O88696 |
| Putative N-acetylglucosamine-6-phosphate deac | Q8JZV7 |
| Putative phospholipase B-like 2 | Q3TCN2 |

| | |
|---|---|
| Putative pre-mRNA-splicing factor ATP-depender | O35286 |
| Putative RNA-binding protein Luc7-like 1 | Q9CYI4 |
| Putative tyrosine-protein phosphatase auxilin | Q80TZ3 |
| Pyridoxal kinase | Q8K183 |
| Pyridoxal phosphate phosphatase | P60487 |
| Pyridoxal-dependent decarboxylase domain-cont | Q99K01 |
| Pyridoxine-5'-phosphate oxidase | Q91XF0 |
| Pyruvate carboxylase, mitochondrial | Q05920 |
| Pyruvate dehydrogenase E1 component subunit | P35486 |
| Pyruvate dehydrogenase E1 component subunit | Q9D051 |
| Pyruvate dehydrogenase phosphatase regulatory | Q7TSQ8 |
| Pyruvate dehydrogenase protein X component, n | Q8BKZ9 |
| Pyruvate kinase isozymes M1/M2 | P52480 |
| Quinone oxidoreductase-like protein 1 | Q921W4 |
| R3H domain-containing protein 2 | Q80TM6 |
| Rab GDP dissociation inhibitor alpha | P50396 |
| Rab GDP dissociation inhibitor beta | Q61598 |
| Rab GTPase-activating protein 1 | A2AWA9 |
| Rab GTPase-activating protein 1-like | A6H6A9 |
| Rab GTPase-binding effector protein 1 | O35551 |
| Rab3 GTPase-activating protein catalytic subunit | Q80UJ7 |
| Rab3 GTPase-activating protein non-catalytic sub | Q8BMG7 |
| Rab5 GDP/GTP exchange factor | Q9JM13 |
| Rabenosyn-5 | Q80Y56 |
| Rab-like protein 1 | Q5U3K5 |
| Rabphilin-3A | P47708 |
| RAC-alpha serine/threonine-protein kinase | P31750 |
| RAC-beta serine/threonine-protein kinase | Q60823 |
| RAC-gamma serine/threonine-protein kinase | Q9WUA6 |
| Radixin | P26043 |
| Ral GTPase-activating protein subunit alpha-1 | Q6GYP7 |
| RalA-binding protein 1 | Q62172 |
| RalBP1-associated Eps domain-containing protei | O54916 |
| Ran GTPase-activating protein 1 | P46061 |
| Ran-binding protein 3 | Q9CT10 |
| Ran-binding protein 9 | P69566 |
| Ran-specific GTPase-activating protein | P34022 |
| Rap guanine nucleotide exchange factor 2 | Q8CHG7 |
| Rap guanine nucleotide exchange factor 4 | Q9EQZ6 |
| Rap1 GTPase-activating protein 2 | Q5SVL6 |
| Ras and Rab interactor 1 | Q921Q7 |
| Ras GTPase-activating protein SynGAP | F6SEU4 |
| Ras GTPase-activating protein-binding protein 1 | P97855 |
| Ras GTPase-activating protein-binding protein 2 | P97379 |
| Ras GTPase-activating-like protein IQGAP1 | Q9JKF1 |
| Ras GTPase-activating-like protein IQGAP2 | Q3UQ44 |
| Ras suppressor protein 1 | Q01730 |
| RasGAP-activating-like protein 1 | Q9Z268 |
| Ras-related C3 botulinum toxin substrate 1 | P63001 |
| Ras-related C3 botulinum toxin substrate 3 | P60764 |
| Ras-related protein Rab-10 | P61027 |
| Ras-related protein Rab-11B | P46638 |
| Ras-related protein Rab-12 | P35283 |
| Ras-related protein Rab-14 | Q91V41 |
| Ras-related protein Rab-15 | Q8K386 |
| Ras-related protein Rab-18 | P35293 |
| Ras-related protein Rab-1A | P62821 |
| Ras-related protein Rab-1B | Q9D1G1 |
| Ras-related protein Rab-21 | P35282 |
| Ras-related protein Rab-2A | P53994 |
| Ras-related protein Rab-35 | Q6PHN9 |
| Ras-related protein Rab-3A | P63011 |
| Ras-related protein Rab-3B | Q9CZT8 |
| Ras-related protein Rab-3C | P62823 |
| Ras-related protein Rab-4B | Q91ZR1 |
| Ras-related protein Rab-5A | Q9CQD1 |
| Ras-related protein Rab-5B | P61021 |
| Ras-related protein Rab-5C | P35278 |
| Ras-related protein Rab-6A | P35279 |
| Ras-related protein Rab-6B | P61294 |
| Ras-related protein Rab-7a | P51150 |
| Ras-related protein Rab-8A | P55258 |
| Ras-related protein Ral-A | P63321 |
| Ras-related protein Rap-1A | P62835 |
| Ras-related protein Rap-2a | Q80ZJ1 |
| Ras-related protein Rap-2b | P61226 |
| Ras-related protein R-Ras2 | P62071 |
| Ras-specific guanine nucleotide-releasing factor | P27671 |
| Receptor expression-enhancing protein 2 | Q8VCD6 |
| Receptor-type tyrosine-protein phosphatase alph | P18052 |
| Receptor-type tyrosine-protein phosphatase delta | Q64487 |
| Receptor-type tyrosine-protein phosphatase eta | Q64455 |
| Receptor-type tyrosine-protein phosphatase N2 | P80560 |
| Receptor-type tyrosine-protein phosphatase S | B0V2N1 |
| Redox-regulatory protein FAM213A | Q9CYH2 |
| Regulating synaptic membrane exocytosis protein | Q99NE5 |
| Regulator of G-protein signaling 6 | Q9Z2H2 |
| Regulator of G-protein signaling 7 | O54829 |
| Regulator of nonsense transcripts 1 | Q9EPU0 |
| Regulatory-associated protein of mTOR | Q8K4Q0 |

| | |
|---|---|
| Reticulocalbin-2 | Q8BP92 |
| Reticulon-1 | Q8K0T0 |
| Reticulon-3 | Q9ES97 |
| Reticulon-4 | Q99P72 |
| Retinal dehydrogenase 1 | P24549 |
| Retinaldehyde-binding protein 1 | Q9Z275 |
| Retinol dehydrogenase 14 | Q9ERI6 |
| Retinol-binding protein 1 | Q00915 |
| Rho GDP-dissociation inhibitor 1 | Q99PT1 |
| Rho GDP-dissociation inhibitor 2 | Q61599 |
| Rho GTPase-activating protein 1 | Q5FWK3 |
| Rho GTPase-activating protein 21 | Q6DFV3 |
| Rho GTPase-activating protein 23 | Q69ZH9 |
| Rho GTPase-activating protein 32 | Q811P8 |
| Rho GTPase-activating protein 35 | Q91YM2 |
| Rho GTPase-activating protein 39 | P59281 |
| Rho guanine nucleotide exchange factor 12 | Q8R4H2 |
| Rho guanine nucleotide exchange factor 2 | Q60875 |
| Rho guanine nucleotide exchange factor 7 | Q9ES28 |
| Rho-associated protein kinase 2 | P70336 |
| Rho-related GTP-binding protein RhoB | P62746 |
| Ribonuclease inhibitor | Q91VI7 |
| Ribonuclease UK114 | P52760 |
| Ribosomal protein S6 kinase alpha-3 | P18654 |
| Ribosomal protein S6 kinase alpha-4 | Q9Z2B9 |
| Ribosome maturation protein SBDS | P70122 |
| Ribosome-binding protein 1 | Q99PL5 |
| Ribosome-recycling factor, mitochondrial | Q9D6S7 |
| RILP-like protein 1 | Q9JJC6 |
| RIMS-binding protein 2 | Q80U40 |
| RING finger protein 214 | Q8BFU3 |
| RNA polymerase II-associated protein 3 | Q9D706 |
| RNA-binding protein 39 | Q8VH51 |
| RNA-binding protein EWS | Q61545 |
| RNA-binding protein FUS | P56959 |
| RUN and FYVE domain-containing protein 2 | Q8R4C2 |
| RUN domain-containing protein 3A | O08576 |
| RuvB-like 1 | P60122 |
| RuvB-like 2 | Q9WTM5 |
| RWD domain-containing protein 1 | Q9CQK7 |
| Ryanodine receptor 2 | E9Q401 |
| Saccharopine dehydrogenase-like oxidoreductase | Q8R127 |
| S-adenosylmethionine synthase isoform type-2 | Q3THS6 |
| Sarcoplasmic/endoplasmic reticulum calcium ATF | O55143 |
| Scavenger mRNA-decapping enzyme DcpS | Q9DAR7 |
| SEC14 domain and spectrin repeat-containing pr | Q80UK0 |
| SEC14-like protein 2 | Q99J08 |
| SEC23-interacting protein | Q6NZC7 |
| Secernin-1 | Q9CZC8 |
| Secretogranin-1 | P16014 |
| Secretogranin-2 | Q03517 |
| Secretogranin-3 | P47867 |
| Secretory carrier-associated membrane protein 1 | Q8K021 |
| Secretory carrier-associated membrane protein 3 | O35609 |
| Seizure 6-like protein | Q6P1D5 |
| Seizure 6-like protein 2 | Q4V9Z5 |
| Selenocysteine lyase | Q9JLI6 |
| Sepiapterin reductase | Q64105 |
| Septin-11 | Q8C1B7 |
| Septin-2 | P42208 |
| Septin-5 | Q9Z2Q6 |
| Septin-6 | Q9R1T4 |
| Septin-7 | O55131 |
| Septin-8 | Q8CHH9 |
| Septin-9 | Q80UG5 |
| Serine racemase | Q9QZX7 |
| Serine/arginine-rich splicing factor 2 | Q62093 |
| Serine/threonine-protein kinase BRSK1 | Q5RJI5 |
| Serine/threonine-protein kinase BRSK2 | Q69Z98 |
| Serine/threonine-protein kinase DCLK1 | Q9JLM8 |
| Serine/threonine-protein kinase DCLK2 | Q6PGN3 |
| Serine/threonine-protein kinase MARK1 | Q8VHJ5 |
| Serine/threonine-protein kinase MRCK alpha | Q3UU96 |
| Serine/threonine-protein kinase MRCK beta | Q7TT50 |
| Serine/threonine-protein kinase mTOR | Q9JLN9 |
| Serine/threonine-protein kinase N1 | P70268 |
| Serine/threonine-protein kinase OSR1 | Q6P9R2 |
| Serine/threonine-protein kinase PAK 1 | O88643 |
| Serine/threonine-protein kinase PAK 2 | Q8CIN4 |
| Serine/threonine-protein kinase PAK 3 | Q61036 |
| Serine/threonine-protein kinase SIK3 | Q6P4S6 |
| Serine/threonine-protein kinase SMG1 | Q8BKX6 |
| Serine/threonine-protein kinase STK11 | Q9WTK7 |
| Serine/threonine-protein kinase TAO1 | Q5F2E8 |
| Serine/threonine-protein kinase WNK1 | P83741 |
| Serine/threonine-protein kinase WNK2 | Q3UH66 |
| Serine/threonine-protein phosphatase 2A 55 kDa | Q6P1F6 |
| Serine/threonine-protein phosphatase 2A 56 kDa | Q6PD03 |
| Serine/threonine-protein phosphatase 2A 56 kDa | Q61151 |
| Serine/threonine-protein phosphatase 2A 56 kDa | Q60996 |

| | |
|---|---|
| Serine/threonine-protein phosphatase 2A 65 kDa | Q76MZ3 |
| Serine/threonine-protein phosphatase 2A activato | P58389 |
| Serine/threonine-protein phosphatase 2A catalyti | P63330 |
| Serine/threonine-protein phosphatase 2B catalyti | P63328 |
| Serine/threonine-protein phosphatase 2B catalyti | P48453 |
| Serine/threonine-protein phosphatase 4 catalytic | P97470 |
| Serine/threonine-protein phosphatase 5 | Q60676 |
| Serine/threonine-protein phosphatase PGAM5, m | Q8BX10 |
| Serine/threonine-protein phosphatase PP1-alpha | P62137 |
| Serine/threonine-protein phosphatase PP1-beta c | P62141 |
| Serine/threonine-protein phosphatase PP1-gamm | P63087 |
| Serine-threonine kinase receptor-associated prot | Q9Z1Z2 |
| Serine--tRNA ligase, cytoplasmic | P26638 |
| Serotransferrin | Q921I1 |
| Serpin B6 | Q60854 |
| Serpin H1 | P19324 |
| Serrate RNA effector molecule homolog | Q99MR6 |
| Serum albumin | P07724 |
| S-formylglutathione hydrolase | Q9R0P3 |
| SH2B adapter protein 1 | Q91ZM2 |
| SH3 and multiple ankyrin repeat domains protein | D3YZU1 |
| SH3 and multiple ankyrin repeat domains protein | Q80Z38 |
| SH3 and multiple ankyrin repeat domains protein | Q4ACU6 |
| SH3 domain-binding glutamic acid-rich-like protei | Q9JJU8 |
| SH3 domain-binding glutamic acid-rich-like protei | Q91VW3 |
| SH3 domain-containing kinase-binding protein 1 | Q8R550 |
| SH3-containing GRB2-like protein 3-interacting pr | Q8VD37 |
| Short coiled-coil protein | Q78YZ6 |
| Sickle tail protein | A2AQ25 |
| Sideroflexin-1 | Q99JR1 |
| Sideroflexin-3 | Q91V61 |
| Sideroflexin-5 | Q925N0 |
| Signal recognition particle 54 kDa protein | P14576 |
| Signal recognition particle 68 kDa protein | Q8BMA6 |
| Signal recognition particle receptor subunit beta | P47758 |
| Signal transducing adapter molecule 1 | P70297 |
| Signal-induced proliferation-associated 1-like prot | Q8C0T5 |
| Single-stranded DNA-binding protein, mitochondr | Q9CYR0 |
| SLIT-ROBO Rho GTPase-activating protein 2 | Q91Z67 |
| SLIT-ROBO Rho GTPase-activating protein 3 | Q812A2 |
| Small G protein signaling modulator 1 | Q8BPQ7 |
| Small glutamine-rich tetratricopeptide repeat-cont | Q8BJU0 |
| Small ubiquitin-related modifier 2 | P61957 |
| Sodium- and chloride-dependent GABA transport | P31650 |
| Sodium bicarbonate cotransporter 3 | Q8BTY2 |
| Sodium/calcium exchanger 1 | P70414 |
| Sodium/potassium-transporting ATPase subunit a | Q8VDN2 |
| Sodium/potassium-transporting ATPase subunit a | Q6PIE5 |
| Sodium/potassium-transporting ATPase subunit a | Q6PIC6 |
| Sodium/potassium-transporting ATPase subunit b | P14094 |
| Sodium/potassium-transporting ATPase subunit b | P14231 |
| Sodium-dependent neutral amino acid transporte | Q8BJI1 |
| Sodium-driven chloride bicarbonate exchanger | Q5DTL9 |
| Solute carrier family 12 member 2 | P55012 |
| Solute carrier family 12 member 5 | Q91V14 |
| Solute carrier family 12 member 9 | Q99MR3 |
| Solute carrier family 2, facilitated glucose transpo | P32037 |
| Sorbin and SH3 domain-containing protein 1 | Q62417 |
| Sorbin and SH3 domain-containing protein 2 | Q3UTJ2 |
| Sorbitol dehydrogenase | Q64442 |
| Sorcin | Q6P069 |
| Sortilin | Q6PHU5 |
| Sortilin-related receptor | O88307 |
| Sorting nexin-1 | Q9WV80 |
| Sorting nexin-12 | O70493 |
| Sorting nexin-2 | Q9CWK8 |
| Sorting nexin-27 | Q3UHD6 |
| Sorting nexin-3 | O70492 |
| Sorting nexin-30 | Q8CE50 |
| Sorting nexin-32 | Q80ZJ7 |
| Sorting nexin-4 | Q91YJ2 |
| Sorting nexin-5 | Q9D8U8 |
| Sorting nexin-6 | Q6P8X1 |
| SPARC-like protein 1 | P70663 |
| Spectrin alpha chain, brain | P16546 |
| Spectrin beta chain, brain 1 | Q62261 |
| Spectrin beta chain, erythrocyte | P15508 |
| Spermidine synthase | Q64674 |
| Spermine synthase | P97355 |
| S-phase kinase-associated protein 1 | Q9WTX5 |
| Spliceosome RNA helicase Ddx39b | Q9Z1N5 |
| Splicing factor 3B subunit 1 | Q99NB9 |
| Splicing factor 3B subunit 3 | Q921M3 |
| Splicing factor U2AF 65 kDa subunit | P26369 |
| Splicing factor, proline- and glutamine-rich | Q8VIJ6 |
| Spondin-1 | Q8VCC9 |
| Squamous cell carcinoma antigen recognized by | Q9JLI8 |
| SRC kinase signaling inhibitor 1 | Q9QWI6 |
| Src substrate cortactin | Q60598 |
| SRSF protein kinase 2 | O54781 |

| | |
|---|---|
| Staphylococcal nuclease domain-containing prote | Q78PY7 |
| Stathmin | P54227 |
| Stathmin-2 | P55821 |
| Stathmin-3 | O70166 |
| STE20/SPS1-related proline-alanine-rich protein | Q9Z1W9 |
| STE20-like serine/threonine-protein kinase | O54988 |
| Sterol-4-alpha-carboxylate 3-dehydrogenase, dec | Q9R1J0 |
| STIP1 homology and U box-containing protein 1 | Q9WUD1 |
| Stomatin-like protein 2 | Q99JB2 |
| Stress-70 protein, mitochondrial | P38647 |
| Stress-induced-phosphoprotein 1 | Q60864 |
| Striatin | O55106 |
| Striatin-3 | Q9ERG2 |
| Striatin-4 | P58404 |
| Stromal membrane-associated protein 2 | Q7TN29 |
| Succinate dehydrogenase [ubiquinone] flavoprote | Q8K2B3 |
| Succinate dehydrogenase [ubiquinone] iron-sulfu | Q9CQA3 |
| Succinate-semialdehyde dehydrogenase, mitocho | Q8BWF0 |
| Succinyl-CoA ligase [ADP-forming] subunit beta, | Q9Z2I9 |
| Succinyl-CoA ligase [GDP-forming] subunit beta, | Q9Z2I8 |
| Succinyl-CoA:3-ketoacid-coenzyme A transferase | Q9D0K2 |
| Sulfated glycoprotein 1 | Q61207 |
| Sulfotransferase 4A1 | P63046 |
| SUMO-activating enzyme subunit 1 | Q9R1T2 |
| SUMO-activating enzyme subunit 2 | Q9Z1F9 |
| Superoxide dismutase [Cu-Zn] | P08228 |
| Superoxide dismutase [Mn], mitochondrial | P09671 |
| Suppressor of G2 allele of SKP1 homolog | Q9CX34 |
| SWI/SNF complex subunit SMARCC2 | Q6PDG5 |
| Switch-associated protein 70 | Q6A028 |
| Synapsin-1 | O88935 |
| Synapsin-2 | Q64332 |
| Synaptic vesicle glycoprotein 2A | Q9JIS5 |
| Synaptic vesicle glycoprotein 2B | Q8BG39 |
| Synaptic vesicle membrane protein VAT-1 homolo | Q62465 |
| Synaptic vesicle membrane protein VAT-1 homolo | Q80TB8 |
| Synaptobrevin homolog YKT6 | Q9CQW1 |
| Synaptogyrin-3 | Q8R191 |
| Synaptojanin-1 | Q8CHC4 |
| Synaptojanin-2-binding protein | Q9D6K5 |
| Synaptophysin | Q62277 |
| Synaptopodin | Q8CC35 |
| Synaptoporin | Q8BGN8 |
| Synaptosomal-associated protein 25 | P60879 |
| Synaptosomal-associated protein 47 | Q8R570 |
| Synaptotagmin-1 | P46096 |
| Synaptotagmin-12 | Q920N7 |
| Synaptotagmin-17 | Q920M7 |
| Synaptotagmin-2 | P46097 |
| Synaptotagmin-3 | O35681 |
| Synaptotagmin-5 | Q9R0N5 |
| Synaptotagmin-7 | Q9R0N7 |
| Synembryn-A | Q3TIR3 |
| Synergin gamma | Q5SV85 |
| Syntaphilin | Q80U23 |
| Syntaxin-12 | Q9ER00 |
| Syntaxin-16 | Q8BVI5 |
| Syntaxin-1A | O35526 |
| Syntaxin-1B | P61264 |
| Syntaxin-6 | Q9JKK1 |
| Syntaxin-7 | O70439 |
| Syntaxin-8 | O88983 |
| Syntaxin-binding protein 1 | O08599 |
| Syntaxin-binding protein 5 | Q8K400 |
| Syntaxin-binding protein 5-like | Q5DQR4 |
| Talin-1 | P26039 |
| Talin-2 | Q71LX4 |
| TAR DNA-binding protein 43 | Q921F2 |
| Target of Myb protein 1 | O88746 |
| Tax1-binding protein 1 homolog | Q3UKC1 |
| TBC1 domain family member 10B | Q8BHL3 |
| TBC1 domain family member 17 | Q8BYH7 |
| TBC1 domain family member 24 | Q3UUG6 |
| TBC1 domain family member 8B | A3KGB4 |
| TBC1 domain family member 9B | Q5SVR0 |
| T-cell immunomodulatory protein | Q99KW9 |
| T-complex protein 1 subunit alpha | P11983 |
| T-complex protein 1 subunit beta | P80314 |
| T-complex protein 1 subunit delta | P80315 |
| T-complex protein 1 subunit epsilon | P80316 |
| T-complex protein 1 subunit eta | P80313 |
| T-complex protein 1 subunit gamma | P80318 |
| T-complex protein 1 subunit theta | P42932 |
| T-complex protein 1 subunit zeta | P80317 |
| T-complex protein 11-like protein 1 | Q8BTG3 |
| Tectonin beta-propeller repeat-containing protein | Q80VP0 |
| Tenascin | Q80YX1 |
| Tenascin-R | Q8BYI9 |
| Teneurin-3 | Q9WTS6 |
| Teneurin-4 | Q3UHK6 |

| | |
|---|---|
| Tether containing UBX domain for GLUT4 | Q8VBT9 |
| Tetratricopeptide repeat protein 1 | Q91Z38 |
| Tetratricopeptide repeat protein 19, mitochondria | Q8CC21 |
| Tetratricopeptide repeat protein 35 | Q9CRD2 |
| Tetratricopeptide repeat protein 38 | A3KMP2 |
| Tetratricopeptide repeat protein 9A | Q3V038 |
| Thiamine-triphosphatase | Q8JZL3 |
| Thimet oligopeptidase | Q8C1A5 |
| Thiomorpholine-carboxylate dehydrogenase | O54983 |
| Thioredoxin | P10639 |
| Thioredoxin domain-containing protein 12 | Q9CQU0 |
| Thioredoxin domain-containing protein 5 | Q91W90 |
| Thioredoxin reductase 1, cytoplasmic | Q9JMH6 |
| Thioredoxin-dependent peroxide reductase, mitoc | P20108 |
| Thioredoxin-like protein 1 | Q8CDN6 |
| Thiosulfate sulfurtransferase | P52196 |
| Threonine--tRNA ligase, cytoplasmic | Q9D0R2 |
| Threonine--tRNA ligase, mitochondrial | Q3UQ84 |
| THUMP domain-containing protein 1 | Q99J36 |
| Thy-1 membrane glycoprotein | P01831 |
| Thymidylate kinase | P97930 |
| Thymosin beta-10 | Q6ZWY8 |
| Thymosin beta-4 | P20065 |
| Tight junction protein ZO-2 | Q9Z0U1 |
| TIP41-like protein | Q8BH58 |
| Titin | A2ASS6 |
| T-lymphoma invasion and metastasis-inducing pr | Q60610 |
| Toll-interacting protein | Q9QZ06 |
| TOM1-like protein 2 | Q5SRX1 |
| TPR and ankyrin repeat-containing protein 1 | Q8BV79 |
| Traf2 and NCK-interacting protein kinase | P83510 |
| Trafficking protein particle complex subunit 12 | Q8K2L8 |
| Trafficking protein particle complex subunit 2-like | Q9JME7 |
| Trafficking protein particle complex subunit 3 | O55013 |
| Trafficking protein particle complex subunit 4 | Q9ES56 |
| Trafficking protein particle complex subunit 5 | Q9CQA1 |
| Trafficking protein particle complex subunit 9 | Q3U0M1 |
| Trans-2-enoyl-CoA reductase, mitochondrial | Q9DCS3 |
| Transaldolase | Q93092 |
| Transcription elongation factor A protein-like 3 | Q8R0A5 |
| Transcription elongation factor A protein-like 5 | Q8CCT4 |
| Transcription elongation factor B polypeptide 1 | P83940 |
| Transcription elongation factor B polypeptide 2 | P62869 |
| Transcription elongation factor SPT5 | O55201 |
| Transcription elongation regulator 1 | Q8CGF7 |
| Transcription factor A, mitochondrial | P40630 |
| Transcription factor BTF3 homolog 4 | Q9CQH7 |
| Transcription intermediary factor 1-beta | Q62318 |
| Transcriptional activator protein Pur-alpha | P42669 |
| Transcriptional activator protein Pur-beta | O35295 |
| Transferrin receptor protein 1 | Q62351 |
| Transforming acidic coiled-coil-containing protein | Q6Y685 |
| Transforming acidic coiled-coil-containing protein | Q9JJG0 |
| Transforming protein RhoA | Q9QUI0 |
| Transgelin-2 | Q9WVA4 |
| Transgelin-3 | Q9R1Q8 |
| Transitional endoplasmic reticulum ATPase | Q01853 |
| Transketolase | P40142 |
| Translation initiation factor eIF-2B subunit alpha | Q99LC8 |
| Translation initiation factor eIF-2B subunit epsilor | Q8CHW4 |
| Translational activator of cytochrome c oxidase 1 | Q8K0Z7 |
| Translationally-controlled tumor protein | P63028 |
| Transmembrane 9 superfamily member 4 | Q8BH24 |
| Transmembrane emp24 domain-containing prote | Q9D1D4 |
| Transmembrane emp24 domain-containing prote | Q99KF1 |
| Transportin-1 | Q8BFY9 |
| Transportin-3 | Q6P2B1 |
| Transthyretin | P07309 |
| Trifunctional enzyme subunit alpha, mitochondria | Q8BMS1 |
| Trifunctional enzyme subunit beta, mitochondrial | Q99JY0 |
| Trifunctional purine biosynthetic protein adenosin | Q64737 |
| Triosephosphate isomerase | P17751 |
| Tripartite motif-containing protein 2 | Q9ESN6 |
| Tripartite motif-containing protein 3 | Q9R1R2 |
| Tripartite motif-containing protein 46 | Q7TNM2 |
| Tripeptidyl-peptidase 2 | Q64514 |
| Triple functional domain protein | Q0KL02 |
| tRNA-splicing ligase RtcB homolog | Q99LF4 |
| Tropomodulin-2 | Q9JKK7 |
| Tropomyosin alpha-1 chain | P58771 |
| Tropomyosin alpha-3 chain | P21107 |
| Tropomyosin alpha-4 chain | Q6IRU2 |
| Tryptophan--tRNA ligase, cytoplasmic | P32921 |
| TSC22 domain family protein 1 | P62500 |
| Tuberin | Q61037 |
| Tubulin alpha chain-like 3 | Q3UX10 |
| Tubulin alpha-4A chain | P68368 |
| Tubulin alpha-8 chain | Q9JJZ2 |
| Tubulin beta-2A chain | Q7TMM9 |
| Tubulin beta-2B chain | Q9CWF2 |

| | |
|---|---|
| Tubulin beta-3 chain | Q9ERD7 |
| Tubulin beta-4A chain | Q9D6F9 |
| Tubulin beta-4B chain | P68372 |
| Tubulin beta-5 chain | P99024 |
| Tubulin polymerization-promoting protein | Q7TQD2 |
| Tubulin-folding cofactor B | Q9D1E6 |
| Tubulin-specific chaperone A | P48428 |
| Tubulin-specific chaperone C | Q8VCN9 |
| Tubulin-specific chaperone D | Q8BYA0 |
| Tubulin-specific chaperone E | Q8CIV8 |
| Tubulin--tyrosine ligase-like protein 12 | Q3UDE2 |
| Tumor necrosis factor alpha-induced protein 8-lik | Q3TBL6 |
| Tumor protein D52 | Q62393 |
| Tumor protein D54 | Q9CYZ2 |
| Tumor susceptibility gene 101 protein | Q61187 |
| Twinfilin-1 | Q91YR1 |
| Twinfilin-2 | Q9Z0P5 |
| Type I inositol 3,4-bisphosphate 4-phosphatase | Q9EPW0 |
| Tyrosine-protein kinase CSK | P41241 |
| Tyrosine-protein kinase Fyn | P39688 |
| Tyrosine-protein phosphatase non-receptor type | P35235 |
| Tyrosine-protein phosphatase non-receptor type | Q6PB44 |
| Tyrosine-protein phosphatase non-receptor type | P97797 |
| Tyrosine--tRNA ligase, cytoplasmic | Q91WQ3 |
| Tyrosine--tRNA ligase, mitochondrial | Q8BYL4 |
| U8 snoRNA-decapping enzyme | Q6P3D0 |
| Ubiquilin-1 | Q8R317 |
| Ubiquilin-2 | Q9QZM0 |
| Ubiquilin-4 | Q99NB8 |
| Ubiquinone biosynthesis protein COQ9, mitochor | Q8K1Z0 |
| Ubiquitin carboxyl-terminal hydrolase 10 | P52479 |
| Ubiquitin carboxyl-terminal hydrolase 14 | Q9JMA1 |
| Ubiquitin carboxyl-terminal hydrolase 15 | Q8R5H1 |
| Ubiquitin carboxyl-terminal hydrolase 24 | B1AY13 |
| Ubiquitin carboxyl-terminal hydrolase 4 | P35123 |
| Ubiquitin carboxyl-terminal hydrolase 47 | Q8BY87 |
| Ubiquitin carboxyl-terminal hydrolase 5 | P56399 |
| Ubiquitin carboxyl-terminal hydrolase 7 | Q6A4J8 |
| Ubiquitin carboxyl-terminal hydrolase CYLD | Q80TQ2 |
| Ubiquitin carboxyl-terminal hydrolase isozyme L1 | Q9R0P9 |
| Ubiquitin carboxyl-terminal hydrolase isozyme L3 | Q9JKB1 |
| Ubiquitin carboxyl-terminal hydrolase isozyme L5 | Q9WUP7 |
| Ubiquitin conjugation factor E4 B | Q9ES00 |
| Ubiquitin fusion degradation protein 1 homolog | P70362 |
| Ubiquitin thioesterase OTUB1 | Q7TQI3 |
| Ubiquitin-associated protein 2-like | Q80X50 |
| Ubiquitin-conjugating enzyme E2 K | P61087 |
| Ubiquitin-conjugating enzyme E2 L3 | P68037 |
| Ubiquitin-conjugating enzyme E2 N | P61089 |
| Ubiquitin-conjugating enzyme E2 O | Q6ZPJ3 |
| Ubiquitin-like modifier-activating enzyme 1 | Q02053 |
| Ubiquitin-like modifier-activating enzyme 6 | Q8C7R4 |
| Ubiquitin-like modifier-activating enzyme ATG7 | Q9D906 |
| Ubiquitin-like protein 4A | P21126 |
| Ubiquitin-like-conjugating enzyme ATG3 | Q9CPX6 |
| Ubiquitin-protein ligase E3A | O08759 |
| Ubiquitin-protein ligase E3C | Q80U95 |
| UBX domain-containing protein 1 | Q922Y1 |
| UBX domain-containing protein 6 | Q99PL6 |
| UDP-glucose:glycoprotein glucosyltransferase 1 | Q6P5E4 |
| UDP-N-acetylglucosamine--peptide N-acetylgluco | Q8CGY8 |
| UDP-N-acetylhexosamine pyrophosphorylase | Q91YN5 |
| UHRF1-binding protein 1-like | A2RSJ4 |
| UMP-CMP kinase | Q9DBP5 |
| UMP-CMP kinase 2, mitochondrial | Q3U5Q7 |
| Uncharacterized protein C19orf52 homolog | Q8BGX2 |
| Uncharacterized protein C1orf198 homolog | Q8C3W1 |
| Uncharacterized protein C1orf96 homolog | Q8QZT2 |
| Uncharacterized protein KIAA0513 | Q8R0A7 |
| Uncharacterized protein KIAA0564 homolog | Q8CC88 |
| Uncharacterized protein KIAA0930 homolog | Q3UE31 |
| Uncharacterized protein KIAA1467 | Q8BYI8 |
| Unconventional myosin-Ic | Q9WTI7 |
| Unconventional myosin-Va | Q99104 |
| Unconventional myosin-VI | Q64331 |
| Unconventional myosin-XVIIIa | Q9JMH9 |
| UPF0160 protein MYG1, mitochondrial | Q9JK81 |
| UPF0364 protein C6orf211 homolog | A6H630 |
| UPF0366 protein C11orf67 homolog | Q8R0P4 |
| UPF0533 protein C5orf44 homolog | Q3TIR1 |
| UPF0556 protein C19orf10 homolog | Q9CPT4 |
| UPF0568 protein C14orf166 homolog | Q9CQE8 |
| UPF0587 protein C1orf123 homolog | Q8BHG2 |
| UPF0606 protein KIAA1549 | Q68FD9 |
| UPF0696 protein C11orf68 homolog | Q8VD62 |
| Uroporphyrinogen decarboxylase | P70697 |
| UTP--glucose-1-phosphate uridylyltransferase | Q91ZJ5 |
| UV excision repair protein RAD23 homolog A | P54726 |
| UV excision repair protein RAD23 homolog B | P54728 |
| Vacuolar protein sorting-associated protein 11 ho | Q91W86 |

| | |
|---|---|
| Vacuolar protein sorting-associated protein 13C | Q8BX70 |
| Vacuolar protein sorting-associated protein 18 ho | Q8R307 |
| Vacuolar protein sorting-associated protein 26A | P40336 |
| Vacuolar protein sorting-associated protein 26B | Q8C0E2 |
| Vacuolar protein sorting-associated protein 29 | Q9QZ88 |
| Vacuolar protein sorting-associated protein 33A | Q9D2N9 |
| Vacuolar protein sorting-associated protein 35 | Q9EQH3 |
| Vacuolar protein sorting-associated protein 45 | P97390 |
| Vacuolar protein sorting-associated protein 4A | Q8VEJ9 |
| Vacuolar protein sorting-associated protein 52 ho | Q8C754 |
| Vacuolar protein-sorting-associated protein 36 | Q91XD6 |
| Valacyclovir hydrolase | Q8R164 |
| Valine--tRNA ligase | Q9Z1Q9 |
| Vascular cell adhesion protein 1 | P29533 |
| Versican core protein | Q62059 |
| Very long-chain specific acyl-CoA dehydrogenase | P50544 |
| Vesicle transport through interaction with t-SNAR | O88384 |
| Vesicle-associated membrane protein 1 | Q62442 |
| Vesicle-associated membrane protein 2 | P63044 |
| Vesicle-associated membrane protein-associated | Q9WV55 |
| Vesicle-associated membrane protein-associated | Q9QY76 |
| Vesicle-fusing ATPase | P46460 |
| Vesicle-trafficking protein SEC22b | O08547 |
| Vesicular glutamate transporter 1 | Q3TXX4 |
| Vesicular inhibitory amino acid transporter | O35633 |
| Vigilin | Q8VDJ3 |
| Vimentin | P20152 |
| Vinculin | Q64727 |
| VIP36-like protein | P59481 |
| Visinin-like protein 1 | P62761 |
| Voltage-dependent anion-selective channel prote | Q60932 |
| Voltage-dependent anion-selective channel prote | Q60930 |
| Voltage-dependent anion-selective channel prote | Q60931 |
| Voltage-dependent calcium channel gamma-8 su | Q8VHW2 |
| Voltage-dependent calcium channel subunit alpha | O08532 |
| Voltage-dependent calcium channel subunit alpha | Q6PHS9 |
| Voltage-dependent calcium channel subunit alpha | Q9Z1L5 |
| Voltage-dependent L-type calcium channel subur | Q8R3Z5 |
| Voltage-dependent L-type calcium channel subur | P54285 |
| Voltage-dependent N-type calcium channel subu | O55017 |
| Voltage-dependent P/Q-type calcium channel sul | P97445 |
| Voltage-dependent R-type calcium channel subu | Q61290 |
| Voltage-gated potassium channel subunit beta-2 | P62482 |
| von Willebrand factor A domain-containing protei | Q99KC8 |
| VPS10 domain-containing receptor SorCS2 | Q9EPR5 |
| V-type proton ATPase 116 kDa subunit a isoform | Q9Z1G4 |
| V-type proton ATPase catalytic subunit A | P50516 |
| V-type proton ATPase subunit B, brain isoform | P62814 |
| V-type proton ATPase subunit C 1 | Q9Z1G3 |
| V-type proton ATPase subunit D | P57746 |
| V-type proton ATPase subunit d 1 | P51863 |
| V-type proton ATPase subunit E 1 | P50518 |
| V-type proton ATPase subunit F | Q9D1K2 |
| V-type proton ATPase subunit G 1 | Q9CR51 |
| V-type proton ATPase subunit G 2 | Q9WTT4 |
| V-type proton ATPase subunit H | Q8BVE3 |
| WAS/WASL-interacting protein family member 3 | P0C7L0 |
| WASH complex subunit FAM21 | Q6PGL7 |
| WASH complex subunit strumpellin | Q8C2E7 |
| WD repeat domain phosphoinositide-interacting p | Q80W47 |
| WD repeat-containing protein 1 | O88342 |
| WD repeat-containing protein 13 | Q91V09 |
| WD repeat-containing protein 37 | Q8CBE3 |
| WD repeat-containing protein 44 | Q6NVE8 |
| WD repeat-containing protein 47 | Q8CGF6 |
| WD repeat-containing protein 54 | Q9R0D8 |
| WD repeat-containing protein 7 | Q920I9 |
| WD repeat-containing protein mio | Q8VE19 |
| Wiskott-Aldrich syndrome protein family member | Q8R5H6 |
| Wiskott-Aldrich syndrome protein family member | Q8VHI6 |
| Wolframin | P56695 |
| WW domain-binding protein 2 | P97765 |
| Xaa-Pro aminopeptidase 1 | Q6P1B1 |
| Xaa-Pro dipeptidase | Q11136 |
| Zinc finger protein ZPR1 | Q62384 |
| Zinc finger RNA-binding protein | O88532 |
| Zinc phosphodiesterase ELAC protein 2 | Q80Y81 |
| Zinc-binding alcohol dehydrogenase domain-cont | Q8BGC4 |
| ZW10 interactor | Q9CQU5 |

lostripain

### Table S4. Proteins detected by iTRAQ in adult

| Detected in all experiments - 1123 proteins | |
|---|---|
| **Protein Name** | **Uniprot ID** |
| 14-3-3 protein beta/alpha | Q9CQV8 |
| 14-3-3 protein epsilon | P62259 |
| 14-3-3 protein eta | P68510 |
| 14-3-3 protein gamma | P61982 |
| 14-3-3 protein theta | P68254 |
| 14-3-3 protein zeta/delta | P63101 |
| 182 kDa tankyrase-1-binding protein | P58871 |
| 1-phosphatidylinositol 4,5-bisphosphate phospho | Q9Z1B3 |
| 1-phosphatidylinositol 4,5-bisphosphate phospho | A2AP18 |
| 2',3'-cyclic-nucleotide 3'-phosphodiesterase | P16330 |
| 26S protease regulatory subunit 4 | P62192 |
| 26S protease regulatory subunit 6A | O88685 |
| 26S protease regulatory subunit 7 | P46471 |
| 26S proteasome non-ATPase regulatory subunit | Q3TXS7 |
| 26S proteasome non-ATPase regulatory subunit | Q8BG32 |
| 26S proteasome non-ATPase regulatory subunit | Q8VDM4 |
| 26S proteasome non-ATPase regulatory subunit | P14685 |
| 26S proteasome non-ATPase regulatory subunit | Q9CX56 |
| 28 kDa heat- and acid-stable phosphoprotein | Q3UHX2 |
| 2-oxoglutarate dehydrogenase, mitochondrial | Q60597 |
| 3-hydroxyisobutyrate dehydrogenase, mitochond | Q99L13 |
| 40S ribosomal protein S13 | P62301 |
| 40S ribosomal protein S14 | P62264 |
| 40S ribosomal protein S16 | P14131 |
| 40S ribosomal protein S18 | P62270 |
| 40S ribosomal protein S28 | P62858 |
| 40S ribosomal protein S3 | P62908 |
| 40S ribosomal protein S3a | P97351 |
| 40S ribosomal protein S4, X isoform | P62702 |
| 40S ribosomal protein S6 | P62754 |
| 40S ribosomal protein S7 | P62082 |
| 40S ribosomal protein S8 | P62242 |
| 40S ribosomal protein S9 | Q6ZWN5 |
| 40S ribosomal protein SA | P14206 |
| 4-aminobutyrate aminotransferase, mitochondria | P61922 |
| 4F2 cell-surface antigen heavy chain | P10852 |
| 4-trimethylaminobutyraldehyde dehydrogenase | Q9JLJ2 |
| 5'-nucleotidase domain-containing protein 3 | Q3UHB1 |
| 60 kDa heat shock protein, mitochondrial | P63038 |
| 60 kDa SS-A/Ro ribonucleoprotein | O08848 |
| 60S acidic ribosomal protein P2 | P99027 |
| 60S ribosomal protein L13 | P47963 |
| 60S ribosomal protein L14 | Q9CR57 |
| 60S ribosomal protein L17 | Q9CPR4 |
| 60S ribosomal protein L18 | P35980 |
| 60S ribosomal protein L18a | P62717 |

| Detected in 2 experiments - 737 proteins | |
|---|---|
| **Protein Name** | **Uniprot ID** |
| **Detected in experiments 1&2 - 144 proteins** | |
| 10 kDa heat shock protein, mitochondrial | Q64433 |
| 2-aminoethanethiol dioxygenase | Q6PDY2 |
| 40S ribosomal protein S21 | Q9CQR2 |
| 60S ribosomal protein L19 | P84099 |
| 60S ribosomal protein L22 | P67984 |
| 60S ribosomal protein L23a | P62751 |
| 60S ribosomal protein L31 | P62900 |
| Acid ceramidase | Q9WV54 |
| Actin, cytoplasmic 1 | P60710 |
| Acyl-CoA-binding protein | P31786 |
| Adenine phosphoribosyltransferase | P08030 |
| Alpha-centractin | P61164 |
| Alpha-taxilin | Q6PAM1 |
| AMP deaminase 2 | Q9DBT5 |
| Amyloid-like protein 2 | Q06335 |
| Apolipoprotein A-IV | P06728 |
| Arfaptin-2 | Q8K221 |
| Arginine and glutamate-rich protein 1 | Q3UL36 |
| Astrocytic phosphoprotein PEA-15 | Q62048 |
| Calsyntenin-1 | Q9EPL2 |
| Cathepsin S | O70370 |
| Cellular nucleic acid-binding protein | P53996 |
| Charged multivesicular body protein 4b | Q9D8B3 |
| Charged multivesicular body protein 5 | Q9D7S9 |
| C-Myc-binding protein | Q9EQS3 |
| Coiled-coil domain-containing protein 9 | Q8VC31 |
| Coiled-coil domain-containing protein 91 | Q9D8L5 |
| Cystatin-B | Q62426 |
| Cytochrome c oxidase subunit 2 | P00405 |
| Cytochrome c oxidase subunit 6B1 | P56391 |
| DDB1- and CUL4-associated factor 7 | P61963 |
| Dickkopf-related protein 3 | Q9QUN9 |
| DNA-(apurinic or apyrimidinic site) lyase | P28352 |
| D-tyrosyl-tRNA(Tyr) deacylase 1 | Q9DD18 |
| Dystroglycan | Q62165 |
| E3 ubiquitin-protein ligase RBX1 | P62878 |
| Endophilin-B1 | Q9JK48 |
| Epsin-2 | Q8CHU3 |
| Eukaryotic translation initiation factor 3 subunit G | Q9Z1D1 |
| Fatty acid-binding protein, epidermal | Q05816 |
| F-box only protein 2 | Q80UW2 |
| FXYD domain-containing ion transport regulator 7 | P59648 |
| Galectin-1 | P16045 |
| Geranylgeranyl transferase type-2 subunit beta | P53612 |
| Glycylpeptide N-tetradecanoyltransferase 2 | O70311 |

| Detected in 1 experiment - 1180 proteins | |
|---|---|
| **Protein Name** | **Uniprot ID** |
| **Detected in experiment 1 only - 147 proteins** | |
| 28S ribosomal protein S31, mitochondrial | Q61733 |
| 39S ribosomal protein L28, mitochondrial | Q9D1B9 |
| 60S ribosomal protein L37 | Q9D823 |
| 80 kDa MCM3-associated protein | Q9WUU9 |
| Acyl-coenzyme A synthetase ACSM4, mitochond | Q80W40 |
| Acyl-coenzyme A thioesterase 10, mitochondrial | Q32MW3 |
| Apolipoprotein D | P51910 |
| Apoptosis-stimulating of p53 protein 1 | Q62415 |
| Arylsulfatase A | P50428 |
| ATP-dependent RNA helicase DDX3X | Q62167 |
| BRI3-binding protein | Q8BXV2 |
| Bridging integrator 2 | D3Z6Q9 |
| Cadherin EGF LAG seven-pass G-type receptor | Q9R0M0 |
| Calcium/calmodulin-dependent protein kinase typ | Q91YS8 |
| Calpastatin | P51125 |
| cAMP-dependent protein kinase inhibitor alpha | P63248 |
| Carboxylesterase 1D | Q8VCT4 |
| Carnitine O-acetyltransferase | P47934 |
| Caspase-3 | P70677 |
| CD81 antigen | P35762 |
| Cerebellar degeneration-related protein 2-like | A2A6T1 |
| Charged multivesicular body protein 2b | Q8BJF9 |
| Chromobox protein homolog 1 | P83917 |
| Cingulin-like protein 1 | Q6AW69 |
| CKLF-like MARVEL transmembrane domain-cont | Q8CJ61 |
| Cofilin-2 | P45591 |
| Coiled-coil domain-containing protein 104 | Q8C6E0 |
| Coiled-coil domain-containing protein 124 | Q9D8X2 |
| Coiled-coil domain-containing protein 136 | Q3TVA9 |
| Coiled-coil domain-containing protein 43 | Q9CR29 |
| Coiled-coil domain-containing protein 96 | Q9CR92 |
| Complexin-3 | Q8R1B5 |
| Cutaneous T-cell lymphoma-associated antigen 5 | Q8R311 |
| Cyclic AMP-responsive element-binding protein 1 | Q01147 |
| Cysteine-rich with EGF-like domain protein 1 | Q91XD7 |
| Diacylglycerol kinase theta | Q6P5E8 |
| DNA-directed RNA polymerase I subunit RPA1 | O35134 |
| DNA-directed RNA polymerase II subunit GRINL1 | Q6P6I6 |
| DnaJ homolog subfamily B member 5 | O89114 |
| DnaJ homolog subfamily C member 2 | P54103 |
| E3 ubiquitin-protein ligase BRE1A | Q5DTM8 |
| E3 ubiquitin-protein ligase HECTD1 | Q69ZR2 |
| E3 ubiquitin-protein ligase RNF187 | Q8BFX1 |
| E3 ubiquitin-protein ligase ZNRF2 | Q71FD5 |
| Eukaryotic peptide chain release factor GTP-bind | Q149F3 |

| | | | | | |
|---|---|---|---|---|---|
| 60S ribosomal protein L26 | P61255 | GMP reductase 2 | Q99L27 | Eukaryotic translation initiation factor 4E | P63073 |
| 60S ribosomal protein L3 | P27659 | Hemoglobin subunit beta-1 | P02088 | FCH domain only protein 2 | Q3UQN2 |
| 60S ribosomal protein L5 | P47962 | Hepatoma-derived growth factor-related protein 2 | Q3UMU9 | FGFR1 oncogene partner 2 homolog | Q9CRA9 |
| 60S ribosomal protein L6 | P47911 | Heterogeneous nuclear ribonucleoprotein U-like p | Q8VDM6 | GDNF family receptor alpha-1 | P97785 |
| 60S ribosomal protein L8 | P62918 | Histone deacetylase 6 | Q9Z2V5 | General transcription factor IIF subunit 2 | Q8R0A0 |
| 6-phosphofructokinase type C | Q9WUA3 | HIV Tat-specific factor 1 homolog | Q8BGC0 | Glucosylceramidase | P17439 |
| 6-phosphofructokinase, muscle type | P47857 | Hydroxyacyl-coenzyme A dehydrogenase, mitoch | Q61425 | Glutamate [NMDA] receptor subunit epsilon-3 | Q01098 |
| 6-phosphogluconate dehydrogenase, decarboxyl | Q9DCD0 | Immunoglobulin superfamily containing leucine-ric | Q5RKR3 | Glutamate carboxypeptidase 2 | O35409 |
| 78 kDa glucose-regulated protein | P20029 | Integrin beta-1 | P09055 | Glutaryl-CoA dehydrogenase, mitochondrial | Q60759 |
| Acetyl-CoA acetyltransferase, mitochondrial | Q8QZT1 | Integrin beta-2 | P11835 | G-protein coupled receptor-associated sorting pro | Q5U4C1 |
| Acidic leucine-rich nuclear phosphoprotein 32 fan | O35381 | Iron-sulfur cluster assembly 2 homolog, mitochon | Q9DCB8 | GRIP and coiled-coil domain-containing protein 1 | Q9D4H2 |
| Aconitate hydratase, mitochondrial | Q99KI0 | Janus kinase and microtubule-interacting protein | Q8BVL9 | Guanine nucleotide-binding protein G(t) subunit a | P20612 |
| Actin-binding LIM protein 1 | Q8K4G5 | Methionine aminopeptidase 2 | O08663 | HD domain-containing protein 2 | Q3SXD3 |
| Actin-binding LIM protein 2 | Q8BL65 | Mitochondrial import inner membrane translocase | P62073 | Heat shock protein beta-1 | P14602 |
| Actin-related protein 2 | P61161 | Mitochondrial import receptor subunit TOM34 | Q9CYG7 | High mobility group protein B2 | P30681 |
| Actin-related protein 2/3 complex subunit 1A | Q9R0Q6 | Mitochondrial peptide methionine sulfoxide reduct | Q9D6Y7 | Histidine triad nucleotide-binding protein 1 | P70349 |
| Actin-related protein 2/3 complex subunit 2 | Q9CVB6 | Moesin | P26041 | Inosine-5'-monophosphate dehydrogenase 1 | P50096 |
| Actin-related protein 2/3 complex subunit 4 | P59999 | Myeloid-associated differentiation marker | O35682 | Inositol hexakisphosphate kinase 1 | Q6PD10 |
| Actin-related protein 3 | Q99JY9 | Myosin regulatory light chain 12B | Q3THE2 | Katanin p80 WD40-containing subunit B1 | Q8BG40 |
| Activator of 90 kDa heat shock protein ATPase h | Q8BK64 | NADH dehydrogenase [ubiquinone] 1 beta subcor | Q9CQZ6 | Kinetochore-associated protein 1 | Q8C3Y4 |
| Active breakpoint cluster region-related protein | Q5SSL4 | NADH dehydrogenase [ubiquinone] 1 beta subcor | Q9CQH3 | Leucine-rich repeat flightless-interacting protein 1 | Q3UZ39 |
| Acylamino-acid-releasing enzyme | Q8R146 | N-alpha-acetyltransferase 38, NatC auxiliary subu | Q6ZWM4 | LIM and cysteine-rich domains protein 1 | Q8VEE1 |
| Acyl-CoA dehydrogenase family member 9, mitoc | Q8JZN5 | Neogenin | P97798 | MAP6 domain-containing protein 1 | Q14BB9 |
| Acyl-CoA synthetase family member 3, mitochon | Q3URE1 | Nuclear fragile X mental retardation-interacting pr | Q5F2E7 | Metabotropic glutamate receptor 1 | P97772 |
| Acyl-CoA-binding domain-containing protein 6 | Q9D061 | Nuclease EXOG, mitochondrial | Q8C163 | MOB kinase activator 2 | Q8VI63 |
| Acylphosphatase-2 | P56375 | Nucleobindin-1 | Q02819 | N-alpha-acetyltransferase 10 | Q9QY36 |
| Adapter molecule crk | Q64010 | Nucleobindin-2 | P81117 | NEDD8 ultimate buster 1 | P54729 |
| Adaptin ear-binding coat-associated protein 1 | Q9CR95 | Nucleolar protein 3 | Q9D1X0 | NEDD8-activating enzyme E1 catalytic subunit | Q8C878 |
| Adenosine kinase | P55264 | Nucleoside diphosphate kinase B | Q01768 | Neuroendocrine convertase 2 | P21661 |
| Adenosylhomocysteinase | P50247 | Partner of Y14 and mago | Q8CHP5 | Neuronal tyrosine-phosphorylated phosphoinositi | Q8BM65 |
| Adenylate cyclase type 9 | P51830 | Parvalbumin alpha | P32848 | NHS-like protein 1 | Q8CAF4 |
| Adenylate kinase isoenzyme 1 | Q9R0Y5 | Piezo-type mechanosensitive ion channel compo | Q8CD54 | Nicalin | Q8VCM8 |
| Adenylate kinase isoenzyme 5 | Q920P5 | PITH domain-containing protein 1 | Q8BWR2 | NIF3-like protein 1 | Q9EQ80 |
| Adenylyl cyclase-associated protein 1 | P40124 | Plastin-2 | Q61233 | NK-tumor recognition protein | P30415 |
| Adenylyl cyclase-associated protein 2 | Q9CYT6 | Poly(ADP-ribose) glycohydrolase ARH3 | Q8CG72 | Nuclear receptor-binding factor 2 | Q8VCQ3 |
| ADP/ATP translocase 1 | P48962 | Potassium voltage-gated channel subfamily D me | Q9Z0V1 | Nucleolar protein 16 | Q9CPT5 |
| ADP-ribosylation factor 5 | P84084 | Prefoldin subunit 1 | Q9CWM4 | OTU domain-containing protein 6B | Q8K2H2 |
| ADP-ribosylation factor-like protein 8B | Q9CQW2 | Prefoldin subunit 3 | P61759 | Palmitoyl-protein thioesterase 1 | O88531 |
| Afadin | Q9QZQ1 | Prostaglandin-H2 D-isomerase | O09114 | Peptidyl-prolyl cis-trans isomerase-like 1 | Q9D0W5 |
| AFG3-like protein 2 | Q8JZQ2 | Proteasomal ubiquitin receptor ADRM1 | Q9JKV1 | Peptidyl-tRNA hydrolase ICT1, mitochondrial | Q8R035 |
| Aggrecan core protein | Q61282 | Proteasome activator complex subunit 1 | P97371 | Phosphatase and actin regulator 4 | Q501J7 |
| AH receptor-interacting protein | O08915 | Proteasome activator complex subunit 2 | P97372 | Phosphoglucomutase-2 | Q7TSV4 |
| A-kinase anchor protein 5 | D3YVF0 | Protein arginine N-methyltransferase 5 | Q8CIG8 | Phosphomannomutase 2 | Q9Z2M7 |
| A-kinase anchor protein 9 | Q70FJ1 | Protein canopy homolog 2 | Q9QXT0 | Phosphopantothenate--cysteine ligase | Q8VDG5 |
| Alanine aminotransferase 1 | Q8QZR5 | Protein dpy-30 homolog | Q99LT0 | Platelet-activating factor acetylhydrolase | Q60963 |
| Alanine--tRNA ligase, cytoplasmic | Q8BGQ7 | Protein FAM192A | Q91WE2 | Poly [ADP-ribose] polymerase 1 | P11103 |
| Alcohol dehydrogenase [NADP(+)] | Q9JII6 | Protein NipSnap homolog 1 | O55125 | Protein bicaudal D homolog 2 | Q921C5 |
| Alcohol dehydrogenase class-3 | P28474 | Protein phosphatase 1 regulatory subunit 1A | Q9ERT9 | Protein disulfide-isomerase TMX3 | Q8BXZ1 |

| | | | | | |
|---|---|---|---|---|---|
| Aldehyde dehydrogenase X, mitochondrial | Q9CZS1 | Protein phosphatase 1B | P36993 | Protein ECT2 | Q07139 |
| Aldehyde dehydrogenase, mitochondrial | P47738 | Protein PRRC2A | Q7TSC1 | Protein FAM171B | Q14CH0 |
| Aldose reductase | P45376 | Protein quaking | Q9QYS9 | Protein HEXIM1 | Q8R409 |
| Alpha-2-macroglobulin receptor-associated prote | P55302 | Putative RNA-binding protein Luc7-like 2 | Q7TNC4 | Protein transport protein Sec61 subunit alpha isof | P61620 |
| Alpha-actinin-1 | Q7TPR4 | Rab GTPase-binding effector protein 2 | Q91WG2 | Protein transport protein Sec61 subunit beta | Q9CQS8 |
| Alpha-actinin-4 | P57780 | Rab-like protein 1 | Q5U3K5 | Protein-L-isoaspartate O-methyltransferase doma | P59913 |
| Alpha-adducin | Q9QYC0 | Ras GTPase-activating protein-binding protein 1 | P97855 | Putative hydrolase RBBP9 | O88851 |
| Alpha-aminoadipic semialdehyde dehydrogenase | Q9DBF1 | Ras-related protein Rab-18 | P35293 | Pyridoxal-dependent decarboxylase domain-conta | Q99K01 |
| Alpha-endosulfine | P60840 | Receptor expression-enhancing protein 5 | Q60870 | Receptor-type tyrosine-protein phosphatase beta | B2RU80 |
| Alpha-enolase | P17182 | Reticulocalbin-1 | Q05186 | Receptor-type tyrosine-protein phosphatase gam | Q05909 |
| Alpha-internexin | P46660 | Rho GDP-dissociation inhibitor 2 | Q61599 | Regulator of G-protein signaling 10 | Q9CQE5 |
| Alpha-soluble NSF attachment protein | Q9DB05 | Ribosome-recycling factor, mitochondrial | Q9D6S7 | Ribosyldihydronicotinamide dehydrogenase [quin | Q9JI75 |
| Amine oxidase [flavin-containing] A | Q64133 | Roundabout homolog 1 | O89026 | RuvB-like 1 | P60122 |
| Amine oxidase [flavin-containing] B | Q8BW75 | Secretogranin-3 | P47867 | RWD domain-containing protein 4 | Q9CPR1 |
| Amphiphysin | Q7TQF7 | Seizure 6-like protein | Q6P1D5 | Segment polarity protein dishevelled homolog DV | Q61062 |
| Amyloid beta A4 precursor protein-binding family | B2RUJ5 | Serine/threonine-protein phosphatase 2A 55 kDa | Q6ZWR4 | Selenium-binding protein 1 | P17563 |
| Amyloid beta A4 protein | P12023 | Serine/threonine-protein phosphatase 6 regulator | Q922D4 | Septin-6 | Q9R1T4 |
| Amyloid-like protein 1 | Q03157 | Serpin B6 | Q60854 | Serine protease inhibitor A3N | Q91WP6 |
| Ankyrin repeat and sterile alpha motif domain-co | Q8BIZ1 | Serrate RNA effector molecule homolog | Q99MR6 | Serine/arginine-rich splicing factor 5 | O35326 |
| Ankyrin-2 | Q8C8R3 | Shootin-1 | Q8K2Q9 | Serine/threonine-protein kinase 4 | Q9JI11 |
| Annexin A11 | P97384 | Single-stranded DNA-binding protein, mitochondr | Q9CYR0 | Serine/threonine-protein kinase Nek9 | Q8K1R7 |
| Annexin A3 | O35639 | Small VCP/p97-interacting protein | Q3UZP4 | Serine/threonine-protein phosphatase 2A 55 kDa | Q925E7 |
| Annexin A6 | P14824 | S-methyl-5'-thioadenosine phosphorylase | Q9CQ65 | SET and MYND domain-containing protein 5 | Q3TYX3 |
| Annexin A7 | Q07076 | Sorbitol dehydrogenase | Q64442 | Signal peptidase complex subunit 2 | Q9CYN2 |
| AP-1 complex subunit beta-1 | O35643 | Sorting and assembly machinery component 50 h | Q8BGH2 | Small glutamine-rich tetratricopeptide repeat-cont | Q8VD33 |
| AP-1 complex subunit gamma-1 | P22892 | Sorting nexin-16 | Q8C080 | Small ubiquitin-related modifier 1 | P63166 |
| AP-2 complex subunit alpha-1 | P17426 | SPARC-like protein 1 | P70663 | Sodium-dependent noradrenaline transporter | O55192 |
| AP-2 complex subunit alpha-2 | P17427 | Splicing factor 1 | Q64213 | Sodium-dependent phosphate transporter 1 | Q61609 |
| AP-2 complex subunit beta | Q9DBG3 | Tax_Id=10090 Gene_Symbol=Krt33b keratin com | S.avidinii | Son of sevenless homolog 2 | Q02384 |
| AP-2 complex subunit mu | P84091 | Thymosin beta-10 | Q6ZWY8 | Stathmin-4 | P63042 |
| AP2-associated protein kinase 1 | Q3UHJ0 | Toll-interacting protein | Q9QZ06 | Steroid receptor RNA activator 1 | Q80VJ2 |
| AP-3 complex subunit beta-2 | Q9JME5 | Transcription factor A, mitochondrial | P40630 | Stromal membrane-associated protein 2 | Q7TN29 |
| AP-3 complex subunit delta-1 | O54774 | TSC22 domain family protein 1 | P62500 | Swi5-dependent recombination DNA repair protei | Q8BP27 |
| Apolipoprotein A-I | Q00623 | TSC22 domain family protein 3 | Q9Z2S7 | Synapse-associated protein 1 | Q9D5V6 |
| Apolipoprotein E | P08226 | Tudor and KH domain-containing protein | Q80VL1 | Target of Myb protein 1 | O88746 |
| Apoptosis-inducing factor 1, mitochondrial | Q9Z0X1 | U6 snRNA-associated Sm-like protein LSm2 | O35900 | TBC1 domain family member 15 | Q9CXF4 |
| Aquaporin-4 | P55088 | Ubiquitin domain-containing protein UBFD1 | Q78JW9 | Tetraspanin-18 | Q80WR1 |
| ARF GTPase-activating protein GIT1 | Q68FF6 | Ubiquitin fusion degradation protein 1 homolog | P70362 | Thioredoxin-related transmembrane protein 1 | Q8VBT0 |
| Arf-GAP with GTPase, ANK repeat and PH doma | Q8BXK8 | Ubiquitin-conjugating enzyme E2 variant 2 | Q9D2M8 | Translational activator of cytochrome c oxidase 1 | Q8K0Z7 |
| Arf-GAP with GTPase, ANK repeat and PH doma | Q3UHD9 | Ubiquitin-fold modifier 1 | P61961 | Translin | Q62348 |
| Arf-GAP with GTPase, ANK repeat and PH doma | Q8VHH5 | UBX domain-containing protein 1 | Q922Y1 | Transmembrane protein 119 | Q8R138 |
| Arf-GAP with SH3 domain, ANK repeat and PH d | Q9QWY8 | Uncharacterized protein C6orf174 homolog | Q6NZL0 | Transmembrane protein 9 | Q9CR23 |
| Arginine--tRNA ligase, cytoplasmic | Q9D0I9 | Uncharacterized protein KIAA1211 | Q5PR69 | tRNA (adenine(58)-N(1))-methyltransferase non-c | Q8CE96 |
| Argininosuccinate lyase | Q91YI0 | UPF0366 protein C11orf67 homolog | Q8R0P4 | Tyrosine-protein kinase BAZ1B | Q9Z277 |
| Argininosuccinate synthase | P16460 | UPF0449 protein C19orf25 homolog | Q9D7E4 | Ubiquilin-4 | Q99NB8 |
| Armadillo repeat-containing protein 6 | Q8BNU0 | UPF0515 protein C19orf66 homolog | Q8CAK3 | Ubiquitin carboxyl-terminal hydrolase 47 | Q8BY87 |
| Asparagine--tRNA ligase, cytoplasmic | Q8BP47 | UPF0552 protein C15orf38 homolog | Q9D0A3 | Unconventional myosin-Vb | P21271 |
| Aspartate aminotransferase, cytoplasmic | P05201 | UPF0587 protein C1orf123 homolog | Q8BHG2 | UPF0600 protein C5orf51 homolog | Q8BR90 |

| | |
|---|---|
| Aspartate aminotransferase, mitochondrial | P05202 |
| Aspartate--tRNA ligase, cytoplasmic | Q922B2 |
| Aspartyl aminopeptidase | Q9Z2W0 |
| Astrotactin-1 | Q61137 |
| Ataxin-10 | P28658 |
| Ataxin-2-like protein | Q7TQH0 |
| ATP synthase subunit alpha, mitochondrial | Q03265 |
| ATP synthase subunit b, mitochondrial | Q9CQQ7 |
| ATP synthase subunit beta, mitochondrial | P56480 |
| ATP synthase subunit d, mitochondrial | Q9DCX2 |
| ATP synthase subunit delta, mitochondrial | Q9D3D9 |
| ATP synthase subunit gamma, mitochondrial | Q91VR2 |
| ATP synthase subunit O, mitochondrial | Q9DB20 |
| ATP synthase-coupling factor 6, mitochondrial | P97450 |
| ATPase family AAA domain-containing protein 3 | Q925I1 |
| ATP-binding cassette sub-family E member 1 | P61222 |
| ATP-binding cassette sub-family F member 1 | Q6P542 |
| ATP-citrate synthase | Q91V92 |
| ATP-dependent RNA helicase DDX1 | Q91VR5 |
| Band 4.1-like protein 1 | Q9Z2H5 |
| Band 4.1-like protein 2 | O70318 |
| Band 4.1-like protein 3 | Q9WV92 |
| Basic leucine zipper and W2 domain-containing | Q9CQC6 |
| Basigin | P18572 |
| Beta-adducin | Q9QYB8 |
| Beta-adrenergic receptor kinase 1 | Q99MK8 |
| Beta-arrestin-1 | Q8BWG8 |
| Beta-centractin | Q8R5C5 |
| Beta-hexosaminidase subunit beta | P20060 |
| Beta-soluble NSF attachment protein | P28663 |
| Beta-synuclein | Q91ZZ3 |
| Bifunctional ATP-dependent dihydroxyacetone ki | Q8VC30 |
| Bifunctional glutamate/proline--tRNA ligase | Q8CGC7 |
| Bifunctional protein NCOAT | Q9EQQ9 |
| Bifunctional purine biosynthesis protein PURH | Q9CWJ9 |
| Biliverdin reductase A | Q9CY64 |
| Brain acid soluble protein 1 | Q91XV3 |
| Brain-enriched guanylate kinase-associated prot | Q68EF6 |
| Brain-specific angiogenesis inhibitor 1-associate | Q8BKX1 |
| Breast carcinoma-amplified sequence 1 homolog | Q80YN3 |
| Brefeldin A-inhibited guanine nucleotide-exchang | Q3UGY8 |
| Brevican core protein | Q61361 |
| BTB/POZ domain-containing protein KCTD12 | Q6WVG3 |
| C-1-tetrahydrofolate synthase, cytoplasmic | Q922D8 |
| Cadherin-13 | Q9WTR5 |
| Calbindin | P12658 |
| Calcineurin subunit B type 1 | Q63810 |
| Calcium uniporter protein, mitochondrial | Q3UMR5 |
| WD repeat-containing protein 37 | Q8CBE3 |
| WD repeat-containing protein 47 | Q8CGF6 |
| Zinc finger protein ZPR1 | Q62384 |
| **Detected in experiments 1&3 - 191 proteins** | |
| 1-phosphatidylinositol 4,5-bisphosphate phospho | Q8R3B1 |
| 26S protease regulatory subunit 10B | P62334 |
| 26S protease regulatory subunit 8 | P62196 |
| 26S proteasome non-ATPase regulatory subunit | Q9WVJ2 |
| 26S proteasome non-ATPase regulatory subunit | Q8BJY1 |
| 3-mercaptopyruvate sulfurtransferase | Q99J99 |
| 40S ribosomal protein S10 | P63325 |
| 40S ribosomal protein S27 | Q6ZWU9 |
| 40S ribosomal protein S27-like | Q6ZWY3 |
| 5'-3' exoribonuclease 1 | P97789 |
| Acetoacetyl-CoA synthetase | Q9D2R0 |
| Actin, aortic smooth muscle | P62737 |
| Acyl-CoA synthetase family member 2, mitochond | Q8VCW8 |
| Acyl-coenzyme A thioesterase 2, mitochondrial | Q9QYR9 |
| ADP-ribosylation factor 2 | Q8BSL7 |
| ADP-ribosylation factor-like protein 8A | Q8VEH3 |
| Aftiphilin | Q80WT5 |
| Alanyl-tRNA editing protein Aarsd1 | Q3THG9 |
| Alpha-parvin | Q9EPC1 |
| Aminoacylase-1 | Q99JW2 |
| Amyloid beta A4 precursor protein-binding family | Q9QXJ1 |
| Asparagine synthetase [glutamine-hydrolyzing] | Q61024 |
| Atlastin-2 | Q6PA06 |
| ATPase inhibitor, mitochondrial | O35143 |
| Autophagy-related protein 16-1 | Q8C0J2 |
| Baculoviral IAP repeat-containing protein 6 | O88738 |
| BAG family molecular chaperone regulator 5 | Q8CI32 |
| Bifunctional 3'-phosphoadenosine 5'-phosphosulf | Q60967 |
| BRISC complex subunit Abro1 | Q3TCJ1 |
| Calmodulin-regulated spectrin-associated protein | Q8C1B1 |
| Calpain-1 catalytic subunit | O35350 |
| Carbonic anhydrase 4 | Q64444 |
| Catechol O-methyltransferase domain-containing | Q8BIG7 |
| CB1 cannabinoid receptor-interacting protein 1 | Q5M8N0 |
| CCA tRNA nucleotidyltransferase 1, mitochondria | Q8K1J6 |
| cGMP-dependent 3',5'-cyclic phosphodiesterase | Q922S4 |
| Choline transporter-like protein 2 | Q8BY89 |
| C-Jun-amino-terminal kinase-interacting protein 3 | Q9ESN9 |
| Clathrin interactor 1 | Q99KN9 |
| Cleavage stimulation factor subunit 2 | Q8BIQ5 |
| Coatomer subunit epsilon | O89079 |
| Coiled-coil domain-containing protein 25 | Q78PG9 |
| Cold shock domain-containing protein E1 | Q91W50 |
| Constitutive coactivator of PPAR-gamma-like pro | Q8C3F2 |
| Vacuolar protein sorting-associated protein 28 ho | Q9D1C8 |
| Vacuolar protein sorting-associated protein 4B | P46467 |
| Vacuolar-sorting protein SNF8 | Q9CZ28 |
| Vesicle transport protein SEC20 | Q6QD59 |
| Voltage-dependent L-type calcium channel subur | P54285 |
| YTH domain family protein 3 | Q8BYK6 |
| **Detected in experiment 2 only - 300 proteins** | |
| 1-phosphatidylinositol 4,5-bisphosphate phospho | P51432 |
| 28S ribosomal protein S30, mitochondrial | Q9D0G0 |
| 39S ribosomal protein L12, mitochondrial | Q9DB15 |
| 39S ribosomal protein L22, mitochondrial | Q8BU88 |
| 39S ribosomal protein L46, mitochondrial | Q9EQI8 |
| 39S ribosomal protein L49, mitochondrial | Q9CQ40 |
| 3-hydroxyacyl-CoA dehydratase 3 | Q8K2C9 |
| 40S ribosomal protein S26 | P62855 |
| 60S ribosomal protein L12 | P35979 |
| Abhydrolase domain-containing protein 3 | Q91ZH7 |
| Actin, alpha skeletal muscle | P68134 |
| Acyl-protein thioesterase 2 | Q9WTL7 |
| Adenylate kinase isoenzyme 4, mitochondrial | Q9WUR9 |
| ADP-ribose pyrophosphatase, mitochondrial | Q8BVU5 |
| ADP-ribosylation factor 3 | P61205 |
| ADP-sugar pyrophosphatase | Q9JKX6 |
| Adrenodoxin-like protein, mitochondrial | Q9CPW2 |
| Aflatoxin B1 aldehyde reductase member 2 | Q8CG76 |
| Aldose reductase-related protein 1 | P21300 |
| Alkaline phosphatase, tissue-nonspecific isozyme | P09242 |
| Alpha-crystallin B chain | P23927 |
| Alpha-ketoglutarate-dependent dioxygenase FTO | Q8BGW1 |
| AN1-type zinc finger protein 6 | Q9DCH6 |
| Anamorsin | Q8WTY4 |
| Ankyrin repeat domain-containing protein 24 | Q80VM7 |
| APC membrane recruitment protein 2 | Q8CCJ4 |
| Ataxin-2 | O70305 |
| ATP synthase mitochondrial F1 complex assembl | Q811I0 |
| ATP synthase subunit e, mitochondrial | Q06185 |
| ATP synthase subunit epsilon, mitochondrial | P56382 |
| ATP-dependent DNA helicase Q1 | Q9Z129 |
| BAG family molecular chaperone regulator 3 | Q9JLV1 |
| B-cell receptor-associated protein 31 | Q61335 |
| Beta-crystallin A1 | P02525 |
| Beta-galactosidase | P23780 |
| BH3-interacting domain death agonist | P70444 |
| BolA-like protein 2 | Q8BGS2 |
| Brain-specific angiogenesis inhibitor 1 | Q3UHD1 |
| Branched-chain-amino-acid aminotransferase, cy | P24288 |
| BTB/POZ domain-containing adapter for CUL3-m | Q8BGV7 |
| C1GALT1-specific chaperone 1 | Q9JMG2 |

| | | | | | |
|---|---|---|---|---|---|
| Calcium/calmodulin-dependent 3',5'-cyclic nucle | Q61481 | COP9 signalosome complex subunit 2 | P61202 | Calcineurin-like phosphoesterase domain-contain | Q8BFS6 |
| Calcium/calmodulin-dependent 3',5'-cyclic nucle | Q01065 | CREB-regulated transcription coactivator 1 | Q68ED7 | Calcium/calmodulin-dependent protein kinase kin | Q8VBY2 |
| Calcium/calmodulin-dependent protein kinase typ | Q8BW96 | Cyclin-dependent kinase 14 | O35495 | Calponin-3 | Q9DAW9 |
| Calcium/calmodulin-dependent protein kinase typ | P11798 | Cytochrome b5 type B | Q9CQX2 | cAMP-regulated phosphoprotein 19 | P56212 |
| Calcium/calmodulin-dependent protein kinase typ | P28652 | DBIRD complex subunit KIAA1967 homolog | Q8VDP4 | Carboxymethylenebutenolidase homolog | Q8R1G2 |
| Calcium/calmodulin-dependent protein kinase typ | Q923T9 | DCC-interacting protein 13-alpha | Q8K3H0 | Cartilage acidic protein 1 | Q8R555 |
| Calcium/calmodulin-dependent protein kinase typ | P08414 | Delta(3,5)-Delta(2,4)-dienoyl-CoA isomerase, mit | O35459 | Casein kinase II subunit alpha' | O54833 |
| Calcium-binding and coiled-coil domain-containir | Q8CGU1 | Dipeptidyl peptidase 9 | Q8BVG4 | Catechol O-methyltransferase | O88587 |
| Calcium-binding protein p22 | P61022 | DnaJ homolog subfamily A member 1 | P63037 | Cathepsin B | P10605 |
| Calcium-dependent secretion activator 1 | Q80TJ1 | DnaJ homolog subfamily C member 11 | Q5U458 | CCR4-NOT transcription complex subunit 1 | Q6ZQ08 |
| Calcium-dependent secretion activator 2 | Q8BYR5 | DnaJ homolog subfamily C member 5 | P60904 | Cell cycle control protein 50A | Q8VEK0 |
| Calcyclin-binding protein | Q9CXW3 | DnaJ homolog subfamily C member 7 | Q9QYI3 | Centrin-2 | Q9R1K9 |
| Calmin | Q8C5W0 | Dynactin subunit 3 | Q9Z0Y1 | Centrosomal protein of 128 kDa | Q8BI22 |
| Calmodulin | P62204 | Dystrophin | P11531 | Centrosomal protein of 290 kDa | Q6A078 |
| Calmodulin-regulated spectrin-associated proteir | A2AHC3 | E3 SUMO-protein ligase RanBP2 | Q9ERU9 | Centrosome-associated protein CEP250 | Q60952 |
| Calnexin | P35564 | E3 ubiquitin-protein ligase LRSAM1 | Q80ZI6 | Ceruloplasmin | Q61147 |
| Calpain small subunit 1 | O88456 | E3 ubiquitin-protein ligase NEDD4-like | Q8CFI0 | Charged multivesicular body protein 1a | Q921W0 |
| Calpain-2 catalytic subunit | O08529 | E3 ubiquitin-protein ligase RNF14 | Q9JI90 | Charged multivesicular body protein 1b-2 | Q9CQD4 |
| Calreticulin | P14211 | Echinoderm microtubule-associated protein-like 1 | Q05BC3 | Charged multivesicular body protein 2a | Q9DB34 |
| Calretinin | Q08331 | Endoplasmic reticulum resident protein 29 | P57759 | Chromogranin-A | P26339 |
| Calumenin | O35887 | Epidermal growth factor receptor kinase substrate | Q08509 | Coiled-coil domain-containing protein 18 | Q640L5 |
| CaM kinase-like vesicle-associated protein | Q3UHL1 | Eukaryotic initiation factor 4A-I | P60843 | Coiled-coil domain-containing protein 39 | Q9D5Y1 |
| cAMP-dependent protein kinase type I-alpha regu | Q9DBC7 | Exocyst complex component 1 | Q8R3S6 | Complement factor H | P06909 |
| cAMP-specific 3',5'-cyclic phosphodiesterase 4D | Q01063 | Exportin-1 | Q6P5F9 | Corticotropin-releasing factor-binding protein | Q60571 |
| CAP-Gly domain-containing linker protein 1 | Q922J3 | F-box/LRR-repeat protein 16 | A2RT62 | Creatine kinase M-type | P07310 |
| CAP-Gly domain-containing linker protein 2 | Q9Z0H8 | FERM, RhoGEF and pleckstrin domain-containing | Q91VS8 | C-terminal-binding protein 2 | P56546 |
| Caprin-1 | Q60865 | Fragile X mental retardation protein 1 homolog | P35922 | CUE domain-containing protein 2 | Q9CXX9 |
| Carbonyl reductase [NADPH] 1 | P48758 | Fragile X mental retardation syndrome-related pro | Q9WVR4 | Cyclin-dependent kinase inhibitor 1B | P46414 |
| Carbonyl reductase [NADPH] 3 | Q8K354 | General receptor for phosphoinositides 1-associa | Q9JJA9 | Cytochrome c, somatic | P62897 |
| Carboxypeptidase E | Q00493 | General transcription factor II-I | Q9ESZ8 | Cytosolic purine 5'-nucleotidase | Q3V1L4 |
| Caskin-1 | Q6P9K8 | Glia maturation factor beta | Q9CQI3 | Density-regulated protein | Q9CQJ6 |
| Catenin alpha-1 | P26231 | Glutamate [NMDA] receptor subunit zeta-1 | P35438 | Deoxyguanosine kinase, mitochondrial | Q9QX60 |
| Catenin alpha-2 | Q61301 | Glutamate decarboxylase 1 | P48318 | DNA polymerase subunit gamma-1 | P54099 |
| Catenin beta-1 | Q02248 | Glutamate receptor 3 | Q9Z2W9 | DNA-binding protein A | Q9JKB3 |
| Catenin delta-2 | O35927 | Glutathione S-transferase omega-1 | O09131 | DnaJ homolog subfamily B member 1 | Q9QYJ3 |
| Cathepsin D | P18242 | Glycerophosphodiester phosphodiesterase doma | Q9CRY7 | Dolichyl-diphosphooligosaccharide--protein glyco | O54734 |
| Caytaxin | Q8BHE3 | Glycogen synthase kinase-3 beta | Q9WV60 | Dynactin subunit 5 | Q9QZB9 |
| CD166 antigen | Q61490 | Growth arrest-specific protein 7 | Q60780 | E3 ubiquitin-protein ligase ARIH1 | Q9Z1K5 |
| Cell adhesion molecule 1 | Q8R5M8 | HEAT repeat-containing protein 5B | Q8C547 | E3 ubiquitin-protein ligase HECW2 | Q6I6G8 |
| Cell adhesion molecule 2 | Q8BLQ9 | Hepatocyte growth factor-regulated tyrosine kinas | Q99LI8 | E3 ubiquitin-protein ligase UBR5 | Q80TP3 |
| Centrosomal protein of 170 kDa | Q6A065 | Histamine N-methyltransferase | Q91VF2 | Ectonucleoside triphosphate diphosphohydrolase | O55026 |
| Charged multivesicular body protein 6 | P0C0A3 | Histone deacetylase 4 | Q6NZM9 | ELAV-like protein 4 | Q61701 |
| Chondroitin sulfate proteoglycan 5 | Q71M36 | Histone H1.0 | P10922 | Embigin | P21995 |
| Chromobox protein homolog 3 | P23198 | Hsp90 co-chaperone Cdc37-like 1 | Q9CZP7 | Enolase-phosphatase E1 | Q8BGB7 |
| Citrate lyase subunit beta-like protein, mitochond | Q8R4N0 | Huntingtin-interacting protein 1 | Q8VD75 | Ephrin type-A receptor 2 | Q03145 |
| Citrate synthase, mitochondrial | Q9CZU6 | Hydroxymethylglutaryl-CoA synthase, cytoplasmic | Q8JZK9 | Ephrin type-A receptor 7 | Q61772 |
| C-Jun-amino-terminal kinase-interacting protein 4 | Q58A65 | Immunoglobulin-binding protein 1 | Q61249 | Epoxide hydrolase 1 | Q9D379 |
| Clathrin coat assembly protein AP180 | Q61548 | Importin-4 | Q8VI75 | ERO1-like protein alpha | Q8R180 |

| | | | | | |
|---|---|---|---|---|---|
| Clathrin heavy chain 1 | Q68FD5 | Importin-5 | Q8BKC5 | Eukaryotic translation initiation factor 2-alpha kina | Q9Z2B5 |
| Clathrin light chain A | O08585 | Inactive phospholipase C-like protein 1 | Q3USB7 | Eukaryotic translation initiation factor 6 | O55135 |
| Clathrin light chain B | Q6IRU5 | Inositol polyphosphate 5-phosphatase K | Q8C5L6 | Excitatory amino acid transporter 1 | P56564 |
| CLIP-associating protein 1 | Q80TV8 | Kinesin-associated protein 3 | P70188 | FERM domain-containing protein 4A | Q8BIE6 |
| CLIP-associating protein 2 | Q8BRT1 | Lambda-crystallin homolog | Q99KP3 | FK506-binding protein 15 | Q6P9Q6 |
| Clusterin | Q06890 | Late secretory pathway protein AVL9 homolog | Q80U56 | Fumarylacetoacetase | P35505 |
| Coactosin-like protein | Q9CQI6 | Leucine-rich repeat-containing protein 47 | Q505F5 | G protein-regulated inducer of neurite outgrowth 3 | Q8BWS5 |
| Coatomer subunit alpha | Q8CIE6 | Lipid phosphate phosphohydrolase 3 | Q99JY8 | GDP-mannose 4,6 dehydratase | Q8K0C9 |
| Coatomer subunit zeta-1 | P61924 | MAP/microtubule affinity-regulating kinase 3 | Q03141 | Glutamate receptor-interacting protein 1 | Q925T6 |
| Cofilin-1 | P18760 | Membrane-associated guanylate kinase, WW and | Q6RHR9 | Glutathione S-transferase Mu 2 | P15626 |
| Coiled-coil and C2 domain-containing protein 1A | Q8K1A6 | Mitogen-activated protein kinase 9 | Q9WTU6 | Glutathione S-transferase P 2 | P46425 |
| Coiled-coil domain-containing protein 127 | Q3TC33 | Monofunctional C1-tetrahydrofolate synthase, mit | Q3V3R1 | Glycerol kinase | Q64516 |
| Coiled-coil domain-containing protein 132 | Q8CI71 | Multifunctional protein ADE2 | Q9DCL9 | Golgi-associated PDZ and coiled-coil motif-contai | Q8BH60 |
| Coiled-coil domain-containing protein 165 | Q3UHU5 | NADH dehydrogenase [ubiquinone] 1 alpha subco | Q99LC3 | Golgin subfamily A member 5 | Q9QYE6 |
| Coiled-coil domain-containing protein 6 | D3YZP9 | NADP-dependent malic enzyme, mitochondrial | Q8BMF3 | GTPase KRas | P32883 |
| Coiled-coil domain-containing protein 93 | Q7TQK5 | N-alpha-acetyltransferase 25, NatB auxiliary subu | Q8BWZ3 | Guanine nucleotide-binding protein G(I)/G(S)/G(O | P63216 |
| Coiled-coil-helix-coiled-coil-helix domain-containi | Q9CRB9 | Nardilysin | Q8BHG1 | Guanine nucleotide-binding protein G(I)/G(S)/G(O | Q61016 |
| Coiled-coil-helix-coiled-coil-helix domain-containi | Q91VN4 | Neuroligin-1 | Q99K10 | Guanine nucleotide-binding protein G(I)/G(S)/G(T | P62880 |
| Complexin-1 | P63040 | Neurolysin, mitochondrial | Q91YP2 | Haloacid dehalogenase-like hydrolase domain-co | Q9CYW4 |
| Complexin-2 | P84086 | Nischarin | Q80TM9 | Hamartin | Q9EP53 |
| Connector enhancer of kinase suppressor of ras | Q80YA9 | NT-3 growth factor receptor | Q6VNS1 | Heat shock factor-binding protein 1 | Q9CQZ1 |
| Contactin-1 | P12960 | Nuclear protein localization protein 4 homolog | P60670 | Hematological and neurological expressed 1 prote | P97825 |
| Contactin-associated protein 1 | O54991 | Opioid growth factor receptor-like protein 1 | Q8VE52 | Hemopexin | Q91X72 |
| COP9 signalosome complex subunit 1 | Q99LD4 | PDZ domain-containing protein GIPC1 | Q9Z0G0 | Hepatocyte cell adhesion molecule | Q640R3 |
| COP9 signalosome complex subunit 3 | O88543 | Peptidyl-prolyl cis-trans isomerase FKBP5 | Q64378 | High mobility group protein B1 | P63158 |
| COP9 signalosome complex subunit 4 | O88544 | Phosphatidylinositide phosphatase SAC2 | Q8CDA1 | Hippocalcin-like protein 1 | P62748 |
| COP9 signalosome complex subunit 7a | Q9CZ04 | Phosphatidylinositol 3,4,5-trisphosphate-depende | Q69ZK0 | Histone-lysine N-methyltransferase setd3 | Q91WC0 |
| COP9 signalosome complex subunit 8 | Q8VBV7 | Phosphatidylinositol transfer protein beta isoform | P53811 | Huntingtin-interacting protein K | Q9CR41 |
| Copine-2 | P59108 | Phospholipase DDHD2 | Q80Y98 | Immunoglobulin superfamily member 21 | Q7TNR6 |
| Copine-6 | Q9Z140 | Plastin-3 | Q99K51 | Immunoglobulin-like and fibronectin type III doma | Q3KNY0 |
| Copine-7 | Q0VE82 | Plexin-B2 | B2RXS4 | Inositol 1,4,5-trisphosphate receptor type 2 | Q9Z329 |
| Coronin-1A | O89053 | Polyadenylate-binding protein-interacting protein | Q8VE62 | Inositol-3-phosphate synthase 1 | Q9JHU9 |
| Coronin-7 | Q9D2V7 | Porphobilinogen deaminase | P22907 | Isopentenyl-diphosphate Delta-isomerase 1 | P58044 |
| Creatine kinase B-type | Q04447 | Probable ATP-dependent RNA helicase DDX17 | Q501J6 | Kelch-like protein 9 | Q6ZPT1 |
| Creatine kinase U-type, mitochondrial | P30275 | Probable ATP-dependent RNA helicase DDX5 | Q61656 | Kinesin-like protein KIF20B | Q80WE4 |
| Cullin-3 | Q9JLV5 | ProSAP-interacting protein 1 | A2AHG0 | Kinesin-like protein KIF3A | P28741 |
| Cullin-5 | Q9D5V5 | Prostamide/prostaglandin F synthase | Q9DB60 | Kv channel-interacting protein 2 | Q9JJ69 |
| Cullin-associated NEDD8-dissociated protein 1 | Q6ZQ38 | Proteasome activator complex subunit 3 | P61290 | Lactadherin | P21956 |
| Cyclin-G-associated kinase | Q80TZ3 | Protein FAM114A2 | Q8VE88 | Latexin | P70202 |
| Cystatin-C | P21460 | Protein FAM177A1 | Q8BR63 | Lethal(2) giant larvae protein homolog 1 | Q80Y17 |
| Cysteine and glycine-rich protein 1 | P97315 | Protein lin-7 homolog C | O88952 | Leucine zipper protein 1 | Q8R4U7 |
| Cysteine desulfurase, mitochondrial | Q9Z1J3 | Protein MON2 homolog | Q80TL7 | Leucine-rich repeat and guanylate kinase domain | Q9D5S7 |
| Cysteine--tRNA ligase, cytoplasmic | Q9ER72 | Protein phosphatase 1 regulatory subunit 12B | Q8BG95 | Leucine-rich repeat flightless-interacting protein 2 | Q91WK0 |
| Cytochrome b5 | P56395 | Protein QIL1 | Q8R404 | Leucine-rich repeat LGI family member 2 | Q8K4Z0 |
| Cytochrome b-c1 complex subunit 1, mitochondr | Q9CZ13 | Pumilio homolog 1 | Q80U78 | Leucine-rich repeat-containing protein 4B | P0C192 |
| Cytochrome b-c1 complex subunit 2, mitochondr | Q9DB77 | Purine nucleoside phosphorylase | P23492 | LIM domain-containing protein 2 | Q8BGB5 |
| Cytochrome b-c1 complex subunit 6, mitochondr | P99028 | Putative ATP-dependent Clp protease proteolytic | O88696 | Lymphocyte antigen 6H | Q9WUC3 |
| Cytochrome b-c1 complex subunit 7 | Q9D855 | Rab5 GDP/GTP exchange factor | Q9JM13 | Lysophospholipase-like protein 1 | Q3UFF7 |

| | |
|---|---|
| Cytochrome c oxidase subunit 4 isoform 1, mitoc | P19783 |
| Cytoplasmic dynein 1 heavy chain 1 | Q9JHU4 |
| Cytoplasmic dynein 1 intermediate chain 1 | O88485 |
| Cytoplasmic dynein 1 intermediate chain 2 | O88487 |
| Cytoplasmic dynein 1 light intermediate chain 2 | Q6PDL0 |
| Cytoplasmic FMR1-interacting protein 2 | Q5SQX6 |
| Cytoskeleton-associated protein 4 | Q8BMK4 |
| Cytoskeleton-associated protein 5 | A2AGT5 |
| Cytosol aminopeptidase | Q9CPY7 |
| Cytosolic 10-formyltetrahydrofolate dehydrogena | Q8R0Y6 |
| Cytosolic acyl coenzyme A thioester hydrolase | Q91V12 |
| Cytosolic non-specific dipeptidase | Q9D1A2 |
| Cytospin-B | Q5SXY1 |
| Dedicator of cytokinesis protein 4 | P59764 |
| Delta-1-pyrroline-5-carboxylate dehydrogenase, i | Q8CHT0 |
| Delta-aminolevulinic acid dehydratase | P10518 |
| Dematin | Q9WV69 |
| Destrin | Q9R0P5 |
| Deubiquitinating protein VCIP135 | Q8CDG3 |
| Developmentally-regulated GTP-binding protein 2 | Q9QXB9 |
| Diacylglycerol kinase zeta | Q80UP3 |
| Dihydrolipoyl dehydrogenase, mitochondrial | O08749 |
| Dihydrolipoyllysine-residue acetyltransferase con | Q8BMF4 |
| Dihydrolipoyllysine-residue succinyltransferase c | Q9D2G2 |
| Dihydropteridine reductase | Q8BVI4 |
| Dihydropyrimidinase-related protein 1 | P97427 |
| Dihydropyrimidinase-related protein 2 | O08553 |
| Dihydropyrimidinase-related protein 3 | Q62188 |
| Dihydropyrimidinase-related protein 4 | O35098 |
| Dihydropyrimidinase-related protein 5 | Q9EQF6 |
| Dipeptidyl aminopeptidase-like protein 6 | Q9Z218 |
| Diphosphoinositol polyphosphate phosphohydrol | Q9JI46 |
| Diphosphoinositol polyphosphate phosphohydrol | P0C027 |
| Disks large homolog 1 | Q811D0 |
| Disks large homolog 2 | Q91XM9 |
| Disks large homolog 3 | P70175 |
| Disks large homolog 4 | Q62108 |
| Disks large-associated protein 1 | Q9D415 |
| DmX-like protein 2 | Q8BPN8 |
| DNA damage-binding protein 1 | Q3U1J4 |
| DnaJ homolog subfamily A member 3, mitochonc | Q99M87 |
| DnaJ homolog subfamily A member 4 | Q9JMC3 |
| DnaJ homolog subfamily B member 12 | Q9QYI4 |
| DnaJ homolog subfamily B member 6 | O54946 |
| DnaJ homolog subfamily C member 8 | Q6NZB0 |
| Dolichyl-diphosphooligosaccharide--protein glycc | Q91YQ5 |
| Drebrin | Q9QXS6 |
| Drebrin-like protein | Q62418 |
| Ras GTPase-activating protein-binding protein 2 | P97379 |
| Ras-related protein Rab-21 | P35282 |
| Redox-regulatory protein FAM213A | Q9CYH2 |
| Regulator of G-protein signaling 6 | Q9Z2H2 |
| Regulator of nonsense transcripts 1 | Q9EPU0 |
| Reticulocalbin-2 | Q8BP92 |
| Rho guanine nucleotide exchange factor 7 | Q9ES28 |
| Ribose-phosphate pyrophosphokinase 1 | Q9D7G0 |
| Ribosome maturation protein SBDS | P70122 |
| RILP-like protein 1 | Q9JJC6 |
| RWD domain-containing protein 1 | Q9CQK7 |
| S-adenosylmethionine synthase isoform type-2 | Q3THS6 |
| Sequestosome-1 | Q64337 |
| Serine/threonine-protein kinase 25 | Q9Z2W1 |
| Serine/threonine-protein kinase PRP4 homolog | Q61136 |
| Serine/threonine-protein kinase TAO1 | Q5F2E8 |
| Sickle tail protein | A2AQ25 |
| Solute carrier family 12 member 2 | P55012 |
| Sorting nexin-4 | Q91YJ2 |
| Spliceosome RNA helicase Ddx39b | Q9Z1N5 |
| Squamous cell carcinoma antigen recognized by | Q9JLI8 |
| SRSF protein kinase 2 | O54781 |
| Striatin-4 | P58404 |
| Stromal interaction molecule 2 | P83093 |
| Sulfated glycoprotein 1 | Q61207 |
| SWI/SNF complex subunit SMARCC2 | Q6PDG5 |
| Synaptic vesicle glycoprotein 2B | Q8BG39 |
| Synaptic vesicle glycoprotein 2C | Q69ZS6 |
| Syntaxin-binding protein 6 | Q8R3T5 |
| Testis-expressed sequence 2 protein | Q6ZPJ0 |
| Thioredoxin-related transmembrane protein 4 | Q8C0L0 |
| T-lymphoma invasion and metastasis-inducing pr | Q60610 |
| T-lymphoma invasion and metastasis-inducing pr | Q6ZPF3 |
| Translationally-controlled tumor protein | P63028 |
| Transthyretin | P07309 |
| tRNA methyltransferase 112 homolog | Q9DCG9 |
| Tuberin | Q61037 |
| Tubulin-specific chaperone D | Q8BYA0 |
| Tyrosine-protein kinase Fyn | P39688 |
| Ubiquitin carboxyl-terminal hydrolase 7 | Q6A4J8 |
| Ubiquitin-conjugating enzyme E2 K | P61087 |
| UBX domain-containing protein 7 | Q6P5G6 |
| UMP-CMP kinase 2, mitochondrial | Q3U5Q7 |
| UPF0505 protein C16orf62 homolog | Q8BWQ6 |
| UPF0533 protein C5orf44 homolog | Q3TIR1 |
| Vacuolar protein sorting-associated protein 45 | P97390 |
| Valacyclovir hydrolase | Q8R164 |
| Voltage-dependent L-type calcium channel subun | Q8R3Z5 |
| Lysosome-associated membrane glycoprotein 1 | P11438 |
| Macoilin | Q7TQE6 |
| Magnesium-dependent phosphatase 1 | Q9D967 |
| Malectin | Q6ZQI3 |
| Malonyl-CoA-acyl carrier protein transacylase, mi | Q8R3F5 |
| Mannose-1-phosphate guanyltransferase beta | Q8BTZ7 |
| Mannose-6-phosphate isomerase | Q924M7 |
| Mast/stem cell growth factor receptor Kit | P05532 |
| Membrane protein MLC1 | Q8VHK5 |
| Membrane-associated guanylate kinase, WW and | Q9EQJ9 |
| Metallo-beta-lactamase domain-containing protei | Q8BL86 |
| Metallothionein-3 | P28184 |
| Methylmalonyl-CoA epimerase, mitochondrial | Q9D1I5 |
| Methylthioribose-1-phosphate isomerase | Q9CQT1 |
| Microtubule-associated protein RP/EB family mer | Q6PER3 |
| Microtubule-associated proteins 1A/1B light chair | Q91VR7 |
| Microtubule-associated tumor suppressor candida | Q3UHD3 |
| Mitochondrial import inner membrane translocase | P62075 |
| Mitochondrial import inner membrane translocase | Q9CQV1 |
| Mitochondrial import receptor subunit TOM22 hor | Q9CPQ3 |
| Mitogen-activated protein kinase-binding protein 1 | Q6NS57 |
| MTSS1-like protein | Q6P9S0 |
| Myotubularin-related protein 2 | Q9Z2D1 |
| NAC-alpha domain-containing protein 1 | Q5SWP3 |
| N-acetylgalactosamine-6-sulfatase | Q571E4 |
| N-acylethanolamine-hydrolyzing acid amidase | Q9D7V9 |
| NAD-dependent protein deacetylase sirtuin-2 | Q8VDQ8 |
| NAD-dependent protein deacylase sirtuin-5, mitoc | Q8K2C6 |
| NADH dehydrogenase [ubiquinone] 1 alpha subcc | Q9JKL4 |
| NADH dehydrogenase [ubiquinone] 1 beta subco | Q9CQC7 |
| NADH dehydrogenase [ubiquinone] 1 beta subco | Q3UIU2 |
| NADH dehydrogenase [ubiquinone] 1 beta subco | Q9CR61 |
| NADH dehydrogenase [ubiquinone] 1 beta subco | Q9CQJ8 |
| Netrin receptor DCC | P70211 |
| Netrin receptor UNC5C | O08747 |
| Neudesin | Q9CQ45 |
| Neuroendocrine protein 7B2 | P12961 |
| Neurogranin | P60761 |
| Neuronal proto-oncogene tyrosine-protein kinase | P05480 |
| Neutral cholesterol ester hydrolase 1 | Q8BLF1 |
| Nexilin | Q7TPW1 |
| Nuclear distribution protein nudE homolog 1 | Q9CZA6 |
| Nuclear receptor coactivator 7 | Q6DFV7 |
| Nuclear receptor corepressor 2 | Q9WU42 |
| Nuclear ubiquitous casein and cyclin-dependent k | Q80XU3 |
| Nucleoside diphosphate kinase 3 | Q9WV85 |
| OCIA domain-containing protein 1 | Q9CRD0 |
| Oligoribonuclease, mitochondrial | Q9D8S4 |

| | |
|---|---|
| Dynactin subunit 1 | O08788 |
| Dynactin subunit 2 | Q99KJ8 |
| Dynamin-1 | P39053 |
| Dynamin-1-like protein | Q8K1M6 |
| Dynamin-3 | Q8BZ98 |
| Dynamin-like 120 kDa protein, mitochondrial | P58281 |
| Dynein light chain 1, cytoplasmic | P63168 |
| Dynein light chain 2, cytoplasmic | Q9D0M5 |
| Dystonin | Q91ZU6 |
| E3 ubiquitin-protein ligase HUWE1 | Q7TMY8 |
| E3 ubiquitin-protein ligase NEDD4 | P46935 |
| E3 ubiquitin-protein ligase UBR4 | A2AN08 |
| Early endosome antigen 1 | Q8BL66 |
| Echinoderm microtubule-associated protein-like 2 | Q7TNG5 |
| EF-hand domain-containing protein D2 | Q9D8Y0 |
| EH domain-containing protein 1 | Q9WVK4 |
| EH domain-containing protein 3 | Q9QXY6 |
| Electron transfer flavoprotein subunit alpha, mito | Q99LC5 |
| Electron transfer flavoprotein subunit beta | Q9DCW4 |
| ELKS/Rab6-interacting/CAST family member 1 | Q99MI1 |
| Elongation factor 1-alpha 1 | P10126 |
| Elongation factor 1-alpha 2 | P62631 |
| Elongation factor 1-delta | P57776 |
| Elongation factor 1-gamma | Q9D8N0 |
| Elongation factor 2 | P58252 |
| Elongation factor Tu, mitochondrial | Q8BFR5 |
| Elongator complex protein 4 | Q9ER73 |
| Endophilin-A1 | Q62420 |
| Endophilin-A2 | Q62419 |
| Endophilin-A3 | Q62421 |
| Endophilin-B2 | Q8R3V5 |
| Endoplasmic reticulum resident protein 44 | Q9D1Q6 |
| Endoplasmin | P08113 |
| Ephrin type-A receptor 4 | Q03137 |
| Epidermal growth factor receptor substrate 15-lik | Q60902 |
| Epsin-1 | Q80VP1 |
| ERC protein 2 | Q6PH08 |
| ES1 protein homolog, mitochondrial | Q9D172 |
| Ester hydrolase C11orf54 homolog | Q91V76 |
| Ethanolamine-phosphate cytidylyltransferase | Q922E4 |
| Eukaryotic initiation factor 4A-II | P10630 |
| Eukaryotic translation initiation factor 3 subunit A | P23116 |
| Eukaryotic translation initiation factor 3 subunit B | Q8JZQ9 |
| Eukaryotic translation initiation factor 3 subunit C | Q8R1B4 |
| Eukaryotic translation initiation factor 3 subunit E | P60229 |
| Eukaryotic translation initiation factor 3 subunit I | Q9QZD9 |
| Eukaryotic translation initiation factor 3 subunit J | Q66JS6 |
| Eukaryotic translation initiation factor 4 gamma 1 | Q6NZJ6 |
| WD repeat and FYVE domain-containing protein 3 | Q6VNB8 |
| WD repeat-containing protein 54 | Q9R0D8 |
| Zinc transporter 9 | Q5IRJ6 |

**Detected in experiments 2&3 - 402 proteins**

| | |
|---|---|
| [Pyruvate dehydrogenase [lipoamide]] kinase isoz | Q922H2 |
| 1,5-anhydro-D-fructose reductase | Q9DCT1 |
| 1-phosphatidylinositol 4,5-bisphosphate phospho | Q62077 |
| 2,4-dienoyl-CoA reductase, mitochondrial | Q9CQ62 |
| 26S proteasome non-ATPase regulatory subunit 9 | Q9CR00 |
| 39S ribosomal protein L9, mitochondrial | Q99N94 |
| 3-hydroxyacyl-CoA dehydrogenase type-2 | O08756 |
| 3-ketoacyl-CoA thiolase, mitochondrial | Q8BWT1 |
| 40S ribosomal protein S19 | Q9CZX8 |
| 40S ribosomal protein S2 | P25444 |
| 40S ribosomal protein S20 | P60867 |
| 40S ribosomal protein S25 | P62852 |
| 40S ribosomal protein S5 | P97461 |
| 5'-AMP-activated protein kinase catalytic subunit | Q5EG47 |
| 5-oxoprolinase | Q8K010 |
| 60S ribosomal protein L28 | P41105 |
| 60S ribosomal protein L30 | P62889 |
| 60S ribosomal protein L7 | P14148 |
| Abelson tyrosine-protein kinase 2 | Q4JIM5 |
| Abl interactor 1 | Q8CBW3 |
| Acetyl-CoA acetyltransferase, cytosolic | Q8CAY6 |
| Acetyl-CoA carboxylase 1 | Q5SWU9 |
| Acetyl-coenzyme A synthetase 2-like, mitochondr | Q99NB1 |
| Actin-related protein 3B | Q641P0 |
| Activated RNA polymerase II transcriptional coact | P11031 |
| Acyl-coenzyme A thioesterase 9, mitochondrial | Q9R0X4 |
| Acyl-protein thioesterase 1 | P97823 |
| Adenomatous polyposis coli protein | Q61315 |
| Adenylosuccinate synthetase isozyme 2 | P46664 |
| Adipocyte plasma membrane-associated protein | Q9D7N9 |
| ADP-ribosylation factor GTPase-activating protein | Q9EPJ9 |
| A-kinase anchor protein SPHKAP | Q6NSW3 |
| Alpha-2-macroglobulin | Q61838 |
| Alpha-actinin-2 | Q9JI91 |
| Aminoacyl tRNA synthase complex-interacting mu | P31230 |
| Aminopeptidase B | Q8VCT3 |
| AMP deaminase 3 | O08739 |
| Ankyrin repeat and MYND domain-containing prot | Q3TPE9 |
| Annexin A5 | P48036 |
| Anoctamin-1 | Q8BHY3 |
| AP-3 complex subunit sigma-1 | Q9DCR2 |
| Apolipoprotein O | Q9DCZ4 |
| Apoptosis inhibitor 5 | O35841 |
| Arf-GAP with coiled-coil, ANK repeat and PH dom | Q6ZQK5 |
| Opalin | Q7M750 |
| Oxysterol-binding protein-related protein 1 | Q91XL9 |
| PDZ and LIM domain protein 5 | Q8CI51 |
| Peroxiredoxin-5, mitochondrial | P99029 |
| Phosphatidylcholine transfer protein | P53808 |
| Phosphatidylinositol 4-phosphate 5-kinase type-1 | P70182 |
| Phosphoprotein associated with glycosphingolipic | Q3U1F9 |
| Phosphoribosyl pyrophosphate synthase-associa | Q8R574 |
| Phosphorylated adapter RNA export protein | Q9JJT9 |
| PI-PLC X domain-containing protein 3 | Q8BLJ3 |
| Platelet-activating factor acetylhydrolase IB subu | Q61205 |
| Potassium voltage-gated channel subfamily D me | Q9Z0V2 |
| PRA1 family protein 2 | Q9JIG8 |
| Prefoldin subunit 6 | Q03958 |
| Prenylcysteine oxidase-like | Q8C7K6 |
| Probable leucine--tRNA ligase, mitochondrial | Q8VDC0 |
| Probable phospholipid-transporting ATPase IC | Q148W0 |
| Proteasome subunit alpha type-2 | P49722 |
| Proteasome subunit beta type-2 | Q9R1P3 |
| Proteasome subunit beta type-6 | Q60692 |
| Protein canopy homolog 3 | Q9DAU1 |
| Protein CDV3 | Q4VAA2 |
| Protein DDI1 homolog 2 | A2ADY9 |
| Protein Dos | Q66L44 |
| Protein Dr1 | Q91WV0 |
| Protein FAM54B | Q9CWE0 |
| Protein kinase C-binding protein NELL2 | Q61220 |
| Protein LZIC | Q8K3C3 |
| Protein phosphatase inhibitor 2 | Q9DCL8 |
| Protein prune homolog 2 | Q52KR3 |
| Protein transport protein Sec23A | Q01405 |
| Protein tweety homolog 1 | Q9D3A9 |
| Pterin-4-alpha-carbinolamine dehydratase | P61458 |
| Rab proteins geranylgeranyltransferase compone | Q9QXG2 |
| RAC-gamma serine/threonine-protein kinase | Q9WUA6 |
| Ral GTPase-activating protein subunit alpha-2 | A3KGS3 |
| RalBP1-associated Eps domain-containing protei | O54916 |
| Ras-related GTP-binding protein B | Q6NTA4 |
| Ras-related protein Rab-33A | P97950 |
| Regulator of G-protein signaling 19 | Q9CX84 |
| Ribosomal biogenesis protein LAS1L | A2BE28 |
| Ribosomal protein S6 kinase alpha-3 | P18654 |
| Ribosomal RNA processing protein 1 homolog A | P56183 |
| RING finger protein 11 | Q9QYK7 |
| RING1 and YY1-binding protein | Q8CCI5 |
| RUN and FYVE domain-containing protein 1 | Q8BIJ7 |
| Secernin-3 | Q3TMH2 |
| Secreted frizzled-related sequence protein 4 | Q9Z1N6 |

| | | | | | |
|---|---|---|---|---|---|
| Eukaryotic translation initiation factor 4 gamma 2 | Q62448 | Arf-GAP with SH3 domain, ANK repeat and PH do | Q7SIG6 | Sel1 repeat-containing protein 1 | Q921H9 |
| Eukaryotic translation initiation factor 4 gamma 3 | Q80XI3 | Aspartoacylase | Q8R3P0 | Semaphorin-4B | Q62179 |
| Eukaryotic translation initiation factor 4B | Q8BGD9 | Atlastin-1 | Q8BH66 | Separin | P60330 |
| Eukaryotic translation initiation factor 4H | Q9WUK2 | ATP-binding cassette sub-family B member 8, mit | Q9CXJ4 | Serine protease HTRA1 | Q9R118 |
| Eukaryotic translation initiation factor 5 | P59325 | ATP-dependent (S)-NAD(P)H-hydrate dehydratas | Q9CZ42 | Serine protease HTRA2, mitochondrial | Q9JIY5 |
| Eukaryotic translation initiation factor 5A-1 | P63242 | ATP-dependent RNA helicase DDX19A | Q61655 | Serine racemase | Q9QZX7 |
| Excitatory amino acid transporter 2 | P43006 | Autophagy-related protein 2 homolog B | Q80XK6 | Serine/arginine repetitive matrix protein 1 | Q52KI8 |
| Exocyst complex component 3 | Q6KAR6 | Bcl-2-associated transcription factor 1 | Q8K019 | Serine/arginine-rich splicing factor 7 | Q8BL97 |
| Exocyst complex component 4 | O35382 | Bcl-2-like protein 13 | P59017 | Serine/threonine-protein kinase 24 | Q99KH8 |
| Exportin-2 | Q9ERK4 | BDNF/NT-3 growth factors receptor | P15209 | Serine/threonine-protein kinase 31 | Q99MW1 |
| Ezrin | P26040 | Beta-hexosaminidase subunit alpha | P29416 | Serine/threonine-protein kinase B-raf | P28028 |
| F-actin-capping protein subunit alpha-1 | P47753 | Bifunctional coenzyme A synthase | Q9DBL7 | Serine/threonine-protein kinase ULK3 | Q3U3Q1 |
| F-actin-capping protein subunit beta | P47757 | Bleomycin hydrolase | Q8R016 | Serine/threonine-protein phosphatase 4 catalytic | P97470 |
| Far upstream element-binding protein 2 | Q3U0V1 | BolA-like protein 1 | Q9D8S9 | Serine/threonine-protein phosphatase 6 regulator | Q505D1 |
| Farnesyl pyrophosphate synthase | Q920E5 | Breakpoint cluster region protein | Q6PAJ1 | Serine/threonine-protein phosphatase PP1-gamm | P63087 |
| Fascin | Q61553 | C2 domain-containing protein 2-like | Q80X80 | SH3 domain-binding glutamic acid-rich-like protei | Q91VW3 |
| Fatty acid synthase | P19096 | Cadherin-11 | P55288 | Short-chain specific acyl-CoA dehydrogenase, mi | Q07417 |
| Fatty acid-binding protein, heart | P11404 | Calcium/calmodulin-dependent protein kinase kin | Q8C078 | Small G protein signaling modulator 1 | Q8BPQ7 |
| Flotillin-1 | O08917 | Calcium-binding mitochondrial carrier protein Aral | Q8BH59 | SNW domain-containing protein 1 | Q9CSN1 |
| Flotillin-2 | Q60634 | Calpain-5 | O08688 | Sodium leak channel non-selective protein | Q8BXR5 |
| Formin-2 | Q9JL04 | cAMP-dependent protein kinase type I-beta regul | P12849 | Somatostatin | P60041 |
| Formin-binding protein 1 | Q80TY0 | cAMP-dependent protein kinase type II-alpha reg | P12367 | Sorcin | Q6P069 |
| Formin-like protein 2 | A2APV2 | Carbonic anhydrase 2 | P00920 | Sphingosine kinase 2 | Q9JIA7 |
| Fragile X mental retardation syndrome-related pr | Q61584 | Carbonyl reductase family member 4 | Q91VT4 | Sterile alpha motif domain-containing protein 14 | Q8K070 |
| Fructose-bisphosphate aldolase A | P05064 | Casein kinase II subunit alpha | Q60737 | SUMO-conjugating enzyme UBC9 | P63280 |
| Fructose-bisphosphate aldolase C | P05063 | Cation-dependent mannose-6-phosphate recepto | P24668 | Superoxide dismutase [Mn], mitochondrial | P09671 |
| FSD1-like protein | Q8BYN5 | Cdc42 effector protein 4 | Q9JM96 | Synapsin-3 | Q8JZP2 |
| Fumarate hydratase, mitochondrial | P97807 | Cell adhesion molecule 3 | Q99N28 | Synaptogyrin-3 | Q8R191 |
| G protein-regulated inducer of neurite outgrowth | Q3UNH4 | Cell cycle exit and neuronal differentiation protein | Q9JKC6 | Synaptotagmin-5 | Q9R0N5 |
| Gamma-enolase | P17183 | Chromobox protein homolog 5 | Q61686 | TATA element modulatory factor | B9EKI3 |
| Gamma-glutamyltransferase 7 | Q99JP7 | Coatomer subunit beta' | O55029 | Tektin-1 | Q9DAJ2 |
| Gamma-soluble NSF attachment protein | Q9CWZ7 | Coatomer subunit gamma-2 | Q9QXK3 | Tether containing UBX domain for GLUT4 | Q8VBT9 |
| Gamma-synuclein | Q9Z0F7 | Coiled-coil domain-containing protein 73 | Q8CDM4 | Tetratricopeptide repeat protein 1 | Q91Z38 |
| Ganglioside-induced differentiation-associated pr | O88741 | Coiled-coil domain-containing protein 92 | Q8VDN4 | Thioesterase superfamily member 4 | Q3UUI3 |
| General vesicular transport factor p115 | Q9Z1Z0 | COMM domain-containing protein 9 | Q8K2Q0 | Thioredoxin reductase 2, mitochondrial | Q9JLT4 |
| Gephyrin | Q8BUV3 | Constitutive coactivator of PPAR-gamma-like prot | Q6A0A9 | Trafficking protein particle complex subunit 2-like | Q9JME7 |
| Geranylgeranyl transferase type-2 subunit alpha | Q9JHK4 | Contactin-2 | Q61330 | Transcription factor BTF3 | Q64152 |
| Girdin | Q5SNZ0 | Contactin-associated protein-like 2 | Q9CPW0 | Trans-Golgi network integral membrane protein 1 | Q62313 |
| Glial fibrillary acidic protein | P03995 | Copine-1 | Q8C166 | Transmembrane protein 126A | Q9D8Y1 |
| Glucose 1,6-bisphosphate synthase | Q8CAA7 | Copine-4 | Q8BLR2 | Tripartite motif-containing protein 46 | Q7TNM2 |
| Glucose-6-phosphate isomerase | P06745 | Copine-8 | Q9DC53 | Tubulin beta-6 chain | Q922F4 |
| Glucosidase 2 subunit beta | O08795 | Coronin-1B | Q9WUM3 | Tubulin--tyrosine ligase-like protein 12 | Q3UDE2 |
| Glutamate [NMDA] receptor subunit epsilon-2 | Q01097 | Coronin-1C | Q9WUM4 | Tumor necrosis factor receptor superfamily meml | Q9EPU5 |
| Glutamate dehydrogenase 1, mitochondrial | P26443 | Cullin-4B | A2A432 | Ubiquilin-2 | Q9QZM0 |
| Glutamate receptor 1 | P23818 | Cyclin-dependent kinase-like 5 | Q3UTQ8 | Ubiquitin carboxyl-terminal hydrolase 4 | P35123 |
| Glutamate receptor 2 | P23819 | Cysteine sulfinic acid decarboxylase | Q9DBE0 | Ubiquitin-conjugating enzyme E2 D2 | P62838 |
| Glutamine synthetase | P15105 | Cysteine-rich protein 2 | Q9DCT8 | Ubiquitin-conjugating enzyme E2 D3 | P61079 |
| Glutaredoxin-3 | Q9CQM9 | Cytochrome b-c1 complex subunit Rieske, mitoch | Q9CR68 | Ubiquitin-like protein 4A | P21126 |

| | | | | | |
|---|---|---|---|---|---|
| Glutathione S-transferase Mu 1 | P10649 | Cytochrome c oxidase subunit 5A, mitochondrial | P12787 | Ubiquitin-like-conjugating enzyme ATG3 | Q9CPX6 |
| Glutathione S-transferase P 1 | P19157 | Cytochrome c oxidase subunit 6C | Q9CPQ1 | Ubiquitin-protein ligase E3C | Q80U95 |
| Glutathione synthetase | P51855 | Cytoplasmic dynein 1 light intermediate chain 1 | Q8R1Q8 | UBX domain-containing protein 2B | Q0KL01 |
| Glyceraldehyde-3-phosphate dehydrogenase | P16858 | Cytoplasmic FMR1-interacting protein 1 | Q7TMB8 | Uncharacterized protein C12orf43 homolog | Q3UY34 |
| Glycerol-3-phosphate dehydrogenase [NAD(+)], | P13707 | Cytoplasmic protein NCK2 | O55033 | Unconventional myosin-XV | Q9QZZ4 |
| Glycerol-3-phosphate dehydrogenase 1-like prot | Q3ULJ0 | D-3-phosphoglycerate dehydrogenase | Q61753 | Unconventionnal myosin-X | F8VQB6 |
| Glycerol-3-phosphate dehydrogenase, mitochon | Q64521 | DCC-interacting protein 13-beta | Q8K3G9 | UPF0184 protein C9orf16 homolog | P58686 |
| Glycine dehydrogenase [decarboxylating], mitoc | Q91W43 | D-dopachrome decarboxylase | O35215 | UPF0498 protein KIAA1191 | Q9DBN4 |
| Glycine--tRNA ligase | Q9CZD3 | Dendrin | Q80TS7 | Vacuolar protein sorting-associated protein 18 ho | Q8R307 |
| Glycogen phosphorylase, brain form | Q8CI94 | Developmentally-regulated GTP-binding protein 1 | P32233 | Vacuolar protein sorting-associated protein VTA1 | Q9CR26 |
| Glycogen phosphorylase, muscle form | Q9WUB3 | Diacylglycerol kinase beta | Q6NS52 | Vesicle-associated membrane protein 3 | P63024 |
| Glycogen synthase kinase-3 alpha | Q2NL51 | Diacylglycerol kinase gamma | Q91WG7 | Vesicular glutamate transporter 1 | Q3TXX4 |
| Glyoxalase domain-containing protein 4 | Q9CPV4 | Disintegrin and metalloproteinase domain-contain | Q9R1V6 | von Willebrand factor A domain-containing protei | Q99KC8 |
| Glyoxylate reductase/hydroxypyruvate reductase | Q91Z53 | Disintegrin and metalloproteinase domain-contain | Q9R1V7 | Xin actin-binding repeat-containing protein 2 | Q4U4S6 |
| Glypican-1 | Q9QZF2 | Disks large-associated protein 2 | Q8BJ42 | Zinc finger CCCH domain-containing protein 15 | Q3TIV5 |
| Glypican-4 | P51655 | Disks large-associated protein 4 | B1AZP2 | Zinc finger protein 40 | Q03172 |
| GMP reductase 1 | Q9DCZ1 | DNA-directed RNA polymerases I, II, and III subu | Q923G2 | Zinc finger ZZ-type and EF-hand domain-containi | Q5SSH7 |
| GMP synthase [glutamine-hydrolyzing] | Q3THK7 | DnaJ homolog subfamily A member 2 | Q9QYJ0 | Zinc transporter 3 | P97441 |
| Golgi apparatus protein 1 | Q61543 | DnaJ homolog subfamily B member 11 | Q99KV1 | Zinc transporter ZIP10 | Q6P5F6 |
| Golgin subfamily A member 2 | Q921M4 | DnaJ homolog subfamily B member 2 | Q9QYI5 | **Detected in experiment 3 only - 733 proteins** | |
| Golgin subfamily A member 3 | P55937 | DnaJ homolog subfamily B member 4 | Q9D832 | [Protein ADP-ribosylarginine] hydrolase | P54923 |
| GRIP1-associated protein 1 | Q8VD04 | Dolichyl-diphosphooligosaccharide--protein glyco | Q9DBG6 | [Pyruvate dehydrogenase [acetyl-transferring]]-p | Q3UV70 |
| Growth factor receptor-bound protein 2 | Q60631 | Dual specificity mitogen-activated protein kinase | P70236 | [Pyruvate dehydrogenase [lipoamide]] kinase iso | Q8BFP9 |
| GrpE protein homolog 1, mitochondrial | Q99LP6 | Dynactin subunit 4 | Q8CBY8 | 1-acyl-sn-glycerol-3-phosphate acyltransferase e | Q9D1E8 |
| GTP:AMP phosphotransferase, mitochondrial | Q9WTP7 | Dystrobrevin alpha | Q9D2N4 | 1-acyl-sn-glycerol-3-phosphate acyltransferase g | Q9D517 |
| GTP-binding protein Di-Ras2 | Q5PR73 | E3 ubiquitin-protein ligase TRIM9 | Q8C7M3 | 1-phosphatidylinositol 3-phosphate 5-kinase | Q9Z1T6 |
| GTP-binding protein Rheb | Q921J2 | ELAV-like protein 1 | P70372 | 26S protease regulatory subunit 6B | P54775 |
| Guanine deaminase | Q9R111 | Electrogenic sodium bicarbonate cotransporter 1 | O88343 | 26S proteasome non-ATPase regulatory subunit | O35226 |
| Guanine nucleotide-binding protein G(i) subunit | B2RSH2 | Electron transfer flavoprotein-ubiquinone oxidore | Q921G7 | 28S ribosomal protein S22, mitochondrial | Q9CXW2 |
| Guanine nucleotide-binding protein G(i) subunit | P08752 | Elongator complex protein 2 | Q91WG4 | 28S ribosomal protein S29, mitochondrial | Q9ER88 |
| Guanine nucleotide-binding protein G(k) subunit | Q9DC51 | Emerin | O08579 | 28S ribosomal protein S35, mitochondrial | Q8BJZ4 |
| Guanine nucleotide-binding protein G(o) subunit | P18872 | Engulfment and cell motility protein 2 | Q8BHL5 | 28S ribosomal protein S9, mitochondrial | Q9D7N3 |
| Guanine nucleotide-binding protein G(q) subunit | P21279 | Enoyl-CoA delta isomerase 1, mitochondrial | P42125 | 2-oxoisovalerate dehydrogenase subunit alpha, n | P50136 |
| Guanine nucleotide-binding protein subunit beta- | P68040 | Enoyl-CoA hydratase, mitochondrial | Q8BH95 | 39S ribosomal protein L27, mitochondrial | Q99N92 |
| Guanylate cyclase soluble subunit beta-1 | O54865 | Ensconsin | O88735 | 39S ribosomal protein L37, mitochondrial | Q921S7 |
| Heat shock 70 kDa protein 12A | Q8K0U4 | Epoxide hydrolase 2 | P34914 | 39S ribosomal protein L38, mitochondrial | Q8K2M0 |
| Heat shock 70 kDa protein 1B | P17879 | Erlin-2 | Q8BFZ9 | 3-hydroxyisobutyryl-CoA hydrolase, mitochondria | Q8QZS1 |
| Heat shock 70 kDa protein 4 | Q61316 | Ermin | Q5EBJ4 | 40S ribosomal protein S12 | P63323 |
| Heat shock 70 kDa protein 4L | P48722 | Eukaryotic translation initiation factor 2 subunit 1 | Q6ZWX6 | 45 kDa calcium-binding protein | Q61112 |
| Heat shock cognate 71 kDa protein | P63017 | Eukaryotic translation initiation factor 2 subunit 3, | Q9Z0N1 | 5'(3')-deoxyribonucleotidase, cytosolic type | Q9JM14 |
| Heat shock protein 105 kDa | Q61699 | Eukaryotic translation initiation factor 3 subunit H | Q91WK2 | 55 kDa erythrocyte membrane protein | P70290 |
| Heat shock protein 75 kDa, mitochondrial | Q9CQN1 | Eukaryotic translation initiation factor 3 subunit K | Q9DBZ5 | 5'-AMP-activated protein kinase subunit beta-2 | Q6PAM0 |
| Heat shock protein HSP 90-alpha | P07901 | Eukaryotic translation initiation factor 3 subunit L | Q8QZY1 | 5'-AMP-activated protein kinase subunit gamma- | Q91WG5 |
| Heat shock protein HSP 90-beta | P11499 | Eukaryotic translation initiation factor 5B | Q05D44 | 5-azacytidine-induced protein 1 | Q62036 |
| Heat shock-related 70 kDa protein 2 | P17156 | Exocyst complex component 5 | Q3TPX4 | 60S acidic ribosomal protein P0 | P14869 |
| Heme oxygenase 2 | O70252 | Exocyst complex component 7 | O35250 | 60S ribosomal protein L10a | P53026 |
| Heme-binding protein 1 | Q9R257 | F-actin-capping protein subunit alpha-2 | P47754 | 60S ribosomal protein L10-like | P86048 |
| Hepatoma-derived growth factor | P51859 | Far upstream element-binding protein 1 | Q91WJ8 | 60S ribosomal protein L13a | P19253 |

| | | | | | |
|---|---|---|---|---|---|
| Heterochromatin protein 1-binding protein 3 | Q3TEA8 | Fatty aldehyde dehydrogenase | P47740 | 60S ribosomal protein L15 | Q9CZM2 |
| Heterogeneous nuclear ribonucleoprotein A/B | Q99020 | F-box only protein 41 | Q6NS60 | 60S ribosomal protein L24 | Q8BP67 |
| Heterogeneous nuclear ribonucleoprotein D0 | Q60668 | Fibronectin type III and SPRY domain-containing | Q7TPM6 | 60S ribosomal protein L27 | P61358 |
| Heterogeneous nuclear ribonucleoprotein K | P61979 | Fibrous sheath-interacting protein 2 | A2ARZ3 | 60S ribosomal protein L34 | Q9D1R9 |
| Heterogeneous nuclear ribonucleoprotein Q | Q7TMK9 | Focal adhesion kinase 1 | P34152 | 60S ribosomal protein L4 | Q9D8E6 |
| Heterogeneous nuclear ribonucleoprotein U | Q8VEK3 | Formin-like protein 1 | Q9JL26 | 60S ribosomal protein L7a | P12970 |
| Hexokinase-1 | P17710 | Galactokinase | Q9R0N0 | 60S ribosomal protein L9 | P51410 |
| Histidine--tRNA ligase, cytoplasmic | Q61035 | Gamma-adducin | Q9QYB5 | 6-phosphofructokinase, liver type | P12382 |
| Histone H3.1 | P68433 | Gamma-aminobutyric acid receptor subunit alpha | P62812 | Abhydrolase domain-containing protein 10, mitoc | Q6PE15 |
| Histone-lysine N-methyltransferase SETD7 | Q8VHL1 | Gamma-aminobutyric acid receptor subunit beta- | P63080 | Abl interactor 2 | P62484 |
| Homer protein homolog 1 | Q9Z2Y3 | Gamma-aminobutyric acid type B receptor subuni | Q80T41 | Acetyl-coenzyme A synthetase, cytoplasmic | Q9QXG4 |
| Homer protein homolog 3 | Q99JP6 | Ganglioside-induced differentiation-associated pr | Q8VE33 | Actin, cytoplasmic 2 | P63260 |
| Hsc70-interacting protein | Q99L47 | General transcription factor IIF subunit 1 | Q3THK3 | Actin-binding LIM protein 3 | Q69ZX8 |
| Hsp90 co-chaperone Cdc37 | Q61081 | Glucosamine-6-phosphate isomerase 1 | O88958 | Actin-like protein 6A | Q9Z2N8 |
| Huntingtin | P42859 | Glucose-6-phosphate 1-dehydrogenase X | Q00612 | Actin-like protein 6B | Q99MR0 |
| Hyaluronan and proteoglycan link protein 1 | Q9QUP5 | Glutamate [NMDA] receptor subunit epsilon-1 | P35436 | Actin-related protein 10 | Q9QZB7 |
| Hydroxyacylglutathione hydrolase, mitochondrial | Q99KB8 | Glutamate receptor 4 | Q9Z2W8 | Actin-related protein 2/3 complex subunit 1B | Q9WV32 |
| Hypoxanthine-guanine phosphoribosyltransferas | P00493 | Glutathione S-transferase Mu 5 | P48774 | Acylglycerol kinase, mitochondrial | Q9ESW4 |
| Immunity-related GTPase family Q protein | Q8VIM9 | Glycerophosphocholine phosphodiesterase GPCI | Q8C0L9 | ADP-ribosylation factor 4 | P61750 |
| Importin subunit alpha-3 | O35344 | Glycolipid transfer protein | Q9JL62 | ADP-ribosylation factor GTPase-activating protei | Q99K28 |
| Importin subunit beta-1 | P70168 | Glycylpeptide N-tetradecanoyltransferase 1 | O70310 | ADP-ribosylation factor-binding protein GGA1 | Q8R0H9 |
| Importin-7 | Q9EPL8 | Glypican-5 | Q8CAL5 | ADP-ribosylation factor-binding protein GGA3 | Q8BMI3 |
| Inactive dipeptidyl peptidase 10 | Q6NXK7 | Golgi reassembly-stacking protein 2 | Q99JX3 | ADP-ribosylation factor-like protein 15 | Q8BGR6 |
| Inactive phospholipase C-like protein 2 | Q8K394 | Golgi resident protein GCP60 | Q8BMP6 | A-kinase anchor protein 10, mitochondrial | O88845 |
| Inorganic pyrophosphatase | Q9D819 | GRIP and coiled-coil domain-containing protein 2 | Q8CHG3 | A-kinase anchor protein 12 | Q9WTQ5 |
| Inorganic pyrophosphatase 2, mitochondrial | Q91VM9 | GTPase NRas | P08556 | Aldehyde dehydrogenase family 3 member B1 | Q80VQ0 |
| Inosine-5'-monophosphate dehydrogenase 2 | P24547 | GTP-binding nuclear protein Ran | P62827 | Aldehyde oxidase | O54754 |
| Inositol 1,4,5-trisphosphate receptor type 1 | P11881 | Guanine nucleotide-binding protein G(I)/G(S)/G(T | P62874 | Alkyldihydroxyacetonephosphate synthase, perox | Q8C0I1 |
| Inositol monophosphatase 1 | O55023 | Guanine nucleotide-binding protein G(s) subunit a | Q6R0H7 | Alpha-1-antitrypsin 1-4 | Q00897 |
| Inositol-trisphosphate 3-kinase A | Q8R071 | Guanine nucleotide-binding protein G(z) subunit a | O70443 | Alpha-1-syntrophin | Q61234 |
| Insulin-degrading enzyme | Q9JHR7 | Guanine nucleotide-binding protein subunit alpha- | P21278 | Alpha-mannosidase 2C1 | Q91W89 |
| Integrin alpha-V | P43406 | Guanine nucleotide-binding protein subunit alpha- | P27601 | Alpha-synuclein | O55042 |
| Intercellular adhesion molecule 5 | Q60625 | Guanine nucleotide-binding protein subunit beta-4 | P29387 | Alpha-tubulin N-acetyltransferase | Q8K341 |
| Intracellular hyaluronan-binding protein 4 | Q9JKS5 | Guanine nucleotide-binding protein subunit beta-5 | P62881 | Aminoacyl tRNA synthase complex-interacting mu | Q8R010 |
| IQ motif and SEC7 domain-containing protein 1 | Q8R0S2 | HBS1-like protein | Q69ZS7 | Amyloid beta A4 precursor protein-binding family | P98084 |
| IQ motif and SEC7 domain-containing protein 2 | Q5DU25 | Hemoglobin subunit alpha | P01942 | Ankyrin repeat and FYVE domain-containing prot | Q810B6 |
| IQ motif and SEC7 domain-containing protein 3 | Q3TES0 | Heterogeneous nuclear ribonucleoprotein U-like p | Q00PI9 | Ankyrin repeat and SAM domain-containing prote | P59672 |
| Iron-sulfur cluster assembly enzyme ISCU, mitoc | Q9D7P6 | Heterogeneous nuclear ribonucleoproteins A2/B1 | O88569 | Ankyrin-1 | Q02357 |
| Islet cell autoantigen 1 | P97411 | Heterogeneous nuclear ribonucleoproteins C1/C2 | Q9Z204 | Anoctamin-4 | Q8C5H1 |
| Isocitrate dehydrogenase [NAD] subunit alpha, m | Q9D6R2 | Hippocalcin-like protein 4 | Q8BGZ1 | AP-1 complex subunit mu-1 | P35585 |
| Isocitrate dehydrogenase [NADP] cytoplasmic | O88844 | Histone H2B type 1-M | P10854 | AP-3 complex subunit beta-1 | Q9Z1T1 |
| Isocitrate dehydrogenase [NADP], mitochondrial | P54071 | Histone H4 | P62806 | Apolipoprotein A-II | P09813 |
| Isoleucine--tRNA ligase, cytoplasmic | Q8BU30 | Huntingtin-interacting protein 1-related protein | Q9JKY5 | Apolipoprotein O-like | Q78IK4 |
| Isovaleryl-CoA dehydrogenase, mitochondrial | Q9JHI5 | Hyaluronan and proteoglycan link protein 4 | Q80WM4 | Apoptosis regulator BAX | Q07813 |
| Janus kinase and microtubule-interacting protein | Q5DTN8 | Hypoxia up-regulated protein 1 | Q9JKR6 | Apoptotic chromatin condensation inducer in the | Q9JIX8 |
| Junctional adhesion molecule C | Q9D8B7 | Immunoglobulin superfamily member 8 | Q8R366 | Arf-GAP domain and FG repeat-containing protei | Q80WC7 |
| Kalirin | A2CG49 | Importin-9 | Q91YE6 | Arginyl-tRNA--protein transferase 1 | Q9Z2A5 |
| Kelch repeat and BTB domain-containing protein | Q8BNW9 | Intersectin-1 | Q9Z0R4 | Armadillo repeat protein deleted in velo-cardio-fa | P98203 |

| | | | | | |
|---|---|---|---|---|---|
| Kinase suppressor of Ras 1 | Q61097 | Intersectin-2 | Q9Z0R6 | Armadillo repeat-containing protein 1 | Q9D7A8 |
| Kinectin | Q61595 | Isocitrate dehydrogenase [NAD] subunit gamma 1 | P70404 | Armadillo repeat-containing protein 8 | Q9DBR3 |
| Kinesin heavy chain isoform 5C | P28738 | Junctophilin-3 | Q9ET77 | Aromatic-L-amino-acid decarboxylase | O88533 |
| Kinesin light chain 1 | O88447 | Kinesin-like protein KIF21A | Q9QXL2 | Arylsulfatase B | P50429 |
| Kinesin light chain 2 | O88448 | Kinesin-like protein KIF3B | Q61771 | ATP-binding cassette sub-family B member 7, mi | Q61102 |
| Kinesin-1 heavy chain | Q61768 | Kininogen-1 | O08677 | ATP-binding cassette sub-family D member 2 | Q61285 |
| Kinesin-like protein KIF1A | P33173 | Kv channel-interacting protein 4 | Q6PHZ8 | ATP-binding cassette sub-family D member 3 | P55096 |
| Kinesin-like protein KIF2A | P28740 | L-2-hydroxyglutarate dehydrogenase, mitochondr | Q91YP0 | ATP-binding cassette sub-family F member 2 | Q99LE6 |
| Kinesin-like protein KIF3C | O35066 | Lactoylglutathione lyase | Q9CPU0 | ATP-binding cassette sub-family F member 3 | Q8K268 |
| LanC-like protein 2 | Q9JJK2 | LanC-like protein 1 | O89112 | ATP-dependent RNA helicase A | O70133 |
| La-related protein 1 | Q6ZQ58 | Large proline-rich protein BAG6 | Q9Z1R2 | ATP-dependent RNA helicase DDX3Y | Q62095 |
| L-asparaginase | Q8C0M9 | Leucine-rich repeat-containing protein 7 | Q80TE7 | ATP-dependent RNA helicase DDX42 | Q810A7 |
| Latrophilin-1 | Q80TR1 | Lipoamide acyltransferase component of branche | P53395 | ATP-dependent RNA helicase Dhx29 | Q6PGC1 |
| Latrophilin-3 | Q80TS3 | Lysophosphatidic acid phosphatase type 6 | Q8BP40 | AT-rich interactive domain-containing protein 1A | A2BH40 |
| LDLR chaperone MESD | Q9ERE7 | MAGUK p55 subfamily member 2 | Q9WV34 | Baculoviral IAP repeat-containing protein 2 | Q62210 |
| LETM1 and EF-hand domain-containing protein 1 | Q9Z2I0 | MAGUK p55 subfamily member 5 | Q9JLB2 | BAG family molecular chaperone regulator 4 | Q8CI61 |
| Leucine zipper transcription factor-like protein 1 | Q9JHQ5 | Matrin-3 | Q8K310 | Barrier-to-autointegration factor | O54962 |
| Leucine-rich glioma-inactivated protein 1 | Q9JIA1 | Megakaryocyte-associated tyrosine-protein kinase | P41242 | Basic leucine zipper and W2 domain-containing p | Q2L4X1 |
| Leucine-rich PPR motif-containing protein, mitocl | Q6PB66 | Membrane-associated phosphatidylinositol transfe | O35954 | B-cell receptor-associated protein 29 | Q61334 |
| Leucine-rich repeat and immunoglobulin-like dom | Q9D1T0 | Membrane-associated phosphatidylinositol transfe | Q6ZPQ6 | Beta-1-syntrophin | Q99L88 |
| Leucine-rich repeat-containing protein 16C | Q3V3V9 | Methionine adenosyltransferase 2 subunit beta | Q99LB6 | Beta-arrestin-2 | Q91YI4 |
| Leucine-rich repeat-containing protein 59 | Q922Q8 | Methyl-CpG-binding protein 2 | Q9Z2D6 | Brain-specific angiogenesis inhibitor 2 | Q8CGM1 |
| Leucine--tRNA ligase, cytoplasmic | Q8BMJ2 | Methylcrotonoyl-CoA carboxylase beta chain, mito | Q3ULD5 | Breast cancer anti-estrogen resistance protein 1 | Q61140 |
| Leukocyte surface antigen CD47 | Q61735 | Methylglutaconyl-CoA hydratase, mitochondrial | Q9JLZ3 | Breast carcinoma-amplified sequence 3 homolog | Q8CCN5 |
| Leukotriene A-4 hydrolase | P24527 | Microtubule-associated serine/threonine-protein k | Q9R1L5 | Bromodomain-containing protein 2 | Q7JJ13 |
| LIM and calponin homology domains-containing p | Q3UH68 | Microtubule-associated serine/threonine-protein k | Q3U214 | Bromodomain-containing protein 4 | Q9ESU6 |
| LIM and SH3 domain protein 1 | Q61792 | Mitochondrial 2-oxoglutarate/malate carrier protei | Q9CR62 | BTB/POZ domain-containing protein KCTD16 | Q5DTY9 |
| Limbic system-associated membrane protein | Q8BLK3 | Mitochondrial chaperone BCS1 | Q9CZP5 | BTB/POZ domain-containing protein KCTD4 | Q9D7X1 |
| Lipid phosphate phosphatase-related protein type | Q7TME0 | Mitochondrial glutamate carrier 1 | Q9D6M3 | CAAX prenyl protease 1 homolog | Q80W54 |
| Liprin-alpha-2 | Q8BSS9 | Mitochondrial-processing peptidase subunit beta | Q9CXT8 | Cadherin-2 | P15116 |
| Liprin-alpha-3 | P60469 | Mitofusin-2 | Q80U63 | Cadherin-8 | P97291 |
| LisH domain and HEAT repeat-containing protein | Q148V7 | Monoacylglycerol lipase ABHD12 | Q99LR1 | Cadherin-9 | P70407 |
| L-lactate dehydrogenase A chain | P06151 | Monocarboxylate transporter 1 | P53986 | Calcium uptake protein 1, mitochondrial | Q8VCX5 |
| L-lactate dehydrogenase B chain | P16125 | Myelin-associated glycoprotein | P20917 | Calcium/calmodulin-dependent protein kinase typ | Q6PHZ2 |
| Lon protease homolog, mitochondrial | Q8CGK3 | Myosin-11 | O08638 | Calcium-activated potassium channel subunit alp | Q08460 |
| Long-chain specific acyl-CoA dehydrogenase, mi | P51174 | Myotubularin-related protein 1 | Q9Z2C4 | Calcium-binding mitochondrial carrier protein SCa | A2ASZ8 |
| Long-chain-fatty-acid--CoA ligase 6 | Q91WC3 | NAD kinase domain-containing protein 1 | Q8C5H8 | Calcium-binding protein 39 | Q06138 |
| Long-chain-fatty-acid--CoA ligase ACSBG1 | Q99PU5 | NADH dehydrogenase [ubiquinone] 1 alpha subco | Q9ERS2 | Calcium-independent phospholipase A2-gamma | Q8K1N1 |
| Low molecular weight phosphotyrosine protein ph | Q9D358 | NADH dehydrogenase [ubiquinone] 1 alpha subco | Q9Z1P6 | Calcium-transporting ATPase type 2C member 1 | Q80XR2 |
| Lupus La protein homolog | P32067 | NADH dehydrogenase [ubiquinone] 1 alpha subco | Q9DC69 | Calmodulin-regulated spectrin-associated protein | Q80VC9 |
| Lysine--tRNA ligase | Q99MN1 | NADH dehydrogenase [ubiquinone] 1 beta subcom | Q9DCS9 | Calponin-1 | Q08091 |
| Lysosomal alpha-glucosidase | P70699 | NADH-cytochrome b5 reductase 1 | Q9DB73 | cAMP-dependent protein kinase catalytic subunit | P05132 |
| MAGUK p55 subfamily member 6 | Q9JLB0 | NADPH--cytochrome P450 reductase | P37040 | cAMP-dependent protein kinase type II-beta regu | P31324 |
| Major prion protein | P04925 | Netrin receptor UNC5A | Q8K1S4 | cAMP-specific 3',5'-cyclic phosphodiesterase 4A | O89084 |
| Malate dehydrogenase, cytoplasmic | P14152 | Neuronal calcium sensor 1 | Q8BNY6 | Cap-specific mRNA (nucleoside-2'-O-)-methyltrar | Q9DBC3 |
| Malate dehydrogenase, mitochondrial | P08249 | Neuronal growth regulator 1 | Q80Z24 | Carbonic anhydrase 7 | Q9ERQ8 |
| Mammalian ependymin-related protein 1 | Q99M71 | Neuropilin-2 | O35375 | Carnitine O-palmitoyltransferase 1, brain isoform | Q8BGD5 |
| MAP kinase-activating death domain protein | Q80U28 | Neutral amino acid transporter A | O35874 | Caseinolytic peptidase B protein homolog | Q60649 |

| | | | | | |
|---|---|---|---|---|---|
| MAP7 domain-containing protein 1 | A2AJI0 | NHS-like protein 2 | B1AXH1 | Catalase | P24270 |
| MAP7 domain-containing protein 2 | A2AG50 | Nicotinamide phosphoribosyltransferase | Q99KQ4 | Catenin delta-1 | P30999 |
| MARCKS-related protein | P28667 | Nitrilase homolog 1 | Q8VDK1 | Cation-independent mannose-6-phosphate recep | Q07113 |
| Medium-chain specific acyl-CoA dehydrogenase, | P45952 | Nodal modulator 1 | Q6GQT9 | CCR4-NOT transcription complex subunit 2 | Q8C5L3 |
| Membrane-associated guanylate kinase, WW an | Q9WVQ1 | Noelin | O88998 | CDGSH iron-sulfur domain-containing protein 1 | Q91WS0 |
| Membrane-associated progesterone receptor co | O55022 | Non-POU domain-containing octamer-binding pro | Q99K48 | Cell adhesion molecule 4 | Q8R464 |
| Membrane-associated progesterone receptor co | Q80UU9 | Nuclear export mediator factor Nemf | Q8CCP0 | Cell differentiation protein RCD1 homolog | Q9JKY0 |
| Metabotropic glutamate receptor 2 | Q14BI2 | Nucleophosmin | Q61937 | Cell division cycle 5-like protein | Q6A068 |
| Metabotropic glutamate receptor 3 | Q9QYS2 | Nucleoside diphosphate kinase A | P15532 | Cell division cycle and apoptosis regulator protein | Q8CH18 |
| Metabotropic glutamate receptor 5 | Q3UVX5 | Omega-amidase NIT2 | Q9JHW2 | Centromere protein V | Q9CXS4 |
| Metaxin-1 | P47802 | Ornithine aminotransferase, mitochondrial | P29758 | Chaperone activity of bc1 complex-like, mitochon | Q60936 |
| Methionine--tRNA ligase, cytoplasmic | Q68FL6 | Palmitoyltransferase ZDHHC5 | Q8VDZ4 | Charged multivesicular body protein 3 | Q9CQ10 |
| Methylcrotonoyl-CoA carboxylase subunit alpha, | Q99MR8 | Phenylalanine--tRNA ligase beta subunit | Q9WUA2 | Charged multivesicular body protein 7 | Q8R1T1 |
| Methylmalonate-semialdehyde dehydrogenase [a | Q9EQ20 | Phosphatase and actin regulator 3 | Q8BYK5 | Chitinase domain-containing protein 1 | Q922Q9 |
| Methylmalonyl-CoA mutase, mitochondrial | P16332 | Phosphatidate cytidylyltransferase 2 | Q99L43 | Chloride channel protein 2 | Q9R0A1 |
| Microtubule-actin cross-linking factor 1 | Q9QXZ0 | Phosphatidylinositol 3-kinase regulatory subunit b | O08908 | Chloride intracellular channel protein 4 | Q9QYB1 |
| Microtubule-associated protein 1A | Q9QYR6 | Phosphatidylinositol 5-phosphate 4-kinase type-2 | Q91XU3 | Cholesterol 24-hydroxylase | Q9WVK8 |
| Microtubule-associated protein 1B | P14873 | Phosphoenolpyruvate carboxykinase [GTP], mito | Q8BH04 | Choline O-acetyltransferase | Q03059 |
| Microtubule-associated protein 2 | P20357 | Phytanoyl-CoA hydroxylase-interacting protein | Q8K0S0 | Chromodomain-helicase-DNA-binding protein 4 | Q6PDQ2 |
| Microtubule-associated protein 4 | P27546 | Plexin-B1 | Q8CJH3 | Chromosome alignment-maintaining phosphoprot | Q8K327 |
| Microtubule-associated protein 6 | Q7TSJ2 | Polypeptide N-acetylgalactosaminyltransferase 2 | Q6PB93 | Citron Rho-interacting kinase | P49025 |
| Microtubule-associated protein tau | P10637 | Potassium/sodium hyperpolarization-activated cy | O88703 | Claudin domain-containing protein 1 | Q9CQX5 |
| Misshapen-like kinase 1 | Q9JM52 | Pre-mRNA-processing factor 19 | Q99KP6 | Clavesin-1 | Q9D4C9 |
| Mitochondrial glutamate carrier 2 | Q9DB41 | Prickle-like protein 2 | Q80Y24 | Cleft lip and palate transmembrane protein 1 hom | Q8VBZ3 |
| Mitochondrial import inner membrane transloca | O35857 | Probable E3 ubiquitin-protein ligase MYCBP2 | Q7TPH6 | Coagulation factor VIII | Q06194 |
| Mitochondrial import inner membrane transloca | Q9D880 | Probable G-protein coupled receptor 158 | Q8C419 | Coatomer subunit beta | Q9JIF7 |
| Mitochondrial import receptor subunit TOM70 | Q9CZW5 | Programmed cell death protein 10 | Q8VE70 | Coatomer subunit delta | Q5XJY5 |
| Mitochondrial inner membrane protein | Q8CAQ8 | Programmed cell death protein 5 | P56812 | Coatomer subunit gamma-1 | Q9QZE5 |
| Mitogen-activated protein kinase 1 | P63085 | Propionyl-CoA carboxylase beta chain, mitochon | Q99MN9 | Cob(I)yrinic acid a,c-diamide adenosyltransferase | Q9D273 |
| Mitogen-activated protein kinase 3 | Q63844 | Proteasome subunit alpha type-5 | Q9Z2U1 | Coiled-coil domain-containing protein 22 | Q9JIG7 |
| MKL/myocardin-like protein 2 | P59759 | Proteasome subunit beta type-4 | P99026 | Coiled-coil domain-containing protein 90B, mitoch | Q8C3X2 |
| MOB-like protein phocein | Q6PEB6 | Protein EFR3 homolog A | Q8BG67 | Collagen alpha-1(VI) chain | Q04857 |
| Monoglyceride lipase | O35678 | Protein enabled homolog | Q03173 | COMM domain-containing protein 3 | Q63829 |
| Myc box-dependent-interacting protein 1 | O08539 | Protein FAM126B | Q8C729 | Complement C3 | P01027 |
| Myelin basic protein | P04370 | Protein FAM169A | Q5XG69 | Condensin-2 complex subunit H2 | Q8BSP2 |
| Myelin-oligodendrocyte glycoprotein | Q61885 | Protein FAM98B | Q80VD1 | Constitutive coactivator of peroxisome proliferato | Q6RI63 |
| Myosin light polypeptide 6 | Q60605 | Protein fat-free homolog | Q3UVL4 | COP9 signalosome complex subunit 6 | O88545 |
| Myosin-10 | Q61879 | Protein flightless-1 homolog | Q9JJ28 | Copine-5 | Q8JZW4 |
| Myosin-14 | Q6URW6 | Protein NipSnap homolog 2 | O55126 | Coronin-2B | Q8BH44 |
| Myosin-9 | Q8VDD5 | Protein TANC2 | A2A690 | Craniofacial development protein 1 | O88271 |
| Myotrophin | P62774 | Protein TSSC1 | Q8K0G5 | C-terminal-binding protein 1 | O88712 |
| Myotubularin-related protein 5 | Q6ZPE2 | Protein VAC14 homolog | Q80WQ2 | CTP synthase 1 | P70698 |
| Myristoylated alanine-rich C-kinase substrate | P26645 | Protein-glutamine gamma-glutamyltransferase 2 | P21981 | Cullin-1 | Q9WTX6 |
| N(G),N(G)-dimethylarginine dimethylaminohydrol | Q9CWS0 | Protein-L-isoaspartate(D-aspartate) O-methyltran | P23506 | Cullin-2 | Q9D4H8 |
| N(G),N(G)-dimethylarginine dimethylaminohydrol | Q99LD8 | Putative phospholipase B-like 2 | Q3TCN2 | Cullin-4A | Q3TCH7 |
| Na(+)/H(+) exchange regulatory cofactor NHE-RI | P70441 | Putative pre-mRNA-splicing factor ATP-depender | O35286 | Cyclin-dependent kinase 13 | Q69ZA1 |
| NADH dehydrogenase [ubiquinone] 1 alpha subc | Q7TMF3 | Putative RNA-binding protein Luc7-like 1 | Q9CYI4 | Cyclin-dependent kinase 16 | Q04735 |
| NADH dehydrogenase [ubiquinone] 1 alpha subc | Q9CQ75 | Putative tyrosine-protein phosphatase auxilin | Q80TZ3 | Cyclin-dependent kinase 17 | Q8K0D0 |

| | | | | | |
|---|---|---|---|---|---|
| NADH dehydrogenase [ubiquinone] 1 alpha subc | Q9CPP6 | Pyridoxine-5'-phosphate oxidase | Q91XF0 | Cyclin-dependent kinase 5 | P49615 |
| NADH dehydrogenase [ubiquinone] 1 beta subcc | Q9D6J5 | Rab11 family-interacting protein 5 | Q8R361 | Cystathionine gamma-lyase | Q8VCN5 |
| NADH dehydrogenase [ubiquinone] flavoprotein | Q91YT0 | RalA-binding protein 1 | Q62172 | Cysteine and histidine-rich domain-containing prc | Q9D1P4 |
| NADH dehydrogenase [ubiquinone] flavoprotein | Q9D6J6 | Ran GTPase-activating protein 1 | P46061 | Cytochrome b-c1 complex subunit 8 | Q9CQ69 |
| NADH dehydrogenase [ubiquinone] iron-sulfur pr | Q91WD5 | Ran-specific GTPase-activating protein | P34022 | Cytochrome c oxidase subunit 5B, mitochondrial | P19536 |
| NADH dehydrogenase [ubiquinone] iron-sulfur pr | Q9DCT2 | Rap guanine nucleotide exchange factor 4 | Q9EQZ6 | Cytochrome c oxidase subunit 7A-related protein, | Q61387 |
| NADH dehydrogenase [ubiquinone] iron-sulfur pr | Q9CXZ1 | Ras suppressor protein 1 | Q01730 | Cytochrome c1, heme protein, mitochondrial | Q9D0M3 |
| NADH dehydrogenase [ubiquinone] iron-sulfur pr | Q8K3J1 | Ras-related protein M-Ras | O08989 | Cytohesin-1 | Q9QX11 |
| NADH-cytochrome b5 reductase 3 | Q9DCN2 | Ras-related protein Rab-10 | P61027 | Cytohesin-2 | P63034 |
| NADH-ubiquinone oxidoreductase 75 kDa subun | Q91VD9 | Ras-related protein Rab-1A | P62821 | Cytohesin-3 | O08967 |
| NADP-dependent malic enzyme | P06801 | Ras-related protein Rab-27B | Q99P58 | Cytoplasmic aconitate hydratase | P28271 |
| N-alpha-acetyltransferase 15, NatA auxiliary sub | Q80UM3 | Ras-related protein Rab-2B | P59279 | Death-associated protein kinase 1 | Q80YE7 |
| Nascent polypeptide-associated complex subunit | P70670 | Ras-related protein Rab-4A | P56371 | Dedicator of cytokinesis protein 10 | Q8BZN6 |
| Nck-associated protein 1 | P28660 | Ras-related protein Rab-5A | Q9CQD1 | Dedicator of cytokinesis protein 11 | A2AF47 |
| NCK-interacting protein with SH3 domain | Q9ESJ4 | Ras-related protein Rab-6B | P61294 | Dedicator of cytokinesis protein 3 | Q8CIQ7 |
| NEDD8 | P29595 | Ras-related protein Rab-7a | P51150 | Dedicator of cytokinesis protein 7 | Q8R1A4 |
| NEDD8-activating enzyme E1 regulatory subunit | Q8VBW6 | Ras-related protein Rab-8B | P61028 | Dedicator of cytokinesis protein 9 | Q8BIK4 |
| NEDD8-conjugating enzyme Ubc12 | P61082 | Ras-related protein Ral-B | Q9JIW9 | Dehydrogenase/reductase SDR family member 1 | Q99L04 |
| Nesprin-1 | Q6ZWR6 | Ras-related protein Rap-2b | P61226 | Deleted in bladder cancer protein 1 homolog | Q920P3 |
| Neurabin-2 | Q6R891 | Ras-related protein R-Ras2 | P62071 | DENN domain-containing protein 1A | Q8K382 |
| Neural cell adhesion molecule 1 | P13595 | Ras-specific guanine nucleotide-releasing factor | P27671 | Desmoplakin | E9Q557 |
| Neural cell adhesion molecule 2 | O35136 | Receptor expression-enhancing protein 2 | Q8VCD6 | Diacylglycerol kinase epsilon | Q9R1C6 |
| Neural cell adhesion molecule L1 | P11627 | Regulator of G-protein signaling 14 | P97492 | Dihydroorotate dehydrogenase (quinone), mitoch | O35435 |
| Neural cell adhesion molecule L1-like protein | P70232 | Regulator of microtubule dynamics protein 3 | Q3UJU9 | Dipeptidyl peptidase 3 | Q99KK7 |
| Neurexin-1-alpha | Q9CS84 | Retinal dehydrogenase 1 | P24549 | Dipeptidyl peptidase 8 | Q80YA7 |
| Neurexin-3-alpha | Q6P9K9 | Retinol dehydrogenase 14 | Q9ERI6 | DIS3-like exonuclease 2 | Q8CI75 |
| Neurobeachin | Q9EPN1 | Rho GTPase-activating protein 26 | Q6ZQ82 | Disabled homolog 2-interacting protein | Q3UHC7 |
| Neurocalcin-delta | Q91X97 | Rho GTPase-activating protein 35 | Q91YM2 | Disheveled-associated activator of morphogenesi | Q8BPM0 |
| Neurocan core protein | P55066 | Rho GTPase-activating protein 39 | P59281 | Disheveled-associated activator of morphogenesi | Q80U19 |
| Neurochondrin | Q9Z0E0 | Rho GTPase-activating protein 44 | Q5SSM3 | Disintegrin and metalloproteinase domain-contair | Q9R1V4 |
| Neurofascin | Q810U3 | Rho-related GTP-binding protein RhoG | P84096 | Disks large-associated protein 3 | Q6PFD5 |
| Neurofibromin | Q04690 | Ribonuclease UK114 | P52760 | DmX-like protein 1 | Q6PNC0 |
| Neurofilament heavy polypeptide | P19246 | Ribosomal protein S6 kinase delta-1 | Q8BLK9 | DNA topoisomerase 1 | Q04750 |
| Neurofilament light polypeptide | P08551 | RNA-binding protein 25 | B2RY56 | DNA-directed RNA polymerase II subunit RPB1 | P08775 |
| Neurofilament medium polypeptide | P08553 | Rootletin | Q8CJ40 | DNA-directed RNA polymerase II subunit RPB2 | Q8CFI7 |
| Neuromodulin | P06837 | Secretory carrier-associated membrane protein 1 | Q8K021 | Dolichol-phosphate mannosyltransferase | O70152 |
| Neuron navigator 1 | Q8CH77 | Seizure 6-like protein 2 | Q4V9Z5 | Dolichyl-diphosphooligosaccharide--protein glyco | P46978 |
| Neuronal cell adhesion molecule | Q810U4 | Selenium-binding protein 2 | Q63836 | Double zinc ribbon and ankyrin repeat-containing | Q8C008 |
| Neuronal membrane glycoprotein M6-a | P35802 | Septin-4 | P28661 | Double-stranded RNA-binding protein Staufen ho | Q9Z108 |
| Neuronal membrane glycoprotein M6-b | P35803 | Serine/arginine repetitive matrix protein 2 | Q8BTI8 | Double-stranded RNA-specific adenosine deamin | Q99MU3 |
| Neuronal pentraxin receptor | Q99J85 | Serine/arginine-rich splicing factor 1 | Q6PDM2 | Dual specificity mitogen-activated protein kinase | P31938 |
| Neuronal pentraxin-1 | Q62443 | Serine/arginine-rich splicing factor 4 | Q8VE97 | Dynamin-2 | P39054 |
| Neuronal-specific septin-3 | Q9Z1S5 | Serine/threonine-protein kinase BRSK1 | Q5RJI5 | Dystrobrevin beta | O70585 |
| Neuron-specific calcium-binding protein hippocal | P84075 | Serine/threonine-protein kinase BRSK2 | Q69Z98 | E3 ubiquitin-protein ligase HECTD3 | Q3U487 |
| Neuroplastin | P97300 | Serine/threonine-protein kinase MARK1 | Q8VHJ5 | E3 ubiquitin-protein ligase HERC2 | Q4U2R1 |
| Neurotrimin | Q99PJ0 | Serine/threonine-protein kinase MARK2 | Q05512 | E3 ubiquitin-protein ligase RBBP6 | P97868 |
| Neutral alpha-glucosidase AB | Q8BHN3 | Serine/threonine-protein kinase MRCK alpha | Q3UU96 | E3 ubiquitin-protein ligase TRIM32 | Q8CH72 |
| NF-kappa-B essential modulator | O88522 | Serine/threonine-protein kinase mTOR | Q9JLN9 | E3 UFM1-protein ligase 1 | Q8CCJ3 |

| | | | | | |
|---|---|---|---|---|---|
| Nicastrin | P57716 | Serine/threonine-protein kinase OSR1 | Q6P9R2 | Echinoderm microtubule-associated protein-like 4 | Q3UMY5 |
| Nitric oxide synthase, brain | Q9Z0J4 | Serine/threonine-protein kinase PAK 3 | Q61036 | Ectonucleotide pyrophosphatase/phosphodiestera | Q8BGN3 |
| NmrA-like family domain-containing protein 1 | Q8K2T1 | Serine/threonine-protein kinase TAO2 | Q6ZQ29 | Ectopic P granules protein 5 homolog | Q80TA9 |
| NSFL1 cofactor p47 | Q9CZ44 | Serine/threonine-protein phosphatase 2A 56 kDa | Q6PD03 | EF-hand domain-containing family member A2 | Q9CTY5 |
| N-terminal EF-hand calcium-binding protein 2 | Q91ZP9 | Serine/threonine-protein phosphatase 6 regulator | Q7TSI3 | Egl nine homolog 1 | Q91YE3 |
| Nuclear distribution protein nudE-like 1 | Q9ERR1 | Short/branched chain specific acyl-CoA dehydrog | Q9DBL1 | ELAV-like protein 2 | Q60899 |
| Nuclear migration protein nudC | O35685 | Signal-induced proliferation-associated 1-like prot | Q8C0T5 | ELAV-like protein 3 | Q60900 |
| Nucleolin | P09405 | SLIT and NTRK-like protein 5 | Q810B7 | Elongation factor G, mitochondrial | Q8K0D5 |
| Nucleosome assembly protein 1-like 4 | Q78ZA7 | Sodium/calcium exchanger 1 | P70414 | Elongator complex protein 1 | Q7TT37 |
| NudC domain-containing protein 3 | Q8R1N4 | Sodium/potassium-transporting ATPase subunit b | P14231 | Endonuclease domain-containing 1 protein | Q8C522 |
| Numb-like protein | O08919 | Sodium-dependent neutral amino acid transporter | Q8BJI1 | Endoplasmic reticulum-Golgi intermediate compa | Q9DC16 |
| Obg-like ATPase 1 | Q9CZ30 | Spermatogenesis-associated protein 2-like protei | Q8BNN1 | Engulfment and cell motility protein 1 | Q8BPU7 |
| Oligodendrocyte-myelin glycoprotein | Q63912 | Spermine synthase | P97355 | Enhancer of mRNA-decapping protein 4 | Q3UJB9 |
| Opioid growth factor receptor | Q99PG2 | Splicing factor U2AF 35 kDa subunit | Q9D883 | Enhancer of rudimentary homolog | P84089 |
| Oxidation resistance protein 1 | Q4KMM3 | Stathmin-2 | P55821 | Ephrin type-A receptor 5 | Q60629 |
| Oxysterol-binding protein 1 | Q3B7Z2 | Sterile alpha and TIR motif-containing protein 1 | Q6PDS3 | Ephrin type-B receptor 1 | Q8CBF3 |
| Palmdelphin | Q9JHU2 | Sulfotransferase 4A1 | P63046 | Ephrin type-B receptor 3 | P54754 |
| Pantothenate kinase 4 | Q80YV4 | Synaptic vesicle 2-related protein | Q8BFT9 | Ephrin-B2 | P52800 |
| Paralemmin-1 | Q9Z0P4 | Synaptosomal-associated protein 29 | Q9ERB0 | Epidermal growth factor receptor substrate 15 | P42567 |
| Paralemmin-2 | Q8BR92 | Synaptotagmin-17 | Q920M7 | Erlin-1 | Q91X78 |
| Paraspeckle component 1 | Q8R326 | Syntaphilin | Q80U23 | Estradiol 17-beta-dehydrogenase 11 | Q9EQ06 |
| Parathymosin | Q9D0J8 | Syntaxin-12 | Q9ER00 | Estradiol 17-beta-dehydrogenase 8 | P50171 |
| PC4 and SFRS1-interacting protein | Q99JF8 | Tenascin | Q80YX1 | Ethylmalonyl-CoA decarboxylase | Q9D9V3 |
| Peptidyl-prolyl cis-trans isomerase A | P17742 | Teneurin-4 | Q3UHK6 | Eukaryotic initiation factor 4A-III | Q91VC3 |
| Peptidyl-prolyl cis-trans isomerase B | P24369 | Thioredoxin reductase 1, cytoplasmic | Q9JMH6 | Eukaryotic translation initiation factor 2 subunit 2 | Q99L45 |
| Peptidyl-prolyl cis-trans isomerase D | Q9CR16 | Thioredoxin-dependent peroxide reductase, mitoc | P20108 | Eukaryotic translation initiation factor 2A | Q8BJW6 |
| Peptidyl-prolyl cis-trans isomerase F, mitochondr | Q99KR7 | Tight junction protein ZO-1 | P39447 | Eukaryotic translation initiation factor 3 subunit D | O70194 |
| Peptidyl-prolyl cis-trans isomerase FKBP3 | Q62446 | Traf2 and NCK-interacting protein kinase | P83510 | Exocyst complex component 2 | Q9D4H1 |
| Peptidyl-prolyl cis-trans isomerase FKBP4 | P30416 | Trafficking protein particle complex subunit 12 | Q8K2L8 | Exocyst complex component 6 | Q8R313 |
| Peptidyl-prolyl cis-trans isomerase FKBP8 | O35465 | Trafficking protein particle complex subunit 5 | Q9CQA1 | Exocyst complex component 6B | A6H5Z3 |
| Peptidyl-prolyl cis-trans isomerase NIMA-interact | Q9QUR7 | Trafficking protein particle complex subunit 9 | Q3U0M1 | Exocyst complex component 8 | Q6PGF7 |
| Peripheral plasma membrane protein CASK | O70589 | Trans-2-enoyl-CoA reductase, mitochondrial | Q9DCS3 | Exportin-5 | Q924C1 |
| Peroxiredoxin-1 | P35700 | Transaldolase | Q93092 | Exportin-7 | Q9EPK7 |
| Peroxiredoxin-2 | Q61171 | Transcriptional activator protein Pur-alpha | P42669 | Extended synaptotagmin-2 | Q3TZZ7 |
| Peroxiredoxin-6 | O08709 | Transferrin receptor protein 1 | Q62351 | FACT complex subunit SPT16 | Q920B9 |
| Peroxisomal acyl-coenzyme A oxidase 1 | Q9R0H0 | Transforming acidic coiled-coil-containing protein | Q9JJG0 | FAD synthase | Q8R123 |
| Peroxisomal biogenesis factor 19 | Q8VCI5 | Transforming protein RhoA | Q9QUI0 | Fasciculation and elongation protein zeta-1 | Q8K0X8 |
| Peroxisomal multifunctional enzyme type 2 | P51660 | Transmembrane protein 132A | Q922P8 | Fatty-acid amide hydrolase 1 | O08914 |
| PEX5-related protein | Q8C437 | tRNA-splicing ligase RtcB homolog | Q99LF4 | F-box only protein 7 | Q3U7U3 |
| PH and SEC7 domain-containing protein 3 | Q2PFD7 | Tubby protein | P50586 | F-box-like/WD repeat-containing protein TBL1X | Q9QXE7 |
| Phenylalanine--tRNA ligase alpha subunit | Q8C0C7 | Tubulin alpha-8 chain | Q9JJZ2 | Fermitin family homolog 2 | Q8CIB5 |
| Phosphatase and actin regulator 1 | Q2M3X8 | Tubulin-folding cofactor B | Q9D1E6 | Ferritin heavy chain | P09528 |
| Phosphate carrier protein, mitochondrial | Q8VEM8 | Tubulin-specific chaperone A | P48428 | Ferrochelatase, mitochondrial | P22315 |
| Phosphatidylethanolamine-binding protein 1 | P70296 | Tubulin-specific chaperone C | Q8VCN9 | Filamin-A | Q8BTM8 |
| Phosphatidylinositide phosphatase SAC1 | Q9EP69 | Tyrosine-protein phosphatase non-receptor type 2 | Q6PB44 | Filamin-B | Q80X90 |
| Phosphatidylinositol 4-phosphate 5-kinase type-1 | O70161 | U1 small nuclear ribonucleoprotein 70 kDa | Q62376 | Forkhead-associated domain-containing protein 1 | A6PWD2 |
| Phosphatidylinositol 5-phosphate 4-kinase type-2 | O70172 | Ubiquinone biosynthesis protein COQ9, mitochon | Q8K1Z0 | FUN14 domain-containing protein 2 | Q9D6K8 |
| Phosphatidylinositol 5-phosphate 4-kinase type-2 | Q80XI4 | Ubiquitin carboxyl-terminal hydrolase 10 | P52479 | FYVE, RhoGEF and PH domain-containing protei | P52734 |

| | |
|---|---|
| Phosphatidylinositol transfer protein alpha isofor | P53810 |
| Phosphatidylinositol-binding clathrin assembly pr | Q7M6Y3 |
| Phosphofurin acidic cluster sorting protein 1 | Q8K212 |
| Phosphoglucomutase-1 | Q9D0F9 |
| Phosphoglycerate kinase 1 | P09411 |
| Phosphoglycerate mutase 1 | Q9DBJ1 |
| Phosphoglycerate mutase 2 | O70250 |
| Phosphoglycolate phosphatase | Q8CHP8 |
| Phospholipase D3 | O35405 |
| Phospholipid hydroperoxide glutathione peroxida | O70325 |
| Phosphomannomutase 1 | O35621 |
| Phosphoribosylformylglycinamidine synthase | Q5SUR0 |
| Phosphoserine aminotransferase | Q99K85 |
| Phytanoyl-CoA hydroxylase-interacting protein-lik | Q8BGT8 |
| Plakophilin-4 | Q68FH0 |
| Plasma membrane calcium-transporting ATPase | Q9R0K7 |
| Plasminogen activator inhibitor 1 RNA-binding pr | Q9CY58 |
| Platelet-activating factor acetylhydrolase IB subu | P63005 |
| Plectin | Q9QXS1 |
| Plexin-A1 | P70206 |
| Plexin-A4 | Q80UG2 |
| Poly(rC)-binding protein 1 | P60335 |
| Poly(rC)-binding protein 2 | Q61990 |
| Polyadenylate-binding protein 1 | P29341 |
| Polyubiquitin-C | P0CG50 |
| Potassium voltage-gated channel subfamily KQT | Q9Z351 |
| Prefoldin subunit 2 | O70591 |
| Presequence protease, mitochondrial | Q8K411 |
| PRKCA-binding protein | Q62083 |
| Probable phospholipid-transporting ATPase IA | P70704 |
| Probable threonine--tRNA ligase 2, cytoplasmic | Q8BLY2 |
| Probable ubiquitin carboxyl-terminal hydrolase F | P70398 |
| Programmed cell death 6-interacting protein | Q9WU78 |
| Prohibitin | P67778 |
| Prohibitin-2 | O35129 |
| Proliferation-associated protein 2G4 | P50580 |
| Proline synthase co-transcribed bacterial homolo | Q9Z2Y8 |
| Proline-rich transmembrane protein 1 | O35449 |
| Proline-rich transmembrane protein 2 | E9PUL5 |
| Prolow-density lipoprotein receptor-related protei | Q91ZX7 |
| Prolyl endopeptidase | Q9QUR6 |
| Prolyl endopeptidase-like | Q8C167 |
| Propionyl-CoA carboxylase alpha chain, mitocho | Q91ZA3 |
| ProSAAS | Q9QXV0 |
| Proteasome subunit alpha type-1 | Q9R1P4 |
| Proteasome subunit alpha type-3 | O70435 |
| Proteasome subunit alpha type-4 | Q9R1P0 |
| Proteasome subunit alpha type-6 | Q9QUM9 |

| | |
|---|---|
| Ubiquitin carboxyl-terminal hydrolase CYLD | Q80TQ2 |
| Ubiquitin-like modifier-activating enzyme 5 | Q8VE47 |
| UBX domain-containing protein 6 | Q99PL6 |
| UHRF1-binding protein 1-like | A2RSJ4 |
| Uncharacterized protein C6orf203 homolog | Q9CQF4 |
| Uncharacterized protein KIAA0090 | Q8C7X2 |
| Uncharacterized protein KIAA0564 homolog | Q8CC88 |
| UPF0606 protein KIAA1549 | Q68FD9 |
| UTP--glucose-1-phosphate uridylyltransferase | Q91ZJ5 |
| Vacuolar protein sorting-associated protein 11 ho | Q91W86 |
| Vacuolar protein sorting-associated protein 26B | Q8C0E2 |
| Vacuolar protein sorting-associated protein 29 | Q9QZ88 |
| Vesicle-associated membrane protein-associated | Q9WV55 |
| Vesicle-trafficking protein SEC22b | O08547 |
| Vimentin | P20152 |
| Voltage-dependent calcium channel subunit alpha | Q6PHS9 |
| Voltage-dependent calcium channel subunit alpha | Q9Z1L5 |
| Voltage-dependent L-type calcium channel subun | Q8CC27 |
| Voltage-dependent N-type calcium channel subur | O55017 |
| Voltage-dependent R-type calcium channel subur | Q61290 |
| V-type proton ATPase subunit H | Q8BVE3 |
| WASH complex subunit FAM21 | Q6PGL7 |

| | |
|---|---|
| Gamma-aminobutyric acid receptor subunit alpha | P26049 |
| Gamma-aminobutyric acid receptor subunit beta- | P63137 |
| Gamma-aminobutyric acid receptor-associated pr | P60521 |
| Gamma-aminobutyric acid type B receptor subun | Q9WV18 |
| Gamma-glutamylcyclotransferase | Q9D7X8 |
| Gamma-tubulin complex component 2 | Q921G8 |
| Gap junction alpha-1 protein | P23242 |
| Gelsolin | P13020 |
| Gem-associated protein 5 | Q8BX17 |
| Glia-derived nexin | Q07235 |
| Glutamate decarboxylase 2 | P48320 |
| Glutamate receptor, ionotropic kainate 2 | P39087 |
| Glutamate--cysteine ligase catalytic subunit | P97494 |
| Glutamate--cysteine ligase regulatory subunit | O09172 |
| Glutathione peroxidase 1 | P11352 |
| Glutathione S-transferase kappa 1 | Q9DCM2 |
| Glutathione S-transferase Mu 7 | Q80W21 |
| Glycine amidinotransferase, mitochondrial | Q9D964 |
| Glycogen phosphorylase, liver form | Q9ET01 |
| Golgi phosphoprotein 3 | Q9CRA5 |
| Golgin subfamily A member 4 | Q91VW5 |
| Golgin subfamily A member 7B | Q9D428 |
| G-protein coupled receptor family C group 5 mem | Q923Z0 |
| GRAM domain-containing protein 1B | Q80TI0 |
| GTPase HRas | Q61411 |
| GTPase-activating protein and VPS9 domain-con | Q6PAR5 |
| GTP-binding protein Di-Ras1 | Q91Z61 |
| GTP-binding protein SAR1a | P36536 |
| Guanine nucleotide-binding protein-like 1 | P36916 |
| HCLS1-associated protein X-1 | O35387 |
| Heat shock 70 kDa protein 14 | Q99M31 |
| Hematopoietic lineage cell-specific protein | P49710 |
| Hepatoma-derived growth factor-related protein 3 | Q9JMG7 |
| Heterogeneous nuclear ribonucleoprotein A0 | Q9CX86 |
| Heterogeneous nuclear ribonucleoprotein A1 | P49312 |
| Heterogeneous nuclear ribonucleoprotein A3 | Q8BG05 |
| Heterogeneous nuclear ribonucleoprotein D-like | Q9Z130 |
| Heterogeneous nuclear ribonucleoprotein H | O35737 |
| Heterogeneous nuclear ribonucleoprotein H2 | P70333 |
| Heterogeneous nuclear ribonucleoprotein L | Q8R081 |
| Heterogeneous nuclear ribonucleoprotein M | Q9D0E1 |
| Hexaprenyldihydroxybenzoate methyltransferase, | Q8BMS4 |
| Histone deacetylase 2 | P70288 |
| Histone deacetylase 5 | Q9Z2V6 |
| Histone H1.4 | P43274 |
| Histone H1.5 | P43276 |
| Histone H2A.x | P27661 |
| Histone lysine demethylase PHF8 | Q80TJ7 |

| | | | |
|---|---|---|---|
| Proteasome subunit alpha type-7 | Q9Z2U0 | Histone-binding protein RBBP4 | Q60972 |
| Proteasome subunit beta type-1 | O09061 | Histone-lysine N-methyltransferase SETD1B | Q8CFT2 |
| Proteasome subunit beta type-5 | O55234 | Huntingtin-associated protein 1 | O35668 |
| Protein arginine N-methyltransferase 1 | Q9JIF0 | Hydroxymethylglutaryl-CoA lyase, mitochondrial | P38060 |
| Protein bassoon | O88737 | Hydroxysteroid dehydrogenase-like protein 2 | Q2TPA8 |
| Protein BRICK1 | Q91VR8 | Immunity-related GTPase family M protein 1 | Q60766 |
| Protein disulfide-isomerase | P09103 | Inositol hexakisphosphate and diphosphoinositol- | A2ARP1 |
| Protein disulfide-isomerase A3 | P27773 | Inositol-tetrakisphosphate 1-kinase | Q8BYN3 |
| Protein disulfide-isomerase A4 | P08003 | Insulin receptor | P15208 |
| Protein disulfide-isomerase A6 | Q922R8 | Insulin receptor substrate 2 | P81122 |
| Protein DJ-1 | Q99LX0 | Integral membrane protein 2B | O89051 |
| Protein FAM131B | Q3TY60 | Inter-alpha-trypsin inhibitor heavy chain H3 | Q61704 |
| Protein FAM136A | Q9CR98 | Interleukin enhancer-binding factor 3 | Q9Z1X4 |
| Protein FAM49A | Q8BHZ0 | Inverted formin-2 | Q0GNC1 |
| Protein FAM49B | Q921M7 | Isobutyryl-CoA dehydrogenase, mitochondrial | Q9D7B6 |
| Protein Hook homolog 3 | Q8BUK6 | Isochorismatase domain-containing protein 1 | Q91V64 |
| Protein IMPACT | O55091 | Isoleucine--tRNA ligase, mitochondrial | Q8BIJ6 |
| Protein KIAA0284 | Q80U49 | Junction plakoglobin | Q02257 |
| Protein KIAA1045 | Q80TL4 | Kelch-like protein 22 | Q99JN2 |
| Protein kinase C alpha type | P20444 | Kelch-like protein 3 | E0CZ16 |
| Protein kinase C and casein kinase substrate in r | Q61644 | Ketosamine-3-kinase | Q8K274 |
| Protein kinase C and casein kinase substrate in r | Q9WVE8 | KH domain-containing, RNA-binding, signal trans | Q9R226 |
| Protein kinase C beta type | P68404 | Kinesin heavy chain isoform 5A | P33175 |
| Protein kinase C epsilon type | P16054 | Kinesin light chain 4 | Q9DBS5 |
| Protein kinase C gamma type | P63318 | Kinesin-like protein KIF16B | B1AVY7 |
| Protein lin-7 homolog A | Q8JZS0 | Kinesin-like protein KIF1B | Q60575 |
| Protein lin-7 homolog B | O88951 | Lamina-associated polypeptide 2, isoforms beta/g | Q61029 |
| Protein lunapark | Q7TQ95 | Lamin-B1 | P14733 |
| Protein LYRIC | Q80WJ7 | Lamin-B2 | P21619 |
| Protein NDRG2 | Q9QYG0 | Large neutral amino acids transporter small subu | Q9Z127 |
| Protein phosphatase 1 regulatory subunit 12A | Q9DBR7 | Leucine-rich repeat and WD repeat-containing pro | Q6P5U7 |
| Protein phosphatase 1 regulatory subunit 12C | Q3UMT1 | Leucine-rich repeat-containing G-protein coupled | Q3UVD5 |
| Protein phosphatase 1 regulatory subunit 1B | Q60829 | Leucine-rich repeat-containing protein 57 | Q9D1G5 |
| Protein phosphatase 1 regulatory subunit 21 | Q3TDD9 | LIM domain and actin-binding protein 1 | Q9ERG0 |
| Protein phosphatase 1 regulatory subunit 29 | Q68FM6 | LMBR1 domain-containing protein 2 | Q8C561 |
| Protein phosphatase 1 regulatory subunit 7 | Q3UM45 | Long-chain fatty acid transport protein 1 | Q60714 |
| Protein phosphatase 1A | P49443 | Long-chain fatty acid transport protein 4 | Q91VE0 |
| Protein phosphatase 1E | Q80TL0 | Long-chain-fatty-acid--CoA ligase 1 | P41216 |
| Protein phosphatase 1G | Q61074 | MAGUK p55 subfamily member 3 | O88910 |
| Protein phosphatase 1H | Q3UYC0 | MAP/microtubule affinity-regulating kinase 4 | Q8CIP4 |
| Protein phosphatase methylesterase 1 | Q8BVQ5 | Melanoma inhibitory activity protein 3 | Q8BI84 |
| Protein piccolo | Q9QYX7 | Melanoma-associated antigen D1 | Q9QYH6 |
| Protein PRRC2C | Q3TLH4 | Membrane-associated phosphatidylinositol transfe | Q3UHE1 |
| Protein prune homolog | Q8BIW1 | Metabotropic glutamate receptor 7 | Q68ED2 |
| Protein RUFY3 | Q9D394 | Metal transporter CNNM1 | Q0GA42 |
| Protein SCAI | Q8C8N2 | Methionine aminopeptidase 1 | Q8BP48 |
| Protein SET | Q9EQU5 | Microtubule-associated protein RP/EB family mer | Q61166 |
| Protein shisa-6 homolog | Q3UH99 | Microtubule-associated protein RP/EB family mer | Q8R001 |

| | |
|---|---|
| Protein shisa-7 | Q8C3Q5 |
| Protein Shroom2 | A2ALU4 |
| Protein transport protein Sec31A | Q3UPL0 |
| Protein tweety homolog 3 | Q6P5F7 |
| Protein unc-13 homolog A | Q4KUS2 |
| Protein-methionine sulfoxide oxidase MICAL3 | Q8CJ19 |
| Protein-tyrosine kinase 2-beta | Q9QVP9 |
| Prothymosin alpha | P26350 |
| Purkinje cell protein 4 | P63054 |
| Puromycin-sensitive aminopeptidase | Q11011 |
| Putative adenosylhomocysteinase 2 | Q80SW1 |
| Putative adenosylhomocysteinase 3 | Q68FL4 |
| Pyridoxal kinase | Q8K183 |
| Pyridoxal phosphate phosphatase | P60487 |
| Pyruvate carboxylase, mitochondrial | Q05920 |
| Pyruvate dehydrogenase E1 component subunit | P35486 |
| Pyruvate dehydrogenase E1 component subunit | Q9D051 |
| Pyruvate dehydrogenase protein X component, r | Q8BKZ9 |
| Pyruvate kinase isozymes M1/M2 | P52480 |
| Rab GDP dissociation inhibitor alpha | P50396 |
| Rab GDP dissociation inhibitor beta | Q61598 |
| Rab GTPase-activating protein 1 | A2AWA9 |
| Rab GTPase-binding effector protein 1 | O35551 |
| Rab3 GTPase-activating protein non-catalytic su | Q8BMG7 |
| Rabphilin-3A | P47708 |
| Radixin | P26043 |
| Ran-binding protein 3 | Q9CT10 |
| Rap guanine nucleotide exchange factor 2 | Q8CHG7 |
| Ras GTPase-activating protein SynGAP | F6SEU4 |
| Ras GTPase-activating-like protein IQGAP2 | Q3UQ44 |
| RasGAP-activating-like protein 1 | Q9Z268 |
| Ras-related C3 botulinum toxin substrate 1 | P63001 |
| Ras-related protein Rab-11B | P46638 |
| Ras-related protein Rab-14 | Q91V41 |
| Ras-related protein Rab-1B | Q9D1G1 |
| Ras-related protein Rab-2A | P53994 |
| Ras-related protein Rab-35 | Q6PHN9 |
| Ras-related protein Rab-3A | P63011 |
| Ras-related protein Rab-3B | Q9CZT8 |
| Ras-related protein Rab-3C | P62823 |
| Ras-related protein Rab-4B | Q91ZR1 |
| Ras-related protein Rab-6A | P35279 |
| Ras-related protein Ral-A | P63321 |
| Ras-related protein Rap-1A | P62835 |
| Receptor-type tyrosine-protein phosphatase delta | Q64487 |
| Receptor-type tyrosine-protein phosphatase N2 | P80560 |
| Receptor-type tyrosine-protein phosphatase S | B0V2N1 |
| Regulating synaptic membrane exocytosis protei | Q99NE5 |

| | |
|---|---|
| Microtubule-associated serine/threonine-protein k | Q811L6 |
| Mitochondrial carrier homolog 1 | Q791T5 |
| Mitochondrial carrier homolog 2 | Q791V5 |
| Mitochondrial carrier triple repeat protein 1 | Q5HZI9 |
| Mitochondrial inner membrane protein OXA1L | Q8BGA9 |
| Mitochondrial intermediate peptidase | A6H611 |
| Mitochondrial Rho GTPase 1 | Q8BG51 |
| Mitochondrial translocator assembly and mainten | Q3TUH1 |
| Mitochondrial-processing peptidase subunit alpha | Q9DC61 |
| Mitogen-activated protein kinase 8 | Q91Y86 |
| Mitogen-activated protein kinase kinase kinase 1( | Q66L42 |
| Mitogen-activated protein kinase kinase kinase 5 | O35099 |
| Mitogen-activated protein kinase kinase kinase ki | P97820 |
| Mitotic checkpoint protein BUB3 | Q9WVA3 |
| Monoacylglycerol lipase ABHD6 | Q8R2Y0 |
| MOSC domain-containing protein 2, mitochondria | Q922Q1 |
| Motile sperm domain-containing protein 2 | Q9CWP6 |
| mRNA cap guanine-N7 methyltransferase | Q9D0L8 |
| Muscarinic acetylcholine receptor M1 | P12657 |
| Myb-binding protein 1A | Q7TPV4 |
| Myelin expression factor 2 | Q8C854 |
| Myelin gene regulatory factor | Q3UR85 |
| Myelin proteolipid protein | P60202 |
| Myeloid leukemia factor 2 | Q99KX1 |
| Myosin light chain 6B | Q8CI43 |
| Myosin phosphatase Rho-interacting protein | P97434 |
| Myosin-IIIb | Q1EG27 |
| Myotubularin-related protein 3 | Q8K296 |
| Myotubularin-related protein 9 | Q9Z2D0 |
| Na(+)/H(+) exchange regulatory cofactor NHE-RF | Q9JHL1 |
| N-acetyl-D-glucosamine kinase | Q9QZ08 |
| N-acetylgalactosamine kinase | Q68FH4 |
| N-acetylglucosamine-6-sulfatase | Q8BFR4 |
| N-acetyltransferase 10 | Q8K224 |
| NAD-dependent malic enzyme, mitochondrial | Q99KE1 |
| NADH dehydrogenase [ubiquinone] 1 alpha subco | Q9CQZ5 |
| NADH dehydrogenase [ubiquinone] 1 beta subco | O09111 |
| NADH dehydrogenase [ubiquinone] 1 subunit C2 | Q9CQ54 |
| NADH dehydrogenase [ubiquinone] iron-sulfur pro | Q9DC70 |
| NADPH:adrenodoxin oxidoreductase, mitochondr | Q61578 |
| Nasal embryonic luteinizing hormone-releasing ho | Q99NF2 |
| N-chimaerin | Q91V57 |
| NEDD4-binding protein 3 | Q8C7U1 |
| Neural Wiskott-Aldrich syndrome protein | Q91YD9 |
| Neurexophilin-1 | Q61200 |
| Neuritin | Q8CFV4 |
| Neuroligin-2 | Q69ZK9 |
| Neuroligin-3 | Q8BYM5 |

| | |
|---|---|
| Regulator of G-protein signaling 7 | O54829 |
| Reticulon-1 | Q8K0T0 |
| Reticulon-3 | Q9ES97 |
| Reticulon-4 | Q99P72 |
| Rho GDP-dissociation inhibitor 1 | Q99PT1 |
| Rho GTPase-activating protein 1 | Q5FWK3 |
| Rho GTPase-activating protein 21 | Q6DFV3 |
| Rho GTPase-activating protein 32 | Q811P8 |
| Rho guanine nucleotide exchange factor 12 | Q8R4H2 |
| Rho guanine nucleotide exchange factor 2 | Q60875 |
| Rho-associated protein kinase 2 | P70336 |
| Ribonuclease inhibitor | Q91VI7 |
| Ribosome-binding protein 1 | Q99PL5 |
| RIMS-binding protein 2 | Q80U40 |
| RING finger protein 214 | Q8BFU3 |
| RNA-binding protein FUS | P56959 |
| Roundabout homolog 2 | Q7TPD3 |
| RuvB-like 2 | Q9WTM5 |
| Ryanodine receptor 2 | E9Q401 |
| Sarcoplasmic/endoplasmic reticulum calcium AT | O55143 |
| Scavenger mRNA-decapping enzyme DcpS | Q9DAR7 |
| SCY1-like protein 2 | Q8CFE4 |
| Secernin-1 | Q9CZC8 |
| Secretogranin-1 | P16014 |
| Secretogranin-2 | Q03517 |
| Sepiapterin reductase | Q64105 |
| Septin-11 | Q8C1B7 |
| Septin-5 | Q9Z2Q6 |
| Septin-7 | O55131 |
| Septin-8 | Q8CHH9 |
| Septin-9 | Q80UG5 |
| Serine protease inhibitor A3K | P07759 |
| Serine/arginine-rich splicing factor 2 | Q62093 |
| Serine/threonine-protein kinase DCLK1 | Q9JLM8 |
| Serine/threonine-protein kinase DCLK2 | Q6PGN3 |
| Serine/threonine-protein kinase PAK 1 | O88643 |
| Serine/threonine-protein kinase PAK 2 | Q8CIN4 |
| Serine/threonine-protein kinase WNK1 | P83741 |
| Serine/threonine-protein kinase WNK2 | Q3UH66 |
| Serine/threonine-protein kinase WNK3 | Q80XP9 |
| Serine/threonine-protein phosphatase 2A 55 kDa | Q6P1F6 |
| Serine/threonine-protein phosphatase 2A 56 kDa | Q61151 |
| Serine/threonine-protein phosphatase 2A 56 kDa | Q60996 |
| Serine/threonine-protein phosphatase 2A 65 kDa | Q76MZ3 |
| Serine/threonine-protein phosphatase 2A activat | P58389 |
| Serine/threonine-protein phosphatase 2A catalyti | P63330 |
| Serine/threonine-protein phosphatase 2B catalyti | P63328 |
| Serine/threonine-protein phosphatase 2B catalyti | P48453 |

| | |
|---|---|
| Neuronal pentraxin-2 | O70340 |
| Neuropilin-1 | P97333 |
| NFU1 iron-sulfur cluster scaffold homolog, mitoch | Q9QZ23 |
| NLR family member X1 | Q3TL44 |
| Non-specific lipid-transfer protein | P32020 |
| Nuclear pore complex protein Nup93 | Q8BJ71 |
| Nuclear receptor coactivator 5 | Q91W39 |
| Nuclear receptor-interacting protein 3 | Q9JJR9 |
| Nuclear RNA export factor 1 | Q99JX7 |
| Nucleolar protein 56 | Q9D6Z1 |
| Nucleolar protein 58 | Q6DFW4 |
| Nucleolar transcription factor 1 | P25976 |
| Nucleolysin TIAR | P70318 |
| Nucleosome assembly protein 1-like 1 | P28656 |
| NudC domain-containing protein 2 | Q9CQ48 |
| O-acetyl-ADP-ribose deacetylase MACROD1 | Q922B1 |
| Oligophrenin-1 | Q99J31 |
| Oxysterol-binding protein 2 | Q5QNQ6 |
| Oxysterol-binding protein-related protein 6 | Q8BXR9 |
| Paraplegin | Q3ULF4 |
| Peflin | Q8BFY6 |
| Pentatricopeptide repeat-containing protein 3, mit | Q14C51 |
| Peptidyl-glycine alpha-amidating monooxygenase | P97467 |
| Perilipin-4 | O88492 |
| Peripherin | P15331 |
| Peroxisomal membrane protein PEX14 | Q9R0A0 |
| PERQ amino acid-rich with GYF domain-containi | Q6Y7W8 |
| Phosphatidylinositol 3-kinase catalytic subunit typ | Q6PF93 |
| Phosphatidylinositol 4,5-bisphosphate 3-kinase ca | Q8BTI9 |
| Phosphatidylinositol 4,5-bisphosphate 5-phospha | P59644 |
| Phosphatidylserine decarboxylase proenzyme | Q8BSF4 |
| Phosphatidylserine synthase 1 | Q99LH2 |
| Phosphoethanolamine/phosphocholine phosphata | Q8R2H9 |
| Phosphoinositide 3-kinase regulatory subunit 4 | Q8VD65 |
| Phospholipase A-2-activating protein | P27612 |
| Phosphorylase b kinase regulatory subunit alpha, | P18826 |
| Phosphorylase b kinase regulatory subunit beta | Q7TSH2 |
| Pinin | O35691 |
| Plasminogen receptor (KT) | Q9D3P8 |
| Pleckstrin homology domain-containing family A r | Q7TQG1 |
| Pleiotropic regulator 1 | Q922V4 |
| Poly(U)-binding-splicing factor PUF60 | Q3UEB3 |
| Polyadenylate-binding protein 2 | Q8CCS6 |
| Polymerase I and transcript release factor | O54724 |
| Polypyrimidine tract-binding protein 2 | Q91Z31 |
| Polypyrimidine tract-binding protein 3 | Q8BHD7 |
| Polyribonucleotide nucleotidyltransferase 1, mitoc | Q8K1R3 |
| Potassium/sodium hyperpolarization-activated cy | O88704 |

| | |
|---|---|
| Serine/threonine-protein phosphatase 5 | Q60676 |
| Serine/threonine-protein phosphatase PGAM5, n | Q8BX10 |
| Serine-threonine kinase receptor-associated prot | Q9Z1Z2 |
| Serine--tRNA ligase, cytoplasmic | P26638 |
| Serotransferrin | Q921I1 |
| Serum albumin | P07724 |
| SH3 and multiple ankyrin repeat domains protein | D3YZU1 |
| SH3 and multiple ankyrin repeat domains protein | Q80Z38 |
| SH3 and multiple ankyrin repeat domains protein | Q4ACU6 |
| SH3 domain-binding glutamic acid-rich-like prote | Q9JJU8 |
| SH3 domain-containing kinase-binding protein 1 | Q8R550 |
| SH3-containing GRB2-like protein 3-interacting p | Q8VD37 |
| Sideroflexin-3 | Q91V61 |
| SLIT-ROBO Rho GTPase-activating protein 2 | Q91Z67 |
| SLIT-ROBO Rho GTPase-activating protein 3 | Q812A2 |
| Small glutamine-rich tetratricopeptide repeat-con | Q8BJU0 |
| Sn1-specific diacylglycerol lipase alpha | Q6WQJ1 |
| Sodium- and chloride-dependent GABA transpor | P31650 |
| Sodium/potassium-transporting ATPase subunit | Q8VDN2 |
| Sodium/potassium-transporting ATPase subunit | Q6PIE5 |
| Sodium/potassium-transporting ATPase subunit | Q6PIC6 |
| Sodium/potassium-transporting ATPase subunit | P14094 |
| Sodium-driven chloride bicarbonate exchanger | Q5DTL9 |
| Solute carrier family 12 member 5 | Q91V14 |
| Sorbin and SH3 domain-containing protein 1 | Q62417 |
| Sortilin | Q6PHU5 |
| Sorting nexin-1 | Q9WV80 |
| Sorting nexin-2 | Q9CWK8 |
| Sorting nexin-27 | Q3UHD6 |
| Sorting nexin-5 | Q9D8U8 |
| Sorting nexin-6 | Q6P8X1 |
| Spectrin alpha chain, brain | P16546 |
| Spectrin beta chain, brain 1 | Q62261 |
| Spectrin beta chain, erythrocyte | P15508 |
| S-phase kinase-associated protein 1 | Q9WTX5 |
| Sphingomyelin phosphodiesterase 3 | Q9JJY3 |
| Splicing factor, proline- and glutamine-rich | Q8VIJ6 |
| SRC kinase signaling inhibitor 1 | Q9QWI6 |
| Src substrate cortactin | Q60598 |
| Staphylococcal nuclease domain-containing prot | Q78PY7 |
| Stathmin | P54227 |
| Stathmin-3 | O70166 |
| STE20/SPS1-related proline-alanine-rich protein | Q9Z1W9 |
| STE20-like serine/threonine-protein kinase | O54988 |
| STIP1 homology and U box-containing protein 1 | Q9WUD1 |
| Stomatin-like protein 2 | Q99JB2 |
| Stress-70 protein, mitochondrial | P38647 |
| Stress-induced-phosphoprotein 1 | Q60864 |

| | |
|---|---|
| Prelamin-A/C | P48678 |
| Pre-mRNA-processing factor 40 homolog A | Q9R1C7 |
| Pre-mRNA-processing factor 6 | Q91YR7 |
| Pre-mRNA-processing-splicing factor 8 | Q99PV0 |
| Pre-mRNA-splicing factor SYF1 | Q9DCD2 |
| Prenylcysteine oxidase | Q9CQF9 |
| Probable arginine--tRNA ligase, mitochondrial | Q3U186 |
| Probable ATP-dependent RNA helicase DDX46 | Q569Z5 |
| Probable ATP-dependent RNA helicase DDX6 | P54823 |
| Probable cationic amino acid transporter | Q8BXR1 |
| Probable global transcription activator SNF2L2 | Q6DIC0 |
| Probable histidine--tRNA ligase, mitochondrial | Q99KK9 |
| Probable hydrolase PNKD | Q69ZP3 |
| Probable phospholipid-transporting ATPase IIB | P98195 |
| Probable proline--tRNA ligase, mitochondrial | Q8CFI5 |
| Profilin-2 | Q9JJV2 |
| Proline dehydrogenase 1, mitochondrial | Q9WU79 |
| Proline-rich transmembrane protein 3 | Q6PE13 |
| Proteasome subunit beta type-7 | P70195 |
| Proteasome-associated protein ECM29 homolog | Q6PDI5 |
| Protein 4.1 | P48193 |
| Protein arginine N-methyltransferase 8 | Q6PAK3 |
| Protein argonaute-3 | Q8CJF9 |
| Protein bicaudal D homolog 1 | Q8BR07 |
| Protein C8orf37 homolog | Q3UJP5 |
| Protein CASC3 | Q8K3W3 |
| Protein cereblon | Q8C7D2 |
| Protein DEK | Q7TNV0 |
| Protein deltex-3 | Q80V91 |
| Protein diaphanous homolog 1 | O08808 |
| Protein diaphanous homolog 2 | O70566 |
| Protein DPCD | Q8BPA8 |
| Protein FAM208A | Q69ZR9 |
| Protein FAM210A | Q8BGY7 |
| Protein FAM59A | Q3UFT3 |
| Protein FAM65A | Q68FE6 |
| Protein LSM12 homolog | Q9D0R8 |
| Protein midA homolog, mitochondrial | Q9CWG8 |
| Protein NDRG3 | Q9QYF9 |
| Protein NDRG4 | Q8BTG7 |
| Protein NipSnap homolog 3B | Q9CQE1 |
| Protein Njmu-R1 | Q9CYI0 |
| Protein N-terminal glutamine amidohydrolase | Q80WB5 |
| Protein odr-4 homolog | Q4PJX1 |
| Protein phosphatase PTC7 homolog | Q6NVE9 |
| Protein Red | Q9Z1M8 |
| Protein SON | Q9QX47 |
| Protein TANC1 | Q0VGY8 |

| | |
|---|---|
| Striated muscle-specific serine/threonine-protein | Q62407 |
| Striatin | O55106 |
| Striatin-3 | Q9ERG2 |
| Succinate dehydrogenase [ubiquinone] flavoprot | Q8K2B3 |
| Succinate dehydrogenase [ubiquinone] iron-sulfu | Q9CQA3 |
| Succinate-semialdehyde dehydrogenase, mitoch | Q8BWF0 |
| Succinyl-CoA ligase [ADP-forming] subunit beta, | Q9Z2I9 |
| Succinyl-CoA ligase [GDP-forming] subunit beta, | Q9Z2I8 |
| Succinyl-CoA:3-ketoacid-coenzyme A transferas | Q9D0K2 |
| SUMO-activating enzyme subunit 1 | Q9R1T2 |
| SUMO-activating enzyme subunit 2 | Q9Z1F9 |
| Superoxide dismutase [Cu-Zn] | P08228 |
| Suppressor of G2 allele of SKP1 homolog | Q9CX34 |
| Synapsin-1 | O88935 |
| Synapsin-2 | Q64332 |
| Synaptic vesicle glycoprotein 2A | Q9JIS5 |
| Synaptic vesicle membrane protein VAT-1 homol | Q80TB8 |
| Synaptobrevin homolog YKT6 | Q9CQW1 |
| Synaptojanin-1 | Q8CHC4 |
| Synaptopodin | Q8CC35 |
| Synaptosomal-associated protein 25 | P60879 |
| Synaptosomal-associated protein 47 | Q8R570 |
| Synaptotagmin-1 | P46096 |
| Synaptotagmin-7 | Q9R0N7 |
| Synergin gamma | Q5SV85 |
| Syntaxin-1A | O35526 |
| Syntaxin-1B | P61264 |
| Syntaxin-7 | O70439 |
| Syntaxin-8 | O88983 |
| Syntaxin-binding protein 1 | O08599 |
| Syntaxin-binding protein 5 | Q8K400 |
| Syntaxin-binding protein 5-like | Q5DQR4 |
| Talin-2 | Q71LX4 |
| TBC1 domain family member 24 | Q3UUG6 |
| T-complex protein 1 subunit alpha | P11983 |
| T-complex protein 1 subunit beta | P80314 |
| T-complex protein 1 subunit delta | P80315 |
| T-complex protein 1 subunit epsilon | P80316 |
| T-complex protein 1 subunit eta | P80313 |
| T-complex protein 1 subunit gamma | P80318 |
| T-complex protein 1 subunit theta | P42932 |
| T-complex protein 1 subunit zeta | P80317 |
| Tectonin beta-propeller repeat-containing protein | Q80VP0 |
| Tenascin-R | Q8BYI9 |
| Thiamine-triphosphatase | Q8JZL3 |
| Thimet oligopeptidase | Q8C1A5 |
| Thiomorpholine-carboxylate dehydrogenase | O54983 |
| Thioredoxin | P10639 |

| | |
|---|---|
| Protein unc-79 homolog | Q0KK59 |
| Protein unc-80 homolog | Q8BLN6 |
| Protein zyg-11 homolog B | Q3UFS0 |
| Protein-arginine deiminase type-2 | Q08642 |
| Protein-tyrosine kinase 6 | Q64434 |
| Protein-tyrosine phosphatase mitochondrial 1 | Q66GT5 |
| Protocadherin-19 | Q80TF3 |
| Proto-oncogene tyrosine-protein kinase ROS | Q78DX7 |
| Putative ATP-dependent RNA helicase DHX30 | Q99PU8 |
| Pyruvate dehydrogenase phosphatase regulatory | Q7TSQ8 |
| Rab GTPase-activating protein 1-like | A6H6A9 |
| Rab3 GTPase-activating protein catalytic subunit | Q80UJ7 |
| Raftlin-2 | Q8CHX7 |
| Ral GTPase-activating protein subunit alpha-1 | Q6GYP7 |
| Ral GTPase-activating protein subunit beta | Q8BQZ4 |
| RalBP1-associated Eps domain-containing protei | Q80XA6 |
| Ran-binding protein 9 | P69566 |
| Rap guanine nucleotide exchange factor-like 1 | Q68EF8 |
| Rap1 GTPase-activating protein 2 | Q5SVL6 |
| Ras and Rab interactor 1 | Q921Q7 |
| Ras GTPase-activating protein 3 | Q60790 |
| Ras GTPase-activating-like protein IQGAP1 | Q9JKF1 |
| RAS guanyl-releasing protein 1 | Q9Z1S3 |
| Ras-related protein Rab-12 | P35283 |
| Ras-related protein Rab-15 | Q8K386 |
| Ras-related protein Rab-24 | P35290 |
| Ras-specific guanine nucleotide-releasing factor 2 | P70392 |
| RB1-inducible coiled-coil protein 1 | Q9ESK9 |
| Regulator of G-protein signaling 7-binding protein | Q8BQP9 |
| Regulator of microtubule dynamics protein 1 | Q9DCV4 |
| Required for meiotic nuclear division protein 1 ho | Q8CI78 |
| Retinal rod rhodopsin-sensitive cGMP 3',5'-cyclic | O55057 |
| Retinal-specific ATP-binding cassette transporter | O35600 |
| Rho GTPase-activating protein 23 | Q69ZH9 |
| Rho GTPase-activating protein 5 | P97393 |
| Rho guanine nucleotide exchange factor 17 | Q80U35 |
| Rho guanine nucleotide exchange factor 9 | Q3UTH8 |
| Rho-associated protein kinase 1 | P70335 |
| Ribonuclease 3 | Q5HZJ0 |
| Ribosomal protein S6 kinase alpha-4 | Q9Z2B9 |
| RNA binding motif protein, X-linked-like-1 | Q91VM5 |
| RNA polymerase II-associated protein 3 | Q9D706 |
| RNA polymerase-associated protein RTF1 homol | A2AQ19 |
| RNA-binding protein 10 | Q99KG3 |
| RNA-binding protein 26 | Q6NZN0 |
| RNA-binding protein 39 | Q8VH51 |
| RNA-binding protein EWS | Q61545 |
| RNA-binding protein Raly | Q64012 |

| | |
|---|---|
| Thioredoxin-like protein 1 | Q8CDN6 |
| Thiosulfate sulfurtransferase | P52196 |
| Threonine synthase-like 1 | Q8BH55 |
| Threonine--tRNA ligase, cytoplasmic | Q9D0R2 |
| Thrombospondin type-1 domain-containing prote | Q69ZU6 |
| THUMP domain-containing protein 1 | Q99J36 |
| Thy-1 membrane glycoprotein | P01831 |
| TIP41-like protein | Q8BH58 |
| Tissue factor | P20352 |
| Titin | A2ASS6 |
| TOM1-like protein 2 | Q5SRX1 |
| Transcription elongation factor A protein 1 | P10711 |
| Transcription elongation factor A protein-like 5 | Q8CCT4 |
| Transcription elongation factor B polypeptide 2 | P62869 |
| Transcription elongation regulator 1 | Q8CGF7 |
| Transcription intermediary factor 1-beta | Q62318 |
| Transcriptional activator protein Pur-beta | O35295 |
| Transforming acidic coiled-coil-containing protein | Q6Y685 |
| Transgelin | P37804 |
| Transgelin-2 | Q9WVA4 |
| Transgelin-3 | Q9R1Q8 |
| Transitional endoplasmic reticulum ATPase | Q01853 |
| Transketolase | P40142 |
| Translin-associated protein X | Q9QZE7 |
| Transmembrane emp24 domain-containing prote | Q99KF1 |
| Trifunctional enzyme subunit alpha, mitochondria | Q8BMS1 |
| Trifunctional enzyme subunit beta, mitochondrial | Q99JY0 |
| Trifunctional purine biosynthetic protein adenosir | Q64737 |
| Triosephosphate isomerase | P17751 |
| Tripeptidyl-peptidase 2 | Q64514 |
| Triple functional domain protein | Q0KL02 |
| Tropomodulin-2 | Q9JKK7 |
| Tropomyosin alpha-1 chain | P58771 |
| Tropomyosin alpha-3 chain | P21107 |
| Tropomyosin alpha-4 chain | Q6IRU2 |
| Tryptophan--tRNA ligase, cytoplasmic | P32921 |
| Tubulin alpha-4A chain | P68368 |
| Tubulin beta-2B chain | Q9CWF2 |
| Tubulin beta-3 chain | Q9ERD7 |
| Tubulin beta-4A chain | Q9D6F9 |
| Tubulin polymerization-promoting protein | Q7TQD2 |
| Tubulin polymerization-promoting protein family r | Q9CRB6 |
| Tumor protein D54 | Q9CYZ2 |
| Twinfilin-1 | Q91YR1 |
| Twinfilin-2 | Q9Z0P5 |
| Type I inositol 3,4-bisphosphate 4-phosphatase | Q9EPW0 |
| Tyrosine-protein phosphatase non-receptor type | P35235 |
| Tyrosine--tRNA ligase, cytoplasmic | Q91WQ3 |
| RNA-binding Raly-like protein | Q8BTF8 |
| rRNA 2'-O-methyltransferase fibrillarin | P35550 |
| Ryanodine receptor 1 | E9PZQ0 |
| Sacsin | Q9JLC8 |
| SAP domain-containing ribonucleoprotein | Q9D1J3 |
| Scaffold attachment factor B1 | D3YXK2 |
| Sec1 family domain-containing protein 1 | Q8BRF7 |
| Sec1 family domain-containing protein 2 | Q8BTY8 |
| SEC14-like protein 2 | Q99J08 |
| SEC23-interacting protein | Q6NZC7 |
| Secretory carrier-associated membrane protein 3 | O35609 |
| Semaphorin-7A | Q9QUR8 |
| Sentrin-specific protease 7 | Q8BUH8 |
| Septin-2 | P42208 |
| Serine beta-lactamase-like protein LACTB, mitocl | Q9EP89 |
| Serine/arginine-rich splicing factor 10 | Q9R0U0 |
| Serine/arginine-rich splicing factor 3 | P84104 |
| Serine/threonine-protein kinase A-Raf | P04627 |
| Serine/threonine-protein kinase LMTK3 | Q5XJV6 |
| Serine/threonine-protein kinase MRCK beta | Q7TT50 |
| Serine/threonine-protein kinase MST4 | Q99JT2 |
| Serine/threonine-protein kinase N1 | P70268 |
| Serine/threonine-protein kinase SIK3 | Q6P4S6 |
| Serine/threonine-protein phosphatase 2A 55 kDa | Q8BG02 |
| Serine/threonine-protein phosphatase 4 regulator | Q8C0Y0 |
| Serine/threonine-protein phosphatase PP1-alpha | P62137 |
| Serine/threonine-protein phosphatase PP1-beta c | P62141 |
| S-formylglutathione hydrolase | Q9R0P3 |
| SH3 and PX domain-containing protein 2A | O89032 |
| SH3 and PX domain-containing protein 2B | A2AAY5 |
| Sideroflexin-1 | Q99JR1 |
| Sideroflexin-5 | Q925N0 |
| Signal recognition particle 68 kDa protein | Q8BMA6 |
| Signal transducer and activator of transcription 1 | P42225 |
| Signal transducing adapter molecule 1 | P70297 |
| Sister chromatid cohesion protein PDS5 homolog | Q6A026 |
| Small ubiquitin-related modifier 2 | P61957 |
| Smith-Magenis syndrome chromosomal region ca | Q3UMB5 |
| Sodium bicarbonate cotransporter 3 | Q8BTY2 |
| Sodium channel protein type 8 subunit alpha | Q9WTU3 |
| Sodium channel protein type 9 subunit alpha | Q62205 |
| Solute carrier family 12 member 6 | Q924N4 |
| Solute carrier family 25 member 46 | Q9CQS4 |
| Sorbin and SH3 domain-containing protein 2 | Q3UTJ2 |
| Sortilin-related receptor | O88307 |
| Sorting nexin-12 | O70493 |
| Sorting nexin-15 | Q91WE1 |
| Sorting nexin-30 | Q8CE50 |

| | |
|---|---|
| Ubiquitin carboxyl-terminal hydrolase 14 | Q9JMA1 |
| Ubiquitin carboxyl-terminal hydrolase 15 | Q8R5H1 |
| Ubiquitin carboxyl-terminal hydrolase 5 | P56399 |
| Ubiquitin carboxyl-terminal hydrolase isozyme L1 | Q9R0P9 |
| Ubiquitin carboxyl-terminal hydrolase isozyme L3 | Q9JKB1 |
| Ubiquitin thioesterase OTUB1 | Q7TQI3 |
| Ubiquitin-associated protein 2-like | Q80X50 |
| Ubiquitin-conjugating enzyme E2 L3 | P68037 |
| Ubiquitin-conjugating enzyme E2 N | P61089 |
| Ubiquitin-conjugating enzyme E2 O | Q6ZPJ3 |
| Ubiquitin-conjugating enzyme E2 variant 1 | Q9CZY3 |
| Ubiquitin-like modifier-activating enzyme 1 | Q02053 |
| Ubiquitin-like modifier-activating enzyme 6 | Q8C7R4 |
| UDP-N-acetylglucosamine--peptide N-acetylgluc | Q8CGY8 |
| UMP-CMP kinase | Q9DBP5 |
| Uncharacterized protein C1orf198 homolog | Q8C3W1 |
| Uncharacterized protein C1orf96 homolog | Q8QZT2 |
| Uncharacterized protein KIAA0513 | Q8R0A7 |
| Uncharacterized protein KIAA1107 | Q80TK0 |
| Uncharacterized protein KIAA1671 | Q8BRV5 |
| Unconventional myosin-Id | Q5SYD0 |
| Unconventional myosin-Va | Q99104 |
| Unconventional myosin-VI | Q64331 |
| Unconventional myosin-XVIIIa | Q9JMH9 |
| UPF0364 protein C6orf211 homolog | A6H630 |
| UPF0568 protein C14orf166 homolog | Q9CQE8 |
| UV excision repair protein RAD23 homolog B | P54728 |
| Vacuolar protein sorting-associated protein 13C | Q8BX70 |
| Vacuolar protein sorting-associated protein 26A | P40336 |
| Vacuolar protein sorting-associated protein 33B | P59016 |
| Vacuolar protein sorting-associated protein 35 | Q9EQH3 |
| Vacuolar protein sorting-associated protein 53 hc | Q8CCB4 |
| Valine--tRNA ligase | Q9Z1Q9 |
| Versican core protein | Q62059 |
| Very long-chain specific acyl-CoA dehydrogenas | P50544 |
| Vesicle transport through interaction with t-SNAR | O89116 |
| Vesicle-associated membrane protein 1 | Q62442 |
| Vesicle-associated membrane protein 2 | P63044 |
| Vesicle-associated membrane protein-associatec | Q9QY76 |
| Vesicle-fusing ATPase | P46460 |
| Vigilin | Q8VDJ3 |
| Vinculin | Q64727 |
| Visinin-like protein 1 | P62761 |
| Voltage-dependent anion-selective channel prote | Q60932 |
| Voltage-dependent anion-selective channel prote | Q60930 |
| Voltage-dependent anion-selective channel prote | Q60931 |
| Voltage-dependent calcium channel gamma-8 su | Q8VHW2 |
| Voltage-dependent calcium channel subunit alph | O08532 |

| | |
|---|---|
| Sorting nexin-32 | Q80ZJ7 |
| Sorting nexin-7 | Q9CY18 |
| Spectrin alpha chain, erythrocyte | P08032 |
| Spermidine synthase | Q64674 |
| Splicing factor 3A subunit 1 | Q8K4Z5 |
| Splicing factor 3A subunit 3 | Q9D554 |
| Splicing factor 3B subunit 1 | Q99NB9 |
| Splicing factor 3B subunit 3 | Q921M3 |
| Splicing factor, arginine/serine-rich 19 | Q5U4C3 |
| Src kinase-associated phosphoprotein 2 | Q3UND0 |
| Stonin-2 | Q8BZ60 |
| Structural maintenance of chromosomes protein : | Q9CU62 |
| Structural maintenance of chromosomes protein : | Q9CW03 |
| Sulfide:quinone oxidoreductase, mitochondrial | Q9R112 |
| Superkiller viralicidic activity 2-like 2 | Q9CZU3 |
| SURP and G-patch domain-containing protein 2 | Q8CH09 |
| SWI/SNF-related matrix-associated actin-depend | Q91ZW3 |
| Synaptophysin | Q62277 |
| Synaptotagmin-12 | Q920N7 |
| Synaptotagmin-2 | P46097 |
| Synaptotagmin-3 | O35681 |
| Synembryn-A | Q3TIR3 |
| Talin-1 | P26039 |
| TAR DNA-binding protein 43 | Q921F2 |
| Tax1-binding protein 1 homolog | Q3UKC1 |
| TBC1 domain family member 10B | Q8BHL3 |
| TBC1 domain family member 5 | Q80XQ2 |
| TBC1 domain family member 8B | A3KGB4 |
| TBC1 domain family member 9B | Q5SVR0 |
| T-cell immunomodulatory protein | Q99KW9 |
| Teneurin-3 | Q9WTS6 |
| Tensin-3 | Q5SSZ5 |
| Testis-specific Y-encoded-like protein 4 | Q8VD63 |
| Tetratricopeptide repeat protein 38 | A3KMP2 |
| THO complex subunit 2 | B1AZI6 |
| THO complex subunit 4 | O08583 |
| Thymidylate kinase | P97930 |
| Thymocyte nuclear protein 1 | Q91YJ3 |
| Thyroid hormone receptor-associated protein 3 | Q569Z6 |
| Tight junction protein ZO-2 | Q9Z0U1 |
| TNF receptor-associated factor 3 | Q60803 |
| Torsin-1A-interacting protein 1 | Q921T2 |
| TOX high mobility group box family member 4 | Q8BU11 |
| TPR and ankyrin repeat-containing protein 1 | Q8BV79 |
| Trans-2,3-enoyl-CoA reductase | Q9CY27 |
| Transcription activator BRG1 | Q3TKT4 |
| Transcription elongation factor B polypeptide 1 | P83940 |
| Transcription elongation factor SPT5 | O55201 |

| | |
|---|---|
| Voltage-gated potassium channel subunit beta-2 | P62482 |
| V-type proton ATPase 116 kDa subunit a isoform | Q9Z1G4 |
| V-type proton ATPase catalytic subunit A | P50516 |
| V-type proton ATPase subunit B, brain isoform | P62814 |
| V-type proton ATPase subunit C 1 | Q9Z1G3 |
| V-type proton ATPase subunit D | P57746 |
| V-type proton ATPase subunit d 1 | P51863 |
| V-type proton ATPase subunit E 1 | P50518 |
| V-type proton ATPase subunit F | Q9D1K2 |
| V-type proton ATPase subunit G 2 | Q9WTT4 |
| WAS/WASL-interacting protein family member 3 | P0C7L0 |
| WD repeat-containing protein 1 | O88342 |
| WD repeat-containing protein 13 | Q91V09 |
| WD repeat-containing protein 44 | Q6NVE8 |
| WD repeat-containing protein 7 | Q920I9 |
| Wiskott-Aldrich syndrome protein family member | Q8R5H6 |
| Wiskott-Aldrich syndrome protein family member | Q8VHI6 |
| Wolframin | P56695 |
| Xaa-Pro aminopeptidase 1 | Q6P1B1 |
| Xaa-Pro dipeptidase | Q11136 |
| ZW10 interactor | Q9CQU5 |
| Transcription factor 20 | Q9EPQ8 |
| Transcription factor BTF3 homolog 4 | Q9CQH7 |
| Transcription factor HIVEP2 | Q3UHF7 |
| Transcriptional repressor p66-beta | Q8VHR5 |
| Transformer-2 protein homolog beta | P62996 |
| Translation initiation factor eIF-2B subunit alpha | Q99LC8 |
| Translation initiation factor eIF-2B subunit beta | Q99LD9 |
| Translation initiation factor eIF-2B subunit delta | Q61749 |
| Transmembrane 9 superfamily member 4 | Q8BH24 |
| Transmembrane emp24 domain-containing protei | Q9D1D4 |
| Transmembrane emp24 domain-containing protei | Q9R0Q3 |
| Transmembrane protein 109 | Q3UBX0 |
| Transmembrane protein 65 | Q4VAE3 |
| Transmembrane protein 85 | Q9CZX9 |
| Tripartite motif-containing protein 2 | Q9ESN6 |
| Tripartite motif-containing protein 3 | Q9R1R2 |
| Tropomodulin-1 | P49813 |
| Tubulin beta-2A chain | Q7TMM9 |
| Tubulin beta-4B chain | P68372 |
| Tubulin beta-5 chain | P99024 |
| Tubulin-specific chaperone cofactor E-like protein | Q8C5W3 |
| Tubulin--tyrosine ligase | P38585 |
| Tumor suppressor p53-binding protein 1 | P70399 |
| Tyrosine-protein kinase Blk | P16277 |
| Tyrosine-protein kinase Fer | P70451 |
| Tyrosine-protein kinase JAK1 | P52332 |
| Tyrosine-protein phosphatase non-receptor type ! | P54830 |
| Tyrosine-protein phosphatase non-receptor type ! | O35239 |
| Tyrosine-protein phosphatase non-receptor type ! | P97797 |
| U2 small nuclear ribonucleoprotein B'' | Q9CQI7 |
| Ubiquilin-1 | Q8R317 |
| Ubiquitin carboxyl-terminal hydrolase 24 | B1AY13 |
| Ubiquitin thioesterase OTUB2 | Q9CQX0 |
| Ubiquitin-conjugating enzyme E2 Q1 | Q7TSS2 |
| Ubiquitin-conjugating enzyme E2 Z | Q3UE37 |
| Ubiquitin-like modifier-activating enzyme ATG7 | Q9D906 |
| Ubiquitin-protein ligase E3A | O08759 |
| UDP-glucose:glycoprotein glucosyltransferase 1 | Q6P5E4 |
| UDP-N-acetylhexosamine pyrophosphorylase | Q91YN5 |
| UDP-N-acetylhexosamine pyrophosphorylase-like | Q3TW96 |
| Uncharacterized protein C10orf35 homolog | Q9D882 |
| Uncharacterized protein C19orf52 homolog | Q8BGX2 |
| Uncharacterized protein C2orf47 homolog, mitoch | Q8BHE8 |
| Uncharacterized protein C9orf135 homolog | Q9CQC3 |
| Uncharacterized protein C9orf172 homolog | A2AJA9 |
| Uncharacterized protein KIAA1467 | Q8BYI8 |
| Unconventional myosin-Ic | Q9WTI7 |
| Unconventional myosin-IXb | Q9QY06 |

| | |
|---|---|
| UPF0160 protein MYG1, mitochondrial | Q9JK81 |
| UPF0556 protein C19orf10 homolog | Q9CPT4 |
| UPF0565 protein C2orf69 homolog | Q9D9H8 |
| UPF0670 protein C8orf55 homolog | Q80ZW2 |
| UPF0696 protein C11orf68 homolog | Q8VD62 |
| Uridine-cytidine kinase 2 | Q99PM9 |
| Uroporphyrinogen decarboxylase | P70697 |
| Vacuolar protein sorting-associated protein 13A | Q5H8C4 |
| Vacuolar protein sorting-associated protein 41 ho | Q5KU39 |
| Vacuolar protein sorting-associated protein 4A | Q8VEJ9 |
| Vacuolar protein sorting-associated protein 52 ho | Q8C754 |
| Vascular cell adhesion protein 1 | P29533 |
| Very long-chain acyl-CoA synthetase | O35488 |
| Vesicle transport through interaction with t-SNAR | O88384 |
| Vesicle-associated membrane protein 7 | P70280 |
| Vesicular inhibitory amino acid transporter | O35633 |
| Voltage-dependent calcium channel gamma-3 su | Q9JJV5 |
| Voltage-dependent P/Q-type calcium channel sul | P97445 |
| VPS10 domain-containing receptor SorCS2 | Q9EPR5 |
| WAS protein family homolog 1 | Q8VDD8 |
| WD repeat-containing protein 11 | Q8K1X1 |
| WD repeat-containing protein 18 | Q4VBE8 |
| WD repeat-containing protein 26 | Q8C6G8 |
| WD repeat-containing protein 48 | Q8BH57 |
| WW domain-binding protein 11 | Q923D5 |
| WW domain-binding protein 2 | P97765 |
| Zinc finger C2HC domain-containing protein 1A | Q8BJH1 |
| Zinc finger E-box-binding homeobox 2 | Q9R0G7 |
| Zinc finger FYVE domain-containing protein 1 | Q810J8 |
| Zinc finger protein 330 | Q922H9 |
| Zinc finger RNA-binding protein | O88532 |
| Zinc transporter 1 | Q60738 |

### Table S5. Comparison of proteins detected by iTRAQ across ages

**Detected at P21 and Adult - 1766 proteins**

| Protein name | Uniprot ID |
|---|---|
| [Pyruvate dehydrogenase [lipoamide]] kinase isozy | Q922H2 |
| 1,5-anhydro-D-fructose reductase | Q9DCT1 |
| 10 kDa heat shock protein, mitochondrial | Q64433 |
| 14-3-3 protein beta/alpha | Q9CQV8 |
| 14-3-3 protein epsilon | P62259 |
| 14-3-3 protein eta | P68510 |
| 14-3-3 protein gamma | P61982 |
| 14-3-3 protein theta | P68254 |
| 14-3-3 protein zeta/delta | P63101 |
| 182 kDa tankyrase-1-binding protein | P58871 |
| 1-phosphatidylinositol 4,5-bisphosphate phosphod | Q9Z1B3 |
| 1-phosphatidylinositol 4,5-bisphosphate phosphod | Q8R3B1 |
| 1-phosphatidylinositol 4,5-bisphosphate phosphod | A2AP18 |
| 1-phosphatidylinositol 4,5-bisphosphate phosphod | Q62077 |
| 2',3'-cyclic-nucleotide 3'-phosphodiesterase | P16330 |
| 2,4-dienoyl-CoA reductase, mitochondrial | Q9CQ62 |
| 26S protease regulatory subunit 10B | P62334 |
| 26S protease regulatory subunit 4 | P62192 |
| 26S protease regulatory subunit 6A | O88685 |
| 26S protease regulatory subunit 7 | P46471 |
| 26S protease regulatory subunit 8 | P62196 |
| 26S proteasome non-ATPase regulatory subunit 1 | Q3TXS7 |
| 26S proteasome non-ATPase regulatory subunit 1 | Q8BG32 |
| 26S proteasome non-ATPase regulatory subunit 1 | Q9WVJ2 |
| 26S proteasome non-ATPase regulatory subunit 2 | Q8VDM4 |
| 26S proteasome non-ATPase regulatory subunit 3 | P14685 |
| 26S proteasome non-ATPase regulatory subunit 5 | Q8BJY1 |
| 26S proteasome non-ATPase regulatory subunit 8 | Q9CX56 |
| 26S proteasome non-ATPase regulatory subunit 9 | Q9CR00 |
| 28 kDa heat- and acid-stable phosphoprotein | Q3UHX2 |
| 2-aminoethanethiol dioxygenase | Q6PDY2 |
| 2-oxoglutarate dehydrogenase, mitochondrial | Q60597 |
| 39S ribosomal protein L9, mitochondrial | Q99N94 |
| 3-hydroxyacyl-CoA dehydrogenase type-2 | O08756 |
| 3-hydroxyisobutyrate dehydrogenase, mitochondri | Q99L13 |
| 3-ketoacyl-CoA thiolase, mitochondrial | Q8BWT1 |
| 3-mercaptopyruvate sulfurtransferase | Q99J99 |
| 40S ribosomal protein S10 | P63325 |
| 40S ribosomal protein S13 | P62301 |
| 40S ribosomal protein S14 | P62264 |
| 40S ribosomal protein S16 | P14131 |
| 40S ribosomal protein S18 | P62270 |
| 40S ribosomal protein S19 | Q9CZX8 |
| 40S ribosomal protein S2 | P25444 |
| 40S ribosomal protein S20 | P60867 |
| 40S ribosomal protein S21 | Q9CQR2 |
| 40S ribosomal protein S25 | P62852 |
| 40S ribosomal protein S27 | Q6ZWU9 |
| 40S ribosomal protein S27-like | Q6ZWY3 |
| 40S ribosomal protein S28 | P62858 |
| 40S ribosomal protein S3 | P62908 |
| 40S ribosomal protein S3a | P97351 |
| 40S ribosomal protein S4, X isoform | P62702 |
| 40S ribosomal protein S5 | P97461 |
| 40S ribosomal protein S6 | P62754 |
| 40S ribosomal protein S7 | P62082 |
| 40S ribosomal protein S8 | P62242 |
| 40S ribosomal protein S9 | Q6ZWN5 |
| 40S ribosomal protein SA | P14206 |
| 4-aminobutyrate aminotransferase, mitochondrial | P61922 |
| 4F2 cell-surface antigen heavy chain | P10852 |
| 4-trimethylaminobutyraldehyde dehydrogenase | Q9JLJ2 |
| 5'-AMP-activated protein kinase catalytic subunit a | Q5EG47 |
| 5'-nucleotidase domain-containing protein 3 | Q3UHB1 |
| 5-oxoprolinase | Q8K010 |
| 60 kDa heat shock protein, mitochondrial | P63038 |
| 60 kDa SS-A/Ro ribonucleoprotein | O08848 |
| 60S acidic ribosomal protein P2 | P99027 |
| 60S ribosomal protein L13 | P47963 |
| 60S ribosomal protein L14 | Q9CR57 |
| 60S ribosomal protein L17 | Q9CPR4 |
| 60S ribosomal protein L18 | P35980 |
| 60S ribosomal protein L18a | P62717 |
| 60S ribosomal protein L19 | P84099 |
| 60S ribosomal protein L22 | P67984 |
| 60S ribosomal protein L23a | P62751 |
| 60S ribosomal protein L26 | P61255 |
| 60S ribosomal protein L28 | P41105 |
| 60S ribosomal protein L3 | P27659 |
| 60S ribosomal protein L30 | P62889 |
| 60S ribosomal protein L31 | P62900 |
| 60S ribosomal protein L5 | P47962 |
| 60S ribosomal protein L6 | P47911 |
| 60S ribosomal protein L7 | P14148 |
| 60S ribosomal protein L8 | P62918 |
| 6-phosphofructokinase type C | Q9WUA3 |
| 6-phosphofructokinase, muscle type | P47857 |
| 6-phosphogluconate dehydrogenase, decarboxylat | Q9DCD0 |
| 78 kDa glucose-regulated protein | P20029 |
| Abelson tyrosine-protein kinase 2 | Q4JIM5 |
| Abl interactor 1 | Q8CBW3 |

**Detected at P21 only - 1450 proteins**

| Protein name | Uniprot ID |
|---|---|
| [Protein ADP-ribosylarginine] hydrolase | P54923 |
| [Pyruvate dehydrogenase [acetyl-transferring]]-pho | Q3UV70 |
| [Pyruvate dehydrogenase [lipoamide]] kinase isozy | Q9JK42 |
| 14 kDa phosphohistidine phosphatase | Q9DAK9 |
| 1-acyl-sn-glycerol-3-phosphate acyltransferase ga | Q9D517 |
| 1-phosphatidylinositol 3-phosphate 5-kinase | Q9Z1T6 |
| 1-phosphatidylinositol 4,5-bisphosphate phosphod | P51432 |
| 1-phosphatidylinositol 4,5-bisphosphate phosphod | Q8CIH5 |
| 26S protease regulatory subunit 6B | P54775 |
| 26S proteasome non-ATPase regulatory subunit 1 | Q9D8W5 |
| 26S proteasome non-ATPase regulatory subunit 1 | O35593 |
| 26S proteasome non-ATPase regulatory subunit 4 | O35226 |
| 26S proteasome non-ATPase regulatory subunit 6 | Q99JI4 |
| 26S proteasome non-ATPase regulatory subunit 7 | P26516 |
| 28S ribosomal protein S16, mitochondrial | Q9CPX7 |
| 28S ribosomal protein S2, mitochondrial | Q924T2 |
| 28S ribosomal protein S22, mitochondrial | Q9CXW2 |
| 28S ribosomal protein S23, mitochondrial | Q8VE22 |
| 28S ribosomal protein S26, mitochondrial | Q80ZS3 |
| 28S ribosomal protein S27, mitochondrial | Q8BK72 |
| 28S ribosomal protein S29, mitochondrial | Q9ER88 |
| 28S ribosomal protein S30, mitochondrial | Q9D0G0 |
| 28S ribosomal protein S31, mitochondrial | Q61733 |
| 28S ribosomal protein S5, mitochondrial | Q99N87 |
| 2-C-methyl-D-erythritol 4-phosphate cytidylyltransf | Q5RJG7 |
| 2-hydroxyacylsphingosine 1-beta-galactosyltransfe | Q64676 |
| 2-methoxy-6-polyprenyl-1,4-benzoquinol methylas | Q9CXI0 |
| 3'(2'),5'-bisphosphate nucleotidase 1 | Q9Z0S1 |
| 39S ribosomal protein L1, mitochondrial | Q99N96 |
| 39S ribosomal protein L11, mitochondrial | Q9CQF0 |
| 39S ribosomal protein L12, mitochondrial | Q9DB15 |
| 39S ribosomal protein L15, mitochondrial | Q9CPR5 |
| 39S ribosomal protein L19, mitochondrial | Q9D338 |
| 39S ribosomal protein L2, mitochondrial | Q9D773 |
| 39S ribosomal protein L22, mitochondrial | Q8BU88 |
| 39S ribosomal protein L37, mitochondrial | Q921S7 |
| 39S ribosomal protein L38, mitochondrial | Q8K2M0 |
| 39S ribosomal protein L39, mitochondrial | Q9JKF7 |
| 39S ribosomal protein L40, mitochondrial | Q9Z2Q5 |
| 39S ribosomal protein L41, mitochondrial | Q9CQN7 |
| 39S ribosomal protein L46, mitochondrial | Q9EQI8 |
| 39S ribosomal protein L49, mitochondrial | Q9CQ40 |
| 3-hydroxyacyl-CoA dehydratase 3 | Q8K2C9 |
| 3-hydroxyisobutyryl-CoA hydrolase, mitochondrial | Q8QZS1 |
| 3-phosphoinositide-dependent protein kinase 1 | Q9Z2A0 |
| 40S ribosomal protein S11 | P62281 |
| 40S ribosomal protein S12 | P63323 |
| 40S ribosomal protein S15a | P62245 |
| 40S ribosomal protein S17 | P63276 |
| 40S ribosomal protein S24 | P62849 |
| 40S ribosomal protein S26 | P62855 |
| 40S ribosomal protein S29 | P62274 |
| 5'(3')-deoxyribonucleotidase, cytosolic type | Q9JM14 |
| 5'(3')-deoxyribonucleotidase, mitochondrial | Q8VCE6 |
| 5'-AMP-activated protein kinase catalytic subunit a | Q8BRK8 |
| 5'-AMP-activated protein kinase subunit gamma-1 | O54950 |
| 5'-AMP-activated protein kinase subunit gamma-2 | Q91WG5 |
| 5-azacytidine-induced protein 1 | Q62036 |
| 60S acidic ribosomal protein P0 | P14869 |
| 60S ribosomal protein L10 | Q6ZWV3 |
| 60S ribosomal protein L10a | P53026 |
| 60S ribosomal protein L11 | Q9CXW4 |
| 60S ribosomal protein L12 | P35979 |
| 60S ribosomal protein L13a | P19253 |
| 60S ribosomal protein L15 | Q9CZM2 |
| 60S ribosomal protein L21 | O09167 |
| 60S ribosomal protein L22-like 1 | Q9D7S7 |
| 60S ribosomal protein L24 | Q8BP67 |
| 60S ribosomal protein L27 | P61358 |
| 60S ribosomal protein L27a | P14115 |
| 60S ribosomal protein L32 | P62911 |
| 60S ribosomal protein L35 | Q6ZWV7 |
| 60S ribosomal protein L35a | O55142 |
| 60S ribosomal protein L36 | P47964 |
| 60S ribosomal protein L38 | Q9JJI8 |
| 60S ribosomal protein L4 | Q9D8E6 |
| 60S ribosomal protein L7a | P12970 |
| 60S ribosomal protein L9 | P51410 |
| 6-phosphofructo-2-kinase/fructose-2,6-bisphospha | P70265 |
| 6-phosphofructokinase, liver type | P12382 |
| 6-phosphogluconolactonase | Q9CQ60 |
| Abhydrolase domain-containing protein 14B | Q8VCR7 |
| Abhydrolase domain-containing protein 16A | Q9Z1Q2 |
| Abhydrolase domain-containing protein 4 | Q8VD66 |
| Abl interactor 2 | P62484 |
| Abnormal spindle-like microcephaly-associated pr | Q8CJ27 |
| Acetyl-coenzyme A synthetase, cytoplasmic | Q9QXG4 |
| Acidic leucine-rich nuclear phosphoprotein 32 fami | Q9EST5 |
| Actin, cytoplasmic 2 | P63260 |
| Actin-related protein 10 | Q9QZB7 |
| Actin-related protein 2/3 complex subunit 1B | Q9WV32 |

**Detected in adult only - 94 proteins**

| Protein name | Uniprot ID |
|---|---|
| 5'-3' exoribonuclease 1 | P97789 |
| Actin, cytoplasmic 1 | P60710 |
| Acyl-protein thioesterase 1 | P97823 |
| Alpha-parvin | Q9EPC1 |
| Aspartoacylase | Q8R3P0 |
| ATP-binding cassette sub-family B member 8, mit | Q9CXJ4 |
| BRISC complex subunit Abro1 | Q3TCJ1 |
| Calmin | Q8C5W0 |
| Calpain-1 catalytic subunit | O35350 |
| Carbonyl reductase family member 4 | Q91VT4 |
| Cathepsin S | O70370 |
| Cleavage stimulation factor subunit 2 | Q8BIQ5 |
| C-Myc-binding protein | Q9EQS3 |
| Coiled-coil domain-containing protein 9 | Q8VC31 |
| COMM domain-containing protein 9 | Q8K2Q0 |
| COP9 signalosome complex subunit 8 | Q8VBV7 |
| Dickkopf-related protein 3 | Q9QUN9 |
| DNA-directed RNA polymerases I, II, and III subun | Q923G2 |
| DnaJ homolog subfamily B member 12 | Q9QYI4 |
| DnaJ homolog subfamily C member 5 | P60904 |
| E3 SUMO-protein ligase RanBP2 | Q9ERU9 |
| E3 ubiquitin-protein ligase RBX1 | P62878 |
| Emerin | O08579 |
| Epidermal growth factor receptor kinase substrate | Q08509 |
| FXYD domain-containing ion transport regulator 7 | P59648 |
| Glutamate receptor 4 | Q9Z2W8 |
| GMP reductase 2 | Q99L27 |
| HEAT repeat-containing protein 5B | Q8C547 |
| Histone H2B type 1-M | P10854 |
| Hsp90 co-chaperone Cdc37-like 1 | Q9CZP7 |
| Hyaluronan and proteoglycan link protein 4 | Q80WM4 |
| Inositol polyphosphate 5-phosphatase K | Q8C5L6 |
| Integrin beta-2 | P11835 |
| Junctional adhesion molecule C | Q9D8B7 |
| Kininogen-1 | O08677 |
| Methyl-CpG-binding protein 2 | Q9Z2D6 |
| Monocarboxylate transporter 1 | P53986 |
| Myosin-11 | O08638 |
| NADP-dependent malic enzyme, mitochondrial | Q8BMF3 |
| Netrin receptor UNC5A | Q8K1S4 |
| Neuroligin-1 | Q99K10 |
| NHS-like protein 2 | B1AXH1 |
| NmrA-like family domain-containing protein 1 | Q8K2T1 |
| Opioid growth factor receptor-like protein 1 | Q8VE52 |
| Phospholipase DDHD2 | Q80Y98 |
| Piezo-type mechanosensitive ion channel compon | Q8CD54 |
| Polypeptide N-acetylgalactosaminyltransferase 2 | Q6PB93 |
| Potassium voltage-gated channel subfamily D mer | Q9Z0V1 |
| Potassium voltage-gated channel subfamily KQT r | Q9Z351 |
| Protein dpy-30 homolog | Q99LT0 |
| Protein FAM131B | Q3TY60 |
| Protein FAM177A1 | Q8BR63 |
| Protein NipSnap homolog 2 | O55126 |
| Protein phosphatase 1 regulatory subunit 12B | Q8BG95 |
| Protein phosphatase 1 regulatory subunit 1B | Q60829 |
| Protein-glutamine gamma-glutamyltransferase 2 | P21981 |
| Protein VAC14 homolog | Q80WQ2 |
| Ras-related protein Rab-2B | P59279 |
| Regulator of microtubule dynamics protein 3 | Q3UJU9 |
| Rho-related GTP-binding protein RhoG | P84096 |
| RNA-binding protein 25 | B2RY56 |
| Septin-4 | P28661 |
| Sequestosome-1 | Q64337 |
| Serine protease inhibitor A3K | P07759 |
| Serine/arginine repetitive matrix protein 2 | Q8BTI8 |
| Serine/threonine-protein kinase 25 | Q9Z2W1 |
| Serine/threonine-protein kinase PRP4 homolog | Q61136 |
| Serine/threonine-protein phosphatase 2A 55 kDa r | Q6ZWR4 |
| Serine/threonine-protein phosphatase 6 regulatory | Q7TSI3 |
| SLIT and NTRK-like protein 5 | Q810B7 |
| Spermatogenesis-associated protein 2-like protein | Q8BNN1 |
| Sphingomyelin phosphodiesterase 3 | Q9JJY3 |
| Splicing factor U2AF 35 kDa subunit | Q9D883 |
| Synaptic vesicle 2-related protein | Q8BFT9 |
| Synaptic vesicle glycoprotein 2C | Q69ZS6 |
| Tax_Id=10090 Gene_Symbol=Krt33b keratin comp | S.avidinii |
| Tight junction protein ZO-1 | P39447 |
| Tissue factor | P20352 |
| T-lymphoma invasion and metastasis-inducing pro | Q6ZPF3 |
| Transmembrane protein 132A | Q922P8 |
| tRNA methyltransferase 112 homolog | Q9DCG9 |
| Tudor and KH domain-containing protein | Q80VL1 |
| Ubiquitin domain-containing protein UBFD1 | Q78JW9 |
| Ubiquitin-conjugating enzyme E2 variant 2 | Q9D2M8 |
| Ubiquitin-fold modifier 1 | P61961 |
| Ubiquitin-like modifier-activating enzyme 5 | Q8VE47 |
| Uncharacterized protein C6orf203 homolog | Q9CQF4 |
| Uncharacterized protein KIAA0090 | Q8C7X2 |
| Uncharacterized protein KIAA1211 | Q5PR69 |
| UPF0449 protein C19orf25 homolog | Q9D7E4 |
| UPF0515 protein C19orf66 homolog | Q8CAK3 |

| | |
|---|---|
| Acetoacetyl-CoA synthetase | Q9D2R0 |
| Acetyl-CoA acetyltransferase, cytosolic | Q8CAY6 |
| Acetyl-CoA acetyltransferase, mitochondrial | Q8QZT1 |
| Acetyl-CoA carboxylase 1 | Q5SWU9 |
| Acetyl-coenzyme A synthetase 2-like, mitochondria | Q99NB1 |
| Acid ceramidase | Q9WV54 |
| Acidic leucine-rich nuclear phosphoprotein 32 fami | O35381 |
| Aconitate hydratase, mitochondrial | Q99KI0 |
| Actin, aortic smooth muscle | P62737 |
| Actin-binding LIM protein 1 | Q8K4G5 |
| Actin-binding LIM protein 2 | Q8BL65 |
| Actin-related protein 2 | P61161 |
| Actin-related protein 2/3 complex subunit 1A | Q9R0Q6 |
| Actin-related protein 2/3 complex subunit 2 | Q9CVB6 |
| Actin-related protein 2/3 complex subunit 4 | P59999 |
| Actin-related protein 3 | Q99JY9 |
| Actin-related protein 3B | Q641P0 |
| Activated RNA polymerase II transcriptional coacti | P11031 |
| Activator of 90 kDa heat shock protein ATPase hor | Q8BK64 |
| Active breakpoint cluster region-related protein | Q5SSL4 |
| Acylamino-acid-releasing enzyme | Q8R146 |
| Acyl-CoA dehydrogenase family member 9, mitocl | Q8JZN5 |
| Acyl-CoA synthetase family member 2, mitochondi | Q8VCW8 |
| Acyl-CoA synthetase family member 3, mitochondi | Q3URE1 |
| Acyl-CoA-binding domain-containing protein 6 | Q9D061 |
| Acyl-CoA-binding protein | P31786 |
| Acyl-coenzyme A thioesterase 2, mitochondrial | Q9QYR9 |
| Acyl-coenzyme A thioesterase 9, mitochondrial | Q9R0X4 |
| Acylphosphatase-2 | P56375 |
| Adapter molecule crk | Q64010 |
| Adaptin ear-binding coat-associated protein 1 | Q9CR95 |
| Adenine phosphoribosyltransferase | P08030 |
| Adenomatous polyposis coli protein | Q61315 |
| Adenosine kinase | P55264 |
| Adenosylhomocysteinase | P50247 |
| Adenylate cyclase type 9 | P51830 |
| Adenylate kinase isoenzyme 1 | Q9R0Y5 |
| Adenylate kinase isoenzyme 5 | Q920P5 |
| Adenylosuccinate synthetase isozyme 2 | P46664 |
| Adenylyl cyclase-associated protein 1 | P40124 |
| Adenylyl cyclase-associated protein 2 | Q9CYT6 |
| Adipocyte plasma membrane-associated protein | Q9D7N9 |
| ADP/ATP translocase 1 | P48962 |
| ADP-ribosylation factor 2 | Q8BSL7 |
| ADP-ribosylation factor 5 | P84084 |
| ADP-ribosylation factor GTPase-activating protein | Q9EPJ9 |
| ADP-ribosylation factor-like protein 8A | Q8VEH3 |
| ADP-ribosylation factor-like protein 8B | Q9CQW2 |
| Afadin | Q9QZQ1 |
| AFG3-like protein 2 | Q8JZQ2 |
| Aftiphilin | Q80WT5 |
| Aggrecan core protein | Q61282 |
| AH receptor-interacting protein | O08915 |
| A-kinase anchor protein 5 | D3YVF0 |
| A-kinase anchor protein 9 | Q70FJ1 |
| A-kinase anchor protein SPHKAP | Q6NSW3 |
| Alanine aminotransferase 1 | Q8QZR5 |
| Alanine--tRNA ligase, cytoplasmic | Q8BGQ7 |
| Alanyl-tRNA editing protein Aarsd1 | Q3THG9 |
| Alcohol dehydrogenase [NADP(+)] | Q9JII6 |
| Alcohol dehydrogenase class-3 | P28474 |
| Aldehyde dehydrogenase X, mitochondrial | Q9CZS1 |
| Aldehyde dehydrogenase, mitochondrial | P47738 |
| Aldose reductase | P45376 |
| Alpha-2-macroglobulin | Q61838 |
| Alpha-2-macroglobulin receptor-associated proteir | P55302 |
| Alpha-actinin-1 | Q7TPR4 |
| Alpha-actinin-2 | Q9JI91 |
| Alpha-actinin-4 | P57780 |
| Alpha-adducin | Q9QYC0 |
| Alpha-aminoadipic semialdehyde dehydrogenase | Q9DBF1 |
| Alpha-centractin | P61164 |
| Alpha-endosulfine | P60840 |
| Alpha-enolase | P17182 |
| Alpha-internexin | P46660 |
| Alpha-soluble NSF attachment protein | Q9DB05 |
| Alpha-taxilin | Q6PAM1 |
| Amine oxidase [flavin-containing] A | Q64133 |
| Amine oxidase [flavin-containing] B | Q8BW75 |
| Aminoacyl tRNA synthase complex-interacting mu | P31230 |
| Aminoacylase-1 | Q99JW2 |
| Aminopeptidase B | Q8VCT3 |
| AMP deaminase 2 | Q9DBT5 |
| AMP deaminase 3 | O08739 |
| Amphiphysin | Q7TQF7 |
| Amyloid beta A4 precursor protein-binding family A | B2RUJ5 |
| Amyloid beta A4 precursor protein-binding family E | Q9QXJ1 |
| Amyloid beta A4 protein | P12023 |
| Amyloid-like protein 1 | Q03157 |
| Amyloid-like protein 2 | Q06335 |
| Ankyrin repeat and MYND domain-containing prote | Q3TPE9 |
| Ankyrin repeat and sterile alpha motif domain-cont | Q8BIZ1 |
| Ankyrin-2 | Q8C8R3 |
| Annexin A11 | P97384 |
| Actin-related protein 2/3 complex subunit 3 | Q9JM76 |
| Actin-related protein 2/3 complex subunit 5 | Q9CPW4 |
| Actin-related protein 2/3 complex subunit 5-like pro | Q9D898 |
| Acyl carrier protein, mitochondrial | Q9CR21 |
| Acyl-coenzyme A thioesterase 1 | O55137 |
| Acyl-coenzyme A thioesterase 13 | Q9CQR4 |
| Acylglycerol kinase, mitochondrial | Q9ESW4 |
| Acylphosphatase-1 | P56376 |
| Acyl-protein thioesterase 2 | Q9WTL7 |
| Adenosine deaminase | P03958 |
| Adenylate kinase 2, mitochondrial | Q9WTP6 |
| Adenylate kinase isoenzyme 4, mitochondrial | Q9WUR9 |
| Adenylosuccinate lyase | P54822 |
| Adenylyltransferase and sulfurtransferase MOCS3 | A2BDX3 |
| ADP/ATP translocase 2 | P51881 |
| ADP/ATP translocase 4 | Q3V132 |
| ADP-ribose pyrophosphatase, mitochondrial | Q8BVU5 |
| ADP-ribosylation factor 3 | P61205 |
| ADP-ribosylation factor 4 | P61750 |
| ADP-ribosylation factor 6 | P62331 |
| ADP-ribosylation factor GTPase-activating protein | Q99K28 |
| ADP-ribosylation factor-binding protein GGA1 | Q8R0H9 |
| ADP-ribosylation factor-binding protein GGA3 | Q8BMI3 |
| ADP-ribosylation factor-like protein 3 | Q9WUL7 |
| Adrenodoxin-like protein, mitochondrial | Q9CPW2 |
| Aflatoxin B1 aldehyde reductase member 2 | Q8CG76 |
| Agrin | A2ASQ1 |
| A-kinase anchor protein 10, mitochondrial | O88845 |
| A-kinase anchor protein 12 | Q9WTQ5 |
| Alkyldihydroxyacetonephosphate synthase, peroxis | Q8C0I1 |
| Alpha-1,3/1,6-mannosyltransferase ALG2 | Q9DBE8 |
| Alpha-1-antitrypsin 1-4 | Q00897 |
| Alpha-1-syntrophin | Q61234 |
| Alpha-2-HS-glycoprotein | P29699 |
| Alpha-ketoglutarate-dependent dioxygenase FTO | Q8BGW1 |
| Alpha-mannosidase 2C1 | Q91W89 |
| Alpha-synuclein | O55042 |
| Alpha-tubulin N-acetyltransferase | Q8K341 |
| Alsin | Q920R0 |
| Aminoacyl tRNA synthase complex-interacting mu | Q8R010 |
| AMSH-like protease | Q76N33 |
| Amyloid beta A4 precursor protein-binding family A | P98084 |
| AN1-type zinc finger protein 2B | Q91X58 |
| Anaphase-promoting complex subunit 2 | Q8BZQ7 |
| Anaphase-promoting complex subunit 4 | Q91W96 |
| Ankyrin repeat and FYVE domain-containing prote | Q810B6 |
| Ankyrin repeat domain-containing protein 13D | Q6PD24 |
| Ankyrin repeat domain-containing protein 17 | Q99NH0 |
| Ankyrin repeat domain-containing protein 32 | Q8R3P9 |
| Ankyrin-1 | Q02357 |
| AP-1 complex subunit mu-1 | P35585 |
| AP-1 complex subunit sigma-1A | P61967 |
| AP-2 complex subunit sigma | P62743 |
| AP-3 complex subunit beta-1 | Q9Z1T1 |
| AP-3 complex subunit mu-1 | Q9JKC8 |
| AP-3 complex subunit mu-2 | Q8R2R9 |
| AP-3 complex subunit sigma-2 | Q8BSZ2 |
| APC membrane recruitment protein 2 | Q8CCJ4 |
| Apoptosis regulator BAX | Q07813 |
| Apoptosis-stimulating of p53 protein 1 | Q62415 |
| Arf-GAP domain and FG repeat-containing protein | Q8K2K6 |
| Arginyl-tRNA--protein transferase 1 | Q9Z2A5 |
| Armadillo repeat protein deleted in velo-cardio-faci | P98203 |
| Armadillo repeat-containing protein 1 | Q9D7A8 |
| Armadillo repeat-containing X-linked protein 3 | Q8BHS6 |
| Aromatic-L-amino-acid decarboxylase | O88533 |
| Arylsulfatase B | P50429 |
| Aspartate--tRNA ligase, mitochondrial | Q8BIP0 |
| Ataxin-2 | O70305 |
| ATP synthase subunit e, mitochondrial | Q06185 |
| ATP synthase subunit epsilon, mitochondrial | P56382 |
| ATP synthase subunit f, mitochondrial | P56135 |
| ATP synthase subunit g, mitochondrial | Q9CPQ8 |
| ATPase Asna1 | O54984 |
| ATPase family AAA domain-containing protein 1 | Q9D5T0 |
| ATPase WRNIP1 | Q91XU0 |
| ATP-binding cassette sub-family A member 2 | P41234 |
| ATP-binding cassette sub-family A member 7 | Q91V24 |
| ATP-binding cassette sub-family D member 2 | Q61285 |
| ATP-binding cassette sub-family D member 3 | P55096 |
| ATP-binding cassette sub-family F member 2 | Q99LE6 |
| ATP-binding cassette sub-family F member 3 | Q8K268 |
| ATP-dependent Clp protease ATP-binding subunit | Q9JHS4 |
| ATP-dependent RNA helicase A | O70133 |
| ATP-dependent RNA helicase DDX3X | Q62167 |
| ATP-dependent RNA helicase Dhx29 | Q6PGC1 |
| Autophagy protein 5 | Q99J83 |
| BAG family molecular chaperone regulator 1 | Q60739 |
| BAG family molecular chaperone regulator 2 | Q91YN9 |
| BAG family molecular chaperone regulator 4 | Q8CI61 |
| Barrier-to-autointegration factor | O54962 |
| Basic leucine zipper and W2 domain-containing pr | Q2L4X1 |
| B-cell receptor-associated protein 29 | Q61334 |
| B-cell receptor-associated protein 31 | Q61335 |
| Vesicle transport through interaction with t-SNARE | O89116 |
| Voltage-dependent L-type calcium channel subuni | Q8CC27 |
| Zinc transporter 9 | Q5IRJ6 |

| | | | |
|---|---|---|---|
| Annexin A3 | O35639 | Bcl-2-like protein 1 | Q64373 |
| Annexin A5 | P48036 | Beta-adrenergic receptor kinase 2 | Q3UYH7 |
| Annexin A6 | P14824 | Beta-crystallin B1 | Q9WVJ5 |
| Annexin A7 | Q07076 | Beta-galactosidase | P23780 |
| Anoctamin-1 | Q8BHY3 | Beta-glucuronidase | P12265 |
| AP-1 complex subunit beta-1 | O35643 | Beta-lactamase-like protein 2 | Q99KR3 |
| AP-1 complex subunit gamma-1 | P22892 | Bifunctional arginine demethylase and lysyl-hydrox | Q9ERI5 |
| AP-2 complex subunit alpha-1 | P17426 | Biorientation of chromosomes in cell division prote | Q5SQY2 |
| AP-2 complex subunit alpha-2 | P17427 | Bis(5'-nucleosyl)-tetraphosphatase [asymmetrical] | P56380 |
| AP-2 complex subunit beta | Q9DBG3 | Bisphosphoglycerate mutase | P15327 |
| AP-2 complex subunit mu | P84091 | BolA-like protein 2 | Q8BGS2 |
| AP2-associated protein kinase 1 | Q3UHJ0 | Brain protein 44-like protein | P63030 |
| AP-3 complex subunit beta-2 | Q9JME5 | Brain-specific angiogenesis inhibitor 1 | Q3UHD1 |
| AP-3 complex subunit delta-1 | O54774 | Brain-specific angiogenesis inhibitor 2 | Q8CGM1 |
| AP-3 complex subunit sigma-1 | Q9DCR2 | Brain-specific angiogenesis inhibitor 3 | Q80ZF8 |
| Apolipoprotein A-I | Q00623 | Branched-chain-amino-acid aminotransferase, cyto | P24288 |
| Apolipoprotein A-IV | P06728 | Breast cancer anti-estrogen resistance protein 1 | Q61140 |
| Apolipoprotein E | P08226 | Breast carcinoma-amplified sequence 3 homolog | Q8CCN5 |
| Apolipoprotein O | Q9DCZ4 | BRI3-binding protein | Q8BXV2 |
| Apoptosis inhibitor 5 | O35841 | BRO1 domain-containing protein BROX | Q8K2Q7 |
| Apoptosis-inducing factor 1, mitochondrial | Q9Z0X1 | BTB/POZ domain-containing protein 17 | Q9DB72 |
| Aquaporin-4 | P55088 | BTB/POZ domain-containing protein KCTD16 | Q5DTY9 |
| ARF GTPase-activating protein GIT1 | Q68FF6 | BTB/POZ domain-containing protein KCTD8 | Q50H33 |
| Arfaptin-2 | Q8K221 | C2 domain-containing protein 3 | Q52KB6 |
| Arf-GAP with coiled-coil, ANK repeat and PH doma | Q6ZQK5 | CAAX prenyl protease 1 homolog | Q80W54 |
| Arf-GAP with GTPase, ANK repeat and PH domain | Q8BXK8 | Cadherin-10 | P70408 |
| Arf-GAP with GTPase, ANK repeat and PH domain | Q3UHD9 | Cadherin-2 | P15116 |
| Arf-GAP with GTPase, ANK repeat and PH domain | Q8VHH5 | Cadherin-8 | P97291 |
| Arf-GAP with SH3 domain, ANK repeat and PH do | Q9QWY8 | Cadherin-9 | P70407 |
| Arf-GAP with SH3 domain, ANK repeat and PH do | Q7SIG6 | Calcineurin-like phosphoesterase domain-containi | Q8BFS6 |
| Arginine and glutamate-rich protein 1 | Q3UL36 | Calcipressin-1 | Q9JHG6 |
| Arginine--tRNA ligase, cytoplasmic | Q9D0I9 | Calcipressin-2 | Q9JHG2 |
| Argininosuccinate lyase | Q91YI0 | Calcium uptake protein 1, mitochondrial | Q8VCX5 |
| Argininosuccinate synthase | P16460 | Calcium/calmodulin-dependent protein kinase II inl | Q6QWF9 |
| Armadillo repeat-containing protein 6 | Q8BNU0 | Calcium/calmodulin-dependent protein kinase kina | Q8VBY2 |
| Asparagine synthetase [glutamine-hydrolyzing] | Q61024 | Calcium/calmodulin-dependent protein kinase type | Q91YS8 |
| Asparagine--tRNA ligase, cytoplasmic | Q8BP47 | Calcium/calmodulin-dependent protein kinase type | Q9QYK9 |
| Aspartate aminotransferase, cytoplasmic | P05201 | Calcium/calmodulin-dependent protein kinase type | Q6PHZ2 |
| Aspartate aminotransferase, mitochondrial | P05202 | Calcium-activated potassium channel subunit alph | Q08460 |
| Aspartate--tRNA ligase, cytoplasmic | Q922B2 | Calcium-binding mitochondrial carrier protein SCa | Q6GQS1 |
| Aspartyl aminopeptidase | Q9Z2W0 | Calcium-binding protein 39 | Q06138 |
| Astrocytic phosphoprotein PEA-15 | Q62048 | Calcium-independent phospholipase A2-gamma | Q8K1N1 |
| Astrotactin-1 | Q61137 | Calcium-transporting ATPase type 2C member 1 | Q80XR2 |
| Ataxin-10 | P28658 | Calmodulin-regulated spectrin-associated protein : | Q80VC9 |
| Ataxin-2-like protein | Q7TQH0 | Calponin-3 | Q9DAW9 |
| Atlastin-1 | Q8BH66 | Calsyntenin-3 | Q99JH7 |
| Atlastin-2 | Q6PA06 | cAMP-dependent protein kinase catalytic subunit b | P68181 |
| ATP synthase subunit alpha, mitochondrial | Q03265 | cAMP-dependent protein kinase inhibitor alpha | P63248 |
| ATP synthase subunit b, mitochondrial | Q9CQQ7 | cAMP-dependent protein kinase type II-beta regula | P31324 |
| ATP synthase subunit beta, mitochondrial | P56480 | cAMP-regulated phosphoprotein 21 | Q9DCB4 |
| ATP synthase subunit d, mitochondrial | Q9DCX2 | CAP-Gly domain-containing linker protein 3 | B9EHT4 |
| ATP synthase subunit delta, mitochondrial | Q9D3D9 | Cap-specific mRNA (nucleoside-2'-O-)-methyltrans | Q9DBC3 |
| ATP synthase subunit gamma, mitochondrial | Q91VR2 | Carboxymethylenebutenolidase homolog | Q8R1G2 |
| ATP synthase subunit O, mitochondrial | Q9DB20 | Carboxypeptidase Q | Q9WVJ3 |
| ATP synthase-coupling factor 6, mitochondrial | P97450 | Carnitine O-acetyltransferase | P47934 |
| ATPase family AAA domain-containing protein 3 | Q925I1 | Carnitine O-palmitoyltransferase 1, brain isoform | Q8BGD5 |
| ATPase inhibitor, mitochondrial | O35143 | Carnitine O-palmitoyltransferase 2, mitochondrial | P52825 |
| ATP-binding cassette sub-family E member 1 | P61222 | Cartilage acidic protein 1 | Q8R555 |
| ATP-binding cassette sub-family F member 1 | Q6P542 | Casein kinase I isoform alpha | Q8BK63 |
| ATP-citrate synthase | Q91V92 | Casein kinase I isoform delta | Q9DC28 |
| ATP-dependent (S)-NAD(P)H-hydrate dehydratase | Q9CZ42 | Casein kinase I isoform epsilon | Q9JMK2 |
| ATP-dependent RNA helicase DDX1 | Q91VR5 | Casein kinase I isoform gamma-1 | Q8BTH8 |
| ATP-dependent RNA helicase DDX19A | Q61655 | Casein kinase II subunit alpha' | O54833 |
| Autophagy-related protein 16-1 | Q8C0J2 | Casein kinase II subunit beta | P67871 |
| Autophagy-related protein 2 homolog B | Q80XK6 | Caspase-3 | P70677 |
| Baculoviral IAP repeat-containing protein 6 | O88738 | Catalase | P24270 |
| BAG family molecular chaperone regulator 5 | Q8CI32 | Catenin delta-1 | P30999 |
| Band 4.1-like protein 1 | Q9Z2H5 | Cathepsin B | P10605 |
| Band 4.1-like protein 2 | O70318 | Cathepsin L1 | P06797 |
| Band 4.1-like protein 3 | Q9WV92 | Cation-independent mannose-6-phosphate recepto | Q07113 |
| Basic leucine zipper and W2 domain-containing pr | Q9CQC6 | CCR4-NOT transcription complex subunit 1 | Q6ZQ08 |
| Basigin | P18572 | CCR4-NOT transcription complex subunit 3 | Q8K0V4 |
| Bcl-2-associated transcription factor 1 | Q8K019 | CDGSH iron-sulfur domain-containing protein 1 | Q91WS0 |
| Bcl-2-like protein 13 | P59017 | CDGSH iron-sulfur domain-containing protein 2 | Q9CQB5 |
| BDNF/NT-3 growth factors receptor | P15209 | CDP-diacylglycerol--glycerol-3-phosphate 3-phosp | Q8BHF7 |
| Beta-adducin | Q9QYB8 | Cell adhesion molecule 4 | Q8R464 |
| Beta-adrenergic receptor kinase 1 | Q99MK8 | Cell division control protein 42 homolog | P60766 |
| Beta-arrestin-1 | Q8BWG8 | Cell division cycle protein 23 homolog | Q8BGZ4 |
| Beta-centractin | Q8R5C5 | Cell division cycle protein 27 homolog | A2A6Q5 |
| Beta-hexosaminidase subunit alpha | P29416 | Cell surface glycoprotein MUC18 | Q8R2Y2 |
| Beta-hexosaminidase subunit beta | P20060 | Centrin-2 | Q9R1K9 |
| Beta-soluble NSF attachment protein | P28663 | Centromere protein V | Q9CXS4 |
| Beta-synuclein | Q91ZZ3 | Centromere-associated protein E | Q6RT24 |
| Bifunctional 3'-phosphoadenosine 5'-phosphosulfa | Q60967 | Centrosomal protein of 128 kDa | Q8BI22 |
| Bifunctional ATP-dependent dihydroxyacetone kina | Q8VC30 | Centrosomal protein of 290 kDa | Q6A078 |
| Bifunctional coenzyme A synthase | Q9DBL7 | Cerebellin-1 | Q9R171 |
| Bifunctional glutamate/proline--tRNA ligase | Q8CGC7 | Charged multivesicular body protein 1a | Q921W0 |
| Bifunctional protein NCOAT | Q9EQQ9 | Charged multivesicular body protein 1b-1 | Q99LU0 |
| Bifunctional purine biosynthesis protein PURH | Q9CWJ9 | Charged multivesicular body protein 2a | Q9DB34 |
| Biliverdin reductase A | Q9CY64 | Charged multivesicular body protein 3 | Q9CQ10 |
| Bleomycin hydrolase | Q8R016 | Charged multivesicular body protein 7 | Q8R1T1 |
| BolA-like protein 1 | Q9D8S9 | Chitinase domain-containing protein 1 | Q922Q9 |
| Brain acid soluble protein 1 | Q91XV3 | Chloride intracellular channel protein 1 | Q9Z1Q5 |
| Brain-enriched guanylate kinase-associated protei | Q68EF6 | Chloride intracellular channel protein 4 | Q9QYB1 |

| | | | |
|---|---|---|---|
| Brain-specific angiogenesis inhibitor 1-associated | Q8BKX1 | Chloride transport protein 6 | O35454 |
| Breakpoint cluster region protein | Q6PAJ1 | Cholecystokinin | P09240 |
| Breast carcinoma-amplified sequence 1 homolog | Q80YN3 | Cholesterol 24-hydroxylase | Q9WVK8 |
| Brefeldin A-inhibited guanine nucleotide-exchange | Q3UGY8 | Choline O-acetyltransferase | Q03059 |
| Brevican core protein | Q61361 | Choline/ethanolamine kinase | O55229 |
| BTB/POZ domain-containing protein KCTD12 | Q6WVG3 | Choline-phosphate cytidylyltransferase A | P49586 |
| C-1-tetrahydrofolate synthase, cytoplasmic | Q922D8 | Choline-phosphate cytidylyltransferase B | Q811Q9 |
| C2 domain-containing protein 2-like | Q80X80 | Chondroitin sulfate proteoglycan 4 | Q8VHY0 |
| Cadherin-11 | P55288 | Chromodomain-helicase-DNA-binding protein 8 | Q09XV5 |
| Cadherin-13 | Q9WTR5 | Chromogranin-A | P26339 |
| Calbindin | P12658 | Citron Rho-interacting kinase | P49025 |
| Calcineurin subunit B type 1 | Q63810 | Clavesin-1 | Q9D4C9 |
| Calcium uniporter protein, mitochondrial | Q3UMR5 | Clavesin-2 | Q8BG92 |
| Calcium/calmodulin-dependent 3',5'-cyclic nucleoti | Q61481 | Clusterin-associated protein 1 | Q8R3P7 |
| Calcium/calmodulin-dependent 3',5'-cyclic nucleoti | Q01065 | Coatomer subunit beta | Q9JIF7 |
| Calcium/calmodulin-dependent protein kinase kina | Q8C078 | Coatomer subunit delta | Q5XJY5 |
| Calcium/calmodulin-dependent protein kinase type | Q8BW96 | Coatomer subunit gamma-1 | Q9QZE5 |
| Calcium/calmodulin-dependent protein kinase type | P11798 | Cob(I)yrinic acid a,c-diamide adenosyltransferase, | Q9D273 |
| Calcium/calmodulin-dependent protein kinase type | P28652 | Cofilin-2 | P45591 |
| Calcium/calmodulin-dependent protein kinase type | Q923T9 | Coiled-coil and C2 domain-containing protein 1B | Q8BRN9 |
| Calcium/calmodulin-dependent protein kinase type | P08414 | Coiled-coil domain-containing protein 104 | Q8C6E0 |
| Calcium-binding and coiled-coil domain-containing | Q8CGU1 | Coiled-coil domain-containing protein 124 | Q9D8X2 |
| Calcium-binding mitochondrial carrier protein Arala | Q8BH59 | Coiled-coil domain-containing protein 136 | Q3TVA9 |
| Calcium-binding protein p22 | P61022 | Coiled-coil domain-containing protein 22 | Q9JIG7 |
| Calcium-dependent secretion activator 1 | Q80TJ1 | Coiled-coil domain-containing protein 43 | Q9CR29 |
| Calcium-dependent secretion activator 2 | Q8BYR5 | Coiled-coil domain-containing protein 47 | Q9D024 |
| Calcyclin-binding protein | Q9CXW3 | Coiled-coil domain-containing protein 50 | Q810U5 |
| Calmodulin | P62204 | Coiled-coil domain-containing protein 58 | Q8R3Q6 |
| Calmodulin-regulated spectrin-associated protein 1 | A2AHC3 | Coiled-coil domain-containing protein 90B, mitocho | Q8C3X2 |
| Calmodulin-regulated spectrin-associated protein 2 | Q8C1B1 | Coiled-coil-helix-coiled-coil-helix domain-containin | Q9D1L0 |
| Calnexin | P35564 | Collagen type IV alpha-3-binding protein | Q9EQG9 |
| Calpain small subunit 1 | O88456 | COMM domain-containing protein 2 | Q8BXC6 |
| Calpain-2 catalytic subunit | O08529 | COMM domain-containing protein 3 | Q63829 |
| Calpain-5 | O08688 | Complement C1q subcomponent subunit B | P14106 |
| Calreticulin | P14211 | Complement C1q tumor necrosis factor-related pro | Q8R066 |
| Calretinin | Q08331 | Complement component 1 Q subcomponent-bindi | O35658 |
| Calsyntenin-1 | Q9EPL2 | Complement regulatory protein Crry | Q64735 |
| Calumenin | O35887 | Conserved oligomeric Golgi complex subunit 5 | Q8C0L8 |
| CaM kinase-like vesicle-associated protein | Q3UHL1 | Conserved oligomeric Golgi complex subunit 7 | Q3UM29 |
| cAMP-dependent protein kinase type I-alpha regul | Q9DBC7 | Contactin-6 | Q9JMB8 |
| cAMP-dependent protein kinase type I-beta regula | P12849 | COP9 signalosome complex subunit 5 | O35864 |
| cAMP-dependent protein kinase type II-alpha regu | P12367 | COP9 signalosome complex subunit 6 | O88545 |
| cAMP-specific 3',5'-cyclic phosphodiesterase 4D | Q01063 | Copper transport protein ATOX1 | O08997 |
| CAP-Gly domain-containing linker protein 1 | Q922J3 | Coproporphyrinogen-III oxidase, mitochondrial | P36552 |
| CAP-Gly domain-containing linker protein 2 | Q9Z0H8 | Coronin-2A | Q8C0P5 |
| Caprin-1 | Q60865 | Coronin-2B | Q8BH44 |
| Carbonic anhydrase 2 | P00920 | Corticotropin-releasing factor-binding protein | Q60571 |
| Carbonic anhydrase 4 | Q64444 | Coxsackievirus and adenovirus receptor homolog | P97792 |
| Carbonyl reductase [NADPH] 1 | P48758 | Craniofacial development protein 1 | O88271 |
| Carbonyl reductase [NADPH] 3 | Q8K354 | Creatine kinase M-type | P07310 |
| Carboxypeptidase E | Q00493 | Crk-like protein | P47941 |
| Casein kinase II subunit alpha | Q60737 | C-terminal-binding protein 1 | O88712 |
| Caskin-1 | Q6P9K8 | C-terminal-binding protein 2 | P56546 |
| Catechol O-methyltransferase domain-containing p | Q8BIG7 | CTP synthase 1 | P70698 |
| Catenin alpha-1 | P26231 | CTP synthase 2 | P70303 |
| Catenin alpha-2 | Q61301 | CUB and sushi domain-containing protein 1 | Q923L3 |
| Catenin beta-1 | Q02248 | CUGBP Elav-like family member 2 | Q9Z0H4 |
| Catenin delta-2 | O35927 | CUGBP Elav-like family member 4 | Q7TSY6 |
| Cathepsin D | P18242 | Cullin-1 | Q9WTX6 |
| Cation-dependent mannose-6-phosphate receptor | P24668 | Cullin-2 | Q9D4H8 |
| Caytaxin | Q8BHE3 | Cullin-4A | Q3TCH7 |
| CB1 cannabinoid receptor-interacting protein 1 | Q5M8N0 | Cullin-9 | Q80TT8 |
| CCA tRNA nucleotidyltransferase 1, mitochondrial | Q8K1J6 | Cyclin-dependent kinase 16 | Q04735 |
| CD166 antigen | Q61490 | Cyclin-dependent kinase 17 | Q8K0D0 |
| Cdc42 effector protein 4 | Q9JM96 | Cyclin-dependent kinase 5 | P49615 |
| Cell adhesion molecule 1 | Q8R5M8 | Cyclin-dependent kinase 9 | Q99J95 |
| Cell adhesion molecule 2 | Q8BLQ9 | Cyclin-Y | Q8BGU5 |
| Cell adhesion molecule 3 | Q99N28 | Cystathionine beta-synthase | Q91WT9 |
| Cell cycle exit and neuronal differentiation protein 1 | Q9JKC6 | Cystathionine gamma-lyase | Q8VCN5 |
| Cellular nucleic acid-binding protein | P53996 | Cysteine and histidine-rich domain-containing prot | Q9D1P4 |
| Centrosomal protein of 170 kDa | Q6A065 | Cysteine-rich with EGF-like domain protein 1 | Q91XD7 |
| cGMP-dependent 3',5'-cyclic phosphodiesterase | Q922S4 | Cytochrome b-c1 complex subunit 9 | Q8R1I1 |
| Charged multivesicular body protein 4b | Q9D8B3 | Cytochrome c oxidase subunit 5B, mitochondrial | P19536 |
| Charged multivesicular body protein 5 | Q9D7S9 | Cytochrome c oxidase subunit 7C, mitochondrial | P17665 |
| Charged multivesicular body protein 6 | P0C0A3 | Cytochrome c, somatic | P62897 |
| Choline transporter-like protein 2 | Q8BY89 | Cytochrome c1, heme protein, mitochondrial | Q9D0M3 |
| Chondroitin sulfate proteoglycan 5 | Q71M36 | Cytohesin-1 | Q9QX11 |
| Chromobox protein homolog 3 | P23198 | Cytohesin-3 | O08967 |
| Chromobox protein homolog 5 | Q61686 | Cytoplasmic aconitate hydratase | P28271 |
| Citrate lyase subunit beta-like protein, mitochondri | Q8R4N0 | Cytoplasmic dynein 2 heavy chain 1 | Q45VK7 |
| Citrate synthase, mitochondrial | Q9CZU6 | Cytoplasmic phosphatidylinositol transfer protein 1 | Q8K4R4 |
| C-Jun-amino-terminal kinase-interacting protein 3 | Q9ESN9 | Cytoplasmic polyadenylation element-binding prot | Q7TN99 |
| C-Jun-amino-terminal kinase-interacting protein 4 | Q58A65 | Cytosolic 5'-nucleotidase III-like protein | Q3UFY7 |
| Clathrin coat assembly protein AP180 | Q61548 | Cytosolic carboxypeptidase 1 | Q641K1 |
| Clathrin heavy chain 1 | Q68FD5 | Cytosolic purine 5'-nucleotidase | Q3V1L4 |
| Clathrin interactor 1 | Q99KN9 | D-beta-hydroxybutyrate dehydrogenase, mitochon | Q80XN0 |
| Clathrin light chain A | O08585 | DBF4-type zinc finger-containing protein 2 homolo | Q5SS00 |
| Clathrin light chain B | Q6IRU5 | DCN1-like protein 1 | Q9QZ73 |
| CLIP-associating protein 1 | Q80TV8 | Death-associated protein kinase 1 | Q80YE7 |
| CLIP-associating protein 2 | Q8BRT1 | Dedicator of cytokinesis protein 10 | Q8BZN6 |
| Clusterin | Q06890 | Dedicator of cytokinesis protein 3 | Q8CIQ7 |
| Coactosin-like protein | Q9CQI6 | Dedicator of cytokinesis protein 7 | Q8R1A4 |
| Coatomer subunit alpha | Q8CIE6 | Dedicator of cytokinesis protein 9 | Q8BIK4 |
| Coatomer subunit beta' | O55029 | Dehydrogenase/reductase SDR family member 1 | Q99L04 |

| | | | |
|---|---|---|---|
| Coatomer subunit epsilon | O89079 | Dehydrogenase/reductase SDR family member 4 | Q99LB2 |
| Coatomer subunit gamma-2 | Q9QXK3 | Dehydrogenase/reductase SDR family member 7E | Q99J47 |
| Coatomer subunit zeta-1 | P61924 | Delta-1-pyrroline-5-carboxylate synthase | Q9Z110 |
| Cofilin-1 | P18760 | Density-regulated protein | Q9CQJ6 |
| Coiled-coil and C2 domain-containing protein 1A | Q8K1A6 | Deoxyhypusine synthase | Q3TXU5 |
| Coiled-coil domain-containing protein 127 | Q3TC33 | Desmoplakin | E9Q557 |
| Coiled-coil domain-containing protein 132 | Q8CI71 | Diablo homolog, mitochondrial | Q9JIQ3 |
| Coiled-coil domain-containing protein 165 | Q3UHU5 | Diacylglycerol kinase epsilon | Q9R1C6 |
| Coiled-coil domain-containing protein 25 | Q78PG9 | Diacylglycerol kinase theta | Q6P5E8 |
| Coiled-coil domain-containing protein 6 | D3YZP9 | Dipeptidyl peptidase 3 | Q99KK7 |
| Coiled-coil domain-containing protein 73 | Q8CDM4 | Dipeptidyl peptidase 8 | Q80YA7 |
| Coiled-coil domain-containing protein 91 | Q9D8L5 | Diphosphomevalonate decarboxylase | Q99JF5 |
| Coiled-coil domain-containing protein 92 | Q8VDN4 | DIS3-like exonuclease 2 | Q8CI75 |
| Coiled-coil domain-containing protein 93 | Q7TQK5 | Disabled homolog 2-interacting protein | Q3UHC7 |
| Coiled-coil-helix-coiled-coil-helix domain-containin | Q9CRB9 | Disco-interacting protein 2 homolog A | Q8BWT5 |
| Coiled-coil-helix-coiled-coil-helix domain-containin | Q91VN4 | Disco-interacting protein 2 homolog B | Q3UH60 |
| Cold shock domain-containing protein E1 | Q91W50 | Disheveled-associated activator of morphogenesis | Q8BPM0 |
| Complexin-1 | P63040 | Disheveled-associated activator of morphogenesis | Q80U19 |
| Complexin-2 | P84086 | Disintegrin and metalloproteinase domain-containi | Q9R1V4 |
| Connector enhancer of kinase suppressor of ras 2 | Q80YA9 | Disks large-associated protein 3 | Q6PFD5 |
| Constitutive coactivator of PPAR-gamma-like prote | Q6A0A9 | DNA topoisomerase 1 | Q04750 |
| Constitutive coactivator of PPAR-gamma-like prote | Q8C3F2 | DNA topoisomerase 2-beta | Q64511 |
| Contactin-1 | P12960 | DNA-binding protein A | Q9JKB3 |
| Contactin-2 | Q61330 | DNA-directed RNA polymerase I subunit RPA1 | O35134 |
| Contactin-associated protein 1 | O54991 | DNA-directed RNA polymerase II subunit RPB2 | Q8CFI7 |
| Contactin-associated protein-like 2 | Q9CPW0 | DnaJ homolog subfamily B member 1 | Q9QYJ3 |
| COP9 signalosome complex subunit 1 | Q99LD4 | DnaJ homolog subfamily B member 14 | Q149L6 |
| COP9 signalosome complex subunit 2 | P61202 | DnaJ homolog subfamily B member 5 | O89114 |
| COP9 signalosome complex subunit 3 | O88543 | DnaJ homolog subfamily C member 10 | Q9DC23 |
| COP9 signalosome complex subunit 4 | O88544 | DnaJ homolog subfamily C member 3 | Q91YW3 |
| COP9 signalosome complex subunit 7a | Q9CZ04 | Dolichol-phosphate mannosyltransferase | O70152 |
| Copine-1 | Q8C166 | Dolichyl-diphosphooligosaccharide--protein glycos | O54734 |
| Copine-2 | P59108 | Dolichyl-diphosphooligosaccharide--protein glycos | P46978 |
| Copine-4 | Q8BLR2 | Dolichyl-diphosphooligosaccharide--protein glycos | Q3TDQ1 |
| Copine-6 | Q9Z140 | Double zinc ribbon and ankyrin repeat-containing p | Q8C008 |
| Copine-7 | Q0VE82 | Down syndrome critical region protein 3 homolog | O35075 |
| Copine-8 | Q9DC53 | Dual specificity mitogen-activated protein kinase k | P31938 |
| Coronin-1A | O89053 | Dual specificity mitogen-activated protein kinase k | Q63932 |
| Coronin-1B | Q9WUM3 | Dual specificity mitogen-activated protein kinase k | P47809 |
| Coronin-1C | Q9WUM4 | Dual specificity mitogen-activated protein kinase k | Q8CE90 |
| Coronin-7 | Q9D2V7 | Dual specificity protein phosphatase 3 | Q9D7X3 |
| Creatine kinase B-type | Q04447 | Dynactin subunit 5 | Q9QZB9 |
| Creatine kinase U-type, mitochondrial | P30275 | Dynamin-2 | P39054 |
| CREB-regulated transcription coactivator 1 | Q68ED7 | Dyslexia-associated protein KIAA0319-like protein | Q8K135 |
| Cullin-3 | Q9JLV5 | Dystrophia myotonica WD repeat-containing prote | Q08274 |
| Cullin-4B | A2A432 | E3 ubiquitin-protein ligase HECTD1 | Q69ZR2 |
| Cullin-5 | Q9D5V5 | E3 ubiquitin-protein ligase HECTD3 | Q3U487 |
| Cullin-associated NEDD8-dissociated protein 1 | Q6ZQ38 | E3 ubiquitin-protein ligase HECW1 | Q8K4P8 |
| Cyclin-dependent kinase 14 | O35495 | E3 ubiquitin-protein ligase HECW2 | Q6I6G8 |
| Cyclin-dependent kinase-like 5 | Q3UTQ8 | E3 ubiquitin-protein ligase MIB2 | Q8R516 |
| Cyclin-G-associated kinase | Q99KY4 | E3 ubiquitin-protein ligase RNF181 | Q9CY62 |
| Cystatin-B | Q62426 | E3 ubiquitin-protein ligase RNF31 | Q924T7 |
| Cystatin-C | P21460 | E3 ubiquitin-protein ligase TRIM23 | Q8BGX0 |
| Cysteine and glycine-rich protein 1 | P97315 | E3 ubiquitin-protein ligase TRIM32 | Q8CH72 |
| Cysteine desulfurase, mitochondrial | Q9Z1J3 | E3 ubiquitin-protein ligase UBR3 | Q5U430 |
| Cysteine sulfinic acid decarboxylase | Q9DBE0 | E3 ubiquitin-protein ligase UBR5 | Q80TP3 |
| Cysteine-rich protein 2 | Q9DCT8 | Echinoderm microtubule-associated protein-like 4 | Q3UMY5 |
| Cysteine--tRNA ligase, cytoplasmic | Q9ER72 | Ectonucleoside triphosphate diphosphohydrolase 2 | O55026 |
| Cytochrome b5 | P56395 | Ectonucleotide pyrophosphatase/phosphodiesteras | Q8BGN3 |
| Cytochrome b5 type B | Q9CQX2 | EF-hand domain-containing family member A2 | Q9CTY5 |
| Cytochrome b-c1 complex subunit 1, mitochondria | Q9CZ13 | EH domain-binding protein 1 | Q69ZW3 |
| Cytochrome b-c1 complex subunit 2, mitochondria | Q9DB77 | ELAV-like protein 2 | Q60899 |
| Cytochrome b-c1 complex subunit 6, mitochondria | P99028 | ELAV-like protein 3 | Q60900 |
| Cytochrome b-c1 complex subunit 7 | Q9D855 | ELAV-like protein 4 | Q61701 |
| Cytochrome b-c1 complex subunit Rieske, mitocho | Q9CR68 | Electroneutral sodium bicarbonate exchanger 1 | Q8JZR6 |
| Cytochrome c oxidase subunit 2 | P00405 | Elongation factor 1-beta | O70251 |
| Cytochrome c oxidase subunit 4 isoform 1, mitoch | P19783 | Elongation factor G, mitochondrial | Q8K0D5 |
| Cytochrome c oxidase subunit 5A, mitochondrial | P12787 | Elongation factor Tu GTP-binding domain-containi | Q8C0D5 |
| Cytochrome c oxidase subunit 6B1 | P56391 | Elongator complex protein 1 | Q7TT37 |
| Cytochrome c oxidase subunit 6C | Q9CPQ1 | Elongator complex protein 3 | Q9CZX0 |
| Cytoplasmic dynein 1 heavy chain 1 | Q9JHU4 | Embigin | P21995 |
| Cytoplasmic dynein 1 intermediate chain 1 | O88485 | Ena/VASP-like protein | P70429 |
| Cytoplasmic dynein 1 intermediate chain 2 | O88487 | Endonuclease domain-containing 1 protein | Q8C522 |
| Cytoplasmic dynein 1 light intermediate chain 1 | Q8R1Q8 | Endoplasmic reticulum metallopeptidase 1 | Q3UVK0 |
| Cytoplasmic dynein 1 light intermediate chain 2 | Q6PDL0 | Endoplasmic reticulum-Golgi intermediate compar | Q9DC16 |
| Cytoplasmic FMR1-interacting protein 1 | Q7TMB8 | Endothelial differentiation-related factor 1 | Q9JMG1 |
| Cytoplasmic FMR1-interacting protein 2 | Q5SQX6 | Enhancer of mRNA-decapping protein 4 | Q3UJB9 |
| Cytoplasmic protein NCK2 | O55033 | Enhancer of rudimentary homolog | P84089 |
| Cytoskeleton-associated protein 4 | Q8BMK4 | Enhancer of yellow 2 transcription factor homolog | Q9JIX0 |
| Cytoskeleton-associated protein 5 | A2AGT5 | Enolase-phosphatase E1 | Q8BGB7 |
| Cytosol aminopeptidase | Q9CPY7 | Enoyl-CoA delta isomerase 2, mitochondrial | Q9WUR2 |
| Cytosolic 10-formyltetrahydrofolate dehydrogenase | Q8R0Y6 | Envoplakin | Q9D952 |
| Cytosolic acyl coenzyme A thioester hydrolase | Q91V12 | Ephexin-1 | Q8CHT1 |
| Cytosolic non-specific dipeptidase | Q9D1A2 | Ephrin type-A receptor 6 | Q62413 |
| Cytospin-B | Q5SXY1 | Ephrin type-B receptor 1 | Q8CBF3 |
| D-3-phosphoglycerate dehydrogenase | Q61753 | Ephrin type-B receptor 2 | P54763 |
| DBIRD complex subunit KIAA1967 homolog | Q8VDP4 | Ephrin-B2 | P52800 |
| DCC-interacting protein 13-alpha | Q8K3H0 | Epidermal growth factor receptor | Q01279 |
| DCC-interacting protein 13-beta | Q8K3G9 | Epidermal growth factor receptor substrate 15 | P42567 |
| DDB1- and CUL4-associated factor 7 | P61963 | EPM2A-interacting protein 1 | Q8VEH5 |
| D-dopachrome decarboxylase | O35215 | Epsin-3 | Q91W69 |
| Dedicator of cytokinesis protein 4 | P59764 | Erlin-1 | Q91X78 |
| Delta(3,5)-Delta(2,4)-dienoyl-CoA isomerase, mito | O35459 | Espin-like protein | Q3UYR4 |
| Delta-1-pyrroline-5-carboxylate dehydrogenase, m | Q8CHT0 | Estradiol 17-beta-dehydrogenase 11 | Q9EQ06 |

| | | | |
|---|---|---|---|
| Delta-aminolevulinic acid dehydratase | P10518 | Estradiol 17-beta-dehydrogenase 12 | O70503 |
| Dematin | Q9WV69 | Estradiol 17-beta-dehydrogenase 8 | P50171 |
| Dendrin | Q80TS7 | Ethylmalonyl-CoA decarboxylase | Q9D9V3 |
| Destrin | Q9R0P5 | Etoposide-induced protein 2.4 | Q61070 |
| Deubiquitinating protein VCIP135 | Q8CDG3 | Eukaryotic initiation factor 4A-III | Q91VC3 |
| Developmentally-regulated GTP-binding protein 1 | P32233 | Eukaryotic peptide chain release factor GTP-bindir | Q8R050 |
| Developmentally-regulated GTP-binding protein 2 | Q9QXB9 | Eukaryotic peptide chain release factor GTP-bindir | Q149F3 |
| Diacylglycerol kinase beta | Q6NS52 | Eukaryotic peptide chain release factor subunit 1 | Q8BWY3 |
| Diacylglycerol kinase gamma | Q91WG7 | Eukaryotic translation initiation factor 1A, X-chromo | Q8BMJ3 |
| Diacylglycerol kinase zeta | Q80UP3 | Eukaryotic translation initiation factor 2 subunit 2 | Q99L45 |
| Dihydrolipoyl dehydrogenase, mitochondrial | O08749 | Eukaryotic translation initiation factor 2A | Q8BJW6 |
| Dihydrolipoyllysine-residue acetyltransferase comp | Q8BMF4 | Eukaryotic translation initiation factor 3 subunit D | O70194 |
| Dihydrolipoyllysine-residue succinyltransferase cor | Q9D2G2 | Eukaryotic translation initiation factor 3 subunit F | Q9DCH4 |
| Dihydropteridine reductase | Q8BVI4 | Eukaryotic translation initiation factor 4E | P63073 |
| Dihydropyrimidinase-related protein 1 | P97427 | Excitatory amino acid transporter 1 | P56564 |
| Dihydropyrimidinase-related protein 2 | O08553 | Exocyst complex component 2 | Q9D4H1 |
| Dihydropyrimidinase-related protein 3 | Q62188 | Exocyst complex component 6 | Q8R313 |
| Dihydropyrimidinase-related protein 4 | O35098 | Exocyst complex component 6B | A6H5Z3 |
| Dihydropyrimidinase-related protein 5 | Q9EQF6 | Exocyst complex component 8 | Q6PGF7 |
| Dipeptidyl aminopeptidase-like protein 6 | Q9Z218 | Exportin-5 | Q924C1 |
| Dipeptidyl peptidase 9 | Q8BVG4 | Exportin-7 | Q9EPK7 |
| Diphosphoinositol polyphosphate phosphohydrolas | Q9JI46 | Exportin-T | Q9CRT8 |
| Diphosphoinositol polyphosphate phosphohydrolas | P0C027 | Extended synaptotagmin-2 | Q3TZZ7 |
| Disintegrin and metalloproteinase domain-containi | Q9R1V6 | FAD synthase | Q8R123 |
| Disintegrin and metalloproteinase domain-containi | Q9R1V7 | Fas apoptotic inhibitory molecule 1 | Q9WUD8 |
| Disks large homolog 1 | Q811D0 | FAS-associated factor 1 | P54731 |
| Disks large homolog 2 | Q91XM9 | Fasciculation and elongation protein zeta-1 | Q8K0X8 |
| Disks large homolog 3 | P70175 | Fascin-2 | Q32M02 |
| Disks large homolog 4 | Q62108 | Fatty acid-binding protein, brain | P51880 |
| Disks large-associated protein 1 | Q9D415 | Fatty acyl-CoA reductase 1 | Q922J9 |
| Disks large-associated protein 2 | Q8BJ42 | Fatty-acid amide hydrolase 1 | O08914 |
| Disks large-associated protein 4 | B1AZP2 | F-box only protein 22 | Q78JE5 |
| DmX-like protein 2 | Q8BPN8 | F-box/SPRY domain-containing protein 1 | Q8K3B1 |
| DNA damage-binding protein 1 | Q3U1J4 | FERM domain-containing protein 4A | Q8BIE6 |
| DNA-(apurinic or apyrimidinic site) lyase | P28352 | Fermitin family homolog 2 | Q8CIB5 |
| DnaJ homolog subfamily A member 1 | P63037 | Ferritin heavy chain | P09528 |
| DnaJ homolog subfamily A member 2 | Q9QYJ0 | Ferrochelatase, mitochondrial | P22315 |
| DnaJ homolog subfamily A member 3, mitochondr | Q99M87 | FGFR1 oncogene partner 2 homolog | Q9CRA9 |
| DnaJ homolog subfamily A member 4 | Q9JMC3 | Fibrinogen C domain-containing protein 1 | A2AV25 |
| DnaJ homolog subfamily B member 11 | Q99KV1 | Fibroblast growth factor 1 | P61148 |
| DnaJ homolog subfamily B member 2 | Q9QYI5 | FK506-binding protein 15 | Q6P9Q6 |
| DnaJ homolog subfamily B member 4 | Q9D832 | Flavin reductase (NADPH) | Q923D2 |
| DnaJ homolog subfamily B member 6 | O54946 | Four and a half LIM domains protein 2 | O70433 |
| DnaJ homolog subfamily C member 11 | Q5U458 | Fractalkine | O35188 |
| DnaJ homolog subfamily C member 7 | Q9QYI3 | FTS and Hook-interacting protein | Q3U2I3 |
| DnaJ homolog subfamily C member 8 | Q6NZB0 | Fucose mutarotase | Q8R2K1 |
| Dolichyl-diphosphooligosaccharide--protein glycos | Q91YQ5 | Fumarylacetoacetase | P35505 |
| Dolichyl-diphosphooligosaccharide--protein glycos | Q9DBG6 | Fumarylacetoacetate hydrolase domain-containing | Q3TC72 |
| Drebrin | Q9QXS6 | FYVE, RhoGEF and PH domain-containing protein | Q91ZT5 |
| Drebrin-like protein | Q62418 | G patch domain-containing protein 8 | A2A6A1 |
| D-tyrosyl-tRNA(Tyr) deacylase 1 | Q9DD18 | G protein-coupled receptor kinase 6 | O70293 |
| Dual specificity mitogen-activated protein kinase k | P70236 | G protein-regulated inducer of neurite outgrowth 3 | Q8BWS5 |
| Dynactin subunit 1 | O08788 | GA-binding protein alpha chain | Q00422 |
| Dynactin subunit 2 | Q99KJ8 | Galectin-related protein A | Q8VED9 |
| Dynactin subunit 3 | Q9Z0Y1 | Gamma-aminobutyric acid receptor subunit alpha- | Q8BHJ7 |
| Dynactin subunit 4 | Q8CBY8 | Gamma-aminobutyric acid receptor subunit beta-2 | P63137 |
| Dynamin-1 | P39053 | Gamma-aminobutyric acid receptor-associated pro | P60521 |
| Dynamin-1-like protein | Q8K1M6 | Gamma-aminobutyric acid type B receptor subunit | Q9WV18 |
| Dynamin-3 | Q8BZ98 | Gamma-glutamylcyclotransferase | Q9D7X8 |
| Dynamin-like 120 kDa protein, mitochondrial | P58281 | Gamma-tubulin complex component 2 | Q921G8 |
| Dynein light chain 1, cytoplasmic | P63168 | Gamma-tubulin complex component 3 | P58854 |
| Dynein light chain 2, cytoplasmic | Q9D0M5 | Gamma-tubulin complex component 4 | Q9D4F8 |
| Dystonin | Q91ZU6 | Gap junction alpha-1 protein | P23242 |
| Dystrobrevin alpha | Q9D2N4 | GDP-D-glucose phosphorylase C15orf58 homolog | Q3TLS3 |
| Dystroglycan | Q62165 | GDP-L-fucose synthase | P23591 |
| Dystrophin | P11531 | GDP-mannose 4,6 dehydratase | Q8K0C9 |
| E3 ubiquitin-protein ligase HUWE1 | Q7TMY8 | Gelsolin | P13020 |
| E3 ubiquitin-protein ligase LRSAM1 | Q80ZI6 | Geranylgeranyl pyrophosphate synthase | Q9WTN0 |
| E3 ubiquitin-protein ligase NEDD4 | P46935 | Glia-derived nexin | Q07235 |
| E3 ubiquitin-protein ligase NEDD4-like | Q8CFI0 | Glucosamine-6-phosphate isomerase 2 | Q9CRC9 |
| E3 ubiquitin-protein ligase RNF14 | Q9JI90 | Glucosamine--fructose-6-phosphate aminotransfer | P47856 |
| E3 ubiquitin-protein ligase TRIM9 | Q8C7M3 | Glucose-fructose oxidoreductase domain-containir | Q3UHD2 |
| E3 ubiquitin-protein ligase UBR4 | A2AN08 | Glucosylceramidase | P17439 |
| Early endosome antigen 1 | Q8BL66 | Glutamate carboxypeptidase 2 | O35409 |
| Echinoderm microtubule-associated protein-like 1 | Q05BC3 | Glutamate decarboxylase 2 | P48320 |
| Echinoderm microtubule-associated protein-like 2 | Q7TNG5 | Glutamate receptor, ionotropic kainate 2 | P39087 |
| EF-hand domain-containing protein D2 | Q9D8Y0 | Glutamate receptor-interacting protein 1 | Q925T6 |
| EH domain-containing protein 1 | Q9WVK4 | Glutamate--cysteine ligase catalytic subunit | P97494 |
| EH domain-containing protein 3 | Q9QXY6 | Glutamate--cysteine ligase regulatory subunit | O09172 |
| ELAV-like protein 1 | P70372 | Glutaredoxin-related protein 5, mitochondrial | Q80Y14 |
| Electrogenic sodium bicarbonate cotransporter 1 | O88343 | Glutaryl-CoA dehydrogenase, mitochondrial | Q60759 |
| Electron transfer flavoprotein subunit alpha, mitoc | Q99LC5 | Glutathione reductase, mitochondrial | P47791 |
| Electron transfer flavoprotein subunit beta | Q9DCW4 | Glutathione S-transferase A4 | P24472 |
| Electron transfer flavoprotein-ubiquinone oxidored | Q921G7 | Glutathione S-transferase Mu 7 | Q80W21 |
| ELKS/Rab6-interacting/CAST family member 1 | Q99MI1 | Glycerol kinase | Q64516 |
| Elongation factor 1-alpha 1 | P10126 | Glycerophosphodiester phosphodiesterase 1 | Q9JL56 |
| Elongation factor 1-alpha 2 | P62631 | Glycine amidinotransferase, mitochondrial | Q9D964 |
| Elongation factor 1-delta | P57776 | Glycogenin-1 | Q9R062 |
| Elongation factor 1-gamma | Q9D8N0 | Golgi membrane protein 1 | Q91XA2 |
| Elongation factor 2 | P58252 | Golgi SNAP receptor complex member 2 | O35166 |
| Elongation factor Tu, mitochondrial | Q8BFR5 | Golgi to ER traffic protein 4 homolog | Q9D1H7 |
| Elongator complex protein 2 | Q91WG4 | Golgin subfamily A member 1 | Q9CW79 |
| Elongator complex protein 4 | Q9ER73 | Golgin subfamily A member 4 | Q91VW5 |
| Endophilin-A1 | Q62420 | G-protein coupled receptor-associated sorting prot | Q5U4C1 |

| | | | |
|---|---|---|---|
| Endophilin-A2 | Q62419 | G-protein-signaling modulator 1 | Q6IR34 |
| Endophilin-A3 | Q62421 | GRAM domain-containing protein 1B | Q80TI0 |
| Endophilin-B1 | Q9JK48 | GRIP and coiled-coil domain-containing protein 1 | Q9D4H2 |
| Endophilin-B2 | Q8R3V5 | Growth arrest and DNA damage-inducible proteins | Q9CR59 |
| Endoplasmic reticulum resident protein 29 | P57759 | GSK3-beta interaction protein | Q8BGR8 |
| Endoplasmic reticulum resident protein 44 | Q9D1Q6 | GTPase HRas | Q61411 |
| Endoplasmin | P08113 | GTPase KRas | P32883 |
| Engulfment and cell motility protein 2 | Q8BHL5 | GTPase-activating protein and VPS9 domain-conta | Q6PAR5 |
| Enoyl-CoA delta isomerase 1, mitochondrial | P42125 | GTPase-activating Rap/Ran-GAP domain-like prot | Q3V0G7 |
| Enoyl-CoA hydratase, mitochondrial | Q8BH95 | GTP-binding protein 1 | O08582 |
| Ensconsin | O88735 | GTP-binding protein Di-Ras1 | Q91Z61 |
| Ephrin type-A receptor 4 | Q03137 | GTP-binding protein REM 2 | Q8VEL9 |
| Epidermal growth factor receptor substrate 15-like | Q60902 | GTP-binding protein SAR1a | P36536 |
| Epoxide hydrolase 2 | P34914 | GTP-binding protein SAR1b | Q9CQC9 |
| Epsin-1 | Q80VP1 | Guanine nucleotide-binding protein G(I)/G(S)/G(O) | Q9DAS9 |
| Epsin-2 | Q8CHU3 | Guanine nucleotide-binding protein G(I)/G(S)/G(O) | P63213 |
| ERC protein 2 | Q6PH08 | Guanine nucleotide-binding protein G(I)/G(S)/G(O) | P63216 |
| Erlin-2 | Q8BFZ9 | Guanine nucleotide-binding protein G(I)/G(S)/G(O) | Q61016 |
| Ermin | Q5EBJ4 | Guanine nucleotide-binding protein G(I)/G(S)/G(T) | P62880 |
| ES1 protein homolog, mitochondrial | Q9D172 | Guanine nucleotide-binding protein G(olf) subunit a | Q8CGK7 |
| Ester hydrolase C11orf54 homolog | Q91V76 | Guanine nucleotide-binding protein-like 1 | P36916 |
| Ethanolamine-phosphate cytidylyltransferase | Q922E4 | Guanylate kinase | Q64520 |
| Eukaryotic initiation factor 4A-I | P60843 | Haloacid dehalogenase-like hydrolase domain-con | Q3UGR5 |
| Eukaryotic initiation factor 4A-II | P10630 | Haloacid dehalogenase-like hydrolase domain-con | Q9CYW4 |
| Eukaryotic translation initiation factor 2 subunit 1 | Q6ZWX6 | Hamartin | Q9EP53 |
| Eukaryotic translation initiation factor 2 subunit 3, X | Q9Z0N1 | HCLS1-associated protein X-1 | O35387 |
| Eukaryotic translation initiation factor 3 subunit A | P23116 | HD domain-containing protein 2 | Q3SXD3 |
| Eukaryotic translation initiation factor 3 subunit B | Q8JZQ9 | Heat shock 70 kDa protein 14 | Q99M31 |
| Eukaryotic translation initiation factor 3 subunit C | Q8R1B4 | Heat shock factor-binding protein 1 | Q9CQZ1 |
| Eukaryotic translation initiation factor 3 subunit E | P60229 | Hemopexin | Q91X72 |
| Eukaryotic translation initiation factor 3 subunit G | Q9Z1D1 | Heparan sulfate 2-O-sulfotransferase 1 | Q8R3H7 |
| Eukaryotic translation initiation factor 3 subunit H | Q91WK2 | Hepatocyte cell adhesion molecule | Q640R3 |
| Eukaryotic translation initiation factor 3 subunit I | Q9QZD9 | Hepatoma-derived growth factor-related protein 3 | Q9JMG7 |
| Eukaryotic translation initiation factor 3 subunit J | Q66JS6 | Heterogeneous nuclear ribonucleoprotein A0 | Q9CX86 |
| Eukaryotic translation initiation factor 3 subunit K | Q9DBZ5 | Heterogeneous nuclear ribonucleoprotein A1 | P49312 |
| Eukaryotic translation initiation factor 3 subunit L | Q8QZY1 | Heterogeneous nuclear ribonucleoprotein A3 | Q8BG05 |
| Eukaryotic translation initiation factor 4 gamma 1 | Q6NZJ6 | Heterogeneous nuclear ribonucleoprotein F | Q9Z2X1 |
| Eukaryotic translation initiation factor 4 gamma 2 | Q62448 | Heterogeneous nuclear ribonucleoprotein H | O35737 |
| Eukaryotic translation initiation factor 4 gamma 3 | Q80XI3 | Heterogeneous nuclear ribonucleoprotein H2 | P70333 |
| Eukaryotic translation initiation factor 4B | Q8BGD9 | Heterogeneous nuclear ribonucleoprotein L | Q8R081 |
| Eukaryotic translation initiation factor 4H | Q9WUK2 | Heterogeneous nuclear ribonucleoprotein L-like | Q921F4 |
| Eukaryotic translation initiation factor 5 | P59325 | Heterogeneous nuclear ribonucleoprotein M | Q9D0E1 |
| Eukaryotic translation initiation factor 5A-1 | P63242 | Hexokinase-2 | O08528 |
| Eukaryotic translation initiation factor 5B | Q05D44 | Hexokinase-3 | Q3TRM8 |
| Excitatory amino acid transporter 2 | P43006 | High mobility group protein B1 | P63158 |
| Exocyst complex component 1 | Q8R3S6 | Hippocalcin-like protein 1 | P62748 |
| Exocyst complex component 3 | Q6KAR6 | Histidine triad nucleotide-binding protein 1 | P70349 |
| Exocyst complex component 4 | O35382 | Histidine triad nucleotide-binding protein 2, mitocho | Q9D0S9 |
| Exocyst complex component 5 | Q3TPX4 | Histone deacetylase 11 | Q91WA3 |
| Exocyst complex component 7 | O35250 | Histone deacetylase 5 | Q9Z2V6 |
| Exportin-1 | Q6P5F9 | Histone H2B type 1-P | Q8CGP2 |
| Exportin-2 | Q9ERK4 | Histone-arginine methyltransferase CARM1 | Q9WVG6 |
| Ezrin | P26040 | Histone-binding protein RBBP4 | Q60972 |
| F-actin-capping protein subunit alpha-1 | P47753 | Histone-lysine N-methyltransferase MLL2 | Q6PDK2 |
| F-actin-capping protein subunit alpha-2 | P47754 | Histone-lysine N-methyltransferase setd3 | Q91WC0 |
| F-actin-capping protein subunit beta | P47757 | Homeodomain-only protein | Q8R1H0 |
| Far upstream element-binding protein 1 | Q91WJ8 | Homer protein homolog 2 | Q9QWW1 |
| Far upstream element-binding protein 2 | Q3U0V1 | Huntingtin-associated protein 1 | O35668 |
| Farnesyl pyrophosphate synthase | Q920E5 | Huntingtin-interacting protein K | Q9CR41 |
| Fascin | Q61553 | Hydroxyacylglutathione hydrolase-like protein | Q9DB32 |
| Fatty acid synthase | P19096 | Hydroxymethylglutaryl-CoA lyase, mitochondrial | P38060 |
| Fatty acid-binding protein, epidermal | Q05816 | Hydroxymethylglutaryl-CoA synthase, mitochondria | P54869 |
| Fatty acid-binding protein, heart | P11404 | Hydroxysteroid dehydrogenase-like protein 2 | Q2TPA8 |
| Fatty aldehyde dehydrogenase | P47740 | IgLON family member 5 | Q8HW98 |
| F-box only protein 2 | Q80UW2 | Immunoglobulin superfamily member 21 | Q7TNR6 |
| F-box only protein 41 | Q6NS60 | Importin subunit alpha-7 | O35345 |
| F-box/LRR-repeat protein 16 | A2RT62 | Inactive hydroxysteroid dehydrogenase-like proteir | Q8BTX9 |
| FERM, RhoGEF and pleckstrin domain-containing | Q91VS8 | Inosine triphosphate pyrophosphatase | Q9D892 |
| Fibronectin type III and SPRY domain-containing p | Q7TPM6 | Inosine-5'-monophosphate dehydrogenase 1 | P50096 |
| Fibrous sheath-interacting protein 2 | A2ARZ3 | Inositol hexakisphosphate and diphosphoinositol-p | A2ARP1 |
| Flotillin-1 | O08917 | Inositol hexakisphosphate and diphosphoinositol-p | Q6ZQB6 |
| Flotillin-2 | Q60634 | Inositol monophosphatase 3 | Q80V26 |
| Focal adhesion kinase 1 | P34152 | Inositol polyphosphate 1-phosphatase | P49442 |
| Formin-2 | Q9JL04 | Inositol-3-phosphate synthase 1 | Q9JHU9 |
| Formin-binding protein 1 | Q80TY0 | Insulin receptor substrate 2 | P81122 |
| Formin-like protein 1 | Q9JL26 | Integrin alpha-6 | Q61739 |
| Formin-like protein 2 | A2APV2 | Integrin alpha-M | P05555 |
| Fragile X mental retardation protein 1 homolog | P35922 | Integrin-alpha FG-GAP repeat-containing protein 2 | Q91WI7 |
| Fragile X mental retardation syndrome-related prot | Q61584 | Interferon-inducible double stranded RNA-depende | Q9WTX2 |
| Fragile X mental retardation syndrome-related prot | Q9WVR4 | Interleukin enhancer-binding factor 2 | Q9CXY6 |
| Fructose-bisphosphate aldolase A | P05064 | Interleukin enhancer-binding factor 3 | Q9Z1X4 |
| Fructose-bisphosphate aldolase C | P05063 | Intraflagellar transport protein 172 homolog | Q6VH22 |
| FSD1-like protein | Q8BYN5 | Inverted formin-2 | Q0GNC1 |
| Fumarate hydratase, mitochondrial | P97807 | Isoamyl acetate-hydrolyzing esterase 1 homolog | Q9DB29 |
| G protein-regulated inducer of neurite outgrowth 1 | Q3UNH4 | Isobutyryl-CoA dehydrogenase, mitochondrial | Q9D7B6 |
| Galactokinase | Q9R0N0 | Isochorismatase domain-containing protein 1 | Q91V64 |
| Galectin-1 | P16045 | Isoleucine--tRNA ligase, mitochondrial | Q8BIJ6 |
| Gamma-adducin | Q9QYB5 | Isopentenyl-diphosphate Delta-isomerase 1 | P58044 |
| Gamma-aminobutyric acid receptor subunit alpha- | P62812 | IST1 homolog | Q9CX00 |
| Gamma-aminobutyric acid receptor subunit beta-3 | P63080 | Junction-mediating and -regulatory protein | Q9QXM1 |
| Gamma-aminobutyric acid type B receptor subunit | Q80T41 | Junctophilin-4 | Q80WT0 |
| Gamma-enolase | P17183 | Katanin p60 ATPase-containing subunit A-like 1 | Q8K0T4 |
| Gamma-glutamyltransferase 7 | Q99JP7 | Katanin p80 WD40-containing subunit B1 | Q8BG40 |
| Gamma-soluble NSF attachment protein | Q9CWZ7 | Kelch-like protein 22 | Q99JN2 |

| | | | |
|---|---|---|---|
| Gamma-synuclein | Q9Z0F7 | Ketosamine-3-kinase | Q8K274 |
| Ganglioside-induced differentiation-associated pro | O88741 | KH domain-containing, RNA-binding, signal transd | Q60749 |
| Ganglioside-induced differentiation-associated pro | Q8VE33 | KH domain-containing, RNA-binding, signal transd | Q9R226 |
| General receptor for phosphoinositides 1-associat | Q9JJA9 | KIF1-binding protein | Q6ZPU9 |
| General transcription factor IIF subunit 1 | Q3THK3 | Kinesin heavy chain isoform 5A | P33175 |
| General transcription factor II-I | Q9ESZ8 | Kinesin-like protein KIF13A | Q9EQW7 |
| General vesicular transport factor p115 | Q9Z1Z0 | Kinesin-like protein KIF21B | Q9QXL1 |
| Gephyrin | Q8BUV3 | Kinesin-like protein KIF3A | P28741 |
| Geranylgeranyl transferase type-2 subunit alpha | Q9JHK4 | Kinesin-like protein KIFC2 | O08672 |
| Geranylgeranyl transferase type-2 subunit beta | P53612 | Kv channel-interacting protein 2 | Q9JJ69 |
| Girdin | Q5SNZ0 | Lanosterol synthase | Q8BLN5 |
| Glia maturation factor beta | Q9CQI3 | La-related protein 4B | Q6A0A2 |
| Glial fibrillary acidic protein | P03995 | La-related protein 6 | Q8BN59 |
| Glucosamine-6-phosphate isomerase 1 | O88958 | La-related protein 7 | Q05CL8 |
| Glucose 1,6-bisphosphate synthase | Q8CAA7 | Latrophilin-2 (Fragment) | Q8JZZ7 |
| Glucose-6-phosphate 1-dehydrogenase X | Q00612 | Legumain | O89017 |
| Glucose-6-phosphate isomerase | P06745 | Lethal(2) giant larvae protein homolog 1 | Q80Y17 |
| Glucosidase 2 subunit beta | O08795 | Leucine zipper protein 1 | Q8R4U7 |
| Glutamate [NMDA] receptor subunit epsilon-1 | P35436 | Leucine zipper protein 2 | Q8BGY3 |
| Glutamate [NMDA] receptor subunit epsilon-2 | Q01097 | Leucine-rich repeat and fibronectin type III domain | Q2WF71 |
| Glutamate [NMDA] receptor subunit zeta-1 | P35438 | Leucine-rich repeat and immunoglobulin-like doma | Q3URE9 |
| Glutamate decarboxylase 1 | P48318 | Leucine-rich repeat and WD repeat-containing pro | Q6P5U7 |
| Glutamate dehydrogenase 1, mitochondrial | P26443 | Leucine-rich repeat flightless-interacting protein 1 | Q3UZ39 |
| Glutamate receptor 1 | P23818 | Leucine-rich repeat flightless-interacting protein 2 | Q91WK0 |
| Glutamate receptor 2 | P23819 | Leucine-rich repeat-containing protein 16A | Q6EDY6 |
| Glutamate receptor 3 | Q9Z2W9 | Leucine-rich repeat-containing protein 16B | Q3UFQ8 |
| Glutamine synthetase | P15105 | Leucine-rich repeat-containing protein 40 | Q9CRC8 |
| Glutaredoxin-3 | Q9CQM9 | Leucine-rich repeat-containing protein 4B | P0C192 |
| Glutathione S-transferase Mu 1 | P10649 | Leucine-rich repeat-containing protein 57 | Q9D1G5 |
| Glutathione S-transferase Mu 5 | P48774 | Leucine-rich repeat-containing protein 8A | Q80WG5 |
| Glutathione S-transferase omega-1 | O09131 | Leucyl-cystinyl aminopeptidase | Q8C129 |
| Glutathione S-transferase P 1 | P19157 | LIM domain and actin-binding protein 1 | Q9ERG0 |
| Glutathione synthetase | P51855 | LIM domain-containing protein 2 | Q8BGB5 |
| Glyceraldehyde-3-phosphate dehydrogenase | P16858 | Lipopolysaccharide-responsive and beige-like ancl | Q9ESE1 |
| Glycerol-3-phosphate dehydrogenase [NAD(+)], cy | P13707 | Long-chain fatty acid transport protein 4 | Q91VE0 |
| Glycerol-3-phosphate dehydrogenase 1-like protei | Q3ULJ0 | Long-chain-fatty-acid--CoA ligase 1 | P41216 |
| Glycerol-3-phosphate dehydrogenase, mitochondr | Q64521 | Long-chain-fatty-acid--CoA ligase 3 | Q9CZW4 |
| Glycerophosphocholine phosphodiesterase GPCP | Q8C0L9 | Long-chain-fatty-acid--CoA ligase 4 | Q9QUJ7 |
| Glycerophosphodiester phosphodiesterase domair | Q9CRY7 | Low-density lipoprotein receptor-related protein 1E | Q9JI18 |
| Glycine dehydrogenase [decarboxylating], mitocho | Q91W43 | Luc7-like protein 3 | Q5SUF2 |
| Glycine--tRNA ligase | Q9CZD3 | Lymphocyte antigen 6H | Q9WUC3 |
| Glycogen phosphorylase, brain form | Q8CI94 | LysM and putative peptidoglycan-binding domain-c | Q9D0E3 |
| Glycogen phosphorylase, muscle form | Q9WUB3 | LysM and putative peptidoglycan-binding domain-c | Q9D7V2 |
| Glycogen synthase kinase-3 alpha | Q2NL51 | Lysophospholipid acyltransferase LPCAT4 | Q6NVG1 |
| Glycogen synthase kinase-3 beta | Q9WV60 | Lysosomal acid phosphatase | P24638 |
| Glycolipid transfer protein | Q9JL62 | Lysosomal-trafficking regulator | P97412 |
| Glycylpeptide N-tetradecanoyltransferase 1 | O70310 | Lysosome membrane protein 2 | O35114 |
| Glycylpeptide N-tetradecanoyltransferase 2 | O70311 | Lysosome-associated membrane glycoprotein 1 | P11438 |
| Glyoxalase domain-containing protein 4 | Q9CPV4 | Macrophage erythroblast attacher | Q4VC33 |
| Glyoxylate reductase/hydroxypyruvate reductase | Q91Z53 | Macrophage migration inhibitory factor | P34884 |
| Glypican-1 | Q9QZF2 | Magnesium-dependent phosphatase 1 | Q9D967 |
| Glypican-4 | P51655 | MAGUK p55 subfamily member 3 | O88910 |
| Glypican-5 | Q8CAL5 | Malcavernin | Q8K2Y9 |
| GMP reductase 1 | Q9DCZ1 | Malectin | Q6ZQI3 |
| GMP synthase [glutamine-hydrolyzing] | Q3THK7 | Mannose-1-phosphate guanyltransferase beta | Q8BTZ7 |
| Golgi apparatus protein 1 | Q61543 | Mannose-6-phosphate isomerase | Q924M7 |
| Golgi reassembly-stacking protein 2 | Q99JX3 | Mannosyl-oligosaccharide glucosidase | Q80UM7 |
| Golgi resident protein GCP60 | Q8BMP6 | MAP/microtubule affinity-regulating kinase 4 | Q8CIP4 |
| Golgin subfamily A member 2 | Q921M4 | Mast/stem cell growth factor receptor Kit | P05532 |
| Golgin subfamily A member 3 | P55937 | Matrilin-4 | O89029 |
| GRIP and coiled-coil domain-containing protein 2 | Q8CHG3 | Melanoma inhibitory activity protein 3 | Q8BI84 |
| GRIP1-associated protein 1 | Q8VD04 | Melanoma-associated antigen E1 | Q6PCZ4 |
| Growth arrest-specific protein 7 | Q60780 | Membrane-associated guanylate kinase, WW and | Q9EQJ9 |
| Growth factor receptor-bound protein 2 | Q60631 | Metabotropic glutamate receptor 1 | P97772 |
| GrpE protein homolog 1, mitochondrial | Q99LP6 | Metabotropic glutamate receptor 7 | Q68ED2 |
| GTP:AMP phosphotransferase, mitochondrial | Q9WTP7 | Metal transporter CNNM3 | Q32NY4 |
| GTPase NRas | P08556 | Metallo-beta-lactamase domain-containing protein | Q8BL86 |
| GTP-binding nuclear protein Ran | P62827 | Metallothionein-3 | P28184 |
| GTP-binding protein Di-Ras2 | Q5PR73 | Metastasis-associated protein MTA3 | Q924K8 |
| GTP-binding protein Rheb | Q921J2 | Metaxin-2 | O88441 |
| Guanine deaminase | Q9R111 | Methionine aminopeptidase 1 | Q8BP48 |
| Guanine nucleotide-binding protein G(i) subunit alp | B2RSH2 | Methylmalonic aciduria type A homolog, mitochond | Q8C7H1 |
| Guanine nucleotide-binding protein G(i) subunit alp | P08752 | Methylosome protein 50 | Q99J09 |
| Guanine nucleotide-binding protein G(I)/G(S)/G(T) | P62874 | Methylosome subunit pICln | Q61189 |
| Guanine nucleotide-binding protein G(k) subunit al | Q9DC51 | Methylthioribose-1-phosphate isomerase | Q9CQT1 |
| Guanine nucleotide-binding protein G(o) subunit al | P18872 | Microtubule-associated protein 1S | Q8C052 |
| Guanine nucleotide-binding protein G(q) subunit al | P21279 | Microtubule-associated protein 9 | Q3TRR0 |
| Guanine nucleotide-binding protein G(s) subunit al | Q6R0H7 | Microtubule-associated protein RP/EB family mem | Q61166 |
| Guanine nucleotide-binding protein G(z) subunit al | O70443 | Microtubule-associated protein RP/EB family mem | Q8R001 |
| Guanine nucleotide-binding protein subunit alpha-1 | P21278 | Microtubule-associated protein RP/EB family mem | Q6PER3 |
| Guanine nucleotide-binding protein subunit alpha-1 | P27601 | Microtubule-associated proteins 1A/1B light chain 3 | Q91VR7 |
| Guanine nucleotide-binding protein subunit beta-2- | P68040 | Microtubule-associated tumor suppressor candidat | Q3UHD3 |
| Guanine nucleotide-binding protein subunit beta-4 | P29387 | Mimitin, mitochondrial | Q59J78 |
| Guanine nucleotide-binding protein subunit beta-5 | P62881 | MIP18 family protein FAM96A | Q9DCL2 |
| Guanylate cyclase soluble subunit beta-1 | O54865 | Mitochondrial 10-formyltetrahydrofolate dehydroge | Q8K009 |
| HBS1-like protein | Q69ZS7 | Mitochondrial carnitine/acylcarnitine carrier protein | Q9Z2Z6 |
| Heat shock 70 kDa protein 12A | Q8K0U4 | Mitochondrial carrier homolog 1 | Q791T5 |
| Heat shock 70 kDa protein 1B | P17879 | Mitochondrial carrier homolog 2 | Q791V5 |
| Heat shock 70 kDa protein 4 | Q61316 | Mitochondrial fission factor | Q6PCP5 |
| Heat shock 70 kDa protein 4L | P48722 | Mitochondrial import inner membrane translocase | P62075 |
| Heat shock cognate 71 kDa protein | P63017 | Mitochondrial import inner membrane translocase | Q9CQV1 |
| Heat shock protein 105 kDa | Q61699 | Mitochondrial import inner membrane translocase | Q9WVA2 |
| Heat shock protein 75 kDa, mitochondrial | Q9CQN1 | Mitochondrial import inner membrane translocase | P62077 |
| Heat shock protein HSP 90-alpha | P07901 | Mitochondrial import inner membrane translocase | Q9WV98 |

| | | | |
|---|---|---|---|
| Heat shock protein HSP 90-beta | P11499 | Mitochondrial import inner membrane translocase | Q9WV96 |
| Heat shock-related 70 kDa protein 2 | P17156 | Mitochondrial import receptor subunit TOM22 hom | Q9CPQ3 |
| Heme oxygenase 2 | O70252 | Mitochondrial import receptor subunit TOM40 hom | Q9QYA2 |
| Heme-binding protein 1 | Q9R257 | Mitochondrial intermediate peptidase | A6H611 |
| Hemoglobin subunit alpha | P01942 | Mitochondrial Rho GTPase 1 | Q8BG51 |
| Hemoglobin subunit beta-1 | P02088 | Mitochondrial-processing peptidase subunit alpha | Q9DC61 |
| Hepatocyte growth factor-regulated tyrosine kinase | Q99LI8 | Mitofusin-1 | Q811U4 |
| Hepatoma-derived growth factor | P51859 | Mitogen-activated protein kinase 10 | Q61831 |
| Hepatoma-derived growth factor-related protein 2 | Q3UMU9 | Mitogen-activated protein kinase 8 | Q91Y86 |
| Heterochromatin protein 1-binding protein 3 | Q3TEA8 | Mitogen-activated protein kinase kinase kinase kin | Q99JP0 |
| Heterogeneous nuclear ribonucleoprotein A/B | Q99020 | Mitogen-activated protein kinase kinase kinase kin | Q8BPM2 |
| Heterogeneous nuclear ribonucleoprotein D0 | Q60668 | Mitotic checkpoint protein BUB3 | Q9WVA3 |
| Heterogeneous nuclear ribonucleoprotein K | P61979 | Mitotic spindle-associated MMXD complex subunit | Q9D187 |
| Heterogeneous nuclear ribonucleoprotein Q | Q7TMK9 | MOB kinase activator 1B | Q8BPB0 |
| Heterogeneous nuclear ribonucleoprotein U | Q8VEK3 | Molybdopterin synthase sulfur carrier subunit | Q9Z224 |
| Heterogeneous nuclear ribonucleoprotein U-like pr | Q8VDM6 | Monoacylglycerol lipase ABHD6 | Q8R2Y0 |
| Heterogeneous nuclear ribonucleoprotein U-like pr | Q00PI9 | MOSC domain-containing protein 2, mitochondrial | Q922Q1 |
| Heterogeneous nuclear ribonucleoproteins A2/B1 | O88569 | mRNA cap guanine-N7 methyltransferase | Q9D0L8 |
| Heterogeneous nuclear ribonucleoproteins C1/C2 | Q9Z204 | mRNA export factor | Q8C570 |
| Hexokinase-1 | P17710 | MTSS1-like protein | Q6P9S0 |
| Hippocalcin-like protein 4 | Q8BGZ1 | Multiple epidermal growth factor-like domains prot | Q8BH27 |
| Histamine N-methyltransferase | Q91VF2 | Myb-binding protein 1A | Q7TPV4 |
| Histidine--tRNA ligase, cytoplasmic | Q61035 | Myelin expression factor 2 | Q8C854 |
| Histone deacetylase 4 | Q6NZM9 | Myelin proteolipid protein | P60202 |
| Histone deacetylase 6 | Q9Z2V5 | Myeloid leukemia factor 2 | Q99KX1 |
| Histone H1.0 | P10922 | Myomegalin | Q80YT7 |
| Histone H3.1 | P68433 | Myosin phosphatase Rho-interacting protein | P97434 |
| Histone H4 | P62806 | Myosin regulatory light polypeptide 9 | Q9CQ19 |
| Histone-lysine N-methyltransferase SETD7 | Q8VHL1 | Myotubularin-related protein 2 | Q9Z2D1 |
| HIV Tat-specific factor 1 homolog | Q8BGC0 | Myotubularin-related protein 3 | Q8K296 |
| Homer protein homolog 1 | Q9Z2Y3 | Myotubularin-related protein 6 | Q8VE11 |
| Homer protein homolog 3 | Q99JP6 | Myotubularin-related protein 7 | Q9Z2C9 |
| Hsc70-interacting protein | Q99L47 | Myotubularin-related protein 9 | Q9Z2D0 |
| Hsp90 co-chaperone Cdc37 | Q61081 | N-acetyl-D-glucosamine kinase | Q9QZ08 |
| Huntingtin | P42859 | N-acetylglucosamine-6-sulfatase | Q8BFR4 |
| Huntingtin-interacting protein 1 | Q8VD75 | N-acetylneuraminate lyase | Q9DCJ9 |
| Huntingtin-interacting protein 1-related protein | Q9JKY5 | N-acylneuraminate cytidylyltransferase | Q99KK2 |
| Hyaluronan and proteoglycan link protein 1 | Q9QUP5 | NAD-dependent malic enzyme, mitochondrial | Q99KE1 |
| Hydroxyacyl-coenzyme A dehydrogenase, mitocho | Q61425 | NAD-dependent protein deacetylase sirtuin-2 | Q8VDQ8 |
| Hydroxyacylglutathione hydrolase, mitochondrial | Q99KB8 | NAD-dependent protein deacylase sirtuin-5, mitoch | Q8K2C6 |
| Hydroxymethylglutaryl-CoA synthase, cytoplasmic | Q8JZK9 | NADH dehydrogenase [ubiquinone] 1 alpha subco | Q9D1H6 |
| Hypoxanthine-guanine phosphoribosyltransferase | P00493 | NADH dehydrogenase [ubiquinone] 1 alpha subco | Q62425 |
| Hypoxia up-regulated protein 1 | Q9JKR6 | NADH dehydrogenase [ubiquinone] 1 alpha subco | Q9CQZ5 |
| Immunity-related GTPase family Q protein | Q8VIM9 | NADH dehydrogenase [ubiquinone] 1 alpha subco | Q9DCJ5 |
| Immunoglobulin superfamily containing leucine-ric | Q5RKR3 | NADH dehydrogenase [ubiquinone] 1 beta subcom | Q9CQC7 |
| Immunoglobulin superfamily member 8 | Q8R366 | NADH dehydrogenase [ubiquinone] 1 beta subcom | Q3UIU2 |
| Immunoglobulin-binding protein 1 | Q61249 | NADH dehydrogenase [ubiquinone] 1 beta subcom | Q9CR61 |
| Importin subunit alpha-3 | O35344 | NADH dehydrogenase [ubiquinone] 1 beta subcom | Q9CQJ8 |
| Importin subunit beta-1 | P70168 | NADH dehydrogenase [ubiquinone] 1 subunit C2 | Q9CQ54 |
| Importin-4 | Q8VI75 | NADH dehydrogenase [ubiquinone] flavoprotein 3, | Q8BK30 |
| Importin-5 | Q8BKC5 | NADH dehydrogenase [ubiquinone] iron-sulfur pro | Q99LY9 |
| Importin-7 | Q9EPL8 | NADH dehydrogenase [ubiquinone] iron-sulfur pro | P52503 |
| Importin-9 | Q91YE6 | NADH dehydrogenase [ubiquinone] iron-sulfur pro | Q9DC70 |
| Inactive dipeptidyl peptidase 10 | Q6NXK7 | NADPH:adrenodoxin oxidoreductase, mitochondria | Q61578 |
| Inactive phospholipase C-like protein 1 | Q3USB7 | Nasal embryonic luteinizing hormone-releasing ho | Q99NF2 |
| Inactive phospholipase C-like protein 2 | Q8K394 | N-chimaerin | Q91V57 |
| Inorganic pyrophosphatase | Q9D819 | NEDD8-activating enzyme E1 catalytic subunit | Q8C878 |
| Inorganic pyrophosphatase 2, mitochondrial | Q91VM9 | Neighbor of COX4 | O70378 |
| Inosine-5'-monophosphate dehydrogenase 2 | P24547 | Netrin receptor DCC | P70211 |
| Inositol 1,4,5-trisphosphate receptor type 1 | P11881 | Neudesin | Q9CQ45 |
| Inositol monophosphatase 1 | O55023 | Neural Wiskott-Aldrich syndrome protein | Q91YD9 |
| Inositol-trisphosphate 3-kinase A | Q8R071 | Neuritin | Q8CFV4 |
| Insulin-degrading enzyme | Q9JHR7 | Neuroendocrine convertase 2 | P21661 |
| Integrin alpha-V | P43406 | Neuroendocrine protein 7B2 | P12961 |
| Integrin beta-1 | P09055 | Neurogranin | P60761 |
| Intercellular adhesion molecule 5 | Q60625 | Neuroligin-2 | Q69ZK9 |
| Intersectin-1 | Q9Z0R4 | Neuroligin-3 | Q8BYM5 |
| Intersectin-2 | Q9Z0R6 | Neuronal migration protein doublecortin | O88809 |
| Intracellular hyaluronan-binding protein 4 | Q9JKS5 | Neuronal pentraxin-2 | O70340 |
| IQ motif and SEC7 domain-containing protein 1 | Q8R0S2 | Neuronal proto-oncogene tyrosine-protein kinase S | P05480 |
| IQ motif and SEC7 domain-containing protein 2 | Q5DU25 | Neuropilin-1 | P97333 |
| IQ motif and SEC7 domain-containing protein 3 | Q3TES0 | Neuroserpin | O35684 |
| Iron-sulfur cluster assembly 2 homolog, mitochon | Q9DCB8 | Neutral cholesterol ester hydrolase 1 | Q8BLF1 |
| Iron-sulfur cluster assembly enzyme ISCU, mitoch | Q9D7P6 | NFU1 iron-sulfur cluster scaffold homolog, mitoch | Q9QZ23 |
| Islet cell autoantigen 1 | P97411 | NHS-like protein 1 | Q8CAF4 |
| Isocitrate dehydrogenase [NAD] subunit alpha, mit | Q9D6R2 | Nicalin | Q8VCM8 |
| Isocitrate dehydrogenase [NAD] subunit gamma 1, | P70404 | NIF3-like protein 1 | Q9EQ80 |
| Isocitrate dehydrogenase [NADP] cytoplasmic | O88844 | NLR family member X1 | Q3TL44 |
| Isocitrate dehydrogenase [NADP], mitochondrial | P54071 | Noelin-2 | Q8BM13 |
| Isoleucine--tRNA ligase, cytoplasmic | Q8BU30 | N-terminal EF-hand calcium-binding protein 1 | Q8BG18 |
| Isovaleryl-CoA dehydrogenase, mitochondrial | Q9JHI5 | Nuclear autoantigenic sperm protein | Q99MD9 |
| Janus kinase and microtubule-interacting protein 1 | Q8BVL9 | Nuclear cap-binding protein subunit 1 | Q3UYV9 |
| Janus kinase and microtubule-interacting protein 3 | Q5DTN8 | Nuclear receptor-binding protein | Q99J45 |
| Junctophilin-3 | Q9ET77 | Nuclear receptor-binding protein 2 | Q91V36 |
| Kalirin | A2CG49 | Nuclear transport factor 2 | P61971 |
| Kelch repeat and BTB domain-containing protein 1 | Q8BNW9 | Nuclear ubiquitous casein and cyclin-dependent ki | Q80XU3 |
| Kinase suppressor of Ras 1 | Q61097 | Nucleolar protein 58 | Q6DFW4 |
| Kinectin | Q61595 | Nucleolysin TIAR | P70318 |
| Kinesin heavy chain isoform 5C | P28738 | Nucleoside diphosphate kinase 3 | Q9WV85 |
| Kinesin light chain 1 | O88447 | Nucleosome assembly protein 1-like 1 | P28656 |
| Kinesin light chain 2 | O88448 | OCIA domain-containing protein 1 | Q9CRD0 |
| Kinesin-1 heavy chain | Q61768 | OCIA domain-containing protein 2 | Q9D8W7 |
| Kinesin-associated protein 3 | P70188 | Oligoribonuclease, mitochondrial | Q9D8S4 |
| Kinesin-like protein KIF1A | P33173 | O-phosphoseryl-tRNA(Sec) selenium transferase | Q6P6M7 |

| | |
|---|---|
| Kinesin-like protein KIF21A | Q9QXL2 |
| Kinesin-like protein KIF2A | P28740 |
| Kinesin-like protein KIF3B | Q61771 |
| Kinesin-like protein KIF3C | O35066 |
| Kv channel-interacting protein 4 | Q6PHZ8 |
| L-2-hydroxyglutarate dehydrogenase, mitochondria | Q91YP0 |
| Lactoylglutathione lyase | Q9CPU0 |
| Lambda-crystallin homolog | Q99KP3 |
| LanC-like protein 1 | O89112 |
| LanC-like protein 2 | Q9JJK2 |
| La-related protein 1 | Q6ZQ58 |
| Large proline-rich protein BAG6 | Q9Z1R2 |
| L-asparaginase | Q8C0M9 |
| Late secretory pathway protein AVL9 homolog | Q80U56 |
| Latrophilin-1 | Q80TR1 |
| Latrophilin-3 | Q80TS3 |
| LDLR chaperone MESD | Q9ERE7 |
| LETM1 and EF-hand domain-containing protein 1, | Q9Z2I0 |
| Leucine zipper transcription factor-like protein 1 | Q9JHQ5 |
| Leucine-rich glioma-inactivated protein 1 | Q9JIA1 |
| Leucine-rich PPR motif-containing protein, mitocho | Q6PB66 |
| Leucine-rich repeat and immunoglobulin-like doma | Q9D1T0 |
| Leucine-rich repeat-containing protein 16C | Q3V3V9 |
| Leucine-rich repeat-containing protein 47 | Q505F5 |
| Leucine-rich repeat-containing protein 59 | Q922Q8 |
| Leucine-rich repeat-containing protein 7 | Q80TE7 |
| Leucine--tRNA ligase, cytoplasmic | Q8BMJ2 |
| Leukocyte surface antigen CD47 | Q61735 |
| Leukotriene A-4 hydrolase | P24527 |
| LIM and calponin homology domains-containing pr | Q3UH68 |
| LIM and SH3 domain protein 1 | Q61792 |
| Limbic system-associated membrane protein | Q8BLK3 |
| Lipid phosphate phosphatase-related protein type | Q7TME0 |
| Lipid phosphate phosphohydrolase 3 | Q99JY8 |
| Lipoamide acyltransferase component of branched | P53395 |
| Liprin-alpha-2 | Q8BSS9 |
| Liprin-alpha-3 | P60469 |
| LisH domain and HEAT repeat-containing protein I | Q148V7 |
| L-lactate dehydrogenase A chain | P06151 |
| L-lactate dehydrogenase B chain | P16125 |
| Lon protease homolog, mitochondrial | Q8CGK3 |
| Long-chain specific acyl-CoA dehydrogenase, mito | P51174 |
| Long-chain-fatty-acid--CoA ligase 6 | Q91WC3 |
| Long-chain-fatty-acid--CoA ligase ACSBG1 | Q99PU5 |
| Low molecular weight phosphotyrosine protein pho | Q9D358 |
| Lupus La protein homolog | P32067 |
| Lysine--tRNA ligase | Q99MN1 |
| Lysophosphatidic acid phosphatase type 6 | Q8BP40 |
| Lysosomal alpha-glucosidase | P70699 |
| MAGUK p55 subfamily member 2 | Q9WV34 |
| MAGUK p55 subfamily member 5 | Q9JLB2 |
| MAGUK p55 subfamily member 6 | Q9JLB0 |
| Major prion protein | P04925 |
| Malate dehydrogenase, cytoplasmic | P14152 |
| Malate dehydrogenase, mitochondrial | P08249 |
| Mammalian ependymin-related protein 1 | Q99M71 |
| MAP kinase-activating death domain protein | Q80U28 |
| MAP/microtubule affinity-regulating kinase 3 | Q03141 |
| MAP7 domain-containing protein 1 | A2AJI0 |
| MAP7 domain-containing protein 2 | A2AG50 |
| MARCKS-related protein | P28667 |
| Matrin-3 | Q8K310 |
| Medium-chain specific acyl-CoA dehydrogenase, r | P45952 |
| Megakaryocyte-associated tyrosine-protein kinase | P41242 |
| Membrane-associated guanylate kinase, WW and | Q6RHR9 |
| Membrane-associated guanylate kinase, WW and | Q9WVQ1 |
| Membrane-associated phosphatidylinositol transfe | O35954 |
| Membrane-associated phosphatidylinositol transfe | Q6ZPQ6 |
| Membrane-associated progesterone receptor com | O55022 |
| Membrane-associated progesterone receptor com | Q80UU9 |
| Metabotropic glutamate receptor 2 | Q14BI2 |
| Metabotropic glutamate receptor 3 | Q9QYS2 |
| Metabotropic glutamate receptor 5 | Q3UVX5 |
| Metaxin-1 | P47802 |
| Methionine adenosyltransferase 2 subunit beta | Q99LB6 |
| Methionine aminopeptidase 2 | O08663 |
| Methionine--tRNA ligase, cytoplasmic | Q68FL6 |
| Methylcrotonoyl-CoA carboxylase beta chain, mito | Q3ULD5 |
| Methylcrotonoyl-CoA carboxylase subunit alpha, m | Q99MR8 |
| Methylglutaconyl-CoA hydratase, mitochondrial | Q9JLZ3 |
| Methylmalonate-semialdehyde dehydrogenase [ac | Q9EQ20 |
| Methylmalonyl-CoA mutase, mitochondrial | P16332 |
| Microtubule-actin cross-linking factor 1 | Q9QXZ0 |
| Microtubule-associated protein 1A | Q9QYR6 |
| Microtubule-associated protein 1B | P14873 |
| Microtubule-associated protein 2 | P20357 |
| Microtubule-associated protein 4 | P27546 |
| Microtubule-associated protein 6 | Q7TSJ2 |
| Microtubule-associated protein tau | P10637 |
| Microtubule-associated serine/threonine-protein ki | Q9R1L5 |
| Microtubule-associated serine/threonine-protein ki | Q3U214 |
| Misshapen-like kinase 1 | Q9JM52 |
| Mitochondrial 2-oxoglutarate/malate carrier protein | Q9CR62 |
| Mitochondrial chaperone BCS1 | Q9CZP5 |
| Osteoclast-stimulating factor 1 | Q62422 |
| OTU domain-containing protein 6B | Q8K2H2 |
| OX-2 membrane glycoprotein | O54901 |
| Oxysterol-binding protein 2 | Q5QNQ6 |
| Oxysterol-binding protein-related protein 1 | Q91XL9 |
| Oxysterol-binding protein-related protein 11 | Q8CI95 |
| Oxysterol-binding protein-related protein 2 | Q8BX94 |
| Oxysterol-binding protein-related protein 6 | Q8BXR9 |
| Oxysterol-binding protein-related protein 9 | A2A8Z1 |
| Palmitoyl-protein thioesterase 1 | O88531 |
| Paraplegin | Q3ULF4 |
| Parkinson disease 7 domain-containing protein 1 | Q8BFQ8 |
| Peflin | Q8BFY6 |
| Pentatricopeptide repeat-containing protein 3, mito | Q14C51 |
| Peptidyl-glycine alpha-amidating monooxygenase | P97467 |
| Peptidyl-prolyl cis-trans isomerase FKBP1A | P26883 |
| Peptidyl-prolyl cis-trans isomerase H | Q9D868 |
| Peptidyl-prolyl cis-trans isomerase-like 1 | Q9D0W5 |
| Peptidyl-tRNA hydrolase 2, mitochondrial | Q8R2Y8 |
| Pericentrin | P48725 |
| Perilipin-3 | Q9DBG5 |
| Peroxiredoxin-4 | O08807 |
| Peroxiredoxin-5, mitochondrial | P99029 |
| Peroxisomal 2,4-dienoyl-CoA reductase | Q9WV68 |
| Peroxisomal carnitine O-octanoyltransferase | Q9DC50 |
| Peroxisomal membrane protein PEX14 | Q9R0A0 |
| Peroxisomal targeting signal 1 receptor | O09012 |
| PEST proteolytic signal-containing nuclear protein | Q6P8I4 |
| PH and SEC7 domain-containing protein 1 | Q5DTT2 |
| PH domain leucine-rich repeat-containing protein p | Q8CHE4 |
| Phosphatidate phosphatase LPIN2 | Q99PI5 |
| Phosphatidylinositol 3,4,5-trisphosphate 3-phospha | O08586 |
| Phosphatidylinositol 3,4,5-trisphosphate-dependen | Q3LAC4 |
| Phosphatidylinositol 3-kinase regulatory subunit alp | P26450 |
| Phosphatidylinositol 4,5-bisphosphate 3-kinase cat | Q8BTI9 |
| Phosphatidylserine decarboxylase proenzyme | Q8BSF4 |
| Phosphoacetylglucosamine mutase | Q9CYR6 |
| Phosphofurin acidic cluster sorting protein 2 | Q3V3Q7 |
| Phosphoglucomutase-2 | Q7TSV4 |
| Phosphoinositide 3-kinase regulatory subunit 4 | Q8VD65 |
| Phospholipase A-2-activating protein | P27612 |
| Phospholipase DDHD1 | Q80YA3 |
| Phosphomannomutase 2 | Q9Z2M7 |
| Phosphoribosyl pyrophosphate synthase-associate | Q9D0M1 |
| Phosphoribosyl pyrophosphate synthase-associate | Q8R574 |
| Phosphorylase b kinase regulatory subunit alpha, s | P18826 |
| Phosphorylase b kinase regulatory subunit beta | Q7TSH2 |
| Pinin | O35691 |
| PI-PLC X domain-containing protein 3 | Q8BLJ3 |
| Plasminogen receptor (KT) | Q9D3P8 |
| Platelet-activating factor acetylhydrolase IB subuni | Q61206 |
| Platelet-activating factor acetylhydrolase IB subuni | Q61205 |
| Platelet-derived growth factor subunit A | P20033 |
| Pleckstrin homology domain-containing family A m | Q7TQG1 |
| Pleiotrophin | P63089 |
| Pleiotropic regulator 1 | Q922V4 |
| Plexin-C1 | Q9QZC2 |
| Poliovirus receptor-related protein 1 | Q9JKF6 |
| Poly [ADP-ribose] polymerase 1 | P11103 |
| Poly(rC)-binding protein 3 | P57722 |
| Poly(U)-binding-splicing factor PUF60 | Q3UEB3 |
| Polymerase delta-interacting protein 2 | Q91VA6 |
| Polypyrimidine tract-binding protein 2 | Q91Z31 |
| Polyribonucleotide nucleotidyltransferase 1, mitoch | Q8K1R3 |
| Potassium voltage-gated channel subfamily A mer | P16390 |
| Potassium voltage-gated channel subfamily A mer | Q61423 |
| Potassium voltage-gated channel subfamily KQT r | Q8K3F6 |
| Potassium/sodium hyperpolarization-activated cyc | O88704 |
| Potassium-transporting ATPase alpha chain 2 | Q9Z1W8 |
| PRA1 family protein 2 | Q9JIG8 |
| PRA1 family protein 3 | Q8R5J9 |
| Prefoldin subunit 5 | Q9WU28 |
| Prefoldin subunit 6 | Q03958 |
| Pre-mRNA branch site protein p14 | P59708 |
| Pre-mRNA-processing factor 6 | Q91YR7 |
| Pre-mRNA-processing-splicing factor 8 | Q99PV0 |
| Prenylcysteine oxidase | Q9CQF9 |
| Prenylcysteine oxidase-like | Q8C7K6 |
| Pre-rRNA-processing protein TSR2 homolog | Q8C8T8 |
| Probable arginine--tRNA ligase, mitochondrial | Q3U186 |
| Probable ATP-dependent RNA helicase DDX46 | Q569Z5 |
| Probable ATP-dependent RNA helicase DDX6 | P54823 |
| Probable ATP-dependent RNA helicase DHX36 | Q8VHK9 |
| Probable cationic amino acid transporter | Q8BXR1 |
| Probable cation-transporting ATPase 13A1 | Q9EPE9 |
| Probable D-tyrosyl-tRNA(Tyr) deacylase 2 | Q8BHA3 |
| Probable E3 ubiquitin-protein ligase HERC4 | Q6PAV2 |
| Probable fructose-2,6-bisphosphatase TIGAR | Q8BZA9 |
| Probable global transcription activator SNF2L2 | Q6DIC0 |
| Probable glutamate--tRNA ligase, mitochondrial | Q9CXJ1 |
| Probable histidine--tRNA ligase, mitochondrial | Q99KK9 |
| Probable phospholipid-transporting ATPase IIA | O70228 |
| Probable tRNA threonylcarbamoyladenosine biosy | Q8BWU5 |
| Profilin-1 | P62962 |

| | | | |
|---|---|---|---|
| Mitochondrial glutamate carrier 1 | Q9D6M3 | Profilin-2 | Q9JJV2 |
| Mitochondrial glutamate carrier 2 | Q9DB41 | Programmed cell death protein 6 | P12815 |
| Mitochondrial import inner membrane translocase | P62073 | Pro-interleukin-16 | O54824 |
| Mitochondrial import inner membrane translocase | O35857 | Prolactin regulatory element-binding protein | Q9WUQ2 |
| Mitochondrial import inner membrane translocase | Q9D880 | Proline dehydrogenase 1, mitochondrial | Q9WU79 |
| Mitochondrial import receptor subunit TOM34 | Q9CYG7 | Proline-rich AKT1 substrate 1 | Q9D1F4 |
| Mitochondrial import receptor subunit TOM70 | Q9CZW5 | Prostaglandin E synthase 2 | Q8BWM0 |
| Mitochondrial inner membrane protein | Q8CAQ8 | Prostaglandin E synthase 3 | Q9R0Q7 |
| Mitochondrial peptide methionine sulfoxide reducta | Q9D6Y7 | Proteasome activator complex subunit 4 | Q5SSW2 |
| Mitochondrial-processing peptidase subunit beta | Q9CXT8 | Proteasome subunit alpha type-2 | P49722 |
| Mitofusin-2 | Q80U63 | Proteasome subunit beta type-2 | Q9R1P3 |
| Mitogen-activated protein kinase 1 | P63085 | Proteasome subunit beta type-3 | Q9R1P1 |
| Mitogen-activated protein kinase 3 | Q63844 | Proteasome subunit beta type-6 | Q60692 |
| Mitogen-activated protein kinase 9 | Q9WTU6 | Proteasome subunit beta type-7 | P70195 |
| MKL/myocardin-like protein 2 | P59759 | Proteasome-associated protein ECM29 homolog | Q6PDI5 |
| MOB-like protein phocein | Q6PEB6 | Protein 4.1 | P48193 |
| Moesin | P26041 | Protein arginine N-methyltransferase 8 | Q6PAK3 |
| Monoacylglycerol lipase ABHD12 | Q99LR1 | Protein argonaute-1 | Q8CJG1 |
| Monofunctional C1-tetrahydrofolate synthase, mito | Q3V3R1 | Protein argonaute-2 | Q8CJG0 |
| Monoglyceride lipase | O35678 | Protein argonaute-3 | Q8CJF9 |
| Multifunctional protein ADE2 | Q9DCL9 | Protein bicaudal D homolog 2 | Q921C5 |
| Myc box-dependent-interacting protein 1 | O08539 | Protein C8orf37 homolog | Q3UJP5 |
| Myelin basic protein | P04370 | Protein canopy homolog 3 | Q9DAU1 |
| Myelin-associated glycoprotein | P20917 | Protein CDV3 | Q4VAA2 |
| Myelin-oligodendrocyte glycoprotein | Q61885 | Protein cereblon | Q8C7D2 |
| Myeloid-associated differentiation marker | O35682 | Protein CutA | Q9CQ89 |
| Myosin light polypeptide 6 | Q60605 | Protein DDI1 homolog 2 | A2ADY9 |
| Myosin regulatory light chain 12B | Q3THE2 | Protein deltex-3 | Q80V91 |
| Myosin-10 | Q61879 | Protein diaphanous homolog 1 | O08808 |
| Myosin-14 | Q6URW6 | Protein diaphanous homolog 2 | O70566 |
| Myosin-9 | Q8VDD5 | Protein dopey-2 | Q3UHQ6 |
| Myotrophin | P62774 | Protein EFR3 homolog B | Q6ZQ18 |
| Myotubularin-related protein 1 | Q9Z2C4 | Protein ETHE1, mitochondrial | Q9DCM0 |
| Myotubularin-related protein 5 | Q6ZPE2 | Protein FAM134C | Q9CQV4 |
| Myristoylated alanine-rich C-kinase substrate | P26645 | Protein FAM13C | Q9DBR2 |
| N(G),N(G)-dimethylarginine dimethylaminohydrola | Q9CWS0 | Protein FAM160B1 | Q8CDM8 |
| N(G),N(G)-dimethylarginine dimethylaminohydrola | Q99LD8 | Protein FAM171B | Q14CH0 |
| Na(+)/H(+) exchange regulatory cofactor NHE-RF: | P70441 | Protein FAM40A | Q8C079 |
| NAD kinase domain-containing protein 1 | Q8C5H8 | Protein FAM54B | Q9CWE0 |
| NADH dehydrogenase [ubiquinone] 1 alpha subco | Q99LC3 | Protein FAM59A | Q3UFT3 |
| NADH dehydrogenase [ubiquinone] 1 alpha subco | Q7TMF3 | Protein FAM5B | Q6DFY8 |
| NADH dehydrogenase [ubiquinone] 1 alpha subco | Q9ERS2 | Protein FAM65A | Q68FE6 |
| NADH dehydrogenase [ubiquinone] 1 alpha subco | Q9CQ75 | Protein FAM65B | Q80U16 |
| NADH dehydrogenase [ubiquinone] 1 alpha subco | Q9CPP6 | Protein FAM81A | Q3UXZ6 |
| NADH dehydrogenase [ubiquinone] 1 alpha subco | Q9Z1P6 | Protein farnesyltransferase subunit beta | Q8K2I1 |
| NADH dehydrogenase [ubiquinone] 1 alpha subco | Q9DC69 | Protein farnesyltransferase/geranylgeranyltransfer | Q61239 |
| NADH dehydrogenase [ubiquinone] 1 beta subcom | Q9DCS9 | Protein HEXIM1 | Q8R409 |
| NADH dehydrogenase [ubiquinone] 1 beta subcom | Q9CQZ6 | Protein Hook homolog 1 | Q8BIL5 |
| NADH dehydrogenase [ubiquinone] 1 beta subcom | Q9CQH3 | Protein KIAA0664 | Q5SW19 |
| NADH dehydrogenase [ubiquinone] 1 beta subcom | Q9D6J5 | Protein kinase C-binding protein NELL2 | Q61220 |
| NADH dehydrogenase [ubiquinone] flavoprotein 1, | Q91YT0 | Protein LSM12 homolog | Q9D0R8 |
| NADH dehydrogenase [ubiquinone] flavoprotein 2, | Q9D6J6 | Protein LZIC | Q8K3C3 |
| NADH dehydrogenase [ubiquinone] iron-sulfur pro | Q91WD5 | Protein misato homolog 1 | Q2YDW2 |
| NADH dehydrogenase [ubiquinone] iron-sulfur pro | Q9DCT2 | Protein NDRG3 | Q9QYF9 |
| NADH dehydrogenase [ubiquinone] iron-sulfur pro | Q9CXZ1 | Protein NDRG4 | Q8BTG7 |
| NADH dehydrogenase [ubiquinone] iron-sulfur pro | Q8K3J1 | Protein NipSnap homolog 3B | Q9CQE1 |
| NADH-cytochrome b5 reductase 1 | Q9DB73 | Protein O-glucosyltransferase 1 | Q8BYB9 |
| NADH-cytochrome b5 reductase 3 | Q9DCN2 | Protein OS-9 | Q8K2C7 |
| NADH-ubiquinone oxidoreductase 75 kDa subunit, | Q91VD9 | Protein phosphatase 1 regulatory inhibitor subunit | Q8VHQ3 |
| NADP-dependent malic enzyme | P06801 | Protein phosphatase 1 regulatory subunit 28 | Q8C8T7 |
| NADPH--cytochrome P450 reductase | P37040 | Protein phosphatase 1 regulatory subunit 37 | Q8BKR5 |
| N-alpha-acetyltransferase 15, NatA auxiliary subur | Q80UM3 | Protein phosphatase 1F | Q8CGA0 |
| N-alpha-acetyltransferase 25, NatB auxiliary subur | Q8BWZ3 | Protein phosphatase Slingshot homolog 2 | Q5SW75 |
| N-alpha-acetyltransferase 38, NatC auxiliary subur | Q6ZWM4 | Protein RCC2 | Q8BK67 |
| Nardilysin | Q8BHG1 | Protein rogdi homolog | Q3TDK6 |
| Nascent polypeptide-associated complex subunit a | P70670 | Protein S100-A1 | P56565 |
| Nck-associated protein 1 | P28660 | Protein S100-A13 | P97352 |
| NCK-interacting protein with SH3 domain | Q9ESJ4 | Protein S100-A6 | P14069 |
| NEDD8 | P29595 | Protein SCO1 homolog, mitochondrial | Q5SUC9 |
| NEDD8-activating enzyme E1 regulatory subunit | Q8VBW6 | Protein SEC13 homolog | Q9D1M0 |
| NEDD8-conjugating enzyme Ubc12 | P61082 | Protein spire homolog 1 | Q52KF3 |
| Neogenin | P97798 | Protein SZT2 | A2A9C3 |
| Nesprin-1 | Q6ZWR6 | Protein TANC1 | Q0VGY8 |
| Neurabin-2 | Q6R891 | Protein transport protein Sec23A | Q01405 |
| Neural cell adhesion molecule 1 | P13595 | Protein transport protein Sec23B | Q9D662 |
| Neural cell adhesion molecule 2 | O35136 | Protein transport protein Sec61 subunit beta | Q9CQS8 |
| Neural cell adhesion molecule L1 | P11627 | Protein tweety homolog 1 | Q9D3A9 |
| Neural cell adhesion molecule L1-like protein | P70232 | Protein unc-79 homolog | Q0KK59 |
| Neurexin-1-alpha | Q9CS84 | Protein very KIND | Q0KK55 |
| Neurexin-3-alpha | Q6P9K9 | Protein WWC2 | Q6NXJ0 |
| Neurobeachin | Q9EPN1 | Protein XRP2 | Q9EPK2 |
| Neurocalcin-delta | Q91X97 | Protein zer-1 homolog | Q80ZJ6 |
| Neurocan core protein | P55066 | Protein-arginine deiminase type-2 | Q08642 |
| Neurochondrin | Q9Z0E0 | Protein-L-isoaspartate O-methyltransferase domai | P59913 |
| Neurofascin | Q810U3 | Protocadherin-19 | Q80TF3 |
| Neurofibromin | Q04690 | Proton myo-inositol cotransporter | Q3UHK1 |
| Neurofilament heavy polypeptide | P19246 | Pterin-4-alpha-carbinolamine dehydratase | P61458 |
| Neurofilament light polypeptide | P08551 | Pumilio homolog 2 | Q80U58 |
| Neurofilament medium polypeptide | P08553 | Purine-rich element-binding protein gamma | Q8R4E6 |
| Neurolysin, mitochondrial | Q91YP2 | Putative ATP-dependent RNA helicase DHX30 | Q99PU8 |
| Neuromodulin | P06837 | Putative N-acetylglucosamine-6-phosphate deacet | Q8JZV7 |
| Neuron navigator 1 | Q8CH77 | Putative peptidyl-tRNA hydrolase PTRHD1 | D3Z4S3 |
| Neuronal calcium sensor 1 | Q8BNY6 | Putative polypeptide N-acetylgalactosaminyltransfe | Q7TT15 |
| Neuronal cell adhesion molecule | Q810U4 | Putative RNA-binding protein 3 | O89086 |

| | | | |
|---|---|---|---|
| Neuronal growth regulator 1 | Q80Z24 | Pyridoxal-dependent decarboxylase domain-contai | Q99K01 |
| Neuronal membrane glycoprotein M6-a | P35802 | Pyrroline-5-carboxylate reductase 2 | Q922Q4 |
| Neuronal membrane glycoprotein M6-b | P35803 | Pyruvate dehydrogenase phosphatase regulatory s | Q7TSQ8 |
| Neuronal pentraxin receptor | Q99J85 | Quinone oxidoreductase | P47199 |
| Neuronal pentraxin-1 | Q62443 | Quinone oxidoreductase-like protein 1 | Q921W4 |
| Neuronal-specific septin-3 | Q9Z1S5 | R3H domain-containing protein 2 | Q80TM6 |
| Neuron-specific calcium-binding protein hippocalci | P84075 | Rab GTPase-activating protein 1-like | A6H6A9 |
| Neuropilin-2 | O35375 | Rab proteins geranylgeranyltransferase componer | Q9QXG2 |
| Neuroplastin | P97300 | Rab3 GTPase-activating protein catalytic subunit | Q80UJ7 |
| Neurotrimin | Q99PJ0 | Rab-3A-interacting protein | Q68EF0 |
| Neutral alpha-glucosidase AB | Q8BHN3 | Rabenosyn-5 | Q80Y56 |
| Neutral amino acid transporter A | O35874 | RAC-alpha serine/threonine-protein kinase | P31750 |
| NF-kappa-B essential modulator | O88522 | RAC-beta serine/threonine-protein kinase | Q60823 |
| Nicastrin | P57716 | RAC-gamma serine/threonine-protein kinase | Q9WUA6 |
| Nicotinamide phosphoribosyltransferase | Q99KQ4 | RAF proto-oncogene serine/threonine-protein kina | Q99N57 |
| Nischarin | Q80TM9 | Ral GTPase-activating protein subunit alpha-1 | Q6GYP7 |
| Nitric oxide synthase, brain | Q9Z0J4 | Ral GTPase-activating protein subunit beta | Q8BQZ4 |
| Nitrilase homolog 1 | Q8VDK1 | RalBP1-associated Eps domain-containing protein | O54916 |
| Nodal modulator 1 | Q6GQT9 | RalBP1-associated Eps domain-containing protein | Q80XA6 |
| Noelin | O88998 | Ran-binding protein 6 | Q8BIV3 |
| Non-POU domain-containing octamer-binding prot | Q99K48 | Ran-binding protein 9 | P69566 |
| NSFL1 cofactor p47 | Q9CZ44 | Rap1 GTPase-activating protein 2 | Q5SVL6 |
| NT-3 growth factor receptor | Q6VNS1 | Rapamycin-insensitive companion of mTOR | Q6QI06 |
| N-terminal EF-hand calcium-binding protein 2 | Q91ZP9 | Ras and Rab interactor 1 | Q921Q7 |
| Nuclear distribution protein nudE-like 1 | Q9ERR1 | Ras GTPase-activating protein 3 | Q60790 |
| Nuclear export mediator factor Nemf | Q8CCP0 | Ras GTPase-activating-like protein IQGAP1 | Q9JKF1 |
| Nuclear fragile X mental retardation-interacting pro | Q5F2E7 | RAS guanyl-releasing protein 1 | Q9Z1S3 |
| Nuclear migration protein nudC | O35685 | Ras-related C3 botulinum toxin substrate 3 | P60764 |
| Nuclear protein localization protein 4 homolog | P60670 | Ras-related GTP-binding protein B | Q6NTA4 |
| Nuclease EXOG, mitochondrial | Q8C163 | Ras-related protein Rab-12 | P35283 |
| Nucleobindin-1 | Q02819 | Ras-related protein Rab-15 | Q8K386 |
| Nucleobindin-2 | P81117 | Ras-related protein Rab-22A | P35285 |
| Nucleolar protein 3 | Q9D1X0 | Ras-related protein Rab-24 | P35290 |
| Nucleolin | P09405 | Ras-related protein Rab-33B | O35963 |
| Nucleophosmin | Q61937 | Ras-related protein Rab-39B | Q8BHC1 |
| Nucleoside diphosphate kinase A | P15532 | Ras-related protein Rab-5B | P61021 |
| Nucleoside diphosphate kinase B | Q01768 | Ras-related protein Rab-5C | P35278 |
| Nucleosome assembly protein 1-like 4 | Q78ZA7 | Ras-related protein Rab-8A | P55258 |
| NudC domain-containing protein 3 | Q8R1N4 | Ras-related protein Rab-9B | Q8BHH2 |
| Numb-like protein | O08919 | Ras-related protein Rap-2a | Q80ZJ1 |
| Obg-like ATPase 1 | Q9CZ30 | Receptor expression-enhancing protein 1 | Q8BGH4 |
| Oligodendrocyte-myelin glycoprotein | Q63912 | Receptor-type tyrosine-protein phosphatase alpha | P18052 |
| Omega-amidase NIT2 | Q9JHW2 | Receptor-type tyrosine-protein phosphatase epsilo | P49446 |
| Opioid growth factor receptor | Q99PG2 | Receptor-type tyrosine-protein phosphatase eta | Q64455 |
| Ornithine aminotransferase, mitochondrial | P29758 | Receptor-type tyrosine-protein phosphatase gamm | Q05909 |
| Oxidation resistance protein 1 | Q4KMM3 | Regulating synaptic membrane exocytosis protein | Q9EQZ7 |
| Oxysterol-binding protein 1 | Q3B7Z2 | Regulation of nuclear pre-mRNA domain-containir | Q8VDS4 |
| Palmdelphin | Q9JHU2 | Regulator of G-protein signaling 10 | Q9CQE5 |
| Palmitoyltransferase ZDHHC5 | Q8VDZ4 | Regulator of G-protein signaling 12 | Q8CGE9 |
| Pantothenate kinase 4 | Q80YV4 | Regulator of microtubule dynamics protein 1 | Q9DCV4 |
| Paralemmin-1 | Q9Z0P4 | Regulatory-associated protein of mTOR | Q8K4Q0 |
| Paralemmin-2 | Q8BR92 | Retinaldehyde-binding protein 1 | Q9Z275 |
| Paraspeckle component 1 | Q8R326 | Retinal-specific ATP-binding cassette transporter | O35600 |
| Parathymosin | Q9D0J8 | Retinol-binding protein 1 | Q00915 |
| Partner of Y14 and mago | Q8CHP5 | Rho GTPase-activating protein 23 | Q69ZH9 |
| Parvalbumin alpha | P32848 | Rho GTPase-activating protein 33 | Q80YF9 |
| PC4 and SFRS1-interacting protein | Q99JF8 | Rho GTPase-activating protein 5 | P97393 |
| PDZ domain-containing protein GIPC1 | Q9Z0G0 | Rho guanine nucleotide exchange factor 9 | Q3UTH8 |
| Peptidyl-prolyl cis-trans isomerase A | P17742 | Rho-related GTP-binding protein RhoB | P62746 |
| Peptidyl-prolyl cis-trans isomerase B | P24369 | Ribosomal protein S6 kinase alpha-2 | Q9WUT3 |
| Peptidyl-prolyl cis-trans isomerase D | Q9CR16 | Ribosomal protein S6 kinase alpha-3 | P18654 |
| Peptidyl-prolyl cis-trans isomerase F, mitochondria | Q99KR7 | Ribosomal protein S6 kinase alpha-4 | Q9Z2B9 |
| Peptidyl-prolyl cis-trans isomerase FKBP3 | Q62446 | Ribosyldihydronicotinamide dehydrogenase [quino | Q9JI75 |
| Peptidyl-prolyl cis-trans isomerase FKBP4 | P30416 | RING finger protein 126 | Q91YL2 |
| Peptidyl-prolyl cis-trans isomerase FKBP5 | Q64378 | RNA polymerase II-associated protein 3 | Q9D706 |
| Peptidyl-prolyl cis-trans isomerase FKBP8 | O35465 | RNA polymerase-associated protein RTF1 homolo | A2AQ19 |
| Peptidyl-prolyl cis-trans isomerase NIMA-interactir | Q9QUR7 | RNA-binding protein 12 | Q8R4X3 |
| Peripheral plasma membrane protein CASK | O70589 | RNA-binding protein 26 | Q6NZN0 |
| Peroxiredoxin-1 | P35700 | RNA-binding protein 39 | Q8VH51 |
| Peroxiredoxin-2 | Q61171 | RNA-binding protein EWS | Q61545 |
| Peroxiredoxin-6 | O08709 | RNA-binding protein Musashi homolog 2 | Q920Q6 |
| Peroxisomal acyl-coenzyme A oxidase 1 | Q9R0H0 | RUN and FYVE domain-containing protein 1 | Q8BIJ7 |
| Peroxisomal biogenesis factor 19 | Q8VCI5 | RUN and FYVE domain-containing protein 2 | Q8R4C2 |
| Peroxisomal multifunctional enzyme type 2 | P51660 | RUN domain-containing protein 3A | O08576 |
| PEX5-related protein | Q8C437 | RuvB-like 1 | P60122 |
| PH and SEC7 domain-containing protein 3 | Q2PFD7 | Saccharopine dehydrogenase-like oxidoreductase | Q8R127 |
| Phenylalanine--tRNA ligase alpha subunit | Q8C0C7 | Sacsin | Q9JLC8 |
| Phenylalanine--tRNA ligase beta subunit | Q9WUA2 | SAM domain and HD domain-containing protein 1 | Q60710 |
| Phosphatase and actin regulator 1 | Q2M3X8 | Sarcolemmal membrane-associated protein | Q3URD3 |
| Phosphatase and actin regulator 3 | Q8BYK5 | Sarcosine dehydrogenase, mitochondrial | Q99LB7 |
| Phosphate carrier protein, mitochondrial | Q8VEM8 | Sec1 family domain-containing protein 1 | Q8BRF7 |
| Phosphatidate cytidylyltransferase 2 | Q99L43 | SEC14 domain and spectrin repeat-containing pro | Q80UK0 |
| Phosphatidylethanolamine-binding protein 1 | P70296 | SEC14-like protein 2 | Q99J08 |
| Phosphatidylinositide phosphatase SAC1 | Q9EP69 | SEC23-interacting protein | Q6NZC7 |
| Phosphatidylinositide phosphatase SAC2 | Q8CDA1 | Secernin-3 | Q3TMH2 |
| Phosphatidylinositol 3,4,5-trisphosphate-dependen | Q69ZK0 | Secretory carrier-associated membrane protein 3 | O35609 |
| Phosphatidylinositol 3-kinase regulatory subunit be | O08908 | Sel1 repeat-containing protein 1 | Q921H9 |
| Phosphatidylinositol 4-phosphate 5-kinase type-1 g | O70161 | Selenocysteine lyase | Q9JLI6 |
| Phosphatidylinositol 5-phosphate 4-kinase type-2 a | O70172 | Selenocysteine-specific elongation factor | Q9JHW4 |
| Phosphatidylinositol 5-phosphate 4-kinase type-2 b | Q80XI4 | Semaphorin-4A | Q62178 |
| Phosphatidylinositol 5-phosphate 4-kinase type-2 g | Q91XU3 | Semaphorin-7A | Q9QUR8 |
| Phosphatidylinositol transfer protein alpha isoform | P53810 | Septin-2 | P42208 |
| Phosphatidylinositol transfer protein beta isoform | P53811 | Septin-6 | Q9R1T4 |
| Phosphatidylinositol-binding clathrin assembly prot | Q7M6Y3 | Serine beta-lactamase-like protein LACTB, mitoch | Q9EP89 |

| | | | |
|---|---|---|---|
| Phosphoenolpyruvate carboxykinase [GTP], mitocl | Q8BH04 | Serine protease HTRA1 | Q9R118 |
| Phosphofurin acidic cluster sorting protein 1 | Q8K212 | Serine racemase | Q9QZX7 |
| Phosphoglucomutase-1 | Q9D0F9 | Serine/arginine-rich splicing factor 3 | P84104 |
| Phosphoglycerate kinase 1 | P09411 | Serine/arginine-rich splicing factor 5 | O35326 |
| Phosphoglycerate mutase 1 | Q9DBJ1 | Serine/arginine-rich splicing factor 7 | Q8BL97 |
| Phosphoglycerate mutase 2 | O70250 | Serine/threonine-protein kinase 24 | Q99KH8 |
| Phosphoglycolate phosphatase | Q8CHP8 | Serine/threonine-protein kinase 38-like | Q7TSE6 |
| Phospholipase D3 | O35405 | Serine/threonine-protein kinase A-Raf | P04627 |
| Phospholipid hydroperoxide glutathione peroxidas | O70325 | Serine/threonine-protein kinase B-raf | P28028 |
| Phosphomannomutase 1 | O35621 | Serine/threonine-protein kinase LMTK2 | Q3TYD6 |
| Phosphoribosylformylglycinamidine synthase | Q5SUR0 | Serine/threonine-protein kinase LMTK3 | Q5XJV6 |
| Phosphoserine aminotransferase | Q99K85 | Serine/threonine-protein kinase MRCK beta | Q7TT50 |
| Phytanoyl-CoA hydroxylase-interacting protein | Q8K0S0 | Serine/threonine-protein kinase N1 | P70268 |
| Phytanoyl-CoA hydroxylase-interacting protein-like | Q8BGT8 | Serine/threonine-protein kinase Nek7 | Q9ES74 |
| PITH domain-containing protein 1 | Q8BWR2 | Serine/threonine-protein kinase SIK3 | Q6P4S6 |
| Plakophilin-4 | Q68FH0 | Serine/threonine-protein kinase SMG1 | Q8BKX6 |
| Plasma membrane calcium-transporting ATPase 2 | Q9R0K7 | Serine/threonine-protein kinase STK11 | Q9WTK7 |
| Plasminogen activator inhibitor 1 RNA-binding pro | Q9CY58 | Serine/threonine-protein kinase TAO3 | Q8BYC6 |
| Plastin-2 | Q61233 | Serine/threonine-protein kinase ULK3 | Q3U3Q1 |
| Plastin-3 | Q99K51 | Serine/threonine-protein phosphatase 4 catalytic s | P97470 |
| Platelet-activating factor acetylhydrolase IB subuni | P63005 | Serine/threonine-protein phosphatase 4 regulatory | Q6P2K6 |
| Plectin | Q9QXS1 | Serine/threonine-protein phosphatase 4 regulatory | Q8C0Y0 |
| Plexin-A1 | P70206 | Serine/threonine-protein phosphatase 6 regulatory | Q8R3Q2 |
| Plexin-A4 | Q80UG2 | Serine/threonine-protein phosphatase PP1-alpha c | P62137 |
| Plexin-B1 | Q8CJH3 | Serine/threonine-protein phosphatase PP1-beta ca | P62141 |
| Plexin-B2 | B2RXS4 | Serine/threonine-protein phosphatase PP1-gamma | P63087 |
| Poly(ADP-ribose) glycohydrolase ARH3 | Q8CG72 | Serine--tRNA ligase, mitochondrial | Q9JJL8 |
| Poly(rC)-binding protein 1 | P60335 | Serpin H1 | P19324 |
| Poly(rC)-binding protein 2 | Q61990 | S-formylglutathione hydrolase | Q9R0P3 |
| Polyadenylate-binding protein 1 | P29341 | SH2B adapter protein 1 | Q91ZM2 |
| Polyadenylate-binding protein-interacting protein 1 | Q8VE62 | SH3 domain-binding glutamic acid-rich-like protein | Q91VW3 |
| Polyubiquitin-C | P0CG50 | SHC-transforming protein 3 | Q61120 |
| Porphobilinogen deaminase | P22907 | Short coiled-coil protein | Q78YZ6 |
| Potassium/sodium hyperpolarization-activated cyc | O88703 | SID1 transmembrane family member 2 | Q8CIF6 |
| Prefoldin subunit 1 | Q9CWM4 | Sideroflexin-1 | Q99JR1 |
| Prefoldin subunit 2 | O70591 | Sideroflexin-5 | Q925N0 |
| Prefoldin subunit 3 | P61759 | Signal peptidase complex catalytic subunit SEC11 | Q9R0P6 |
| Pre-mRNA-processing factor 19 | Q99KP6 | Signal peptidase complex catalytic subunit SEC11 | Q9D8V7 |
| Presequence protease, mitochondrial | Q8K411 | Signal peptidase complex subunit 2 | Q9CYN2 |
| Prickle-like protein 2 | Q80Y24 | Signal recognition particle 54 kDa protein | P14576 |
| PRKCA-binding protein | Q62083 | Signal recognition particle 68 kDa protein | Q8BMA6 |
| Probable ATP-dependent RNA helicase DDX17 | Q501J6 | Signal recognition particle receptor subunit beta | P47758 |
| Probable ATP-dependent RNA helicase DDX5 | Q61656 | Signal transducer and activator of transcription 5B | P42232 |
| Probable E3 ubiquitin-protein ligase MYCBP2 | Q7TPH6 | Signal transducing adapter molecule 1 | P70297 |
| Probable G-protein coupled receptor 158 | Q8C419 | Signal transducing adapter molecule 2 | O88811 |
| Probable phospholipid-transporting ATPase IA | P70704 | Signal-induced proliferation-associated 1-like prote | Q80TE4 |
| Probable threonine--tRNA ligase 2, cytoplasmic | Q8BLY2 | Sister chromatid cohesion protein PDS5 homolog | Q4VA53 |
| Probable ubiquitin carboxyl-terminal hydrolase FAI | P70398 | SLIT and NTRK-like protein 1 | Q810C1 |
| Programmed cell death 6-interacting protein | Q9WU78 | SLIT and NTRK-like protein 3 | Q810B9 |
| Programmed cell death protein 10 | Q8VE70 | SLIT-ROBO Rho GTPase-activating protein 1 | Q91Z69 |
| Programmed cell death protein 5 | P56812 | Small acidic protein | Q9R0P4 |
| Prohibitin | P67778 | Small G protein signaling modulator 1 | Q8BPQ7 |
| Prohibitin-2 | O35129 | Small glutamine-rich tetratricopeptide repeat-conta | Q8VD33 |
| Proliferation-associated protein 2G4 | P50580 | Small ubiquitin-related modifier 2 | P61957\|Q9Z172 |
| Proline synthase co-transcribed bacterial homolog | Q9Z2Y8 | Small ubiquitin-related modifier 3 | Q9Z172 |
| Proline-rich transmembrane protein 1 | O35449 | Smith-Magenis syndrome chromosomal region can | Q3UMB5 |
| Proline-rich transmembrane protein 2 | E9PUL5 | SNARE-associated protein Snapin | Q9Z266 |
| Prolow-density lipoprotein receptor-related protein | Q91ZX7 | SNF-related serine/threonine-protein kinase | Q8VDU5 |
| Prolyl endopeptidase | Q9QUR6 | Sodium- and chloride-dependent GABA transporte | P31648 |
| Prolyl endopeptidase-like | Q8C167 | Sodium bicarbonate cotransporter 3 | Q8BTY2 |
| Propionyl-CoA carboxylase alpha chain, mitochono | Q91ZA3 | Sodium channel protein type 9 subunit alpha | Q62205 |
| Propionyl-CoA carboxylase beta chain, mitochondr | Q99MN9 | Sodium/potassium/calcium exchanger 4 | Q8CGQ8 |
| ProSAAS | Q9QXV0 | Solute carrier family 12 member 6 | Q924N4 |
| ProSAP-interacting protein 1 | A2AHG0 | Solute carrier family 12 member 9 | Q99MR3 |
| Prostaglandin-H2 D-isomerase | O09114 | Solute carrier family 2, facilitated glucose transpor | P17809 |
| Prostamide/prostaglandin F synthase | Q9DB60 | Solute carrier family 2, facilitated glucose transpor | P32037 |
| Proteasomal ubiquitin receptor ADRM1 | Q9JKV1 | Sorbin and SH3 domain-containing protein 2 | Q3UTJ2 |
| Proteasome activator complex subunit 1 | P97371 | Sorcin | Q6P069 |
| Proteasome activator complex subunit 2 | P97372 | Sortilin-related receptor | O88307 |
| Proteasome activator complex subunit 3 | P61290 | Sorting nexin-12 | O70493 |
| Proteasome subunit alpha type-1 | Q9R1P4 | Sorting nexin-17 | Q8BVL3 |
| Proteasome subunit alpha type-3 | O70435 | Sorting nexin-3 | O70492 |
| Proteasome subunit alpha type-4 | Q9R1P0 | Sorting nexin-30 | Q8CE50 |
| Proteasome subunit alpha type-5 | Q9Z2U1 | Sorting nexin-32 | Q80ZJ7 |
| Proteasome subunit alpha type-6 | Q9QUM9 | SPARC | P07214 |
| Proteasome subunit alpha type-7 | Q9Z2U0 | Spartin | Q8R1X6 |
| Proteasome subunit beta type-1 | O09061 | Spastin | Q9QYY8 |
| Proteasome subunit beta type-4 | P99026 | Spermatogenesis-associated protein 7 homolog | Q80VP2 |
| Proteasome subunit beta type-5 | O55234 | Spermatogenesis-associated serine-rich protein 2 | Q8K1N4 |
| Protein arginine N-methyltransferase 1 | Q9JIF0 | Spermatogenesis-defective protein 39 homolog | Q8BGQ1 |
| Protein arginine N-methyltransferase 5 | Q8CIG8 | Spermidine synthase | Q64674 |
| Protein bassoon | O88737 | Sphingomyelin phosphodiesterase | Q04519 |
| Protein BRICK1 | Q91VR8 | Splicing factor 3A subunit 1 | Q8K4Z5 |
| Protein canopy homolog 2 | Q9QXT0 | Splicing factor 3A subunit 2 | Q62203 |
| Protein disulfide-isomerase | P09103 | Splicing factor 3A subunit 3 | Q9D554 |
| Protein disulfide-isomerase A3 | P27773 | Splicing factor 3B subunit 1 | Q99NB9 |
| Protein disulfide-isomerase A4 | P08003 | Splicing factor 3B subunit 3 | Q921M3 |
| Protein disulfide-isomerase A6 | Q922R8 | Splicing factor U2AF 65 kDa subunit | P26369 |
| Protein DJ-1 | Q99LX0 | Spondin-1 | Q8VCC9 |
| Protein EFR3 homolog A | Q8BG67 | SPRY domain-containing protein 4 | Q91WK1 |
| Protein enabled homolog | Q03173 | SRA stem-loop-interacting RNA-binding protein, m | Q9D8T7 |
| Protein FAM114A2 | Q8VE88 | Steroid receptor RNA activator 1 | Q80VJ2 |
| Protein FAM126B | Q8C729 | Sterol regulatory element-binding protein 2 | Q3U1N2 |
| Protein FAM136A | Q9CR98 | Sterol-4-alpha-carboxylate 3-dehydrogenase, deca | Q9R1J0 |

| | | | |
|---|---|---|---|
| Protein FAM169A | Q5XG69 | Stromal interaction molecule 1 | P70302 |
| Protein FAM192A | Q91WE2 | Stromal membrane-associated protein 2 | Q7TN29 |
| Protein FAM49A | Q8BHZ0 | Structural maintenance of chromosomes protein 3 | Q9CW03 |
| Protein FAM49B | Q921M7 | Succinate dehydrogenase assembly factor 1, mito | Q3U276 |
| Protein FAM98B | Q80VD1 | Sulfite oxidase, mitochondrial | Q8R086 |
| Protein fat-free homolog | Q3UVL4 | Superoxide dismutase [Mn], mitochondrial | P09671 |
| Protein flightless-1 homolog | Q9JJ28 | Switch-associated protein 70 | Q6A028 |
| Protein Hook homolog 3 | Q8BUK6 | Synapse-associated protein 1 | Q9D5V6 |
| Protein IMPACT | O55091 | Synapsin-3 | Q8JZP2 |
| Protein KIAA0284 | Q80U49 | Synaptic vesicle membrane protein VAT-1 homolo | Q62465 |
| Protein KIAA1045 | Q80TL4 | Synaptogyrin-3 | Q8R191 |
| Protein kinase C alpha type | P20444 | Synaptojanin-2-binding protein | Q9D6K5 |
| Protein kinase C and casein kinase substrate in ne | Q61644 | Synaptophysin | Q62277 |
| Protein kinase C and casein kinase substrate in ne | Q9WVE8 | Synaptoporin | Q8BGN8 |
| Protein kinase C beta type | P68404 | Synaptotagmin-12 | Q920N7 |
| Protein kinase C epsilon type | P16054 | Synaptotagmin-13 | Q9EQT6 |
| Protein kinase C gamma type | P63318 | Synaptotagmin-2 | P46097 |
| Protein lin-7 homolog A | Q8JZS0 | Synaptotagmin-3 | O35681 |
| Protein lin-7 homolog B | O88951 | Synaptotagmin-5 | Q9R0N5 |
| Protein lin-7 homolog C | O88952 | Synembryn-A | Q3TIR3 |
| Protein lunapark | Q7TQ95 | Syntabulin | Q8BHS8 |
| Protein LYRIC | Q80WJ7 | Syntaxin-16 | Q8BVI5 |
| Protein MON2 homolog | Q80TL7 | Syntaxin-6 | Q9JKK1 |
| Protein NDRG2 | Q9QYG0 | Syntaxin-binding protein 3 | Q60770 |
| Protein NipSnap homolog 1 | O55125 | Syntenin-1 | O08992 |
| Protein phosphatase 1 regulatory subunit 12A | Q9DBR7 | Talin-1 | P26039 |
| Protein phosphatase 1 regulatory subunit 12C | Q3UMT1 | TAR DNA-binding protein 43 | Q921F2 |
| Protein phosphatase 1 regulatory subunit 1A | Q9ERT9 | Target of Myb protein 1 | O88746 |
| Protein phosphatase 1 regulatory subunit 21 | Q3TDD9 | Target of rapamycin complex subunit LST8 | Q9DCJ1 |
| Protein phosphatase 1 regulatory subunit 29 | Q68FM6 | Tax1-binding protein 1 homolog | Q3UKC1 |
| Protein phosphatase 1 regulatory subunit 7 | Q3UM45 | TBC1 domain family member 10B | Q8BHL3 |
| Protein phosphatase 1A | P49443 | TBC1 domain family member 15 | Q9CXF4 |
| Protein phosphatase 1B | P36993 | TBC1 domain family member 17 | Q8BYH7 |
| Protein phosphatase 1E | Q80TL0 | TBC1 domain family member 7 | Q9D0K0 |
| Protein phosphatase 1G | Q61074 | TBC1 domain family member 8B | A3KGB4 |
| Protein phosphatase 1H | Q3UYC0 | TBC1 domain family member 9B | Q5SVR0 |
| Protein phosphatase methylesterase 1 | Q8BVQ5 | T-cell immunomodulatory protein | Q99KW9 |
| Protein piccolo | Q9QYX7 | T-complex protein 11-like protein 1 | Q8BTG3 |
| Protein PRRC2A | Q7TSC1 | Teneurin-2 | Q9WTS5 |
| Protein PRRC2C | Q3TLH4 | Teneurin-3 | Q9WTS6 |
| Protein prune homolog | Q8BIW1 | Tensin-3 | Q5SSZ5 |
| Protein QIL1 | Q8R404 | Terminal uridylyltransferase 7 | Q5BLK4 |
| Protein quaking | Q9QYS9 | Tescalcin | Q9JKL5 |
| Protein RUFY3 | Q9D394 | Testican-2 | Q9ER58 |
| Protein SCAI | Q8C8N2 | Testis-specific Y-encoded-like protein 4 | Q8VD63 |
| Protein SET | Q9EQU5 | Tether containing UBX domain for GLUT4 | Q8VBT9 |
| Protein shisa-6 homolog | Q3UH99 | Tetratricopeptide repeat protein 1 | Q91Z38 |
| Protein shisa-7 | Q8C3Q5 | Tetratricopeptide repeat protein 19, mitochondrial | Q8CC21 |
| Protein Shroom2 | A2ALU4 | Tetratricopeptide repeat protein 35 | Q9CRD2 |
| Protein TANC2 | A2A690 | Tetratricopeptide repeat protein 38 | A3KMP2 |
| Protein transport protein Sec31A | Q3UPL0 | Tetratricopeptide repeat protein 9A | Q3V038 |
| Protein TSSC1 | Q8K0G5 | Tetratricopeptide repeat protein 9C | Q810A3 |
| Protein tweety homolog 3 | Q6P5F7 | TGF-beta-activated kinase 1 and MAP3K7-binding | Q8CF89 |
| Protein unc-13 homolog A | Q4KUS2 | Thioesterase superfamily member 4 | Q3UUI3 |
| Protein-L-isoaspartate(D-aspartate) O-methyltrans | P23506 | Thiopurine S-methyltransferase | O55060 |
| Protein-methionine sulfoxide oxidase MICAL3 | Q8CJ19 | Thioredoxin domain-containing protein 12 | Q9CQU0 |
| Protein-tyrosine kinase 2-beta | Q9QVP9 | Thioredoxin domain-containing protein 5 | Q91W90 |
| Prothymosin alpha | P26350 | Thioredoxin reductase 2, mitochondrial | Q9JLT4 |
| Pumilio homolog 1 | Q80U78 | Thioredoxin-related transmembrane protein 2 | Q9D710 |
| Purine nucleoside phosphorylase | P23492 | THO complex subunit 4 | O08583 |
| Purkinje cell protein 4 | P63054 | Threonine--tRNA ligase, mitochondrial | Q3UQ84 |
| Puromycin-sensitive aminopeptidase | Q11011 | Thymidylate kinase | P97930 |
| Putative adenosylhomocysteinase 2 | Q80SW1 | Thymocyte nuclear protein 1 | Q91YJ3 |
| Putative adenosylhomocysteinase 3 | Q68FL4 | Thymosin beta-4 | P20065 |
| Putative ATP-dependent Clp protease proteolytic s | O88696 | Thyroid hormone receptor-associated protein 3 | Q569Z6 |
| Putative phospholipase B-like 2 | Q3TCN2 | Thyrotropin-releasing hormone-degrading ectoenz | Q8K093 |
| Putative pre-mRNA-splicing factor ATP-dependent | O35286 | Tight junction protein ZO-2 | Q9Z0U1 |
| Putative RNA-binding protein Luc7-like 1 | Q9CYI4 | TNF receptor-associated factor 3 | Q60803 |
| Putative RNA-binding protein Luc7-like 2 | Q7TNC4 | Tomoregulin-1 | Q6PFE7 |
| Putative tyrosine-protein phosphatase auxilin | Q80TZ3 | TP53-regulated inhibitor of apoptosis 1 | Q9D8Z2 |
| Pyridoxal kinase | Q8K183 | TPR and ankyrin repeat-containing protein 1 | Q8BV79 |
| Pyridoxal phosphate phosphatase | P60487 | Trafficking protein particle complex subunit 10 | Q3TLI0 |
| Pyridoxine-5'-phosphate oxidase | Q91XF0 | Trafficking protein particle complex subunit 2-like p | Q9JME7 |
| Pyruvate carboxylase, mitochondrial | Q05920 | Trafficking protein particle complex subunit 3 | O55013 |
| Pyruvate dehydrogenase E1 component subunit al | P35486 | Trafficking protein particle complex subunit 4 | Q9ES56 |
| Pyruvate dehydrogenase E1 component subunit be | Q9D051 | Transcription elongation factor A protein-like 3 | Q8R0A5 |
| Pyruvate dehydrogenase protein X component, mi | Q8BKZ9 | Transcription elongation factor B polypeptide 1 | P83940 |
| Pyruvate kinase isozymes M1/M2 | P52480 | Transcription elongation factor SPT5 | O55201 |
| Rab GDP dissociation inhibitor alpha | P50396 | Transcription factor BTF3 homolog 4 | Q9CQH7 |
| Rab GDP dissociation inhibitor beta | Q61598 | Transcription factor E3 | Q64092 |
| Rab GTPase-activating protein 1 | A2AWA9 | Transcription factor HIVEP2 | Q3UHF7 |
| Rab GTPase-binding effector protein 1 | O35551 | Transcription factor RFX3 | P48381 |
| Rab GTPase-binding effector protein 2 | Q91WG2 | Transcriptional regulator ATRX | Q61687 |
| Rab11 family-interacting protein 5 | Q8R361 | Transformer-2 protein homolog beta | P62996 |
| Rab3 GTPase-activating protein non-catalytic subu | Q8BMG7 | Transient receptor potential cation channel subfam | Q9WTR1 |
| Rab5 GDP/GTP exchange factor | Q9JM13 | Translation initiation factor eIF-2B subunit alpha | Q99LC8 |
| Rab-like protein 1 | Q5U3K5 | Translation initiation factor eIF-2B subunit beta | Q99LD9 |
| Rabphilin-3A | P47708 | Translation initiation factor eIF-2B subunit delta | Q61749 |
| Radixin | P26043 | Translation initiation factor eIF-2B subunit epsilon | Q8CHW4 |
| RalA-binding protein 1 | Q62172 | Translational activator of cytochrome c oxidase 1 | Q8K0Z7 |
| Ran GTPase-activating protein 1 | P46061 | Translin | Q62348 |
| Ran-binding protein 3 | Q9CT10 | Translocon-associated protein subunit delta | Q62186 |
| Ran-specific GTPase-activating protein | P34022 | Transmembrane 9 superfamily member 4 | Q8BH24 |
| Rap guanine nucleotide exchange factor 2 | Q8CHG7 | Transmembrane emp24 domain-containing protein | Q9D1D4 |

| | | | |
|---|---|---|---|
| Rap guanine nucleotide exchange factor 4 | Q9EQZ6 | Transmembrane protein 2 | Q5FWI3 |
| Ras GTPase-activating protein SynGAP | F6SEU4 | Transmembrane protein 65 | Q4VAE3 |
| Ras GTPase-activating protein-binding protein 1 | P97855 | Transportin-1 | Q8BFY9 |
| Ras GTPase-activating protein-binding protein 2 | P97379 | Transportin-2 | Q99LG2 |
| Ras GTPase-activating-like protein IQGAP2 | Q3UQ44 | Transportin-3 | Q6P2B1 |
| Ras suppressor protein 1 | Q01730 | Tripartite motif-containing protein 2 | Q9ESN6 |
| RasGAP-activating-like protein 1 | Q9Z268 | Tripartite motif-containing protein 3 | Q9R1R2 |
| Ras-related C3 botulinum toxin substrate 1 | P63001 | Tripartite motif-containing protein 46 | Q7TNM2 |
| Ras-related protein M-Ras | O08989 | tRNA (adenine(58)-N(1))-methyltransferase non-ca | Q8CE96 |
| Ras-related protein Rab-10 | P61027 | tRNA (cytosine(34)-C(5))-methyltransferase | Q1HFZ0 |
| Ras-related protein Rab-11B | P46638 | tRNA-dihydrouridine(47) synthase [NAD(P)(+)]-like | Q91XI1 |
| Ras-related protein Rab-14 | Q91V41 | Trophoblast glycoprotein | Q9Z0L0 |
| Ras-related protein Rab-18 | P35293 | Tropomodulin-1 | P49813 |
| Ras-related protein Rab-1A | P62821 | Trypsin - Sus scrofa (Pig). | P00761 |
| Ras-related protein Rab-1B | Q9D1G1 | Tryptophan 5-hydroxylase 2 | Q8CGV2 |
| Ras-related protein Rab-21 | P35282 | Tubulin alpha chain-like 3 | Q3UX10 |
| Ras-related protein Rab-27B | Q99P58 | Tubulin beta-2A chain | Q7TMM9 |
| Ras-related protein Rab-2A | P53994 | Tubulin beta-4B chain | P68372 |
| Ras-related protein Rab-35 | Q6PHN9 | Tubulin beta-5 chain | P99024 |
| Ras-related protein Rab-3A | P63011 | Tubulin beta-6 chain | Q922F4 |
| Ras-related protein Rab-3B | Q9CZT8 | Tubulin-specific chaperone cofactor E-like protein | Q8C5W3 |
| Ras-related protein Rab-3C | P62823 | Tubulin-specific chaperone E | Q8CIV8 |
| Ras-related protein Rab-4A | P56371 | Tubulin--tyrosine ligase | P38585 |
| Ras-related protein Rab-4B | Q91ZR1 | Tubulin--tyrosine ligase-like protein 12 | Q3UDE2 |
| Ras-related protein Rab-5A | Q9CQD1 | Tumor necrosis factor alpha-induced protein 8-like | Q3TBL6 |
| Ras-related protein Rab-6A | P35279 | Tumor protein D52 | Q62393 |
| Ras-related protein Rab-6B | P61294 | Tumor suppressor p53-binding protein 1 | P70399 |
| Ras-related protein Rab-7a | P51150 | Tumor susceptibility gene 101 protein | Q61187 |
| Ras-related protein Rab-8B | P61028 | Tyrosine-protein kinase CSK | P41241 |
| Ras-related protein Ral-A | P63321 | Tyrosine-protein phosphatase non-receptor type 9 | O35239 |
| Ras-related protein Ral-B | Q9JIW9 | Tyrosine-protein phosphatase non-receptor type su | P97797 |
| Ras-related protein Rap-1A | P62835 | Tyrosine--tRNA ligase, mitochondrial | Q8BYL4 |
| Ras-related protein Rap-2b | P61226 | U2 small nuclear ribonucleoprotein B'' | Q9CQI7 |
| Ras-related protein R-Ras2 | P62071 | U6 snRNA-associated Sm-like protein LSm1 | Q8VC85 |
| Ras-specific guanine nucleotide-releasing factor 1 | P27671 | U6 snRNA-associated Sm-like protein LSm3 | P62311 |
| Receptor expression-enhancing protein 2 | Q8VCD6 | U6 snRNA-associated Sm-like protein LSm4 | Q9QXA5 |
| Receptor expression-enhancing protein 5 | Q60870 | U6 snRNA-associated Sm-like protein LSm7 | Q9CQQ8 |
| Receptor-type tyrosine-protein phosphatase delta | Q64487 | U8 snoRNA-decapping enzyme | Q6P3D0 |
| Receptor-type tyrosine-protein phosphatase N2 | P80560 | Ubiquilin-1 | Q8R317 |
| Receptor-type tyrosine-protein phosphatase S | B0V2N1 | Ubiquilin-2 | Q9QZM0 |
| Redox-regulatory protein FAM213A | Q9CYH2 | Ubiquilin-4 | Q99NB8 |
| Regulating synaptic membrane exocytosis protein | Q99NE5 | Ubiquitin carboxyl-terminal hydrolase 11 | Q99K46 |
| Regulator of G-protein signaling 14 | P97492 | Ubiquitin carboxyl-terminal hydrolase 24 | B1AY13 |
| Regulator of G-protein signaling 6 | Q9Z2H2 | Ubiquitin carboxyl-terminal hydrolase 4 | P35123 |
| Regulator of G-protein signaling 7 | O54829 | Ubiquitin carboxyl-terminal hydrolase 47 | Q8BY87 |
| Regulator of nonsense transcripts 1 | Q9EPU0 | Ubiquitin carboxyl-terminal hydrolase isozyme L5 | Q9WUP7 |
| Reticulocalbin-1 | Q05186 | Ubiquitin conjugation factor E4 B | Q9ES00 |
| Reticulocalbin-2 | Q8BP92 | Ubiquitin-associated protein 2 | Q91VX2 |
| Reticulon-1 | Q8K0T0 | Ubiquitin-conjugating enzyme E2 Q1 | Q7TSS2 |
| Reticulon-3 | Q9ES97 | Ubiquitin-fold modifier-conjugating enzyme 1 | Q9CR09 |
| Reticulon-4 | Q99P72 | Ubiquitin-like modifier-activating enzyme ATG7 | Q9D906 |
| Retinal dehydrogenase 1 | P24549 | Ubiquitin-like protein 4A | P21126 |
| Retinol dehydrogenase 14 | Q9ERI6 | Ubiquitin-like-conjugating enzyme ATG3 | Q9CPX6 |
| Rho GDP-dissociation inhibitor 1 | Q99PT1 | Ubiquitin-protein ligase E3A | O08759 |
| Rho GDP-dissociation inhibitor 2 | Q61599 | Ubiquitin-protein ligase E3B | Q9ES34 |
| Rho GTPase-activating protein 1 | Q5FWK3 | Ubiquitin-protein ligase E3C | Q80U95 |
| Rho GTPase-activating protein 21 | Q6DFV3 | UBX domain-containing protein 2B | Q0KL01 |
| Rho GTPase-activating protein 26 | Q6ZQ82 | UDP-glucose:glycoprotein glucosyltransferase 1 | Q6P5E4 |
| Rho GTPase-activating protein 32 | Q811P8 | UDP-N-acetylhexosamine pyrophosphorylase | Q91YN5 |
| Rho GTPase-activating protein 35 | Q91YM2 | UDP-N-acetylhexosamine pyrophosphorylase-like | Q3TW96 |
| Rho GTPase-activating protein 39 | P59281 | Uncharacterized glycosyltransferase AGO61 | Q8BW41 |
| Rho GTPase-activating protein 44 | Q5SSM3 | Uncharacterized membrane protein C1orf95 homo | Q0VBF8 |
| Rho guanine nucleotide exchange factor 12 | Q8R4H2 | Uncharacterized protein C11orf96 homolog | Q3UPL5 |
| Rho guanine nucleotide exchange factor 2 | Q60875 | Uncharacterized protein C18orf25 homolog | Q8BH50 |
| Rho guanine nucleotide exchange factor 7 | Q9ES28 | Uncharacterized protein C19orf52 homolog | Q8BGX2 |
| Rho-associated protein kinase 2 | P70336 | Uncharacterized protein C1orf43 homolog | Q8R092 |
| Ribonuclease inhibitor | Q91VI7 | Uncharacterized protein KIAA0930 homolog | Q3UE31 |
| Ribonuclease UK114 | P52760 | Uncharacterized protein KIAA1467 | Q8BYI8 |
| Ribose-phosphate pyrophosphokinase 1 | Q9D7G0 | Unconventional myosin-Ib | P46735 |
| Ribosomal protein S6 kinase delta-1 | Q8BLK9 | Unconventional myosin-Ic | Q9WTI7 |
| Ribosome maturation protein SBDS | P70122 | Unconventional myosin-IXa | Q8C170 |
| Ribosome-binding protein 1 | Q99PL5 | UPF0160 protein MYG1, mitochondrial | Q9JK81 |
| Ribosome-recycling factor, mitochondrial | Q9D6S7 | UPF0556 protein C19orf10 homolog | Q9CPT4 |
| RILP-like protein 1 | Q9JJC6 | UPF0600 protein C5orf51 homolog | Q8BR90 |
| RIMS-binding protein 2 | Q80U40 | UPF0670 protein C8orf55 homolog | Q80ZW2 |
| RING finger protein 214 | Q8BFU3 | UPF0687 protein C20orf27 homolog | Q9D1K7 |
| RNA-binding protein FUS | P56959 | UPF0696 protein C11orf68 homolog | Q8VD62 |
| Rootletin | Q8CJ40 | Uridine-cytidine kinase-like 1 | Q91YL3 |
| Roundabout homolog 1 | O89026 | Uroporphyrinogen decarboxylase | P70697 |
| Roundabout homolog 2 | Q7TPD3 | UV excision repair protein RAD23 homolog A | P54726 |
| RuvB-like 2 | Q9WTM5 | Vacuolar protein sorting-associated protein 13A | Q5H8C4 |
| RWD domain-containing protein 1 | Q9CQK7 | Vacuolar protein sorting-associated protein 16 hon | Q920Q4 |
| Ryanodine receptor 2 | E9Q401 | Vacuolar protein sorting-associated protein 18 hon | Q8R307 |
| S-adenosylmethionine synthase isoform type-2 | Q3THS6 | Vacuolar protein sorting-associated protein 28 hon | Q9D1C8 |
| Sarcoplasmic/endoplasmic reticulum calcium ATP | O55143 | Vacuolar protein sorting-associated protein 33A | Q9D2N9 |
| Scavenger mRNA-decapping enzyme DcpS | Q9DAR7 | Vacuolar protein sorting-associated protein 37C | Q8R105 |
| SCY1-like protein 2 | Q8CFE4 | Vacuolar protein sorting-associated protein 41 hon | Q5KU39 |
| Secernin-1 | Q9CZC8 | Vacuolar protein sorting-associated protein 4A | Q8VEJ9 |
| Secretogranin-1 | P16014 | Vacuolar protein sorting-associated protein 4B | P46467 |
| Secretogranin-2 | Q03517 | Vacuolar protein sorting-associated protein 52 hon | Q8C754 |
| Secretogranin-3 | P47867 | Vacuolar protein sorting-associated protein 8 homo | Q0P5W1 |
| Secretory carrier-associated membrane protein 1 | Q8K021 | Vacuolar protein sorting-associated protein VTA1 h | Q9CR26 |
| Seizure 6-like protein | Q6P1D5 | Vacuolar protein-sorting-associated protein 36 | Q91XD6 |
| Seizure 6-like protein 2 | Q4V9Z5 | Vacuolar-sorting protein SNF8 | Q9CZ28 |

| | |
|---|---|
| Selenium-binding protein 2 | Q63836 |
| Sepiapterin reductase | Q64105 |
| Septin-11 | Q8C1B7 |
| Septin-5 | Q9Z2Q6 |
| Septin-7 | O55131 |
| Septin-8 | Q8CHH9 |
| Septin-9 | Q80UG5 |
| Serine/arginine-rich splicing factor 1 | Q6PDM2 |
| Serine/arginine-rich splicing factor 2 | Q62093 |
| Serine/arginine-rich splicing factor 4 | Q8VE97 |
| Serine/threonine-protein kinase BRSK1 | Q5RJI5 |
| Serine/threonine-protein kinase BRSK2 | Q69Z98 |
| Serine/threonine-protein kinase DCLK1 | Q9JLM8 |
| Serine/threonine-protein kinase DCLK2 | Q6PGN3 |
| Serine/threonine-protein kinase MARK1 | Q8VHJ5 |
| Serine/threonine-protein kinase MARK2 | Q05512 |
| Serine/threonine-protein kinase MRCK alpha | Q3UU96 |
| Serine/threonine-protein kinase mTOR | Q9JLN9 |
| Serine/threonine-protein kinase OSR1 | Q6P9R2 |
| Serine/threonine-protein kinase PAK 1 | O88643 |
| Serine/threonine-protein kinase PAK 2 | Q8CIN4 |
| Serine/threonine-protein kinase PAK 3 | Q61036 |
| Serine/threonine-protein kinase TAO1 | Q5F2E8 |
| Serine/threonine-protein kinase TAO2 | Q6ZQ29 |
| Serine/threonine-protein kinase WNK1 | P83741 |
| Serine/threonine-protein kinase WNK2 | Q3UH66 |
| Serine/threonine-protein kinase WNK3 | Q80XP9 |
| Serine/threonine-protein phosphatase 2A 55 kDa r | Q6P1F6 |
| Serine/threonine-protein phosphatase 2A 56 kDa r | Q6PD03 |
| Serine/threonine-protein phosphatase 2A 56 kDa r | Q61151 |
| Serine/threonine-protein phosphatase 2A 56 kDa r | Q60996 |
| Serine/threonine-protein phosphatase 2A 65 kDa r | Q76MZ3 |
| Serine/threonine-protein phosphatase 2A activator | P58389 |
| Serine/threonine-protein phosphatase 2A catalytic | P63330 |
| Serine/threonine-protein phosphatase 2B catalytic | P63328 |
| Serine/threonine-protein phosphatase 2B catalytic | P48453 |
| Serine/threonine-protein phosphatase 5 | Q60676 |
| Serine/threonine-protein phosphatase 6 regulatory | Q922D4 |
| Serine/threonine-protein phosphatase PGAM5, mit | Q8BX10 |
| Serine-threonine kinase receptor-associated protei | Q9Z1Z2 |
| Serine--tRNA ligase, cytoplasmic | P26638 |
| Serotransferrin | Q921I1 |
| Serpin B6 | Q60854 |
| Serrate RNA effector molecule homolog | Q99MR6 |
| Serum albumin | P07724 |
| SH3 and multiple ankyrin repeat domains protein 1 | D3YZU1 |
| SH3 and multiple ankyrin repeat domains protein 2 | Q80Z38 |
| SH3 and multiple ankyrin repeat domains protein 3 | Q4ACU6 |
| SH3 domain-binding glutamic acid-rich-like protein | Q9JJU8 |
| SH3 domain-containing kinase-binding protein 1 | Q8R550 |
| SH3-containing GRB2-like protein 3-interacting pro | Q8VD37 |
| Shootin-1 | Q8K2Q9 |
| Short/branched chain specific acyl-CoA dehydroge | Q9DBL1 |
| Sickle tail protein | A2AQ25 |
| Sideroflexin-3 | Q91V61 |
| Signal-induced proliferation-associated 1-like prote | Q8C0T5 |
| Single-stranded DNA-binding protein, mitochondria | Q9CYR0 |
| SLIT-ROBO Rho GTPase-activating protein 2 | Q91Z67 |
| SLIT-ROBO Rho GTPase-activating protein 3 | Q812A2 |
| Small glutamine-rich tetratricopeptide repeat-conta | Q8BJU0 |
| Small VCP/p97-interacting protein | Q3UZP4 |
| S-methyl-5'-thioadenosine phosphorylase | Q9CQ65 |
| Sn1-specific diacylglycerol lipase alpha | Q6WQJ1 |
| Sodium- and chloride-dependent GABA transporte | P31650 |
| Sodium/calcium exchanger 1 | P70414 |
| Sodium/potassium-transporting ATPase subunit al | Q8VDN2 |
| Sodium/potassium-transporting ATPase subunit al | Q6PIE5 |
| Sodium/potassium-transporting ATPase subunit al | Q6PIC6 |
| Sodium/potassium-transporting ATPase subunit be | P14094 |
| Sodium/potassium-transporting ATPase subunit be | P14231 |
| Sodium-dependent neutral amino acid transporter | Q8BJI1 |
| Sodium-driven chloride bicarbonate exchanger | Q5DTL9 |
| Solute carrier family 12 member 2 | P55012 |
| Solute carrier family 12 member 5 | Q91V14 |
| Sorbin and SH3 domain-containing protein 1 | Q62417 |
| Sorbitol dehydrogenase | Q64442 |
| Sortilin | Q6PHU5 |
| Sorting and assembly machinery component 50 ho | Q8BGH2 |
| Sorting nexin-1 | Q9WV80 |
| Sorting nexin-16 | Q8C080 |
| Sorting nexin-2 | Q9CWK8 |
| Sorting nexin-27 | Q3UHD6 |
| Sorting nexin-4 | Q91YJ2 |
| Sorting nexin-5 | Q9D8U8 |
| Sorting nexin-6 | Q6P8X1 |
| SPARC-like protein 1 | P70663 |
| Spectrin alpha chain, brain | P16546 |
| Spectrin beta chain, brain 1 | Q62261 |
| Spectrin beta chain, erythrocyte | P15508 |
| Spermine synthase | P97355 |
| S-phase kinase-associated protein 1 | Q9WTX5 |
| Spliceosome RNA helicase Ddx39b | Q9Z1N5 |
| Splicing factor 1 | Q64213 |
| Splicing factor, proline- and glutamine-rich | Q8VIJ6 |
| Vascular cell adhesion protein 1 | P29533 |
| Vesicle transport through interaction with t-SNARE | O88384 |
| Vesicle-associated membrane protein 7 | P70280 |
| Vesicular glutamate transporter 1 | Q3TXX4 |
| Vesicular inhibitory amino acid transporter | O35633 |
| Vesicular integral-membrane protein VIP36 | Q9DBH5 |
| VIP36-like protein | P59481 |
| Voltage-dependent L-type calcium channel subunit | P54285 |
| Voltage-dependent L-type calcium channel subunit | Q8R0S4 |
| Voltage-dependent P/Q-type calcium channel subu | P97445 |
| von Willebrand factor A domain-containing protein | Q99KC8 |
| VPS10 domain-containing receptor SorCS2 | Q9EPR5 |
| VPS10 domain-containing receptor SorCS3 | Q8VI51 |
| V-type proton ATPase subunit G 1 | Q9CR51 |
| V-type proton ATPase subunit S1 | Q9R1Q9 |
| WASH complex subunit 7 | Q3UMB9 |
| WASH complex subunit strumpellin | Q8C2E7 |
| WD and tetratricopeptide repeats protein 1 | Q80ZK9 |
| WD repeat domain phosphoinositide-interacting pr | Q80W47 |
| WD repeat domain phosphoinositide-interacting pr | Q9CR39 |
| WD repeat domain phosphoinositide-interacting pr | Q91VM3 |
| WD repeat-containing protein 11 | Q8K1X1 |
| WD repeat-containing protein 18 | Q4VBE8 |
| WD repeat-containing protein 26 | Q8C6G8 |
| WD repeat-containing protein 34 | Q5U4F6 |
| WD repeat-containing protein 48 | Q8BH57 |
| WD repeat-containing protein 59 | Q8C0M0 |
| WD repeat-containing protein 70 | Q3TWF6 |
| WD repeat-containing protein 82 | Q8BFQ4 |
| WD repeat-containing protein 92 | Q8BGF3 |
| WD repeat-containing protein mio | Q8VE19 |
| WW domain-binding protein 11 | Q923D5 |
| WW domain-binding protein 2 | P97765 |
| XK-related protein 4 | Q5GH67 |
| X-linked retinitis pigmentosa GTPase regulator-inte | Q9EPQ2 |
| YTH domain family protein 3 | Q8BYK6 |
| Zinc finger C2HC domain-containing protein 1A | Q8BJH1 |
| Zinc finger FYVE domain-containing protein 1 | Q810J8 |
| Zinc finger RNA-binding protein | O88532 |
| Zinc finger ZZ-type and EF-hand domain-containin | Q5SSH7 |
| Zinc phosphodiesterase ELAC protein 2 | Q80Y81 |
| Zinc transporter SLC39A7 | Q31125 |
| Zinc-binding alcohol dehydrogenase domain-conta | Q8BGC4 |

| | |
|---|---|
| Squamous cell carcinoma antigen recognized by T | Q9JLI8 |
| SRC kinase signaling inhibitor 1 | Q9QWI6 |
| Src substrate cortactin | Q60598 |
| SRSF protein kinase 2 | O54781 |
| Staphylococcal nuclease domain-containing protei | Q78PY7 |
| Stathmin | P54227 |
| Stathmin-2 | P55821 |
| Stathmin-3 | O70166 |
| STE20/SPS1-related proline-alanine-rich protein ki | Q9Z1W9 |
| STE20-like serine/threonine-protein kinase | O54988 |
| Sterile alpha and TIR motif-containing protein 1 | Q6PDS3 |
| STIP1 homology and U box-containing protein 1 | Q9WUD1 |
| Stomatin-like protein 2 | Q99JB2 |
| Stress-70 protein, mitochondrial | P38647 |
| Stress-induced-phosphoprotein 1 | Q60864 |
| Striated muscle-specific serine/threonine-protein k | Q62407 |
| Striatin | O55106 |
| Striatin-3 | Q9ERG2 |
| Striatin-4 | P58404 |
| Stromal interaction molecule 2 | P83093 |
| Succinate dehydrogenase [ubiquinone] flavoprotei | Q8K2B3 |
| Succinate dehydrogenase [ubiquinone] iron-sulfur | Q9CQA3 |
| Succinate-semialdehyde dehydrogenase, mitochor | Q8BWF0 |
| Succinyl-CoA ligase [ADP-forming] subunit beta, n | Q9Z2I9 |
| Succinyl-CoA ligase [GDP-forming] subunit beta, r | Q9Z2I8 |
| Succinyl-CoA:3-ketoacid-coenzyme A transferase | Q9D0K2 |
| Sulfated glycoprotein 1 | Q61207 |
| Sulfotransferase 4A1 | P63046 |
| SUMO-activating enzyme subunit 1 | Q9R1T2 |
| SUMO-activating enzyme subunit 2 | Q9Z1F9 |
| Superoxide dismutase [Cu-Zn] | P08228 |
| Suppressor of G2 allele of SKP1 homolog | Q9CX34 |
| SWI/SNF complex subunit SMARCC2 | Q6PDG5 |
| Synapsin-1 | O88935 |
| Synapsin-2 | Q64332 |
| Synaptic vesicle glycoprotein 2A | Q9JIS5 |
| Synaptic vesicle glycoprotein 2B | Q8BG39 |
| Synaptic vesicle membrane protein VAT-1 homolo | Q80TB8 |
| Synaptobrevin homolog YKT6 | Q9CQW1 |
| Synaptojanin-1 | Q8CHC4 |
| Synaptopodin | Q8CC35 |
| Synaptosomal-associated protein 25 | P60879 |
| Synaptosomal-associated protein 29 | Q9ERB0 |
| Synaptosomal-associated protein 47 | Q8R570 |
| Synaptotagmin-1 | P46096 |
| Synaptotagmin-17 | Q920M7 |
| Synaptotagmin-7 | Q9R0N7 |
| Synergin gamma | Q5SV85 |
| Syntaphilin | Q80U23 |
| Syntaxin-12 | Q9ER00 |
| Syntaxin-1A | O35526 |
| Syntaxin-1B | P61264 |
| Syntaxin-7 | O70439 |
| Syntaxin-8 | O88983 |
| Syntaxin-binding protein 1 | O08599 |
| Syntaxin-binding protein 5 | Q8K400 |
| Syntaxin-binding protein 5-like | Q5DQR4 |
| Syntaxin-binding protein 6 | Q8R3T5 |
| Talin-2 | Q71LX4 |
| TBC1 domain family member 24 | Q3UUG6 |
| T-complex protein 1 subunit alpha | P11983 |
| T-complex protein 1 subunit beta | P80314 |
| T-complex protein 1 subunit delta | P80315 |
| T-complex protein 1 subunit epsilon | P80316 |
| T-complex protein 1 subunit eta | P80313 |
| T-complex protein 1 subunit gamma | P80318 |
| T-complex protein 1 subunit theta | P42932 |
| T-complex protein 1 subunit zeta | P80317 |
| Tectonin beta-propeller repeat-containing protein 1 | Q80VP0 |
| Tenascin | Q80YX1 |
| Tenascin-R | Q8BYI9 |
| Teneurin-4 | Q3UHK6 |
| Testis-expressed sequence 2 protein | Q6ZPJ0 |
| Thiamine-triphosphatase | Q8JZL3 |
| Thimet oligopeptidase | Q8C1A5 |
| Thiomorpholine-carboxylate dehydrogenase | O54983 |
| Thioredoxin | P10639 |
| Thioredoxin reductase 1, cytoplasmic | Q9JMH6 |
| Thioredoxin-dependent peroxide reductase, mitoch | P20108 |
| Thioredoxin-like protein 1 | Q8CDN6 |
| Thioredoxin-related transmembrane protein 4 | Q8C0L0 |
| Thiosulfate sulfurtransferase | P52196 |
| Threonine synthase-like 1 | Q8BH55 |
| Threonine--tRNA ligase, cytoplasmic | Q9D0R2 |
| Thrombospondin type-1 domain-containing protein | Q69ZU6 |
| THUMP domain-containing protein 1 | Q99J36 |
| Thy-1 membrane glycoprotein | P01831 |
| Thymosin beta-10 | Q6ZWY8 |
| TIP41-like protein | Q8BH58 |
| Titin | A2ASS6 |
| T-lymphoma invasion and metastasis-inducing pro | Q60610 |
| Toll-interacting protein | Q9QZ06 |
| TOM1-like protein 2 | Q5SRX1 |
| Traf2 and NCK-interacting protein kinase | P83510 |

| | |
|---|---|
| Trafficking protein particle complex subunit 12 | Q8K2L8 |
| Trafficking protein particle complex subunit 5 | Q9CQA1 |
| Trafficking protein particle complex subunit 9 | Q3U0M1 |
| Trans-2-enoyl-CoA reductase, mitochondrial | Q9DCS3 |
| Transaldolase | Q93092 |
| Transcription elongation factor A protein 1 | P10711 |
| Transcription elongation factor A protein-like 5 | Q8CCT4 |
| Transcription elongation factor B polypeptide 2 | P62869 |
| Transcription elongation regulator 1 | Q8CGF7 |
| Transcription factor A, mitochondrial | P40630 |
| Transcription intermediary factor 1-beta | Q62318 |
| Transcriptional activator protein Pur-alpha | P42669 |
| Transcriptional activator protein Pur-beta | O35295 |
| Transferrin receptor protein 1 | Q62351 |
| Transforming acidic coiled-coil-containing protein 1 | Q6Y685 |
| Transforming acidic coiled-coil-containing protein 2 | Q9JJG0 |
| Transforming protein RhoA | Q9QUI0 |
| Transgelin | P37804 |
| Transgelin-2 | Q9WVA4 |
| Transgelin-3 | Q9R1Q8 |
| Transitional endoplasmic reticulum ATPase | Q01853 |
| Transketolase | P40142 |
| Translationally-controlled tumor protein | P63028 |
| Translin-associated protein X | Q9QZE7 |
| Transmembrane emp24 domain-containing protein | Q99KF1 |
| Transthyretin | P07309 |
| Trifunctional enzyme subunit alpha, mitochondrial | Q8BMS1 |
| Trifunctional enzyme subunit beta, mitochondrial | Q99JY0 |
| Trifunctional purine biosynthetic protein adenosine | Q64737 |
| Triosephosphate isomerase | P17751 |
| Tripeptidyl-peptidase 2 | Q64514 |
| Triple functional domain protein | Q0KL02 |
| tRNA-splicing ligase RtcB homolog | Q99LF4 |
| Tropomodulin-2 | Q9JKK7 |
| Tropomyosin alpha-1 chain | P58771 |
| Tropomyosin alpha-3 chain | P21107 |
| Tropomyosin alpha-4 chain | Q6IRU2 |
| Tryptophan--tRNA ligase, cytoplasmic | P32921 |
| TSC22 domain family protein 1 | P62500 |
| TSC22 domain family protein 3 | Q9Z2S7 |
| Tubby protein | P50586 |
| Tuberin | Q61037 |
| Tubulin alpha-4A chain | P68368 |
| Tubulin alpha-8 chain | Q9JJZ2 |
| Tubulin beta-2B chain | Q9CWF2 |
| Tubulin beta-3 chain | Q9ERD7 |
| Tubulin beta-4A chain | Q9D6F9 |
| Tubulin polymerization-promoting protein | Q7TQD2 |
| Tubulin polymerization-promoting protein family m | Q9CRB6 |
| Tubulin-folding cofactor B | Q9D1E6 |
| Tubulin-specific chaperone A | P48428 |
| Tubulin-specific chaperone C | Q8VCN9 |
| Tubulin-specific chaperone D | Q8BYA0 |
| Tumor protein D54 | Q9CYZ2 |
| Twinfilin-1 | Q91YR1 |
| Twinfilin-2 | Q9Z0P5 |
| Type I inositol 3,4-bisphosphate 4-phosphatase | Q9EPW0 |
| Tyrosine-protein kinase Fyn | P39688 |
| Tyrosine-protein phosphatase non-receptor type 1 | P35235 |
| Tyrosine-protein phosphatase non-receptor type 2 | Q6PB44 |
| Tyrosine--tRNA ligase, cytoplasmic | Q91WQ3 |
| U1 small nuclear ribonucleoprotein 70 kDa | Q62376 |
| U6 snRNA-associated Sm-like protein LSm2 | O35900 |
| Ubiquinone biosynthesis protein COQ9, mitochond | Q8K1Z0 |
| Ubiquitin carboxyl-terminal hydrolase 10 | P52479 |
| Ubiquitin carboxyl-terminal hydrolase 14 | Q9JMA1 |
| Ubiquitin carboxyl-terminal hydrolase 15 | Q8R5H1 |
| Ubiquitin carboxyl-terminal hydrolase 5 | P56399 |
| Ubiquitin carboxyl-terminal hydrolase 7 | Q6A4J8 |
| Ubiquitin carboxyl-terminal hydrolase CYLD | Q80TQ2 |
| Ubiquitin carboxyl-terminal hydrolase isozyme L1 | Q9R0P9 |
| Ubiquitin carboxyl-terminal hydrolase isozyme L3 | Q9JKB1 |
| Ubiquitin fusion degradation protein 1 homolog | P70362 |
| Ubiquitin thioesterase OTUB1 | Q7TQI3 |
| Ubiquitin-associated protein 2-like | Q80X50 |
| Ubiquitin-conjugating enzyme E2 K | P61087 |
| Ubiquitin-conjugating enzyme E2 L3 | P68037 |
| Ubiquitin-conjugating enzyme E2 N | P61089 |
| Ubiquitin-conjugating enzyme E2 O | Q6ZPJ3 |
| Ubiquitin-conjugating enzyme E2 variant 1 | Q9CZY3 |
| Ubiquitin-like modifier-activating enzyme 1 | Q02053 |
| Ubiquitin-like modifier-activating enzyme 6 | Q8C7R4 |
| UBX domain-containing protein 1 | Q922Y1 |
| UBX domain-containing protein 6 | Q99PL6 |
| UBX domain-containing protein 7 | Q6P5G6 |
| UDP-N-acetylglucosamine--peptide N-acetylglucos | Q8CGY8 |
| UHRF1-binding protein 1-like | A2RSJ4 |
| UMP-CMP kinase | Q9DBP5 |
| UMP-CMP kinase 2, mitochondrial | Q3U5Q7 |
| Uncharacterized protein C1orf198 homolog | Q8C3W1 |
| Uncharacterized protein C1orf96 homolog | Q8QZT2 |
| Uncharacterized protein C6orf174 homolog | Q6NZL0 |
| Uncharacterized protein KIAA0513 | Q8R0A7 |
| Uncharacterized protein KIAA0564 homolog | Q8CC88 |

| | |
|---|---|
| Uncharacterized protein KIAA1107 | Q80TK0 |
| Uncharacterized protein KIAA1671 | Q8BRV5 |
| Unconventional myosin-Id | Q5SYD0 |
| Unconventional myosin-Va | Q99104 |
| Unconventional myosin-VI | Q64331 |
| Unconventional myosin-XVIIIa | Q9JMH9 |
| UPF0364 protein C6orf211 homolog | A6H630 |
| UPF0366 protein C11orf67 homolog | Q8R0P4 |
| UPF0505 protein C16orf62 homolog | Q8BWQ6 |
| UPF0533 protein C5orf44 homolog | Q3TIR1 |
| UPF0552 protein C15orf38 homolog | Q9D0A3 |
| UPF0568 protein C14orf166 homolog | Q9CQE8 |
| UPF0587 protein C1orf123 homolog | Q8BHG2 |
| UPF0606 protein KIAA1549 | Q68FD9 |
| UTP--glucose-1-phosphate uridylyltransferase | Q91ZJ5 |
| UV excision repair protein RAD23 homolog B | P54728 |
| Vacuolar protein sorting-associated protein 11 hom | Q91W86 |
| Vacuolar protein sorting-associated protein 13C | Q8BX70 |
| Vacuolar protein sorting-associated protein 26A | P40336 |
| Vacuolar protein sorting-associated protein 26B | Q8C0E2 |
| Vacuolar protein sorting-associated protein 29 | Q9QZ88 |
| Vacuolar protein sorting-associated protein 33B | P59016 |
| Vacuolar protein sorting-associated protein 35 | Q9EQH3 |
| Vacuolar protein sorting-associated protein 45 | P97390 |
| Vacuolar protein sorting-associated protein 53 hom | Q8CCB4 |
| Valacyclovir hydrolase | Q8R164 |
| Valine--tRNA ligase | Q9Z1Q9 |
| Versican core protein | Q62059 |
| Very long-chain specific acyl-CoA dehydrogenase, | P50544 |
| Vesicle-associated membrane protein 1 | Q62442 |
| Vesicle-associated membrane protein 2 | P63044 |
| Vesicle-associated membrane protein-associated | Q9WV55 |
| Vesicle-associated membrane protein-associated | Q9QY76 |
| Vesicle-fusing ATPase | P46460 |
| Vesicle-trafficking protein SEC22b | O08547 |
| Vigilin | Q8VDJ3 |
| Vimentin | P20152 |
| Vinculin | Q64727 |
| Visinin-like protein 1 | P62761 |
| Voltage-dependent anion-selective channel proteir | Q60932 |
| Voltage-dependent anion-selective channel proteir | Q60930 |
| Voltage-dependent anion-selective channel proteir | Q60931 |
| Voltage-dependent calcium channel gamma-8 sub | Q8VHW2 |
| Voltage-dependent calcium channel subunit alpha- | O08532 |
| Voltage-dependent calcium channel subunit alpha- | Q6PHS9 |
| Voltage-dependent calcium channel subunit alpha- | Q9Z1L5 |
| Voltage-dependent L-type calcium channel subunit | Q8R3Z5 |
| Voltage-dependent N-type calcium channel subuni | O55017 |
| Voltage-dependent R-type calcium channel subuni | Q61290 |
| Voltage-gated potassium channel subunit beta-2 | P62482 |
| V-type proton ATPase 116 kDa subunit a isoform 1 | Q9Z1G4 |
| V-type proton ATPase catalytic subunit A | P50516 |
| V-type proton ATPase subunit B, brain isoform | P62814 |
| V-type proton ATPase subunit C 1 | Q9Z1G3 |
| V-type proton ATPase subunit D | P57746 |
| V-type proton ATPase subunit d 1 | P51863 |
| V-type proton ATPase subunit E 1 | P50518 |
| V-type proton ATPase subunit F | Q9D1K2 |
| V-type proton ATPase subunit G 2 | Q9WTT4 |
| V-type proton ATPase subunit H | Q8BVE3 |
| WAS/WASL-interacting protein family member 3 | P0C7L0 |
| WASH complex subunit FAM21 | Q6PGL7 |
| WD repeat and FYVE domain-containing protein 3 | Q6VNB8 |
| WD repeat-containing protein 1 | O88342 |
| WD repeat-containing protein 13 | Q91V09 |
| WD repeat-containing protein 37 | Q8CBE3 |
| WD repeat-containing protein 44 | Q6NVE8 |
| WD repeat-containing protein 47 | Q8CGF6 |
| WD repeat-containing protein 54 | Q9R0D8 |
| WD repeat-containing protein 7 | Q920I9 |
| Wiskott-Aldrich syndrome protein family member 1 | Q8R5H6 |
| Wiskott-Aldrich syndrome protein family member 3 | Q8VHI6 |
| Wolframin | P56695 |
| Xaa-Pro aminopeptidase 1 | Q6P1B1 |
| Xaa-Pro dipeptidase | Q11136 |
| Zinc finger protein ZPR1 | Q62384 |
| ZW10 interactor | Q9CQU5 |

### Table S6. Differentially expressed proteins at P21

ND, not detected; [a]detected in the same comparison in adults; [b]differential expression between sexes in control and ELS reared hippocampus; [c]altered in males and females in response to ELS

| Male ELS vs Male control - 8 proteins | | | | | |
|---|---|---|---|---|---|
| **Protein Name** | **Uniprot ID** | **Experiment 1** | **Experiment 2** | **Experiment 3** | **Average** |
| **Upregulated in stress - 6 proteins** | | | | | |
| Arginine and glutamate-rich protein 1 | Q3UL36 | ND | 1.3888 | 1.2167 | **1.3027** |
| Heterogeneous nuclear ribonucleoprotein L | Q8R081 | 1.2336 | 1.1948 | 1.3851 | **1.2712** |
| Histone H3.1 | P68433 | 1.2789 | 1.2516 | ND | **1.2653** |
| Histone H4 | P62806 | 1.1793 | 1.3003 | 1.5659 | **1.3485** |
| Parvalbumin alpha | P32848 | 1.4378 | 0.9099 | 1.2825 | **1.2101** |
| Pinin | O35691 | ND | 1.2770 | 1.2728 | **1.2749** |
| **Downregulated in stress - 2 proteins** | | | | | |
| Activated RNA polymerase II transcriptional coacti | P11031 | 0.7396 | 0.7324 | 0.6926 | **0.7215** |
| [c]Cytochrome c, somatic | P62897 | 0.6515 | 0.9297 | 0.5913 | **0.7242** |

| Female ELS vs Female control - 60 proteins | | | | | |
|---|---|---|---|---|---|
| **Protein Name** | **Uniprot ID** | **Experiment 1** | **Experiment 2** | **Experiment 3** | **Average** |
| **Upregulated in stress - 56 proteins** | | | | | |
| Acetyl-CoA acetyltransferase, cytosolic | Q8CAY6 | 1.0913 | 1.3574 | 1.2370 | **1.2286** |
| Acylphosphatase-1 | P56376 | 1.3186 | 1.1784 | 1.2613 | **1.2528** |
| [a]Alpha-2-macroglobulin receptor-associated protein | P55302 | 1.2614 | 1.2023 | 1.2833 | **1.2490** |
| Aspartate aminotransferase, cytoplasmic | P05201 | 1.1761 | 1.4377 | 1.2107 | **1.2749** |
| [a]Aspartyl aminopeptidase | Q9Z2W0 | 1.2684 | 1.2647 | 1.1908 | **1.2413** |
| BAG family molecular chaperone regulator 4 | Q8CI61 | 1.2763 | 1.2242 | 1.2150 | **1.2385** |
| Beta-synuclein | Q91ZZ3 | 1.3585 | 1.2031 | 1.1234 | **1.2283** |
| [a]Cellular nucleic acid-binding protein | P53996 | ND | 1.2499 | 1.2175 | **1.2337** |
| Charged multivesicular body protein 1a | Q921W0 | 1.3463 | 1.2318 | ND | **1.2891** |
| [a]Charged multivesicular body protein 4b | Q9D8B3 | 1.4191 | 1.2284 | 1.1320 | **1.2599** |
| Coiled-coil domain-containing protein 58 | Q8R3Q6 | ND | 1.2895 | 1.2754 | **1.2824** |
| [a]Cysteine and glycine-rich protein 1 | P97315 | 1.4499 | 1.2770 | 1.1258 | **1.2842** |
| [c]Cytochrome c, somatic | P62897 | 1.3140 | 1.3994 | 2.9037 | **1.8724** |
| Diablo homolog, mitochondrial | Q9JIQ3 | ND | 1.2048 | 1.2243 | **1.2145** |
| ES1 protein homolog, mitochondrial | Q9D172 | 1.5637 | 1.3687 | 1.2099 | **1.3808** |
| Gamma-synuclein | Q9Z0F7 | 1.1810 | 1.2378 | 1.2431 | **1.2206** |
| Gamma-tubulin complex component 4 | Q9D4F8 | 1.2825 | ND | 1.2613 | **1.2719** |
| Haloacid dehalogenase-like hydrolase domain-cont | Q3UGR5 | ND | 1.2149 | 1.3898 | **1.3023** |
| Hepatoma-derived growth factor | P51859 | 1.4083 | 1.3166 | 1.1064 | **1.2771** |
| Hepatoma-derived growth factor-related protein 3 | Q9JMG7 | 1.2167 | 1.2656 | 1.2878 | **1.2567** |
| Hydroxyacylglutathione hydrolase, mitochondrial | Q99KB8 | 1.2816 | 1.2284 | 1.1273 | **1.2125** |
| Inorganic pyrophosphatase 2, mitochondrial | Q91VM9 | 1.3113 | 1.1414 | 1.2457 | **1.2328** |
| Inositol monophosphatase 1 | O55023 | 1.2614 | 1.1319 | 1.2622 | **1.2185** |
| Lymphocyte antigen 6H | Q9WUC3 | ND | 1.5050 | 1.3556 | **1.4303** |
| Magnesium-dependent phosphatase 1 | Q9D967 | 1.1439 | 1.2967 | 1.2353 | **1.2253** |
| [a]Malate dehydrogenase, mitochondrial | P08249 | 1.5105 | 1.1438 | 1.2192 | **1.2911** |
| Metallothionein-3 | P28184 | ND | 1.2430 | 1.3258 | **1.2844** |
| [a]Mitochondrial import inner membrane translocase : | P62073 | 1.3361 | 1.0739 | 1.2482 | **1.2194** |
| Mitochondrial import inner membrane translocase s | Q9WVA2 | ND | 1.2048 | 1.2066 | **1.2057** |
| [a]Neuronal cell adhesion molecule | Q810U4 | 1.4661 | 1.2056 | 1.1360 | **1.2692** |
| Neuronal pentraxin-2 | O70340 | 1.3928 | 1.1179 | 1.2413 | **1.2507** |
| NHS-like protein 1 | Q8CAF4 | 1.2311 | ND | 1.3378 | **1.2845** |
| [a]Nitrilase homolog 1 | Q8VDK1 | 1.2243 | 1.3303 | 1.2217 | **1.2588** |
| [a]Parathymosin | Q9D0J8 | 1.3784 | 1.3285 | 1.1510 | **1.2860** |
| Peptidyl-prolyl cis-trans isomerase F, mitochondrial | Q99KR7 | 1.4794 | 1.1613 | 1.2396 | **1.2935** |
| [a]Prefoldin subunit 1 | Q9CWM4 | 1.3068 | 1.2123 | 1.1157 | **1.2116** |
| Prefoldin subunit 2 | O70591 | 1.2423 | 1.1581 | 1.2082 | **1.2029** |
| Prefoldin subunit 5 | Q9WU28 | 1.0770 | 1.2832 | 1.2561 | **1.2054** |
| Proline synthase co-transcribed bacterial homolog p | Q9Z2Y8 | 1.4692 | 1.4268 | 1.1041 | **1.3334** |
| Protein HEXIM1 | Q8R409 | ND | 1.2098 | 1.3556 | **1.2827** |
| [a]Rho GDP-dissociation inhibitor 1 | Q99PT1 | 1.2702 | 1.2293 | 1.1558 | **1.2184** |
| Ribonuclease UK114 | P52760 | 1.2666 | 1.2242 | 1.1630 | **1.2179** |
| [a]Ribosome-recycling factor, mitochondrial | Q9D6S7 | 1.6133 | 1.0791 | 1.3481 | **1.3468** |
| Secretogranin-2 | Q03517 | 1.7195 | 1.1398 | 1.2192 | **1.3595** |
| Small ubiquitin-related modifier 2 | P61957 | 1.2596 | 1.4357 | 1.1817 | **1.2924** |
| SPARC-like protein 1 | P70663 | 1.3491 | 1.2404 | 1.1423 | **1.2439** |
| Spermatogenesis-associated protein 7 homolog | Q80VP2 | 2.2942 | 1.3030 | ND | **1.7986** |
| Stathmin | P54227 | 1.3159 | 1.2895 | 1.1622 | **1.2559** |
| Stathmin-2 | P55821 | 1.4359 | 1.2656 | 1.1273 | **1.2763** |
| Thioredoxin reductase 2, mitochondrial | Q9JLT4 | 1.4064 | 1.2815 | ND | **1.3439** |
| [a]Transcription elongation factor A protein 1 | P10711 | ND | 1.2242 | 1.2336 | **1.2289** |
| Transcription elongation factor A protein-like 3 | Q8R0A5 | 1.2125 | 1.2595 | 1.1542 | **1.2087** |
| Transketolase | P40142 | 1.2058 | 1.6723 | 1.2569 | **1.3783** |
| Triosephosphate isomerase | P17751 | 1.2816 | 1.3517 | 1.1312 | **1.2549** |
| [a]U6 snRNA-associated Sm-like protein LSm2 | O35900 | ND | 1.2647 | 1.3575 | **1.3111** |
| Uncharacterized protein C1orf198 homolog | Q8C3W1 | 1.3068 | 1.2395 | 1.0844 | **1.2102** |
| **Downregulated in stress - 4 proteins** | | | | | |
| 60S ribosomal protein L38 | Q9JJI8 | 0.3560 | 0.7747 | 0.8875 | **0.6728** |
| Carnitine O-palmitoyltransferase 1, brain isoform | Q8BGD5 | 0.6620 | 0.7949 | 0.9375 | **0.7981** |
| [a]Diacylglycerol kinase beta | Q6NS52 | 0.6281 | 0.7758 | 0.8473 | **0.7504** |
| Phosphatidate cytidylyltransferase 2 | Q99L43 | 0.5905 | 0.6987 | 0.9408 | **0.7433** |

| Male control vs Female control - 4 proteins | | | | | |
|---|---|---|---|---|---|
| **Protein Name** | **Uniprot ID** | **Experiment 1** | **Experiment 2** | **Experiment 3** | **Average** |
| **Lower in females - 2 proteins** | | | | | |
| [b]Cytochrome c, somatic | P62897 | 2.5377 | 0.9219 | 2.2673 | **1.9090** |
| Sodium/potassium-transporting ATPase subunit b | P14231 | 1.3943 | 0.8992 | 1.3928 | **1.2288** |
| **Lower in males - 2 proteins** | | | | | |
| Ferritin heavy chain | P09528 | 0.6683 | 0.8548 | 0.7955 | **0.7729** |
| Parvalbumin alpha | P32848 | 0.7991 | 0.9664 | 0.6134 | **0.7930** |

| Male ELS vs Female ELS - 91 proteins | | | | | |
|---|---|---|---|---|---|
| **Protein Name** | **Uniprot ID** | **Experiment 1** | **Experiment 2** | **Experiment 3** | **Average** |
| **Lower in females - 32 proteins** | | | | | |
| 5'-AMP-activated protein kinase subunit gamma-1 | O54950 | 1.2879 | 1.2158 | ND | **1.2518** |
| 60S ribosomal protein L38 | Q9JJI8 | 2.7397 | 1.3518 | 0.8433 | **1.6449** |
| ADP/ATP translocase 1 | P48962 | 1.7219 | 1.3249 | 1.0318 | **1.3595** |
| ADP/ATP translocase 2 | P51881 | 1.5519 | 1.2413 | 1.0455 | **1.2796** |
| ADP/ATP translocase 4 | Q3V132 | 1.4340 | 1.3149 | 1.0183 | **1.2557** |
| Amine oxidase [flavin-containing] A | Q64133 | 1.4211 | 1.2024 | 1.0616 | **1.2283** |
| ATP synthase subunit b, mitochondrial | Q9CQQ7 | 1.7351 | 1.2082 | 1.0557 | **1.3330** |
| ATPase family AAA domain-containing protein 3 | Q925I1 | 1.6982 | 1.2630 | 1.1174 | **1.3596** |
| BTB/POZ domain-containing protein KCTD8 | Q50H33 | 1.7605 | 1.2388 | ND | **1.4996** |
| Carnitine O-palmitoyltransferase 1, brain isoform | Q8BGD5 | 1.2311 | 1.2869 | 1.1960 | **1.2380** |
| Cholesterol 24-hydroxylase | Q9WVK8 | 1.5508 | 1.2217 | 1.0050 | **1.2592** |
| Cytochrome c oxidase subunit 2 | P00405 | 1.3031 | 1.1916 | 1.2118 | **1.2355** |
| D-dopachrome decarboxylase | O35215 | 1.5497 | 1.4731 | 0.8877 | **1.3035** |
| Disabled homolog 2-interacting protein | Q3UHC7 | 1.4123 | 1.2032 | 1.0155 | **1.2103** |
| Glutamate [NMDA] receptor subunit zeta-1 | P35438 | 1.6369 | 1.2277 | 1.1528 | **1.3391** |
| Glutamate receptor 1 | P23818 | 1.1704 | 1.2192 | 1.2110 | **1.2002** |
| Histone H2B type 1-P | Q8CGP2 | 1.2440 | 1.0020 | 1.7008 | **1.3156** |
| Histone H4 | P62806 | 1.6644 | 1.3831 | 1.5607 | **1.5361** |
| Kalirin | A2CG49 | 1.3122 | 1.2639 | 1.0719 | **1.2160** |
| Mitofusin-1 | Q811U4 | ND | 1.2387 | 1.2564 | **1.2476** |
| Myelin proteolipid protein | P60202 | 1.8188 | 1.2277 | 1.0226 | **1.3563** |
| Phosphatidate cytidylyltransferase 2 | Q99L43 | 2.0591 | 1.3388 | 1.1457 | **1.5145** |
| Phosphatidylinositol 4,5-bisphosphate 3-kinase cata | Q8BTI9 | 1.5476 | 1.2007 | ND | **1.3741** |
| Potassium voltage-gated channel subfamily A meml | Q61423 | 1.6911 | 1.2860 | ND | **1.4886** |
| Protein QIL1 | Q8R404 | ND | 1.2370 | 1.2245 | **1.2308** |
| Protein TANC2 | A2A690 | 1.4570 | 1.2032 | 1.0914 | **1.2505** |
| Saccharopine dehydrogenase-like oxidoreductase | Q8R127 | 1.4897 | 1.0259 | 1.2952 | **1.2703** |
| Sideroflexin-5 | Q925N0 | 1.6575 | 1.2482 | 1.0899 | **1.3319** |
| Solute carrier family 2, facilitated glucose transporte | P32037 | 1.3660 | 1.2099 | 1.1120 | **1.2293** |
| Synaptic vesicle glycoprotein 2B | Q8BG39 | 1.5790 | 1.2166 | 1.0727 | **1.2894** |
| Thioredoxin-related transmembrane protein 2 | Q9D710 | ND | 1.2869 | 1.2169 | **1.2519** |
| Trafficking protein particle complex subunit 12 | Q8K2L8 | 1.5422 | 1.3139 | 1.0204 | **1.2922** |
| **Lower in males - 59 proteins** | | | | | |
| 10 kDa heat shock protein, mitochondrial | Q64433 | 0.6736 | 0.7737 | 0.8581 | **0.7685** |
| 39S ribosomal protein L12, mitochondrial | Q9DB15 | 0.8462 | 0.7797 | 0.7352 | **0.7870** |
| [a]Acyl-CoA-binding protein | P31786 | 0.9760 | 0.7120 | 0.6347 | **0.7742** |
| Acylphosphatase-1 | P56376 | 0.7371 | 0.7335 | 0.7664 | **0.7457** |
| [a]Alpha-endosulfine | P60840 | 0.8415 | 0.7159 | 0.7201 | **0.7592** |
| Aspartate aminotransferase, cytoplasmic | P05201 | 0.9025 | 0.6272 | 0.7638 | **0.7645** |
| Aspartyl aminopeptidase | Q9Z2W0 | 0.7647 | 0.7189 | 0.7831 | **0.7556** |
| [a]ATPase inhibitor, mitochondrial | O35143 | 0.5133 | 0.7494 | 0.9735 | **0.7454** |
| BAG family molecular chaperone regulator 4 | Q8CI61 | 0.8901 | 0.7075 | 0.7151 | **0.7709** |
| Charged multivesicular body protein 1a | Q921W0 | 0.6849 | 0.7407 | ND | **0.7128** |
| Charged multivesicular body protein 4b | Q9D8B3 | 0.7051 | 0.7249 | 0.7352 | **0.7218** |
| Coiled-coil domain-containing protein 124 | Q9D8X2 | 0.5684 | 0.7994 | ND | **0.6839** |
| Coiled-coil domain-containing protein 58 | Q8R3Q6 | ND | 0.7552 | 0.7728 | **0.7640** |
| [b]Cytochrome c, somatic | P62897 | 1.0725 | 0.6058 | 0.5181 | **0.7321** |
| [a]Dematin | Q9WV69 | 0.6983 | 0.7999 | 0.9001 | **0.7995** |
| FAD synthase | Q8R123 | 0.6547 | ND | 0.7643 | **0.7095** |
| General transcription factor IIF subunit 1 | Q3THK3 | ND | 0.7484 | 0.7787 | **0.7636** |
| Haloacid dehalogenase-like hydrolase domain-conta | Q3UGR5 | ND | 0.6797 | 0.6994 | **0.6895** |
| Hepatoma-derived growth factor | P51859 | 0.6926 | 0.6887 | 0.8451 | **0.7421** |
| Hepatoma-derived growth factor-related protein 3 | Q9JMG7 | 0.7180 | 0.6675 | 0.7744 | **0.7200** |
| Integrin alpha-M | P05555 | 0.3558 | 0.7775 | 0.7868 | **0.6400** |
| Lymphocyte antigen 6H | Q9WUC3 | ND | 0.6293 | 0.7696 | **0.6995** |
| Macrophage migration inhibitory factor | P34884 | 0.742545636 | 0.7177 | 0.7826 | **0.7476** |
| Metallothionein-3 | P28184 | ND | 0.6806 | 0.6299 | **0.6553** |
| Mitochondrial import inner membrane translocase s | P62073 | 0.8000 | 0.7562 | 0.6980 | **0.7514** |
| Mitochondrial import inner membrane translocase s | Q9WVA2 | ND | 0.7149 | 0.7717 | **0.7433** |
| Mitochondrial import inner membrane translocase s | P62077 | 0.8895 | 0.6834 | 0.7206 | **0.7645** |
| Na(+)/H(+) exchange regulatory cofactor NHE-RF1 | P70441 | 0.6760 | 0.7547 | 0.8599 | **0.7635** |
| Neurogranin | P60761 | ND | 0.7179 | 0.6737 | **0.6958** |
| Nitrilase homolog 1 | Q8VDK1 | 0.8803 | 0.7179 | 0.7744 | **0.7909** |
| [a]Parathymosin | Q9D0J8 | 0.7245 | 0.5629 | 0.7691 | **0.6855** |
| Peptidyl-prolyl cis-trans isomerase NIMA-interacting | Q9QUR7 | 0.7056 | 0.7960 | 0.8510 | **0.7842** |
| Prefoldin subunit 1 | Q9CWM4 | 0.7485 | 0.6792 | 0.8197 | **0.7491** |
| Pre-rRNA-processing protein TSR2 homolog | Q8C8T8 | 0.8316 | 0.7100 | 0.7755 | **0.7724** |
| Programmed cell death protein 5 | P56812 | 0.7140 | 0.7300 | 0.7622 | **0.7354** |
| Proline synthase co-transcribed bacterial homolog p | Q9Z2Y8 | 0.7851 | 0.6466 | 0.9159 | **0.7825** |
| ProSAAS | Q9QXV0 | 0.7195 | 0.8027 | 0.7067 | **0.7430** |
| Protein CDV3 | Q4VAA2 | 0.6448 | 0.6983 | 0.7654 | **0.7028** |
| [a]Protein phosphatase 1 regulatory subunit 1A | Q9ERT9 | 0.8000 | 0.5751 | 0.6617 | **0.6789** |
| Protein S100-A13 | P97352 | 0.8190 | 0.7916 | 0.7685 | **0.7931** |
| Protein SET | Q9EQU5 | 0.8827 | 0.6012 | 0.7777 | **0.7538** |
| Purkinje cell protein 4 | P63054 | 0.7642 | 0.7489 | 0.7429 | **0.7520** |
| Ribonuclease UK114 | P52760 | 0.7190 | 0.7269 | 0.8995 | **0.7818** |
| [a]Secretogranin-2 | Q03517 | 0.6507 | 0.7775 | 0.6746 | **0.7009** |
| Sel1 repeat-containing protein 1 | Q921H9 | ND | 0.7625 | 0.7831 | **0.7728** |
| Selenocysteine lyase | Q9JLI6 | 0.6435 | 0.7944 | 0.8243 | **0.7541** |
| [a]Serum albumin | P07724 | 0.8106 | 0.6386 | 0.7181 | **0.7224** |
| Short coiled-coil protein | Q78YZ6 | 0.7584 | 0.8190 | 0.6557 | **0.7444** |
| Small ubiquitin-related modifier 2 | P61957 | 0.7469 | 0.6778 | 0.8079 | **0.7442** |
| SPARC-like protein 1 | P70663 | 0.7341 | 0.7125 | 0.7585 | **0.7350** |
| Spermatogenesis-associated protein 7 homolog | Q80VP2 | 0.5445 | 0.7604 | ND | **0.6525** |
| Stathmin | P54227 | 0.7695 | 0.6745 | 0.7912 | **0.7451** |
| Stathmin-2 | P55821 | 0.7584 | 0.6778 | 0.8375 | **0.7579** |
| Transcription elongation factor A protein 1 | P10711 | ND | 0.7056 | 0.7956 | **0.7506** |
| Transcription factor BTF3 homolog 4 | Q9CQH7 | 0.7448 | 0.7305 | 0.8141 | **0.7631** |
| Transketolase | P40142 | 0.8473 | 0.5937 | 0.7836 | **0.7416** |
| U6 snRNA-associated Sm-like protein LSm2 | O35900 | ND | 0.7417 | 0.6756 | **0.7086** |
| Uncharacterized protein C11orf96 homolog | Q3UPL5 | ND | 0.7090 | 0.6737 | **0.6914** |
| WAS/WASL-interacting protein family member 3 | P0C7L0 | 0.7521 | 0.7391 | 0.7171 | **0.7361** |

**Table S7. Differentially expressed proteins in adult**

ND, not detected; [a]detected in the same comparison at P21; [b]differential expression between sexes in control and ELS reared hippocampus; [c]altered in males and females in response to ELS

| Male ELS vs Male control - 20 proteins | | | | | |
|---|---|---|---|---|---|
| **Protein Name** | **Uniprot ID** | **Experiment 1** | **Experiment 2** | **Experiment 3** | **Average** |
| **Upregulated in stress - 8 proteins** | | | | | |
| Alpha-internexin | P46660 | 1.5850 | 1.2061 | 0.9174 | **1.2361** |
| [c]Coiled-coil domain-containing protein 93 | Q7TQK5 | 1.5205 | 1.3034 | 1.0025 | **1.2754** |
| [c]Cytochrome c oxidase subunit 2 | P00405 | 1.6016 | 1.2599 | ND | **1.4307** |
| Eukaryotic translation initiation factor 4 gamma 2 | Q62448 | 1.5546 | 1.2220 | 1.0278 | **1.2681** |
| Inactive phospholipase C-like protein 2 | Q8K394 | 1.5236 | 1.2128 | 0.9250 | **1.2205** |
| Neurofilament heavy polypeptide | P19246 | 1.6976 | 1.2408 | 1.0767 | **1.3384** |
| Superoxide dismutase [Cu-Zn] | P08228 | 1.7000 | 1.2069 | 1.0523 | **1.3197** |
| [c]Syntaxin-binding protein 5 | Q8K400 | 1.3045 | 1.3152 | 1.0530 | **1.2243** |
| **Downregulated in stress - 12 proteins** | | | | | |
| 1,5-anhydro-D-fructose reductase | Q9DCT1 | ND | 0.7996 | 0.7693 | **0.7844** |
| [c]60S ribosomal protein L22 | P67984 | 0.7482 | 0.7591 | ND | **0.7536** |
| Adapter molecule crk | Q64010 | 0.7741 | 0.7809 | 0.8194 | **0.7915** |
| Aspartate aminotransferase, cytoplasmic | P05201 | 0.5107 | 0.7649 | 0.9199 | **0.7318** |
| Hydroxyacyl-coenzyme A dehydrogenase, mitocho | Q61425 | 0.6536 | 0.7974 | ND | **0.7255** |
| Phosphatidylethanolamine-binding protein 1 | P70296 | 0.7518 | 0.7455 | 0.8542 | **0.7838** |
| Prothymosin alpha | P26350 | 0.5989 | 0.9210 | 0.7344 | **0.7514** |
| Sorbitol dehydrogenase | Q64442 | 0.6771 | 0.7935 | ND | **0.7353** |
| Threonine synthase-like 1 | Q8BH55 | 0.6464 | 0.9905 | 0.7493 | **0.7954** |
| [c]UPF0366 protein C11orf67 homolog | Q8R0P4 | 0.6509 | 0.7266 | ND | **0.6888** |
| WD repeat-containing protein 1 | O88342 | 0.6823 | 0.7974 | 0.8941 | **0.7913** |
| [c]Xaa-Pro dipeptidase | Q11136 | 0.5718 | 0.7256 | 0.9942 | **0.7639** |

| Female ELS vs Female control - 221 proteins | | | | | |
|---|---|---|---|---|---|
| **Protein Name** | **Uniprot ID** | **Experiment 1** | **Experiment 2** | **Experiment 3** | **Average** |
| **Upregulated in stress - 90 proteins** | | | | | |
| 26S proteasome non-ATPase regulatory subunit 2 | Q8VDM4 | 1.2124 | 1.6600 | 1.1316 | **1.3347** |
| 60S ribosomal protein L28 | P41105 | ND | 1.7583 | 1.2556 | **1.5070** |
| Aconitate hydratase, mitochondrial | Q99KI0 | 1.2328 | 1.3568 | 1.0617 | **1.2171** |
| Actin-related protein 2 | P61161 | 1.3575 | 1.3106 | 1.0931 | **1.2537** |
| Adenylate cyclase type 9 | P51830 | 1.3622 | 1.7779 | 0.9906 | **1.3769** |
| ADP/ATP translocase 1 | P48962 | 1.5421 | 1.6635 | 0.9211 | **1.3756** |
| AFG3-like protein 2 | Q8JZQ2 | 1.3745 | 1.3606 | 1.1846 | **1.3066** |
| Alanine--tRNA ligase, cytoplasmic | Q8BGQ7 | 1.3716 | 1.3549 | 1.1137 | **1.2801** |
| Alpha-adducin | Q9QYC0 | 1.2552 | 1.3133 | 1.0848 | **1.2178** |
| AP-2 complex subunit alpha-1 | P17426 | 1.4339 | 1.4858 | 0.9865 | **1.3020** |
| AP-2 complex subunit alpha-2 | P17427 | 1.3022 | 1.4612 | 1.0728 | **1.2787** |
| AP-2 complex subunit mu | P84091 | 1.4329 | 1.6890 | 1.1395 | **1.4205** |
| ATP synthase subunit b, mitochondrial | Q9CQQ7 | 1.4171 | 1.6057 | 1.1846 | **1.4025** |
| ATP synthase subunit O, mitochondrial | Q9DB20 | 1.2226 | 1.3911 | 1.0691 | **1.2276** |
| Calcium uniporter protein, mitochondrial | Q3UMR5 | 1.3917 | 1.3748 | 1.0220 | **1.2628** |
| Calcium/calmodulin-dependent protein kinase type | P11798 | 1.2285 | 1.6236 | 1.0893 | **1.3138** |
| Calcium-dependent secretion activator 1 | Q80TJ1 | 1.2578 | 1.5629 | 1.0728 | **1.2978** |
| Calcium-dependent secretion activator 2 | Q8BYR5 | 1.3203 | 1.5650 | 1.0654 | **1.3169** |
| Citrate synthase, mitochondrial | Q9CZU6 | 1.4210 | 1.3316 | 0.9496 | **1.2341** |
| Clathrin coat assembly protein AP180 | Q61548 | 1.2328 | 1.4582 | 0.9463 | **1.2124** |
| Clathrin heavy chain 1 | Q68FD5 | 1.2630 | 1.4421 | 1.0017 | **1.2356** |
| CLIP-associating protein 1 | Q80TV8 | 1.1671 | 1.5159 | 1.2972 | **1.3267** |
| CLIP-associating protein 2 | Q8BRT1 | 1.3397 | 1.2925 | 1.0639 | **1.2321** |
| [c]Coiled-coil domain-containing protein 93 | Q7TQK5 | 1.2234 | 1.5456 | 1.0810 | **1.2834** |
| Coronin-1B | Q9WUM3 | ND | 1.2330 | 1.2179 | **1.2255** |
| Cyclin-G-associated kinase | Q80TZ3 | 1.2561 | 1.3911 | 1.0728 | **1.2400** |
| [c]Cytochrome c oxidase subunit 2 | P00405 | 1.3603 | 1.6113 | ND | **1.4858** |
| Cytochrome c oxidase subunit 6C | Q9CPQ1 | ND | 1.2271 | 1.4235 | **1.3253** |
| Cytoplasmic dynein 1 intermediate chain 2 | O88487 | 1.2869 | 1.5201 | 1.1538 | **1.3203** |
| [a]Diacylglycerol kinase beta | Q6NS52 | ND | 1.4411 | 1.2444 | **1.3427** |
| DmX-like protein 2 | Q8BPN8 | 1.4014 | 1.3796 | 1.0413 | **1.2741** |
| Dynamin-1 | P39053 | 1.2754 | 1.4951 | 1.1114 | **1.2939** |
| Dynamin-3 | Q8BZ98 | 1.4151 | 1.5107 | 1.0632 | **1.3297** |
| Geranylgeranyl transferase type-2 subunit beta | P53612 | 1.3121 | 1.2186 | ND | **1.2653** |
| Glutamate receptor 1 | P23818 | 1.3622 | 1.3160 | 1.0320 | **1.2367** |
| Guanine nucleotide-binding protein G(i) subunit alp | B2RSH2 | 1.4589 | 1.5781 | 0.8371 | **1.2914** |
| Guanine nucleotide-binding protein G(i) subunit alp | P08752 | 1.4589 | 1.5414 | 0.8290 | **1.2764** |
| Guanine nucleotide-binding protein G(k) subunit al | Q9DC51 | 1.3908 | 1.4125 | 0.8702 | **1.2245** |
| Guanine nucleotide-binding protein G(o) subunit al | P18872 | 1.2941 | 1.5339 | 0.9065 | **1.2448** |
| Guanylate cyclase soluble subunit beta-1 | O54865 | 1.4053 | 1.3133 | 0.9790 | **1.2325** |
| Huntingtin | P42859 | 1.2896 | 1.3796 | 0.9388 | **1.2026** |
| Immunity-related GTPase family Q protein | Q8VIM9 | 1.0784 | 1.4633 | 1.2652 | **1.2690** |
| Lipoamide acyltransferase component of branched | P53395 | ND | 1.4213 | 1.2247 | **1.3230** |
| Metabotropic glutamate receptor 2 | Q14BI2 | 1.3139 | 1.7305 | 0.8261 | **1.2902** |
| Methionine--tRNA ligase, cytoplasmic | Q68FL6 | 1.2259 | 1.5013 | 1.0241 | **1.2505** |
| Mitochondrial glutamate carrier 2 | Q9DB41 | 1.2561 | 1.6774 | 0.9179 | **1.2838** |
| Mitochondrial import receptor subunit TOM70 | Q9CZW5 | 1.2745 | 1.5467 | 1.0234 | **1.2815** |
| Mitogen-activated protein kinase 1 | P63085 | 1.3176 | 1.2890 | 1.0384 | **1.2150** |
| Myeloid-associated differentiation marker | O35682 | 1.2158 | 1.3465 | ND | **1.2811** |
| Myotubularin-related protein 5 | Q6ZPE2 | 1.3841 | 1.3940 | 1.0370 | **1.2717** |
| NADH dehydrogenase [ubiquinone] 1 alpha subco | Q7TMF3 | 1.3277 | 1.2907 | 0.9836 | **1.2007** |
| NADH dehydrogenase [ubiquinone] 1 alpha subco | Q9DC69 | ND | 1.2563 | 1.2954 | **1.2759** |
| NADH-cytochrome b5 reductase 3 | Q9DCN2 | 1.7837 | 1.3549 | 1.0384 | **1.3924** |
| Nck-associated protein 1 | P28660 | 1.4043 | 1.2093 | 1.0522 | **1.2220** |
| NEDD8-activating enzyme E1 regulatory subunit | Q8VBW6 | 1.2657 | 1.8028 | 1.0721 | **1.3802** |
| Pantothenate kinase 4 | Q80YV4 | 1.2932 | 1.6543 | 0.9968 | **1.3148** |
| Phosphate carrier protein, mitochondrial | Q8VEM8 | 1.4428 | 1.5489 | 1.1301 | **1.3739** |
| Phosphatidylinositol-binding clathrin assembly prot | Q7M6Y3 | 1.3697 | 1.6797 | 1.0370 | **1.3621** |
| Piezo-type mechanosensitive ion channel compone | Q8CD54 | 1.2328 | 1.3252 | ND | **1.2790** |
| Plasma membrane calcium-transporting ATPase 2 | Q9R0K7 | 1.2789 | 1.3930 | 1.0878 | **1.2532** |
| Plexin-A1 | P70206 | 1.2049 | 1.3559 | 1.1498 | **1.2369** |
| Plexin-A4 | Q80UG2 | 1.3268 | 1.3587 | 1.0044 | **1.2300** |
| Prolyl endopeptidase-like | Q8C167 | 1.0438 | 1.4272 | 1.2392 | **1.2367** |
| Protein FAM49B | Q921M7 | 1.2448 | 1.4982 | 1.2556 | **1.3329** |
| Protein kinase C beta type | P68404 | 1.2914 | 1.5759 | 0.9886 | **1.2853** |
| Rap guanine nucleotide exchange factor 2 | Q8CHG7 | 1.2413 | 1.3437 | 1.0661 | **1.2171** |
| Rho GTPase-activating protein 1 | Q5FWK3 | 1.3249 | 1.5902 | 1.1153 | **1.3435** |
| Ryanodine receptor 2 | E9Q401 | 1.2066 | 1.5683 | 0.9756 | **1.2502** |
| Serine/threonine-protein kinase DCLK1 | Q9JLM8 | 1.3995 | 1.3549 | 1.0348 | **1.2631** |
| Serine/threonine-protein kinase TAO2 | Q6ZQ29 | ND | 1.3988 | 1.2095 | **1.3042** |
| Serine/threonine-protein phosphatase 2A 65 kDa r | Q76MZ3 | 1.2824 | 1.3843 | 1.0916 | **1.2528** |
| Serpin B6 | Q60854 | 1.2183 | 1.2988 | ND | **1.2586** |
| Sodium-driven chloride bicarbonate exchanger | Q5DTL9 | 1.3258 | 1.3930 | 1.1675 | **1.2954** |
| Solute carrier family 12 member 5 | Q91V14 | 1.4498 | 1.4562 | 0.9886 | **1.2982** |
| Spectrin beta chain, erythrocyte | P15508 | 1.3612 | 1.6046 | 0.9696 | **1.3118** |
| Sphingomyelin phosphodiesterase 3 | Q9JJY3 | 1.4448 | 1.6079 | 1.0573 | **1.3700** |
| Striated muscle-specific serine/threonine-protein k | Q62407 | 1.0927 | 1.3261 | 1.3392 | **1.2527** |
| Succinyl-CoA ligase [ADP-forming] subunit beta, r | Q9Z2I9 | 1.2243 | 1.3681 | 1.0356 | **1.2093** |
| Synaptic vesicle glycoprotein 2A | Q9JIS5 | 1.4005 | 1.5286 | 1.0199 | **1.3163** |
| Syntaxin-binding protein 1 | O08599 | 1.3946 | 1.3409 | 1.0999 | **1.2785** |
| [c]Syntaxin-binding protein 5 | Q8K400 | 1.4112 | 1.2271 | 1.0961 | **1.2448** |
| Syntaxin-binding protein 5-like | Q5DQR4 | 1.2789 | 1.8483 | 1.0999 | **1.4090** |
| Tectonin beta-propeller repeat-containing protein 1 | Q80VP0 | 1.6188 | 1.3521 | 0.9777 | **1.3162** |
| Trifunctional enzyme subunit alpha, mitochondrial | Q8BMS1 | 1.3295 | 1.5607 | 1.0566 | **1.3156** |
| Uncharacterized protein KIAA0564 homolog | Q8CC88 | ND | 1.2520 | 1.2740 | **1.2630** |
| Unconventional myosin-Id | Q5SYD0 | 1.3659 | 1.5661 | 1.1498 | **1.3606** |
| UPF0364 protein C6orf211 homolog | A6H630 | 1.2914 | 1.3748 | 1.0370 | **1.2344** |
| Vacuolar protein sorting-associated protein 53 hon | Q8CCB4 | 1.2285 | 1.5489 | 1.1458 | **1.3077** |
| V-type proton ATPase subunit d 1 | P51863 | 1.2639 | 1.5138 | 1.1246 | **1.3008** |
| WD repeat-containing protein 7 | Q920I9 | 1.6065 | 1.6046 | 1.0320 | **1.4143** |
| **Downregulated in stress - 131 proteins** | | | | | |
| 28 kDa heat- and acid-stable phosphoprotein | Q3UHX2 | 0.6844 | 0.6927 | 0.8052 | **0.7274** |
| 40S ribosomal protein S28 | P62858 | 0.5633 | 0.7413 | 0.3534 | **0.5527** |
| 60S acidic ribosomal protein P2 | P99027 | 0.4979 | 0.6030 | 0.9109 | **0.6706** |
| [c]60S ribosomal protein L22 | P67984 | 0.7960 | 0.6269 | ND | **0.7114** |
| Acid ceramidase | Q9WV54 | 0.7229 | 0.6893 | ND | **0.7061** |
| Activated RNA polymerase II transcriptional coacti | P11031 | ND | 0.7347 | 0.5364 | **0.6355** |
| Acyl-CoA-binding protein | P31786 | 0.7641 | 0.7553 | ND | **0.7597** |
| [a]Alpha-2-macroglobulin receptor-associated protei | P55302 | 0.6293 | 0.6812 | 0.8860 | **0.7322** |
| Alpha-taxilin | Q6PAM1 | 0.6117 | 0.7654 | ND | **0.6885** |
| Amyloid beta A4 protein | P12023 | 0.7484 | 0.7455 | 0.7197 | **0.7378** |
| Annexin A11 | P97384 | 0.7988 | 0.7327 | 0.8074 | **0.7796** |
| Annexin A3 | O35639 | 0.7818 | 0.6230 | 0.9179 | **0.7742** |
| [a]Aspartyl aminopeptidase | Q9Z2W0 | 0.7753 | 0.6449 | 0.9589 | **0.7930** |
| Brain acid soluble protein 1 | Q91XV3 | 0.6497 | 0.7347 | 0.8660 | **0.7501** |
| Brevican core protein | Q61361 | 0.7366 | 0.7038 | 0.9166 | **0.7857** |
| Calbindin | P12658 | 0.7224 | 0.9184 | 0.7358 | **0.7922** |
| Calcineurin subunit B type 1 | Q63810 | 0.6825 | 0.8214 | 0.7242 | **0.7427** |
| Calcium/calmodulin-dependent protein kinase type | Q8BW96 | 0.7259 | 0.8068 | 0.7581 | **0.7636** |
| Calcium-binding protein p22 | P61022 | 0.6346 | 0.7067 | 0.6027 | **0.6480** |

| | | | | | |
|---|---|---|---|---|---|
| Calmodulin | P62204 | 0.7625 | 0.7072 | 0.8452 | **0.7717** |
| Calretinin | Q08331 | 0.9131 | 0.7111 | 0.7607 | **0.7950** |
| Cell adhesion molecule 1 | Q8R5M8 | 0.6906 | 0.6431 | 0.8052 | **0.7130** |
| [a]Cellular nucleic acid-binding protein | P53996 | 0.6565 | 0.7522 | ND | **0.7044** |
| [a]Charged multivesicular body protein 4b | Q9D8B3 | 0.5803 | 0.6123 | ND | **0.5963** |
| Charged multivesicular body protein 5 | Q9D7S9 | 0.6220 | 0.6999 | ND | **0.6609** |
| Chondroitin sulfate proteoglycan 5 | Q71M36 | 0.6671 | 0.8028 | 0.7991 | **0.7563** |
| Clathrin light chain A | O08585 | 0.7922 | 0.5552 | 0.9021 | **0.7499** |
| Clathrin light chain B | Q6IRU5 | 0.6250 | 0.7506 | 0.8891 | **0.7549** |
| C-Myc-binding protein | Q9EQS3 | 0.6949 | 0.7962 | ND | **0.7456** |
| Complexin-2 | P84086 | 0.8362 | 0.6626 | 0.5858 | **0.6949** |
| COP9 signalosome complex subunit 8 | Q8VBV7 | 0.5799 | 0.7486 | 0.6477 | **0.6587** |
| Cystatin-C | P21460 | 0.7135 | 0.6157 | 0.9046 | **0.7446** |
| [a]Cysteine and glycine-rich protein 1 | P97315 | 0.8845 | 0.6718 | 0.4162 | **0.6575** |
| Cytochrome b-c1 complex subunit 6, mitochondria | P99028 | 0.4438 | 0.6047 | 0.6976 | **0.5820** |
| Cytochrome c oxidase subunit 6B1 | P56391 | 0.6570 | 0.6383 | ND | **0.6476** |
| D-dopachrome decarboxylase | O35215 | ND | 0.7455 | 0.5694 | **0.6574** |
| Dihydrolipoyl dehydrogenase, mitochondrial | O08749 | 0.7748 | 0.7146 | 0.6504 | **0.7133** |
| Dihydropteridine reductase | Q8BVI4 | 0.7759 | 0.6603 | 0.5117 | **0.6493** |
| DnaJ homolog subfamily B member 2 | Q9QYI5 | 0.6671 | 0.7728 | 0.7772 | **0.7390** |
| D-tyrosyl-tRNA(Tyr) deacylase 1 | Q9DD18 | 0.6488 | 0.6965 | ND | **0.6727** |
| Dynactin subunit 2 | Q99KJ8 | 0.7396 | 0.7023 | 0.7729 | **0.7383** |
| Endoplasmic reticulum resident protein 44 | Q9D1Q6 | 0.6355 | 0.7643 | 0.7602 | **0.7200** |
| Eukaryotic translation initiation factor 4H | Q9WUK2 | 0.6877 | 0.7669 | 0.7217 | **0.7254** |
| Far upstream element-binding protein 2 | Q3U0V1 | 0.7905 | 0.7367 | 0.8541 | **0.7938** |
| F-box only protein 2 | Q80UW2 | 0.6925 | 0.5780 | ND | **0.6353** |
| GMP reductase 2 | Q99L27 | 0.7982 | 0.5152 | ND | **0.6567** |
| Heterogeneous nuclear ribonucleoprotein A/B | Q99020 | 0.5370 | 0.7707 | 0.9615 | **0.7564** |
| Heterogeneous nuclear ribonucleoprotein D0 | Q60668 | 0.7071 | 0.5579 | 0.8768 | **0.7140** |
| Integrin beta-1 | P09055 | 0.5347 | 0.6414 | ND | **0.5880** |
| Integrin beta-2 | P11835 | 0.6484 | 0.7929 | ND | **0.7206** |
| Lactoylglutathione lyase | Q9CPU0 | ND | 0.7347 | 0.6701 | **0.7024** |
| LDLR chaperone MESD | Q9ERE7 | 0.7071 | 0.6110 | 0.6747 | **0.6643** |
| Leucine-rich repeat-containing protein 16C | Q3V3V9 | 0.6666 | 0.5724 | 0.9609 | **0.7333** |
| Leucine-rich repeat-containing protein 59 | Q922Q8 | 0.5727 | 0.7251 | 1.0428 | **0.7802** |
| Limbic system-associated membrane protein | Q8BLK3 | 0.6745 | 0.7979 | 0.6490 | **0.7071** |
| Lysosomal alpha-glucosidase | P70699 | 0.6935 | 0.5943 | 0.9529 | **0.7469** |
| [a]Malate dehydrogenase, mitochondrial | P08249 | 0.8645 | 0.5289 | 0.7297 | **0.7077** |
| Membrane-associated progesterone receptor com | Q80UU9 | 0.6872 | 0.6199 | 0.9913 | **0.7662** |
| Methionine aminopeptidase 2 | O08663 | 0.6470 | 0.5976 | ND | **0.6223** |
| Microtubule-associated protein 4 | P27546 | 0.5715 | 0.7934 | 1.0031 | **0.7893** |
| [a]Mitochondrial import inner membrane translocase | P62073 | 0.6920 | 0.7755 | ND | **0.7338** |
| NADH dehydrogenase [ubiquinone] 1 alpha subco | Q9CQ75 | 0.7775 | 0.6621 | 0.6410 | **0.6935** |
| NADH dehydrogenase [ubiquinone] flavoprotein 2, | Q9D6J6 | 0.6147 | 0.5976 | 0.9503 | **0.7208** |
| NADH dehydrogenase [ubiquinone] iron-sulfur pro | Q9CXZ1 | 0.6267 | 0.7146 | 0.9968 | **0.7794** |
| Nascent polypeptide-associated complex subunit a | P70670 | 0.7310 | 0.6585 | 0.7783 | **0.7226** |
| Neuronal calcium sensor 1 | Q8BNY6 | ND | 0.5812 | 0.6187 | **0.6000** |
| [a]Neuronal cell adhesion molecule | Q810U4 | 0.7479 | 0.6654 | 0.9702 | **0.7945** |
| Neuron-specific calcium-binding protein hippocalci | P84075 | 0.6272 | 0.7077 | 0.9982 | **0.7777** |
| Neuroplastin | P97300 | 0.6680 | 0.6975 | 0.8612 | **0.7422** |
| [a]Nitrilase homolog 1 | Q8VDK1 | ND | 0.6217 | 0.6222 | **0.6219** |
| NSFL1 cofactor p47 | Q9CZ44 | 0.7376 | 0.6672 | 0.9294 | **0.7781** |
| Nuclear fragile X mental retardation-interacting pro | Q5F2E7 | 0.7526 | 0.5363 | ND | **0.6445** |
| Nucleobindin-1 | Q02819 | 0.6008 | 0.6955 | ND | **0.6482** |
| Nucleolar protein 3 | Q9D1X0 | 0.5045 | 0.6951 | ND | **0.5998** |
| Nucleolin | P09405 | 0.6717 | 0.7532 | 0.8866 | **0.7705** |
| Paralemmin-1 | Q9Z0P4 | 0.7110 | 0.7072 | 0.8922 | **0.7701** |
| [a]Parathymosin | Q9D0J8 | 0.7290 | 0.6485 | 0.7848 | **0.7208** |
| Peptidyl-prolyl cis-trans isomerase B | P24369 | 0.7100 | 0.5689 | 1.1068 | **0.7952** |
| Peptidyl-prolyl cis-trans isomerase FKBP3 | Q62446 | 0.5629 | 0.6161 | 0.8267 | **0.6685** |
| Peptidyl-prolyl cis-trans isomerase NIMA-interactir | Q9QUR7 | 0.6863 | 0.8623 | 0.7348 | **0.7611** |
| Peroxiredoxin-6 | O08709 | 0.6968 | 0.6234 | 0.7676 | **0.6959** |
| Peroxisomal biogenesis factor 19 | Q8VCI5 | 0.6529 | 0.6135 | 0.8836 | **0.7167** |
| Phosphoglycerate mutase 1 | Q9DBJ1 | 0.8339 | 0.7664 | 0.6586 | **0.7530** |
| PITH domain-containing protein 1 | Q8BWR2 | 0.6194 | 0.6658 | ND | **0.6426** |
| Plasminogen activator inhibitor 1 RNA-binding prot | Q9CY58 | 0.7878 | 0.6195 | 0.8535 | **0.7536** |
| [a]Prefoldin subunit 1 | Q9CWM4 | 0.7531 | 0.6893 | ND | **0.7212** |
| Prefoldin subunit 3 | P61759 | 0.5688 | 0.6714 | ND | **0.6201** |
| ProSAAS | Q9QXV0 | 0.7046 | 0.6903 | 0.9682 | **0.7877** |
| Proteasome subunit alpha type-1 | Q9R1P4 | 0.8444 | 0.7564 | 0.7892 | **0.7966** |
| Proteasome subunit beta type-4 | P99026 | ND | 0.7616 | 0.6696 | **0.7156** |
| Protein canopy homolog 2 | Q9QXT0 | 0.6293 | 0.7327 | ND | **0.6810** |
| Protein DJ-1 | Q99LX0 | 0.7479 | 0.6989 | 0.7222 | **0.7230** |
| Protein FAM192A | Q91WE2 | 0.6797 | 0.6746 | ND | **0.6772** |
| Protein phosphatase 1 regulatory subunit 1A | Q9ERT9 | 0.5509 | 0.7038 | ND | **0.6273** |
| Protein phosphatase 1 regulatory subunit 1B | Q60829 | 0.7684 | 0.6999 | 0.8452 | **0.7712** |
| Protein SET | Q9EQU5 | 0.6346 | 0.5800 | 0.9015 | **0.7054** |
| Protein Shroom2 | A2ALU4 | 0.4201 | 0.6789 | 0.8762 | **0.6584** |
| Radixin | P26043 | 0.7095 | 0.7004 | 0.9476 | **0.7858** |
| Ran-specific GTPase-activating protein | P34022 | ND | 0.7501 | 0.7581 | **0.7541** |
| Reticulocalbin-1 | Q05186 | 0.7295 | 0.7486 | ND | **0.7390** |
| Reticulocalbin-2 | Q8BP92 | 0.7335 | ND | 0.5294 | **0.6315** |
| [a]Rho GDP-dissociation inhibitor 1 | Q99PT1 | 0.7856 | 0.6526 | 0.8013 | **0.7465** |
| [a]Ribosome-recycling factor, mitochondrial | Q9D6S7 | 0.4776 | 0.7480 | ND | **0.6128** |
| RILP-like protein 1 | Q9JJC6 | 0.7279 | ND | 0.7157 | **0.7218** |
| RNA-binding protein FUS | P56959 | 0.2720 | 0.7383 | 1.0529 | **0.6877** |
| RWD domain-containing protein 1 | Q9CQK7 | 0.7125 | ND | 0.7676 | **0.7400** |
| Secretogranin-3 | P47867 | 0.7594 | 0.7156 | ND | **0.7375** |
| Serine/arginine-rich splicing factor 1 | Q6PDM2 | ND | 0.7216 | 0.7697 | **0.7456** |
| Serum albumin | P07724 | 0.6324 | 0.5626 | 0.6432 | **0.6127** |
| SH3 domain-binding glutamic acid-rich-like protein | Q9JJU8 | 0.6552 | 0.7470 | 0.7267 | **0.7096** |
| SH3 domain-containing kinase-binding protein 1 | Q8R550 | 0.6868 | 0.8192 | 0.7628 | **0.7562** |
| Small glutamine-rich tetratricopeptide repeat-conta | Q8BJU0 | 0.7007 | 0.9455 | 0.5964 | **0.7476** |
| S-phase kinase-associated protein 1 | Q9WTX5 | 0.5613 | 0.6718 | 0.8505 | **0.6946** |
| Synaptotagmin-17 | Q920M7 | ND | 0.7885 | 0.6179 | **0.7032** |
| Syntaxin-8 | O88983 | 0.6615 | 0.8786 | 0.6410 | **0.7270** |
| Thioredoxin | P10639 | 0.6615 | 0.7057 | 0.8301 | **0.7325** |
| Thrombospondin type-1 domain-containing protein | Q69ZU6 | 0.7684 | 0.7627 | 0.8452 | **0.7921** |
| [a]Transcription elongation factor A protein 1 | P10711 | 0.7484 | 0.7316 | 0.7991 | **0.7597** |
| Tropomodulin-2 | Q9JKK7 | 0.7264 | 0.6709 | 0.7985 | **0.7320** |
| Tropomyosin alpha-1 chain | P58771 | 0.6588 | 0.7111 | 0.6761 | **0.6820** |
| Tropomyosin alpha-3 chain | P21107 | 0.5567 | 0.8068 | 0.7379 | **0.7004** |
| [a]U6 snRNA-associated Sm-like protein LSm2 | O35900 | 0.6448 | 0.6635 | ND | **0.6542** |
| Uncharacterized protein C6orf203 homolog | Q9CQF4 | ND | 0.4698 | 0.7740 | **0.6219** |
| [c]UPF0366 protein C11orf67 homolog | Q8R0P4 | 0.7046 | 0.7744 | ND | **0.7395** |
| UPF0449 protein C19orf25 homolog | Q9D7E4 | 0.6008 | 0.6798 | ND | **0.6403** |
| UPF0515 protein C19orf66 homolog | Q8CAK3 | 0.6024 | 0.6553 | ND | **0.6289** |
| UV excision repair protein RAD23 homolog B | P54728 | 0.7110 | 0.6677 | 0.9314 | **0.7700** |
| Vesicle-associated membrane protein 2 | P63044 | 0.7599 | 0.8289 | 0.7450 | **0.7779** |
| Visinin-like protein 1 | P62761 | 0.6643 | 0.6903 | 1.0065 | **0.7870** |
| V-type proton ATPase subunit F | Q9D1K2 | 0.6045 | 0.7256 | 0.9968 | **0.7756** |
| [c]Xaa-Pro dipeptidase | Q11136 | 0.8603 | 0.5930 | 0.7466 | **0.7333** |

| Male control vs Female control - 56 proteins | | | | | |
|---|---|---|---|---|---|
| **Protein Name** | **Uniprot ID** | **Experiment 1** | **Experiment 2** | **Experiment 3** | **Average** |
| **Lower in females - 19 proteins** | | | | | |
| [b]Acidic leucine-rich nuclear phosphoprotein 32 fam | O35381 | 2.2012 | 1.2693 | 0.9481 | **1.4729** |
| [b]Alpha-endosulfine | P60840 | 1.8561 | 1.2226 | 1.0534 | **1.3774** |
| [b]Apolipoprotein A-I | Q00623 | 1.6047 | 0.9520 | 1.3737 | **1.3101** |
| [b]ATP synthase-coupling factor 6, mitochondrial | P97450 | 1.5947 | 1.0154 | 1.2338 | **1.2813** |
| Bifunctional coenzyme A synthase | Q9DBL7 | ND | 1.2235 | 1.2117 | **1.2176** |
| Bifunctional protein NCOAT | Q9EQQ9 | 1.9350 | 1.3689 | 0.9700 | **1.4246** |
| Carbonic anhydrase 4 | Q64444 | 1.2011 | ND | 1.2025 | **1.2018** |
| [b]Dematin | Q9WV69 | 1.5576 | 1.2150 | 1.0035 | **1.2587** |
| Diacylglycerol kinase beta | Q6NS52 | ND | 1.2075 | 1.2193 | **1.2134** |
| Elongation factor 1-alpha 1 | P10126 | 1.5171 | 1.2303 | 1.1843 | **1.3106** |
| MKL/myocardin-like protein 2 | P59759 | 1.4776 | 1.2041 | 1.0541 | **1.2453** |
| NF-kappa-B essential modulator | O88522 | 1.1245 | 1.2329 | 1.2746 | **1.2107** |
| Pre-mRNA-processing factor 19 | Q99KP6 | ND | 1.3140 | 1.2261 | **1.2701** |
| Prothymosin alpha | P26350 | 2.7289 | 1.0403 | 1.3205 | **1.6966** |
| RING finger protein 214 | Q8BFU3 | 1.4746 | 1.2252 | 1.0267 | **1.2422** |
| Sortilin | Q6PHU5 | 1.2616 | 1.1769 | 1.3150 | **1.2512** |
| Thymosin beta-10 | Q6ZWY8 | 2.9166 | 1.2772 | ND | **2.0969** |
| Translin-associated protein X | Q9QZE7 | 1.5076 | 1.2588 | 1.0246 | **1.2637** |
| UMP-CMP kinase | Q9DBP5 | 1.4797 | 1.2363 | 1.0225 | **1.2461** |
| **Lower in males - 37 proteins** | | | | | |
| 39S ribosomal protein L28, mitochondrial | Q9D1B9 | 0.6990 | 0.4061 | ND | **0.5525** |
| 40S ribosomal protein S14 | P62264 | 0.7919 | 0.9952 | 0.5772 | **0.7881** |
| 60S ribosomal protein L18 | P35980 | 0.6454 | 0.8645 | 0.7961 | **0.7687** |
| 60S ribosomal protein L18a | P62717 | 0.3165 | 1.0512 | 0.7803 | **0.7160** |
| Acetoacetyl-CoA synthetase | Q9D2R0 | 0.7302 | ND | 0.5704 | **0.6503** |
| Acylamino-acid-releasing enzyme | Q8R146 | 0.6039 | 0.7743 | 1.0161 | **0.7981** |
| A-kinase anchor protein SPHKAP | Q6NSW3 | ND | 0.7989 | 0.7989 | **0.7989** |
| Bifunctional 3'-phosphoadenosine 5'-phosphosulfat | Q60967 | 0.7352 | ND | 0.7595 | **0.7474** |
| Biliverdin reductase A | Q9CY64 | 0.6719 | 0.7215 | 0.9693 | **0.7876** |
| BTB/POZ domain-containing protein KCTD12 | Q6WVG3 | 0.8493 | 0.7105 | 0.6325 | **0.7308** |
| Calpain-1 catalytic subunit | O35350 | 0.7429 | ND | 0.6760 | **0.7095** |
| Coatomer subunit epsilon | O89079 | 0.5599 | ND | 0.6812 | **0.6205** |
| Coatomer subunit gamma-2 | Q9QXK3 | ND | 0.7115 | 0.6998 | **0.7057** |
| COP9 signalosome complex subunit 7a | Q9CZ04 | 0.5096 | 0.8803 | 0.7934 | **0.7277** |
| COP9 signalosome complex subunit 8 | Q8VBV7 | 0.5623 | 0.7873 | 0.5732 | **0.6409** |
| Cullin-5 | Q9D5V5 | 0.5454 | 0.9196 | 0.7781 | **0.7477** |
| Cysteine-rich protein 2 | Q9DCT8 | ND | 0.5396 | 0.7945 | **0.6670** |
| D-dopachrome decarboxylase | O35215 | ND | 0.4481 | 0.7568 | **0.6025** |
| Dihydropteridine reductase | Q8BVI4 | 0.7087 | 0.8833 | 0.7469 | **0.7797** |
| Epoxide hydrolase 2 | P34914 | ND | 0.8000 | 0.7690 | **0.7845** |
| Importin-7 | Q9EPL8 | 0.5246 | 0.7160 | 1.0476 | **0.7627** |
| Kv channel-interacting protein 4 | Q6PHZ8 | ND | 0.6207 | 0.7879 | **0.7043** |
| NCK-interacting protein with SH3 domain | Q9ESJ4 | 0.7146 | 0.7180 | 0.9593 | **0.7973** |
| Peptidyl-prolyl cis-trans isomerase FKBP8 | O35465 | | 0.7351 | 0.7475 | **0.7413** |
| Phenylalanine--tRNA ligase alpha subunit | Q8C0C7 | 0.6540 | 0.5245 | 0.9120 | **0.6968** |
| Protein prune homolog | Q8BIW1 | 0.4545 | 0.6959 | 0.6732 | **0.6079** |
| Rab-like protein 1 | Q5U3K5 | 0.6463 | 0.7017 | ND | **0.6740** |
| Selenium-binding protein 2 | Q63836 | 0.8977 | 0.7260 | 0.7454 | **0.7897** |
| Sn1-specific diacylglycerol lipase alpha | Q6WQJ1 | 0.5890 | 0.7679 | 0.9019 | **0.7529** |
| Sodium/potassium-transporting ATPase subunit al | Q8VDN2 | 0.5046 | 0.8362 | 0.7994 | **0.7134** |
| Sorting nexin-4 | Q91YJ2 | 0.3935 | ND | 0.7972 | **0.5953** |
| Synaptotagmin-17 | Q920M7 | ND | 0.7047 | 0.5897 | **0.6472** |
| Tubulin beta-3 chain | Q9ERD7 | 0.4751 | 1.0644 | 0.7950 | **0.7782** |
| Ubiquitin carboxyl-terminal hydrolase 15 | Q8R5H1 | 0.6682 | 0.5996 | 1.0418 | **0.7698** |
| Vacuolar protein sorting-associated protein 33B | P59016 | 0.3772 | 0.7983 | 1.0823 | **0.7526** |
| Valine--tRNA ligase | Q9Z1Q9 | 0.6379 | 0.7652 | 0.9781 | **0.7937** |
| Voltage-dependent calcium channel gamma-8 subu | Q8VHW2 | 0.5213 | 0.4652 | 1.0267 | **0.6711** |

| Male ELS vs Female ELS - 70 proteins | | | | | |
|---|---|---|---|---|---|
| **Protein Name** | **Uniprot I** | **Experiment 1** | **Experiment 2** | **Experiment 3** | **Average** |
| **Lower in females - 68 proteins** | | | | | |
| 40S ribosomal protein S28 | P62858 | 2.8284 | 1.2761 | 1.9914 | **2.0320** |
| 60S acidic ribosomal protein P2 | P99027 | 3.5259 | 1.2559 | 1.0806 | **1.9541** |
| [c]Acidic leucine-rich nuclear phosphoprotein 32 fam | O35381 | 3.4082 | 1.2039 | 1.0694 | **1.8938** |
| [a]Acyl-CoA-binding protein | P31786 | 3.1888 | 1.2796 | ND | **2.2342** |
| Alpha-2-macroglobulin receptor-associated protein | P55302 | 2.5280 | 1.1758 | 1.2115 | **1.6384** |
| [a,c]Alpha-endosulfine | P60840 | 2.5263 | 1.1629 | 1.2779 | **1.6557** |
| [c]Apolipoprotein A-I | Q00623 | 3.9834 | 1.5387 | 1.0089 | **2.1770** |
| [c]ATP synthase-coupling factor 6, mitochondrial | P97450 | 2.3983 | 1.2014 | 1.0026 | **1.5341** |
| [a]ATPase inhibitor, mitochondrial | O35143 | 2.8859 | ND | 1.2344 | **2.0601** |
| Basigin | P18572 | 1.8391 | 1.3762 | 1.0481 | **1.4211** |
| Cadherin-13 | Q9WTR5 | 2.3768 | 1.2533 | 1.0503 | **1.5601** |
| Calcineurin subunit B type 1 | Q63810 | 3.2266 | 1.2594 | 1.4710 | **1.9856** |
| Calmodulin | P62204 | 2.2910 | 1.1807 | 1.2967 | **1.5895** |
| Calretinin | Q08331 | 1.5823 | 1.2131 | 1.3368 | **1.3774** |
| Calumenin | O35887 | 2.4318 | 1.2014 | 1.1327 | **1.5886** |
| Charged multivesicular body protein 5 | Q9D7S9 | 2.9876 | 1.3229 | ND | **2.1553** |
| Choline transporter-like protein 2 | Q8BY89 | 1.4330 | ND | 1.3102 | **1.3716** |
| Chondroitin sulfate proteoglycan 5 | Q71M36 | 2.3834 | 1.2620 | 1.1965 | **1.6139** |
| Complexin-2 | P84086 | 2.6851 | 1.2814 | 1.8971 | **1.9545** |
| Cysteine and glycine-rich protein 1 | P97315 | 2.0139 | 1.1334 | 1.3012 | **1.4828** |
| Cytochrome b5 | P56395 | 1.6483 | 1.2131 | 0.9902 | **1.2839** |
| Cytochrome b-c1 complex subunit 6, mitochondria | P99028 | 3.2378 | 1.1381 | 1.2182 | **1.8647** |
| Cytochrome c oxidase subunit 6B1 | P56391 | 2.7019 | 1.2446 | ND | **1.9733** |
| [a,c]Dematin | Q9WV69 | 1.8113 | 1.2576 | 0.9847 | **1.3512** |
| Dihydrolipoyl dehydrogenase, mitochondrial | O08749 | 1.8442 | 1.1178 | 1.2040 | **1.3887** |
| DnaJ homolog subfamily A member 1 | P63037 | 1.7195 | ND | 1.2242 | **1.4718** |
| DnaJ homolog subfamily C member 5 | P60904 | 2.0279 | ND | 1.2259 | **1.6269** |
| Heterogeneous nuclear ribonucleoprotein A/B | Q99020 | 4.1267 | 1.3312 | 1.0012 | **2.1530** |
| Integrin beta-1 | P09055 | 2.7001 | 1.2139 | ND | **1.9570** |
| LDLR chaperone MESD | Q9ERE7 | 1.9292 | 1.1750 | 1.3564 | **1.4869** |
| MARCKS-related protein | P28667 | 2.3768 | 1.2309 | 1.1485 | **1.5854** |
| Membrane-associated progesterone receptor comp | O55022 | 2.0392 | 1.2039 | 0.9800 | **1.4077** |
| Methionine aminopeptidase 2 | O08663 | 2.5775 | 1.2014 | ND | **1.8895** |
| NADH dehydrogenase [ubiquinone] 1 beta subcom | Q9D6J5 | 1.6245 | 1.2224 | 1.1063 | **1.3177** |
| NADH dehydrogenase [ubiquinone] flavoprotein 2, | Q9D6J6 | 2.2673 | 1.4022 | 0.9099 | **1.5265** |
| NADH dehydrogenase [ubiquinone] iron-sulfur prot | Q9CXZ1 | 2.3230 | 1.2072 | 0.9861 | **1.5054** |
| Nascent polypeptide-associated complex subunit a | P70670 | 1.9279 | 1.1303 | 1.2040 | **1.4207** |
| Neural cell adhesion molecule 1 | P13595 | 1.9092 | 1.2123 | 1.0518 | **1.3911** |
| Neurocan core protein | P55066 | 2.2736 | 1.2334 | 1.0627 | **1.5233** |
| Neuromodulin | P06837 | 2.3457 | 1.2039 | 1.0569 | **1.5355** |
| Neuronal calcium sensor 1 | Q8BNY6 | ND | 1.3137 | 1.7098 | **1.5118** |
| Noelin | O88998 | ND | 1.2681 | 1.2753 | **1.2717** |
| Nucleolin | P09405 | 3.1514 | 1.2131 | 1.0315 | **1.7987** |
| Oligodendrocyte-myelin glycoprotein | Q63912 | 1.9319 | 1.2481 | 1.0828 | **1.4209** |
| [a]Parathymosin | Q9D0J8 | 2.2768 | 1.2412 | 1.1858 | **1.5679** |
| Peptidyl-prolyl cis-trans isomerase FKBP3 | Q62446 | 3.1755 | 1.2300 | 1.1257 | **1.8437** |
| Peroxiredoxin-6 | O08709 | 1.8661 | 1.1319 | 1.2031 | **1.4004** |
| Phosphoglycerate mutase 1 | Q9DBJ1 | 2.1615 | 1.2123 | 1.0724 | **1.4820** |
| Phosphoglycerate mutase 2 | O70250 | 2.1615 | 1.2123 | 0.9220 | **1.4319** |
| Proline-rich transmembrane protein 2 | E9PUL5 | 1.8125 | 1.2097 | 1.0709 | **1.3644** |
| Protein FAM136A | Q9CR98 | 2.3916 | 1.2156 | 1.1557 | **1.5877** |
| Protein kinase C and casein kinase substrate in ne | Q61644 | 2.1750 | 1.2343 | 1.0672 | **1.4922** |
| [a]Protein phosphatase 1 regulatory subunit 1A | Q9ERT9 | 5.2343 | 1.2047 | ND | **3.2195** |
| Protein phosphatase 1G | Q61074 | 2.0237 | 1.0336 | 1.2378 | **1.4317** |
| Reticulocalbin-2 | Q8BP92 | 2.6703 | ND | 1.2216 | **1.9460** |
| [a]Secretogranin-2 | Q03517 | 2.1931 | 1.2064 | 1.0459 | **1.4818** |
| [a]Serum albumin | P07724 | 3.5162 | 1.2993 | 1.0753 | **1.9636** |
| Small glutamine-rich tetratricopeptide repeat-conta | Q8BJU0 | 1.6935 | 1.0229 | 1.2770 | **1.3311** |
| Src substrate cortactin | Q60598 | 2.6135 | 1.2437 | 1.0569 | **1.6380** |
| Synaptopodin | Q8CC35 | 1.4948 | 1.2446 | 0.9578 | **1.2324** |
| Transcription elongation regulator 1 | Q8CGF7 | 1.3794 | 1.0634 | 1.2682 | **1.2370** |
| Transcription factor A, mitochondrial | P40630 | 2.1780 | 1.2005 | ND | **1.6893** |
| Transmembrane emp24 domain-containing protein | Q99KF1 | 1.3547 | 1.1677 | 1.2534 | **1.2586** |
| Tropomodulin-2 | Q9JKK7 | 2.2501 | 1.2072 | 1.0627 | **1.5067** |
| Tropomyosin alpha-1 chain | P58771 | 2.4016 | 1.2097 | 1.2225 | **1.6113** |
| Tumor protein D54 | Q9CYZ2 | 1.7387 | 1.2039 | 0.9459 | **1.2962** |
| Repressor of yield of DENV | Q8CAK3 | 2.7094 | 1.2215 | ND | **1.9655** |
| Visinin-like protein 1 | P62761 | 2.4453 | 1.2275 | 1.0679 | **1.5802** |
| **Lower in males - 2 proteins** | | | | | |
| Syntaxin-binding protein 5-like | Q5DQR4 | 0.5415 | 0.7926 | 1.0258 | **0.7866** |
| UMP-CMP kinase 2, mitochondrial | Q3U5Q7 | 0.4945 | ND | 0.7769 | **0.6357** |

### Table S8. Enriched GO terms following ELS

Terms related to mitochondria in bold

| P21 Male ELS vs Male control | | |
|---|---|---|
| **Go term** | **Benjamini-corrected P-value** | **# Proteins** |
| **Total proteins - 0 terms** | | |
| **Upregulated proteins - 0 terms** | | |
| **Downregulated proteins - 0 terms** | | |

| P21 Female ELS vs Female control | | |
|---|---|---|
| **Go term** | **Benjamini-corrected P-value** | **# Proteins** |
| **Total proteins - 5 terms** | | |
| **Mitochondrion** | 2.5E-05 | 20 |
| Extracellular exosome | 6.3E-04 | 22 |
| Prefoldin complex | 6.0E-03 | 3 |
| Axon | 1.9E-02 | 7 |
| Cytosol | 3.5E-02 | 14 |
| **Upregulated proteins - 4 terms** | | |
| **Mitochondrion** | 1.9E-04 | 18 |
| Extracellular exosome | 1.6E-04 | 22 |
| Prefoldin complex | 5.0E-03 | 3 |
| Cytosol | 2.1E-02 | 14 |
| **Downregulated proteins - 0 terms** | | |

| Adult Male ELS vs Male control | | |
|---|---|---|
| Go term | Benjamini-corrected P-value | # Proteins |
| **Total proteins - 2 terms** | | |
| Myelin sheath | 2.0E-03 | 5 |
| Extracellular exosome | 2.0E-03 | 11 |
| **Upregulated proteins - 0 terms** | | |
| **Downregulated proteins - 1 terms** | | |
| Extracellular exosome | 5.2E-04 | 9 |

| Adult Female ELS vs Female control | | |
|---|---|---|
| Go term | Benjamini-corrected P-value | # Proteins |
| **Total proteins - 41 terms** | | |
| Extracellular exosome | 2.1E-23 | 93 |
| Myelin sheath | 7.1E-08 | 17 |
| Cytosol | 3.6E-07 | 49 |
| Membrane | 5.2E-07 | 119 |
| Synapse | 5.4E-07 | 24 |
| Neuron projection | 2.2E-06 | 21 |
| **Mitochondrial inner membrane** | 2.6E-06 | 20 |
| **Mitochondrion** | 3.6E-06 | 45 |
| Synaptic vesicle | 5.2E-06 | 12 |
| Clathrin-coated pit | 9.3E-06 | 9 |
| Dendrite | 1.5E-05 | 21 |
| Axon | 2.1E-05 | 18 |
| Cell-cell adherens junction | 5.9E-05 | 16 |
| Terminal bouton | 1.0E-04 | 10 |
| Postynaptic density | 3.1E-04 | 13 |
| Focal adhesion | 6.1E-04 | 16 |
| **Respiratory chain** | 8.1E-04 | 7 |
| Clarin coat of coated pit | 1.3E-03 | 4 |
| Vesicle | 1.5E-03 | 10 |
| Neuronal cell body | 1.5E-03 | 18 |
| Cytoplasmic vesicle | 1.6E-03 | 20 |
| **Mitochondrial respiratory chain complex I** | 2.5E-03 | 6 |
| Protein complex | 3.1E-03 | 19 |
| Cytoskeleton | 3.3E-03 | 27 |
| Perinuclear region of cytoplasm | 3.2E-03 | 20 |
| Membrane coat | 4.0E-03 | 5 |
| Clathrin-coated vesicle | 5.9E-03 | 6 |
| Clathrin coat of trans-Golgi network vesicle | 8.6E-03 | 3 |
| Dendritic spine | 1.5E-02 | 8 |
| Presynaptic membrane | 1.6E-02 | 6 |
| Secretory granule | 1.6E-02 | 7 |
| Cell-cell junction | 1.8E-02 | 9 |
| SNARE complex | 2.1E-02 | 5 |
| Trans-Golgi network | 2.3E-02 | 8 |
| Clarthrin coat | 2.4E-02 | 3 |
| Dendrite membrane | 2.7E-02 | 4 |
| Midbody | 2.9E-02 | 7 |
| Centrosome | 2.9E-02 | 13 |
| Smooth endoplasmic reticulum | 3.4E-02 | 4 |
| Dendritic spine head | 3.5E-02 | 3 |
| Presynaspe | 4.2E-02 | 5 |
| **Upregulated proteins - 25 terms** | | |
| Membrane | 4.90E-08 | 61 |
| Extracellular exosome | 7.2E-08 | 36 |
| Myelin sheath | 4.4E-07 | 11 |
| **Mitochondrion** | 7.3E-07 | 27 |
| Synaptic vesicle | 3.7E-06 | 9 |
| **Mitochondrial inner membrane** | 4.7E-06 | 13 |
| Synapse | 9.8E-06 | 14 |
| Protein complex | 1.6E-05 | 15 |
| Clathrin-coated pit | 1.6E-04 | 6 |
| Postynaptic density | 2.3E-04 | 9 |
| Dendrite | 1.2E-05 | 11 |
| Plasma membrane | 3.7E-03 | 38 |
| Membrane coat | 5.0E-03 | 4 |
| Presynaptic membrane | 5.1E-03 | 5 |
| Cytoskeleton | 5.0E-03 | 15 |
| Cytoplasmic vesicle | 7.3E-03 | 11 |
| Heterotrimeric G-protein complex | 7.8E-03 | 4 |
| Secretory granule | 1.8E-02 | 5 |
| Terminal bouton | 1.7E-02 | 5 |
| Focal adhesion | 1.8E-02 | 8 |
| Clathrin-coated vesicle | 2.3E-02 | 4 |
| Neuronal cell body | 2.4E-02 | 9 |
| Neuron projection | 2.3E-02 | 8- |
| Cytosol | 3.9E-02 | 17 |
| Axon | 4.7E-02 | 7 |
| **Downregulated proteins - 6 terms** | | |
| Extracellular exosome | 1.4E-14 | 57 |
| Cytosol | 2.7E-05 | 32 |
| Cell-cell adherens junction | 4.9E-04 | 12 |
| Neuron projection | 1.1E-03 | 13 |
| Axon | 7.1E-03 | 11 |
| Vesicle | 2.8E-02 | 7 |

**Table S9. Differentially expressed mitochondrial proteins at P21 following ELS**

[a]detected in the same comparison in adults, but opposite direction; [b]altered in males and females in response to ELS, but in opposite direction

| Male ELS vs Male control - 2 proteins | | |
|---|---|---|
| **Protein Name** | **Uniprot ID** | **Function** |
| **Upregulated in stress - 1 protein** | | |
| Arginine and glutamate-rich protein 1 | Q3UL36 | Proposed to be a modulator of glucocorticoid signaling in CNS |
| **Downregulated in stress - 1 protein** | | |
| [b]Cytochrome c, somatic | P62897 | Electron carrier protein in mitochondrial electron transport chain; plays a role in apoptosis |

| Female ELS vs Female control - 20 proteins | |
|---|---|
| **Protein Name** | **Uniprot ID** |
| **Upregulated in stress - 18 proteins** | |
| Acetyl-CoA acetyltransferase 2 | Q8CAY6 |
| Beta-synuclein | Q91ZZ3 |
| Coiled-coil domain-containing protein 58 | Q8R3Q6 |
| [b]Cytochrome c, somatic | P62897 |
| Diablo homolog, mitochondrial | Q9JIQ3 |
| ES1 protein homolog, mitochondrial | Q9D172 |
| Hydroxyacylglutathione hydrolase, mitochondrial | Q99KB8 |
| Inorganic pyrophosphatase 2, mitochondrial | Q91VM9 |
| [a]Malate dehydrogenase, mitochondrial | P08249 |
| [a]Mitochondrial import inner membrane translocase subunit Tim10 | P62073 |
| Mitochondrial import inner membrane translocase subunit Tim8 A | Q9WVA2 |
| [a]Nitrilase homolog 1 | Q8VDK1 |
| Peptidyl-prolyl cis-trans isomerase F, mitochondrial | Q99KR7 |
| Prefoldin subunit 2 | O70591 |
| Proline synthase co-transcribed bacterial homolog protein | Q9Z2Y8 |
| Ribonuclease UK114 | P52760 |
| [a]Ribosome-recycling factor, mitochondrial | Q9D6S7 |
| Thioredoxin reductase 2, mitochondrial | Q9JLT4 |
| **Downregulated in stress - 2 proteins** | |
| Carnitine O-palmitoyltransferase 1, brain isoform | Q8BGD5 |
| Phosphatidate cytidylyltransferase 2 | Q99L43 |

| Function |
|---|
| Cholesterol metabolism |
| Primarily expressed in brain, with enrichment in presynaptic terminals; some evidence that it inhibits alpha-synuclein aggregation |
| Unknown |
| Electron carrier protein in mitochondrial electron transport chain; plays a role in apoptosis |
| Promotes apoptosis |
| [illegible] enlargement in cone cells and protein is increased in Down Syndrome brains |
| Member of glyoxalase system with broad substrate specificty for glutathione thiol esters - prevents accumulation of methylglyoxal |
| Hyrolyzes inorganic pyrophosphate, essential for regulation of mitochondrial membrane potential |
| Key enzyme in TCA cycle, catalyzing the interconversion of malate and oxaloacetate |
| Mitochondrial intermembrane chaperone (component of TIMM9-TIMM10 complex) that facilitates transport of proteins across the intermembrane space and insertion into inner mitochondrial membrane |
| Mitochondrial intermembrane chaperone (component of TIMM8-TIMM13 complex) that facilitates transport of proteins across the intermembrane space and insertion into inner mitochondrial membrane |
| Amidase which is highly active toward deaminated glutathione |
| Also known as cyclophilin D - an essential component of the mitochondrial permeability transition pore |
| Subunit of the molecular chaperone prefoldin that facilitates proper protein folding |
| Involved in homeostatic regulation of intracellular pyridoxal 5'-phosphate, the active form of vitamin $B_6$ |
| Hydrolyzes enamine/imine intermediates generated during normal metabolism |
| Component of mitochondrial translation machinery that mediates disassembly of ribosomes from mRNA |
| Plays key role in redox homeostasis - maintains thioredoxin in a reduced state |
| Regulates beta-oxidation and transport of long-chain fatty acids into mitochondria |
| In mitochondrion, thought to be involved in synthesis of phosphatidylglycerol and cardiolipin |

**Table S10. Differentially expressed mitochondrial proteins in adults following ELS**

[a]detected in the same comparison at P21, but opposite direction; [b]altered in males and females in response to ELS

| Male ELS vs Male control - 7 proteins | | |
|---|---|---|
| **Protein Name** | **Uniprot ID** | **Function** |
| **Upregulated in stress - 3 proteins** | | |
| [b]Cytochrome c oxidase subunit 2 | P00405 | Subunit of complex IV in the respiratory chain - this subunit transfers the electrons from cytochrome c |
| Neurofilament heavy polypeptide | P19246 | Identified in a proteomic survey of mitochondria from mouse tissues |
| Superoxide dismutase [Cu-Zn] | P08228 | Destroys free superoxide radicals in cells |
| **Downregulated in stress - 4 proteins** | | |
| Hydroxyacyl-coenzyme A dehydrogenase, mitochondrial | Q61425 | Plays essential role in mitochondrial beta-oxidation of short chain fatty acids |
| Phosphatidylethanolamine-binding protein 1 | P70296 | Modulator of multiple signaling pathways |
| Sorbitol dehydrogenase | Q64442 | Key enzyme in the sorbitol pathway that intraconvers glucose and fructose |
| Threonine synthase-like 1 | Q8BH55 | Predicted PLP-dependent enzyme |

| Female ELS vs Female control - 45 proteins | |
|---|---|
| **Protein Name** | **Uniprot ID** |
| **Upregulated in stress - 27 proteins** | |
| Aconitate hydratase, mitochondrial | Q99KI0 |
| ADP/ATP translocase 1 | P48962 |
| AGF3-like protein 2 | Q8JZQ2 |
| AP-2 complex subunit mu | P84091 |
| ATP synthase subunit b, mitochondrial | Q9CQQ7 |
| ATP synthase subunit O, mitochondrial | Q9DB20 |
| Calcium uniporter protein, mitochondrial | Q3UMR5 |
| Calcium/calmodulin-dependent protein kinase type II subunit alpha | P11798 |
| Citrate synthase, mitochondrial | Q9CZU6 |
| Clathrin heavy chain 1 | Q61548 |
| [b]Cytochrome c oxidase subunit 2 | P00405 |
| Cytochrome c oxidase subunit 6C | Q9CPQ1 |
| Dynamin-3 | Q8BZ98 |
| Huntingtin | P42859 |
| Lipoamide acyltransferase component of branched-chain alpha-ket | P53395 |
| Methionine--tRNA ligase, cytoplasmic | Q68FL6 |
| Mitochondrial glutamate carrier 2 | Q9DB41 |
| Mitochondrial import receptor subunit TOM70 | Q9CZW5 |
| Mitogen-activated protein kinase 1 | P63085 |
| NADH-cytochrome b5 reductase 3 | Q9DCN2 |
| NADH dehydrogenase [ubiquinone] 1 alpha subcomplex subunit 12 | Q7TMF3 |
| NADH dehydrogenase [ubiquinone] 1 alpha subcomplex subunit 9, | Q9DC69 |
| Phosphate carrier protein, mitochondrial | Q8VEM8 |
| Succinyl-CoA ligase [ADP-forming] subunit beta, mitochondrial | Q9Z2I9 |
| Syntaxin-binding protein 1 | O08599 |
| Trifunctional enzyme subunit alpha, mitochondrial | Q8BMS1 |
| Uncharacterized protein KIAA0564 homolog | Q8CC88 |
| **Downregulated in stress - 18 proteins** | |
| Acyl-CoA-binding protein | P31786 |
| Cytochrome b-c1 complex subunit 6, mitochondrial | P99028 |
| Cytochrome c oxidase subunit 6B1 | P56391 |
| Dihydrolipoyl dehydrogenase, mitochondrial | O08749 |
| Dihydropteridine reductase | Q8BVI4 |
| [a]Malate dehydrogenase, mitochondrial | P08249 |
| [a]Mitochondrial import inner membrane translocase subunit Tim10 | P62073 |
| NADH dehydrogenase [ubiquinone] 1 alpha subcomplex subunit 2 | Q9CQ75 |
| NADH dehydrogenase [ubiquinone] iron-sulfur protein 4, mitochond | Q9CXZ1 |
| NADH dehydrogenase [ubiquinone] flavoprotein 2, mitochondrial | Q9D6J6 |
| [a]Nitrilase homolog 1 | Q8VDK1 |
| Nucleolar protein 3 | Q9D1X0 |
| Peptidyl-prolyl cis-trans isomerase NIMA-interacting 1 | Q9QUR7 |
| Peroxiredoxin-6 | O08709 |
| Protein DJ-1 | Q99LX0 |
| [a]Ribosome-recycling factor, mitochondrial | Q9D6S7 |
| Thioredoxin | P10639 |
| Uncharacterized protein C6orf203 homolog | Q9CQF4 |

| Function |
| --- |
| Enzyme in TCA cycle, catalyzing the isomerization of citrate to isocitrate |
| Catalyzes exchange of cytoplasmic ADP with mitochondrial ATP across inner mitochondrial membrane |
| A subunit of the mitochondrion m-AAA protease |
| Subunit of the adaptor protein complex 2 |
| Subunit of ATP synthase (complex V) |
| Subunit of ATP synthase (complex V) |
| Mitochondrial inner membrane calcium uniporter - mediates calcium uptake into mitochondria |
| Subunit of CaMKII - a multifunctional serine/threonine kinase |
| Enzyme in TCA cycle, synthesizing isocitrate from oxaloacetate |
| Component of the adaptor complexes - these link clatherin to recepotrs in coated vesicles |
| Subunit of complex IV in the respiratory chain - this subunit transfers the electrons from cytochrome c |
| Subunit of complex IV in the respiratory chain |
| Associated with microtubules and involved in vesicular transport |
| Primarily associated with vesicles and microtubules - may have a role in transport of mitochondria |
| Subunit of the branched chain alpha-ketoacid dehydrogenase complex - this complex converts alpha-keto acids to acyl-CoA |
| Catalyzes the attachment of an amino acid to its cognate tRNA |
| Co-transports glutamate and $H^+$ across the inner mitochondrial membrane |
| Subunit of outer mitochondrial membrane translocase - plays major role in import of mitochondrial precursor proteins |
| Serine/threonine kinase |
| Functions in the desaturation and elongation of fatty acids, cholesterol biosynthesis and drug metabolism |
| Accessory subunit of complex I of respiratory chain |
| Accessory subunit of complex I of respiratory chain |
| Catalyzes transport of phosphate into mitochondrial matrix |
| Subunit of enzyme in TCA cycle (ATP-specific succinyl-CoA synthetase) that hydrolyzes ATP to convert succinate to succinyl-CoA |
| Plays role in release of neurotransmitters via regulation of syntaxin |
| Subunit of the trifuncional protein that catalyzes the last steps of beta-oxidation of long chain fatty acids |
| Also known as von Willebrand factor A domain-containing protein 8 - unknown function |
| Lipid metabolism and modulation of $GABA_A$ receptor signaling |
| Subunit of complex III of respiratory chain - this subunit modulates cytochrome c1 and cytochdrome c complex formation |
| Subunit of complex IV of respiratory chain |
| $NAD^+$-dependent oxidoreductase - component of three mitochondrial alpha keto acid dehydrogenase complexes |
| Enzyme that catalyzes NADH-mediated reduction of quinonoid dihydrobiopterin - results in regeneration of tetrhydrobiopterin, a cofactor for monoamine synthesis |
| Key enzyme in TCA cycle, catalyzing the interconversion of malate and oxaloacetate |
| Mitochondrial intermembrane chaperone (component of TIMM9-TIMM10 complex) that facilitates transport of proteins across the intermembrane space and insertion into inner mitochondrial membrane |
| Accessory subunit of complex I of respiratory chain |
| Accessory subunit of complex I of respiratory chain |
| Core subunit of complex I of respiratory chain |
| Amidase which is highly active toward deaminated glutathione |
| Anti-apoptotic function - downregulates activity of caspases 2 and 8 |
| Multifunctional protein - functions include mitochondrial-mediated apoptosis |
| Member of thiol-specific antioxidant family |
| Multifunctional protein - functions include antioxidant activity and maintenance of mitochondrial homeostasis |
| Component of mitochondrial translation machinery that mediates disassembly of ribosomes from mRNA |
| Involved in redox reactions |
| Unknown |

### Table S11. Enriched KEGG pathways following ELS

| P21 Male ELS vs Male control | | |
|---|---|---|
| Pathway | Benjamini-corrected P-value | # Proteins |
| **Total proteins - 0 pathways** | | |
| **Upregulated proteins - 0 pathways** | | |
| **Downregulated proteins - 0 pathways** | | |

| P21 Female ELS vs Female control | | |
|---|---|---|
| Pathway | Benjamini-corrected P-value | # Proteins |
| **Total proteins - 2 pathways** | | |
| Carbon metabolism | 1.2E-02 | 5 |
| Pyruvate metabolism | 1.3E-02 | 4 |
| **Upregulated proteins - 3 pathways** | | |
| Carbon metabolism | 4.7E-03 | 5 |
| Pyruvate metabolism | 6.5E-03 | 4 |
| Biosynthesis of antibiotics | 3.1E-02 | 5 |
| **Downregulated proteins - 0 pathways** | | |

| Adult Male ELS vs Male control | | |
|---|---|---|
| Pathway | Benjamini-corrected P-value | # Proteins |
| **Total proteins - 0 pathways** | | |
| **Upregulated proteins - 0 pathways** | | |
| **Downregulated proteins - 0 pathways** | | |

| Adult Female ELS vs Female control | | |
|---|---|---|
| Pathway | Benjamini-corrected P-value | # Proteins |
| **Total proteins - 40 pathways** | | |
| Huntington's disease | 6.2E-09 | 20 |
| Synaptic vesicle cycle | 4.4E-09 | 13 |
| Endocrine and other factor-regulated calcium reabsorption | 1.4E-07 | 11 |
| Parkinson's disease | 1.1E-07 | 16 |
| Alzheimer's disease | 9.7E-07 | 16 |
| Circadian entrainment | 3.8E-06 | 12 |
| Oxidative phosphorylation | 1.6E-05 | 13 |
| Adrenergic signaling in cardiomyocytes | 1.7E-05 | 13 |
| Oxytocin signaling pathway | 2.8E-05 | 13 |
| Long-term depression | 4.0E-05 | 9 |
| Glutamatergic synapse | 6.1E-04 | 10 |
| Melanogenesis | 1.2E-03 | 9 |
| Dopaminergic synapse | 1.7E-03 | 10 |
| Long-term potentiation | 4.5E-03 | 7 |
| cAMP signaling pathway | 6.1E-03 | 11 |
| cGMP-PKG signaling pathway | 6.0E-03 | 10 |
| Gastric acid secretion | 5.9E-03 | 7 |
| Pertussis | 6.5E-03 | 7 |
| Retrograde endocannabinoid signaling | 6.4E-03 | 8 |
| Cardiac muscle contraction | 7.2E-03 | 7 |
| Salivary secretion | 7.2E-03 | 7 |
| Rap1 signaling pathway | 8.3E-03 | 11 |
| Citrate cycle (TCA cycle) | 8.9E-03 | 5 |
| Cholinergic synapse | 9.1E-03 | 8 |
| Endocytosis | 9.4E-03 | 12 |
| Gap junction | 1.0E-02 | 7 |
| Non-alcholic fatty liver disease | 1.3E-02 | 9 |
| Axon guidance | 1.7E-02 | 8 |
| Amphetamine addiction | 1.7E-02 | 6 |
| Estrogen signaling pathway | 1.7E-02 | 7 |
| Vasopressin-regulated water reabsorption | 2.0E-02 | 5 |
| Renin secretion | 2.0E-02 | 6 |
| Bacterial invasion of epithelial ells | 2.9E-02 | 6 |
| Cocaine addiction | 2.9E-02 | 5 |
| Oocyte meiosis | 2.8E-02 | 7 |
| Carbon metabolism | 3.3E-02 | 7 |
| Glyoxylate and dicarboxyalte metabolism | 4.1E-02 | 4 |
| GABAergic synapse | 4.0E-02 | 6 |
| Sphigolipid signaling pathway | 4.1E-02 | 7 |
| Morphine addiction | 4.9E-02 | 6 |
| **Upregulated proteins - 35 pathways** | | |
| Circadian entrainment | 1.1E-07 | 11 |
| Long-term depression | 4.4E-07 | 9 |
| Endocrine and other factor-regulated calcium reabsorption | 2.4E-06 | 8 |
| Huntington's disease | 2.4E-06 | 12 |
| Synaptic vesicle cycle | 5.0E-06 | 8 |
| Parkinson's disease | 1.4E-05 | 10 |
| Oxytocin signaling pathway | 1.3E-05 | 10 |
| Glutamatergic synapse | 1.7E-05 | 9 |
| Melanogenesis | 7.0E-05 | 8 |
| Adrenergic signaling in cardiomyocytes | 6.7E-05 | 9 |
| Retrograde endocannabinoid signaling | 7.4E-05 | 8 |
| Cholinergic synapse | 1.3E-04 | 8 |
| Gap junction | 2.7E-04 | 7 |
| Dopaminergic synapse | 3.3E-04 | 8 |
| cAMP signaling pathway | 4.9E-04 | 9 |
| cGMP-PKG signaling pathway | 1.0E-03 | 8 |
| Gastric acid secretion | 1.1E-03 | 6 |
| Alzheimer's disease | 1.5E-03 | 8 |
| GABAergic synapse | 2.4E-03 | 6 |
| Oxidative phosphorylation | 2.6E-03 | 7 |

| | | |
|---|---|---|
| Morphine addiction | 2.9E-03 | 6 |
| Estrogen signaling pathway | 3.5E-03 | 6 |
| Rap1 signaling pathway | 3.7E-03 | 8 |
| Chagas disease | 4.1E-03 | 6 |
| Sphigolipid signaling pathway | 9.0E-03 | 6 |
| Axon guidance | 1.0E-02 | 6 |
| Platelet activation | 1.1E-02 | 6 |
| Serotonergic synapse | 1.1E-02 | 6 |
| Progesterone-mediated oocyte maturation | 1.5E-02 | 5 |
| Cocaine addiction | 2.3E-02 | 5 |
| Toxoplasmosis | 2.7E-02 | 5 |
| Regulation of liolysis in adipocytes | 3.2E-02 | 4 |
| Calcium signaling pathway | 3.4E-02 | 6 |
| Endocytosis | 3.7E-02 | 7 |
| Long-term potentiation | 4.4E-02 | 4 |
| **Downregulated proteins - 3 terms** | | |
| Alzheimer's disease | 3.5E-02 | 8 |
| Huntington's disease | 3.4E-02 | 8 |
| Synaptic vesicle cycle | 4.2E-02 | 5 |

**Table S12. P21 Complex I data and statistics**

**Complex I activity (mOD/min/mg protein)**

**Hippocampus**

| | Control | | ELS | |
|---|---|---|---|---|
| | Male | Female | Male | Female |
| **Experiment** | | | | |
| 1 | 6.69 | 5.28 | 7.99 | 7.74 |
| 2 | 6.49 | 7.92 | 8.98 | 13.39 |
| 3 | 6.39 | 5.04 | 6.68 | 7.98 |
| 4 | 6.65 | 7.44 | 7.30 | 11.19 |
| 5 | 4.96 | 5.66 | 7.11 | 6.30 |
| 6 | 5.03 | 6.05 | 7.92 | 10.63 |
| 7 | 3.24 | 4.17 | 5.07 | 5.74 |
| 8 | 6.05 | 5.95 | 7.78 | 8.68 |
| 9 | 6.76 | 5.54 | 7.68 | 8.38 |
| 10 | 3.27 | 4.60 | 4.53 | 5.64 |
| 11 | 2.94 | 3.44 | 6.21 | 5.08 |
| 12 | 4.13 | 4.49 | 5.77 | 10.09 |
| **Mean** | **5.22** | **5.47** | **6.92** | **8.40** |
| **Standard deviation** | **1.49** | **1.29** | **1.31** | **2.55** |
| **95% CI** | **4.27, 6.17** | **4.65, 6.29** | **6.09, 7.75** | **6.78, 10.02** |

**Two-way ANOVA (sex and rearing between subjects factors)**

| | DF | SS | MS | F-value | [a]p-value | $h^2$ | [b]$w^2$ |
|---|---|---|---|---|---|---|---|
| **Sex** | 1 | 9.013 | 9.013 | 2.983 | 0.091 | 0.043 | **0.028** |
| **Rearing** | 1 | 64.589 | 64.589 | 21.378 | **<0.001** | 0.306 | **0.288** |
| **Sex*Rearing** | 1 | 4.588 | 4.588 | 1.519 | 0.224 | 0.022 | 0.007 |
| **Residual** | 44 | 132.936 | 3.021 | | | | |

[a]values in red = significant
[b]values in red = large effect size; values in green = small effect size

### Liver

| | Control | | ELS | |
|---|---|---|---|---|
| | Male | Female | Male | Female |
| **Experiment** | | | | |
| 1 | 6.26 | 4.88 | 5.88 | 6.03 |
| 2 | 5.32 | 5.23 | 6.24 | 8.31 |
| 3 | 5.89 | 4.96 | 6.88 | 8.29 |
| 4 | 5.69 | 5.79 | 7.16 | 8.28 |
| 5 | 4.97 | 5.33 | 6.20 | 7.36 |
| 6 | 5.80 | 7.45 | 6.53 | 9.74 |
| 7 | 4.84 | 5.04 | 3.34 | 3.46 |
| 8 | 6.46 | 6.24 | 6.32 | 7.69 |
| 9 | 3.88 | 4.58 | 4.17 | 8.40 |
| 10 | 6.99 | 4.12 | 5.79 | 5.08 |
| 11 | 5.04 | 5.76 | 7.54 | 9.21 |
| 12 | 5.24 | 5.21 | 8.04 | 7.99 |
| **Mean** | **5.53** | **5.38** | **6.17** | **7.49** |
| **Standard deviation** | **0.83** | **0.86** | **1.32** | **1.79** |
| **95% CI** | **5.00, 6.06** | **4.83, 5.93** | **5.33, 7.01** | **6.35, 8.63** |

### Two-way ANOVA (sex and rearing between subjects factors)

| | DF | SS | MS | F-value | [a]p-value | $h^2$ | [b]$w^2$ |
|---|---|---|---|---|---|---|---|
| **Sex** | 1 | 4.060 | 4.060 | 2.540 | 0.118 | 0.039 | **0.023** |
| **Rearing** | 1 | 22.633 | 22.633 | 14.161 | **<0.001** | 0.219 | **0.200** |
| **Sex*Rearing** | 1 | 6.409 | 6.409 | 4.010 | 0.051 | 0.062 | **0.046** |
| **Residual** | 44 | 70.324 | 1.598 | | | | |

[a]values in red = significant
[b]values in red = large effect size; values in green = small effect size

### Table S13. Adult Complex I data and statistics

### Complex I activity (mOD/min/mg protein)

### Hippocampus

| | Control | | ELS | |
|---|---|---|---|---|
| | Male | Female | Male | Female |
| **Experiment** | | | | |
| 1 | 6.96 | 7.73 | 7.79 | 3.86 |
| 2 | 5.79 | 5.73 | 4.64 | 3.64 |
| 3 | 6.40 | 6.92 | 6.29 | 5.44 |
| 4 | 5.77 | 5.79 | 5.56 | 4.18 |
| 5 | 5.28 | 5.88 | 5.68 | 4.65 |
| 6 | 6.40 | 10.30 | 3.74 | 4.80 |
| 7 | 6.58 | 8.98 | 4.63 | 5.36 |
| 8 | 9.44 | 7.44 | 6.82 | 3.62 |
| 9 | 5.61 | 8.10 | 4.29 | 4.03 |
| 10 | 8.53 | 9.49 | 5.54 | 4.45 |
| 11 | 5.47 | 8.47 | 3.76 | 3.82 |
| **Mean** | **6.57** | **7.71** | **5.34** | **4.35** |
| **Standard deviation** | **1.32** | **1.55** | **1.28** | **0.65** |
| **95% CI** | **5.68, 7.46** | **6.67, 8.75** | **4.48, 6.20** | **3.91, 4.79** |

### Two-way ANOVA (sex and rearing between subjects factors)

| | DF | SS | MS | F-value | [a]p-value | $h^2$ | [b]$w^2$ |
|---|---|---|---|---|---|---|---|
| **Sex** | 1 | 0.066 | 0.066 | 0.043 | 0.837 | <0.001 | -0.011 |
| **Rearing** | 1 | 57.891 | 57.891 | 37.436 | **<0.001** | 0.437 | **0.421** |
| **Sex*Rearing** | 1 | 12.540 | 12.540 | 8.109 | **0.007** | 0.095 | **0.082** |
| **Residual** | 40 | 61.857 | 1.546 | | | | |

[a]values in red = significant
[b]values in red = large effect size; values in blue = medium effect size

### Bonferroni/Dunn post-hoc analyses

**Sex*rearing**

| | Mean difference | Critical difference | [a]p-value | Cohen's *d* | Hedge's *g* |
|---|---|---|---|---|---|
| MC, MELS | 1.23 | 1.47 | **0.026** | 0.943 | 0.907 |
| FC, FELS | 3.36 | 1.47 | **<0.001** | 2.838 | 2.730 |
| MC, FC | 1.15 | 1.47 | **0.037** | 0.798 | 0.768 |
| MELS, FELS | 0.99 | 1.47 | 0.069 | 0.974 | 0.937 |
| FC, MELS | 2.37 | 1.47 | **<0.001** | 1.670 | 1.607 |
| MC, FELS | 2.22 | 1.47 | **<0.001** | 2.135 | 2.054 |

[a]values in red = significant; values in blue = not significant when corrected for multiple comparisons

### Liver

| | Control | | ELS | |
|---|---|---|---|---|
| | Male | Female | Male | Female |
| Experiment | | | | |
| 1 | 6.73 | 7.80 | 7.56 | 7.61 |
| 2 | 9.06 | 7.57 | 8.15 | 7.32 |
| 3 | 8.66 | 6.43 | 7.02 | 6.61 |
| 4 | 5.25 | 6.55 | 9.02 | 5.67 |
| 5 | 5.42 | 5.63 | 6.09 | 6.09 |
| 6 | 8.30 | 8.50 | 6.08 | 10.02 |
| 7 | 9.06 | 12.46 | 7.58 | 11.21 |
| 8 | 9.87 | 7.99 | 6.08 | 4.99 |
| 9 | 6.82 | 6.18 | 5.02 | 8.05 |
| 10 | 5.60 | 5.69 | 10.90 | 7.83 |
| 11 | 5.79 | 6.85 | 6.37 | 5.01 |
| **Mean** | **7.32** | **7.42** | **7.26** | **7.31** |
| **Standard deviation** | **1.71** | **1.92** | **1.65** | **1.97** |
| **95% CI** | **6.17, 8.47** | **6.13, 8.71** | **6.15, 8.37** | **5.99, 8.63** |

### Two-way ANOVA (sex and rearing between subjects factors)

| | DF | SS | MS | F-value | p-Value | $h^2$ | $w^2$ |
|---|---|---|---|---|---|---|---|
| **Sex** | 1 | 0.060 | 0.060 | 0.018 | 0.893 | <0.001 | -0.024 |
| **Rearing** | 1 | 0.085 | 0.085 | 0.026 | 0.874 | <0.001 | -0.024 |
| **Sex*Rearing** | 1 | 0.007 | 0.007 | 0.002 | 0.964 | <0.001 | -0.024 |
| **Residual** | 40 | 132.117 | 3.303 | | | | |

### Table S14. P21 respiration data and statistics

### Oxygen consumption rate (pmol/min/mg protein)

### Hippocampus

**Basal**

| | Control | | ELS | |
|---|---|---|---|---|
| | Male | Female | Male | Female |
| **Experiment** | | | | |
| 1 | ND | ND | ND | ND |
| 2 | ND | ND | ND | ND |
| 3 | 54.36 | 55.10 | 78.77 | 45.07 |
| 4 | 73.88 | 75.21 | 68.26 | 62.14 |
| 5 | 41.73 | 37.43 | 37.99 | 36.93 |
| 6 | 68.16 | 66.09 | 62.48 | 58.59 |
| 7 | 48.48 | 29.86 | 57.77 | 56.76 |
| 8 | 57.89 | 66.47 | 82.45 | 65.43 |
| 9 | 63.33 | 55.27 | 60.81 | 66.42 |
| **Mean** | **58.26** | **55.06** | **64.08** | **55.91** |
| **Standard deviation** | **11.19** | **16.35** | **14.74** | **11.00** |
| **95% CI** | **47.91, 68.61** | **39.94, 70.18** | **50.45, 77.71** | **45.74, 66.08** |

### Two-way ANOVA (sex and rearing between subjects factors)

| | DF | SS | MS | F-value | p-value | $h^2$ | $w^2$ |
|---|---|---|---|---|---|---|---|
| **Sex** | 1 | 226.235 | 226.235 | 1.238 | 0.277 | 0.048 | 0.009 |
| **Rearing** | 1 | 77.589 | 77.589 | 0.425 | 0.521 | 0.017 | -0.021 |
| **Sex*Rearing** | 1 | 43.227 | 43.227 | 0.237 | 0.631 | 0.009 | -0.028 |
| **Residual** | 24 | 4385.746 | 182.739 | | | | |

**ADP**

| | Control | | ELS | |
|---|---|---|---|---|
| | Male | Female | Male | Female |
| **Experiment** | | | | |
| 1 | 118.82 | 110.76 | 175.90 | 159.72 |
| 2 | 140.13 | 163.97 | 185.20 | 195.58 |
| 3 | 206.60 | 168.86 | 322.90 | 234.28 |
| 4 | 206.85 | 165.46 | 252.57 | 260.98 |
| 5 | 98.12 | 70.56 | 146.11 | 151.80 |
| 6 | 139.36 | 124.14 | 166.93 | 156.05 |
| 7 | 93.77 | 68.59 | 122.08 | 139.45 |
| 8 | 136.64 | 150.30 | 195.45 | 214.54 |
| 9 | 106.10 | 100.02 | 119.49 | 136.61 |
| **Mean** | **138.49** | **124.74** | **187.40** | **183.22** |
| **Standard deviation** | **42.40** | **39.83** | **65.01** | **44.94** |
| **95% CI** | **103.04, 173.94** | **91.44, 158.04** | **133.05, 241.75** | **145.65, 220.79** |

### Two-way ANOVA (sex and rearing between subjects factors)

| | DF | SS | MS | F-value | [a]p-value | $h^2$ | [b]$w^2$ |
|---|---|---|---|---|---|---|---|
| **Sex** | 1 | 723.162 | 723.162 | 0.300 | 0.588 | 0.007 | -0.016 |
| **Rearing** | 1 | 25952.673 | 25952.673 | 10.780 | **0.003** | 0.250 | **0.221** |
| **Sex*Rearing** | 1 | 205.970 | 205.970 | 0.086 | 0.772 | 0.002 | -0.021 |
| **Residual** | 32 | 77041.800 | 2407.556 | | | | |

[a]values in red = significant
[b]values in red = large effect size

**Olig**

| | Control | | ELS | |
|---|---|---|---|---|
| | **Male** | **Female** | **Male** | **Female** |
| **Experiment** | | | | |
| 1 | 23.10 | 29.60 | 38.93 | 29.62 |
| 2 | 25.06 | 31.23 | 34.44 | 34.38 |
| 3 | 31.91 | 29.06 | 39.99 | 29.62 |
| 4 | 33.80 | 29.46 | 52.84 | 33.11 |
| 5 | 33.53 | 24.53 | 32.56 | 36.14 |
| 6 | 37.89 | 31.09 | 37.44 | 40.79 |
| 7 | 20.10 | 13.38 | 24.34 | 34.10 |
| 8 | 28.07 | 33.20 | 30.65 | 27.15 |
| 9 | 15.07 | 21.64 | 22.02 | 30.91 |
| **Mean** | **27.61** | **27.02** | **34.80** | **32.87** |
| **Standard deviation** | **7.40** | **6.23** | **9.16** | **4.11** |
| **95% CI** | **21.42, 33.80** | **21.81, 32.23** | **27.14, 42.46** | **29.43, 36.31** |

### Two-way ANOVA (sex and rearing between subjects factors)

| | DF | SS | MS | F-value | [a]p-value | $h^2$ | [b]$w^2$ |
|---|---|---|---|---|---|---|---|
| **Sex** | 1 | 14.351 | 14.351 | 0.295 | 0.591 | 0.007 | -0.017 |
| **Rearing** | 1 | 382.268 | 382.268 | 7.866 | **0.009** | 0.195 | **0.166** |
| **Sex*Rearing** | 1 | 4.033 | 4.033 | 0.083 | 0.775 | 0.002 | -0.022 |
| **Residual** | 32 | 1555.152 | 48.599 | | | | |

[a]values in red = significant
[b]values in red = large effect size

**FCCP**

| | Control | | ELS | |
|---|---|---|---|---|
| | **Male** | **Female** | **Male** | **Female** |
| **Experiment** | | | | |
| 1 | 95.71 | 113.85 | 166.85 | 149.07 |
| 2 | 130.73 | 151.60 | 143.53 | 186.61 |
| 3 | 184.11 | 156.19 | 297.70 | 188.64 |
| 4 | 192.47 | 168.71 | 262.92 | 254.64 |
| 5 | 96.17 | 72.05 | 137.65 | 147.98 |
| 6 | 124.65 | 124.65 | 137.83 | 160.74 |
| 7 | 88.07 | 51.87 | 109.37 | 128.74 |
| 8 | 92.63 | 153.67 | 171.65 | 186.52 |
| 9 | 93.42 | 84.73 | 92.38 | 127.13 |
| **Mean** | **122.00** | **119.70** | **168.88** | **170.01** |
| **Standard deviation** | **40.46** | **41.92** | **68.40** | **39.70** |
| **95% CI** | **88.17, 155.83** | **84.65, 154.75** | **111.70, 226.06** | **136.82, 203.20** |

### Two-way ANOVA (sex and rearing between subjects factors)

| | DF | SS | MS | F-value | [a]p-value | $h^2$ | [b]$w^2$ |
|---|---|---|---|---|---|---|---|
| **Sex** | 1 | 3.192 | 3.192 | 0.001 | 0.972 | <.001 | -0.024 |
| **Rearing** | 1 | 20277.760 | 20277.760 | 8.262 | **0.007** | 0.205 | **0.176** |
| **Sex*Rearing** | 1 | 3.098 | 3.098 | 0.001 | 0.972 | <0.001 | -0.024 |
| **Residual** | 32 | 78541.940 | 2454.436 | | | | |

**AA**

| | Control | | ELS | |
|---|---|---|---|---|
| | Male | Female | Male | Female |
| Experiment | | | | |
| 1 | 0.03 | 5.65 | 7.35 | 1.81 |
| 2 | 12.52 | 16.80 | 12.89 | 14.33 |
| 3 | 8.57 | 7.96 | 9.04 | 7.73 |
| 4 | 2.29 | 8.17 | 9.50 | 4.16 |
| 5 | 11.14 | 7.18 | 0.96 | 0.89 |
| 6 | 14.79 | 14.79 | 6.69 | 7.59 |
| 7 | 16.28 | 4.69 | 10.87 | 13.61 |
| 8 | 10.51 | 15.55 | 13.32 | 10.91 |
| 9 | 6.27 | 5.76 | 7.08 | 10.31 |
| **Mean** | **9.16** | **9.62** | **8.63** | **7.93** |
| **Standard deviation** | **5.46** | **4.73** | **3.75** | **4.86** |
| **95% CI** | **4.60, 13.72** | **5.67, 13.57** | **5.49, 11.77** | **3.87, 11.99** |

### Two-way ANOVA (sex and rearing between subjects factors)

| | DF | SS | MS | F-value | p-value | $h^2$ | $w^2$ |
|---|---|---|---|---|---|---|---|
| **Sex** | 1 | 0.136 | 0.136 | 0.006 | 0.939 | <0.001 | -0.030 |
| **Rearing** | 1 | 11.011 | 11.011 | 0.490 | 0.489 | 0.015 | -0.015 |
| **Sex*Rearing** | 1 | 3.068 | 3.068 | 0.136 | 0.714 | 0.004 | -0.026 |
| **Residual** | 32 | 719.772 | 22.493 | | | | |

### Liver

#### Basal

| | Control | | ELS | |
|---|---|---|---|---|
| | **Male** | **Female** | **Male** | **Female** |
| **Experiment** | | | | |
| 1 | ND | ND | ND | ND |
| 2 | ND | ND | ND | ND |
| 3 | 16.84 | 15.13 | 25.36 | 14.98 |
| 4 | 13.97 | 14.34 | 15.96 | 13.58 |
| 5 | 17.1 | 10.54 | 17.57 | 11.4 |
| 6 | 26.95 | 21.21 | 30.14 | 15.83 |
| 7 | 9.85 | 13.94 | 17.21 | 17.32 |
| 8 | 17.31 | 20.93 | 33.74 | 22.99 |
| 9 | 21.48 | 17.32 | 32.08 | 30.36 |
| **Mean** | **17.64** | **16.20** | **24.58** | **18.07** |
| **Standard deviatior** | **5.43** | **3.88** | **7.63** | **6.52** |
| **95% CI** | **12.62, 22.66** | **12.61, 19.79** | **17.52, 31.64** | **12.04, 24.10** |

### Two-way ANOVA (sex and rearing between subjects factors)

| | DF | SS | MS | F-value | p-value | $h^2$ | [b]$w^2$ |
|---|---|---|---|---|---|---|---|
| **Sex** | 1 | 110.763 | 110.763 | 3.050 | 0.094 | 0.095 | **0.062** |
| **Rearing** | 1 | 135.464 | 135.464 | 3.733 | 0.065 | 0.117 | **0.083** |
| **Sex*Rearing** | 1 | 45.034 | 45.034 | 1.240 | 0.277 | 0.039 | 0.007 |
| **Residual** | 24 | 871.557 | 36.315 | | | | |

[b] values in blue = medium effect size

**ADP**

| | Control | | ELS | |
|---|---|---|---|---|
| | Male | Female | Male | Female |
| **Experiment** | | | | |
| 1 | ND | ND | ND | ND |
| 2 | 25.07 | 33.61 | 32.87 | 28.89 |
| 3 | 52.21 | 52.94 | 65.96 | 38.95 |
| 4 | 30.74 | 41.45 | 38.31 | 35.3 |
| 5 | 26.68 | 32.87 | 31.64 | 25.07 |
| 6 | 49.4 | 87.56 | 80.99 | 47.3 |
| 7 | 105.75 | 78.26 | 141.24 | 99.42 |
| 8 | 47.37 | 76.79 | 51.5 | 94.31 |
| 9 | 31.75 | 37.87 | 52.35 | 52.71 |
| **Mean** | **46.12** | **55.17** | **61.86** | **52.74** |
| **Standard deviation** | **26.37** | **22.37** | **36.23** | **28.70** |
| **95% CI** | **24.07, 68.17** | **34.47, 73.87** | **31.57, 92.15** | **28.75, 76.73** |

### Two-way ANOVA (sex and rearing between subjects factors)

| | DF | SS | MS | F-value | p-value | $h^2$ | $w^2$ |
|---|---|---|---|---|---|---|---|
| **Sex** | 1 | 9.538 | 9.538 | 0.011 | 0.916 | 0.000 | -0.032 |
| **Rearing** | 1 | 250.600 | 250.600 | 0.295 | 0.591 | 0.010 | -0.023 |
| **Sex*Rearing** | 1 | 822.455 | 822.455 | 0.969 | 0.333 | 0.033 | -0.001 |
| **Residual** | 28 | 23764.483 | 848.732 | | | | |

**Olig**

| | Control | | ELS | |
|---|---|---|---|---|
| | Male | Female | Male | Female |
| **Experiment** | | | | |
| 1 | ND | ND | ND | ND |
| 2 | 7.53 | 3.26 | 8.06 | 6.31 |
| 3 | 6.78 | 5.45 | 5.16 | 0.64 |
| 4 | 6.53 | 4 | 7.62 | 5.68 |
| 5 | 6.96 | 4.22 | 8.23 | 3.15 |
| 6 | 9.95 | 6.31 | 8.28 | 5.65 |
| 7 | 2.75 | 12.61 | 7.18 | 10.44 |
| 8 | 8.57 | 13.22 | 10.95 | 6.7 |
| 9 | 4.6 | 7.09 | 5.63 | 10.01 |
| **Mean** | **6.71** | **7.02** | **7.64** | **6.07** |
| **Standard deviation** | **2.23** | **3.85** | **1.78** | **3.24** |
| **95% CI** | **4.85, 8.57** | **3.80, 10.24** | **6.15, 9.13** | **3.36, 8.78** |

### Two-way ANOVA (sex and rearing between subjects factors)

| | DF | SS | MS | F-value | p-value | $h^2$ | $w^2$ |
|---|---|---|---|---|---|---|---|
| **Sex** | 1 | 42.828 | 42.828 | 0.562 | 0.460 | 0.018 | -0.014 |
| **Rearing** | 1 | 68.796 | 68.796 | 0.903 | 0.350 | 0.030 | -0.003 |
| **Sex*Rearing** | 1 | 32.080 | 32.080 | 0.421 | 0.522 | 0.014 | -0.019 |
| **Residual** | 28 | 2132.378 | 76.156 | | | | |

**FCCP**

| | Control | | ELS | |
|---|---|---|---|---|
| | **Male** | **Female** | **Male** | **Female** |
| **Experiment** | | | | |
| 1 | ND | ND | ND | ND |
| 2 | 26.84 | 32.42 | 35.81 | 30.44 |
| 3 | 45.31 | 35.07 | 53.62 | 31.26 |
| 4 | 25.15 | 44.35 | 35.06 | 35.01 |
| 5 | 29.12 | 33.04 | 31.12 | 20.84 |
| 6 | 56.07 | 104.35 | 102.94 | 45.15 |
| 7 | 127.81 | 75.52 | 126.53 | 124.2 |
| 8 | 41.02 | 84.8 | 55.05 | 75.83 |
| 9 | 26.87 | 37.15 | 47.5 | 50.15 |
| **Mean** | **47.27** | **55.84** | **60.95** | **51.61** |
| **Standard deviation** | **34.35** | **28.18** | **34.90** | **33.77** |
| **95% CI** | **18.55, 75.99** | **32.28, 79.40** | **31.77, 90.13** | **23.38, 79.84** |

### Two-way ANOVA (sex and rearing between subjects factors)

| | DF | SS | MS | F-value | p-value | $h^2$ | $w^2$ |
|---|---|---|---|---|---|---|---|
| **Sex** | 1 | 53.510 | 53.510 | 0.047 | 0.830 | 0.002 | -0.032 |
| **Rearing** | 1 | 51.207 | 51.207 | 0.045 | 0.834 | 0.002 | -0.032 |
| **Sex*Rearing** | 1 | 994.580 | 994.580 | 0.874 | 0.358 | 0.030 | -0.004 |
| **Residual** | 28 | 31863.945 | 1137.998 | | | | |

**AA**

| | Control | | ELS | |
|---|---|---|---|---|
| | **Male** | **Female** | **Male** | **Female** |
| **Experiment** | | | | |
| 1 | **AA** | ND | ND | ND |
| 2 | ND | 1.31 | 0.18 | 2.47 |
| 3 | 0.07 | 1.84 | 0.3 | 0.48 |
| 4 | 0.79 | 0.81 | 0.71 | 0.54 |
| 5 | 0.72 | 0.18 | 0.75 | 0.07 |
| 6 | 2.45 | 9.87 | 3.71 | 5.98 |
| 7 | 4.34 | 5.28 | 15.41 | 7.06 |
| 8 | 7.72 | 5.67 | 3.9 | 9.09 |
| 9 | 8.19 | 2.04 | 6.97 | 4.91 |
| | 2.85 | | | |
| **Mean** | | **3.38** | **3.99** | **3.83** |
| **Standard deviation** | **3.39** | **3.30** | **5.19** | **3.42** |
| **95% CI** | **3.13** | **0.62, 6.14** | **-0.35, 8.33** | **0.97, 6.69** |
| | **0.77, 6.01** | | | |

### Two-way ANOVA (sex and rearing between subjects factors)

| | DF | SS | MS | F-value | p-value | $h^2$ | $w^2$ |
|---|---|---|---|---|---|---|---|
| **Sex** | 1 | 57.005 | 57.005 | 0.690 | 0.413 | 0.023 | -0.010 |
| **Rearing** | 1 | 86.363 | 86.363 | 1.045 | 0.315 | 0.034 | 0.001 |
| **Sex*Rearing** | 1 | 57.701 | 57.701 | 0.698 | 0.410 | 0.023 | -0.010 |
| **Residual** | 28 | 2313.231 | 82.615 | | | | |

**Table S15. Adult respiration data and statistics**

**Oxygen consumption rate (pmol/min/mg protein)**

**Hippocampus**

**Basal**

| | Control | | ELS | |
|---|---|---|---|---|
| | Male | Female | Male | Female |
| Experiment | | | | |
| 1 | ND | ND | ND | ND |
| 2 | ND | ND | ND | ND |
| 3 | 89.85 | 84.95 | 78.16 | 55.92 |
| 4 | 84.08 | 87.02 | 64.96 | 71.12 |
| 5 | 43.84 | 58.12 | 43.68 | 55.38 |
| 6 | 39.47 | 74.11 | 34.62 | 32.22 |
| 7 | 49.22 | 62.79 | 41.07 | 50 |
| 8 | 69.07 | 55.5 | 66.21 | 63.55 |
| 9 | 60.31 | 61.85 | 77.27 | 43.83 |
| 10 | 80.7 | 53.96 | 62.4 | 81.53 |
| **Mean** | **64.57** | **67.29** | **58.55** | **56.69** |
| **Standard deviation** | **19.32** | **13.08** | **16.69** | **15.51** |
| **95% CI** | **48.42, 80.72** | **56.36, 78.22** | **44.60, 72.50** | **43.72, 69.66** |

**Two-way ANOVA (sex and rearing between subjects factors)**

| | DF | SS | MS | F-value | p-value | $h^2$ | $w^2$ |
|---|---|---|---|---|---|---|---|
| **Sex** | 1 | 1.505 | 1.505 | 0.006 | 0.941 | <0.001 | -0.032 |
| **Rearing** | 1 | 552.116 | 552.116 | 2.077 | 0.161 | 0.069 | **0.034** |
| **Sex*Rearing** | 1 | 41.816 | 41.816 | 0.157 | 0.695 | 0.005 | -0.027 |
| **Residual** | 28 | 7444.369 | 265.87 | | | | |

[b] values in green = small effect size

**ADP**

| | Control | | ELS | |
|---|---|---|---|---|
| | Male | Female | Male | Female |
| Experiment | | | | |
| 1 | 127.25 | 131.81 | 131.03 | 97.18 |
| 2 | 136.82 | 181.7 | 123.87 | 87.11 |
| 3 | 195.27 | 186.72 | 221.86 | 101.94 |
| 4 | 154.86 | 212.05 | 102.76 | 158.92 |
| 5 | 137.06 | 214.06 | 148.69 | 133.41 |
| 6 | 106.38 | 162.71 | 131.39 | 86.51 |
| 7 | 101.72 | 134.49 | 105.83 | 78.53 |
| 8 | 202 | 217.05 | 184.56 | 156.74 |
| 9 | 136.39 | 141.2 | 107.2 | 89.14 |
| 10 | 163.15 | 157.32 | 149.17 | 112.72 |
| **Mean** | **146.09** | **173.91** | **140.64** | **110.22** |
| **Standard deviation** | **33.46** | **33.25** | **37.84** | **29.56** |
| **95% CI** | **122.15, 170.03** | **150.12, 197.70** | **113.57, 167.71** | **89.07, 131.37** |

### Two-way ANOVA (sex and rearing between subjects factors)

| | DF | SS | MS | F-value | [a]p-value | $h^2$ | [b]$w^2$ |
|---|---|---|---|---|---|---|---|
| **Sex** | 1 | 16.835 | 16.835 | 0.015 | 0.904 | <0.001 | -0.018 |
| **Rearing** | 1 | 11952.578 | 11952.578 | 10.553 | **0.003** | 0.195 | **0.174** |
| **Sex*Rearing** | 1 | 8478.870 | 8478.870 | 7.486 | **0.009** | 0.138 | **0.118** |
| **Residual** | 36 | 40774.643 | 1132.629 | | | | |

[a]values in red = significant
[b]values in red = large effect size; values in blue = medium effect size

### Bonferroni/Dunn post-hoc analyses for significant effects

**Sex*rearing**

| | Mean difference | Critical difference | [a]p-value | Cohen's *d* | Hedge's *g* |
|---|---|---|---|---|---|
| FC, FELS | 63.691 | 42.021 | **<0.001** | 2.025 | 1.939 |
| FC, MC | 27.821 | 42.021 | 0.073 | 0.834 | 0.799 |
| FC, MELS | 33.275 | 42.021 | **0.034** | 0.934 | 0.895 |
| FELS, MC | 35.870 | 42.021 | **0.023** | 1.136 | 1.088 |
| FELS, MELS | 30.416 | 42.021 | 0.051 | 0.896 | 0.858 |
| MC, MELS | 5.454 | 42.021 | 0.712 | 0.153 | 0.146 |

[a]values in red = significant; values in blue = not significant when corrected for multiple comparisons

**Olig**

| | Control | | ELS | |
|---|---|---|---|---|
| | **Male** | **Female** | **Male** | **Female** |
| **Experiment** | | | | |
| 1 | 11.65 | 11.44 | 10.09 | 12.16 |
| 2 | 16.31 | 15.63 | 17.39 | 14.89 |
| 3 | 45.47 | 35.73 | 31.68 | 12.82 |
| 4 | 16.67 | 35.45 | 22.45 | 18.51 |
| 5 | 36.84 | 15.08 | 37 | 31.44 |
| 6 | 23.53 | 41.5 | 34.68 | 36.52 |
| 7 | 20.81 | 25.55 | 25.73 | 18.86 |
| 8 | 25.23 | 34.36 | 35.9 | 35.21 |
| 9 | 23.66 | 28.17 | 33.64 | 20.52 |
| 10 | 37.6 | 39.72 | 41.59 | 21.39 |
| **Mean** | **25.78** | **28.26** | **29.02** | **22.23** |
| **Standard deviation** | **10.84** | **10.93** | **9.87** | **9.01** |
| **95% CI** | **18.03, 33.53** | **20.44, 36.08** | **21.96, 36.08** | **15.79, 26.69** |

### Two-way ANOVA (sex and rearing between subjects factors)

| | DF | SS | MS | F-value | p-value | $h^2$ | $w^2$ |
|---|---|---|---|---|---|---|---|
| **Sex** | 1 | 46.161 | 46.161 | 0.338 | 0.566 | 0.009 | -0.017 |
| **Rearing** | 1 | 19.502 | 19.502 | 0.143 | 0.708 | 0.004 | -0.022 |
| **Sex*Rearing** | 1 | 214.786 | 214.786 | 1.572 | 0.218 | 0.041 | **0.015** |
| **Residual** | 36 | 4920.020 | 136.667 | | | | |

[b] values in green = small effect size

**FCCP**

| | Control | | ELS | |
|---|---|---|---|---|
| | Male | Female | Male | Female |
| **Experiment** | | | | |
| 1 | 205.98 | 143.7 | 197.27 | 86.54 |
| 2 | 146.45 | 236.75 | 113.03 | 127.12 |
| 3 | 221.67 | 149.6 | 186.25 | 61.88 |
| 4 | 142.87 | 175.08 | 132.11 | 145.03 |
| 5 | 114.64 | 194.32 | 128.38 | 101.46 |
| 6 | 104.85 | 132.34 | 128.19 | 88.4 |
| 7 | 104.75 | 131.65 | 100.77 | 72.3 |
| 8 | 153.54 | 183.68 | 143.44 | 132.37 |
| 9 | 94.83 | 129.9 | 97.4 | 78.89 |
| 10 | 136.73 | 136.96 | 127.38 | 104.24 |
| **Mean** | **142.63** | **161.40** | **135.42** | **99.82** |
| **Standard deviation** | **42.63** | **35.30** | **33.00** | **27.51** |
| **95% CI** | **112.13, 173.13** | **136.15, 186.65** | **111.81, 159.03** | **80.14, 119.50** |

### Two-way ANOVA (sex and rearing between subjects factors)

| | DF | SS | MS | F-value | [a]p-value | $h^2$ | [b]$w^2$ |
|---|---|---|---|---|---|---|---|
| **Sex** | 1 | 1303.251 | 1303.251 | 0.947 | 0.337 | 0.020 | -0.001 |
| **Rearing** | 1 | 9854.577 | 9854.577 | 7.160 | **0.011** | 0.148 | **0.125** |
| **Sex*Rearing** | 1 | 5848.174 | 5848.174 | 4.249 | **0.047** | 0.088 | **0.066** |
| **Residual** | 36 | 49550.740 | 1376.409 | | | | |

[a]values in red = significant
[b] values in blue = medium effect size

**AA**

| | Control | | ELS | |
|---|---|---|---|---|
| | **Male** | **Female** | **Male** | **Female** |
| **Experiment** | | | | |
| 1 | 9.73 | 8.61 | 8.6 | 8.99 |
| 2 | 1.66 | 1.58 | 1.16 | 3.24 |
| 3 | 23.15 | 19.7 | 16.31 | 16.53 |
| 4 | 14.51 | 22.63 | 14.74 | 13.56 |
| 5 | 15.1 | 7.98 | 8.6 | 12.13 |
| 6 | 15.13 | 18.61 | 25.85 | 14.65 |
| 7 | 3.43 | 7.48 | 1.81 | 5.15 |
| 8 | 9.23 | 9.36 | 6.76 | 9.71 |
| 9 | 9.08 | 11.22 | 11.93 | 6.16 |
| 10 | 15.13 | 16.87 | 16.87 | 12.2 |
| **Mean** | **11.62** | **12.40** | **11.26** | **10.23** |
| **Standard deviation** | **6.33** | **6.69** | **7.51** | **4.36** |
| **95% CI** | **7.09, 16.15** | **7.61, 17.19** | **5.89, 16.63** | **7.11, 13.35** |

### Two-way ANOVA (sex and rearing between subjects factors)

| | DF | SS | MS | F-value | p-value | $h^2$ | $^{b}w^2$ |
|---|---|---|---|---|---|---|---|
| **Sex** | 1 | 0.146 | 0.146 | 0.004 | 0.952 | <0.001 | -0.026 |
| **Rearing** | 1 | 15.926 | 15.926 | 0.398 | 0.532 | 0.011 | -0.016 |
| **Sex*Rearing** | 1 | 8.281 | 8.281 | 0.207 | 0.652 | 0.006 | -0.021 |
| **Residual** | 36 | 1441.319 | 40.037 | | | | |

[b] values in green = small effect size

### Liver

**Basal**

| | Control | | ELS | |
|---|---|---|---|---|
| | Male | Female | Male | Female |
| **Experiment** | | | | |
| 1 | ND | ND | ND | ND |
| 2 | ND | ND | ND | ND |
| 3 | 14.75 | 17.2 | 14.11 | 17.78 |
| 4 | 14.28 | 16.46 | 9.3 | 14.66 |
| 5 | 15.73 | 12.54 | 11.03 | 7.81 |
| 6 | 19.74 | 22.72 | 20.33 | 18.49 |
| 7 | 22.81 | 26.08 | 24.77 | 24.72 |
| 8 | 13.53 | 22.27 | 20.17 | 39.12 |
| 9 | 60.31 | 14.97 | 16.45 | 10.65 |
| 10 | 22.95 | 18.88 | 22.87 | 24.69 |
| **Mean** | **23.01** | **18.89** | **17.38** | **19.74** |
| **Standard deviation** | **15.53** | **4.51** | **5.59** | **9.86** |
| **95% CI** | **10.03, 35.99** | **15.12, 22.66** | **12.71, 22.05** | **11.50, 27.98** |

#### Two-way ANOVA (sex and rearing between subjects factors)

| | DF | SS | MS | F-value | p-value | $h^2$ | [b]$W^2$ |
|---|---|---|---|---|---|---|---|
| **Sex** | 1 | 150.338 | 150.338 | 1.591 | 0.218 | 0.053 | **0.019** |
| **Rearing** | 1 | 2.892 | 2.892 | 0.031 | 0.864 | 0.001 | -0.031 |
| **Sex*Rearing** | 1 | 33.456 | 33.456 | 0.354 | 0.557 | 0.012 | -0.021 |
| **Residual** | 28 | 2646.192 | 94.507 | | | | |

[b] values in green = small effect size

**ADP**

| | Control | | ELS | |
|---|---|---|---|---|
| | **Male** | **Female** | **Male** | **Female** |
| **Experiment** | | | | |
| 1 | 75.53 | 73.99 | 57.69 | 56.54 |
| 2 | ND | ND | ND | ND |
| 3 | 49.15 | 57.45 | 33.03 | 48.17 |
| 4 | 86.48 | 98.01 | 51.23 | 63.41 |
| 5 | 85.09 | 74.55 | 62.97 | 56.83 |
| 6 | 83.4 | 88.92 | 63.67 | 67.09 |
| 7 | 59.84 | 48.2 | 95.44 | 84.37 |
| 8 | 55.09 | 78.89 | 49.15 | 68.41 |
| 9 | 136.39 | 113.97 | 75.74 | 73.53 |
| 10 | 96.04 | 81.76 | 69.82 | 76.81 |
| **Mean** | **80.78** | **79.53** | **62.08** | **66.13** |
| **Standard deviation** | **26.24** | **19.83** | **17.72** | **11.27** |
| **95% CI** | **60.61, 100.95** | **64.29, 94.77** | **48.46, 75.70** | **57.47, 74.79** |

### Two-way ANOVA (sex and rearing between subjects factors)

| | DF | SS | MS | F-value | [a]p-value | $h^2$ | [b]$w^2$ |
|---|---|---|---|---|---|---|---|
| **Sex** | 1 | 7.775 | 7.775 | 0.021 | 0.886 | <0.001 | -0.025 |
| **Rearing** | 1 | 2184.472 | 2184.472 | 5.857 | **0.021** | 0.154 | **0.124** |
| **Sex*Rearing** | 1 | 87.454 | 87.454 | 0.234 | 0.632 | 0.006 | -0.020 |
| **Residual** | 32 | 11934.128 | 372.941 | | | | |

[a]values in red = significant
[b] values in blue = medium effect size

**Olig**

| | Control | | ELS | |
|---|---|---|---|---|
| | **Male** | **Female** | **Male** | **Female** |
| **Experiment** | | | | |
| 1 | 8.81 | 6.8 | 5.92 | 4.76 |
| 2 | ND | ND | ND | ND |
| 3 | 9.26 | 6.08 | 6.52 | 6.25 |
| 4 | 7.53 | 7.21 | 6.57 | 5.63 |
| 5 | 6.42 | 6.98 | 5.47 | 4.38 |
| 6 | 5.36 | 9.54 | 5.64 | 7.27 |
| 7 | 3.61 | 6.64 | 2.42 | 18.76 |
| 8 | 10.19 | 8.95 | 8.35 | 10.27 |
| 9 | 8.22 | 2.47 | 9.1 | 3.71 |
| 10 | 11.71 | 8.28 | 10.46 | 10.19 |
| **Mean** | **7.90** | **6.99** | **6.72** | **7.91** |
| **Standard deviation** | **2.49** | **2.05** | **2.35** | **4.70** |
| **95% CI** | **5.99, 9.81** | **5.41, 8.57** | **4.91, 8.53** | **4.30, 11.52** |

### Two-way ANOVA (sex and rearing between subjects factors)

| | DF | SS | MS | F-value | p-value | $h^2$ | $^bW^2$ |
|---|---|---|---|---|---|---|---|
| **Sex** | 1 | 0.189 | 0.189 | 0.020 | 0.888 | <.001 | -0.029 |
| **Rearing** | 1 | 0.159 | 0.159 | 0.017 | 0.898 | <.001 | -0.029 |
| **Sex*Rearing** | 1 | 9.954 | 9.954 | 0.047 | 0.314 | 0.032 | 0.001 |
| **Residual** | 32 | 304.305 | 9.510 | | | | |

**FCCP**

| | Control | | ELS | |
|---|---|---|---|---|
| | **Male** | **Female** | **Male** | **Female** |
| **Experiment** | | | | |
| 1 | 112.57 | 89.69 | 75.6 | 69.41 |
| 2 | ND | ND | ND | ND |
| 3 | 75.45 | 72.23 | 35.31 | 41.29 |
| 4 | 118.75 | 101.69 | 63.92 | 75.27 |
| 5 | 97.31 | 70.22 | 81.03 | 54.61 |
| 6 | 85.38 | 94.65 | 77.63 | 75.01 |
| 7 | 64.21 | 45.49 | 64.25 | 86.02 |
| 8 | 44.29 | 64.61 | 42.03 | 73.14 |
| 9 | 117.57 | 114.77 | 94.61 | 48.89 |
| 10 | 80.49 | 68.06 | 75.28 | 72.88 |
| **Mean** | **88.45** | **80.16** | **67.74** | **66.28** |
| **Standard deviation** | **25.51** | **21.55** | **18.89** | **14.62** |
| **95% CI** | **68.84, 108.06** | **63.60, 96.72** | **53.22, 82.26** | **55.04, 75.52** |

### Two-way ANOVA (sex and rearing between subjects factors)

| | DF | SS | MS | F-value | [a]p-value | $h^2$ | [b]$w^2$ |
|---|---|---|---|---|---|---|---|
| **Sex** | 1 | 265.418 | 265.418 | 0.619 | 0.437 | 0.016 | -0.010 |
| **Rearing** | 1 | 2520.877 | 2520.877 | 5.874 | **0.021** | 0.151 | **0.122** |
| **Sex*Rearing** | 1 | 141.888 | 141.888 | 0.331 | 0.569 | 0.009 | -0.017 |
| **Residual** | 32 | 13732.078 | 429.127 | | | | |

[a]values in red = significant
[b] values in blue = medium effect size

**AA**

| Experiment | Control | | ELS | |
|---|---|---|---|---|
| | Male | Female | Male | Female |
| 1 | 8.69 | 2.31 | 2.52 | 3.73 |
| 2 | ND | ND | ND | ND |
| 3 | 9.58 | 2.17 | 3.39 | 4.42 |
| 4 | 3.26 | 8.53 | 3.61 | 3.27 |
| 5 | 6.25 | 4.75 | 1.16 | 2.23 |
| 6 | 3.25 | 3.34 | 4.71 | 3.25 |
| 7 | 1.04 | 0.21 | 0.13 | 5.88 |
| 8 | 2.72 | 6.44 | 4.6 | 2.64 |
| 9 | 19.35 | 6.93 | 7.44 | 7.21 |
| 10 | 14.81 | 9.94 | 7.22 | 8.59 |
| **Mean** | **7.66** | **4.96** | **3.86** | **4.58** |
| **Standard deviation** | **6.14** | **3.23** | **2.47** | **2.19** |
| **95% CI** | **2.94, 12.38** | **2.48, 7.44** | **1.96, 5.76** | **2.90, 6.26** |

### Two-way ANOVA (sex and rearing between subjects factors)

| | DF | SS | MS | F-value | p-value | h$^2$ | [b]w$^2$ |
|---|---|---|---|---|---|---|---|
| **Sex** | 1 | 8.890 | 8.890 | 0.603 | 0.443 | 0.016 | -0.010 |
| **Rearing** | 1 | 39.208 | 39.208 | 2.658 | 0.113 | 0.072 | **0.044** |
| **Sex*Rearing** | 1 | 26.300 | 26.300 | 1.783 | 0.191 | 0.048 | **0.021** |
| **Residual** | 32 | 472.068 | 14.752 | | | | |

[b] values in green = small effect size
